# Single cell transcriptomics reveals the effect of PD-L1/TGF-β blockade on the tumor microenvironment

**DOI:** 10.1101/2020.11.13.382358

**Authors:** Yoong Wearn Lim, Garry L. Coles, Savreet K. Sandhu, David S. Johnson, Adam S. Adler, Erica L. Stone

## Abstract

**Background:** The anti-tumor activity of anti-PD-1/PD-L1 therapies correlates with T cell infiltration in tumors. Thus, a major goal in oncology is to find strategies that enhance T cell infiltration and efficacy of anti-PD-1/PD-L1 therapy. TGF-β has been shown to contribute to T cell exclusion and anti-TGF-β improves anti-PD-L1 efficacy *in vivo*. However, TGF-β inhibition has frequently been shown to induce toxicity in the clinic, and the clinical efficacy of combination PD-L1 and TGF-β blockade has not yet been proven. To identify strategies to overcome resistance to PD-L1 blockade, the transcriptional programs associated with PD-L1 and/or TGF-β blockade in the tumor microenvironment should be further elucidated.

**Results:** For the first time, we used single-cell RNA sequencing to characterize the transcriptomic effects of PD-L1 and/or TGF-β blockade on nearly 30,000 single cells in the tumor and surrounding microenvironment. Combination treatment led to upregulation of immune response genes, including multiple chemokine genes such as CCL5, in CD45+ cells, and down-regulation of extracellular matrix genes in CD45-cells. Analysis of publicly available tumor transcriptome profiles showed that the chemokine CCL5 was strongly associated with immune cell infiltration in various human cancers. Further investigation with *in vivo* models showed that intratumorally administered CCL5 enhanced cytotoxic lymphocytes and the anti-tumor activity of anti-PD-L1.

**Conclusions:** Taken together, our data could be leveraged translationally to improve anti-PD-L1 plus anti-TGF-β combination therapy, for example through companion biomarkers, and/or to identify novel targets that could be modulated to overcome resistance.

## Background

Antibodies blocking the PD-1/PD-L1 immune checkpoint pathway have been approved in the first-line setting for a range of cancer types including non-small-cell lung carcinoma (NSCLC), urothelial cancer, triple negative breast cancer, colorectal cancer, head and neck squamous cell carcinoma (HNSCC), microsatellite instability-high cancer, and melanoma. A major goal in oncology is to improve the response rate of these agents to benefit more patients. T cell excluded tumors are less likely to respond to immune checkpoint blockade targeting the PD-1/PD-L1 pathway [1-3]. Furthermore, in metastatic urothelial cancer and some murine tumor models, T cell exclusion correlates with TGF-β signaling, which is thought to induce expression of collagen and other factors leading to a physical barrier that prevents immune infiltration [3-5]. Consistently, in pre-clinical models, combination therapy with anti-PD-L1 plus anti-TGF-β has been shown to induce T cell infiltration and synergistic anti-tumor efficacy [3,5-10]. However, TGF-β blockade can lead to significant toxicity, which has led to termination of several clinical trials testing TGF-β/TGF-βRII inhibitors [11,12], and clinical efficacy has not yet been demonstrated conclusively. Deeper understanding of the transcriptional programs involved in anti-PD-L1 plus anti-TGF-β combination therapy could lead to improved approaches that address these shortcomings.

The transcriptional programs induced by PD-L1 plus TGF-β signaling blockade remain poorly understood. Single-cell RNA sequencing (scRNA-seq) is a powerful tool that provides transcriptional profiles of tens of thousands of cells, enabling comprehensive analysis of the tumor microenvironment. Thus, for the first time, we used scRNA-seq to profile the transcriptional changes induced at the single cell level in the tumor microenvironment after combination anti-PD-L1 plus anti-TGF-β treatment. In particular, we wanted to identify genes that are altered during a productive anti-tumor immune response that may enhance T cell infiltration and efficacy of PD-1/PD-L1 blockade, with the ultimate goal of discovering novel therapeutic strategies to overcome resistance to PD-1/PD-L1 blockade.

We found that anti-TGF-β led to reduced expression of collagen and other matrix remodeling genes by cancer-associated fibroblasts and that combination treatment with anti-PD-L1 (atezolizumab) plus anti-TGF-β further enhanced this downregulation. Anti-PD-L1 induced the expression of several chemokines that are associated with the recruitment of cytotoxic T cells, with a further increase in expression after combination therapy. Analysis of The Cancer Genome Atlas (TCGA) transcriptome data confirmed that multiple chemokines are associated with immune cell infiltration in human tumors, and revealed C-C Motif Chemokine Ligand 5 (CCL5) as the chemokine most highly correlated with immune infiltration across several tumor types. Finally, intratumoral administration of CCL5 increased the frequency of CCR5+ CD8+ T cells and mature CD11b+ NK cells within the tumor, and administration of CCL5 plus anti-PD-L1 (atezolizumab) induced tumor growth inhibition over anti-PD-L1 alone in the murine colon tumor model MC38. Taken together, our data could be leveraged translationally to improve anti-PD-L1 plus anti-TGF-β combination therapy, for example through companion biomarkers, and/or to identify novel targets that could be modulated to overcome resistance.

## Results

### PD-L1 plus TGF-β blockade reduces tumor growth and enhances immune cell infiltration

Anti-TGF-β has been shown to enhance the efficacy of PD-L1 blockade in several murine models [3,5,6,8,13,14]. To confirm these results, and to investigate whether anti-TGF-β enhances the anti-tumor efficacy of the FDA-approved anti-PD-L1 antibody atezolizumab, human PD-L1 knock-in mice bearing subcutaneous colon carcinoma MC38 tumors expressing Hu-PD-L1 (Hu-PD-L1-MC38) were treated with vehicle, anti-PD-L1, or a combination of anti-PD-L1 plus anti-TGF-β. As expected, anti-PD-L1 significantly limited tumor growth (p = 0.0087; Figure 1a, b), while anti-TGF-β alone was not efficacious (Additional file 1, Figure S1a, b). Combination treatment of anti-PD-L1 plus anti-TGF-β was significantly more efficacious than anti-PD-L1 alone (p = 0.038) and led to tumor regression in four of six (66.67%) animals by day 28 (Figure 1a, b). Both anti-PD-L1 and anti-PD-L1 plus anti-TGF-β also improved survival relative to control (Figure 1c). One of six (16.67%) mice from the anti-PD-L1 group and three of six (50%) mice from the anti-PD-L1 plus anti-TGF-β group had a complete response and were re-challenged with Hu-PD-L1 MC38 cells, implanted subcutaneously (s.c.) on the opposite flank from the original tumor. Tumor-naïve, wild-type C57BL/6 mice were used as controls. All previously cured mice, but not the naïve mice, were protected from tumor re-challenge, indicating the presence of anti-tumor immune memory (p = 0.026; Additional file 1, Figure S1c). To investigate if PD-L1 plus TGF-β blockade had an effect on T cell infiltration, CD3 immunohistochemistry (IHC) was performed on tumors from the treated mice. Anti-PD-L1 plus anti-TGF-β significantly increased T cell infiltration while anti-PD-L1 alone did not (Figure 1d). In a different mouse colon carcinoma tumor model, CT26, significant tumor growth inhibition was also observed upon anti-PD-L1 (atezolizumab, which cross reacts with murine PD-L1) plus anti-TGF-β treatment, whether anti-TGF-β was given intraperitoneally (I.P.) or intratumorally (I.T.) (Additional file 1, Figure S2).

**Figure 1.**
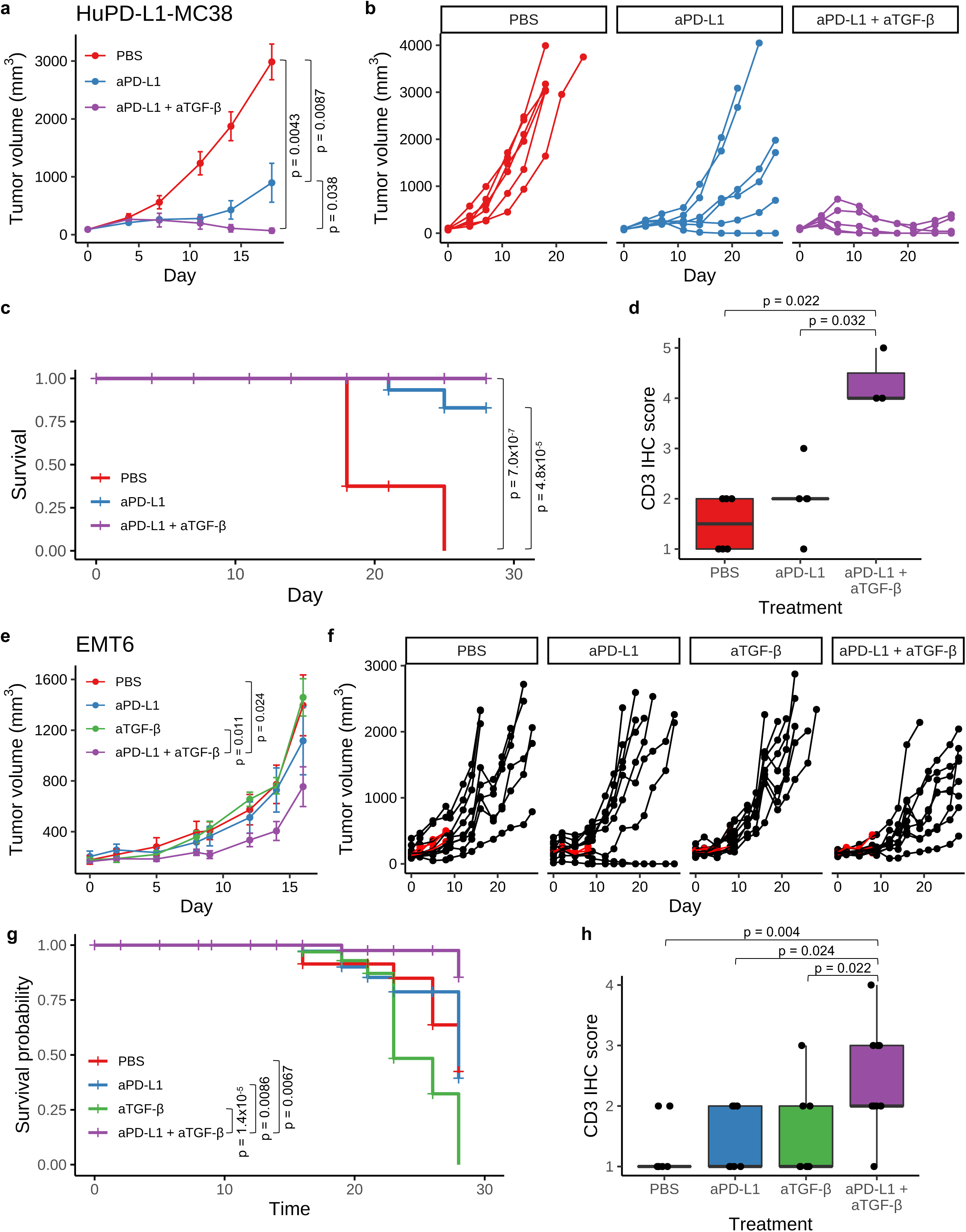
Anti-PD-L1 plus anti-TGF-β leads to tumor regression in mice. **(a-d)** HuPD-L1 KI mice bearing s.c. HuPD-L1-MC38 tumors (n = 6 per group) were dosed I.P. biweekly for three weeks with PBS, aPD-L1 (2 mg/kg; atezolizumab), or aPD-L1 plus aTGF-β (10 mg/kg; 1D11). **(a)** Average MC38 tumor volume ± SEM is shown. P-values were determined using Wilcoxon rank sum test, comparing tumor sizes on day 18. **(b)** Spider plots showing MC38 tumor volume for individual mice over time. **(c)** Survival plot for the MC38 study. P-values were determined using log-rank test. **(d)** CD3 IHC score in MC38 tumors by immunohistochemistry (IHC). Each point is the IHC score representing the density of CD3+ cells from an individual mouse. The horizontal black bars of the box plots indicate median score, while the lower and upper hinges correspond to the first and third quartiles, respectively. P-values were determined using Wilcoxon rank sum test. **(e-h)** Mice bearing orthotopic EMT6 tumors were treated with PBS, aPD-L1 (10 mg/kg for the first dose with each subsequent dose at 5 mg/kg; atezolizumab), aTGF-β (10 mg/kg; 1D11), or aPD-L1 plus aTGF-β. For all treatments, the first dose was administered I.V. on day 0 and the eight subsequent doses were administered I.P. three times per week. n = 12 mice per group, including 3 mice per group taken down early on day 8 for scRNA-seq analysis. **(e)** Average EMT6 tumor volume ± SEM is shown. P-values were determined using Wilcoxon rank sum test. Mice removed early for scRNA-seq analysis were not included in the average. **(f)** Spider plots showing EMT6 tumor volume for individual mice over time. Mice sacrificed for scRNA-seq (n = 3 per group) are shown in red. **(g)** Survival plot for the EMT6 study. P-values were determined using log-rank test. **(h)** CD3 IHC score in EMT6 tumors. Each point is the IHC score representing the density of CD3+ cells from an individual mouse. The horizontal black bars of the box plots indicate median score, while the lower and upper hinges correspond to the first and third quartiles, respectively. P-values were determined using Wilcoxon rank sum test. Non-significant p-values are not shown for all panels.

To study this therapeutic regimen in an orthotopic tumor type, we treated mice harboring EMT6 breast tumors orthotopically grown in the mammary fat pad with PBS, anti-PD-L1 (atezolizumab), anti-TGF-β, or anti-PD-L1 (atezolizumab) plus anti-TGF-β. While we did not observe efficacy from either single agent, anti-PD-L1 plus anti-TGF-β significantly reduced tumor size relative to PBS or anti-TGF-β alone (p<0.03; Figure 1e, f). Further, the combination treatment improved the survival relative to all individual arms (p<0.01; Figure 1g). Anti-PD-L1 plus anti-TGF-β in combination, but neither monotherapy alone, enhanced CD3 immune cell infiltration relative to all individual arms (p<0.03; Figure 1h).

Together, these results established that dual blockade of PD-L1 and TGF-β effectively controlled tumor growth and improved T cell infiltration into the tumor in multiple mouse models. We thus reasoned that *in vivo* inhibition of PD-L1 plus TGF-β could be used as an experimental model to identify genes important for immune cell infiltration and anti-tumor response, and this information could be used to uncover strategies to overcome resistance to anti-PD-L1 therapy. The EMT6 orthotopic tumor model provided us with an opportunity to characterize the single cell molecular responses to anti-PD-L1 ± anti-TGF-β treatment in a setting where anti-TGF-β addition overcomes resistance to anti-PD-L1 therapy. Eight days after the first dose, three representative tumors per group were harvested (Figure 1f, in red), dissociated, flow sorted for CD45+ immune cells and CD45-non-immune cells, and subjected to single-cell RNA sequencing (scRNA-seq).

### Single-cell RNA sequencing of tumors from anti-PD-L1 ± anti-TGF-β treated mice

Single-cell transcriptomic profiles for 27,797 high-quality cells were generated, including 18,002 CD45+ and 9,795 CD45-cells. Uniform Manifold Approximation and Projection (UMAP) dimensionality reduction of the transcriptomes revealed distinct clusters of cells present in all treatment groups (Figure 2a, b). We assigned cell type labels using SingleR, which annotated cell type based on reference transcriptomes of pure cell types in the ImmGen database [15]. To discern tumor cells from host stromal cells, we further performed fine UMAP clustering on the CD45-cells, generating 15 distinct clusters (Additional file 1, Figure S3a). Cells in cluster 7 had high expression of the fibroblast marker *Fap* and were thus annotated as fibroblasts (Additional file 1, Figure S3b). The remaining cells were annotated as tumor cells or assigned their original SingleR labels as endothelial or epithelial cells. Copy number analysis confirmed that the annotated tumor cells exhibited aberrant copy number while the fibroblasts did not (Additional file 1, Figure S3c). When comparing the treatment groups, we observed differences in cell type composition. For example, the anti-PD-L1, anti-TGF-β, and anti-PD-L1 plus anti-TGF-β samples had lower numbers of macrophages (9.2 – 20.3% of CD45+ cells) compared to the PBS control (31% of CD45+ cells; Figure 2c). Conversely, relative to the PBS sample where B cells represented 8.9% of the CD45+ cells, the three treatment groups had higher numbers of B cells (23.7% – 31.9%).

**Figure 2.**
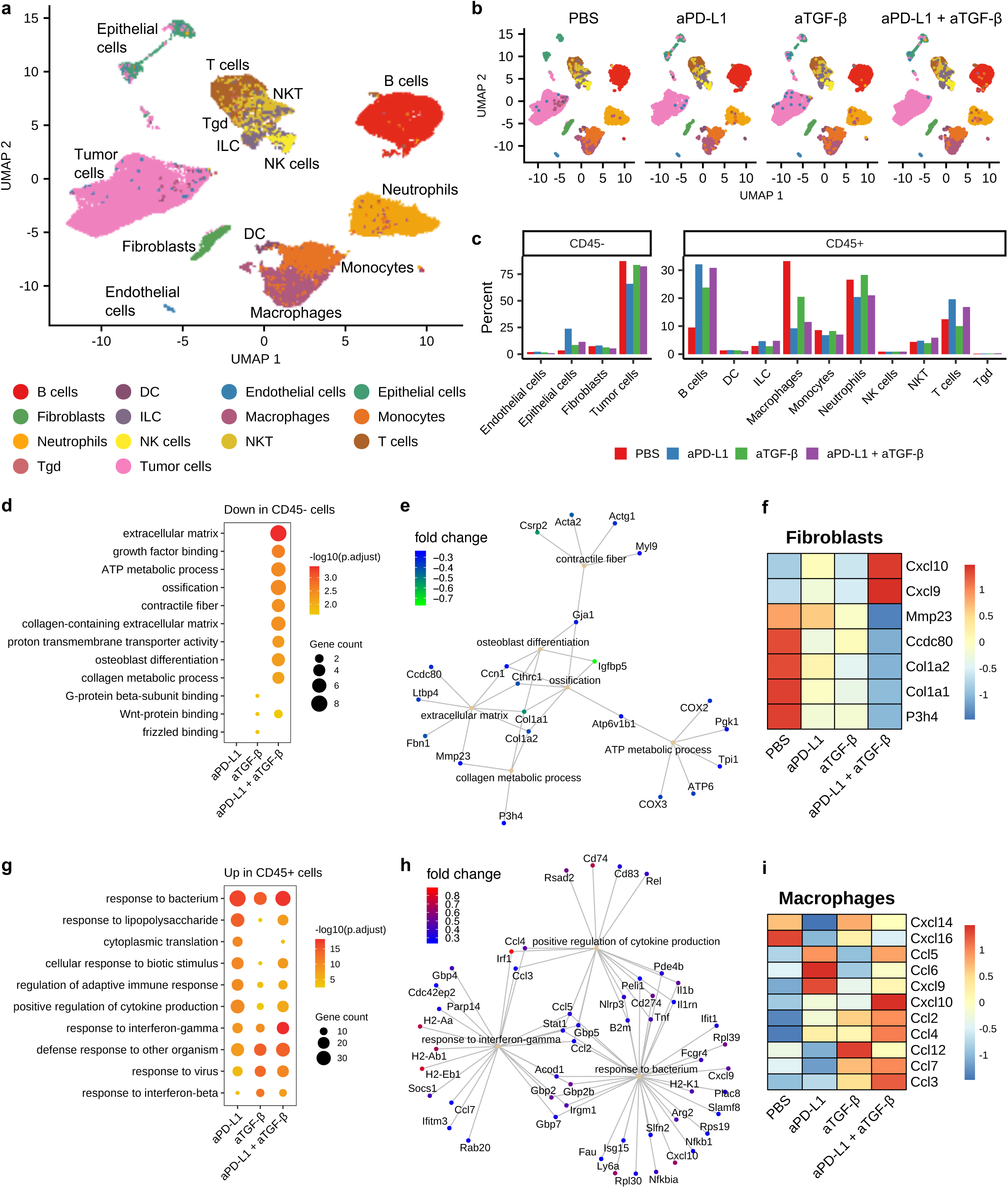
scRNA-seq of EMT6 tumors from mice treated with anti-PD-L1 ± anti-TGF-β. **(a)** Single cell transcriptomes for all cells visualized on a Uniform Manifold Approximation and Projection (UMAP) plot. Cells are colored based on cell type annotations, as indicated in the legend. **(b)** UMAP plots for cells from individual treatment groups. Cell types are annotated with the same color scheme as **a. (c)** Percent composition of cells types within the CD45-cells (left) and the CD45+ (right) cells. The different treatment groups are shown in different colors, as indicated in the legend. **(d)** Functional enrichment analysis of genes downregulated in CD45-cells, for each of the treatment group when compared to the PBS group. The size and color of the circles indicate the number of downregulated genes and the −log_10_ adjusted p-value, respectively, for the enriched terms indicated on the y-axis. **(e)** Gene concept network showing representative enriched terms for the genes downregulated in CD45-cells in the aPD-L1 plus aTGF-β group. The downregulated genes associated with the pathway are shown, with the color of the nodes representing log_2_ fold change in gene expression relative to the PBS group. **(f)** Heatmap showing relative (z-scored per row) gene expression levels for representative differentially expressed genes in fibroblasts. **(g)** Functional enrichment analysis of genes upregulated in CD45+ cells, for each of the treatment group when compared to the PBS group. The size and color of the circles indicate the number of upregulated genes and the −log_10_ adjusted p-value, respectively, for the enriched terms indicated on the y-axis. **(h)** Gene concept network showing representative enriched terms for the genes upregulated in CD45+ cells in the aPD-L1 plus aTGF-β group. The upregulated genes associated with the pathway are shown, with the color of the nodes representing log_2_ fold change in gene expression relative to the PBS group. **(i)** Heatmap showing relative (z-scored per row) gene expression levels for differentially expressed chemokine genes in macrophages.

Next, we performed differential gene expression analysis between each drug treatment group and the control sample, within each annotated cell type. Anti-PD-L1, anti-TGF-β, and anti-PD-L1 plus anti-TGF-β treatments resulted in 598, 132, and 386 differentially expressed genes across different cell types, respectively (Additional file 2, Table S1). To assess the overall impact of the drug treatments on the non-immune and immune compartments, we performed functional enrichment analysis of the differentially expressed genes within the CD45- and CD45+ cells, respectively. While single agent inhibition of PD-L1 did not result in downregulation of specific functional categories for the CD45-cells, TGF-β blockade led to downregulation of WNT-signaling related pathways including Wnt-protein binding and frizzled binding (Figure 2d; Additional file 2, Table S2). Interestingly, WNT/β-catenin pathway was previously shown to contribute to T cell exclusion [16]. More strikingly, dual blockade of PD-L1 plus TGF-β led to significant downregulation of matrix remodeling associated functional categories including extracellular matrix, contractile fiber, and collagen-containing extracellular matrix (Figure 2d). For example, collagen and metalloproteinase genes such as *Col1a1, Col1a2, Cthrc1, P3h4*, and *Mmp23* were downregulated in fibroblasts following combination blockade (Figure 2e, f). This is consistent with previous observations suggesting that TGF-β blockade synergizes with anti-PD-L1 to reprogram the peritumoral stromal fibroblasts [3].

To elucidate the interactions between various cell types involved in TGF-β signaling, we generated a cell-cell communication network involving all TGF-β ligands and receptors using CellPhoneDB [17]. CellPhoneDB analysis implicated multiple cell types as the source of TGF-β ligands, including B cells, dendritic cells, macrophages, NK cells, and intriguingly, tumor cells (Additional file 1, Figure S4a). Tumor cell-expressed TGF-β molecules were predicted to interact with TGF-β receptors on fibroblast cells, macrophages, and tumor cells themselves. Interestingly, these interactions were largely abolished following anti-TGF-β treatment, alone or in combination with anti-PD-L1. This suggests that tumor cell produced TGF-β can induce fibroblasts to express collagen and other extracellular matrix genes resulting in a physical barrier to T cell exclusion, and that TGF-β blockade counteracts this immune suppression.

Functional enrichment analysis of the upregulated genes within the CD45+ cells showed enhanced immune responses following anti-PD-L1 and anti-TGF-β treatments, administered alone or in combination. For example, anti-PD-L1 plus anti-TGF-β induced upregulation of genes involved in response to bacterium (eg. *H2-K1, B2m, Cd274, Il1b*), response to interferon-gamma (eg. *H2-Aa, Stat1, Irf1*) and positive regulation of cytokine production (eg. *Cd74, Cd83, Ccl4*) (Figure 2g, h; Additional file 2, Table S2). Notably, multiple chemokines were upregulated in the macrophages of the anti-PD-L1 sample (*Ccl5, Cxcl9, Ccl4, Cxcl14*), the anti-TGF-β sample (*Ccl12, Ccl4*), and the anti-PD-L1 plus anti-TGF-β sample (*Ccl5, Cxcl10, Cxcl9, Ccl4, Ccl3, Ccl2, Ccl7*) (Figure 2i). This is consistent with previous *in vivo* studies showing anti-PD-1/PD-L1 treatment favors polarization of macrophages towards a more immunostimulatory state [18,19].

Further analysis revealed that chemokines were upregulated in fibroblasts, neutrophils, and T cells in anti-PD-L1 and anti-PD-L1 plus anti-TGF-β treated tumors (Additional file 1, Figure S4b). Additionally, anti-PD-L1 plus anti-TGF-β combination treatment led to upregulation of additional chemokines such as *Cxcl1* and *Cxcl2* in tumor cells above and beyond those induced by either single treatment alone (Additional file 2, Table S1).

Chemokines lay a critical role in recruiting various immune cells to create an inflamed tumor microenvironment [20,21]. The enhanced chemokine expression in samples after anti-PD-L1 plus anti-TGF-β combination treatment suggests that chemokines may facilitate the success of checkpoint blockade treatments. Indeed, both pre- and post-treatment *CXCL9* and *CXCL10* expression levels correlate with response to anti-PD-1 treatment in melanoma patients and anti-PD-L1 treatment in urothelial cancer patients [22-24].

A major goal in immunotherapy is to find strategies to enhance immune cell infiltration. While TGF-β monoclonal antibodies in combination with anti-PD-L1s have been shown to enhance immune infiltration and efficacy in preclinical models, TGF-β targeting molecules have been plagued by toxicity, leading to the halting of several clinical trials with these agents. As an alternative, our data suggest the possibility that a chemokine administered therapeutically may improve immune cell infiltration and efficacy in combination with anti-PD-L1 antibodies.

### CCL5 expression is associated with immune cell infiltration across human cancer types

The scRNA-seq data suggested that the enhanced expression of chemokines in tumors treated with anti-PD-L1 plus anti-TGF-β may contribute to the ability of anti-TGF-β to enhance the anti-tumor efficacy of anti-PD-L1. To further investigate the chemokines that may be most important for enhancing efficacy of anti-PD-L1 therapy, we took a computational approach to identify chemokines associated with immune cell infiltration in different human cancer types using The Cancer Genome Atlas (TCGA). For this we utilized tumor-infiltrating lymphocyte (TIL) scores for TCGA tumor samples, which have been previously quantified using two distinct methods. For one method, Saltz et al. used a deep learning image recognition algorithm to quantify TILs from hematoxylin and eosin (H&E)-stained pathology images for 13 TCGA cancer types [25,26]. Independently, Aran et al. developed a gene signature-based method termed xCell to infer immune cell abundance based on tumor gene expression profiles [27]. Specifically, we focused on the immune cell abundance scores for three cytotoxicity relevant cell types: CD8+ T cells, dendritic cells, and NK cells. Using linear regression models, we tested for association between gene expression levels for a panel of chemokine genes (i.e. genes starting with CXC, CCL, CX3C, or XC) with both the image-based and gene signature-based immune infiltration scoring methods (Figure 3a). Chemokines significantly associated with immune infiltration scores were further ranked based on their association strengths (i.e. regression coefficients) across cancer types (multiple testing adjusted p-values <0.05).

**Figure 3.**
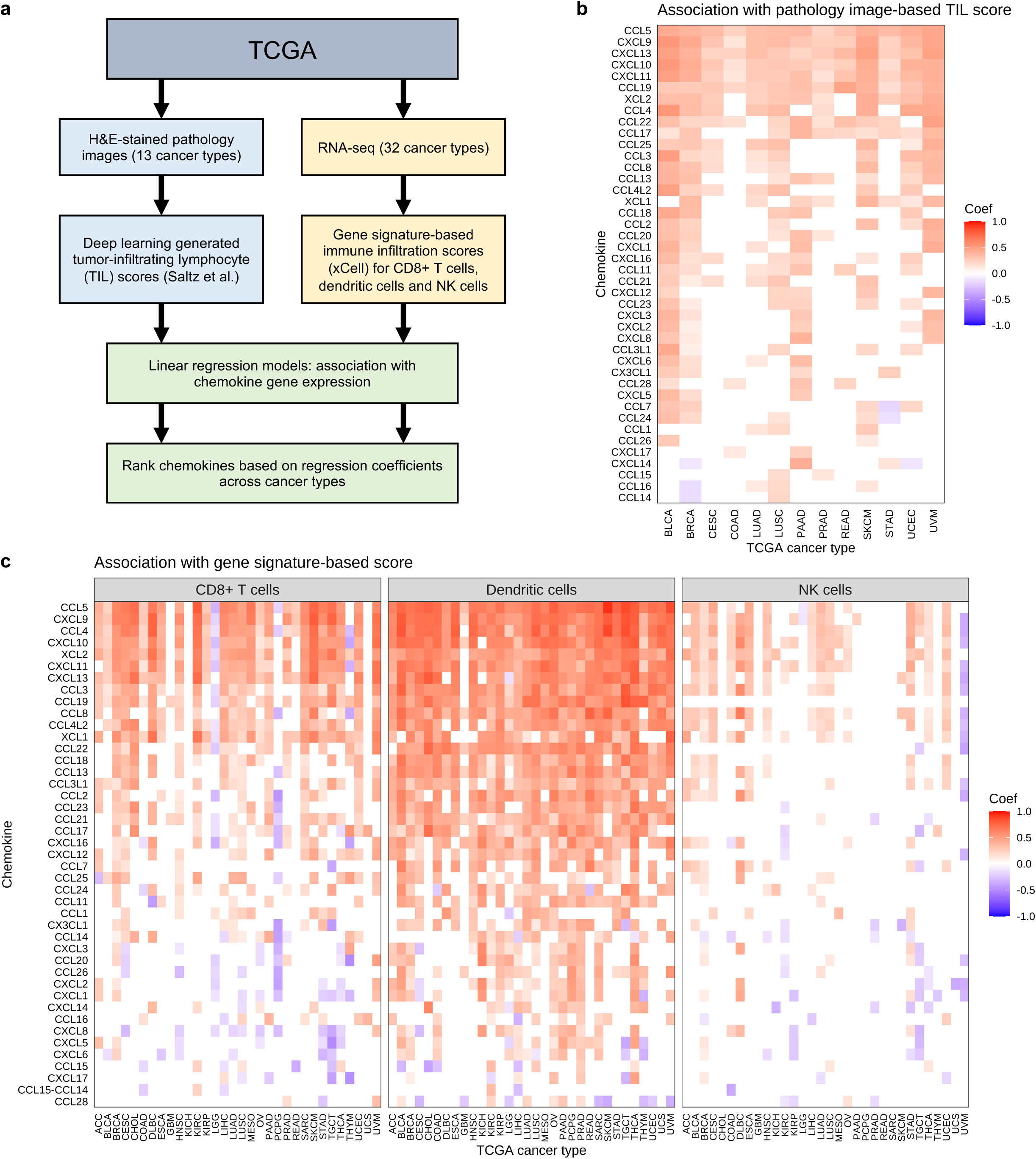
Association of chemokine gene expression with inferred immune cell infiltration in TCGA. **(a)** Analysis workflow to identify chemokine genes associated with immune cell abundance in TCGA tumor samples. Two parallel methods were used to infer immune cell abundance: image-based estimation from pathology slides (left) and gene signature-based estimation from RNA-seq (right). Linear regression models were used to find chemokine genes whose expression levels are associated with inferred immune infiltration. **(b)** Heatmap showing chemokines (rows) significantly associated with the pathology image-based TIL scores across different TCGA cancer types (columns). The color indicates regression coefficients from the linear models (red/orange = positive association between chemokine expression and immune infiltration). **(c)** Heatmap showing chemokines (rows) significantly associated with the gene signature-based immune scores across different TCGA cancer types (columns). The panels represent associations with CD8+ T cells, dendritic cells, and NK cells scores. The chemokines (rows) are ordered descendingly by the sum of the regression coefficients across all cancer types and all 3 immune cell types (i.e. all columns).

The expression levels of six chemokine genes (*CCL5, CXCL9, CXCL13, CXCL10, CXCL11*, and *CCL19*) were significantly associated with the image-based immune scores across all 13 tested cancer types (Figure 3b). For example, correlation of *CCL5* expression in breast cancer (BRCA) with the image-based infiltration scores is shown in Figure S5 (Additional file 1). *CCL5, CXCL9, CXCL13, CXCL10*, and *CXCL11* were also among the top chemokine genes associated with the gene expression-based immune scores across multiple cancer types (Figure 3c). Notably, *CCL5* was consistently the top chemokine associated with immune infiltration scores, as determined by both the image-based and the gene signature-based methods. *CCL5* expression levels was significantly associated with inferred CD8+ T cell, dendritic cell, and NK cell abundance across multiple cancer types (Figure 3c). These findings are consistent with the chemotactic roles of CCL5 in regulating immune cell trafficking in tumors via interactions with its receptor CCR5 [28].

### Combination therapy of anti-PD-L1 plus intratumorally administered CCL5 leads to tumor growth inhibition

Given its chemotactic properties, we hypothesized that intratumoral delivery of recombinant CCL5 protein may enhance immune cell recruitment and control tumor growth. While multiple intratumoral cell subsets express CCR5 (Figure 4a; Additional file 1, Figure S6) [29,30], intratumorally administering recombinant CCL5 to s.c. MC38 tumors resulted in an increased frequency of CD8+ T cells in the tumor that were CCR5 positive twelve days after starting dosing (p = 0.03; Figure 4a, b; Additional file 1, Figure S6). There was also a trend towards an increase in the frequency of NK cells expressing CCR5 (p = 0.09; Figure 4a, b; Additional file 1, Figure S6). Furthermore, we found that intratumoral CCL5 resulted in an increased frequency of NK cells expressing CD11b+ (p = 0.04; Figure 4c, d; Additional file 1, Figure S6). CD11b expression by NK cells has been reported to mark mature NK cells with enhanced effector function [31]. Together these data suggest that intratumorally administered CCL5 altered the tumor microenvironment, including by recruiting cytotoxic lymphocytes.

**Figure 4.**
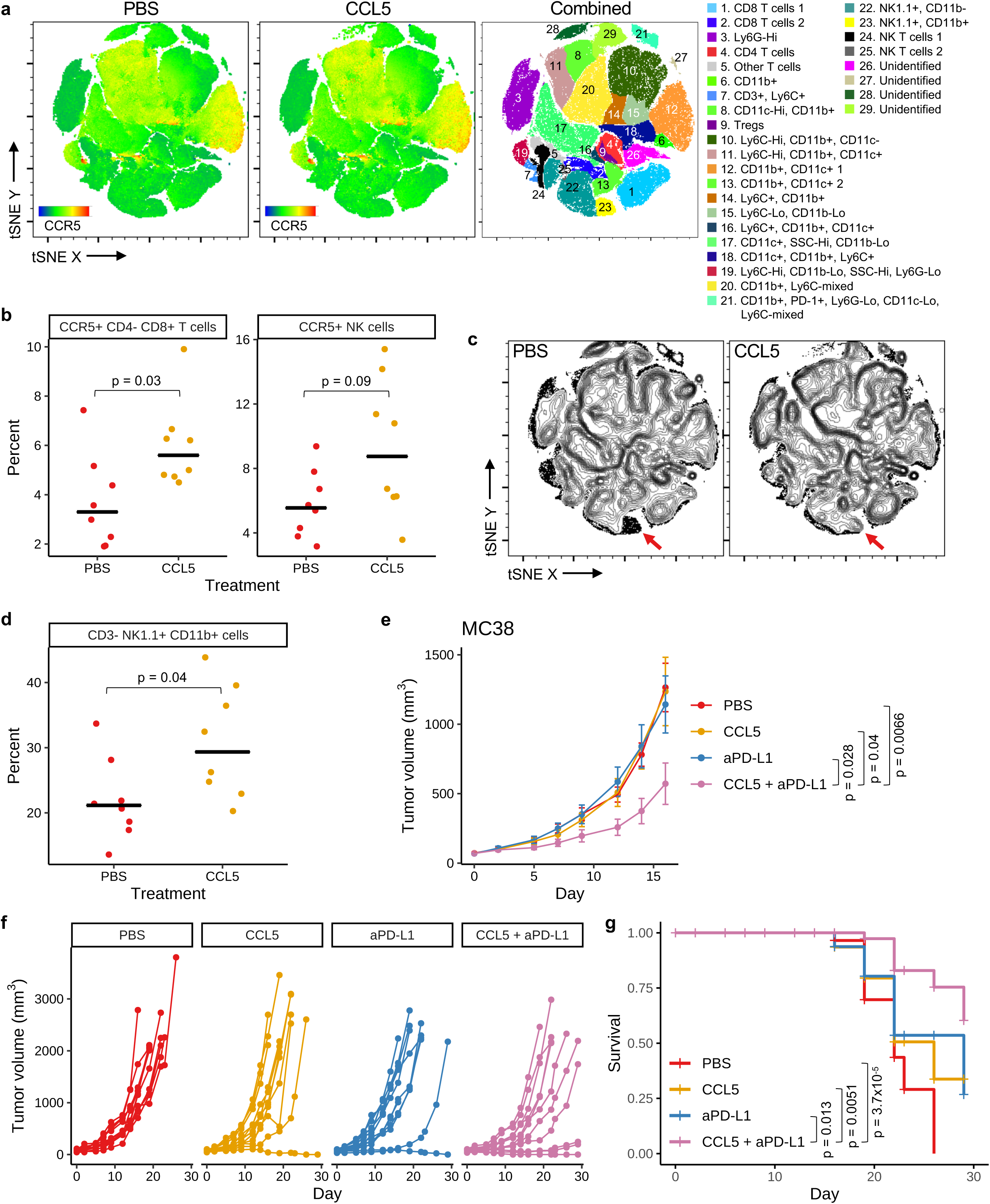
Combination therapy of anti-PD-L1 plus intratumorally administered CCL5 leads to tumor growth inhibition. **(a-d)** Mice bearing s.c. MC38 tumors were administered PBS or CCL5 (1 µg/dose) I.T. three times per a week for 5 doses, and on day 12 tumors were harvested and analyzed by flow cytometry. **(a)** tSNE plots of flow data showing the expression of CCR5 within PBS control (left) or CCL5 treated tumors (middle), or where populations are colored according to expression of marker genes as indicated in the key (right). Heat maps showing expression of each flow marker are in Additional file 1, Figure S6. **(b)** Left: percent of CCR5+ cells within the CD8+ T cell population (live cells, CD4-, CD8+). Right: percent of CCR5+ cells within the NK cell population (live cells, NK1.1+, CD3-). Black cross bars represent median values. P-values were determined using Wilcoxon rank sum test. **(c)** Density plots (with outlier cells indicated as dots) of populations from the tSNE analysis in **a** within control or CCL5 treated samples. Red arrows indicate CD11b+ NK cells (population 23 from **a**). **(d)** Frequency of NK cells (live, single cells, NK1.1+ CD3-) that express CD11b as determined by flow cytometry. Black cross bars represent median values. P-value was determined using Wilcoxon rank sum test. **(e-g)** Mice bearing s.c. MC38 tumors (n = 12 per group) were administered PBS (I.T., 3 times a week for 3 weeks), CCL5 (I.T., 1 µg/animal, 3 times a week for 3 weeks), aPD-L1 (atezolizumab; I.P., 1 mg/kg, twice a week for 3 weeks), or a combination of CCL5 and aPD-L1. **(e)** Average tumor volume ± SEM for each treatment group is shown. P-values were determined using Wilcoxon rank sum test, comparing tumor sizes on day 16. **(f)** Spider plots showing tumor volume for individual mice over time. **(g)** Survival plot for the same study. P-values were determined using log-rank test. Non-significant p-values are not shown for all panels.

Lastly, we determined if intratumorally administered CCL5 was able to enhance the efficacy of anti-PD-L1 therapy using the same tumor model. While neither intratumoral CCL5 nor anti-PD-L1 (atezolizumab) alone inhibited tumor growth, CCL5 in combination with anti-PD-L1 significantly reduced tumor growth relative to all individual arms (p<0.05; Figure 4e, f). Mice that received CCL5 plus anti-PD-L1 combination therapy also had significantly prolonged survival compared to all individual arms (p<0.02; Figure 4g). Thus, intratumorally administered CCL5 is able to overcome resistance to anti-PD-L1 therapy.

## Discussion

For the first time, we used scRNA-seq to investigate gene expression changes in tens of thousands of single cells associated with anti-PD-L1 and/or anti-TGF-β therapy. Anti-PD-L1 treatment alone or in combination with anti-TGF-β induced immune response related genes in both fibroblasts and immune cells, including multiple chemokines associated with the recruitment of cytotoxic cells. The combination of anti-PD-L1 plus anti-TGF-β enhanced the expression of many of these genes over anti-PD-L1 alone, and thus the combination appears to work together to alter the expression of fibroblasts in a way that may enhance immune infiltration into tumors.

Our scRNA-seq data set comprises almost 30,000 single cell transcriptomes across four treatment groups. This is a significant community resource for discovery of novel biomarkers and therapeutic targets. Though patient response to PD-1/PD-L1 checkpoint inhibition remains difficult to predict, biomarkers for positive outcomes in anti-PD-L1 therapy include IHC for tumor PD-L1 and tumor mutational burden [32], while biomarkers for anti-TGF-β are unknown. Our data suggest that extracellular matrix or chemokine receptor genes could be used to identify patients more likely to achieve immune cell infiltration for anti-PD-L1 and/or anti-TGF-β therapy. Our work also identified dozens of genes that could be further investigated as targets for cancer therapy. For example, collagen and metalloproteinase genes such as *Col1a1, Col1a2, Cthrc1, P3h4*, and *Mmp23*, which were downregulated in fibroblasts, could be targeted therapeutically to encourage immune cell infiltration. Our work also suggested that chemokines, cytokines, and their receptors (e.g., *Ccl5, Cxcl10, Cxcl9, Ccl4, Ccl3, Ccl2, Ccl7*) were involved in immune cell infiltration, and therefore represent attractive therapeutic targets. Such molecules could be blocked or activated by monoclonal antibodies, or attached to tumor-directed antibodies as bi-functional molecules [33]. Though there is extensive research on the involvement of chemokines, cytokines, and their receptors in tumor models [20,21], our work uniquely identifies molecules specific to anti-PD-L1 and/or anti-TGF-β therapy, suggesting possibilities for combination therapies. However, all such targets require significant further research. For example, not all chemokines enhance immune responses, and instead some may actually inhibit anti-tumor immunity [34,35].

The particular case of CCL5 highlights the challenges of understanding and modulating the complex pathways underlying the tumor microenvironment. Specifically, conflicting data exist as to whether the CCL5 pathway should be viewed as a target for activation or a target for inhibition. For example, CCL5 expression has been associated with better survival in patients treated with immunotherapy, and CCL5 and its receptor CCR5 are required for productive anti-tumor responses following immunotherapy [30,36-40]. More specifically, blockade of CCL5 after tumor establishment limited the anti-tumor effect of CD40 agonism plus immune checkpoint blockade [30]. In contrast, CCL5 enhances homing of Tregs in some tumor models, and the transplantable pancreatic tumor model KPC has been shown to grow less aggressively in mice lacking CCL5 [41-44]. Thus, CCL5 may have a different role in controlling the makeup of the tumor microenvironment in developing versus established tumors, especially in the context of immunotherapy. Our data showed that intratumoral injection of CCL5 into MC38 tumors might enhance the fraction of CD8+ T cells positive for the CCL5 receptor CCR5, could increase intratumoral levels of mature CD11b+ NK cells, and, in combination with an otherwise ineffective dose of anti-PD-L1, limited tumor progression. Taken together, data from our group and others suggest that CCL5 may have an important role in modulating immune cells in the tumor microenvironment, but specific mechanisms and potential therapeutic approaches remain elusive.

## Methods

### Murine models

Murine experiments were approved by the Institutional Animal Care and Use Committees of Explora BioLabs, Crown Bioscience, or Champions Oncology.

Experiments using Hu-PD-L1 KI mice were performed at Crown Bioscience (Taicang Jiangsu Province, China), and mice were obtained from Shanghai Model Organisms Center, Inc. Crown Bioscience acquired MC38 cells from FDCC (The Institutes of Biomedical Sciences (IBS), Fudan University) and authenticated cell identity by SNP analysis. MC38 cells expressing human PD-L1 (Hu-PD-L1 MC38) were created by knocking out murine *B7h1* (expressing PD-L1) and transgenically expressing human *B7H1* driven by the CMV promoter. MC38 and Hu-PD-L1 MC38 cells were tested for mycoplasma after cell banking. Two independent *in vivo* Hu-PD-L1 MC38 tumor experiments were performed. In the first study, 1×10^6^ Hu-PD-L1 MC38 were implanted s.c. into the flank of HuPD-L1 knock-in C57BL/6 mice (female, 9-11 weeks old). The mice were randomized into treatment groups (n = 6 per group) when average tumor volume reached 90.48 mm^3^. Dosing was initiated on the same day (day 0). The mice were dosed I.P. biweekly for 3 weeks with PBS, anti-PD-L1 (atezolizumab, Roche, 2 mg/kg), or a combination of anti-PD-L1 (2 mg/kg) and anti-TGF-β (mouse IgG1 clone 1D11 from BioXCell, 10 mg/kg). Tumor volumes and body weight were measured in a blinded fashion twice a week. Tumor volumes were calculated using the formula: V = (L x W x W)/2, where V is tumor volume, L is tumor length (the longest tumor dimension), and W is tumor width (the longest tumor dimension perpendicular to L). Individual animals were removed from the study as their tumor volumes measured greater than 3000 mm^3^. All mice that did not have a complete response within the initial 28-day observation period were euthanized, and tumors were collected as formalin-fixed paraffin-embedded (FFPE) blocks. Mice with full tumor regression (tumor volume of 0.00 mm^3^) after 4 weeks were re-challenged on study day 49. Six wildtype C57BL/6 mice (16-18 weeks of age) from Shanghai Lingchang Biotechnology Co., Ltd were used as a naïve control for the re-challenge experiment. Mice for the re-challenge experiment were inoculated s.c. at the opposite flank (left lower flank) with 1×10^6^ HuPD-L1-MC38 cells. Following tumor implantation, tumor volume and body weight were measured twice a week in a blinded manner for an additional 4 weeks.

In the second Hu-PD-L1 MC38 study, 1×10^6^ Hu-PD-L1 MC38 were implanted s.c. into the flank of HuPD-L1 knock-in C57BL/6 mice (female, 9-11 weeks old). The mice were randomized into treatment groups (n = 6 per group) when average tumor volume reached 99.3 mm^3^. Dosing was initiated on the same day (day 0). The mice were dosed I.P. biweekly for 3 weeks with PBS, anti-PD-L1 (atezolizumab, Roche, 2 mg/kg), anti-TGF-β (mouse IgG1 clone 1D11 from BioXCell, 10 mg/kg), or a combination of anti-PD-L1 (2 mg/kg) and anti-TGF-β (10 mg/kg). Tumor volumes and body weight were measured in the same manner as in the first study, and individual mice were removed from the study as their tumor volumes measured >3000 mm^3^. In both studies, two-sided Wilcoxon rank sum test was used to determine if the differences in average tumor volume are statistically significant. Survival analysis was performed using the R package survminer (version 0.4.5) and p-values were determined using log-rank test.

The EMT6 tumor study was performed at Champions Oncology, Inc. Champions Oncology acquired EMT6 cells from ATCC and routinely have STR analysis and pathogen testing, including for mycoplasma, of these cells completed by IDEXX BioAnalytics. For the *in vivo* EMT6 tumor experiment, 2.5×10^5^ EMT6 cells suspended in 0.1 mL 1x PBS were implanted orthotopically in the 4th right mammary fat pad of BALB/C mice (female, 7.5-12 weeks old; Taconic). When tumors reached an average tumor volume of 176.56 mm^3^ with a range of 15 – 400 mm^3^, animals were matched by tumor volume into four blinded groups (n = 12 mice per group) on day 0. The four treatment groups were PBS, anti-PD-L1 (10 mg/kg for the first dose with each subsequent dose at 5 mg/kg; atezolizumab, Roche), anti-TGF-β (10 mg/kg; mouse IgG1 clone 1D11, BioXCell), or a combination of anti-PD-L1 plus anti-TGF-β. For all treatments, the first dose was administered intravenously (I.V.) on day 0 and the eight subsequent doses were administered I.P. using a dose volume of 5 mL/kg three times per a week. Tumor volumes were measured three times weekly by a person blinded to treatment group. Mice were observed for 28 days post initiation of dosing, with individual animals being removed from the study as their tumor volumes measured >2000 mm^3^ and collected for FFPE. On day 8, three mice per group (chosen based on day 6 tumor volumes that were representative of the entire group) were sacrificed. Whole tumors from the selected mice were sterilely harvested, removing adjacent skin but leaving the exterior surface of the tumor intact to preserve the tumor microenvironment. If there was a large amount of mammary fat attached to the tumor or if the tumor had invaded the adjacent tissue, the tumor was cut away from the tissue and approximately 1-2 mm of the tissues was kept attached to the tumor, so that the boundary between the tumor and non-tumor tissues was not disturbed. Tumors were placed into MACS tissue storage solution buffer (Miltenyi Biotec) on ice packs and shipped overnight for scRNA-seq processing.

The CT26 tumor study was performed at Explora BioLabs by GigaGen staff. CT26 cells were obtained from ATCC and expanded to generate a research cell bank which was used for murine experiments after a representative vial was found to be mycoplasma negative. 5×10^5^ CT26 cells suspended in 0.1 mL 1x PBS were implanted s.c. into the right hind flank of BALB/c mice (female, 8-10 weeks old; Taconic). On day 10 post implantation when tumors were an average tumor volume of 40.01 mm^3^ (range 25.5 – 63.89 mm^3^), animals were randomized into seven blinded groups based on tumor volume. Group 1 received PBS I.P. (n = 10). Group 2 received anti-PD-L1 I.P. (2 mg/kg; atezolizumab, Roche; n = 12). Group 3 received anti-TGF-β I.P. (10 mg/kg; clone 1D11, BioXcell; n = 10). Group 4 received anti-PD-L1 I.P. + anti-TGF-β I.P. (n = 12). Group 5 received anti-PD-L1 I.P. (2 mg/kg) + PBS I.T. (n = 12). Group 6 received anti-TGF-β I.T. (40 µg/dose; n = 6). Group 7 received anti-PD-L1 I.P. (2 mg/kg) + anti-TGF-β I.T. (40 ug/dose) (n = 12). Starting on day 11, mice were treated two times a week for three weeks. Tumors were measured three times weekly. Mice were observed for 35 days post implantation, with individual animals being removed from the study as their tumor volumes measured greater than 2000 mm^3^.

The MC38 tumors study was performed at Champions Oncology, Inc. 5×10^5^ MC38 cells suspended in 100 ul of PBS were injected s.c. into the left flank of C57BL/6 mice (female, 7.5-12 weeks old; Taconic). When tumors reached an average tumor volume of 71.1 mm^3^ with a range of 32 – 113 mm^3^, animals were randomized into four blinded groups (n = 12 mice per group) on day 0. The treatment groups were PBS, recombinant human CCL5 (R&D systems, 278-RN-050/CF), PBS plus anti-PD-L1 (atezolizumab, Roche), and CCL5 plus anti-PD-L1. PBS and CCL5 (1 μg/animal) were administered I.T., three times a week for three weeks. Anti-PD-L1 (1 mg/kg) was administered I.P., twice a week for three weeks. Dosing started on day 1. Tumor volumes were measured three times weekly. Mice were observed for 28 days, with individual animals being removed from the study as their tumor volumes measured >2000 mm^3^, and the tumors were collected as FFPE. Two-sided Wilcoxon rank sum test was used to determine statistical significance in difference in average tumor volume.

### Immunohistochemistry

All immunohistochemistry (IHC) was performed by Allele Biotechnology. 4 μm sections were cut and placed on glass slides for CD3 antibody staining (clone SP7, Abcam). An IHC score was blindly assigned to each stained tumor section based on the observed density of CD3 cells across the entire slide (1 to 5, where 1 = very low density of CD3+ staining and 5 = very high density of CD3+ cells). Statistical significance in the difference in CD3+ cell score was determined using Wilcoxon rank sum test.

### Single cell RNA sequencing

EMT6 mammary fat pad tumors and the neighboring tissue were collected from mice such that the entire tumor and the interface between tumor and normal tissue was preserved, and the tissue was stored overnight at 4°C in MACS Tissue Storage Solution (Miltenyi Biotec). Tumor tissue was minced and dissociated using the mouse Tumor Dissociation Kit (Miltenyi Biotec) and gentleMACS Octo Dissociator with Heaters (Miltenyi Biotec). The “37C_m_TDK_2” and “m_imptumor_01” programs were used for the primary and secondary gentle MACS dissociation programs. Tumor suspensions were filtered through a 70 µm MACS Smart Strainers (Miltenyi Biotec) and washed with PBS containing 0.5% BSA and 2 mM EDTA. Cells were pelleted by centrifugation at 500xg for 7 min. A red blood cell lysis step was performed by incubating cells for 2 minutes with 1x Red Blood Cell Lysis Solution (Miltenyi Biotec). Cells were washed again and then cell viability was measured using the Nexcelom Cellometer K2 Fluorescent Viability Cell Counter and the ViaStain AOPI Staining Solution (Nexcelom). Single cell suspensions were then cryopreserved using CryoStor CS10 solution (STEMCELL Technologies) and stored in liquid nitrogen.

Cells were thawed and FACS sorted for DAPI-CD45+ (live, hematopoietic cells) and DAPI-CD45-(live, non-hematopoietic cells) populations. Briefly, cells were thawed in a 37°C water bath, resuspended in 10 mL of RPMI (Gibco) with 10% FBS (Gibco), centrifuged at 500xg for 5 minutes, washed in PBS containing 0.5% BSA and 2 mM EDTA, and counted using the Nexcelom Cellometer. Approximately 10×10^6^ cells were transferred to a new tube, washed in PBS with 0.5% BSA and 2 mM EDTA, and centrifuged at 500xg for 5 minutes. The liquid was poured off from the cells (leaving approximately 100 µl in each tube) and 5 µl of mouse Fc Blocking Reagent (rat anti-Mouse CD16/CD32 BD) was added to samples and incubated for 5 minutes at 4°C. 162.5 µl of Brilliant Stain Buffer (BD) containing 12.5 µl of mouse CD45-PE clone 30-F11 (BioLegend) was added to cells and incubated on ice for 30 minutes protected from light. Cells were washed, resuspended in 1x DAPI solution (in PBS plus 0.5% BSA and 2 mM EDTA; BioLegend) at 5×10^6^ cells/ml, and filtered into 5 mL FACS tubes with cell-strainer caps (Falcon). Cells were sorted using the BD FACSMelody using the 100 µm nozzle. Mouse splenocytes were used as a positive control to determine the voltage needed to identify CD45+ immune cells in the initial side scatter area and forward scatter area gate that would be used for tumor cell suspensions. Singlet and DAPI-(live cell) gates were then applied. Stained single cell suspensions from EMT6 tumors were then sorted and approximately 1×10^5^ live CD45+ and 1×10^5^ live CD45-cells were collected at 4°C. Cells were washed in RPMI with 10% FBS, centrifuged at 500xg for 5 minutes, and counted as described above. Cells were resuspended in RPMI with 10% FBS at 7×10^5^ cells/ml and were placed on ice. CD45+ and CD45-sorted cells from each of the four treatment groups were run on the 10X Genomics Chromium device using the Single Cell 3’ Reagent Kit v3 (work performed at SeqMatic). Libraries were sequenced at SeqMatic with an Illumina NovaSeq SP 100 cycle kit (28 bp for read 1, 91 bp for read 2).

### Single cell RNA-seq data analysis

Raw sequencing FASTQ files were processed using the Cell Ranger (version 3.1.0) analysis pipeline. Briefly, alignment and filtering of sequencing reads, barcode counting, and UMI counting were performed using the *cellranger count* command. The reads were aligned to the *Mus musculus* reference genome assembly GRCm38 (mm10). The outputs of *cellranger count* for all samples were aggregated using the command *cellranger aggr*, normalizing the runs to the same sequencing depth. Secondary analyses were performed using the Seurat package (version 3.1.2) in R (version 3.6.0) [45]. First, we performed quality control by removing cells with fewer than 200 or more than 5,000 expressed genes, as low and high number of gene counts may indicate low quality cells and cell multiplets, respectively. We also removed cells with more than 10% mitochondrial gene expression. This resulted in 27,797 cells, including 18,002 CD45+ and 9,795 CD45-cells. We used *sctransform* to normalize the expression data and to regress out mitochondrial mapping percentage as a confounding source of variation [46]. We performed principal component analysis using the *RunPCA* function in Seurat. Principal components 1 to 30 were provided as an input for non-linear dimensionality reduction via Uniform Manifold Approximation and Projection (UMAP) using the *RunUMAP* function.

For cell type annotation, we used the R package SingleR, which leveraged reference transcriptomic datasets of pure cell types to infer the cell of origin of each of the single cells independently [47]. The ImmGen (www.immgen.org) reference dataset was used. The cells were first annotated using the main cell type labels generated by SingleR. Since the ImmGen reference did not include tumor cell as a cell type, we needed to manually discern tumor cells from host non-immune cells within the CD45-cells. We therefore reanalyzed the CD45-cells using *RunUMAP*, followed by clustering using the *FindNeighbors* and *FindClusters* functions in Seurat. These functions applied shared nearest neighbor (SNN) based clustering on the scRNA-seq data, identifying 15 clusters of cells. Cells in cluster 7 were annotated as fibroblasts based on expression of fibroblast marker gene *Fap*. Cells previously annotated by SingleR as endothelial cells or epithelial cells were assigned their original SingleR labels. All other CD45-cells were annotated as tumor cells. To validate the annotations, we analyzed copy number alterations for the CD45-cells using InferCNV [48], using epithelial and endothelial cells as reference cells.

### Differential gene expression and functional enrichment analysis

We performed differential gene expression analysis within each cell type using the *FindMarkers* function in Seurat. We compared each treatment group to the PBS control, using Wilcoxon rank sum test to identify differentially expressed (DE) genes (log fold change >0.25, adjusted p-value <0.05) between the two groups of cells. We also compared the anti-PD-L1 plus anti-TGF-β treatment group to the single treatment groups. We used ClusterProfiler (version 3.12.0) [49] to perform functional enrichment analysis. For each treatment group, we generated foreground gene lists containing genes upregulated or downregulated in any CD45+ and CD45-cell types, respectively. These gene lists were tested against background gene lists containing all expressed genes (expressed in at least 5 cells) in CD45+ and CD45-cells, respectively. The enrichment analysis was performed using the *enrichGO* function in ClusterProfiler using the org.Mm.eg.db database and the gene ontology (GO) categories biological process (BP), molecular function (MF), and cellular component (CC). Multiple testing correction was performed using the Benjamini and Hochberg method. The enriched pathways were visualized as gene concept networks using the *cnetplot* function of the enrichplot package (version 1.4.0) [50].

### Cell-cell communication using CellphoneDB

Cell-cell communication analysis was performed using CellphoneDB [17], a repository of curated receptors, ligands, and their interactions. First, all mouse genes from this study were mapped to their human orthologs using biomaRt [51]. scRNA-seq count tables and cell type annotations for each treatment group were used as inputs for running CellphoneDB, using the method “statistical analysis” and default parameters. Receptor-ligand interactions were visualized as dot plots using ggplot2 (version 3.2.1) [52].

### Association of chemokine expression and immune cell abundance in TCGA

We inferred intratumoral immune cell abundance using either an image-based approach or a gene signature-based approach. For the image-based approach, we downloaded 4,612 published TIL scores estimated from TCGA H&E images [25,26]. Briefly, Saltz et al. developed deep learning-based image recognition to classify and quantify lymphocytes on H&E diagnostic images from 13 TCGA cancer types (LUAD, BRCA, PAAD, COAD, LUSC, PRAD, UCEC, READ, BLCA, STAD, CESC, SKCM and UVM). The deep learning model training process was repeated until pathologists judged that the lymphocyte classification was adequate. The TIL scores were reported as “TIL Regional Fraction” in Table S1 by Thorsson et al. [26]. For the gene signature-based approach, we downloaded pre-calculated xCell immune cell scores for 9,358 TCGA samples (https://xcell.ucsf.edu/) [27] and filtered for immune cell types of interest: CD8+ T cells, dendritic cells (aDC), and NK cells.

For TCGA tumor gene expression data, we downloaded RNA-seq quantifications (in Fragments Per Kilobase of transcript per Million mapped reads, FPKM) from the TCGA portal (https://portal.gdc.cancer.gov/) for 32 cancer types (ACC, BLCA, BRCA, CESC, CHOL, COAD, DLBC, ESCA, GBM, HNSC, KICH, KIRC, KIRP, LGG, LIHC, LUAD, LUSC, MESO, OV, PAAD, PCPG, PRAD, READ, SARC, SKCM, STAD, TGCT, THCA, THYM, UCEC, UCS, UVM). We filtered for chemokine genes by retaining only expression data for genes whose names started with CXC, CCL, CX3C, or XC. Within each cancer type, genes that were not or lowly expressed (median expression across all samples <0.01 FPKM) were excluded from further analysis.

For each TCGA cancer type, the chemokine gene expression data frame was combined with either the image-based or the gene signature-based immune score data frame. Only samples with data present in both data frames were analyzed. A pseudo count of 0.01 was added to the expression and immune scores, followed by log_2_ and z-score transformation, in preparation for linear regression analysis. We performed linear regression analysis for each chemokine gene using immune score as the dependent variable and chemokine gene expression as the independent variable. P-values were adjusted for multiple testing using the Benjamini and Hochberg method. Chemokines significantly associated with immune scores were further ranked by the sum of their regression coefficients across different cancer types. The results were visualized as heatmaps using ggplot2 (version 3.2.1) [52]. All data manipulations and analyses were performed in R 3.6.0.

### Flow cytometry analysis

An additional cohort of C57BL/6 mice bearing MC38 tumors were treated with 5 doses I.T. PBS or CCL5 (1 μg/animal) on days 1, 3, 5, 8, 10 as indicated in the murine models section above. Tumors were harvested on day 12, cut into small pieces approximately of 2-4 mm, and dissociated using the Miltenyi Biotec MACS Tumor Dissociation Kit according to manufacturer instructions (all performed by Champions Oncology). Dissociated cell suspensions were filtered through a 70 µm strainer with 10 mL RPMI 1640 (Thermo Fisher Scientific) and centrifuged at 300xg for 7 minutes. Cells were resuspended in MACS Quant running buffer (Miltenyi Biotec) and red blood cells were lysed using RBC Pharm Lyse (BD). Cells were pelleted, washed, and resuspended with MACs buffer. Cells were then put into 96 well plates and stained with fluorescently labelled antibodies and FVS780 viability stain. Cells were analyzed using BD FACSymphony. FlowJo 10 was used to generate tSNE plots of cells pre-gated based on live, single cells and taking into account data from side scatter, CD11b (BUV395, clone M1/70, BD Biosciences), CD4 (BUV496, clone RM4-5, BD Biosciences), CD8 (9BUV661, clone 53-6.7, BD Biosciences), CD3 (BUV805, clone 17A2, BD Biosciences), CD11c (BV421, clone N418, BD Biosciences), Ly6G (BV605, clone 1A8, BD Biosciences), CD44 (BV786, clone IM7, BD Biosciences), Ly6C (AF488, clone HK1.4, BioLegend), PD-1 (BB700, clone j43, BD Biosciences), FOXP3 (PE, clone FJK-16S, Fisher Scientific), NK1.1 (PECy7, clone PK136, BioLegend), and CCR5 (CD195; APC, Clone 2D7, BD Biosciences) staining. Populations were named based on expression of the indicated marker(s).

### Availability of data and materials

Single cell RNA-seq fastq sequence files are deposited at the Sequence Read Archive (https://www.ncbi.nlm.nih.gov/sra/), under BioProject ID PRJNA615238.

## Acknowledgements

The authors appreciate thoughtful discussions from Matthew Spindler, Kyle Carter, Jan Simons, Ellen Wagner, Nicholas Wayham, and Jasmeen Saini (GigaGen). The authors also thank Penny Chen and Michael Boice (Crown Bioscience) and Mary Topalovski and Monika Buczek (Champions Oncology) for assistance with efficacy studies, and Abbas Hussain and Anthony Wong (Allele Biotechnology) for assistance with IHC.

## Author information

### Contributions

Y.W.L., A.S.A., and E.L.S. conceived and designed the study; Y.W.L., G.L.C., and S.K.S. performed the experiments; Y.W.L., S.K.S., A.S.A., and E.L.S. analyzed the data; Y.W.L., D.S.J., A.S.A., and E.L.S. wrote and edited the manuscript; D.S.J., A.S.A., and E.L.S. provided supervision and project administration; D.S.J. acquired the funding.

## Ethics declarations

### Ethics approval and consent to participate

Not applicable.

### Consent for publication

Not applicable.

## Additional files

### Additional file 1

Supplementary figures.

### Additional file 2

Supplementary tables.

