## Additional file 1 (supplementary figures) for "Single cell transcriptomics reveals the effect of PD-L1/TGF-β blockade on the tumor microenvironment"

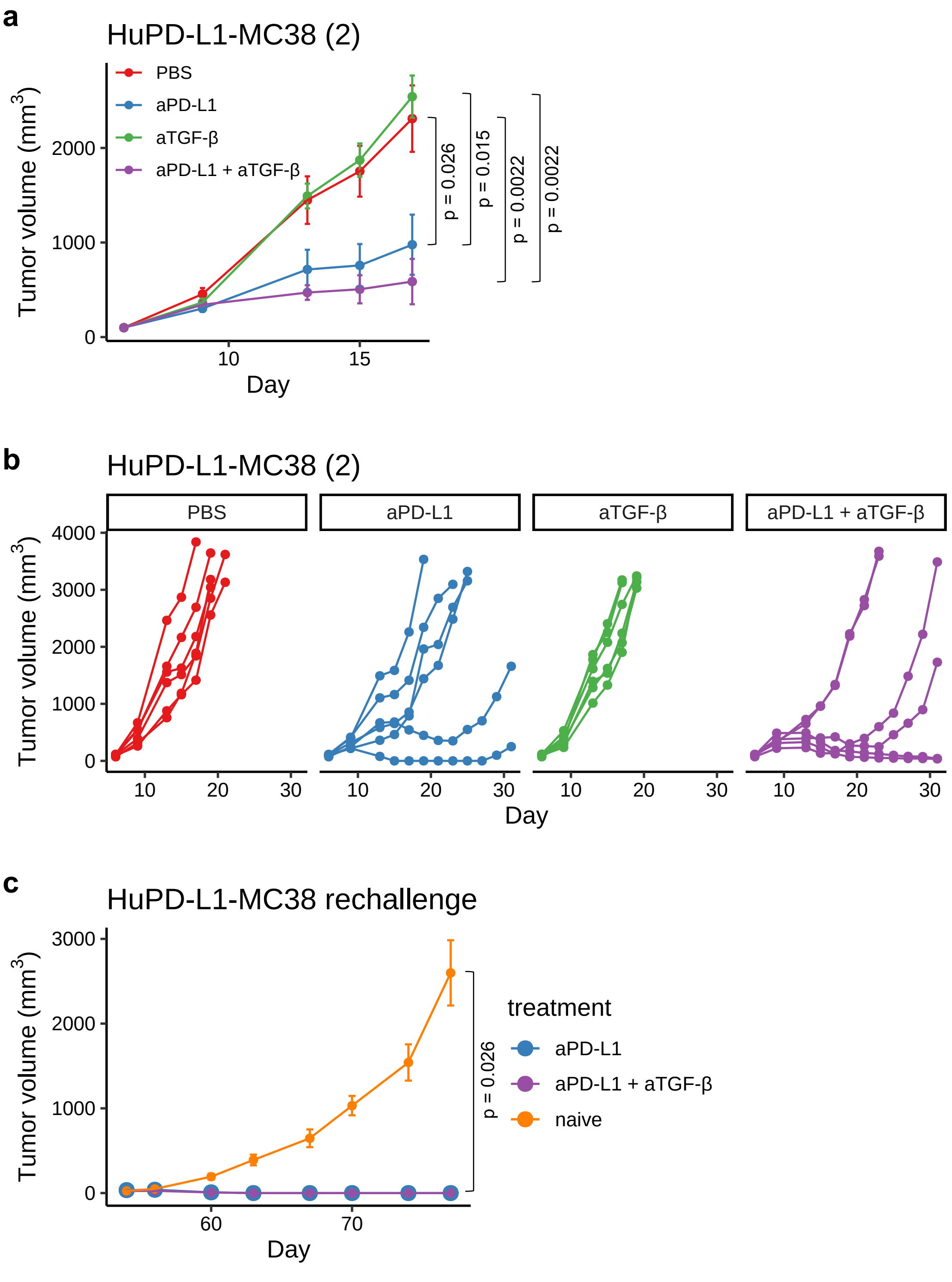


**Figure S1. Anti-PD-L1 plus anti-TGF-β in the murine tumor model HuPD-L1-MC38. (a-b)** Mice bearing s.c. HuPD-L1-MC38 tumors (n = 6 per group) were dosed I.P. biweekly for three weeks with PBS, aPD-L1 (2 mg/kg; atezolizumab), aTGF-β (10 mg/kg; 1D11), or aPD-L1 plus aTGF-β. **(a)** Average MC38 tumor volume ± SEM is shown. P-values were determined using Wilcoxon rank sum test, comparing tumor sizes on day 17. Non-significant p-values are not shown. **(b)** Spider plots showing tumor volume for individual mice over time. **(c)** Mice from Figure 1a that resulted in a complete response (aPD-L1, n = 1; aPD-L1 + aTGF-β, n = 3) were re-challenged with s.c. HuPD-L1-MC38 tumor cells at the opposite flank. Treatment naïve, wildtype mice (n = 6) were used as a control. Average tumor volume ± SEM is shown. P-value was determined using Wilcoxon rank sum test, comparing tumor sizes on day 77 between naïve and aPD-L1 + aTGF-β groups.

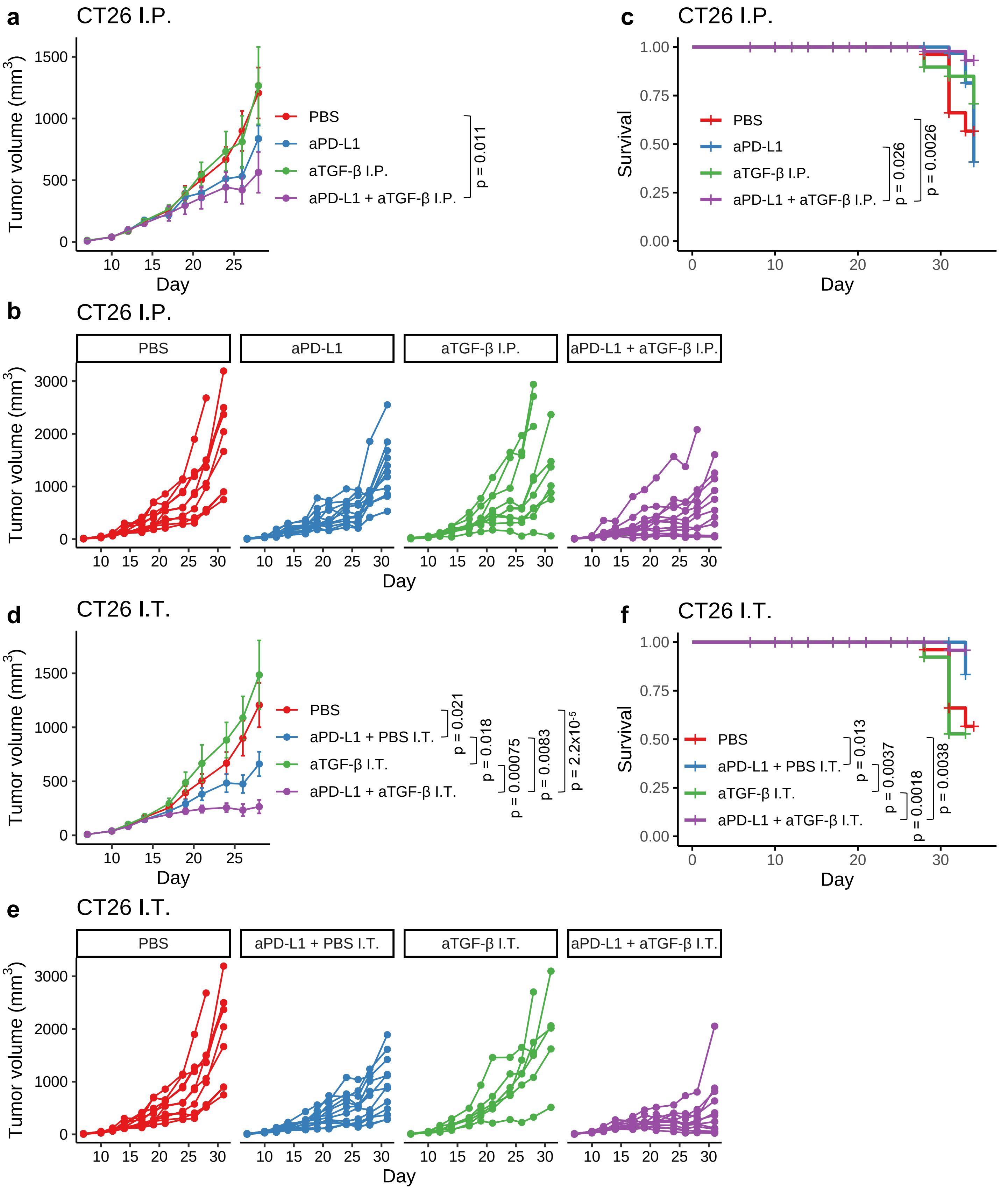


**Figure S2. Anti-PD-L1 plus anti-TGF-β in the murine tumor model CT26. (a-c)** Mice bearing CT26 tumors were dosed intraperitoneally (I.P.) two times a week for 3 weeks with PBS (n = 10), aPD-L1 (2 mg/kg; atezolizumab; n = 12), aTGF-β (10 mg/kg; 1D11; n = 10), or aPD-L1 + aTGF-β (n = 12). **(a)** Average tumor volume ± SEM is shown. P-value was determined using Wilcoxon rank sum test, using tumor sizes on day 28. **(b)** Spider plots showing CT26 tumor volume for individual mice over time. **(c)** Survival plot for the study. P-values were determined using log-rank test. **(d-f)** Mice bearing CT26 tumors were dosed two times a week for 3 weeks with PBS (I.P.; n = 10; same PBS arm as in **a-c**), aPD-L1 (2 mg/kg; atezolizumab; I.P.) + PBS (intratumorally, I.T.) (n = 12), aTGF-β (10 mg/kg; 1D11; I.T.; n = 6), or aPD-L1 (I.P.) + aTGF-β (I.T.) (n = 12). **(d)** Average tumor volume ± SEM is shown. P-values were determined using Wilcoxon rank sum test, using tumor sizes on day 28. **(e)** Spider plots showing CT26 tumor volume for individual mice over time. **(f)** Survival plot for the study. P-values were determined using log-rank test. Non-significant p-values are not shown for all panels.

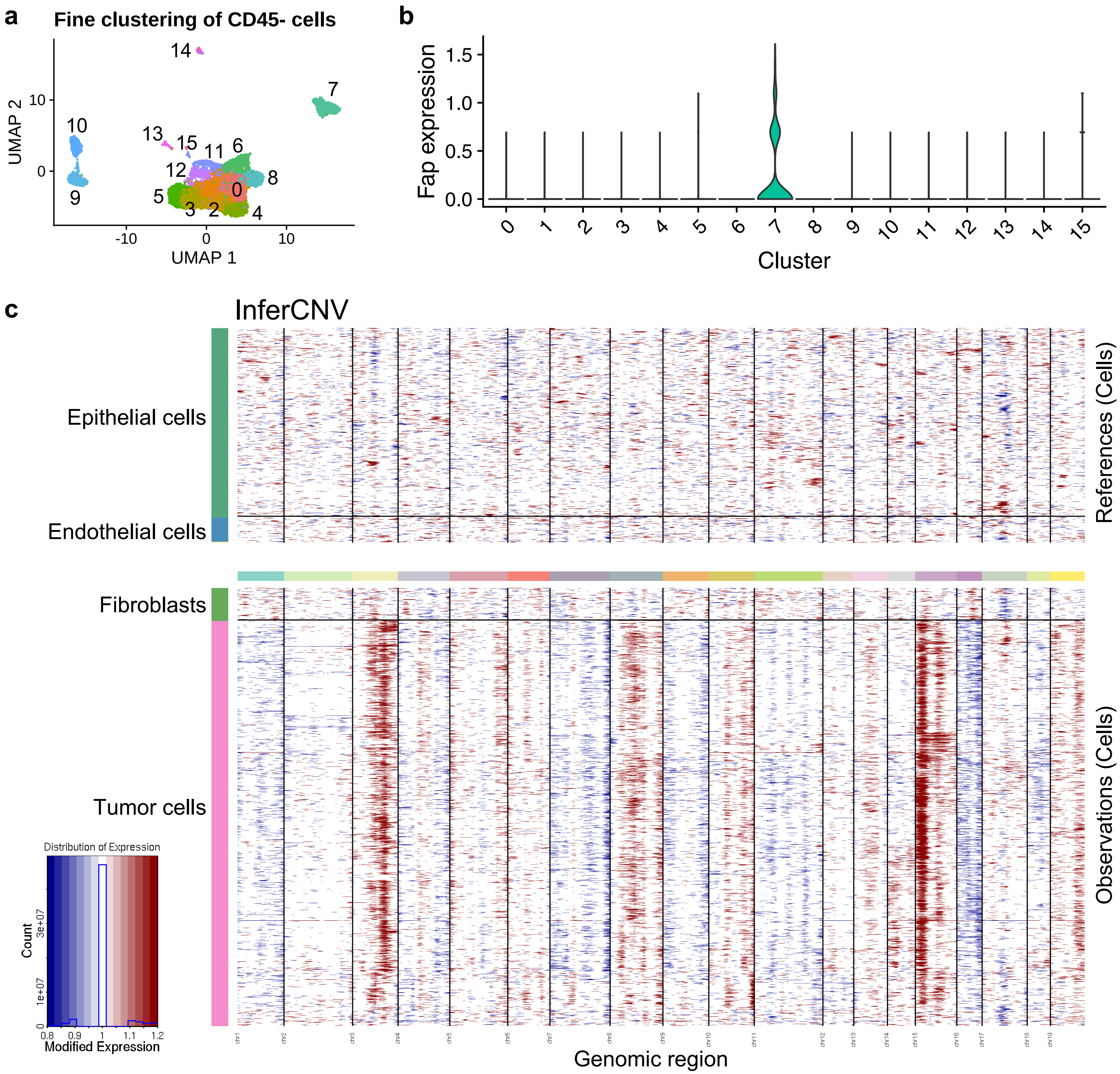


**Figure S3. CD45- cell analysis and InferCNV of anti-PD-L1 ± anti-TGF-β treated EMT6 tumor-bearing mice. (a)** Single cell transcriptomes for all CD45- cells visualized on a UMAP plot. Cell clusters are colored and numbered. **(b)** Violin plot showing *Fap* expression level in the different CD45- cell clusters. **(c)** Inferred copy number for individual cells of the different CD45- cell types (y-axis) along the mouse genome (x-axis). Regions with gain and loss of copy number are shown in red and blue, respectively.

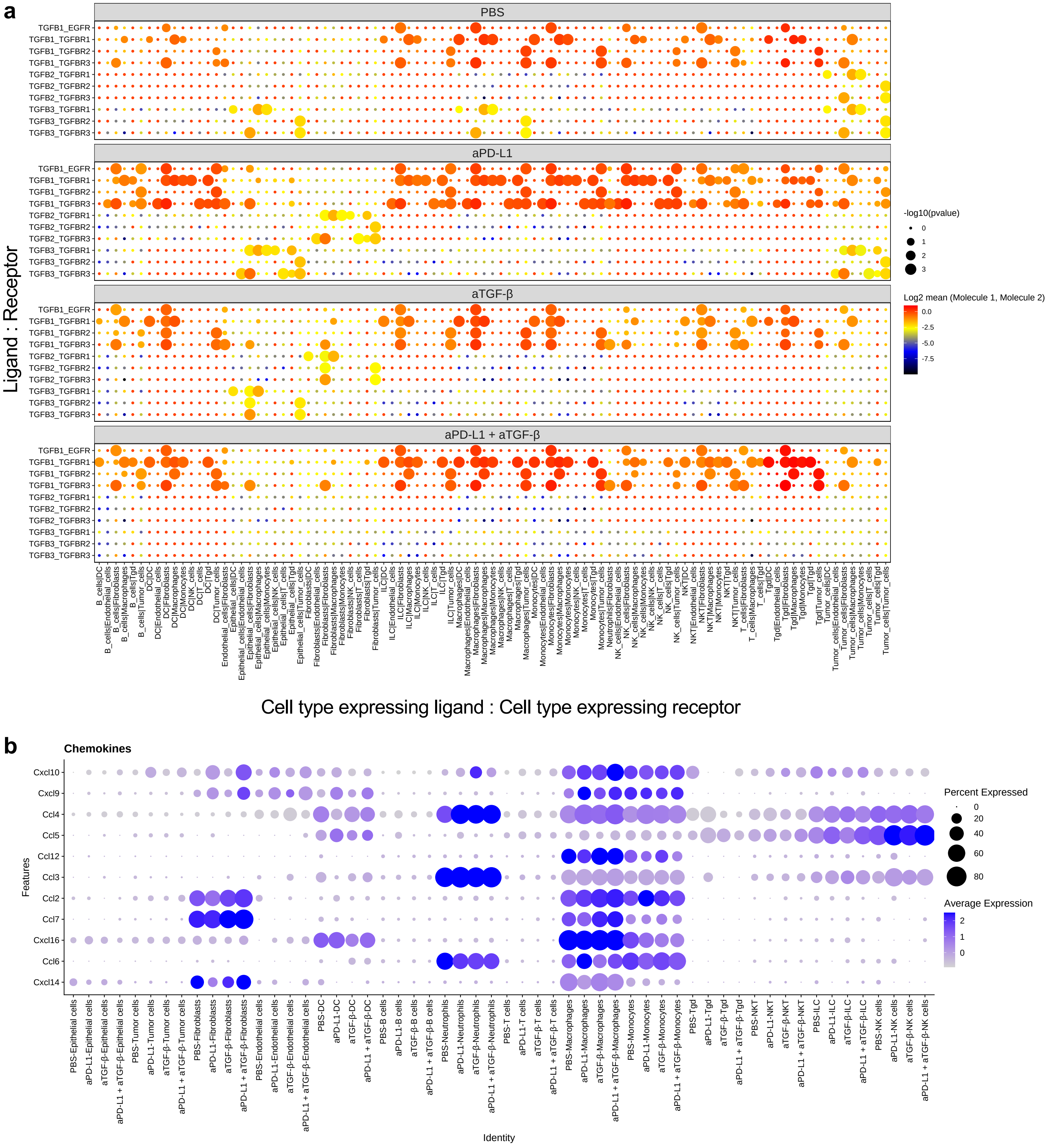

**Figure S4. CellPhoneDB and chemokine expression of anti-PD-L1 ± anti-TGF-β treated EMT6 tumor-bearing mice. (a)** Cell-cell communication network involving all TGF-β ligands and receptors, as analyzed using CellPhoneDB. The x-axis displays the cell type pairs expressing the ligand and the receptor, respectively, while the y-axis displays the corresponding ligand and receptor pairs expressed by the cell types indicated on the x-axis. The size and color of the points show the significance of the interactions and $\log_2$ mean expression values of the ligand-receptor pairs, respectively, as indicated in the key. **(b)** Dot plot showing chemokine gene expression in different cell types under different treatments. The size of the dots indicates percent cells expressing the chemokine gene, while the color of the dots indicates average gene expression level.

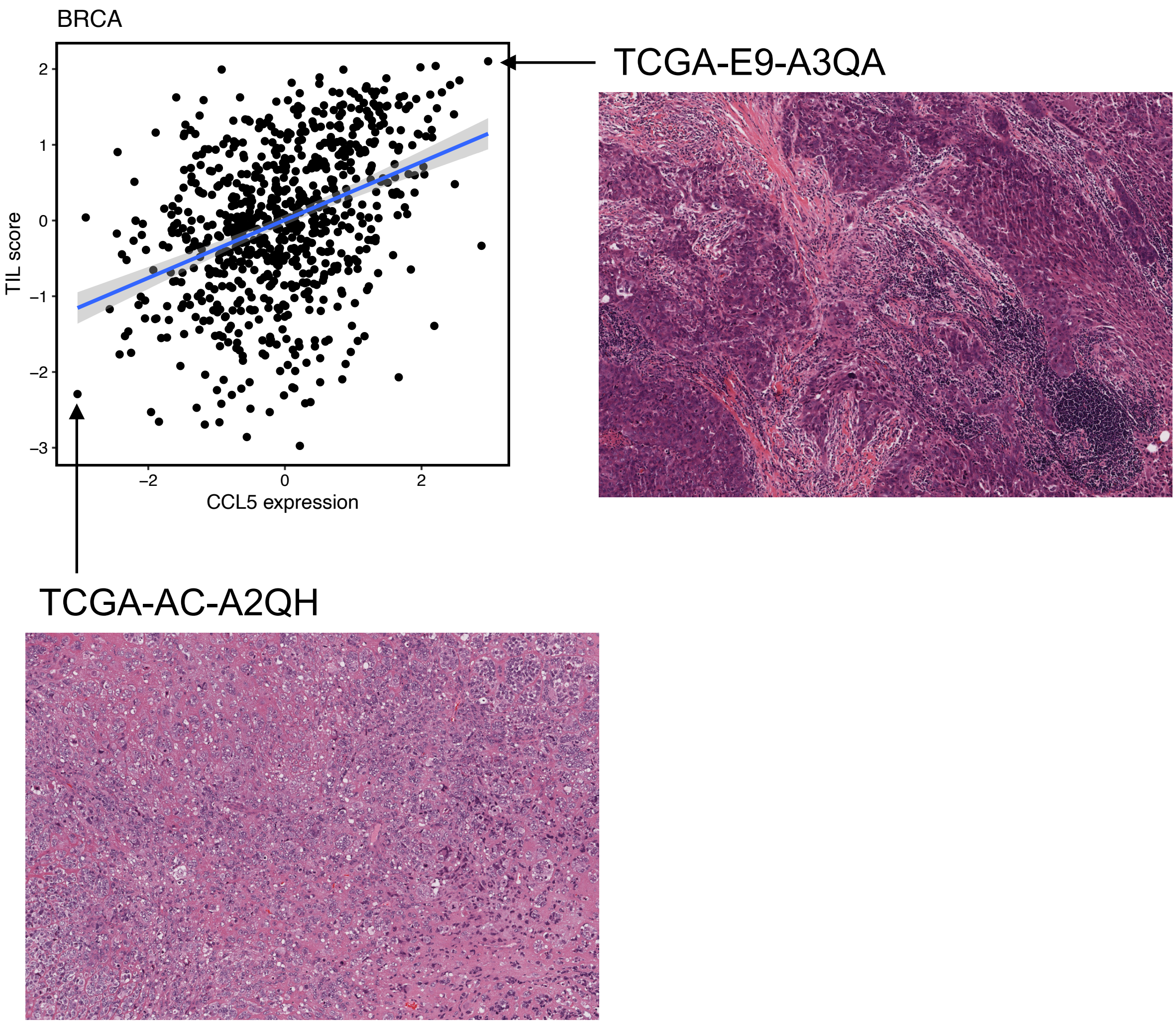


**Figure S5. *CCL5* expression correlates with TIL score.** The scatter plot shows the correlation between *CCL5* expression and pathology image-based tumor-infiltrating lymphocyte (TIL) score in breast cancer (BRCA) in TCGA. Each point represents a tumor sample. Both axes are shown in Z-score space. TIL score is inferred computationally from H&E-stained pathology images of the tumor samples. Representative images for a high TIL sample and a low TIL sample are shown.

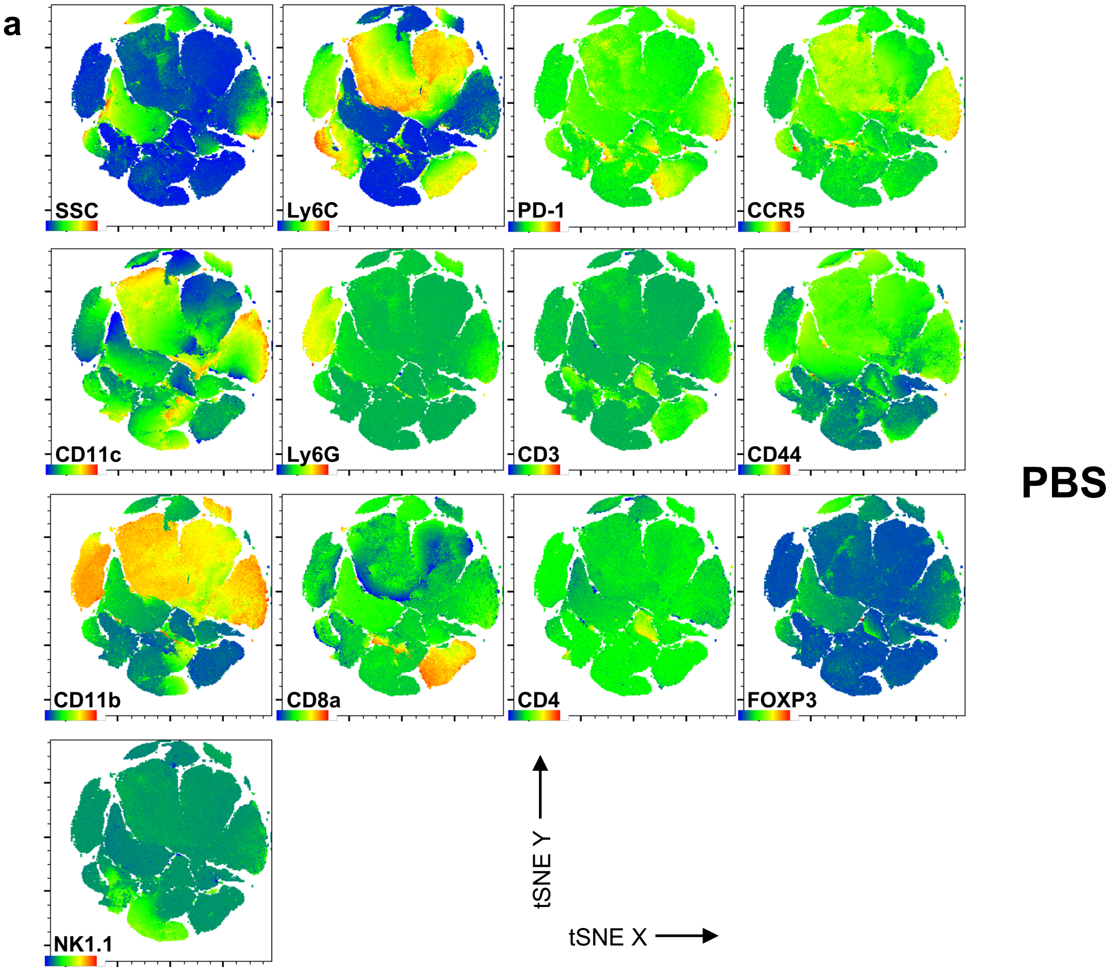


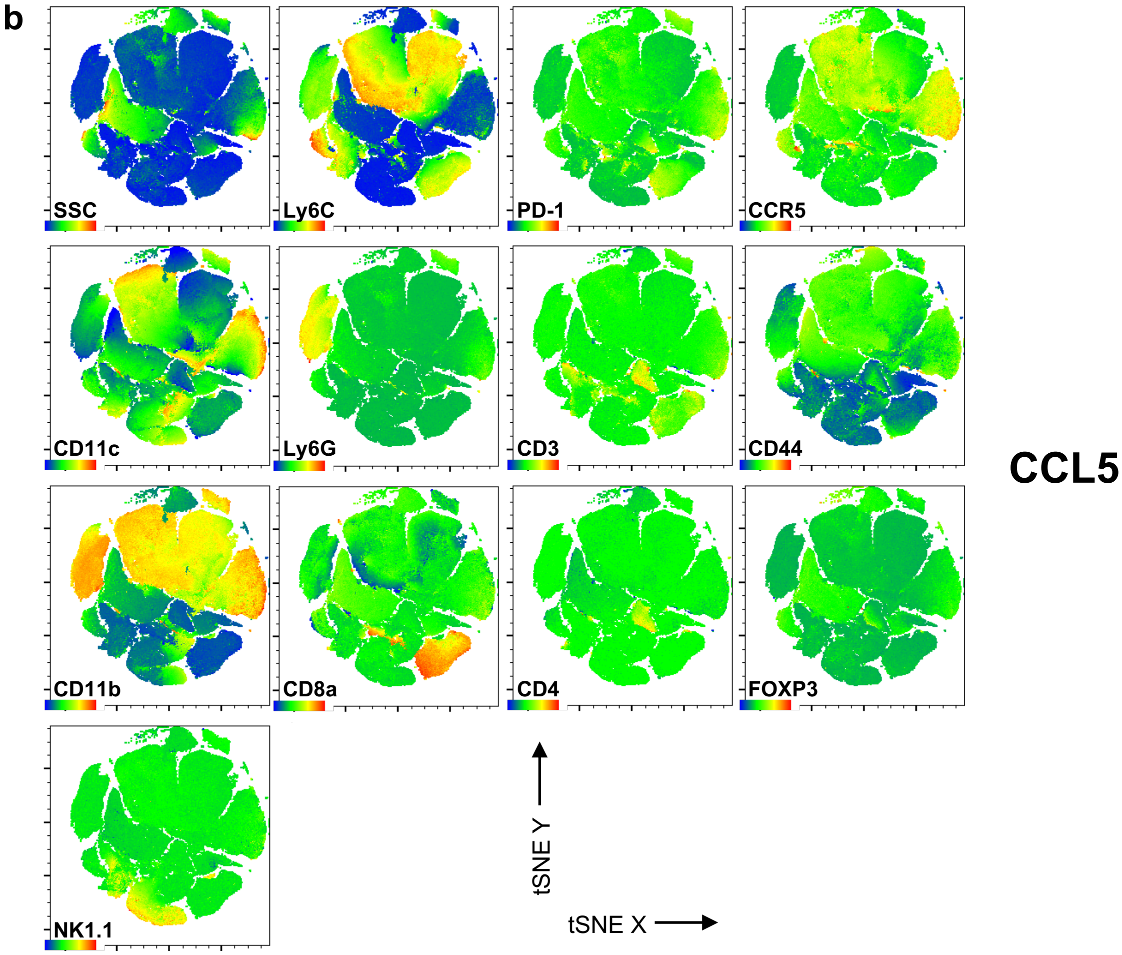


**Figure S6. Flow cytometry-based expression of key genes in tSNE identified populations.** Plots show heat maps of the indicated marker for each parameter used in tSNE analysis in PBS control **(a)** or recombinant CCL5 **(b)** treated samples.
