## Additional file 2, Table S1 for "Single cell transcriptomics reveals the effect of PD-L1/TGF-β blockade on the tumor microenvironment"

Table S1. ***Differentially expressed (DE) genes determined from the EMT6 single cell RNA-seq experiment.*** Differential expression analysis was performed within each cell type, indicated in the "cell" column. The "comparison" column indicates the treatment group comparison in which a given DE gene was observed. The columns "pct.1" and "pct.2" represent percent cells expressing the gene in the first treatment group and the second treatment group, respectively. The column "avg_logFC" represents log fold-change of the average expression between the two groups. Positive values indicate that the gene is more highly expressed in the first treatment group. P-values, as shown in the "p_val" column, were determined using Wilcoxon rank sum test. The column "p_val_adj" shows adjusted p-value, based on Bonferroni correction using all genes in the dataset.

| **gene** | **p_val** | **p_val_adj** | **avg_logFC** | **pct.1** | **pct.2** | **cell** | **comparison** |
|---|---|---|---|---|---|---|---|
| Igkc | 1.68E-14 | 2.99E-10 | 1.93 | 0.296 | 0.044 | Monocytes | aPD-L1 vs PBS |
| Ighg1 | 1.97E-12 | 3.52E-08 | 1.40 | 0.158 | 0.003 | Monocytes | aPD-L1 vs PBS |
| Wfdc17 | 5.29E-10 | 9.43E-06 | 1.30 | 0.407 | 0.259 | Macrophages | aPD-L1 vs PBS |
| AY036118 | 3.30E-08 | 5.88E-04 | 1.29 | 1 | 0.788 | NK cells | aPD-L1 vs PBS |
| AY036118 | 2.96E-133 | 5.28E-129 | 1.28 | 1 | 0.767 | T cells | aPD-L1 vs PBS |
| AY036118 | 8.25E-10 | 1.47E-05 | 1.22 | 1 | 0.792 | DC | aPD-L1 vs PBS |
| AY036118 | 3.12E-115 | 5.56E-111 | 1.22 | 0.998 | 0.707 | B cells | aPD-L1 vs PBS |
| Igfbp5 | 7.88E-12 | 1.40E-07 | 1.15 | 0.667 | 0.242 | Fibroblasts | aPD-L1 vs PBS |
| Igkc | 5.32E-24 | 9.49E-20 | 1.12 | 0.179 | 0.025 | Macrophages | aPD-L1 vs PBS |
| AY036118 | 4.28E-23 | 7.63E-19 | 1.10 | 1 | 0.71 | ILC | aPD-L1 vs PBS |
| AY036118 | 2.65E-20 | 4.72E-16 | 1.02 | 1 | 0.85 | NKT | aPD-L1 vs PBS |
| AY036118 | 1.41E-139 | 2.52E-135 | 1.01 | 0.988 | 0.904 | Neutrophils | aPD-L1 vs PBS |
| H2-Eb1 | 7.19E-41 | 1.28E-36 | 0.96 | 0.84 | 0.502 | Macrophages | aPD-L1 vs PBS |
| Rpl39 | 7.52E-12 | 1.34E-07 | 0.95 | 1 | 1 | DC | aPD-L1 vs PBS |
| AY036118 | 9.43E-34 | 1.68E-29 | 0.91 | 1 | 0.937 | Monocytes | aPD-L1 vs PBS |
| H2-Aa | 1.73E-39 | 3.08E-35 | 0.87 | 0.869 | 0.537 | Macrophages | aPD-L1 vs PBS |
| Klf2 | 3.68E-09 | 6.56E-05 | 0.86 | 0.789 | 0.5 | Epithelial cells | aPD-L1 vs PBS |
| Rps27 | 1.40E-11 | 2.50E-07 | 0.86 | 1 | 1 | DC | aPD-L1 vs PBS |
| Gm42418 | 9.69E-103 | 1.73E-98 | 0.85 | 1 | 1 | Tumor_cells | aPD-L1 vs PBS |
| Ighg1 | 3.19E-16 | 5.69E-12 | 0.83 | 0.187 | 0.009 | B cells | aPD-L1 vs PBS |
| Cd74 | 3.03E-35 | 5.41E-31 | 0.81 | 0.948 | 0.805 | Macrophages | aPD-L1 vs PBS |
| Rps27 | 2.90E-33 | 5.17E-29 | 0.81 | 0.995 | 0.94 | Monocytes | aPD-L1 vs PBS |
| Irf1 | 4.70E-19 | 8.38E-15 | 0.80 | 0.714 | 0.396 | Monocytes | aPD-L1 vs PBS |
| Rps21 | 5.25E-13 | 9.35E-09 | 0.80 | 1 | 1 | DC | aPD-L1 vs PBS |
| Cxcl9 | 1.04E-09 | 1.86E-05 | 0.79 | 0.351 | 0.192 | Macrophages | aPD-L1 vs PBS |
| Rpl39 | 2.01E-117 | 3.59E-113 | 0.77 | 1 | 1 | B cells | aPD-L1 vs PBS |
| Rps27 | 2.69E-123 | 4.79E-119 | 0.76 | 1 | 1 | B cells | aPD-L1 vs PBS |
| Rpl37 | 7.46E-12 | 1.33E-07 | 0.76 | 1 | 1 | DC | aPD-L1 vs PBS |
| Tmsb10 | 1.25E-46 | 2.23E-42 | 0.74 | 0.97 | 0.917 | Macrophages | aPD-L1 vs PBS |
| Rpl39 | 9.41E-110 | 1.68E-105 | 0.73 | 1 | 1 | T cells | aPD-L1 vs PBS |
| Rpl37a | 1.39E-11 | 2.48E-07 | 0.71 | 1 | 1 | DC | aPD-L1 vs PBS |
| Rps27 | 6.04E-108 | 1.08E-103 | 0.71 | 1 | 1 | T cells | aPD-L1 vs PBS |
| Cxcl10 | 1.45E-09 | 2.59E-05 | 0.71 | 0.414 | 0.081 | Fibroblasts | aPD-L1 vs PBS |
| H2-Ab1 | 4.38E-32 | 7.80E-28 | 0.70 | 0.858 | 0.586 | Macrophages | aPD-L1 vs PBS |
| Irf1 | 6.57E-44 | 1.17E-39 | 0.70 | 0.694 | 0.375 | Neutrophils | aPD-L1 vs PBS |
| Rpl37a | 4.14E-112 | 7.38E-108 | 0.70 | 1 | 1 | B cells | aPD-L1 vs PBS |
| Lmna | 4.96E-07 | 8.84E-03 | 0.70 | 0.748 | 0.495 | ILC | aPD-L1 vs PBS |
| Rps21 | 1.62E-113 | 2.89E-109 | 0.70 | 0.999 | 1 | B cells | aPD-L1 vs PBS |
| Rpl35a | 3.76E-11 | 6.71E-07 | 0.70 | 1 | 1 | DC | aPD-L1 vs PBS |
| Rps27 | 2.78E-11 | 4.95E-07 | 0.68 | 1 | 1 | NKT | aPD-L1 vs PBS |
| Rpl37a | 2.32E-106 | 4.14E-102 | 0.68 | 1 | 1 | T cells | aPD-L1 vs PBS |
| Rpl37a | 2.80E-36 | 4.99E-32 | 0.68 | 0.995 | 0.962 | Monocytes | aPD-L1 vs PBS |
| Rpl34 | 4.41E-10 | 7.87E-06 | 0.66 | 1 | 1 | DC | aPD-L1 vs PBS |
| Dnaja1 | 4.49E-09 | 8.00E-05 | 0.65 | 0.963 | 0.841 | ILC | aPD-L1 vs PBS |

| | | | | | | | |
|---|---|---|---|---|---|---|---|
| Rps29 | 5.09E-10 | 9.07E-06 | 0.64 | 1 | 1 | DC | aPD-L1 vs PBS |
| Crip1 | 4.22E-12 | 7.52E-08 | 0.64 | 0.866 | 0.83 | Macrophages | aPD-L1 vs PBS |
| Rps27 | 9.18E-51 | 1.64E-46 | 0.64 | 0.993 | 0.974 | Macrophages | aPD-L1 vs PBS |
| Rpl37 | 3.20E-94 | 5.71E-90 | 0.64 | 1 | 1 | B cells | aPD-L1 vs PBS |
| Rpl39 | 8.56E-26 | 1.53E-21 | 0.63 | 0.98 | 0.943 | Monocytes | aPD-L1 vs PBS |
| Rpl38 | 4.66E-10 | 8.30E-06 | 0.63 | 1 | 1 | DC | aPD-L1 vs PBS |
| AY036118 | 4.22E-40 | 7.52E-36 | 0.63 | 0.989 | 0.836 | Macrophages | aPD-L1 vs PBS |
| Hilpda | 4.69E-07 | 8.36E-03 | 0.62 | 0.515 | 0.316 | Monocytes | aPD-L1 vs PBS |
| Gm10076 | 1.18E-54 | 2.10E-50 | 0.62 | 0.913 | 0.49 | B cells | aPD-L1 vs PBS |
| Rpl37 | 1.89E-91 | 3.37E-87 | 0.61 | 1 | 1 | T cells | aPD-L1 vs PBS |
| Rps21 | 2.24E-93 | 4.00E-89 | 0.61 | 1 | 1 | T cells | aPD-L1 vs PBS |
| Irf1 | 7.48E-10 | 1.33E-05 | 0.61 | 0.678 | 0.295 | Fibroblasts | aPD-L1 vs PBS |
| Rps28 | 3.28E-75 | 5.85E-71 | 0.61 | 0.998 | 0.987 | T cells | aPD-L1 vs PBS |
| Gm10076 | 1.67E-06 | 2.99E-02 | 0.60 | 0.854 | 0.625 | DC | aPD-L1 vs PBS |
| Rpl35a | 1.67E-100 | 2.97E-96 | 0.60 | 0.999 | 0.997 | B cells | aPD-L1 vs PBS |
| Rps27 | 8.32E-12 | 1.48E-07 | 0.60 | 1 | 1 | ILC | aPD-L1 vs PBS |
| Tnfaip2 | 1.98E-26 | 3.54E-22 | 0.59 | 0.627 | 0.386 | Neutrophils | aPD-L1 vs PBS |
| Rps27 | 1.80E-63 | 3.21E-59 | 0.59 | 0.992 | 0.983 | Neutrophils | aPD-L1 vs PBS |
| Gm10076 | 3.50E-55 | 6.25E-51 | 0.59 | 0.921 | 0.588 | T cells | aPD-L1 vs PBS |
| Rpl41 | 4.18E-84 | 7.45E-80 | 0.58 | 1 | 0.997 | B cells | aPD-L1 vs PBS |
| Gm47283 | 3.09E-45 | 5.52E-41 | 0.58 | 0.861 | 0.525 | T cells | aPD-L1 vs PBS |
| Ifi27l2a | 6.98E-26 | 1.24E-21 | 0.58 | 0.716 | 0.495 | Neutrophils | aPD-L1 vs PBS |
| Lmna | 8.24E-07 | 1.47E-02 | 0.58 | 0.655 | 0.394 | NKT | aPD-L1 vs PBS |
| Rpl35 | 3.88E-69 | 6.91E-65 | 0.58 | 1 | 0.985 | T cells | aPD-L1 vs PBS |
| Rpl35 | 2.84E-73 | 5.06E-69 | 0.57 | 0.998 | 0.986 | B cells | aPD-L1 vs PBS |
| Rps29 | 1.58E-109 | 2.82E-105 | 0.57 | 1 | 1 | B cells | aPD-L1 vs PBS |
| Rpl36 | 4.58E-86 | 8.16E-82 | 0.57 | 1 | 0.997 | B cells | aPD-L1 vs PBS |
| Rpl36 | 5.78E-89 | 1.03E-84 | 0.57 | 1 | 1 | T cells | aPD-L1 vs PBS |
| Inhba | 1.17E-07 | 2.09E-03 | 0.57 | 0.112 | 0.036 | Macrophages | aPD-L1 vs PBS |
| Rpl37a | 7.03E-46 | 1.25E-41 | 0.56 | 0.954 | 0.914 | Neutrophils | aPD-L1 vs PBS |
| Rps29 | 7.88E-99 | 1.40E-94 | 0.56 | 1 | 1 | T cells | aPD-L1 vs PBS |
| Rpl38 | 5.49E-82 | 9.79E-78 | 0.56 | 0.999 | 0.991 | B cells | aPD-L1 vs PBS |
| Rpl27 | 7.77E-08 | 1.38E-03 | 0.56 | 0.927 | 0.521 | DC | aPD-L1 vs PBS |
| Marcks | 7.24E-09 | 1.29E-04 | 0.56 | 0.897 | 0.745 | Fibroblasts | aPD-L1 vs PBS |
| Atf3 | 2.02E-06 | 3.59E-02 | 0.56 | 0.77 | 0.497 | Fibroblasts | aPD-L1 vs PBS |
| Timp2 | 5.12E-09 | 9.13E-05 | 0.55 | 0.954 | 0.866 | Fibroblasts | aPD-L1 vs PBS |
| Rps24 | 5.45E-44 | 9.71E-40 | 0.55 | 0.965 | 0.923 | Neutrophils | aPD-L1 vs PBS |
| Rpl38 | 1.46E-77 | 2.59E-73 | 0.55 | 1 | 0.993 | T cells | aPD-L1 vs PBS |
| Rps29 | 2.01E-56 | 3.58E-52 | 0.55 | 0.995 | 0.977 | Neutrophils | aPD-L1 vs PBS |
| Rps28 | 1.17E-07 | 2.08E-03 | 0.55 | 1 | 0.938 | DC | aPD-L1 vs PBS |
| Ly6a | 8.24E-17 | 1.47E-12 | 0.55 | 0.638 | 0.42 | Macrophages | aPD-L1 vs PBS |
| Cd274 | 7.33E-30 | 1.31E-25 | 0.55 | 0.735 | 0.475 | Neutrophils | aPD-L1 vs PBS |
| Malat1 | 2.46E-09 | 4.38E-05 | 0.54 | 1 | 0.991 | ILC | aPD-L1 vs PBS |
| Rpl36a | 1.22E-58 | 2.18E-54 | 0.54 | 0.991 | 0.954 | B cells | aPD-L1 vs PBS |
| Irf1 | 1.24E-31 | 2.21E-27 | 0.54 | 0.537 | 0.22 | Macrophages | aPD-L1 vs PBS |
| Irgm1 | 4.97E-35 | 8.87E-31 | 0.54 | 0.508 | 0.226 | Neutrophils | aPD-L1 vs PBS |
| Gm42418 | 5.66E-76 | 1.01E-71 | 0.54 | 1 | 1 | Neutrophils | aPD-L1 vs PBS |
| Rpl35 | 3.07E-07 | 5.48E-03 | 0.54 | 1 | 0.979 | DC | aPD-L1 vs PBS |
| Tmsb10 | 7.79E-20 | 1.39E-15 | 0.54 | 1 | 0.981 | Monocytes | aPD-L1 vs PBS |
| Rpl37a | 6.44E-14 | 1.15E-09 | 0.54 | 1 | 1 | ILC | aPD-L1 vs PBS |
| Rps21 | 5.75E-24 | 1.03E-19 | 0.53 | 0.985 | 0.943 | Monocytes | aPD-L1 vs PBS |
| B2m | 1.54E-08 | 2.74E-04 | 0.53 | 0.977 | 0.882 | Epithelial cells | aPD-L1 vs PBS |
| Cxcl10 | 1.16E-39 | 2.06E-35 | 0.53 | 0.227 | 0.051 | Tumor_cells | aPD-L1 vs PBS |
| Lars2 | 2.96E-13 | 5.27E-09 | 0.53 | 0.842 | 0.608 | Monocytes | aPD-L1 vs PBS |
| Gm10076 | 1.19E-11 | 2.13E-07 | 0.53 | 0.874 | 0.495 | ILC | aPD-L1 vs PBS |

| | | | | | | | |
|---|---|---|---|---|---|---|---|
| Tnf | 1.06E-17 | 1.89E-13 | 0.53 | 0.309 | 0.137 | Neutrophils | aPD-L1 vs PBS |
| Rps21 | 3.82E-11 | 6.80E-07 | 0.53 | 1 | 1 | NKT | aPD-L1 vs PBS |
| Rpl41 | 1.14E-78 | 2.03E-74 | 0.53 | 1 | 1 | T cells | aPD-L1 vs PBS |
| Uba52 | 5.26E-46 | 9.38E-42 | 0.53 | 0.786 | 0.399 | T cells | aPD-L1 vs PBS |
| Rpl36 | 2.25E-08 | 4.02E-04 | 0.53 | 1 | 1 | DC | aPD-L1 vs PBS |
| Rpl37 | 1.12E-24 | 2.00E-20 | 0.52 | 0.99 | 0.94 | Monocytes | aPD-L1 vs PBS |
| Rps29 | 7.35E-12 | 1.31E-07 | 0.52 | 1 | 1 | ILC | aPD-L1 vs PBS |
| Igkc | 5.38E-12 | 9.58E-08 | 0.52 | 0.309 | 0.019 | NKT | aPD-L1 vs PBS |
| Rps28 | 1.31E-57 | 2.34E-53 | 0.52 | 0.997 | 0.949 | B cells | aPD-L1 vs PBS |
| Rpl41 | 1.88E-08 | 3.35E-04 | 0.51 | 1 | 1 | DC | aPD-L1 vs PBS |
| Rpl39 | 3.11E-08 | 5.54E-04 | 0.51 | 1 | 0.994 | NKT | aPD-L1 vs PBS |
| Rplp2 | 2.99E-07 | 5.33E-03 | 0.51 | 1 | 1 | DC | aPD-L1 vs PBS |
| Rps21 | 4.03E-11 | 7.18E-07 | 0.51 | 1 | 0.981 | ILC | aPD-L1 vs PBS |
| Rpl35a | 8.44E-77 | 1.51E-72 | 0.51 | 1 | 0.998 | T cells | aPD-L1 vs PBS |
| Uba52 | 2.21E-06 | 3.94E-02 | 0.50 | 0.659 | 0.188 | DC | aPD-L1 vs PBS |
| Rpl39 | 1.10E-40 | 1.96E-36 | 0.50 | 0.938 | 0.914 | Neutrophils | aPD-L1 vs PBS |
| Samsn1 | 2.58E-23 | 4.59E-19 | 0.50 | 0.558 | 0.262 | B cells | aPD-L1 vs PBS |
| Rpl39 | 3.32E-33 | 5.92E-29 | 0.50 | 0.993 | 0.98 | Macrophages | aPD-L1 vs PBS |
| Rpl39 | 7.05E-09 | 1.26E-04 | 0.50 | 1 | 1 | ILC | aPD-L1 vs PBS |
| Nr4a1 | 3.85E-09 | 6.87E-05 | 0.49 | 0.617 | 0.386 | Monocytes | aPD-L1 vs PBS |
| Ubc | 4.29E-43 | 7.64E-39 | 0.49 | 0.855 | 0.718 | Tumor_cells | aPD-L1 vs PBS |
| Arg2 | 1.85E-14 | 3.31E-10 | 0.49 | 0.643 | 0.326 | Monocytes | aPD-L1 vs PBS |
| Rpl37a | 9.28E-57 | 1.65E-52 | 0.49 | 1 | 1 | Macrophages | aPD-L1 vs PBS |
| Plaur | 1.22E-13 | 2.17E-09 | 0.49 | 0.719 | 0.43 | Monocytes | aPD-L1 vs PBS |
| Gm42418 | 9.53E-10 | 1.70E-05 | 0.49 | 1 | 1 | Fibroblasts | aPD-L1 vs PBS |
| Fosb | 3.34E-37 | 5.95E-33 | 0.49 | 0.701 | 0.465 | Tumor_cells | aPD-L1 vs PBS |
| Rpl34 | 2.85E-82 | 5.08E-78 | 0.49 | 1 | 1 | B cells | aPD-L1 vs PBS |
| Rpl35 | 4.26E-10 | 7.60E-06 | 0.49 | 0.978 | 0.869 | NKT | aPD-L1 vs PBS |
| Rpl36a | 2.84E-51 | 5.07E-47 | 0.49 | 1 | 0.978 | T cells | aPD-L1 vs PBS |
| H2-DMb1 | 1.32E-22 | 2.35E-18 | 0.49 | 0.638 | 0.356 | Macrophages | aPD-L1 vs PBS |
| Uba52 | 1.54E-39 | 2.75E-35 | 0.49 | 0.718 | 0.279 | B cells | aPD-L1 vs PBS |
| Rps20 | 2.18E-36 | 3.89E-32 | 0.49 | 0.948 | 0.918 | Neutrophils | aPD-L1 vs PBS |
| Lars2 | 2.27E-26 | 4.05E-22 | 0.49 | 0.699 | 0.379 | T cells | aPD-L1 vs PBS |
| Rpl37a | 7.79E-11 | 1.39E-06 | 0.48 | 1 | 0.994 | NKT | aPD-L1 vs PBS |
| Rpl35a | 8.59E-19 | 1.53E-14 | 0.48 | 0.99 | 0.934 | Monocytes | aPD-L1 vs PBS |
| Rsad2 | 9.17E-10 | 1.64E-05 | 0.48 | 0.399 | 0.232 | Macrophages | aPD-L1 vs PBS |
| Rps29 | 1.94E-27 | 3.46E-23 | 0.48 | 1 | 1 | Monocytes | aPD-L1 vs PBS |
| Rpl36 | 3.78E-09 | 6.73E-05 | 0.48 | 1 | 0.981 | NKT | aPD-L1 vs PBS |
| Lars2 | 2.19E-31 | 3.91E-27 | 0.48 | 0.571 | 0.283 | Neutrophils | aPD-L1 vs PBS |
| Rpl37 | 7.08E-12 | 1.26E-07 | 0.48 | 1 | 0.991 | ILC | aPD-L1 vs PBS |
| Fcgr4 | 9.75E-17 | 1.74E-12 | 0.47 | 0.69 | 0.496 | Macrophages | aPD-L1 vs PBS |
| Rps21 | 1.06E-38 | 1.89E-34 | 0.47 | 1 | 0.982 | Macrophages | aPD-L1 vs PBS |
| Rpl27 | 3.31E-35 | 5.91E-31 | 0.47 | 0.852 | 0.551 | T cells | aPD-L1 vs PBS |
| Cd274 | 8.71E-11 | 1.55E-06 | 0.47 | 0.628 | 0.373 | Monocytes | aPD-L1 vs PBS |
| Gm10076 | 5.41E-37 | 9.64E-33 | 0.47 | 0.821 | 0.558 | Macrophages | aPD-L1 vs PBS |
| Gm10076 | 8.50E-23 | 1.52E-18 | 0.47 | 0.816 | 0.446 | Monocytes | aPD-L1 vs PBS |
| mt-Atp8 | 2.95E-11 | 5.26E-07 | 0.47 | 0.568 | 0.2 | NKT | aPD-L1 vs PBS |
| Rps29 | 1.34E-06 | 2.39E-02 | 0.46 | 1 | 0.994 | NKT | aPD-L1 vs PBS |
| Gm42418 | 1.54E-45 | 2.74E-41 | 0.46 | 1 | 1 | T cells | aPD-L1 vs PBS |
| Rps28 | 8.56E-18 | 1.53E-13 | 0.46 | 0.944 | 0.848 | Monocytes | aPD-L1 vs PBS |
| Rpl35a | 1.26E-09 | 2.24E-05 | 0.46 | 1 | 0.972 | ILC | aPD-L1 vs PBS |
| Lars2 | 2.10E-12 | 3.75E-08 | 0.46 | 0.728 | 0.59 | Macrophages | aPD-L1 vs PBS |
| Rpl30 | 1.89E-06 | 3.36E-02 | 0.46 | 1 | 0.979 | DC | aPD-L1 vs PBS |
| AW112010 | 1.60E-09 | 2.85E-05 | 0.46 | 0.709 | 0.544 | Macrophages | aPD-L1 vs PBS |
| Ccl4 | 3.35E-13 | 5.97E-09 | 0.45 | 0.758 | 0.591 | Neutrophils | aPD-L1 vs PBS |

| | | | | | | | |
|---|---|---|---|---|---|---|---|
| mt-Atp8 | 1.20E-37 | 2.15E-33 | 0.45 | 0.671 | 0.288 | T cells | aPD-L1 vs PBS |
| Uba52 | 1.73E-08 | 3.08E-04 | 0.45 | 0.547 | 0.262 | NKT | aPD-L1 vs PBS |
| Scara5 | 1.33E-08 | 2.36E-04 | 0.45 | 0.563 | 0.228 | Fibroblasts | aPD-L1 vs PBS |
| Hsp90ab1 | 4.39E-08 | 7.82E-04 | 0.45 | 1 | 1 | ILC | aPD-L1 vs PBS |
| Rpl35a | 7.79E-08 | 1.39E-03 | 0.45 | 1 | 0.988 | NKT | aPD-L1 vs PBS |
| Gm42418 | 4.24E-48 | 7.55E-44 | 0.44 | 1 | 1 | B cells | aPD-L1 vs PBS |
| Rpl38 | 4.91E-10 | 8.75E-06 | 0.44 | 1 | 0.991 | ILC | aPD-L1 vs PBS |
| Atp5e | 1.71E-32 | 3.05E-28 | 0.44 | 0.867 | 0.513 | B cells | aPD-L1 vs PBS |
| Igkc | 1.15E-56 | 2.05E-52 | 0.44 | 0.282 | 0.015 | Neutrophils | aPD-L1 vs PBS |
| Rpl36 | 1.45E-10 | 2.59E-06 | 0.44 | 1 | 1 | ILC | aPD-L1 vs PBS |
| Gbp2b | 2.93E-24 | 5.22E-20 | 0.44 | 0.476 | 0.293 | Tumor_cells | aPD-L1 vs PBS |
| Sub1 | 1.25E-09 | 2.22E-05 | 0.44 | 0.885 | 0.625 | NKT | aPD-L1 vs PBS |
| Ifrd1 | 4.31E-10 | 7.69E-06 | 0.44 | 0.867 | 0.623 | Monocytes | aPD-L1 vs PBS |
| Rps15a | 8.92E-35 | 1.59E-30 | 0.44 | 0.939 | 0.905 | Neutrophils | aPD-L1 vs PBS |
| Emilin2 | 8.12E-07 | 1.45E-02 | 0.44 | 0.69 | 0.45 | Fibroblasts | aPD-L1 vs PBS |
| Rpl38 | 2.84E-18 | 5.07E-14 | 0.43 | 0.964 | 0.956 | Monocytes | aPD-L1 vs PBS |
| Rpl36 | 9.54E-17 | 1.70E-12 | 0.43 | 0.98 | 0.927 | Monocytes | aPD-L1 vs PBS |
| Rpl22l1 | 9.05E-12 | 1.61E-07 | 0.43 | 0.745 | 0.513 | Monocytes | aPD-L1 vs PBS |
| H2-T23 | 5.75E-28 | 1.03E-23 | 0.43 | 0.724 | 0.44 | Macrophages | aPD-L1 vs PBS |
| Rpl37 | 1.45E-08 | 2.58E-04 | 0.43 | 1 | 0.981 | NKT | aPD-L1 vs PBS |
| Cd274 | 7.23E-17 | 1.29E-12 | 0.43 | 0.418 | 0.199 | Macrophages | aPD-L1 vs PBS |
| Iigp1 | 6.94E-37 | 1.24E-32 | 0.43 | 0.331 | 0.12 | Tumor_cells | aPD-L1 vs PBS |
| Tmsb10 | 5.05E-52 | 9.01E-48 | 0.43 | 1 | 0.998 | T cells | aPD-L1 vs PBS |
| Selenom | 7.25E-10 | 1.29E-05 | 0.43 | 0.47 | 0.316 | Macrophages | aPD-L1 vs PBS |
| Klf2 | 1.14E-06 | 2.04E-02 | 0.43 | 0.713 | 0.409 | Fibroblasts | aPD-L1 vs PBS |
| Sec61g | 2.97E-34 | 5.30E-30 | 0.43 | 0.701 | 0.276 | B cells | aPD-L1 vs PBS |
| Lst1 | 7.00E-13 | 1.25E-08 | 0.43 | 0.739 | 0.631 | Macrophages | aPD-L1 vs PBS |
| Atf3 | 1.17E-08 | 2.09E-04 | 0.43 | 0.554 | 0.231 | NKT | aPD-L1 vs PBS |
| Sub1 | 1.05E-22 | 1.88E-18 | 0.42 | 0.933 | 0.709 | B cells | aPD-L1 vs PBS |
| Rplp2 | 5.22E-57 | 9.31E-53 | 0.42 | 1 | 1 | B cells | aPD-L1 vs PBS |
| Htra3 | 1.26E-06 | 2.25E-02 | 0.42 | 0.874 | 0.685 | Fibroblasts | aPD-L1 vs PBS |
| Rps21 | 1.13E-27 | 2.01E-23 | 0.42 | 0.902 | 0.904 | Neutrophils | aPD-L1 vs PBS |
| Gm47283 | 1.19E-15 | 2.11E-11 | 0.42 | 0.601 | 0.389 | Macrophages | aPD-L1 vs PBS |
| Rpl38 | 2.54E-37 | 4.52E-33 | 0.42 | 0.996 | 0.987 | Macrophages | aPD-L1 vs PBS |
| Irf1 | 1.82E-39 | 3.24E-35 | 0.42 | 0.579 | 0.334 | Tumor_cells | aPD-L1 vs PBS |
| Gbp2 | 3.58E-18 | 6.39E-14 | 0.42 | 0.586 | 0.468 | Tumor_cells | aPD-L1 vs PBS |
| B2m | 4.74E-50 | 8.46E-46 | 0.42 | 1 | 0.998 | Neutrophils | aPD-L1 vs PBS |
| Rpl34 | 1.44E-65 | 2.56E-61 | 0.42 | 1 | 1 | T cells | aPD-L1 vs PBS |
| Mdm2 | 7.44E-09 | 1.33E-04 | 0.42 | 0.51 | 0.291 | Monocytes | aPD-L1 vs PBS |
| Gbp2 | 9.16E-29 | 1.63E-24 | 0.41 | 0.551 | 0.266 | Neutrophils | aPD-L1 vs PBS |
| Dusp10 | 2.66E-18 | 4.75E-14 | 0.41 | 0.701 | 0.453 | T cells | aPD-L1 vs PBS |
| Gbp2b | 1.69E-25 | 3.02E-21 | 0.41 | 0.451 | 0.169 | Macrophages | aPD-L1 vs PBS |
| Zbp1 | 5.06E-27 | 9.03E-23 | 0.41 | 0.468 | 0.208 | Neutrophils | aPD-L1 vs PBS |
| Snrpg | 2.82E-30 | 5.03E-26 | 0.41 | 0.578 | 0.246 | T cells | aPD-L1 vs PBS |
| Ly6a | 2.25E-13 | 4.01E-09 | 0.41 | 0.718 | 0.639 | Tumor_cells | aPD-L1 vs PBS |
| Acod1 | 1.46E-19 | 2.61E-15 | 0.41 | 0.377 | 0.148 | Macrophages | aPD-L1 vs PBS |
| Dennd4a | 1.11E-10 | 1.98E-06 | 0.41 | 0.51 | 0.256 | Monocytes | aPD-L1 vs PBS |
| Rps28 | 2.18E-31 | 3.89E-27 | 0.41 | 0.981 | 0.946 | Macrophages | aPD-L1 vs PBS |
| Ubc | 3.69E-15 | 6.57E-11 | 0.41 | 0.858 | 0.719 | Macrophages | aPD-L1 vs PBS |
| H2-D1 | 6.39E-37 | 1.14E-32 | 0.41 | 1 | 0.999 | Neutrophils | aPD-L1 vs PBS |
| Neat1 | 3.80E-18 | 6.77E-14 | 0.41 | 0.746 | 0.639 | Tumor_cells | aPD-L1 vs PBS |
| Slc7a11 | 2.77E-18 | 4.94E-14 | 0.41 | 0.721 | 0.532 | Neutrophils | aPD-L1 vs PBS |
| Tnfaip2 | 3.10E-13 | 5.52E-09 | 0.41 | 0.495 | 0.193 | Monocytes | aPD-L1 vs PBS |
| Rpl27 | 4.29E-27 | 7.64E-23 | 0.41 | 0.822 | 0.493 | B cells | aPD-L1 vs PBS |
| mt-Atp8 | 2.00E-27 | 3.57E-23 | 0.41 | 0.632 | 0.291 | B cells | aPD-L1 vs PBS |

| | | | | | | | |
|---|---|---|---|---|---|---|---|
| Hspa8 | 1.23E-06 | 2.20E-02 | 0.40 | 1 | 0.981 | ILC | aPD-L1 vs PBS |
| mt-Nd3 | 5.16E-23 | 9.19E-19 | 0.40 | 0.571 | 0.288 | T cells | aPD-L1 vs PBS |
| Rps29 | 2.96E-43 | 5.28E-39 | 0.40 | 1 | 1 | Macrophages | aPD-L1 vs PBS |
| Rpl35 | 1.02E-06 | 1.83E-02 | 0.40 | 0.978 | 0.916 | ILC | aPD-L1 vs PBS |
| mt-Atp8 | 1.81E-26 | 3.22E-22 | 0.40 | 0.649 | 0.375 | Macrophages | aPD-L1 vs PBS |
| Man1a | 2.75E-06 | 4.90E-02 | 0.40 | 0.655 | 0.403 | Fibroblasts | aPD-L1 vs PBS |
| H2-K1 | 1.36E-38 | 2.42E-34 | 0.40 | 0.998 | 0.989 | Neutrophils | aPD-L1 vs PBS |
| Rpl37 | 2.54E-35 | 4.53E-31 | 0.40 | 0.996 | 0.996 | Macrophages | aPD-L1 vs PBS |
| Ubc | 1.11E-12 | 1.98E-08 | 0.40 | 0.934 | 0.826 | Monocytes | aPD-L1 vs PBS |
| Serpina3g | 4.55E-19 | 8.11E-15 | 0.39 | 0.272 | 0.083 | Macrophages | aPD-L1 vs PBS |
| Malat1 | 1.78E-09 | 3.18E-05 | 0.39 | 1 | 1 | NKT | aPD-L1 vs PBS |
| H2-D1 | 1.63E-09 | 2.90E-05 | 0.39 | 1 | 1 | Epithelial cells | aPD-L1 vs PBS |
| Rpl32 | 1.08E-16 | 1.93E-12 | 0.39 | 0.9 | 0.912 | Neutrophils | aPD-L1 vs PBS |
| Tnfaip3 | 4.12E-12 | 7.34E-08 | 0.39 | 0.602 | 0.304 | Monocytes | aPD-L1 vs PBS |
| Ccl5 | 7.01E-08 | 1.25E-03 | 0.39 | 0.194 | 0.083 | Macrophages | aPD-L1 vs PBS |
| Cd36 | 4.84E-12 | 8.64E-08 | 0.39 | 0.347 | 0.17 | Macrophages | aPD-L1 vs PBS |
| S100a8 | 2.79E-07 | 4.97E-03 | 0.39 | 0.931 | 0.935 | Neutrophils | aPD-L1 vs PBS |
| Igkc | 1.07E-36 | 1.91E-32 | 0.39 | 0.299 | 0.002 | T cells | aPD-L1 vs PBS |
| Tnfaip2 | 2.13E-31 | 3.79E-27 | 0.39 | 0.317 | 0.072 | Macrophages | aPD-L1 vs PBS |
| Hk3 | 2.76E-18 | 4.92E-14 | 0.39 | 0.549 | 0.325 | Macrophages | aPD-L1 vs PBS |
| Samhd1 | 2.45E-14 | 4.37E-10 | 0.39 | 0.567 | 0.361 | Macrophages | aPD-L1 vs PBS |
| Rplp2 | 1.47E-55 | 2.62E-51 | 0.39 | 1 | 1 | T cells | aPD-L1 vs PBS |
| Tnfaip3 | 2.09E-21 | 3.73E-17 | 0.38 | 0.437 | 0.178 | Macrophages | aPD-L1 vs PBS |
| Gm47283 | 5.27E-20 | 9.40E-16 | 0.38 | 0.842 | 0.587 | B cells | aPD-L1 vs PBS |
| Rpl41 | 4.37E-19 | 7.78E-15 | 0.38 | 0.995 | 0.987 | Monocytes | aPD-L1 vs PBS |
| Tmsb10 | 1.06E-31 | 1.88E-27 | 0.38 | 0.984 | 0.972 | B cells | aPD-L1 vs PBS |
| Gbp4 | 1.02E-23 | 1.81E-19 | 0.38 | 0.22 | 0.052 | Neutrophils | aPD-L1 vs PBS |
| Rps28 | 2.05E-06 | 3.65E-02 | 0.38 | 0.985 | 0.916 | ILC | aPD-L1 vs PBS |
| mt-Nd3 | 4.75E-25 | 8.47E-21 | 0.38 | 0.613 | 0.279 | B cells | aPD-L1 vs PBS |
| Malt1 | 1.43E-09 | 2.56E-05 | 0.38 | 0.622 | 0.361 | Monocytes | aPD-L1 vs PBS |
| Rplp1 | 7.09E-26 | 1.26E-21 | 0.38 | 0.973 | 0.965 | Neutrophils | aPD-L1 vs PBS |
| Egr1 | 3.29E-22 | 5.87E-18 | 0.38 | 0.747 | 0.649 | Tumor_cells | aPD-L1 vs PBS |
| Nr4a3 | 4.87E-07 | 8.69E-03 | 0.38 | 0.424 | 0.162 | NKT | aPD-L1 vs PBS |
| Rpl35a | 2.18E-22 | 3.88E-18 | 0.38 | 0.9 | 0.904 | Neutrophils | aPD-L1 vs PBS |
| mt-Atp8 | 1.54E-14 | 2.74E-10 | 0.37 | 0.597 | 0.272 | Monocytes | aPD-L1 vs PBS |
| Rpl30 | 4.57E-54 | 8.15E-50 | 0.37 | 0.999 | 1 | B cells | aPD-L1 vs PBS |
| H2-K1 | 3.24E-07 | 5.77E-03 | 0.37 | 0.992 | 0.956 | Epithelial cells | aPD-L1 vs PBS |
| Zfand5 | 3.08E-31 | 5.50E-27 | 0.37 | 0.655 | 0.473 | Tumor_cells | aPD-L1 vs PBS |
| Irf1 | 3.85E-19 | 6.86E-15 | 0.37 | 0.641 | 0.375 | T cells | aPD-L1 vs PBS |
| Samhd1 | 1.75E-14 | 3.12E-10 | 0.37 | 0.65 | 0.476 | Neutrophils | aPD-L1 vs PBS |
| Rps15a | 4.35E-60 | 7.75E-56 | 0.37 | 0.999 | 1 | B cells | aPD-L1 vs PBS |
| Nfkbia | 7.40E-10 | 1.32E-05 | 0.37 | 0.908 | 0.807 | Monocytes | aPD-L1 vs PBS |
| Rpl35a | 8.17E-25 | 1.46E-20 | 0.37 | 0.978 | 0.981 | Macrophages | aPD-L1 vs PBS |
| Rps19 | 9.75E-20 | 1.74E-15 | 0.37 | 0.914 | 0.909 | Neutrophils | aPD-L1 vs PBS |
| Ifrd1 | 2.43E-09 | 4.32E-05 | 0.37 | 0.597 | 0.445 | Macrophages | aPD-L1 vs PBS |
| Lars2 | 3.72E-22 | 6.62E-18 | 0.37 | 0.612 | 0.276 | B cells | aPD-L1 vs PBS |
| Dcn | 1.19E-08 | 2.12E-04 | 0.37 | 0.191 | 0.106 | Tumor_cells | aPD-L1 vs PBS |
| Rpl23 | 1.00E-23 | 1.79E-19 | 0.37 | 0.968 | 0.965 | Neutrophils | aPD-L1 vs PBS |
| Atp5e | 3.96E-26 | 7.06E-22 | 0.36 | 0.918 | 0.849 | Macrophages | aPD-L1 vs PBS |
| Rpl22l1 | 2.41E-24 | 4.30E-20 | 0.36 | 0.966 | 0.909 | B cells | aPD-L1 vs PBS |
| Rpl35 | 3.34E-24 | 5.95E-20 | 0.36 | 0.97 | 0.968 | Macrophages | aPD-L1 vs PBS |
| Gbp7 | 5.48E-17 | 9.77E-13 | 0.36 | 0.448 | 0.309 | Tumor_cells | aPD-L1 vs PBS |
| mt-Nd3 | 3.95E-07 | 7.03E-03 | 0.36 | 0.518 | 0.256 | NKT | aPD-L1 vs PBS |
| Sgk1 | 1.16E-22 | 2.06E-18 | 0.36 | 0.671 | 0.501 | Tumor_cells | aPD-L1 vs PBS |
| Stat1 | 1.57E-15 | 2.79E-11 | 0.36 | 0.571 | 0.353 | Macrophages | aPD-L1 vs PBS |

| | | | | | | | |
|---|---|---|---|---|---|---|---|
| Gm10260 | 3.12E-31 | 5.56E-27 | 0.36 | 0.991 | 0.937 | T cells | aPD-L1 vs PBS |
| mt-Nd1 | 5.84E-07 | 1.04E-02 | 0.36 | 0.971 | 0.962 | NKT | aPD-L1 vs PBS |
| Jund | 4.86E-08 | 8.67E-04 | 0.36 | 0.881 | 0.817 | Macrophages | aPD-L1 vs PBS |
| Hk2 | 3.22E-07 | 5.73E-03 | 0.36 | 0.526 | 0.329 | Monocytes | aPD-L1 vs PBS |
| Rpl13 | 1.66E-18 | 2.97E-14 | 0.36 | 0.958 | 0.939 | Neutrophils | aPD-L1 vs PBS |
| Cd83 | 3.17E-07 | 5.65E-03 | 0.35 | 0.602 | 0.373 | Monocytes | aPD-L1 vs PBS |
| Rpl36a | 4.09E-08 | 7.28E-04 | 0.35 | 0.77 | 0.62 | Monocytes | aPD-L1 vs PBS |
| Cox7c | 5.05E-23 | 9.01E-19 | 0.35 | 0.744 | 0.396 | B cells | aPD-L1 vs PBS |
| Vps37b | 6.34E-14 | 1.13E-09 | 0.35 | 0.817 | 0.643 | T cells | aPD-L1 vs PBS |
| Hal | 2.00E-06 | 3.57E-02 | 0.35 | 0.399 | 0.278 | Macrophages | aPD-L1 vs PBS |
| Rpl30 | 1.00E-17 | 1.78E-13 | 0.35 | 0.926 | 0.918 | Neutrophils | aPD-L1 vs PBS |
| Zfp36 | 2.45E-09 | 4.36E-05 | 0.35 | 0.836 | 0.69 | Macrophages | aPD-L1 vs PBS |
| Zfand5 | 2.70E-06 | 4.81E-02 | 0.35 | 0.719 | 0.573 | Monocytes | aPD-L1 vs PBS |
| Snrpg | 6.35E-28 | 1.13E-23 | 0.35 | 0.561 | 0.194 | B cells | aPD-L1 vs PBS |
| Scara3 | 7.45E-07 | 1.33E-02 | 0.35 | 0.621 | 0.295 | Fibroblasts | aPD-L1 vs PBS |
| Gbp4 | 6.92E-14 | 1.23E-09 | 0.35 | 0.34 | 0.158 | Macrophages | aPD-L1 vs PBS |
| mt-Nd4l | 5.93E-24 | 1.06E-19 | 0.35 | 0.671 | 0.349 | T cells | aPD-L1 vs PBS |
| Tomm7 | 6.24E-25 | 1.11E-20 | 0.35 | 0.668 | 0.384 | Macrophages | aPD-L1 vs PBS |
| Gm47283 | 4.42E-07 | 7.89E-03 | 0.35 | 0.663 | 0.475 | Monocytes | aPD-L1 vs PBS |
| Bola2 | 9.29E-25 | 1.66E-20 | 0.35 | 0.599 | 0.285 | T cells | aPD-L1 vs PBS |
| mt-Atp6 | 2.37E-08 | 4.22E-04 | 0.34 | 1 | 1 | NKT | aPD-L1 vs PBS |
| Rpl34 | 3.77E-19 | 6.72E-15 | 0.34 | 0.941 | 0.922 | Neutrophils | aPD-L1 vs PBS |
| Psap | 2.81E-16 | 5.00E-12 | 0.34 | 0.981 | 0.978 | Macrophages | aPD-L1 vs PBS |
| Rpl34 | 2.40E-10 | 4.27E-06 | 0.34 | 0.974 | 0.975 | Monocytes | aPD-L1 vs PBS |
| Ccl4 | 1.72E-07 | 3.07E-03 | 0.34 | 0.791 | 0.653 | Macrophages | aPD-L1 vs PBS |
| Rpl37 | 8.47E-24 | 1.51E-19 | 0.34 | 0.941 | 0.929 | Neutrophils | aPD-L1 vs PBS |
| H2-Q7 | 3.22E-19 | 5.74E-15 | 0.34 | 0.617 | 0.405 | Neutrophils | aPD-L1 vs PBS |
| Rps23 | 2.44E-20 | 4.35E-16 | 0.34 | 0.939 | 0.923 | Neutrophils | aPD-L1 vs PBS |
| mt-Cytb | 6.15E-07 | 1.10E-02 | 0.34 | 1 | 0.994 | NKT | aPD-L1 vs PBS |
| Fos | 6.16E-18 | 1.10E-13 | 0.34 | 0.839 | 0.694 | Tumor_cells | aPD-L1 vs PBS |
| Vps37b | 4.32E-10 | 7.70E-06 | 0.34 | 0.595 | 0.396 | B cells | aPD-L1 vs PBS |
| Polr2l | 1.70E-09 | 3.02E-05 | 0.34 | 0.403 | 0.174 | Monocytes | aPD-L1 vs PBS |
| Sec61g | 9.22E-22 | 1.64E-17 | 0.34 | 0.791 | 0.616 | Macrophages | aPD-L1 vs PBS |
| Nampt | 6.65E-08 | 1.18E-03 | 0.33 | 0.633 | 0.462 | Monocytes | aPD-L1 vs PBS |
| Rps27rt | 1.31E-24 | 2.34E-20 | 0.33 | 0.501 | 0.177 | B cells | aPD-L1 vs PBS |
| Fosb | 2.68E-07 | 4.78E-03 | 0.33 | 0.627 | 0.492 | Macrophages | aPD-L1 vs PBS |
| Rps15a | 1.14E-49 | 2.03E-45 | 0.33 | 1 | 1 | T cells | aPD-L1 vs PBS |
| Rpl41 | 1.53E-22 | 2.72E-18 | 0.33 | 0.965 | 0.959 | Neutrophils | aPD-L1 vs PBS |
| Gm31243 | 1.96E-18 | 3.50E-14 | 0.33 | 0.606 | 0.353 | B cells | aPD-L1 vs PBS |
| Sod2 | 6.64E-13 | 1.18E-08 | 0.33 | 0.345 | 0.185 | Neutrophils | aPD-L1 vs PBS |
| Socs3 | 9.67E-07 | 1.72E-02 | 0.33 | 0.755 | 0.589 | Monocytes | aPD-L1 vs PBS |
| Icam1 | 4.68E-11 | 8.34E-07 | 0.33 | 0.215 | 0.099 | Neutrophils | aPD-L1 vs PBS |
| Pde4b | 1.08E-08 | 1.93E-04 | 0.33 | 0.52 | 0.291 | Monocytes | aPD-L1 vs PBS |
| Rpl35 | 3.72E-10 | 6.63E-06 | 0.33 | 0.923 | 0.816 | Monocytes | aPD-L1 vs PBS |
| Fam26f | 2.45E-20 | 4.37E-16 | 0.33 | 0.325 | 0.11 | Macrophages | aPD-L1 vs PBS |
| Igtp | 2.91E-21 | 5.18E-17 | 0.33 | 0.432 | 0.209 | Neutrophils | aPD-L1 vs PBS |
| Rps27rt | 3.18E-23 | 5.67E-19 | 0.33 | 0.547 | 0.257 | T cells | aPD-L1 vs PBS |
| Rpl22l1 | 9.50E-22 | 1.69E-17 | 0.33 | 0.984 | 0.915 | T cells | aPD-L1 vs PBS |
| Rpsa | 2.80E-14 | 5.00E-10 | 0.33 | 0.909 | 0.91 | Neutrophils | aPD-L1 vs PBS |
| Gbp2b | 7.98E-21 | 1.42E-16 | 0.33 | 0.557 | 0.312 | Neutrophils | aPD-L1 vs PBS |
| Socs1 | 4.26E-19 | 7.59E-15 | 0.33 | 0.37 | 0.168 | Neutrophils | aPD-L1 vs PBS |
| Rpl22l1 | 1.29E-14 | 2.30E-10 | 0.32 | 0.795 | 0.602 | Macrophages | aPD-L1 vs PBS |
| Rpl41 | 8.69E-28 | 1.55E-23 | 0.32 | 1 | 0.999 | Macrophages | aPD-L1 vs PBS |
| Atp5e | 2.85E-22 | 5.08E-18 | 0.32 | 0.882 | 0.617 | T cells | aPD-L1 vs PBS |
| Fgl2 | 8.18E-10 | 1.46E-05 | 0.32 | 0.44 | 0.277 | Macrophages | aPD-L1 vs PBS |

| | | | | | | | |
|---|---|---|---|---|---|---|---|
| Sem1 | 4.64E-25 | 8.28E-21 | 0.32 | 0.937 | 0.847 | Macrophages | aPD-L1 vs PBS |
| Irf1 | 6.97E-18 | 1.24E-13 | 0.32 | 0.492 | 0.197 | B cells | aPD-L1 vs PBS |
| Rpl27a | 1.86E-17 | 3.31E-13 | 0.32 | 0.949 | 0.938 | Neutrophils | aPD-L1 vs PBS |
| Acod1 | 1.87E-13 | 3.34E-09 | 0.32 | 0.86 | 0.75 | Neutrophils | aPD-L1 vs PBS |
| Ccnl1 | 2.82E-32 | 5.03E-28 | 0.32 | 0.51 | 0.285 | Tumor_cells | aPD-L1 vs PBS |
| Trib1 | 2.03E-34 | 3.62E-30 | 0.32 | 0.472 | 0.237 | Tumor_cells | aPD-L1 vs PBS |
| Atp5k | 6.02E-18 | 1.07E-13 | 0.32 | 0.731 | 0.547 | Macrophages | aPD-L1 vs PBS |
| Cox7c | 7.70E-22 | 1.37E-17 | 0.32 | 0.888 | 0.773 | Macrophages | aPD-L1 vs PBS |
| Rpl36 | 4.62E-22 | 8.23E-18 | 0.32 | 0.996 | 0.992 | Macrophages | aPD-L1 vs PBS |
| Ifi27l2a | 1.10E-10 | 1.95E-06 | 0.32 | 0.703 | 0.468 | T cells | aPD-L1 vs PBS |
| Ndufa1 | 1.02E-23 | 1.82E-19 | 0.32 | 0.563 | 0.281 | Macrophages | aPD-L1 vs PBS |
| Clec4e | 5.02E-10 | 8.95E-06 | 0.32 | 0.794 | 0.669 | Neutrophils | aPD-L1 vs PBS |
| Nlrp3 | 4.79E-07 | 8.54E-03 | 0.32 | 0.536 | 0.329 | Monocytes | aPD-L1 vs PBS |
| Lst1 | 4.42E-11 | 7.88E-07 | 0.32 | 0.556 | 0.412 | Neutrophils | aPD-L1 vs PBS |
| Rps23 | 3.12E-46 | 5.55E-42 | 0.32 | 1 | 1 | B cells | aPD-L1 vs PBS |
| Blvrb | 2.41E-08 | 4.30E-04 | 0.31 | 0.601 | 0.464 | Macrophages | aPD-L1 vs PBS |
| mt-Nd3 | 8.45E-13 | 1.51E-08 | 0.31 | 0.608 | 0.436 | Macrophages | aPD-L1 vs PBS |
| Gadd45b | 3.25E-11 | 5.78E-07 | 0.31 | 0.543 | 0.329 | T cells | aPD-L1 vs PBS |
| Rpl36al | 2.56E-22 | 4.56E-18 | 0.31 | 0.473 | 0.16 | B cells | aPD-L1 vs PBS |
| H2-T23 | 6.91E-19 | 1.23E-14 | 0.31 | 0.622 | 0.402 | Neutrophils | aPD-L1 vs PBS |
| Rel | 5.00E-09 | 8.91E-05 | 0.31 | 0.541 | 0.291 | Monocytes | aPD-L1 vs PBS |
| Sem1 | 4.29E-10 | 7.64E-06 | 0.31 | 0.944 | 0.902 | Monocytes | aPD-L1 vs PBS |
| Cox7c | 7.40E-22 | 1.32E-17 | 0.31 | 0.759 | 0.42 | T cells | aPD-L1 vs PBS |
| Ly6e | 2.98E-10 | 5.31E-06 | 0.31 | 0.854 | 0.759 | Macrophages | aPD-L1 vs PBS |
| Ahnak | 3.80E-21 | 6.77E-17 | 0.30 | 0.732 | 0.584 | Tumor_cells | aPD-L1 vs PBS |
| Nampt | 7.39E-15 | 1.32E-10 | 0.30 | 0.511 | 0.294 | Macrophages | aPD-L1 vs PBS |
| Gm10260 | 4.58E-21 | 8.17E-17 | 0.30 | 0.962 | 0.843 | B cells | aPD-L1 vs PBS |
| Gbp4 | 3.00E-29 | 5.35E-25 | 0.30 | 0.229 | 0.071 | Tumor_cells | aPD-L1 vs PBS |
| Cox17 | 1.95E-18 | 3.47E-14 | 0.30 | 0.424 | 0.219 | Neutrophils | aPD-L1 vs PBS |
| Spp1 | 6.05E-09 | 1.08E-04 | 0.30 | 0.965 | 0.968 | Tumor_cells | aPD-L1 vs PBS |
| Sec61g | 1.42E-08 | 2.53E-04 | 0.30 | 0.806 | 0.633 | Monocytes | aPD-L1 vs PBS |
| Igfbp4 | 9.59E-13 | 1.71E-08 | 0.30 | 0.597 | 0.486 | Tumor_cells | aPD-L1 vs PBS |
| Itgb7 | 3.30E-17 | 5.88E-13 | 0.30 | 0.507 | 0.258 | Macrophages | aPD-L1 vs PBS |
| Kdm6b | 2.65E-06 | 4.72E-02 | 0.30 | 0.597 | 0.44 | Monocytes | aPD-L1 vs PBS |
| Nampt | 6.18E-14 | 1.10E-09 | 0.30 | 0.476 | 0.299 | Neutrophils | aPD-L1 vs PBS |
| Nfkb1 | 1.29E-12 | 2.31E-08 | 0.30 | 0.511 | 0.298 | T cells | aPD-L1 vs PBS |
| mt-Nd4l | 2.27E-15 | 4.05E-11 | 0.29 | 0.658 | 0.37 | B cells | aPD-L1 vs PBS |
| Rpl38 | 5.92E-17 | 1.06E-12 | 0.29 | 0.89 | 0.878 | Neutrophils | aPD-L1 vs PBS |
| Rplp0 | 8.75E-12 | 1.56E-07 | 0.29 | 0.926 | 0.926 | Neutrophils | aPD-L1 vs PBS |
| Mmp2 | 7.37E-09 | 1.31E-04 | 0.29 | 0.319 | 0.223 | Tumor_cells | aPD-L1 vs PBS |
| Rpl30 | 2.56E-39 | 4.56E-35 | 0.29 | 1 | 1 | T cells | aPD-L1 vs PBS |
| Rps26 | 1.74E-23 | 3.09E-19 | 0.29 | 0.993 | 0.989 | T cells | aPD-L1 vs PBS |
| Tnfaip3 | 5.25E-10 | 9.36E-06 | 0.29 | 0.568 | 0.396 | Neutrophils | aPD-L1 vs PBS |
| Txn1 | 2.55E-08 | 4.54E-04 | 0.29 | 0.955 | 0.944 | Macrophages | aPD-L1 vs PBS |
| Rpl31 | 1.47E-18 | 2.61E-14 | 0.29 | 0.951 | 0.778 | B cells | aPD-L1 vs PBS |
| Gramd3 | 3.74E-12 | 6.67E-08 | 0.29 | 0.541 | 0.346 | T cells | aPD-L1 vs PBS |
| Tnfaip3 | 3.94E-10 | 7.03E-06 | 0.29 | 0.714 | 0.519 | T cells | aPD-L1 vs PBS |
| Adgre5 | 4.00E-12 | 7.14E-08 | 0.29 | 0.328 | 0.157 | Macrophages | aPD-L1 vs PBS |
| Rpl36 | 6.04E-12 | 1.08E-07 | 0.29 | 0.894 | 0.908 | Neutrophils | aPD-L1 vs PBS |
| Rplp2 | 7.32E-14 | 1.30E-09 | 0.29 | 0.912 | 0.909 | Neutrophils | aPD-L1 vs PBS |
| Rpl35 | 6.62E-10 | 1.18E-05 | 0.29 | 0.785 | 0.824 | Neutrophils | aPD-L1 vs PBS |
| Nos2 | 1.47E-08 | 2.62E-04 | 0.29 | 0.179 | 0.032 | Monocytes | aPD-L1 vs PBS |
| Tmsb10 | 3.55E-09 | 6.33E-05 | 0.29 | 0.856 | 0.879 | Neutrophils | aPD-L1 vs PBS |
| Zfp36l2 | 2.05E-08 | 3.65E-04 | 0.29 | 0.642 | 0.499 | Macrophages | aPD-L1 vs PBS |
| Uba52 | 8.56E-11 | 1.53E-06 | 0.28 | 0.454 | 0.199 | Monocytes | aPD-L1 vs PBS |

| | | | | | | | |
|---|---|---|---|---|---|---|---|
| Sec61g | 1.15E-18 | 2.05E-14 | 0.28 | 0.617 | 0.336 | T cells | aPD-L1 vs PBS |
| Fst | 1.78E-11 | 3.18E-07 | 0.28 | 0.388 | 0.27 | Tumor_cells | aPD-L1 vs PBS |
| Gm26825 | 3.34E-12 | 5.95E-08 | 0.28 | 0.525 | 0.318 | T cells | aPD-L1 vs PBS |
| Dnaja1 | 8.90E-08 | 1.59E-03 | 0.28 | 0.928 | 0.847 | T cells | aPD-L1 vs PBS |
| Gm11808 | 3.13E-23 | 5.58E-19 | 0.28 | 0.496 | 0.194 | T cells | aPD-L1 vs PBS |
| Tra2b | 2.09E-13 | 3.72E-09 | 0.28 | 0.847 | 0.641 | T cells | aPD-L1 vs PBS |
| Mt2 | 3.73E-23 | 6.66E-19 | 0.28 | 0.743 | 0.561 | Tumor_cells | aPD-L1 vs PBS |
| Rplp2 | 2.29E-11 | 4.09E-07 | 0.28 | 0.99 | 0.981 | Monocytes | aPD-L1 vs PBS |
| Slc11a1 | 1.63E-09 | 2.91E-05 | 0.28 | 0.526 | 0.378 | Macrophages | aPD-L1 vs PBS |
| Ighg1 | 7.28E-07 | 1.30E-02 | 0.28 | 0.144 | 0 | NKT | aPD-L1 vs PBS |
| Klf4 | 9.97E-07 | 1.78E-02 | 0.28 | 0.466 | 0.332 | Macrophages | aPD-L1 vs PBS |
| Il1f9 | 1.25E-06 | 2.23E-02 | 0.28 | 0.515 | 0.407 | Neutrophils | aPD-L1 vs PBS |
| Rpl31 | 3.62E-19 | 6.46E-15 | 0.28 | 0.981 | 0.839 | T cells | aPD-L1 vs PBS |
| Crip1 | 6.11E-08 | 1.09E-03 | 0.28 | 0.485 | 0.288 | B cells | aPD-L1 vs PBS |
| Itgb2 | 3.53E-08 | 6.30E-04 | 0.28 | 0.683 | 0.586 | Macrophages | aPD-L1 vs PBS |
| H2-D1 | 6.96E-18 | 1.24E-13 | 0.28 | 1 | 1 | Macrophages | aPD-L1 vs PBS |
| Ubb | 4.85E-31 | 8.64E-27 | 0.28 | 0.997 | 0.99 | Tumor_cells | aPD-L1 vs PBS |
| Phlda1 | 3.72E-08 | 6.64E-04 | 0.28 | 0.809 | 0.772 | Tumor_cells | aPD-L1 vs PBS |
| Arid5a | 5.94E-08 | 1.06E-03 | 0.28 | 0.485 | 0.256 | Monocytes | aPD-L1 vs PBS |
| Hnrnpa0 | 1.23E-35 | 2.19E-31 | 0.28 | 0.839 | 0.692 | Tumor_cells | aPD-L1 vs PBS |
| Zfand5 | 1.68E-08 | 2.99E-04 | 0.28 | 0.612 | 0.494 | Macrophages | aPD-L1 vs PBS |
| Rps8 | 8.21E-12 | 1.46E-07 | 0.28 | 0.932 | 0.928 | Neutrophils | aPD-L1 vs PBS |
| Gbp2 | 1.03E-15 | 1.84E-11 | 0.28 | 0.332 | 0.135 | Macrophages | aPD-L1 vs PBS |
| Egr1 | 1.13E-07 | 2.02E-03 | 0.27 | 0.343 | 0.198 | Macrophages | aPD-L1 vs PBS |
| Ifrd1 | 4.63E-12 | 8.26E-08 | 0.27 | 0.612 | 0.399 | T cells | aPD-L1 vs PBS |
| Junb | 9.53E-08 | 1.70E-03 | 0.27 | 0.896 | 0.798 | Macrophages | aPD-L1 vs PBS |
| Csf1 | 5.08E-11 | 9.05E-07 | 0.27 | 0.444 | 0.279 | Neutrophils | aPD-L1 vs PBS |
| Atf3 | 1.97E-13 | 3.50E-09 | 0.27 | 0.388 | 0.248 | Tumor_cells | aPD-L1 vs PBS |
| Rpl19 | 1.30E-13 | 2.32E-09 | 0.27 | 0.927 | 0.924 | Neutrophils | aPD-L1 vs PBS |
| Cox6c | 2.88E-16 | 5.14E-12 | 0.27 | 0.724 | 0.458 | T cells | aPD-L1 vs PBS |
| Ets2 | 2.10E-06 | 3.75E-02 | 0.27 | 0.556 | 0.364 | Monocytes | aPD-L1 vs PBS |
| Snrpf | 6.94E-19 | 1.24E-14 | 0.27 | 0.48 | 0.2 | T cells | aPD-L1 vs PBS |
| Il1a | 1.52E-07 | 2.71E-03 | 0.27 | 0.203 | 0.107 | Neutrophils | aPD-L1 vs PBS |
| Lars2 | 4.12E-31 | 7.35E-27 | 0.27 | 0.364 | 0.156 | Tumor_cells | aPD-L1 vs PBS |
| Klf2 | 4.55E-09 | 8.11E-05 | 0.27 | 0.795 | 0.626 | Macrophages | aPD-L1 vs PBS |
| Nfkbia | 3.78E-09 | 6.74E-05 | 0.27 | 0.869 | 0.711 | Macrophages | aPD-L1 vs PBS |
| B2m | 5.77E-23 | 1.03E-18 | 0.27 | 1 | 1 | Macrophages | aPD-L1 vs PBS |
| Rpl27 | 1.31E-15 | 2.33E-11 | 0.27 | 0.69 | 0.473 | Macrophages | aPD-L1 vs PBS |
| Btg1 | 6.86E-08 | 1.22E-03 | 0.27 | 0.98 | 0.953 | Monocytes | aPD-L1 vs PBS |
| Smad7 | 7.91E-11 | 1.41E-06 | 0.27 | 0.882 | 0.767 | T cells | aPD-L1 vs PBS |
| Cox6c | 5.89E-15 | 1.05E-10 | 0.26 | 0.646 | 0.359 | B cells | aPD-L1 vs PBS |
| mt-Nd3 | 1.18E-06 | 2.11E-02 | 0.26 | 0.48 | 0.291 | Monocytes | aPD-L1 vs PBS |
| Il1rn | 9.76E-08 | 1.74E-03 | 0.26 | 0.745 | 0.652 | Neutrophils | aPD-L1 vs PBS |
| Rpl36al | 7.57E-17 | 1.35E-12 | 0.26 | 0.567 | 0.317 | Macrophages | aPD-L1 vs PBS |
| Hmgb2 | 2.73E-10 | 4.87E-06 | 0.26 | 0.782 | 0.564 | B cells | aPD-L1 vs PBS |
| Tnf | 5.03E-07 | 8.97E-03 | 0.26 | 0.422 | 0.28 | Macrophages | aPD-L1 vs PBS |
| Gbp3 | 2.35E-12 | 4.20E-08 | 0.26 | 0.388 | 0.26 | Tumor_cells | aPD-L1 vs PBS |
| Hcst | 1.03E-13 | 1.84E-09 | 0.26 | 0.671 | 0.464 | T cells | aPD-L1 vs PBS |
| Rpl12 | 2.68E-16 | 4.78E-12 | 0.26 | 0.895 | 0.902 | Neutrophils | aPD-L1 vs PBS |
| Rps3a1 | 1.37E-10 | 2.45E-06 | 0.26 | 0.909 | 0.926 | Neutrophils | aPD-L1 vs PBS |
| Prkca | 1.04E-09 | 1.85E-05 | 0.26 | 0.448 | 0.272 | T cells | aPD-L1 vs PBS |
| Jun | 1.44E-09 | 2.57E-05 | 0.26 | 0.89 | 0.834 | Tumor_cells | aPD-L1 vs PBS |
| Nr4a1 | 1.03E-10 | 1.84E-06 | 0.26 | 0.295 | 0.14 | Macrophages | aPD-L1 vs PBS |
| Igkc | 1.63E-07 | 2.90E-03 | 0.26 | 0.311 | 0.037 | ILC | aPD-L1 vs PBS |
| Son | 2.54E-13 | 4.53E-09 | 0.26 | 0.791 | 0.551 | T cells | aPD-L1 vs PBS |

| | | | | | | | |
|---|---|---|---|---|---|---|---|
| Adamts1 | 4.42E-24 | 7.87E-20 | 0.26 | 0.394 | 0.206 | Tumor_cells | aPD-L1 vs PBS |
| Klf6 | 2.30E-07 | 4.11E-03 | 0.26 | 0.925 | 0.814 | Macrophages | aPD-L1 vs PBS |
| Ptgs2 | 2.44E-07 | 4.35E-03 | 0.26 | 0.25 | 0.126 | Macrophages | aPD-L1 vs PBS |
| Rpl21 | 3.03E-24 | 5.40E-20 | 0.26 | 0.999 | 1 | B cells | aPD-L1 vs PBS |
| Tomm7 | 1.05E-16 | 1.88E-12 | 0.26 | 0.483 | 0.202 | B cells | aPD-L1 vs PBS |
| Crem | 4.54E-11 | 8.10E-07 | 0.26 | 0.3 | 0.125 | B cells | aPD-L1 vs PBS |
| Dusp2 | 5.96E-10 | 1.06E-05 | 0.26 | 0.261 | 0.119 | Macrophages | aPD-L1 vs PBS |
| mt-Co2 | 2.18E-06 | 3.89E-02 | 0.26 | 1 | 1 | NKT | aPD-L1 vs PBS |
| Cd52 | 6.39E-11 | 1.14E-06 | 0.25 | 0.823 | 0.757 | Neutrophils | aPD-L1 vs PBS |
| Zfp36 | 4.14E-08 | 7.38E-04 | 0.25 | 0.978 | 0.989 | Neutrophils | aPD-L1 vs PBS |
| Brd2 | 5.64E-23 | 1.00E-18 | 0.25 | 0.469 | 0.282 | Tumor_cells | aPD-L1 vs PBS |
| H2-Q7 | 2.43E-10 | 4.33E-06 | 0.25 | 0.534 | 0.356 | Macrophages | aPD-L1 vs PBS |
| Dusp6 | 2.39E-23 | 4.26E-19 | 0.25 | 0.37 | 0.188 | Tumor_cells | aPD-L1 vs PBS |
| Polr2l | 4.51E-09 | 8.04E-05 | 0.25 | 0.362 | 0.203 | Macrophages | aPD-L1 vs PBS |
| F10 | 4.81E-07 | 8.57E-03 | 0.25 | 0.408 | 0.203 | Monocytes | aPD-L1 vs PBS |
| Rpl28 | 1.68E-09 | 3.00E-05 | 0.25 | 0.91 | 0.925 | Neutrophils | aPD-L1 vs PBS |
| Klf4 | 9.33E-21 | 1.66E-16 | 0.25 | 0.475 | 0.29 | Tumor_cells | aPD-L1 vs PBS |
| Rps8 | 5.15E-35 | 9.18E-31 | 0.25 | 1 | 1 | B cells | aPD-L1 vs PBS |
| Tmx4 | 8.28E-07 | 1.48E-02 | -0.25 | 0.224 | 0.374 | Macrophages | aPD-L1 vs PBS |
| Ntpcr | 1.89E-09 | 3.37E-05 | -0.25 | 0.369 | 0.557 | Macrophages | aPD-L1 vs PBS |
| Nfil3 | 1.24E-07 | 2.21E-03 | -0.25 | 0.38 | 0.487 | Neutrophils | aPD-L1 vs PBS |
| Bsg | 1.92E-09 | 3.42E-05 | -0.25 | 0.6 | 0.73 | Neutrophils | aPD-L1 vs PBS |
| Park7 | 3.38E-07 | 6.03E-03 | -0.25 | 0.255 | 0.468 | Monocytes | aPD-L1 vs PBS |
| Hexb | 4.96E-08 | 8.84E-04 | -0.25 | 0.481 | 0.635 | Macrophages | aPD-L1 vs PBS |
| Ier2 | 1.07E-08 | 1.90E-04 | -0.25 | 0.777 | 0.855 | B cells | aPD-L1 vs PBS |
| Ltb | 1.77E-06 | 3.15E-02 | -0.25 | 0.761 | 0.815 | T cells | aPD-L1 vs PBS |
| Ostc | 2.92E-22 | 5.20E-18 | -0.25 | 0.638 | 0.774 | Tumor_cells | aPD-L1 vs PBS |
| Ppia | 1.76E-12 | 3.13E-08 | -0.26 | 0.993 | 0.998 | Macrophages | aPD-L1 vs PBS |
| Tmed9 | 4.57E-07 | 8.15E-03 | -0.26 | 0.321 | 0.509 | Monocytes | aPD-L1 vs PBS |
| Akr1a1 | 1.07E-08 | 1.91E-04 | -0.26 | 0.622 | 0.791 | Monocytes | aPD-L1 vs PBS |
| Fth1 | 1.41E-12 | 2.52E-08 | -0.26 | 1 | 1 | Neutrophils | aPD-L1 vs PBS |
| Cap1 | 2.36E-06 | 4.21E-02 | -0.26 | 0.556 | 0.706 | Monocytes | aPD-L1 vs PBS |
| Shisa5 | 9.24E-11 | 1.65E-06 | -0.26 | 0.81 | 0.821 | T cells | aPD-L1 vs PBS |
| Atp5f1 | 8.27E-30 | 1.47E-25 | -0.26 | 0.891 | 0.961 | Tumor_cells | aPD-L1 vs PBS |
| Rps6 | 1.24E-13 | 2.22E-09 | -0.26 | 0.933 | 0.98 | Macrophages | aPD-L1 vs PBS |
| Atp5f1 | 1.32E-07 | 2.35E-03 | -0.26 | 0.5 | 0.709 | Monocytes | aPD-L1 vs PBS |
| Pkm | 2.97E-21 | 5.30E-17 | -0.26 | 0.383 | 0.675 | B cells | aPD-L1 vs PBS |
| Rps3 | 6.07E-15 | 1.08E-10 | -0.26 | 0.978 | 0.991 | Macrophages | aPD-L1 vs PBS |
| Klf2 | 3.54E-08 | 6.31E-04 | -0.26 | 0.923 | 0.966 | B cells | aPD-L1 vs PBS |
| Dusp1 | 8.70E-12 | 1.55E-07 | -0.26 | 0.672 | 0.806 | B cells | aPD-L1 vs PBS |
| Ftl1 | 3.10E-09 | 5.52E-05 | -0.26 | 0.977 | 0.994 | B cells | aPD-L1 vs PBS |
| Mcl1 | 3.58E-18 | 6.37E-14 | -0.26 | 1 | 1 | Neutrophils | aPD-L1 vs PBS |
| Clic1 | 5.34E-12 | 9.52E-08 | -0.27 | 0.554 | 0.667 | T cells | aPD-L1 vs PBS |
| Arpc1b | 3.00E-19 | 5.35E-15 | -0.27 | 0.963 | 0.991 | Macrophages | aPD-L1 vs PBS |
| Lgmn | 8.42E-09 | 1.50E-04 | -0.27 | 0.854 | 0.921 | Macrophages | aPD-L1 vs PBS |
| Park7 | 7.62E-13 | 1.36E-08 | -0.27 | 0.466 | 0.673 | Macrophages | aPD-L1 vs PBS |
| Oaz1 | 7.14E-09 | 1.27E-04 | -0.27 | 0.867 | 0.953 | Monocytes | aPD-L1 vs PBS |
| Rpl5 | 3.19E-15 | 5.68E-11 | -0.27 | 0.998 | 0.998 | T cells | aPD-L1 vs PBS |
| Hist1h1c | 1.86E-06 | 3.32E-02 | -0.27 | 0.496 | 0.601 | B cells | aPD-L1 vs PBS |
| Eid1 | 1.68E-14 | 3.00E-10 | -0.27 | 0.373 | 0.617 | Macrophages | aPD-L1 vs PBS |
| Clic1 | 4.76E-08 | 8.49E-04 | -0.27 | 0.862 | 0.946 | Monocytes | aPD-L1 vs PBS |
| Atp5g2 | 2.43E-15 | 4.34E-11 | -0.27 | 0.687 | 0.854 | Macrophages | aPD-L1 vs PBS |
| Lgals1 | 5.18E-41 | 9.23E-37 | -0.27 | 0.996 | 0.999 | Tumor_cells | aPD-L1 vs PBS |
| Alox5ap | 2.16E-06 | 3.85E-02 | -0.27 | 0.443 | 0.528 | Neutrophils | aPD-L1 vs PBS |
| Junb | 3.02E-09 | 5.39E-05 | -0.27 | 0.98 | 0.986 | B cells | aPD-L1 vs PBS |

| | | | | | | | |
|---|---|---|---|---|---|---|---|
| Tmsb4x | 3.83E-08 | 6.83E-04 | -0.28 | 1 | 1 | Monocytes | aPD-L1 vs PBS |
| Slc2a1 | 9.20E-09 | 1.64E-04 | -0.28 | 0.492 | 0.592 | Neutrophils | aPD-L1 vs PBS |
| Btg2 | 7.75E-12 | 1.38E-07 | -0.28 | 0.905 | 0.926 | B cells | aPD-L1 vs PBS |
| Rpl14 | 1.25E-18 | 2.23E-14 | -0.28 | 0.951 | 0.984 | Macrophages | aPD-L1 vs PBS |
| Rps3a1 | 1.20E-13 | 2.15E-09 | -0.28 | 0.978 | 0.985 | Macrophages | aPD-L1 vs PBS |
| Ifitm3 | 4.20E-07 | 7.48E-03 | -0.28 | 0.99 | 0.997 | Monocytes | aPD-L1 vs PBS |
| Srrm2 | 1.11E-11 | 1.99E-07 | -0.28 | 0.687 | 0.76 | T cells | aPD-L1 vs PBS |
| Ppp1r14b | 7.18E-12 | 1.28E-07 | -0.28 | 0.422 | 0.613 | Macrophages | aPD-L1 vs PBS |
| Rps2 | 5.32E-15 | 9.48E-11 | -0.29 | 0.955 | 0.984 | Macrophages | aPD-L1 vs PBS |
| Fth1 | 3.91E-10 | 6.97E-06 | -0.29 | 1 | 1 | Macrophages | aPD-L1 vs PBS |
| S100a6 | 9.61E-41 | 1.71E-36 | -0.29 | 0.177 | 0.59 | B cells | aPD-L1 vs PBS |
| Fcgr3 | 1.40E-09 | 2.49E-05 | -0.29 | 0.662 | 0.743 | Neutrophils | aPD-L1 vs PBS |
| Erp29 | 2.11E-16 | 3.76E-12 | -0.29 | 0.765 | 0.888 | Macrophages | aPD-L1 vs PBS |
| Apoe | 1.64E-06 | 2.93E-02 | -0.29 | 0.453 | 0.561 | B cells | aPD-L1 vs PBS |
| Ndel1 | 2.83E-10 | 5.04E-06 | -0.29 | 0.498 | 0.611 | Neutrophils | aPD-L1 vs PBS |
| Ppp1r3b | 3.61E-12 | 6.44E-08 | -0.30 | 0.328 | 0.472 | Neutrophils | aPD-L1 vs PBS |
| Pgam1 | 4.28E-10 | 7.63E-06 | -0.30 | 0.848 | 0.923 | Neutrophils | aPD-L1 vs PBS |
| Pfn1 | 1.81E-17 | 3.23E-13 | -0.30 | 0.972 | 0.972 | T cells | aPD-L1 vs PBS |
| Rpl22 | 1.94E-06 | 3.46E-02 | -0.30 | 0.969 | 0.985 | Epithelial cells | aPD-L1 vs PBS |
| Clic1 | 1.34E-06 | 2.39E-02 | -0.30 | 0.881 | 0.944 | ILC | aPD-L1 vs PBS |
| Ppia | 5.26E-07 | 9.37E-03 | -0.31 | 0.986 | 0.994 | NKT | aPD-L1 vs PBS |
| Rpl5 | 3.87E-18 | 6.89E-14 | -0.31 | 0.918 | 0.964 | Macrophages | aPD-L1 vs PBS |
| Atp5g2 | 2.34E-06 | 4.17E-02 | -0.31 | 0.8 | 0.86 | ILC | aPD-L1 vs PBS |
| Aprt | 2.27E-07 | 4.05E-03 | -0.31 | 0.648 | 0.791 | Monocytes | aPD-L1 vs PBS |
| Ms4a6b | 2.24E-07 | 3.99E-03 | -0.31 | 0.582 | 0.747 | Monocytes | aPD-L1 vs PBS |
| Gng11 | 1.83E-06 | 3.27E-02 | -0.31 | 0.282 | 0.362 | Tumor_cells | aPD-L1 vs PBS |
| Arpc1b | 9.63E-12 | 1.72E-07 | -0.31 | 0.944 | 0.991 | Monocytes | aPD-L1 vs PBS |
| Ppia | 3.10E-08 | 5.53E-04 | -0.32 | 0.993 | 1 | ILC | aPD-L1 vs PBS |
| Psme2 | 5.79E-08 | 1.03E-03 | -0.32 | 0.571 | 0.737 | Monocytes | aPD-L1 vs PBS |
| Ifitm3 | 5.05E-43 | 9.00E-39 | -0.32 | 0.148 | 0.533 | B cells | aPD-L1 vs PBS |
| Sh3bgrl3 | 4.31E-10 | 7.69E-06 | -0.32 | 0.852 | 0.908 | Monocytes | aPD-L1 vs PBS |
| Rgs10 | 4.33E-16 | 7.73E-12 | -0.32 | 0.455 | 0.707 | Macrophages | aPD-L1 vs PBS |
| Ifitm2 | 1.46E-11 | 2.60E-07 | -0.32 | 0.905 | 0.965 | Neutrophils | aPD-L1 vs PBS |
| Gnai2 | 2.60E-06 | 4.63E-02 | -0.32 | 0.856 | 0.875 | NKT | aPD-L1 vs PBS |
| Rpl13 | 2.27E-26 | 4.05E-22 | -0.32 | 0.996 | 1 | Macrophages | aPD-L1 vs PBS |
| Actg1 | 6.01E-33 | 1.07E-28 | -0.32 | 0.972 | 0.992 | Tumor_cells | aPD-L1 vs PBS |
| Rpl3 | 5.35E-18 | 9.53E-14 | -0.33 | 0.951 | 0.989 | Macrophages | aPD-L1 vs PBS |
| Cfp | 2.76E-06 | 4.92E-02 | -0.33 | 0.485 | 0.636 | Monocytes | aPD-L1 vs PBS |
| Pid1 | 1.78E-09 | 3.18E-05 | -0.33 | 0.687 | 0.772 | Macrophages | aPD-L1 vs PBS |
| Srgn | 4.85E-27 | 8.64E-23 | -0.33 | 1 | 1 | Neutrophils | aPD-L1 vs PBS |
| Rpl18 | 2.84E-07 | 5.06E-03 | -0.33 | 1 | 1 | ILC | aPD-L1 vs PBS |
| Rpl18 | 4.26E-29 | 7.59E-25 | -0.33 | 0.985 | 0.994 | Macrophages | aPD-L1 vs PBS |
| Rpl8 | 4.02E-22 | 7.17E-18 | -0.33 | 0.996 | 0.996 | Macrophages | aPD-L1 vs PBS |
| Rhob | 7.25E-08 | 1.29E-03 | -0.34 | 0.733 | 0.777 | Neutrophils | aPD-L1 vs PBS |
| S100a10 | 1.79E-12 | 3.18E-08 | -0.34 | 0.548 | 0.686 | T cells | aPD-L1 vs PBS |
| Actb | 2.60E-23 | 4.63E-19 | -0.34 | 1 | 1 | T cells | aPD-L1 vs PBS |
| Psmb8 | 7.67E-08 | 1.37E-03 | -0.34 | 0.935 | 0.956 | NKT | aPD-L1 vs PBS |
| Rpl12 | 4.90E-18 | 8.73E-14 | -0.34 | 0.933 | 0.986 | Macrophages | aPD-L1 vs PBS |
| Ccl6 | 1.16E-06 | 2.06E-02 | -0.34 | 0.493 | 0.586 | Neutrophils | aPD-L1 vs PBS |
| Ier3 | 1.44E-13 | 2.56E-09 | -0.34 | 0.985 | 0.999 | Neutrophils | aPD-L1 vs PBS |
| Rps4x | 1.74E-19 | 3.11E-15 | -0.34 | 0.981 | 0.993 | Macrophages | aPD-L1 vs PBS |
| Rpl15 | 3.44E-24 | 6.13E-20 | -0.34 | 0.966 | 0.993 | Macrophages | aPD-L1 vs PBS |
| Gnai2 | 1.11E-14 | 1.98E-10 | -0.34 | 0.929 | 0.978 | Monocytes | aPD-L1 vs PBS |
| Gapdh | 1.26E-15 | 2.25E-11 | -0.34 | 0.936 | 0.981 | Neutrophils | aPD-L1 vs PBS |
| Ltb | 1.99E-11 | 3.54E-07 | -0.35 | 0.62 | 0.704 | B cells | aPD-L1 vs PBS |

| | | | | | | | |
|---|---|---|---|---|---|---|---|
| Tgfbi | 1.28E-11 | 2.27E-07 | -0.35 | 0.761 | 0.874 | Macrophages | aPD-L1 vs PBS |
| Rps5 | 1.53E-07 | 2.73E-03 | -0.35 | 1 | 1 | ILC | aPD-L1 vs PBS |
| Itm2b | 1.09E-08 | 1.95E-04 | -0.35 | 0.893 | 0.953 | Monocytes | aPD-L1 vs PBS |
| Cfl1 | 1.27E-13 | 2.27E-09 | -0.35 | 0.944 | 0.994 | Monocytes | aPD-L1 vs PBS |
| Ms4a6d | 3.50E-08 | 6.24E-04 | -0.35 | 0.883 | 0.924 | Monocytes | aPD-L1 vs PBS |
| Cxcl14 | 3.62E-12 | 6.45E-08 | -0.35 | 0.418 | 0.633 | Macrophages | aPD-L1 vs PBS |
| Tspo | 1.83E-09 | 3.26E-05 | -0.35 | 0.903 | 0.953 | Monocytes | aPD-L1 vs PBS |
| Ppia | 1.88E-09 | 3.35E-05 | -0.35 | 0.939 | 0.962 | Monocytes | aPD-L1 vs PBS |
| Klf2 | 5.17E-10 | 9.22E-06 | -0.35 | 0.947 | 0.959 | T cells | aPD-L1 vs PBS |
| Ms4a6d | 1.95E-14 | 3.48E-10 | -0.35 | 0.825 | 0.896 | Macrophages | aPD-L1 vs PBS |
| Lgals1 | 2.00E-42 | 3.57E-38 | -0.35 | 0.192 | 0.615 | B cells | aPD-L1 vs PBS |
| Gapdh | 2.95E-18 | 5.26E-14 | -0.35 | 0.955 | 0.995 | Macrophages | aPD-L1 vs PBS |
| Cst3 | 2.84E-22 | 5.06E-18 | -0.36 | 0.647 | 0.835 | B cells | aPD-L1 vs PBS |
| Gatm | 1.62E-11 | 2.88E-07 | -0.36 | 0.638 | 0.792 | Macrophages | aPD-L1 vs PBS |
| Pfn1 | 1.77E-16 | 3.16E-12 | -0.36 | 0.985 | 0.994 | Monocytes | aPD-L1 vs PBS |
| Plau | 1.34E-08 | 2.39E-04 | -0.36 | 0.366 | 0.536 | Macrophages | aPD-L1 vs PBS |
| Actg1 | 1.45E-15 | 2.58E-11 | -0.36 | 0.997 | 1 | Neutrophils | aPD-L1 vs PBS |
| Acta2 | 9.36E-15 | 1.67E-10 | -0.36 | 0.575 | 0.722 | Tumor_cells | aPD-L1 vs PBS |
| Rpl10a | 1.06E-20 | 1.89E-16 | -0.36 | 0.918 | 0.976 | Macrophages | aPD-L1 vs PBS |
| Fos | 3.34E-08 | 5.95E-04 | -0.37 | 0.849 | 0.867 | T cells | aPD-L1 vs PBS |
| Rps5 | 1.16E-20 | 2.07E-16 | -0.37 | 0.978 | 0.989 | Macrophages | aPD-L1 vs PBS |
| Ifitm2 | 6.84E-09 | 1.22E-04 | -0.37 | 0.903 | 0.962 | Monocytes | aPD-L1 vs PBS |
| G0s2 | 9.44E-08 | 1.68E-03 | -0.37 | 0.836 | 0.865 | Neutrophils | aPD-L1 vs PBS |
| Cfl1 | 2.09E-06 | 3.72E-02 | -0.37 | 0.956 | 0.981 | ILC | aPD-L1 vs PBS |
| Pfn1 | 1.09E-23 | 1.94E-19 | -0.37 | 0.908 | 0.966 | B cells | aPD-L1 vs PBS |
| Prelid1 | 9.78E-13 | 1.74E-08 | -0.38 | 0.372 | 0.649 | Monocytes | aPD-L1 vs PBS |
| Nfkbia | 2.42E-10 | 4.31E-06 | -0.38 | 0.795 | 0.886 | B cells | aPD-L1 vs PBS |
| Lgals1 | 5.64E-22 | 1.00E-17 | -0.38 | 0.966 | 0.987 | Macrophages | aPD-L1 vs PBS |
| Gimap7 | 1.11E-06 | 1.98E-02 | -0.38 | 0.209 | 0.469 | NKT | aPD-L1 vs PBS |
| Eif5a | 5.77E-13 | 1.03E-08 | -0.38 | 0.658 | 0.858 | Monocytes | aPD-L1 vs PBS |
| Itm2b | 5.99E-08 | 1.07E-03 | -0.39 | 0.928 | 0.95 | NKT | aPD-L1 vs PBS |
| Ms4a6c | 6.41E-11 | 1.14E-06 | -0.39 | 0.832 | 0.905 | Monocytes | aPD-L1 vs PBS |
| Sept1 | 1.43E-06 | 2.55E-02 | -0.39 | 0.474 | 0.71 | ILC | aPD-L1 vs PBS |
| Actb | 3.43E-07 | 6.11E-03 | -0.39 | 1 | 1 | NKT | aPD-L1 vs PBS |
| Fos | 5.96E-10 | 1.06E-05 | -0.39 | 0.971 | 0.994 | Neutrophils | aPD-L1 vs PBS |
| Dab2 | 2.71E-21 | 4.83E-17 | -0.41 | 0.638 | 0.834 | Macrophages | aPD-L1 vs PBS |
| Arl5a | 1.16E-06 | 2.08E-02 | -0.42 | 0.101 | 0.338 | NKT | aPD-L1 vs PBS |
| Ldha | 9.11E-19 | 1.62E-14 | -0.42 | 0.833 | 0.937 | Neutrophils | aPD-L1 vs PBS |
| Actb | 8.54E-10 | 1.52E-05 | -0.42 | 1 | 1 | ILC | aPD-L1 vs PBS |
| Sh3bgrl3 | 1.47E-07 | 2.61E-03 | -0.43 | 0.881 | 0.935 | ILC | aPD-L1 vs PBS |
| Jun | 1.62E-20 | 2.89E-16 | -0.43 | 0.922 | 0.994 | B cells | aPD-L1 vs PBS |
| Ms4a7 | 8.76E-13 | 1.56E-08 | -0.45 | 0.455 | 0.681 | Macrophages | aPD-L1 vs PBS |
| Cthrc1 | 3.06E-32 | 5.46E-28 | -0.47 | 0.417 | 0.635 | Tumor_cells | aPD-L1 vs PBS |
| Psmb8 | 3.63E-11 | 6.47E-07 | -0.47 | 0.837 | 0.953 | ILC | aPD-L1 vs PBS |
| Lgals1 | 1.08E-09 | 1.93E-05 | -0.47 | 0.781 | 0.911 | Monocytes | aPD-L1 vs PBS |
| Pglyrp1 | 3.43E-08 | 6.11E-04 | -0.51 | 0.108 | 0.375 | NKT | aPD-L1 vs PBS |
| Pcolce | 2.13E-06 | 3.80E-02 | -0.53 | 0.759 | 0.879 | Fibroblasts | aPD-L1 vs PBS |
| Pmepa1 | 7.28E-23 | 1.30E-18 | -0.54 | 0.384 | 0.701 | Macrophages | aPD-L1 vs PBS |
| AW112010 | 9.95E-07 | 1.77E-02 | -0.55 | 0.748 | 0.916 | ILC | aPD-L1 vs PBS |
| Cdkn1a | 1.16E-28 | 2.07E-24 | -0.56 | 0.699 | 0.817 | Neutrophils | aPD-L1 vs PBS |
| Pid1 | 1.42E-12 | 2.53E-08 | -0.57 | 0.628 | 0.835 | Monocytes | aPD-L1 vs PBS |
| Pfn1 | 2.43E-13 | 4.34E-09 | -0.58 | 0.993 | 0.988 | NKT | aPD-L1 vs PBS |
| Pfn1 | 3.29E-15 | 5.87E-11 | -0.59 | 0.978 | 0.991 | ILC | aPD-L1 vs PBS |
| Pf4 | 6.69E-12 | 1.19E-07 | -0.60 | 0.239 | 0.483 | Macrophages | aPD-L1 vs PBS |
| Hspa1b | 3.32E-11 | 5.91E-07 | -0.60 | 0.45 | 0.593 | B cells | aPD-L1 vs PBS |

| | | | | | | | |
|---|---|---|---|---|---|---|---|
| Fos | 8.87E-34 | 1.58E-29 | -0.61 | 0.892 | 0.969 | B cells | aPD-L1 vs PBS |
| Spp1 | 1.49E-25 | 2.65E-21 | -0.63 | 0.5 | 0.794 | Macrophages | aPD-L1 vs PBS |
| Ly6a | 3.88E-07 | 6.92E-03 | -1.04 | 0.288 | 0.5 | NKT | aPD-L1 vs PBS |
| Acta2 | 1.19E-06 | 2.13E-02 | -1.31 | 0.02 | 0.162 | Epithelial cells | aPD-L1 vs PBS |
| Cxcl10 | 1.69E-17 | 3.01E-13 | 1.27 | 0.511 | 0.081 | Fibroblasts | aPD-L1+aTGF-β vs PBS |
| Irf1 | 1.95E-40 | 3.48E-36 | 0.88 | 0.806 | 0.396 | Monocytes | aPD-L1+aTGF-β vs PBS |
| Igkc | 1.57E-27 | 2.79E-23 | 0.87 | 0.162 | 0.025 | Macrophages | aPD-L1+aTGF-β vs PBS |
| Klf2 | 1.56E-12 | 2.78E-08 | 0.87 | 0.883 | 0.5 | Epithelial cells | aPD-L1+aTGF-β vs PBS |
| Mt1 | 1.38E-09 | 2.45E-05 | 0.84 | 0.794 | 0.412 | Epithelial cells | aPD-L1+aTGF-β vs PBS |
| Cxcl9 | 2.04E-07 | 3.64E-03 | 0.79 | 0.324 | 0.094 | Fibroblasts | aPD-L1+aTGF-β vs PBS |
| H2-Eb1 | 1.04E-34 | 1.86E-30 | 0.78 | 0.709 | 0.502 | Macrophages | aPD-L1+aTGF-β vs PBS |
| Irf1 | 4.81E-63 | 8.57E-59 | 0.72 | 0.686 | 0.375 | Neutrophils | aPD-L1+aTGF-β vs PBS |
| H2-Aa | 3.48E-30 | 6.20E-26 | 0.70 | 0.712 | 0.537 | Macrophages | aPD-L1+aTGF-β vs PBS |
| Cd74 | 1.57E-30 | 2.79E-26 | 0.70 | 0.868 | 0.805 | Macrophages | aPD-L1+aTGF-β vs PBS |
| H2-Ab1 | 1.95E-27 | 3.48E-23 | 0.67 | 0.728 | 0.586 | Macrophages | aPD-L1+aTGF-β vs PBS |
| Ly6a | 7.37E-08 | 1.31E-03 | 0.63 | 0.687 | 0.324 | Epithelial cells | aPD-L1+aTGF-β vs PBS |
| Cxcl10 | 3.96E-25 | 7.06E-21 | 0.63 | 0.593 | 0.37 | Macrophages | aPD-L1+aTGF-β vs PBS |
| Ly6a | 2.63E-18 | 4.69E-14 | 0.62 | 0.558 | 0.42 | Macrophages | aPD-L1+aTGF-β vs PBS |
| Gm10076 | 4.35E-66 | 7.75E-62 | 0.61 | 0.899 | 0.588 | T cells | aPD-L1+aTGF-β vs PBS |
| Gm10076 | 1.40E-16 | 2.50E-12 | 0.61 | 0.854 | 0.495 | ILC | aPD-L1+aTGF-β vs PBS |
| B2m | 1.44E-11 | 2.56E-07 | 0.61 | 0.993 | 0.882 | Epithelial cells | aPD-L1+aTGF-β vs PBS |
| Tmsb10 | 2.93E-61 | 5.23E-57 | 0.61 | 0.971 | 0.917 | Macrophages | aPD-L1+aTGF-β vs PBS |
| Gm10076 | 3.69E-52 | 6.58E-48 | 0.60 | 0.831 | 0.49 | B cells | aPD-L1+aTGF-β vs PBS |
| Irf1 | 2.52E-12 | 4.50E-08 | 0.60 | 0.633 | 0.295 | Fibroblasts | aPD-L1+aTGF-β vs PBS |
| Dnaja1 | 1.62E-07 | 2.88E-03 | 0.59 | 0.897 | 0.841 | ILC | aPD-L1+aTGF-β vs PBS |
| Tmsb4x | 7.38E-07 | 1.32E-02 | 0.58 | 0.993 | 0.971 | Epithelial cells | aPD-L1+aTGF-β vs PBS |
| Cxcl2 | 5.44E-57 | 9.70E-53 | 0.58 | 0.476 | 0.267 | Tumor_cells | aPD-L1+aTGF-β vs PBS |
| AW112010 | 7.08E-22 | 1.26E-17 | 0.57 | 0.696 | 0.544 | Macrophages | aPD-L1+aTGF-β vs PBS |
| Wfdc17 | 1.77E-09 | 3.16E-05 | 0.57 | 0.372 | 0.259 | Macrophages | aPD-L1+aTGF-β vs PBS |
| Irf1 | 2.74E-57 | 4.89E-53 | 0.56 | 0.548 | 0.22 | Macrophages | aPD-L1+aTGF-β vs PBS |
| Cxcl1 | 2.38E-96 | 4.23E-92 | 0.56 | 0.797 | 0.537 | Tumor_cells | aPD-L1+aTGF-β vs PBS |
| Samhd1 | 2.69E-40 | 4.80E-36 | 0.55 | 0.69 | 0.476 | Neutrophils | aPD-L1+aTGF-β vs PBS |
| Psmb9 | 3.96E-08 | 7.06E-04 | 0.55 | 0.618 | 0.279 | Epithelial cells | aPD-L1+aTGF-β vs PBS |
| Rpl39 | 5.74E-25 | 1.02E-20 | 0.54 | 1 | 0.994 | NKT | aPD-L1+aTGF-β vs PBS |
| H2-D1 | 4.27E-13 | 7.61E-09 | 0.53 | 0.995 | 0.946 | Fibroblasts | aPD-L1+aTGF-β vs PBS |
| Phlda1 | 5.67E-08 | 1.01E-03 | 0.53 | 0.654 | 0.362 | Fibroblasts | aPD-L1+aTGF-β vs PBS |
| Rsad2 | 6.31E-21 | 1.12E-16 | 0.53 | 0.426 | 0.232 | Macrophages | aPD-L1+aTGF-β vs PBS |
| Cd274 | 4.68E-18 | 8.34E-14 | 0.52 | 0.666 | 0.373 | Monocytes | aPD-L1+aTGF-β vs PBS |
| Mgp | 4.19E-25 | 7.48E-21 | 0.52 | 0.339 | 0.199 | Tumor_cells | aPD-L1+aTGF-β vs PBS |
| Cd274 | 6.92E-35 | 1.23E-30 | 0.52 | 0.701 | 0.475 | Neutrophils | aPD-L1+aTGF-β vs PBS |
| Rps27 | 8.46E-30 | 1.51E-25 | 0.52 | 0.988 | 0.94 | Monocytes | aPD-L1+aTGF-β vs PBS |
| Gbp2b | 6.25E-44 | 1.11E-39 | 0.51 | 0.603 | 0.312 | Neutrophils | aPD-L1+aTGF-β vs PBS |
| H2-D1 | 7.85E-14 | 1.40E-09 | 0.50 | 1 | 1 | Epithelial cells | aPD-L1+aTGF-β vs PBS |
| Gm10076 | 8.07E-94 | 1.44E-89 | 0.50 | 0.874 | 0.558 | Macrophages | aPD-L1+aTGF-β vs PBS |
| Selenom | 4.44E-11 | 7.92E-07 | 0.50 | 0.423 | 0.316 | Macrophages | aPD-L1+aTGF-β vs PBS |
| Psmb8 | 2.66E-08 | 4.74E-04 | 0.50 | 0.935 | 0.75 | Epithelial cells | aPD-L1+aTGF-β vs PBS |
| Gbp2 | 1.78E-44 | 3.17E-40 | 0.50 | 0.554 | 0.266 | Neutrophils | aPD-L1+aTGF-β vs PBS |
| B2m | 2.95E-11 | 5.26E-07 | 0.49 | 1 | 0.919 | Fibroblasts | aPD-L1+aTGF-β vs PBS |
| Oasl1 | 8.49E-27 | 1.51E-22 | 0.49 | 0.48 | 0.264 | Macrophages | aPD-L1+aTGF-β vs PBS |
| Il1b | 6.45E-10 | 1.15E-05 | 0.49 | 0.823 | 0.699 | Monocytes | aPD-L1+aTGF-β vs PBS |
| Rpl39 | 8.18E-09 | 1.46E-04 | 0.49 | 1 | 1 | DC | aPD-L1+aTGF-β vs PBS |
| Cxcl9 | 1.35E-14 | 2.40E-10 | 0.48 | 0.344 | 0.192 | Macrophages | aPD-L1+aTGF-β vs PBS |
| Gm10076 | 2.24E-14 | 3.99E-10 | 0.48 | 0.793 | 0.562 | NKT | aPD-L1+aTGF-β vs PBS |
| Ccl4 | 5.02E-22 | 8.94E-18 | 0.47 | 0.818 | 0.653 | Macrophages | aPD-L1+aTGF-β vs PBS |
| Crip1 | 5.46E-33 | 9.74E-29 | 0.47 | 0.919 | 0.83 | Macrophages | aPD-L1+aTGF-β vs PBS |

| | | | | | | | |
|---|---|---|---|---|---|---|---|
| Tmsb10 | 8.89E-18 | 1.58E-13 | 0.47 | 0.99 | 0.981 | Monocytes | aPD-L1+aTGF-β vs PBS |
| Rps27 | 5.75E-18 | 1.02E-13 | 0.47 | 0.997 | 1 | NKT | aPD-L1+aTGF-β vs PBS |
| H2-K1 | 1.93E-12 | 3.45E-08 | 0.47 | 0.989 | 0.913 | Fibroblasts | aPD-L1+aTGF-β vs PBS |
| H2-K1 | 1.14E-10 | 2.03E-06 | 0.47 | 0.998 | 0.956 | Epithelial cells | aPD-L1+aTGF-β vs PBS |
| Gm10076 | 2.64E-31 | 4.70E-27 | 0.46 | 0.794 | 0.446 | Monocytes | aPD-L1+aTGF-β vs PBS |
| Spp1 | 5.50E-51 | 9.80E-47 | 0.46 | 0.989 | 0.968 | Tumor_cells | aPD-L1+aTGF-β vs PBS |
| Rpl39 | 2.75E-88 | 4.91E-84 | 0.46 | 1 | 1 | T cells | aPD-L1+aTGF-β vs PBS |
| Irgm1 | 4.47E-37 | 7.96E-33 | 0.46 | 0.475 | 0.226 | Neutrophils | aPD-L1+aTGF-β vs PBS |
| Ifi27l2a | 1.14E-13 | 2.02E-09 | 0.46 | 0.65 | 0.468 | T cells | aPD-L1+aTGF-β vs PBS |
| H2-K1 | 2.47E-58 | 4.40E-54 | 0.45 | 0.998 | 0.989 | Neutrophils | aPD-L1+aTGF-β vs PBS |
| Arg2 | 8.17E-16 | 1.46E-11 | 0.44 | 0.603 | 0.326 | Monocytes | aPD-L1+aTGF-β vs PBS |
| Rpl39 | 4.99E-15 | 8.89E-11 | 0.44 | 1 | 1 | ILC | aPD-L1+aTGF-β vs PBS |
| Rps27 | 8.94E-20 | 1.59E-15 | 0.44 | 0.996 | 1 | ILC | aPD-L1+aTGF-β vs PBS |
| Icam1 | 2.49E-24 | 4.44E-20 | 0.44 | 0.265 | 0.099 | Neutrophils | aPD-L1+aTGF-β vs PBS |
| Rpl37 | 7.61E-20 | 1.36E-15 | 0.44 | 1 | 0.981 | NKT | aPD-L1+aTGF-β vs PBS |
| Rpl35a | 4.52E-08 | 8.06E-04 | 0.44 | 1 | 1 | DC | aPD-L1+aTGF-β vs PBS |
| Rpl37a | 1.08E-17 | 1.93E-13 | 0.43 | 1 | 1 | ILC | aPD-L1+aTGF-β vs PBS |
| Rpl39 | 1.81E-65 | 3.22E-61 | 0.43 | 0.996 | 1 | B cells | aPD-L1+aTGF-β vs PBS |
| Rpl36 | 9.32E-22 | 1.66E-17 | 0.42 | 0.994 | 0.981 | NKT | aPD-L1+aTGF-β vs PBS |
| Ifrd1 | 3.70E-07 | 6.59E-03 | 0.42 | 0.707 | 0.463 | Fibroblasts | aPD-L1+aTGF-β vs PBS |
| Rpl37a | 1.06E-72 | 1.89E-68 | 0.42 | 1 | 1 | T cells | aPD-L1+aTGF-β vs PBS |
| Gbp4 | 1.13E-24 | 2.01E-20 | 0.42 | 0.2 | 0.052 | Neutrophils | aPD-L1+aTGF-β vs PBS |
| Socs1 | 1.09E-12 | 1.94E-08 | 0.42 | 0.453 | 0.218 | Monocytes | aPD-L1+aTGF-β vs PBS |
| Nfkbia | 3.53E-93 | 6.28E-89 | 0.42 | 0.919 | 0.747 | Tumor_cells | aPD-L1+aTGF-β vs PBS |
| Zfand5 | 1.64E-92 | 2.93E-88 | 0.42 | 0.712 | 0.473 | Tumor_cells | aPD-L1+aTGF-β vs PBS |
| Irf1 | 8.74E-100 | 1.56E-95 | 0.41 | 0.639 | 0.334 | Tumor_cells | aPD-L1+aTGF-β vs PBS |
| Irf1 | 4.13E-21 | 7.37E-17 | 0.41 | 0.612 | 0.375 | T cells | aPD-L1+aTGF-β vs PBS |
| Ctla4 | 1.15E-09 | 2.06E-05 | 0.41 | 0.205 | 0.081 | T cells | aPD-L1+aTGF-β vs PBS |
| Upp1 | 4.63E-07 | 8.25E-03 | 0.41 | 0.511 | 0.354 | Monocytes | aPD-L1+aTGF-β vs PBS |
| Rpl39 | 1.37E-21 | 2.44E-17 | 0.41 | 0.978 | 0.943 | Monocytes | aPD-L1+aTGF-β vs PBS |
| Rpl37a | 2.99E-20 | 5.32E-16 | 0.41 | 1 | 0.994 | NKT | aPD-L1+aTGF-β vs PBS |
| Tnf | 4.36E-27 | 7.78E-23 | 0.41 | 0.502 | 0.28 | Macrophages | aPD-L1+aTGF-β vs PBS |
| Rps21 | 2.92E-68 | 5.21E-64 | 0.41 | 0.998 | 1 | B cells | aPD-L1+aTGF-β vs PBS |
| Rps27 | 1.20E-06 | 2.15E-02 | 0.41 | 1 | 1 | DC | aPD-L1+aTGF-β vs PBS |
| Irf1 | 6.40E-21 | 1.14E-16 | 0.41 | 0.477 | 0.197 | B cells | aPD-L1+aTGF-β vs PBS |
| Rps21 | 6.22E-18 | 1.11E-13 | 0.41 | 1 | 1 | NKT | aPD-L1+aTGF-β vs PBS |
| Rpl37 | 2.39E-63 | 4.25E-59 | 0.40 | 1 | 1 | T cells | aPD-L1+aTGF-β vs PBS |
| Sgk1 | 5.10E-74 | 9.08E-70 | 0.40 | 0.751 | 0.501 | Tumor_cells | aPD-L1+aTGF-β vs PBS |
| Rpl37 | 2.03E-17 | 3.62E-13 | 0.40 | 1 | 0.991 | ILC | aPD-L1+aTGF-β vs PBS |
| Fcgr4 | 3.16E-13 | 5.63E-09 | 0.40 | 0.593 | 0.496 | Macrophages | aPD-L1+aTGF-β vs PBS |
| Rpl37a | 3.09E-60 | 5.51E-56 | 0.40 | 0.998 | 1 | B cells | aPD-L1+aTGF-β vs PBS |
| Nfe2l2 | 1.00E-11 | 1.78E-07 | 0.40 | 0.554 | 0.316 | Monocytes | aPD-L1+aTGF-β vs PBS |
| Tnfaip2 | 4.24E-22 | 7.56E-18 | 0.40 | 0.563 | 0.386 | Neutrophils | aPD-L1+aTGF-β vs PBS |
| Mt1 | 1.43E-06 | 2.55E-02 | 0.40 | 0.957 | 0.832 | Fibroblasts | aPD-L1+aTGF-β vs PBS |
| Nfkbia | 1.77E-14 | 3.15E-10 | 0.39 | 0.944 | 0.807 | Monocytes | aPD-L1+aTGF-β vs PBS |
| Rpl34 | 9.12E-19 | 1.63E-14 | 0.39 | 1 | 1 | NKT | aPD-L1+aTGF-β vs PBS |
| Serpina3g | 1.13E-37 | 2.01E-33 | 0.39 | 0.308 | 0.083 | Macrophages | aPD-L1+aTGF-β vs PBS |
| Rpl35 | 1.63E-10 | 2.90E-06 | 0.39 | 0.963 | 0.869 | NKT | aPD-L1+aTGF-β vs PBS |
| Rpl37a | 5.16E-24 | 9.21E-20 | 0.39 | 0.99 | 0.962 | Monocytes | aPD-L1+aTGF-β vs PBS |
| Rps27 | 3.61E-77 | 6.44E-73 | 0.39 | 0.999 | 1 | B cells | aPD-L1+aTGF-β vs PBS |
| Gbp2b | 2.18E-12 | 3.89E-08 | 0.39 | 0.646 | 0.396 | Monocytes | aPD-L1+aTGF-β vs PBS |
| Rps21 | 1.82E-14 | 3.25E-10 | 0.39 | 1 | 0.981 | ILC | aPD-L1+aTGF-β vs PBS |
| Dusp5 | 2.07E-12 | 3.69E-08 | 0.39 | 0.554 | 0.31 | Monocytes | aPD-L1+aTGF-β vs PBS |
| Tmsb10 | 7.95E-13 | 1.42E-08 | 0.39 | 1 | 1 | NKT | aPD-L1+aTGF-β vs PBS |
| H2-D1 | 3.39E-45 | 6.04E-41 | 0.38 | 1 | 0.999 | Neutrophils | aPD-L1+aTGF-β vs PBS |

| | | | | | | | |
|---|---|---|---|---|---|---|---|
| Vps37b | 1.12E-12 | 2.00E-08 | 0.38 | 0.593 | 0.396 | B cells | aPD-L1+aTGF-β vs PBS |
| Nlrp3 | 2.05E-17 | 3.66E-13 | 0.38 | 0.707 | 0.59 | Neutrophils | aPD-L1+aTGF-β vs PBS |
| Cxcl10 | 7.04E-07 | 1.26E-02 | 0.38 | 0.109 | 0.033 | T cells | aPD-L1+aTGF-β vs PBS |
| Rpl36 | 2.50E-60 | 4.46E-56 | 0.38 | 1 | 1 | T cells | aPD-L1+aTGF-β vs PBS |
| Gm10260 | 6.53E-11 | 1.16E-06 | 0.38 | 0.92 | 0.819 | NKT | aPD-L1+aTGF-β vs PBS |
| Dcn | 2.26E-33 | 4.03E-29 | 0.38 | 0.252 | 0.106 | Tumor_cells | aPD-L1+aTGF-β vs PBS |
| Plaur | 5.36E-10 | 9.56E-06 | 0.38 | 0.634 | 0.43 | Monocytes | aPD-L1+aTGF-β vs PBS |
| Rpl39 | 1.31E-61 | 2.34E-57 | 0.38 | 0.991 | 0.98 | Macrophages | aPD-L1+aTGF-β vs PBS |
| Hspa8 | 1.57E-06 | 2.80E-02 | 0.38 | 0.993 | 0.981 | ILC | aPD-L1+aTGF-β vs PBS |
| Cxcl10 | 8.54E-35 | 1.52E-30 | 0.37 | 0.176 | 0.051 | Tumor_cells | aPD-L1+aTGF-β vs PBS |
| Rel | 1.31E-11 | 2.33E-07 | 0.37 | 0.525 | 0.291 | Monocytes | aPD-L1+aTGF-β vs PBS |
| Rpl37 | 1.09E-47 | 1.94E-43 | 0.37 | 0.999 | 1 | B cells | aPD-L1+aTGF-β vs PBS |
| Ccl5 | 7.23E-10 | 1.29E-05 | 0.37 | 0.178 | 0.083 | Macrophages | aPD-L1+aTGF-β vs PBS |
| Ifrd1 | 2.91E-69 | 5.19E-65 | 0.37 | 0.802 | 0.614 | Tumor_cells | aPD-L1+aTGF-β vs PBS |
| S100a8 | 4.74E-14 | 8.45E-10 | 0.37 | 0.946 | 0.935 | Neutrophils | aPD-L1+aTGF-β vs PBS |
| Rpl34 | 2.78E-12 | 4.95E-08 | 0.36 | 1 | 1 | ILC | aPD-L1+aTGF-β vs PBS |
| Dennd4a | 2.35E-09 | 4.19E-05 | 0.36 | 0.438 | 0.256 | Monocytes | aPD-L1+aTGF-β vs PBS |
| Rps15a | 3.16E-12 | 5.64E-08 | 0.36 | 1 | 1 | NKT | aPD-L1+aTGF-β vs PBS |
| Txn1 | 6.97E-32 | 1.24E-27 | 0.36 | 0.969 | 0.944 | Macrophages | aPD-L1+aTGF-β vs PBS |
| Sub1 | 1.15E-06 | 2.04E-02 | 0.36 | 0.744 | 0.625 | NKT | aPD-L1+aTGF-β vs PBS |
| Cytip | 1.01E-10 | 1.80E-06 | 0.36 | 0.668 | 0.468 | Monocytes | aPD-L1+aTGF-β vs PBS |
| Gm10260 | 8.30E-14 | 1.48E-09 | 0.36 | 0.705 | 0.513 | Monocytes | aPD-L1+aTGF-β vs PBS |
| Pde4b | 3.62E-12 | 6.44E-08 | 0.35 | 0.518 | 0.291 | Monocytes | aPD-L1+aTGF-β vs PBS |
| Rpl36 | 1.66E-45 | 2.96E-41 | 0.35 | 0.997 | 0.997 | B cells | aPD-L1+aTGF-β vs PBS |
| Kdm6b | 5.82E-14 | 1.04E-09 | 0.35 | 0.671 | 0.44 | Monocytes | aPD-L1+aTGF-β vs PBS |
| Rps27 | 3.58E-45 | 6.38E-41 | 0.35 | 0.987 | 0.974 | Macrophages | aPD-L1+aTGF-β vs PBS |
| Nampt | 4.35E-21 | 7.75E-17 | 0.35 | 0.483 | 0.299 | Neutrophils | aPD-L1+aTGF-β vs PBS |
| Ifi47 | 2.67E-27 | 4.76E-23 | 0.35 | 0.498 | 0.272 | Neutrophils | aPD-L1+aTGF-β vs PBS |
| Tmsb10 | 5.01E-44 | 8.93E-40 | 0.35 | 0.999 | 0.998 | T cells | aPD-L1+aTGF-β vs PBS |
| Rpl35a | 4.80E-50 | 8.56E-46 | 0.35 | 0.997 | 0.997 | B cells | aPD-L1+aTGF-β vs PBS |
| Mt2 | 4.90E-55 | 8.74E-51 | 0.35 | 0.763 | 0.561 | Tumor_cells | aPD-L1+aTGF-β vs PBS |
| Rpl36a | 1.60E-31 | 2.86E-27 | 0.34 | 0.982 | 0.954 | B cells | aPD-L1+aTGF-β vs PBS |
| Fosb | 3.70E-43 | 6.59E-39 | 0.34 | 0.642 | 0.465 | Tumor_cells | aPD-L1+aTGF-β vs PBS |
| Fos | 2.70E-52 | 4.82E-48 | 0.34 | 0.881 | 0.694 | Tumor_cells | aPD-L1+aTGF-β vs PBS |
| Klf2 | 1.48E-29 | 2.65E-25 | 0.34 | 0.427 | 0.272 | Tumor_cells | aPD-L1+aTGF-β vs PBS |
| Gm10260 | 7.18E-35 | 1.28E-30 | 0.34 | 0.98 | 0.937 | T cells | aPD-L1+aTGF-β vs PBS |
| Ifrd1 | 7.75E-12 | 1.38E-07 | 0.34 | 0.843 | 0.623 | Monocytes | aPD-L1+aTGF-β vs PBS |
| Tnfaip2 | 2.05E-12 | 3.66E-08 | 0.34 | 0.438 | 0.193 | Monocytes | aPD-L1+aTGF-β vs PBS |
| Rpl41 | 1.40E-45 | 2.49E-41 | 0.34 | 1 | 1 | T cells | aPD-L1+aTGF-β vs PBS |
| Zbp1 | 2.52E-24 | 4.49E-20 | 0.34 | 0.412 | 0.208 | Neutrophils | aPD-L1+aTGF-β vs PBS |
| Cd83 | 1.83E-09 | 3.27E-05 | 0.34 | 0.571 | 0.373 | Monocytes | aPD-L1+aTGF-β vs PBS |
| Acod1 | 5.12E-26 | 9.12E-22 | 0.34 | 0.355 | 0.148 | Macrophages | aPD-L1+aTGF-β vs PBS |
| Rpl36 | 1.58E-11 | 2.82E-07 | 0.34 | 1 | 1 | ILC | aPD-L1+aTGF-β vs PBS |
| Gm10260 | 3.30E-08 | 5.88E-04 | 0.34 | 0.94 | 0.832 | ILC | aPD-L1+aTGF-β vs PBS |
| B2m | 2.23E-43 | 3.97E-39 | 0.34 | 1 | 0.998 | Neutrophils | aPD-L1+aTGF-β vs PBS |
| Plac8 | 2.22E-18 | 3.96E-14 | 0.34 | 0.364 | 0.197 | Neutrophils | aPD-L1+aTGF-β vs PBS |
| Ifi27l2a | 1.35E-21 | 2.40E-17 | 0.33 | 0.887 | 0.799 | Macrophages | aPD-L1+aTGF-β vs PBS |
| Rpl37 | 5.72E-22 | 1.02E-17 | 0.33 | 0.99 | 0.94 | Monocytes | aPD-L1+aTGF-β vs PBS |
| Fgl2 | 1.90E-21 | 3.39E-17 | 0.33 | 0.468 | 0.277 | Macrophages | aPD-L1+aTGF-β vs PBS |
| Gbp2b | 8.39E-24 | 1.50E-19 | 0.33 | 0.369 | 0.169 | Macrophages | aPD-L1+aTGF-β vs PBS |
| Rpl41 | 9.21E-40 | 1.64E-35 | 0.33 | 1 | 0.997 | B cells | aPD-L1+aTGF-β vs PBS |
| Samhd1 | 9.96E-09 | 1.78E-04 | 0.33 | 0.867 | 0.772 | Monocytes | aPD-L1+aTGF-β vs PBS |
| Rpl22 | 8.44E-13 | 1.50E-08 | 0.33 | 0.986 | 0.975 | NKT | aPD-L1+aTGF-β vs PBS |
| Hcst | 1.19E-06 | 2.13E-02 | 0.33 | 0.811 | 0.617 | ILC | aPD-L1+aTGF-β vs PBS |
| Ccl4 | 9.79E-13 | 1.74E-08 | 0.33 | 0.722 | 0.591 | Neutrophils | aPD-L1+aTGF-β vs PBS |

| | | | | | | | |
|---|---|---|---|---|---|---|---|
| Rps27 | 1.65E-56 | 2.94E-52 | 0.33 | 1 | 1 | T cells | aPD-L1+aTGF-β vs PBS |
| Upp1 | 3.22E-15 | 5.73E-11 | 0.33 | 0.686 | 0.547 | Neutrophils | aPD-L1+aTGF-β vs PBS |
| Rpl35 | 9.73E-37 | 1.73E-32 | 0.33 | 0.997 | 0.985 | T cells | aPD-L1+aTGF-β vs PBS |
| Acod1 | 1.20E-20 | 2.14E-16 | 0.33 | 0.856 | 0.75 | Neutrophils | aPD-L1+aTGF-β vs PBS |
| Rpl35a | 9.99E-11 | 1.78E-06 | 0.33 | 0.997 | 0.988 | NKT | aPD-L1+aTGF-β vs PBS |
| Gbp5 | 1.37E-26 | 2.44E-22 | 0.33 | 0.311 | 0.118 | Neutrophils | aPD-L1+aTGF-β vs PBS |
| Rps21 | 2.14E-17 | 3.82E-13 | 0.32 | 0.961 | 0.943 | Monocytes | aPD-L1+aTGF-β vs PBS |
| Rpl37a | 1.88E-72 | 3.35E-68 | 0.32 | 0.997 | 1 | Macrophages | aPD-L1+aTGF-β vs PBS |
| Adgre5 | 7.74E-10 | 1.38E-05 | 0.32 | 0.521 | 0.291 | Monocytes | aPD-L1+aTGF-β vs PBS |
| Ifit1 | 2.17E-17 | 3.87E-13 | 0.32 | 0.258 | 0.113 | Macrophages | aPD-L1+aTGF-β vs PBS |
| Rps21 | 9.86E-49 | 1.76E-44 | 0.32 | 1 | 1 | T cells | aPD-L1+aTGF-β vs PBS |
| F10 | 3.23E-11 | 5.76E-07 | 0.32 | 0.438 | 0.203 | Monocytes | aPD-L1+aTGF-β vs PBS |
| Fcgr4 | 8.35E-21 | 1.49E-16 | 0.32 | 0.393 | 0.215 | Neutrophils | aPD-L1+aTGF-β vs PBS |
| Rpl41 | 2.52E-11 | 4.50E-07 | 0.32 | 1 | 0.994 | NKT | aPD-L1+aTGF-β vs PBS |
| Parp14 | 1.59E-19 | 2.83E-15 | 0.32 | 0.514 | 0.324 | Neutrophils | aPD-L1+aTGF-β vs PBS |
| H2-DMb1 | 9.58E-17 | 1.71E-12 | 0.32 | 0.515 | 0.356 | Macrophages | aPD-L1+aTGF-β vs PBS |
| Slc7a11 | 1.13E-15 | 2.01E-11 | 0.32 | 0.669 | 0.532 | Neutrophils | aPD-L1+aTGF-β vs PBS |
| Phlda1 | 1.22E-43 | 2.17E-39 | 0.32 | 0.916 | 0.772 | Tumor_cells | aPD-L1+aTGF-β vs PBS |
| AW112010 | 2.16E-10 | 3.85E-06 | 0.32 | 0.57 | 0.392 | T cells | aPD-L1+aTGF-β vs PBS |
| Gbp7 | 8.83E-19 | 1.57E-14 | 0.31 | 0.43 | 0.258 | Neutrophils | aPD-L1+aTGF-β vs PBS |
| Uba52 | 4.97E-21 | 8.86E-17 | 0.31 | 0.645 | 0.399 | T cells | aPD-L1+aTGF-β vs PBS |
| Rpl35a | 7.51E-17 | 1.34E-12 | 0.31 | 0.969 | 0.934 | Monocytes | aPD-L1+aTGF-β vs PBS |
| Rpl38 | 1.98E-10 | 3.53E-06 | 0.31 | 0.996 | 0.991 | ILC | aPD-L1+aTGF-β vs PBS |
| Nupr1 | 7.55E-50 | 1.35E-45 | 0.31 | 0.779 | 0.606 | Tumor_cells | aPD-L1+aTGF-β vs PBS |
| Nr4a1 | 1.83E-08 | 3.26E-04 | 0.31 | 0.596 | 0.386 | Monocytes | aPD-L1+aTGF-β vs PBS |
| Stat1 | 1.81E-17 | 3.22E-13 | 0.31 | 0.509 | 0.27 | T cells | aPD-L1+aTGF-β vs PBS |
| Sod2 | 1.41E-15 | 2.51E-11 | 0.31 | 0.331 | 0.185 | Neutrophils | aPD-L1+aTGF-β vs PBS |
| Igtp | 1.16E-21 | 2.06E-17 | 0.31 | 0.392 | 0.209 | Neutrophils | aPD-L1+aTGF-β vs PBS |
| Neat1 | 1.52E-43 | 2.71E-39 | 0.31 | 0.792 | 0.639 | Tumor_cells | aPD-L1+aTGF-β vs PBS |
| Blvrb | 1.66E-18 | 2.97E-14 | 0.31 | 0.615 | 0.464 | Macrophages | aPD-L1+aTGF-β vs PBS |
| Marcksl1 | 3.08E-12 | 5.49E-08 | 0.31 | 0.593 | 0.479 | Neutrophils | aPD-L1+aTGF-β vs PBS |
| Socs1 | 1.49E-20 | 2.65E-16 | 0.31 | 0.343 | 0.168 | Neutrophils | aPD-L1+aTGF-β vs PBS |
| Samsn1 | 1.20E-10 | 2.14E-06 | 0.31 | 0.437 | 0.262 | B cells | aPD-L1+aTGF-β vs PBS |
| Tnf | 1.83E-10 | 3.27E-06 | 0.31 | 0.24 | 0.137 | Neutrophils | aPD-L1+aTGF-β vs PBS |
| Cxcl10 | 1.60E-12 | 2.86E-08 | 0.31 | 0.172 | 0.07 | Neutrophils | aPD-L1+aTGF-β vs PBS |
| Ubc | 6.63E-08 | 1.18E-03 | 0.31 | 0.908 | 0.826 | Monocytes | aPD-L1+aTGF-β vs PBS |
| Ccl3 | 1.31E-11 | 2.34E-07 | 0.30 | 0.615 | 0.461 | Macrophages | aPD-L1+aTGF-β vs PBS |
| Rps20 | 1.70E-08 | 3.02E-04 | 0.30 | 1 | 0.994 | NKT | aPD-L1+aTGF-β vs PBS |
| Rpl35a | 1.95E-08 | 3.47E-04 | 0.30 | 0.996 | 0.972 | ILC | aPD-L1+aTGF-β vs PBS |
| Igkc | 8.52E-08 | 1.52E-03 | 0.30 | 0.179 | 0.044 | Monocytes | aPD-L1+aTGF-β vs PBS |
| Rpl36a | 9.12E-27 | 1.62E-22 | 0.30 | 0.989 | 0.978 | T cells | aPD-L1+aTGF-β vs PBS |
| Spag9 | 6.87E-11 | 1.23E-06 | 0.30 | 0.719 | 0.519 | Monocytes | aPD-L1+aTGF-β vs PBS |
| Rpl30 | 5.24E-11 | 9.34E-07 | 0.30 | 1 | 0.994 | NKT | aPD-L1+aTGF-β vs PBS |
| Rpl34 | 6.64E-15 | 1.18E-10 | 0.30 | 0.985 | 0.975 | Monocytes | aPD-L1+aTGF-β vs PBS |
| Rpl35 | 1.01E-25 | 1.81E-21 | 0.30 | 0.989 | 0.986 | B cells | aPD-L1+aTGF-β vs PBS |
| Uba52 | 3.74E-08 | 6.67E-04 | 0.30 | 0.529 | 0.262 | NKT | aPD-L1+aTGF-β vs PBS |
| Rpl34 | 2.45E-41 | 4.38E-37 | 0.30 | 1 | 1 | B cells | aPD-L1+aTGF-β vs PBS |
| Mirt1 | 3.94E-13 | 7.03E-09 | 0.30 | 0.547 | 0.422 | Neutrophils | aPD-L1+aTGF-β vs PBS |
| Sqstm1 | 4.10E-61 | 7.31E-57 | 0.30 | 0.686 | 0.465 | Tumor_cells | aPD-L1+aTGF-β vs PBS |
| Socs3 | 2.56E-09 | 4.57E-05 | 0.30 | 0.765 | 0.589 | Monocytes | aPD-L1+aTGF-β vs PBS |
| Gadd45b | 2.33E-12 | 4.16E-08 | 0.30 | 0.525 | 0.329 | T cells | aPD-L1+aTGF-β vs PBS |
| Rpl35 | 2.51E-06 | 4.48E-02 | 0.30 | 0.975 | 0.916 | ILC | aPD-L1+aTGF-β vs PBS |
| Rps15a | 1.81E-07 | 3.22E-03 | 0.30 | 1 | 1 | ILC | aPD-L1+aTGF-β vs PBS |
| Rps21 | 5.13E-45 | 9.15E-41 | 0.30 | 0.988 | 0.982 | Macrophages | aPD-L1+aTGF-β vs PBS |
| Klf2 | 1.58E-13 | 2.81E-09 | 0.30 | 0.764 | 0.626 | Macrophages | aPD-L1+aTGF-β vs PBS |

| | | | | | | | |
|---|---|---|---|---|---|---|---|
| Nfkbiz | 5.88E-57 | 1.05E-52 | 0.29 | 0.528 | 0.305 | Tumor_cells | aPD-L1+aTGF-β vs PBS |
| Ubc | 5.37E-16 | 9.57E-12 | 0.29 | 0.827 | 0.719 | Macrophages | aPD-L1+aTGF-β vs PBS |
| Rpl37 | 7.52E-52 | 1.34E-47 | 0.29 | 0.993 | 0.996 | Macrophages | aPD-L1+aTGF-β vs PBS |
| Dusp1 | 7.32E-35 | 1.30E-30 | 0.29 | 0.822 | 0.664 | Tumor_cells | aPD-L1+aTGF-β vs PBS |
| Crem | 3.34E-14 | 5.95E-10 | 0.29 | 0.323 | 0.125 | B cells | aPD-L1+aTGF-β vs PBS |
| Gbp4 | 1.75E-21 | 3.13E-17 | 0.29 | 0.336 | 0.158 | Macrophages | aPD-L1+aTGF-β vs PBS |
| Atp5e | 2.23E-14 | 3.98E-10 | 0.29 | 0.695 | 0.513 | B cells | aPD-L1+aTGF-β vs PBS |
| Rpl23 | 3.11E-10 | 5.54E-06 | 0.29 | 1 | 1 | NKT | aPD-L1+aTGF-β vs PBS |
| Zfand5 | 7.52E-09 | 1.34E-04 | 0.29 | 0.753 | 0.573 | Monocytes | aPD-L1+aTGF-β vs PBS |
| Nampt | 1.95E-08 | 3.48E-04 | 0.29 | 0.627 | 0.462 | Monocytes | aPD-L1+aTGF-β vs PBS |
| Ccl2 | 2.64E-10 | 4.71E-06 | 0.29 | 0.692 | 0.565 | Macrophages | aPD-L1+aTGF-β vs PBS |
| Rpl36a | 1.39E-08 | 2.48E-04 | 0.29 | 0.77 | 0.62 | Monocytes | aPD-L1+aTGF-β vs PBS |
| Nfkb1 | 2.53E-11 | 4.51E-07 | 0.29 | 0.469 | 0.298 | T cells | aPD-L1+aTGF-β vs PBS |
| Ubc | 2.54E-51 | 4.53E-47 | 0.29 | 0.869 | 0.718 | Tumor_cells | aPD-L1+aTGF-β vs PBS |
| Rps15a | 1.44E-08 | 2.57E-04 | 0.29 | 0.956 | 0.934 | Monocytes | aPD-L1+aTGF-β vs PBS |
| Rpl36 | 7.34E-14 | 1.31E-09 | 0.29 | 0.983 | 0.927 | Monocytes | aPD-L1+aTGF-β vs PBS |
| Mt1 | 8.76E-51 | 1.56E-46 | 0.28 | 0.942 | 0.836 | Tumor_cells | aPD-L1+aTGF-β vs PBS |
| Rps29 | 7.93E-10 | 1.41E-05 | 0.28 | 1 | 1 | ILC | aPD-L1+aTGF-β vs PBS |
| Ly6a | 4.83E-08 | 8.62E-04 | 0.28 | 0.666 | 0.639 | Tumor_cells | aPD-L1+aTGF-β vs PBS |
| Ier3 | 1.06E-41 | 1.89E-37 | 0.28 | 0.934 | 0.799 | Tumor_cells | aPD-L1+aTGF-β vs PBS |
| Peli1 | 1.52E-22 | 2.70E-18 | 0.28 | 0.454 | 0.25 | Macrophages | aPD-L1+aTGF-β vs PBS |
| Egr1 | 6.26E-46 | 1.12E-41 | 0.28 | 0.805 | 0.649 | Tumor_cells | aPD-L1+aTGF-β vs PBS |
| Uba52 | 5.85E-15 | 1.04E-10 | 0.28 | 0.516 | 0.279 | B cells | aPD-L1+aTGF-β vs PBS |
| Rpl34 | 9.67E-42 | 1.72E-37 | 0.28 | 1 | 1 | T cells | aPD-L1+aTGF-β vs PBS |
| Rpl36a | 1.41E-06 | 2.51E-02 | 0.28 | 0.917 | 0.881 | NKT | aPD-L1+aTGF-β vs PBS |
| Nfkbia | 1.59E-16 | 2.84E-12 | 0.28 | 0.836 | 0.711 | Macrophages | aPD-L1+aTGF-β vs PBS |
| Fam26f | 1.16E-26 | 2.06E-22 | 0.27 | 0.3 | 0.11 | Macrophages | aPD-L1+aTGF-β vs PBS |
| Rab20 | 3.82E-08 | 6.80E-04 | 0.27 | 0.557 | 0.38 | Monocytes | aPD-L1+aTGF-β vs PBS |
| Rpl38 | 2.04E-09 | 3.64E-05 | 0.27 | 0.994 | 0.981 | NKT | aPD-L1+aTGF-β vs PBS |
| Tmsb10 | 4.91E-18 | 8.76E-14 | 0.27 | 0.985 | 0.972 | B cells | aPD-L1+aTGF-β vs PBS |
| Rpl30 | 5.80E-08 | 1.03E-03 | 0.27 | 1 | 0.981 | ILC | aPD-L1+aTGF-β vs PBS |
| Stat1 | 2.51E-20 | 4.48E-16 | 0.27 | 0.554 | 0.353 | Macrophages | aPD-L1+aTGF-β vs PBS |
| Ccrl2 | 4.41E-16 | 7.86E-12 | 0.27 | 0.898 | 0.794 | Neutrophils | aPD-L1+aTGF-β vs PBS |
| Samhd1 | 7.31E-12 | 1.30E-07 | 0.27 | 0.499 | 0.361 | Macrophages | aPD-L1+aTGF-β vs PBS |
| Ccnl1 | 4.48E-56 | 7.99E-52 | 0.27 | 0.507 | 0.285 | Tumor_cells | aPD-L1+aTGF-β vs PBS |
| Il1rn | 9.06E-12 | 1.62E-07 | 0.27 | 0.75 | 0.652 | Neutrophils | aPD-L1+aTGF-β vs PBS |
| Rpl35 | 5.71E-40 | 1.02E-35 | 0.27 | 0.979 | 0.968 | Macrophages | aPD-L1+aTGF-β vs PBS |
| Rpl27a | 1.97E-10 | 3.51E-06 | 0.27 | 1 | 1 | NKT | aPD-L1+aTGF-β vs PBS |
| Cd274 | 2.11E-16 | 3.77E-12 | 0.27 | 0.358 | 0.199 | Macrophages | aPD-L1+aTGF-β vs PBS |
| Sec61g | 1.92E-13 | 3.43E-09 | 0.27 | 0.49 | 0.276 | B cells | aPD-L1+aTGF-β vs PBS |
| Cdc42ep2 | 3.58E-18 | 6.38E-14 | 0.27 | 0.27 | 0.121 | Neutrophils | aPD-L1+aTGF-β vs PBS |
| Rps29 | 9.22E-08 | 1.64E-03 | 0.27 | 0.997 | 0.994 | NKT | aPD-L1+aTGF-β vs PBS |
| Pnrc1 | 4.68E-46 | 8.35E-42 | 0.27 | 0.677 | 0.502 | Tumor_cells | aPD-L1+aTGF-β vs PBS |
| Rpl22l1 | 6.17E-10 | 1.10E-05 | 0.26 | 0.707 | 0.513 | Monocytes | aPD-L1+aTGF-β vs PBS |
| Slfn2 | 2.09E-08 | 3.72E-04 | 0.26 | 0.872 | 0.775 | Monocytes | aPD-L1+aTGF-β vs PBS |
| Isg15 | 6.09E-08 | 1.09E-03 | 0.26 | 0.515 | 0.417 | Macrophages | aPD-L1+aTGF-β vs PBS |
| Rpl36 | 5.46E-07 | 9.74E-03 | 0.26 | 0.979 | 0.98 | Fibroblasts | aPD-L1+aTGF-β vs PBS |
| Rps24 | 1.14E-08 | 2.04E-04 | 0.26 | 1 | 1 | NKT | aPD-L1+aTGF-β vs PBS |
| Slamf8 | 1.11E-25 | 1.98E-21 | 0.26 | 0.363 | 0.162 | Macrophages | aPD-L1+aTGF-β vs PBS |
| Sqstm1 | 1.26E-13 | 2.25E-09 | 0.26 | 0.449 | 0.299 | Macrophages | aPD-L1+aTGF-β vs PBS |
| Ifitm3 | 1.75E-14 | 3.12E-10 | 0.26 | 0.962 | 0.947 | Macrophages | aPD-L1+aTGF-β vs PBS |
| Rpl22 | 2.04E-07 | 3.64E-03 | 0.26 | 0.982 | 0.981 | ILC | aPD-L1+aTGF-β vs PBS |
| Rpl35a | 1.87E-37 | 3.34E-33 | 0.26 | 0.988 | 0.981 | Macrophages | aPD-L1+aTGF-β vs PBS |
| Atp5e | 8.82E-34 | 1.57E-29 | 0.26 | 0.94 | 0.849 | Macrophages | aPD-L1+aTGF-β vs PBS |
| Gadd45b | 1.48E-06 | 2.65E-02 | 0.26 | 0.396 | 0.303 | Macrophages | aPD-L1+aTGF-β vs PBS |

| | | | | | | | |
|---|---|---|---|---|---|---|---|
| Ninj1 | 1.88E-06 | 3.35E-02 | 0.26 | 0.666 | 0.522 | Monocytes | aPD-L1+aTGF-β vs PBS |
| Rps19 | 6.65E-08 | 1.19E-03 | 0.26 | 0.997 | 1 | NKT | aPD-L1+aTGF-β vs PBS |
| Rpl41 | 7.46E-07 | 1.33E-02 | 0.26 | 1 | 1 | ILC | aPD-L1+aTGF-β vs PBS |
| Rilpl2 | 9.68E-10 | 1.72E-05 | 0.25 | 0.446 | 0.234 | Monocytes | aPD-L1+aTGF-β vs PBS |
| Csrnp1 | 4.54E-08 | 8.10E-04 | 0.25 | 0.588 | 0.408 | Monocytes | aPD-L1+aTGF-β vs PBS |
| Gbp2 | 1.04E-07 | 1.85E-03 | 0.25 | 0.574 | 0.37 | Monocytes | aPD-L1+aTGF-β vs PBS |
| Nfkbia | 1.59E-09 | 2.83E-05 | 0.25 | 0.902 | 0.858 | T cells | aPD-L1+aTGF-β vs PBS |
| Fau | 1.94E-12 | 3.46E-08 | 0.25 | 1 | 1 | NKT | aPD-L1+aTGF-β vs PBS |
| Rpl21 | 3.65E-07 | 6.51E-03 | 0.25 | 0.994 | 0.988 | NKT | aPD-L1+aTGF-β vs PBS |
| Ccl7 | 1.38E-06 | 2.46E-02 | 0.25 | 0.512 | 0.399 | Macrophages | aPD-L1+aTGF-β vs PBS |
| P3h4 | 3.43E-07 | 6.12E-03 | -0.25 | 0.176 | 0.423 | Fibroblasts | aPD-L1+aTGF-β vs PBS |
| Gadd45g | 7.98E-14 | 1.42E-09 | -0.25 | 0.32 | 0.41 | Tumor_cells | aPD-L1+aTGF-β vs PBS |
| Myl9 | 3.76E-23 | 6.70E-19 | -0.25 | 0.235 | 0.354 | Tumor_cells | aPD-L1+aTGF-β vs PBS |
| Btg2 | 2.05E-09 | 3.66E-05 | -0.25 | 0.89 | 0.913 | T cells | aPD-L1+aTGF-β vs PBS |
| Fosb | 2.66E-06 | 4.74E-02 | -0.25 | 0.827 | 0.858 | Neutrophils | aPD-L1+aTGF-β vs PBS |
| Eno1 | 2.10E-52 | 3.75E-48 | -0.26 | 0.7 | 0.84 | Tumor_cells | aPD-L1+aTGF-β vs PBS |
| Gm42418 | 1.12E-59 | 1.99E-55 | -0.26 | 0.998 | 1 | Tumor_cells | aPD-L1+aTGF-β vs PBS |
| Gsn | 7.00E-07 | 1.25E-02 | -0.26 | 0.462 | 0.614 | Monocytes | aPD-L1+aTGF-β vs PBS |
| Tpi1 | 2.00E-47 | 3.57E-43 | -0.26 | 0.918 | 0.961 | Tumor_cells | aPD-L1+aTGF-β vs PBS |
| Il7r | 8.33E-09 | 1.48E-04 | -0.26 | 0.516 | 0.647 | T cells | aPD-L1+aTGF-β vs PBS |
| P4hb | 1.88E-09 | 3.35E-05 | -0.27 | 0.406 | 0.498 | Neutrophils | aPD-L1+aTGF-β vs PBS |
| Fn1 | 7.62E-08 | 1.36E-03 | -0.27 | 0.348 | 0.488 | Macrophages | aPD-L1+aTGF-β vs PBS |
| Ccr2 | 7.62E-07 | 1.36E-02 | -0.27 | 0.663 | 0.804 | Monocytes | aPD-L1+aTGF-β vs PBS |
| Dusp1 | 3.28E-13 | 5.85E-09 | -0.27 | 0.642 | 0.806 | B cells | aPD-L1+aTGF-β vs PBS |
| Grn | 9.16E-08 | 1.63E-03 | -0.27 | 0.685 | 0.804 | Monocytes | aPD-L1+aTGF-β vs PBS |
| Atp6v1b1 | 2.43E-06 | 4.34E-02 | -0.27 | 0.107 | 0.324 | Epithelial cells | aPD-L1+aTGF-β vs PBS |
| Iglc2 | 1.55E-10 | 2.76E-06 | -0.27 | 0.768 | 0.858 | B cells | aPD-L1+aTGF-β vs PBS |
| Rpl5 | 2.12E-16 | 3.79E-12 | -0.27 | 0.959 | 0.991 | B cells | aPD-L1+aTGF-β vs PBS |
| Actb | 1.21E-58 | 2.16E-54 | -0.27 | 1 | 1 | Tumor_cells | aPD-L1+aTGF-β vs PBS |
| Cdkn1a | 1.17E-12 | 2.08E-08 | -0.27 | 0.778 | 0.817 | Neutrophils | aPD-L1+aTGF-β vs PBS |
| Wls | 1.09E-07 | 1.94E-03 | -0.27 | 0.261 | 0.537 | Fibroblasts | aPD-L1+aTGF-β vs PBS |
| Gatm | 9.22E-10 | 1.64E-05 | -0.27 | 0.705 | 0.792 | Macrophages | aPD-L1+aTGF-β vs PBS |
| Cxcr4 | 4.78E-07 | 8.53E-03 | -0.27 | 0.562 | 0.624 | B cells | aPD-L1+aTGF-β vs PBS |
| Tshz2 | 3.55E-07 | 6.33E-03 | -0.27 | 0.33 | 0.604 | Fibroblasts | aPD-L1+aTGF-β vs PBS |
| Dab2 | 3.65E-21 | 6.50E-17 | -0.28 | 0.714 | 0.834 | Macrophages | aPD-L1+aTGF-β vs PBS |
| Cyr61 | 3.42E-11 | 6.10E-07 | -0.28 | 0.492 | 0.543 | Tumor_cells | aPD-L1+aTGF-β vs PBS |
| Ltb | 1.09E-09 | 1.94E-05 | -0.28 | 0.603 | 0.704 | B cells | aPD-L1+aTGF-β vs PBS |
| Ccr2 | 2.67E-10 | 4.76E-06 | -0.28 | 0.376 | 0.491 | Macrophages | aPD-L1+aTGF-β vs PBS |
| Mmp23 | 1.26E-06 | 2.24E-02 | -0.28 | 0.426 | 0.678 | Fibroblasts | aPD-L1+aTGF-β vs PBS |
| Pid1 | 5.97E-15 | 1.07E-10 | -0.28 | 0.678 | 0.772 | Macrophages | aPD-L1+aTGF-β vs PBS |
| Pgk1 | 2.04E-46 | 3.63E-42 | -0.28 | 0.91 | 0.959 | Tumor_cells | aPD-L1+aTGF-β vs PBS |
| Lgmn | 1.77E-06 | 3.16E-02 | -0.28 | 0.603 | 0.725 | Monocytes | aPD-L1+aTGF-β vs PBS |
| Ldha | 2.35E-13 | 4.20E-09 | -0.29 | 0.912 | 0.937 | Neutrophils | aPD-L1+aTGF-β vs PBS |
| Gng11 | 6.68E-09 | 1.19E-04 | -0.29 | 0.304 | 0.362 | Tumor_cells | aPD-L1+aTGF-β vs PBS |
| Ppp1r3b | 2.52E-19 | 4.48E-15 | -0.30 | 0.303 | 0.472 | Neutrophils | aPD-L1+aTGF-β vs PBS |
| Pmepa1 | 3.79E-17 | 6.75E-13 | -0.30 | 0.529 | 0.701 | Macrophages | aPD-L1+aTGF-β vs PBS |
| Ier2 | 9.34E-08 | 1.66E-03 | -0.30 | 0.732 | 0.789 | T cells | aPD-L1+aTGF-β vs PBS |
| Zfp36 | 1.72E-11 | 3.07E-07 | -0.30 | 0.708 | 0.823 | B cells | aPD-L1+aTGF-β vs PBS |
| Junb | 9.71E-12 | 1.73E-07 | -0.30 | 0.979 | 0.986 | B cells | aPD-L1+aTGF-β vs PBS |
| Rpl5 | 2.73E-24 | 4.88E-20 | -0.30 | 0.987 | 0.998 | T cells | aPD-L1+aTGF-β vs PBS |
| Smc6 | 4.03E-13 | 7.18E-09 | -0.30 | 0.724 | 0.802 | T cells | aPD-L1+aTGF-β vs PBS |
| Gja1 | 1.78E-06 | 3.18E-02 | -0.31 | 0.181 | 0.416 | Fibroblasts | aPD-L1+aTGF-β vs PBS |
| Grb10 | 2.83E-07 | 5.05E-03 | -0.31 | 0.213 | 0.477 | Fibroblasts | aPD-L1+aTGF-β vs PBS |
| Ier2 | 6.49E-13 | 1.16E-08 | -0.31 | 0.713 | 0.855 | B cells | aPD-L1+aTGF-β vs PBS |
| Ier5 | 5.61E-18 | 1.00E-13 | -0.31 | 0.79 | 0.906 | B cells | aPD-L1+aTGF-β vs PBS |

| | | | | | | | |
|---|---|---|---|---|---|---|---|
| Fdps | 1.29E-39 | 2.30E-35 | -0.32 | 0.411 | 0.556 | Tumor_cells | aPD-L1+aTGF-β vs PBS |
| Hspa1b | 1.33E-10 | 2.38E-06 | -0.32 | 0.408 | 0.593 | B cells | aPD-L1+aTGF-β vs PBS |
| Celsr2 | 1.16E-07 | 2.06E-03 | -0.33 | 0.141 | 0.397 | Epithelial cells | aPD-L1+aTGF-β vs PBS |
| Tgfbi | 4.59E-17 | 8.18E-13 | -0.33 | 0.815 | 0.874 | Macrophages | aPD-L1+aTGF-β vs PBS |
| Jun | 1.71E-12 | 3.05E-08 | -0.34 | 0.871 | 0.941 | T cells | aPD-L1+aTGF-β vs PBS |
| Smad7 | 5.80E-13 | 1.03E-08 | -0.34 | 0.674 | 0.767 | T cells | aPD-L1+aTGF-β vs PBS |
| Ccdc80 | 1.54E-06 | 2.74E-02 | -0.34 | 0.628 | 0.819 | Fibroblasts | aPD-L1+aTGF-β vs PBS |
| Jun | 9.87E-12 | 1.76E-07 | -0.34 | 0.974 | 0.988 | Neutrophils | aPD-L1+aTGF-β vs PBS |
| Prnp | 2.64E-07 | 4.70E-03 | -0.34 | 0.441 | 0.691 | Fibroblasts | aPD-L1+aTGF-β vs PBS |
| Dusp2 | 2.28E-12 | 4.06E-08 | -0.34 | 0.606 | 0.715 | B cells | aPD-L1+aTGF-β vs PBS |
| Egr1 | 5.96E-08 | 1.06E-03 | -0.34 | 0.424 | 0.544 | B cells | aPD-L1+aTGF-β vs PBS |
| Actg1 | 4.91E-74 | 8.76E-70 | -0.34 | 0.981 | 0.992 | Tumor_cells | aPD-L1+aTGF-β vs PBS |
| mt-Co2 | 5.38E-09 | 9.60E-05 | -0.35 | 0.995 | 1 | Fibroblasts | aPD-L1+aTGF-β vs PBS |
| Ms4a7 | 3.94E-14 | 7.03E-10 | -0.35 | 0.527 | 0.681 | Macrophages | aPD-L1+aTGF-β vs PBS |
| Cxcl16 | 2.06E-08 | 3.68E-04 | -0.36 | 0.383 | 0.554 | Monocytes | aPD-L1+aTGF-β vs PBS |
| Ltbp4 | 4.09E-07 | 7.28E-03 | -0.36 | 0.117 | 0.336 | Fibroblasts | aPD-L1+aTGF-β vs PBS |
| Nfix | 2.02E-07 | 3.61E-03 | -0.37 | 0.33 | 0.647 | Epithelial cells | aPD-L1+aTGF-β vs PBS |
| Nfkbia | 1.34E-12 | 2.38E-08 | -0.37 | 0.782 | 0.886 | B cells | aPD-L1+aTGF-β vs PBS |
| mt-Co3 | 2.81E-09 | 5.00E-05 | -0.37 | 0.995 | 1 | Fibroblasts | aPD-L1+aTGF-β vs PBS |
| mt-Atp6 | 1.16E-09 | 2.07E-05 | -0.37 | 0.995 | 1 | Fibroblasts | aPD-L1+aTGF-β vs PBS |
| Bst2 | 6.88E-11 | 1.23E-06 | -0.37 | 0.634 | 0.785 | Monocytes | aPD-L1+aTGF-β vs PBS |
| Ccl6 | 4.17E-09 | 7.43E-05 | -0.38 | 0.502 | 0.586 | Neutrophils | aPD-L1+aTGF-β vs PBS |
| Tmsb4x | 8.65E-16 | 1.54E-11 | -0.38 | 0.103 | 0.184 | Tumor_cells | aPD-L1+aTGF-β vs PBS |
| Hist1h2bc | 2.58E-09 | 4.60E-05 | -0.38 | 0.174 | 0.305 | B cells | aPD-L1+aTGF-β vs PBS |
| Fos | 2.76E-16 | 4.91E-12 | -0.38 | 0.989 | 0.994 | Neutrophils | aPD-L1+aTGF-β vs PBS |
| Smad7 | 9.22E-07 | 1.64E-02 | -0.38 | 0.583 | 0.756 | NKT | aPD-L1+aTGF-β vs PBS |
| Col1a2 | 1.41E-07 | 2.52E-03 | -0.39 | 0.989 | 0.98 | Fibroblasts | aPD-L1+aTGF-β vs PBS |
| Lgals1 | 1.27E-10 | 2.27E-06 | -0.40 | 0.84 | 0.911 | Monocytes | aPD-L1+aTGF-β vs PBS |
| Cthrc1 | 3.20E-53 | 5.70E-49 | -0.40 | 0.447 | 0.635 | Tumor_cells | aPD-L1+aTGF-β vs PBS |
| Klf2 | 1.35E-18 | 2.40E-14 | -0.42 | 0.871 | 0.959 | T cells | aPD-L1+aTGF-β vs PBS |
| Klf2 | 5.20E-11 | 9.28E-07 | -0.43 | 0.967 | 0.976 | Neutrophils | aPD-L1+aTGF-β vs PBS |
| Fbn1 | 1.53E-06 | 2.73E-02 | -0.43 | 0.782 | 0.879 | Fibroblasts | aPD-L1+aTGF-β vs PBS |
| Acta2 | 2.80E-34 | 4.99E-30 | -0.43 | 0.592 | 0.722 | Tumor_cells | aPD-L1+aTGF-β vs PBS |
| Tgfbi | 1.03E-10 | 1.84E-06 | -0.43 | 0.768 | 0.877 | Monocytes | aPD-L1+aTGF-β vs PBS |
| Arg1 | 2.09E-09 | 3.72E-05 | -0.44 | 0.291 | 0.42 | Macrophages | aPD-L1+aTGF-β vs PBS |
| Fn1 | 2.43E-07 | 4.34E-03 | -0.45 | 0.237 | 0.408 | Monocytes | aPD-L1+aTGF-β vs PBS |
| Gm42418 | 1.30E-08 | 2.32E-04 | -0.45 | 1 | 1 | Monocytes | aPD-L1+aTGF-β vs PBS |
| Fxyd3 | 2.30E-07 | 4.10E-03 | -0.46 | 0.744 | 0.882 | Epithelial cells | aPD-L1+aTGF-β vs PBS |
| Fos | 9.83E-26 | 1.75E-21 | -0.47 | 0.917 | 0.969 | B cells | aPD-L1+aTGF-β vs PBS |
| AY036118 | 3.11E-08 | 5.55E-04 | -0.47 | 0.766 | 0.899 | Fibroblasts | aPD-L1+aTGF-β vs PBS |
| Klf2 | 1.92E-22 | 3.42E-18 | -0.50 | 0.885 | 0.966 | B cells | aPD-L1+aTGF-β vs PBS |
| Gm42418 | 6.63E-23 | 1.18E-18 | -0.50 | 1 | 1 | Macrophages | aPD-L1+aTGF-β vs PBS |
| Jun | 1.22E-29 | 2.17E-25 | -0.52 | 0.922 | 0.994 | B cells | aPD-L1+aTGF-β vs PBS |
| Hist1h1c | 1.18E-20 | 2.10E-16 | -0.53 | 0.385 | 0.601 | B cells | aPD-L1+aTGF-β vs PBS |
| Col1a1 | 4.93E-07 | 8.79E-03 | -0.53 | 0.968 | 0.946 | Fibroblasts | aPD-L1+aTGF-β vs PBS |
| Tcim | 2.95E-07 | 5.26E-03 | -0.54 | 0.231 | 0.515 | Epithelial cells | aPD-L1+aTGF-β vs PBS |
| Pid1 | 5.77E-17 | 1.03E-12 | -0.54 | 0.685 | 0.835 | Monocytes | aPD-L1+aTGF-β vs PBS |
| Ms4a7 | 1.08E-08 | 1.93E-04 | -0.55 | 0.111 | 0.269 | Monocytes | aPD-L1+aTGF-β vs PBS |
| Csrp2 | 9.30E-07 | 1.66E-02 | -0.55 | 0.628 | 0.799 | Fibroblasts | aPD-L1+aTGF-β vs PBS |
| Cthrc1 | 1.03E-07 | 1.84E-03 | -0.57 | 0.415 | 0.658 | Fibroblasts | aPD-L1+aTGF-β vs PBS |
| AY036118 | 1.14E-07 | 2.04E-03 | -0.60 | 0.648 | 0.838 | Epithelial cells | aPD-L1+aTGF-β vs PBS |
| Gm42418 | 3.67E-08 | 6.54E-04 | -0.61 | 1 | 1 | Fibroblasts | aPD-L1+aTGF-β vs PBS |
| Pf4 | 8.92E-11 | 1.59E-06 | -0.63 | 0.348 | 0.483 | Macrophages | aPD-L1+aTGF-β vs PBS |
| AY036118 | 7.61E-15 | 1.36E-10 | -0.65 | 0.835 | 0.937 | Monocytes | aPD-L1+aTGF-β vs PBS |
| AY036118 | 7.58E-174 | 1.35E-169 | -0.70 | 0.588 | 0.876 | Tumor_cells | aPD-L1+aTGF-β vs PBS |

| | | | | | | | |
|---|---|---|---|---|---|---|---|
| AY036118 | 3.01E-41 | 5.37E-37 | -0.71 | 0.652 | 0.836 | Macrophages | aPD-L1+aTGF-β vs PBS |
| Gm42418 | 9.70E-09 | 1.73E-04 | -0.74 | 0.993 | 1 | Epithelial cells | aPD-L1+aTGF-β vs PBS |
| Igfbp5 | 7.57E-08 | 1.35E-03 | -0.76 | 0.355 | 0.691 | Epithelial cells | aPD-L1+aTGF-β vs PBS |
| Spp1 | 1.04E-08 | 1.86E-04 | -0.82 | 0.324 | 0.494 | Monocytes | aPD-L1+aTGF-β vs PBS |
| Apoe | 5.16E-20 | 9.19E-16 | -1.07 | 0.542 | 0.769 | Monocytes | aPD-L1+aTGF-β vs PBS |
| Cxcl10 | 8.62E-39 | 1.54E-34 | 1.12 | 0.282 | 0.07 | Neutrophils | aTGF-β vs PBS |
| Rsad2 | 1.07E-38 | 1.90E-34 | 0.75 | 0.716 | 0.502 | Neutrophils | aTGF-β vs PBS |
| Ifit1 | 6.22E-35 | 1.11E-30 | 0.75 | 0.525 | 0.301 | Neutrophils | aTGF-β vs PBS |
| Ifit2 | 4.10E-33 | 7.31E-29 | 0.65 | 0.349 | 0.139 | Neutrophils | aTGF-β vs PBS |
| Gbp2b | 3.78E-37 | 6.74E-33 | 0.52 | 0.562 | 0.312 | Neutrophils | aTGF-β vs PBS |
| Parp14 | 4.05E-36 | 7.22E-32 | 0.49 | 0.563 | 0.324 | Neutrophils | aTGF-β vs PBS |
| Dnajb1 | 1.36E-54 | 2.43E-50 | 0.48 | 0.803 | 0.647 | Tumor_cells | aTGF-β vs PBS |
| Gbp2 | 6.19E-41 | 1.10E-36 | 0.48 | 0.533 | 0.266 | Neutrophils | aTGF-β vs PBS |
| Plac8 | 5.87E-22 | 1.05E-17 | 0.46 | 0.373 | 0.197 | Neutrophils | aTGF-β vs PBS |
| Ifit3 | 2.17E-25 | 3.87E-21 | 0.46 | 0.326 | 0.143 | Neutrophils | aTGF-β vs PBS |
| Ifitm3 | 2.62E-32 | 4.66E-28 | 0.45 | 0.988 | 0.975 | Neutrophils | aTGF-β vs PBS |
| Gbp5 | 1.02E-34 | 1.82E-30 | 0.45 | 0.336 | 0.118 | Neutrophils | aTGF-β vs PBS |
| Irf1 | 3.77E-23 | 6.72E-19 | 0.44 | 0.563 | 0.375 | Neutrophils | aTGF-β vs PBS |
| Isg15 | 1.06E-32 | 1.89E-28 | 0.44 | 0.965 | 0.916 | Neutrophils | aTGF-β vs PBS |
| Ifi47 | 1.45E-32 | 2.59E-28 | 0.43 | 0.507 | 0.272 | Neutrophils | aTGF-β vs PBS |
| Cmpk2 | 2.69E-28 | 4.79E-24 | 0.43 | 0.303 | 0.117 | Neutrophils | aTGF-β vs PBS |
| Dcn | 1.85E-22 | 3.30E-18 | 0.42 | 0.219 | 0.106 | Tumor_cells | aTGF-β vs PBS |
| Slfn4 | 1.97E-27 | 3.51E-23 | 0.42 | 0.853 | 0.754 | Neutrophils | aTGF-β vs PBS |
| Isg20 | 5.02E-18 | 8.94E-14 | 0.42 | 0.618 | 0.486 | Neutrophils | aTGF-β vs PBS |
| Irgm1 | 2.90E-29 | 5.17E-25 | 0.40 | 0.44 | 0.226 | Neutrophils | aTGF-β vs PBS |
| Irf7 | 4.56E-21 | 8.12E-17 | 0.39 | 0.567 | 0.404 | Neutrophils | aTGF-β vs PBS |
| Hspa1a | 7.30E-20 | 1.30E-15 | 0.39 | 0.613 | 0.445 | Macrophages | aTGF-β vs PBS |
| Hspa1b | 7.83E-65 | 1.40E-60 | 0.39 | 0.818 | 0.616 | Tumor_cells | aTGF-β vs PBS |
| Klf2 | 1.95E-29 | 3.48E-25 | 0.39 | 0.427 | 0.272 | Tumor_cells | aTGF-β vs PBS |
| Atf3 | 4.55E-10 | 8.11E-06 | 0.39 | 0.913 | 0.826 | Monocytes | aTGF-β vs PBS |
| Ccl4 | 7.26E-11 | 1.29E-06 | 0.38 | 0.7 | 0.591 | Neutrophils | aTGF-β vs PBS |
| Hspb1 | 6.61E-49 | 1.18E-44 | 0.38 | 0.924 | 0.87 | Tumor_cells | aTGF-β vs PBS |
| Cd52 | 1.20E-16 | 2.14E-12 | 0.37 | 0.923 | 0.855 | B cells | aTGF-β vs PBS |
| Hsp90ab1 | 1.19E-07 | 2.12E-03 | 0.36 | 1 | 1 | ILC | aTGF-β vs PBS |
| Dusp1 | 6.81E-49 | 1.21E-44 | 0.36 | 0.821 | 0.664 | Tumor_cells | aTGF-β vs PBS |
| Fos | 9.99E-50 | 1.78E-45 | 0.35 | 0.856 | 0.694 | Tumor_cells | aTGF-β vs PBS |
| AW112010 | 1.11E-09 | 1.98E-05 | 0.35 | 0.579 | 0.392 | T cells | aTGF-β vs PBS |
| Slfn5 | 4.66E-20 | 8.30E-16 | 0.35 | 0.474 | 0.298 | Neutrophils | aTGF-β vs PBS |
| Samhd1 | 2.35E-21 | 4.18E-17 | 0.35 | 0.645 | 0.476 | Neutrophils | aTGF-β vs PBS |
| Acod1 | 5.38E-23 | 9.59E-19 | 0.35 | 0.861 | 0.75 | Neutrophils | aTGF-β vs PBS |
| Ccl12 | 1.77E-12 | 3.16E-08 | 0.35 | 0.589 | 0.451 | Macrophages | aTGF-β vs PBS |
| Irf1 | 1.46E-07 | 2.60E-03 | 0.34 | 0.562 | 0.396 | Monocytes | aTGF-β vs PBS |
| Cd274 | 1.10E-19 | 1.96E-15 | 0.33 | 0.647 | 0.475 | Neutrophils | aTGF-β vs PBS |
| Klf2 | 2.86E-07 | 5.10E-03 | 0.33 | 0.739 | 0.579 | Monocytes | aTGF-β vs PBS |
| Ifit3b | 1.91E-13 | 3.41E-09 | 0.33 | 0.308 | 0.179 | Neutrophils | aTGF-β vs PBS |
| Ccrl2 | 3.15E-21 | 5.62E-17 | 0.33 | 0.901 | 0.794 | Neutrophils | aTGF-β vs PBS |
| Oasl1 | 7.64E-17 | 1.36E-12 | 0.32 | 0.537 | 0.386 | Neutrophils | aTGF-β vs PBS |
| Ccl4 | 3.05E-11 | 5.43E-07 | 0.32 | 0.757 | 0.653 | Macrophages | aTGF-β vs PBS |
| H2-D1 | 1.11E-34 | 1.99E-30 | 0.32 | 1 | 0.999 | Neutrophils | aTGF-β vs PBS |
| H2-K1 | 1.03E-27 | 1.84E-23 | 0.32 | 0.997 | 0.989 | Neutrophils | aTGF-β vs PBS |
| Ddx60 | 1.97E-20 | 3.51E-16 | 0.32 | 0.399 | 0.221 | Neutrophils | aTGF-β vs PBS |
| Trafd1 | 1.76E-20 | 3.13E-16 | 0.32 | 0.422 | 0.254 | Neutrophils | aTGF-β vs PBS |
| Fosb | 1.02E-40 | 1.82E-36 | 0.32 | 0.645 | 0.465 | Tumor_cells | aTGF-β vs PBS |
| Klf2 | 7.05E-18 | 1.26E-13 | 0.32 | 0.751 | 0.626 | Macrophages | aTGF-β vs PBS |
| Fcgr4 | 2.71E-21 | 4.83E-17 | 0.31 | 0.392 | 0.215 | Neutrophils | aTGF-β vs PBS |

| | | | | | | | |
|---|---|---|---|---|---|---|---|
| Ifi27l2a | 2.25E-22 | 4.02E-18 | 0.31 | 0.883 | 0.799 | Macrophages | aTGF-β vs PBS |
| Trim30a | 8.87E-20 | 1.58E-15 | 0.31 | 0.586 | 0.424 | Neutrophils | aTGF-β vs PBS |
| Sod2 | 5.27E-13 | 9.40E-09 | 0.31 | 0.311 | 0.185 | Neutrophils | aTGF-β vs PBS |
| Herc6 | 6.59E-19 | 1.17E-14 | 0.31 | 0.378 | 0.218 | Neutrophils | aTGF-β vs PBS |
| Klf6 | 1.00E-07 | 1.79E-03 | 0.31 | 0.925 | 0.873 | Monocytes | aTGF-β vs PBS |
| Tmsb10 | 2.23E-06 | 3.98E-02 | 0.31 | 0.95 | 0.94 | Fibroblasts | aTGF-β vs PBS |
| Slfn1 | 1.87E-18 | 3.33E-14 | 0.30 | 0.593 | 0.426 | Neutrophils | aTGF-β vs PBS |
| Hspa1b | 3.53E-13 | 6.28E-09 | 0.30 | 0.51 | 0.373 | Macrophages | aTGF-β vs PBS |
| Gm47283 | 9.27E-12 | 1.65E-07 | 0.30 | 0.701 | 0.525 | T cells | aTGF-β vs PBS |
| Zbp1 | 2.84E-18 | 5.05E-14 | 0.30 | 0.375 | 0.208 | Neutrophils | aTGF-β vs PBS |
| Sgk1 | 2.14E-40 | 3.81E-36 | 0.30 | 0.683 | 0.501 | Tumor_cells | aTGF-β vs PBS |
| B2m | 1.99E-26 | 3.55E-22 | 0.30 | 0.998 | 0.998 | Neutrophils | aTGF-β vs PBS |
| Marcksl1 | 8.12E-12 | 1.45E-07 | 0.30 | 0.59 | 0.479 | Neutrophils | aTGF-β vs PBS |
| Ikzf2 | 5.20E-09 | 9.28E-05 | 0.29 | 0.151 | 0.039 | T cells | aTGF-β vs PBS |
| Clec2d | 1.32E-18 | 2.35E-14 | 0.29 | 0.353 | 0.19 | Neutrophils | aTGF-β vs PBS |
| Usp18 | 2.37E-19 | 4.23E-15 | 0.29 | 0.318 | 0.157 | Neutrophils | aTGF-β vs PBS |
| 9930111J21Rik2 | 2.66E-18 | 4.74E-14 | 0.29 | 0.353 | 0.196 | Neutrophils | aTGF-β vs PBS |
| Gbp7 | 3.48E-17 | 6.20E-13 | 0.29 | 0.42 | 0.258 | Neutrophils | aTGF-β vs PBS |
| Icam1 | 4.90E-12 | 8.74E-08 | 0.28 | 0.2 | 0.099 | Neutrophils | aTGF-β vs PBS |
| Rnf213 | 1.15E-20 | 2.06E-16 | 0.27 | 0.319 | 0.151 | Neutrophils | aTGF-β vs PBS |
| Upp1 | 4.06E-14 | 7.25E-10 | 0.27 | 0.691 | 0.547 | Neutrophils | aTGF-β vs PBS |
| Igtp | 2.76E-20 | 4.92E-16 | 0.27 | 0.385 | 0.209 | Neutrophils | aTGF-β vs PBS |
| Gbp3 | 2.16E-23 | 3.85E-19 | 0.27 | 0.237 | 0.082 | Neutrophils | aTGF-β vs PBS |
| Nampt | 1.26E-17 | 2.24E-13 | 0.27 | 0.465 | 0.299 | Neutrophils | aTGF-β vs PBS |
| Thbs1 | 1.55E-10 | 2.76E-06 | 0.27 | 0.138 | 0.056 | Neutrophils | aTGF-β vs PBS |
| Samsn1 | 6.82E-08 | 1.21E-03 | 0.27 | 0.407 | 0.262 | B cells | aTGF-β vs PBS |
| Sppl2a | 1.16E-18 | 2.07E-14 | 0.27 | 0.313 | 0.157 | Neutrophils | aTGF-β vs PBS |
| Slfn2 | 1.88E-13 | 3.35E-09 | 0.27 | 0.684 | 0.587 | Neutrophils | aTGF-β vs PBS |
| Cr2 | 1.90E-11 | 3.38E-07 | 0.26 | 0.473 | 0.265 | B cells | aTGF-β vs PBS |
| Cd52 | 5.34E-09 | 9.51E-05 | 0.26 | 0.934 | 0.889 | T cells | aTGF-β vs PBS |
| Izumo1r | 4.51E-08 | 8.05E-04 | 0.25 | 0.217 | 0.092 | T cells | aTGF-β vs PBS |
| Samd9l | 8.01E-19 | 1.43E-14 | 0.25 | 0.425 | 0.251 | Neutrophils | aTGF-β vs PBS |
| Slc38a2 | 6.82E-07 | 1.22E-02 | -0.25 | 0.538 | 0.651 | T cells | aTGF-β vs PBS |
| Cstb | 3.97E-08 | 7.07E-04 | -0.26 | 0.686 | 0.744 | Neutrophils | aTGF-β vs PBS |
| Btg2 | 2.55E-11 | 4.54E-07 | -0.26 | 0.838 | 0.926 | B cells | aTGF-β vs PBS |
| Ccr7 | 1.05E-09 | 1.87E-05 | -0.26 | 0.58 | 0.735 | B cells | aTGF-β vs PBS |
| Ctsc | 8.35E-07 | 1.49E-02 | -0.27 | 0.854 | 0.934 | Monocytes | aTGF-β vs PBS |
| Hsp90aa1 | 1.75E-09 | 3.11E-05 | -0.27 | 0.94 | 0.969 | Neutrophils | aTGF-β vs PBS |
| Tsc22d3 | 1.06E-07 | 1.90E-03 | -0.27 | 0.673 | 0.761 | B cells | aTGF-β vs PBS |
| Pmepa1 | 3.86E-09 | 6.89E-05 | -0.27 | 0.059 | 0.199 | Monocytes | aTGF-β vs PBS |
| Pim1 | 1.59E-07 | 2.84E-03 | -0.27 | 0.33 | 0.462 | B cells | aTGF-β vs PBS |
| Zfp36 | 1.46E-09 | 2.60E-05 | -0.28 | 0.708 | 0.823 | B cells | aTGF-β vs PBS |
| Bcl2a1b | 1.16E-07 | 2.08E-03 | -0.28 | 0.247 | 0.421 | Monocytes | aTGF-β vs PBS |
| Pid1 | 7.60E-07 | 1.35E-02 | -0.29 | 0.696 | 0.835 | Monocytes | aTGF-β vs PBS |
| Cxcr4 | 6.53E-09 | 1.16E-04 | -0.29 | 0.497 | 0.624 | B cells | aTGF-β vs PBS |
| Gng11 | 2.86E-07 | 5.10E-03 | -0.29 | 0.312 | 0.362 | Tumor_cells | aTGF-β vs PBS |
| Dennd4a | 9.43E-11 | 1.68E-06 | -0.29 | 0.528 | 0.684 | B cells | aTGF-β vs PBS |
| Hsp90aa1 | 1.95E-11 | 3.48E-07 | -0.30 | 0.979 | 0.989 | B cells | aTGF-β vs PBS |
| Btg1 | 8.77E-13 | 1.56E-08 | -0.30 | 0.928 | 0.98 | T cells | aTGF-β vs PBS |
| Dusp2 | 3.63E-09 | 6.46E-05 | -0.30 | 0.612 | 0.715 | B cells | aTGF-β vs PBS |
| Jun | 7.05E-09 | 1.26E-04 | -0.30 | 0.979 | 0.988 | Neutrophils | aTGF-β vs PBS |
| Tshz2 | 6.48E-08 | 1.15E-03 | -0.31 | 0.342 | 0.604 | Fibroblasts | aTGF-β vs PBS |
| Klf2 | 1.92E-09 | 3.42E-05 | -0.31 | 0.931 | 0.966 | B cells | aTGF-β vs PBS |
| Il7r | 2.90E-09 | 5.17E-05 | -0.31 | 0.501 | 0.647 | T cells | aTGF-β vs PBS |
| Ier2 | 1.46E-12 | 2.61E-08 | -0.31 | 0.677 | 0.855 | B cells | aTGF-β vs PBS |

| | | | | | | | |
|---|---|---|---|---|---|---|---|
| Srgn | 4.23E-11 | 7.54E-07 | -0.31 | 0.885 | 0.905 | Macrophages | aTGF-β vs PBS |
| Vps37b | 9.10E-07 | 1.62E-02 | -0.31 | 0.526 | 0.643 | T cells | aTGF-β vs PBS |
| Socs3 | 7.66E-11 | 1.37E-06 | -0.31 | 0.693 | 0.754 | Neutrophils | aTGF-β vs PBS |
| Junb | 1.52E-13 | 2.72E-09 | -0.32 | 0.964 | 0.986 | B cells | aTGF-β vs PBS |
| Cxcl16 | 7.75E-08 | 1.38E-03 | -0.32 | 0.388 | 0.554 | Monocytes | aTGF-β vs PBS |
| Acta2 | 5.73E-21 | 1.02E-16 | -0.32 | 0.624 | 0.722 | Tumor_cells | aTGF-β vs PBS |
| Hspa8 | 2.42E-17 | 4.32E-13 | -0.32 | 0.992 | 1 | B cells | aTGF-β vs PBS |
| Btg2 | 7.18E-13 | 1.28E-08 | -0.32 | 0.847 | 0.913 | T cells | aTGF-β vs PBS |
| Ier2 | 1.70E-08 | 3.03E-04 | -0.33 | 0.701 | 0.789 | T cells | aTGF-β vs PBS |
| Hist1h1c | 5.95E-13 | 1.06E-08 | -0.35 | 0.404 | 0.601 | B cells | aTGF-β vs PBS |
| Cd83 | 5.81E-14 | 1.03E-09 | -0.36 | 0.834 | 0.872 | B cells | aTGF-β vs PBS |
| Klf2 | 2.47E-13 | 4.40E-09 | -0.36 | 0.874 | 0.959 | T cells | aTGF-β vs PBS |
| Jund | 1.50E-24 | 2.67E-20 | -0.37 | 0.994 | 1 | B cells | aTGF-β vs PBS |
| Cthrc1 | 2.55E-41 | 4.54E-37 | -0.38 | 0.482 | 0.635 | Tumor_cells | aTGF-β vs PBS |
| Ccl6 | 3.53E-07 | 6.29E-03 | -0.39 | 0.539 | 0.586 | Neutrophils | aTGF-β vs PBS |
| Smad7 | 7.69E-07 | 1.37E-02 | -0.40 | 0.527 | 0.756 | NKT | aTGF-β vs PBS |
| Hist1h2bc | 1.50E-10 | 2.68E-06 | -0.40 | 0.157 | 0.305 | B cells | aTGF-β vs PBS |
| Nr4a1 | 1.24E-16 | 2.21E-12 | -0.40 | 0.451 | 0.672 | B cells | aTGF-β vs PBS |
| Fn1 | 2.18E-06 | 3.89E-02 | -0.42 | 0.249 | 0.408 | Monocytes | aTGF-β vs PBS |
| Nfkbia | 8.45E-14 | 1.51E-09 | -0.43 | 0.82 | 0.886 | B cells | aTGF-β vs PBS |
| Smad7 | 2.50E-16 | 4.46E-12 | -0.44 | 0.612 | 0.767 | T cells | aTGF-β vs PBS |
| Gm42418 | 9.98E-105 | 1.78E-100 | -0.44 | 0.996 | 1 | Tumor_cells | aTGF-β vs PBS |
| Ms4a7 | 1.72E-06 | 3.06E-02 | -0.46 | 0.139 | 0.269 | Monocytes | aTGF-β vs PBS |
| AY036118 | 1.12E-26 | 2.00E-22 | -0.48 | 0.73 | 0.836 | Macrophages | aTGF-β vs PBS |
| AY036118 | 5.50E-96 | 9.80E-92 | -0.50 | 0.701 | 0.876 | Tumor_cells | aTGF-β vs PBS |
| Gm42418 | 5.20E-07 | 9.26E-03 | -0.61 | 1 | 1 | Fibroblasts | aTGF-β vs PBS |
| Arg1 | 4.46E-25 | 7.95E-21 | -0.74 | 0.224 | 0.42 | Macrophages | aTGF-β vs PBS |
| Spp1 | 6.24E-12 | 1.11E-07 | 0.95 | 0.675 | 0.5 | Macrophages | aPD-L1+aTGF-β vs aPD-L1 |
| AW112010 | 3.13E-08 | 5.57E-04 | 0.65 | 0.787 | 0.604 | NKT | aPD-L1+aTGF-β vs aPD-L1 |
| Tmsb10 | 6.67E-14 | 1.19E-09 | 0.63 | 0.952 | 0.724 | Fibroblasts | aPD-L1+aTGF-β vs aPD-L1 |
| Pfn1 | 5.80E-20 | 1.03E-15 | 0.58 | 0.989 | 0.993 | NKT | aPD-L1+aTGF-β vs aPD-L1 |
| AW112010 | 4.90E-10 | 8.73E-06 | 0.53 | 0.922 | 0.748 | ILC | aPD-L1+aTGF-β vs aPD-L1 |
| S100a6 | 5.19E-09 | 9.26E-05 | 0.49 | 0.968 | 0.887 | Epithelial cells | aPD-L1+aTGF-β vs aPD-L1 |
| Pfn1 | 1.23E-14 | 2.20E-10 | 0.48 | 0.986 | 0.978 | ILC | aPD-L1+aTGF-β vs aPD-L1 |
| Lgals1 | 1.10E-07 | 1.97E-03 | 0.47 | 0.984 | 1 | Fibroblasts | aPD-L1+aTGF-β vs aPD-L1 |
| Ier3 | 6.61E-38 | 1.18E-33 | 0.45 | 0.934 | 0.829 | Tumor_cells | aPD-L1+aTGF-β vs aPD-L1 |
| Rac2 | 1.69E-11 | 3.01E-07 | 0.45 | 0.868 | 0.612 | NKT | aPD-L1+aTGF-β vs aPD-L1 |
| C1qa | 3.25E-11 | 5.80E-07 | 0.44 | 0.822 | 0.69 | Macrophages | aPD-L1+aTGF-β vs aPD-L1 |
| Rpl22 | 1.75E-10 | 3.12E-06 | 0.44 | 0.957 | 0.885 | Fibroblasts | aPD-L1+aTGF-β vs aPD-L1 |
| Cxcl2 | 1.13E-17 | 2.01E-13 | 0.42 | 0.476 | 0.321 | Tumor_cells | aPD-L1+aTGF-β vs aPD-L1 |
| Cxcl1 | 1.01E-31 | 1.80E-27 | 0.41 | 0.797 | 0.607 | Tumor_cells | aPD-L1+aTGF-β vs aPD-L1 |
| Psme2 | 2.35E-14 | 4.18E-10 | 0.41 | 0.787 | 0.571 | Monocytes | aPD-L1+aTGF-β vs aPD-L1 |
| Lgals1 | 2.69E-14 | 4.79E-10 | 0.41 | 0.653 | 0.34 | Epithelial cells | aPD-L1+aTGF-β vs aPD-L1 |
| S100a10 | 2.74E-06 | 4.88E-02 | 0.40 | 0.908 | 0.928 | NKT | aPD-L1+aTGF-β vs aPD-L1 |
| Ifitm3 | 4.62E-07 | 8.23E-03 | 0.40 | 0.995 | 1 | Fibroblasts | aPD-L1+aTGF-β vs aPD-L1 |
| Rac2 | 1.24E-09 | 2.22E-05 | 0.40 | 0.875 | 0.719 | ILC | aPD-L1+aTGF-β vs aPD-L1 |
| C1qc | 8.74E-10 | 1.56E-05 | 0.39 | 0.812 | 0.69 | Macrophages | aPD-L1+aTGF-β vs aPD-L1 |
| Tmsb4x | 1.12E-10 | 1.99E-06 | 0.39 | 1 | 1 | NKT | aPD-L1+aTGF-β vs aPD-L1 |
| S100a4 | 2.05E-53 | 3.65E-49 | 0.38 | 0.966 | 0.912 | Tumor_cells | aPD-L1+aTGF-β vs aPD-L1 |
| Pfn1 | 1.50E-30 | 2.67E-26 | 0.38 | 0.975 | 0.972 | T cells | aPD-L1+aTGF-β vs aPD-L1 |
| Tmsb10 | 8.04E-09 | 1.43E-04 | 0.37 | 0.821 | 0.66 | Epithelial cells | aPD-L1+aTGF-β vs aPD-L1 |
| Rps15a | 1.91E-09 | 3.41E-05 | 0.37 | 0.995 | 1 | Fibroblasts | aPD-L1+aTGF-β vs aPD-L1 |
| Rpl41 | 1.31E-138 | 2.33E-134 | 0.37 | 1 | 1 | Tumor_cells | aPD-L1+aTGF-β vs aPD-L1 |
| Rps5 | 5.33E-11 | 9.50E-07 | 0.36 | 1 | 1 | NKT | aPD-L1+aTGF-β vs aPD-L1 |
| Rpl41 | 7.88E-12 | 1.40E-07 | 0.36 | 1 | 1 | Fibroblasts | aPD-L1+aTGF-β vs aPD-L1 |

| | | | | | | | |
|---|---|---|---|---|---|---|---|
| Sept1 | 8.03E-09 | 1.43E-04 | 0.36 | 0.751 | 0.474 | ILC | aPD-L1+aTGF-β vs aPD-L1 |
| Rps4x | 4.34E-11 | 7.73E-07 | 0.36 | 1 | 1 | NKT | aPD-L1+aTGF-β vs aPD-L1 |
| Arhgdib | 2.19E-07 | 3.90E-03 | 0.36 | 0.801 | 0.57 | ILC | aPD-L1+aTGF-β vs aPD-L1 |
| AW112010 | 2.58E-07 | 4.60E-03 | 0.36 | 0.57 | 0.496 | T cells | aPD-L1+aTGF-β vs aPD-L1 |
| Gapdh | 8.26E-07 | 1.47E-02 | 0.36 | 0.966 | 0.95 | NKT | aPD-L1+aTGF-β vs aPD-L1 |
| S100a6 | 2.56E-93 | 4.56E-89 | 0.36 | 1 | 1 | Tumor_cells | aPD-L1+aTGF-β vs aPD-L1 |
| Rpl30 | 1.58E-07 | 2.82E-03 | 0.35 | 0.984 | 0.989 | Fibroblasts | aPD-L1+aTGF-β vs aPD-L1 |
| Rpl36 | 4.74E-08 | 8.45E-04 | 0.35 | 0.979 | 0.954 | Fibroblasts | aPD-L1+aTGF-β vs aPD-L1 |
| Arhgdib | 5.75E-07 | 1.03E-02 | 0.35 | 0.727 | 0.489 | NKT | aPD-L1+aTGF-β vs aPD-L1 |
| Cd3d | 2.30E-07 | 4.09E-03 | 0.34 | 0.836 | 0.604 | NKT | aPD-L1+aTGF-β vs aPD-L1 |
| Actg1 | 1.02E-06 | 1.82E-02 | 0.34 | 0.991 | 0.993 | NKT | aPD-L1+aTGF-β vs aPD-L1 |
| Ifitm2 | 2.70E-08 | 4.81E-04 | 0.34 | 0.932 | 0.903 | Monocytes | aPD-L1+aTGF-β vs aPD-L1 |
| Ly6a | 1.05E-06 | 1.87E-02 | 0.34 | 0.687 | 0.539 | Epithelial cells | aPD-L1+aTGF-β vs aPD-L1 |
| Ppia | 3.33E-09 | 5.93E-05 | 0.33 | 0.986 | 0.986 | NKT | aPD-L1+aTGF-β vs aPD-L1 |
| C1qb | 4.31E-08 | 7.69E-04 | 0.33 | 0.819 | 0.683 | Macrophages | aPD-L1+aTGF-β vs aPD-L1 |
| Rps4x | 1.05E-09 | 1.88E-05 | 0.33 | 0.996 | 1 | ILC | aPD-L1+aTGF-β vs aPD-L1 |
| Chchd2 | 8.92E-07 | 1.59E-02 | 0.33 | 0.91 | 0.885 | Fibroblasts | aPD-L1+aTGF-β vs aPD-L1 |
| Tmsb4x | 1.56E-07 | 2.79E-03 | 0.33 | 1 | 1 | ILC | aPD-L1+aTGF-β vs aPD-L1 |
| Rpl18 | 3.05E-12 | 5.43E-08 | 0.33 | 1 | 1 | ILC | aPD-L1+aTGF-β vs aPD-L1 |
| Rps5 | 6.48E-11 | 1.16E-06 | 0.33 | 1 | 1 | ILC | aPD-L1+aTGF-β vs aPD-L1 |
| Rpl30 | 1.07E-115 | 1.90E-111 | 0.33 | 1 | 0.999 | Tumor_cells | aPD-L1+aTGF-β vs aPD-L1 |
| Nfkbia | 1.93E-33 | 3.44E-29 | 0.32 | 0.919 | 0.808 | Tumor_cells | aPD-L1+aTGF-β vs aPD-L1 |
| Rpl18a | 1.77E-11 | 3.16E-07 | 0.32 | 1 | 1 | NKT | aPD-L1+aTGF-β vs aPD-L1 |
| Serf2 | 7.55E-12 | 1.35E-07 | 0.32 | 0.923 | 0.762 | Epithelial cells | aPD-L1+aTGF-β vs aPD-L1 |
| Rpl22l1 | 2.16E-67 | 3.86E-63 | 0.32 | 0.989 | 0.952 | Tumor_cells | aPD-L1+aTGF-β vs aPD-L1 |
| Rpl22 | 5.25E-12 | 9.36E-08 | 0.32 | 0.978 | 0.969 | Epithelial cells | aPD-L1+aTGF-β vs aPD-L1 |
| Trac | 7.99E-07 | 1.42E-02 | 0.31 | 0.759 | 0.525 | NKT | aPD-L1+aTGF-β vs aPD-L1 |
| Rps11 | 7.04E-10 | 1.25E-05 | 0.31 | 1 | 1 | NKT | aPD-L1+aTGF-β vs aPD-L1 |
| Rpl35a | 2.48E-13 | 4.42E-09 | 0.31 | 0.995 | 0.98 | Epithelial cells | aPD-L1+aTGF-β vs aPD-L1 |
| Cfl1 | 3.06E-07 | 5.45E-03 | 0.31 | 0.954 | 0.942 | NKT | aPD-L1+aTGF-β vs aPD-L1 |
| Rpl34 | 9.33E-08 | 1.66E-03 | 0.31 | 0.984 | 0.966 | Fibroblasts | aPD-L1+aTGF-β vs aPD-L1 |
| Psme2 | 1.41E-14 | 2.52E-10 | 0.31 | 0.724 | 0.466 | Macrophages | aPD-L1+aTGF-β vs aPD-L1 |
| Rpl22 | 1.00E-09 | 1.78E-05 | 0.30 | 0.986 | 0.978 | NKT | aPD-L1+aTGF-β vs aPD-L1 |
| Rps5 | 1.37E-12 | 2.45E-08 | 0.30 | 0.988 | 0.978 | Macrophages | aPD-L1+aTGF-β vs aPD-L1 |
| Rpl18 | 8.13E-10 | 1.45E-05 | 0.30 | 1 | 1 | NKT | aPD-L1+aTGF-β vs aPD-L1 |
| Rpl41 | 4.42E-17 | 7.88E-13 | 0.30 | 1 | 1 | Epithelial cells | aPD-L1+aTGF-β vs aPD-L1 |
| Eef1a1 | 2.30E-08 | 4.09E-04 | 0.30 | 1 | 1 | ILC | aPD-L1+aTGF-β vs aPD-L1 |
| Rpl39 | 1.25E-13 | 2.23E-09 | 0.30 | 1 | 0.992 | Epithelial cells | aPD-L1+aTGF-β vs aPD-L1 |
| Rpl27a | 5.50E-08 | 9.80E-04 | 0.29 | 1 | 1 | Fibroblasts | aPD-L1+aTGF-β vs aPD-L1 |
| Rps15a | 4.18E-83 | 7.46E-79 | 0.29 | 1 | 0.994 | Tumor_cells | aPD-L1+aTGF-β vs aPD-L1 |
| Cox8a | 5.33E-10 | 9.50E-06 | 0.29 | 0.973 | 0.895 | Epithelial cells | aPD-L1+aTGF-β vs aPD-L1 |
| Rps4x | 1.71E-13 | 3.05E-09 | 0.29 | 0.987 | 0.981 | Macrophages | aPD-L1+aTGF-β vs aPD-L1 |
| Rpl37 | 1.33E-06 | 2.37E-02 | 0.29 | 0.989 | 0.954 | Fibroblasts | aPD-L1+aTGF-β vs aPD-L1 |
| Rpl35a | 1.05E-76 | 1.88E-72 | 0.29 | 0.997 | 0.989 | Tumor_cells | aPD-L1+aTGF-β vs aPD-L1 |
| Rpl18a | 2.83E-10 | 5.05E-06 | 0.29 | 1 | 1 | ILC | aPD-L1+aTGF-β vs aPD-L1 |
| Lgals1 | 4.02E-09 | 7.17E-05 | 0.29 | 0.99 | 0.966 | Macrophages | aPD-L1+aTGF-β vs aPD-L1 |
| Cfl1 | 1.22E-10 | 2.18E-06 | 0.29 | 0.973 | 0.944 | Monocytes | aPD-L1+aTGF-β vs aPD-L1 |
| Psmb8 | 4.35E-07 | 7.76E-03 | 0.28 | 0.879 | 0.837 | ILC | aPD-L1+aTGF-β vs aPD-L1 |
| Eif3h | 6.80E-07 | 1.21E-02 | 0.28 | 0.891 | 0.899 | NKT | aPD-L1+aTGF-β vs aPD-L1 |
| Pfn1 | 8.14E-11 | 1.45E-06 | 0.28 | 0.998 | 0.985 | Monocytes | aPD-L1+aTGF-β vs aPD-L1 |
| Cdkn1a | 2.39E-09 | 4.26E-05 | 0.28 | 0.778 | 0.699 | Neutrophils | aPD-L1+aTGF-β vs aPD-L1 |
| Rpl10a | 1.74E-11 | 3.10E-07 | 0.28 | 0.96 | 0.918 | Macrophages | aPD-L1+aTGF-β vs aPD-L1 |
| Myl12b | 2.74E-06 | 4.88E-02 | 0.28 | 0.744 | 0.541 | ILC | aPD-L1+aTGF-β vs aPD-L1 |
| Gapdh | 4.13E-10 | 7.36E-06 | 0.27 | 0.99 | 0.955 | Macrophages | aPD-L1+aTGF-β vs aPD-L1 |
| Cox5a | 2.02E-12 | 3.61E-08 | 0.27 | 0.806 | 0.608 | Macrophages | aPD-L1+aTGF-β vs aPD-L1 |

| | | | | | | | |
|---|---|---|---|---|---|---|---|
| Rpl36al | 4.37E-42 | 7.79E-38 | 0.27 | 0.538 | 0.254 | Tumor_cells | aPD-L1+aTGF-β vs aPD-L1 |
| Ifitm2 | 2.43E-12 | 4.33E-08 | 0.27 | 0.963 | 0.905 | Neutrophils | aPD-L1+aTGF-β vs aPD-L1 |
| Rps27l | 2.44E-33 | 4.34E-29 | 0.27 | 0.834 | 0.689 | Tumor_cells | aPD-L1+aTGF-β vs aPD-L1 |
| Gm11361 | 7.85E-40 | 1.40E-35 | 0.27 | 0.583 | 0.315 | Tumor_cells | aPD-L1+aTGF-β vs aPD-L1 |
| Uqcrb | 6.71E-39 | 1.20E-34 | 0.27 | 0.858 | 0.702 | Tumor_cells | aPD-L1+aTGF-β vs aPD-L1 |
| Fth1 | 4.25E-09 | 7.58E-05 | 0.27 | 1 | 1 | Macrophages | aPD-L1+aTGF-β vs aPD-L1 |
| Gapdh | 2.03E-08 | 3.62E-04 | 0.27 | 0.845 | 0.815 | T cells | aPD-L1+aTGF-β vs aPD-L1 |
| Tuba1b | 1.88E-25 | 3.35E-21 | 0.26 | 0.65 | 0.452 | Tumor_cells | aPD-L1+aTGF-β vs aPD-L1 |
| Rps11 | 4.39E-08 | 7.82E-04 | 0.26 | 1 | 1 | ILC | aPD-L1+aTGF-β vs aPD-L1 |
| Supt4a | 1.97E-06 | 3.51E-02 | 0.26 | 0.651 | 0.4 | ILC | aPD-L1+aTGF-β vs aPD-L1 |
| Psme1 | 1.22E-13 | 2.17E-09 | 0.26 | 0.728 | 0.643 | T cells | aPD-L1+aTGF-β vs aPD-L1 |
| Prelid1 | 9.89E-08 | 1.76E-03 | 0.26 | 0.571 | 0.372 | Monocytes | aPD-L1+aTGF-β vs aPD-L1 |
| Sept1 | 2.30E-06 | 4.10E-02 | 0.26 | 0.641 | 0.388 | NKT | aPD-L1+aTGF-β vs aPD-L1 |
| Bcl2a1d | 9.71E-08 | 1.73E-03 | 0.26 | 0.511 | 0.336 | Macrophages | aPD-L1+aTGF-β vs aPD-L1 |
| Rpl18 | 4.13E-17 | 7.36E-13 | 0.26 | 0.996 | 0.985 | Macrophages | aPD-L1+aTGF-β vs aPD-L1 |
| Lgals1 | 2.51E-41 | 4.48E-37 | 0.26 | 1 | 0.996 | Tumor_cells | aPD-L1+aTGF-β vs aPD-L1 |
| Cxcl2 | 3.50E-12 | 6.24E-08 | 0.26 | 0.439 | 0.176 | Epithelial cells | aPD-L1+aTGF-β vs aPD-L1 |
| Snx3 | 4.07E-07 | 7.26E-03 | 0.26 | 0.509 | 0.252 | ILC | aPD-L1+aTGF-β vs aPD-L1 |
| Pfn1 | 9.80E-16 | 1.75E-11 | 0.26 | 0.997 | 0.996 | Macrophages | aPD-L1+aTGF-β vs aPD-L1 |
| Ndufa4 | 2.69E-31 | 4.79E-27 | 0.26 | 0.869 | 0.746 | Tumor_cells | aPD-L1+aTGF-β vs aPD-L1 |
| Rgs10 | 9.02E-12 | 1.61E-07 | 0.26 | 0.695 | 0.455 | Macrophages | aPD-L1+aTGF-β vs aPD-L1 |
| Ly6a | 1.05E-09 | 1.87E-05 | 0.25 | 0.166 | 0.088 | B cells | aPD-L1+aTGF-β vs aPD-L1 |
| Snrpe | 8.52E-36 | 1.52E-31 | 0.25 | 0.723 | 0.5 | Tumor_cells | aPD-L1+aTGF-β vs aPD-L1 |
| Rps16 | 2.97E-08 | 5.30E-04 | 0.25 | 1 | 1 | NKT | aPD-L1+aTGF-β vs aPD-L1 |
| Srgn | 1.06E-17 | 1.89E-13 | 0.25 | 1 | 1 | Neutrophils | aPD-L1+aTGF-β vs aPD-L1 |
| Upp1 | 8.92E-07 | 1.59E-02 | 0.25 | 0.686 | 0.596 | Neutrophils | aPD-L1+aTGF-β vs aPD-L1 |
| Isg20 | 1.15E-06 | 2.04E-02 | 0.25 | 0.546 | 0.458 | Neutrophils | aPD-L1+aTGF-β vs aPD-L1 |
| Crip1 | 1.21E-16 | 2.16E-12 | 0.25 | 0.697 | 0.575 | Tumor_cells | aPD-L1+aTGF-β vs aPD-L1 |
| Cxcl14 | 2.60E-06 | 4.63E-02 | 0.25 | 0.564 | 0.418 | Macrophages | aPD-L1+aTGF-β vs aPD-L1 |
| Rac2 | 4.28E-15 | 7.64E-11 | 0.25 | 0.855 | 0.784 | T cells | aPD-L1+aTGF-β vs aPD-L1 |
| Pfn1 | 3.77E-28 | 6.73E-24 | 0.25 | 0.935 | 0.908 | B cells | aPD-L1+aTGF-β vs aPD-L1 |
| Rpl30 | 8.13E-10 | 1.45E-05 | 0.25 | 1 | 0.992 | Epithelial cells | aPD-L1+aTGF-β vs aPD-L1 |
| Elob | 2.59E-08 | 4.62E-04 | 0.25 | 0.752 | 0.574 | Epithelial cells | aPD-L1+aTGF-β vs aPD-L1 |
| Rpl15 | 2.58E-13 | 4.60E-09 | 0.25 | 0.985 | 0.966 | Macrophages | aPD-L1+aTGF-β vs aPD-L1 |
| Zfp36 | 3.14E-07 | 5.60E-03 | -0.25 | 0.982 | 0.978 | Neutrophils | aPD-L1+aTGF-β vs aPD-L1 |
| mt-Atp8 | 4.59E-08 | 8.18E-04 | -0.25 | 0.38 | 0.597 | Monocytes | aPD-L1+aTGF-β vs aPD-L1 |
| Myh9 | 1.79E-18 | 3.18E-14 | -0.25 | 0.439 | 0.578 | Tumor_cells | aPD-L1+aTGF-β vs aPD-L1 |
| Rpl17 | 6.87E-11 | 1.22E-06 | -0.25 | 0.995 | 1 | Epithelial cells | aPD-L1+aTGF-β vs aPD-L1 |
| Rpl5 | 3.88E-13 | 6.92E-09 | -0.25 | 0.993 | 0.996 | Epithelial cells | aPD-L1+aTGF-β vs aPD-L1 |
| Rplp2 | 4.95E-62 | 8.82E-58 | -0.25 | 0.998 | 1 | B cells | aPD-L1+aTGF-β vs aPD-L1 |
| Rpl41 | 3.81E-56 | 6.79E-52 | -0.25 | 1 | 1 | B cells | aPD-L1+aTGF-β vs aPD-L1 |
| Rplp0 | 1.04E-12 | 1.85E-08 | -0.25 | 0.902 | 0.926 | Neutrophils | aPD-L1+aTGF-β vs aPD-L1 |
| Rpl37a | 1.52E-45 | 2.72E-41 | -0.25 | 1 | 1 | T cells | aPD-L1+aTGF-β vs aPD-L1 |
| Hist1h1c | 1.99E-09 | 3.55E-05 | -0.25 | 0.385 | 0.496 | B cells | aPD-L1+aTGF-β vs aPD-L1 |
| Gm10260 | 1.13E-10 | 2.02E-06 | -0.26 | 0.794 | 0.914 | Epithelial cells | aPD-L1+aTGF-β vs aPD-L1 |
| Rpl35a | 3.57E-74 | 6.37E-70 | -0.26 | 0.997 | 0.999 | B cells | aPD-L1+aTGF-β vs aPD-L1 |
| Rpsa | 1.52E-15 | 2.70E-11 | -0.26 | 0.888 | 0.909 | Neutrophils | aPD-L1+aTGF-β vs aPD-L1 |
| Rpl13a | 2.92E-20 | 5.20E-16 | -0.26 | 0.502 | 0.696 | T cells | aPD-L1+aTGF-β vs aPD-L1 |
| Malat1 | 2.51E-06 | 4.47E-02 | -0.26 | 1 | 1 | Monocytes | aPD-L1+aTGF-β vs aPD-L1 |
| Cldn3 | 1.82E-07 | 3.24E-03 | -0.26 | 0.737 | 0.82 | Epithelial cells | aPD-L1+aTGF-β vs aPD-L1 |
| Rpl37 | 1.14E-68 | 2.02E-64 | -0.26 | 0.999 | 1 | B cells | aPD-L1+aTGF-β vs aPD-L1 |
| Rps3a1 | 6.04E-16 | 1.08E-11 | -0.26 | 0.9 | 0.909 | Neutrophils | aPD-L1+aTGF-β vs aPD-L1 |
| mt-Nd3 | 5.05E-34 | 9.01E-30 | -0.26 | 0.37 | 0.613 | B cells | aPD-L1+aTGF-β vs aPD-L1 |
| AC160336.1 | 1.68E-11 | 2.99E-07 | -0.27 | 0.525 | 0.648 | B cells | aPD-L1+aTGF-β vs aPD-L1 |
| Rpl37 | 8.77E-18 | 1.56E-13 | -0.27 | 0.924 | 0.941 | Neutrophils | aPD-L1+aTGF-β vs aPD-L1 |

| | | | | | | | |
|---|---|---|---|---|---|---|---|
| Ablim1 | 6.74E-07 | 1.20E-02 | -0.27 | 0.181 | 0.483 | Fibroblasts | aPD-L1+aTGF-β vs aPD-L1 |
| Rplp2 | 6.36E-14 | 1.13E-09 | -0.27 | 0.901 | 0.912 | Neutrophils | aPD-L1+aTGF-β vs aPD-L1 |
| Hist1h2bc | 4.75E-11 | 8.47E-07 | -0.27 | 0.174 | 0.278 | B cells | aPD-L1+aTGF-β vs aPD-L1 |
| Ier5 | 1.87E-16 | 3.33E-12 | -0.27 | 0.669 | 0.83 | T cells | aPD-L1+aTGF-β vs aPD-L1 |
| mt-Co3 | 5.78E-07 | 1.03E-02 | -0.27 | 1 | 1 | NKT | aPD-L1+aTGF-β vs aPD-L1 |
| Lars2 | 1.98E-31 | 3.53E-27 | -0.27 | 0.175 | 0.364 | Tumor_cells | aPD-L1+aTGF-β vs aPD-L1 |
| Rpl35a | 1.51E-45 | 2.68E-41 | -0.27 | 1 | 1 | T cells | aPD-L1+aTGF-β vs aPD-L1 |
| Fdps | 4.01E-13 | 7.14E-09 | -0.27 | 0.411 | 0.508 | Tumor_cells | aPD-L1+aTGF-β vs aPD-L1 |
| Rpl39 | 1.16E-49 | 2.06E-45 | -0.27 | 1 | 1 | T cells | aPD-L1+aTGF-β vs aPD-L1 |
| mt-Nd3 | 5.30E-15 | 9.45E-11 | -0.27 | 0.409 | 0.571 | T cells | aPD-L1+aTGF-β vs aPD-L1 |
| Gadd45g | 3.70E-08 | 6.59E-04 | -0.27 | 0.32 | 0.41 | Tumor_cells | aPD-L1+aTGF-β vs aPD-L1 |
| Gm26825 | 1.34E-13 | 2.38E-09 | -0.27 | 0.361 | 0.525 | T cells | aPD-L1+aTGF-β vs aPD-L1 |
| Rpl35 | 1.94E-59 | 3.46E-55 | -0.28 | 0.989 | 0.998 | B cells | aPD-L1+aTGF-β vs aPD-L1 |
| Rpl23 | 2.48E-18 | 4.43E-14 | -0.28 | 0.964 | 0.968 | Neutrophils | aPD-L1+aTGF-β vs aPD-L1 |
| Tspan5 | 3.70E-07 | 6.59E-03 | -0.28 | 0.112 | 0.368 | Fibroblasts | aPD-L1+aTGF-β vs aPD-L1 |
| Rpl27a | 1.69E-18 | 3.02E-14 | -0.28 | 0.919 | 0.949 | Neutrophils | aPD-L1+aTGF-β vs aPD-L1 |
| Eno1 | 9.79E-24 | 1.74E-19 | -0.28 | 0.7 | 0.777 | Tumor_cells | aPD-L1+aTGF-β vs aPD-L1 |
| mt-Co2 | 1.00E-08 | 1.78E-04 | -0.28 | 1 | 1 | NKT | aPD-L1+aTGF-β vs aPD-L1 |
| Ctsl | 2.19E-14 | 3.91E-10 | -0.28 | 0.138 | 0.46 | NKT | aPD-L1+aTGF-β vs aPD-L1 |
| Atp5k | 1.03E-08 | 1.84E-04 | -0.28 | 0.492 | 0.673 | Monocytes | aPD-L1+aTGF-β vs aPD-L1 |
| Zfp385a | 2.55E-07 | 4.55E-03 | -0.28 | 0.106 | 0.368 | Fibroblasts | aPD-L1+aTGF-β vs aPD-L1 |
| Rps29 | 6.07E-23 | 1.08E-18 | -0.28 | 1 | 1 | Macrophages | aPD-L1+aTGF-β vs aPD-L1 |
| Rpl35a | 8.23E-18 | 1.47E-13 | -0.28 | 0.888 | 0.9 | Neutrophils | aPD-L1+aTGF-β vs aPD-L1 |
| AC160336.1 | 2.74E-06 | 4.89E-02 | -0.28 | 0.286 | 0.422 | Macrophages | aPD-L1+aTGF-β vs aPD-L1 |
| Rps19 | 8.65E-19 | 1.54E-14 | -0.28 | 0.886 | 0.914 | Neutrophils | aPD-L1+aTGF-β vs aPD-L1 |
| Rps27 | 4.38E-12 | 7.81E-08 | -0.29 | 0.987 | 0.993 | Macrophages | aPD-L1+aTGF-β vs aPD-L1 |
| Socs3 | 6.54E-07 | 1.17E-02 | -0.29 | 0.201 | 0.424 | NKT | aPD-L1+aTGF-β vs aPD-L1 |
| Rpl37a | 1.30E-11 | 2.32E-07 | -0.29 | 0.99 | 0.995 | Monocytes | aPD-L1+aTGF-β vs aPD-L1 |
| Rpl34 | 9.47E-18 | 1.69E-13 | -0.29 | 0.907 | 0.941 | Neutrophils | aPD-L1+aTGF-β vs aPD-L1 |
| Rps21 | 3.92E-56 | 6.99E-52 | -0.29 | 1 | 1 | T cells | aPD-L1+aTGF-β vs aPD-L1 |
| Rps21 | 5.13E-99 | 9.15E-95 | -0.29 | 0.998 | 0.999 | B cells | aPD-L1+aTGF-β vs aPD-L1 |
| Hspb1 | 1.19E-15 | 2.13E-11 | -0.29 | 0.028 | 0.296 | ILC | aPD-L1+aTGF-β vs aPD-L1 |
| Rpl38 | 8.76E-11 | 1.56E-06 | -0.29 | 0.959 | 0.964 | Monocytes | aPD-L1+aTGF-β vs aPD-L1 |
| Gm10260 | 8.68E-41 | 1.55E-36 | -0.29 | 0.866 | 0.936 | Tumor_cells | aPD-L1+aTGF-β vs aPD-L1 |
| Rps23 | 9.16E-22 | 1.63E-17 | -0.30 | 0.909 | 0.939 | Neutrophils | aPD-L1+aTGF-β vs aPD-L1 |
| Rps28 | 9.37E-16 | 1.67E-11 | -0.30 | 0.95 | 0.981 | Macrophages | aPD-L1+aTGF-β vs aPD-L1 |
| Nfix | 2.14E-10 | 3.81E-06 | -0.30 | 0.33 | 0.594 | Epithelial cells | aPD-L1+aTGF-β vs aPD-L1 |
| Rpl37a | 5.62E-98 | 1.00E-93 | -0.30 | 0.998 | 1 | B cells | aPD-L1+aTGF-β vs aPD-L1 |
| mt-Cytb | 2.77E-07 | 4.94E-03 | -0.30 | 0.971 | 1 | NKT | aPD-L1+aTGF-β vs aPD-L1 |
| Rps28 | 6.14E-10 | 1.09E-05 | -0.30 | 0.923 | 0.944 | Monocytes | aPD-L1+aTGF-β vs aPD-L1 |
| mt-Atp8 | 1.78E-06 | 3.18E-02 | -0.31 | 0.365 | 0.568 | NKT | aPD-L1+aTGF-β vs aPD-L1 |
| mt-Atp8 | 1.59E-41 | 2.83E-37 | -0.31 | 0.388 | 0.632 | B cells | aPD-L1+aTGF-β vs aPD-L1 |
| Malat1 | 8.14E-30 | 1.45E-25 | -0.31 | 1 | 1 | T cells | aPD-L1+aTGF-β vs aPD-L1 |
| Gda | 8.76E-07 | 1.56E-02 | -0.32 | 0.324 | 0.632 | Fibroblasts | aPD-L1+aTGF-β vs aPD-L1 |
| Hmgb2 | 2.16E-31 | 3.84E-27 | -0.32 | 0.568 | 0.782 | B cells | aPD-L1+aTGF-β vs aPD-L1 |
| Rpl30 | 7.36E-19 | 1.31E-14 | -0.32 | 0.902 | 0.926 | Neutrophils | aPD-L1+aTGF-β vs aPD-L1 |
| mt-Atp8 | 3.22E-28 | 5.75E-24 | -0.32 | 0.415 | 0.671 | T cells | aPD-L1+aTGF-β vs aPD-L1 |
| Pdpn | 1.42E-06 | 2.53E-02 | -0.33 | 0.484 | 0.793 | Fibroblasts | aPD-L1+aTGF-β vs aPD-L1 |
| Rpl13 | 2.04E-25 | 3.64E-21 | -0.33 | 0.92 | 0.958 | Neutrophils | aPD-L1+aTGF-β vs aPD-L1 |
| AC160336.1 | 9.84E-14 | 1.75E-09 | -0.33 | 0.298 | 0.469 | T cells | aPD-L1+aTGF-β vs aPD-L1 |
| Rpl38 | 2.63E-57 | 4.69E-53 | -0.33 | 0.998 | 1 | T cells | aPD-L1+aTGF-β vs aPD-L1 |
| mt-Atp8 | 3.35E-15 | 5.97E-11 | -0.33 | 0.454 | 0.649 | Macrophages | aPD-L1+aTGF-β vs aPD-L1 |
| Rpl38 | 2.61E-93 | 4.66E-89 | -0.33 | 0.997 | 0.999 | B cells | aPD-L1+aTGF-β vs aPD-L1 |
| mt-Cytb | 2.25E-06 | 4.01E-02 | -0.33 | 0.972 | 0.97 | ILC | aPD-L1+aTGF-β vs aPD-L1 |
| Rps29 | 1.24E-17 | 2.22E-13 | -0.33 | 0.995 | 1 | Monocytes | aPD-L1+aTGF-β vs aPD-L1 |

| | | | | | | | |
|---|---|---|---|---|---|---|---|
| Antxr2 | 1.97E-08 | 3.51E-04 | -0.33 | 0.186 | 0.506 | Fibroblasts | aPD-L1+aTGF-β vs aPD-L1 |
| Fosb | 6.67E-07 | 1.19E-02 | -0.33 | 0.767 | 0.942 | NKT | aPD-L1+aTGF-β vs aPD-L1 |
| Pcsk6 | 2.92E-08 | 5.21E-04 | -0.33 | 0.085 | 0.345 | Fibroblasts | aPD-L1+aTGF-β vs aPD-L1 |
| mt-Atp8 | 5.91E-07 | 1.05E-02 | -0.33 | 0.345 | 0.541 | ILC | aPD-L1+aTGF-β vs aPD-L1 |
| Rpl32 | 3.06E-18 | 5.45E-14 | -0.33 | 0.884 | 0.9 | Neutrophils | aPD-L1+aTGF-β vs aPD-L1 |
| Rps21 | 1.37E-22 | 2.45E-18 | -0.34 | 0.876 | 0.902 | Neutrophils | aPD-L1+aTGF-β vs aPD-L1 |
| Rps20 | 2.38E-31 | 4.24E-27 | -0.34 | 0.909 | 0.948 | Neutrophils | aPD-L1+aTGF-β vs aPD-L1 |
| mt-Co2 | 9.30E-09 | 1.66E-04 | -0.34 | 1 | 1 | ILC | aPD-L1+aTGF-β vs aPD-L1 |
| Lars2 | 1.86E-13 | 3.31E-09 | -0.34 | 0.161 | 0.422 | Epithelial cells | aPD-L1+aTGF-β vs aPD-L1 |
| Rpl39 | 8.09E-113 | 1.44E-108 | -0.34 | 0.996 | 1 | B cells | aPD-L1+aTGF-β vs aPD-L1 |
| Cavin1 | 2.49E-06 | 4.43E-02 | -0.34 | 0.452 | 0.724 | Fibroblasts | aPD-L1+aTGF-β vs aPD-L1 |
| Rpl37a | 1.49E-06 | 2.65E-02 | -0.34 | 0.985 | 1 | DC | aPD-L1+aTGF-β vs aPD-L1 |
| Osr1 | 5.77E-08 | 1.03E-03 | -0.34 | 0.229 | 0.54 | Fibroblasts | aPD-L1+aTGF-β vs aPD-L1 |
| Rps15a | 4.08E-32 | 7.27E-28 | -0.35 | 0.881 | 0.939 | Neutrophils | aPD-L1+aTGF-β vs aPD-L1 |
| Lars2 | 6.56E-51 | 1.17E-46 | -0.35 | 0.313 | 0.612 | B cells | aPD-L1+aTGF-β vs aPD-L1 |
| B2m | 1.15E-59 | 2.05E-55 | -0.35 | 1 | 0.999 | Tumor_cells | aPD-L1+aTGF-β vs aPD-L1 |
| Prelp | 2.68E-06 | 4.77E-02 | -0.35 | 0.441 | 0.713 | Fibroblasts | aPD-L1+aTGF-β vs aPD-L1 |
| Rps29 | 1.69E-174 | 3.01E-170 | -0.35 | 1 | 1 | B cells | aPD-L1+aTGF-β vs aPD-L1 |
| Lama4 | 1.41E-06 | 2.51E-02 | -0.35 | 0.441 | 0.736 | Fibroblasts | aPD-L1+aTGF-β vs aPD-L1 |
| mt-Nd1 | 1.85E-07 | 3.29E-03 | -0.35 | 0.89 | 0.956 | ILC | aPD-L1+aTGF-β vs aPD-L1 |
| Grb10 | 8.62E-07 | 1.54E-02 | -0.35 | 0.213 | 0.494 | Fibroblasts | aPD-L1+aTGF-β vs aPD-L1 |
| Ifi27l2a | 4.23E-11 | 7.54E-07 | -0.36 | 0.59 | 0.716 | Neutrophils | aPD-L1+aTGF-β vs aPD-L1 |
| Rpl34 | 1.67E-06 | 2.97E-02 | -0.36 | 1 | 1 | DC | aPD-L1+aTGF-β vs aPD-L1 |
| Rpl38 | 7.51E-07 | 1.34E-02 | -0.36 | 1 | 1 | DC | aPD-L1+aTGF-β vs aPD-L1 |
| Rpl36 | 1.34E-06 | 2.39E-02 | -0.36 | 0.985 | 1 | DC | aPD-L1+aTGF-β vs aPD-L1 |
| Gm47283 | 6.30E-43 | 1.12E-38 | -0.36 | 0.628 | 0.842 | B cells | aPD-L1+aTGF-β vs aPD-L1 |
| Fosb | 7.36E-07 | 1.31E-02 | -0.37 | 0.858 | 0.948 | ILC | aPD-L1+aTGF-β vs aPD-L1 |
| mt-Co3 | 6.75E-10 | 1.20E-05 | -0.37 | 0.996 | 1 | ILC | aPD-L1+aTGF-β vs aPD-L1 |
| Rplp1 | 2.35E-33 | 4.19E-29 | -0.37 | 0.946 | 0.973 | Neutrophils | aPD-L1+aTGF-β vs aPD-L1 |
| Rps27 | 3.49E-166 | 6.22E-162 | -0.38 | 0.999 | 1 | B cells | aPD-L1+aTGF-β vs aPD-L1 |
| Rps27 | 2.14E-90 | 3.82E-86 | -0.38 | 1 | 1 | T cells | aPD-L1+aTGF-β vs aPD-L1 |
| Rpl39 | 1.43E-30 | 2.56E-26 | -0.38 | 0.91 | 0.938 | Neutrophils | aPD-L1+aTGF-β vs aPD-L1 |
| Btg1 | 6.77E-09 | 1.21E-04 | -0.38 | 0.743 | 0.866 | Macrophages | aPD-L1+aTGF-β vs aPD-L1 |
| Cmah | 2.73E-11 | 4.87E-07 | -0.39 | 0.165 | 0.54 | Fibroblasts | aPD-L1+aTGF-β vs aPD-L1 |
| mt-Atp6 | 1.31E-12 | 2.33E-08 | -0.39 | 1 | 1 | NKT | aPD-L1+aTGF-β vs aPD-L1 |
| Rpl37a | 1.58E-29 | 2.81E-25 | -0.39 | 0.914 | 0.954 | Neutrophils | aPD-L1+aTGF-β vs aPD-L1 |
| Gm47283 | 3.62E-17 | 6.46E-13 | -0.39 | 0.414 | 0.785 | Epithelial cells | aPD-L1+aTGF-β vs aPD-L1 |
| Rps29 | 2.18E-102 | 3.88E-98 | -0.40 | 1 | 1 | T cells | aPD-L1+aTGF-β vs aPD-L1 |
| Osr2 | 7.54E-08 | 1.34E-03 | -0.40 | 0.138 | 0.425 | Fibroblasts | aPD-L1+aTGF-β vs aPD-L1 |
| Rps28 | 8.97E-98 | 1.60E-93 | -0.40 | 0.962 | 0.997 | B cells | aPD-L1+aTGF-β vs aPD-L1 |
| mt-Atp6 | 2.17E-11 | 3.87E-07 | -0.40 | 1 | 1 | ILC | aPD-L1+aTGF-β vs aPD-L1 |
| mt-Nd2 | 4.07E-11 | 7.26E-07 | -0.40 | 0.931 | 0.993 | NKT | aPD-L1+aTGF-β vs aPD-L1 |
| Rps28 | 5.86E-65 | 1.05E-60 | -0.40 | 0.992 | 0.998 | T cells | aPD-L1+aTGF-β vs aPD-L1 |
| Antxr1 | 6.17E-08 | 1.10E-03 | -0.41 | 0.309 | 0.621 | Fibroblasts | aPD-L1+aTGF-β vs aPD-L1 |
| Thbs3 | 1.15E-07 | 2.05E-03 | -0.42 | 0.404 | 0.69 | Fibroblasts | aPD-L1+aTGF-β vs aPD-L1 |
| Ddr2 | 1.20E-08 | 2.14E-04 | -0.42 | 0.372 | 0.655 | Fibroblasts | aPD-L1+aTGF-β vs aPD-L1 |
| Rps24 | 4.71E-41 | 8.40E-37 | -0.42 | 0.904 | 0.965 | Neutrophils | aPD-L1+aTGF-β vs aPD-L1 |
| Rps27 | 3.74E-42 | 6.67E-38 | -0.42 | 0.979 | 0.992 | Neutrophils | aPD-L1+aTGF-β vs aPD-L1 |
| Rnase4 | 2.68E-06 | 4.78E-02 | -0.42 | 0.718 | 0.839 | Fibroblasts | aPD-L1+aTGF-β vs aPD-L1 |
| Lars2 | 9.38E-09 | 1.67E-04 | -0.42 | 0.335 | 0.585 | ILC | aPD-L1+aTGF-β vs aPD-L1 |
| Ahnak | 5.70E-07 | 1.02E-02 | -0.43 | 0.777 | 0.931 | Fibroblasts | aPD-L1+aTGF-β vs aPD-L1 |
| mt-Nd1 | 3.05E-12 | 5.44E-08 | -0.43 | 0.891 | 0.971 | NKT | aPD-L1+aTGF-β vs aPD-L1 |
| Wnt2 | 1.17E-07 | 2.09E-03 | -0.43 | 0.186 | 0.471 | Fibroblasts | aPD-L1+aTGF-β vs aPD-L1 |
| Dcn | 1.01E-08 | 1.80E-04 | -0.44 | 0.957 | 0.989 | Fibroblasts | aPD-L1+aTGF-β vs aPD-L1 |
| Tgfbr2 | 5.35E-07 | 9.53E-03 | -0.44 | 0.58 | 0.805 | Fibroblasts | aPD-L1+aTGF-β vs aPD-L1 |

| | | | | | | | |
|---|---|---|---|---|---|---|---|
| Gm47283 | 2.71E-21 | 4.83E-17 | -0.44 | 0.316 | 0.601 | Macrophages | aPD-L1+aTGF-β vs aPD-L1 |
| Man1a | 1.08E-08 | 1.93E-04 | -0.44 | 0.319 | 0.655 | Fibroblasts | aPD-L1+aTGF-β vs aPD-L1 |
| Pamr1 | 2.46E-10 | 4.39E-06 | -0.45 | 0.17 | 0.529 | Fibroblasts | aPD-L1+aTGF-β vs aPD-L1 |
| Rps27 | 3.07E-07 | 5.47E-03 | -0.45 | 1 | 1 | DC | aPD-L1+aTGF-β vs aPD-L1 |
| Rps21 | 4.01E-09 | 7.16E-05 | -0.45 | 1 | 1 | DC | aPD-L1+aTGF-β vs aPD-L1 |
| Mmp2 | 1.14E-06 | 2.04E-02 | -0.46 | 0.824 | 0.885 | Fibroblasts | aPD-L1+aTGF-β vs aPD-L1 |
| Rpl39 | 6.18E-08 | 1.10E-03 | -0.46 | 1 | 1 | DC | aPD-L1+aTGF-β vs aPD-L1 |
| Prss23 | 3.02E-07 | 5.38E-03 | -0.46 | 0.452 | 0.736 | Fibroblasts | aPD-L1+aTGF-β vs aPD-L1 |
| Scara5 | 5.52E-10 | 9.84E-06 | -0.46 | 0.213 | 0.563 | Fibroblasts | aPD-L1+aTGF-β vs aPD-L1 |
| Tppp3 | 9.30E-07 | 1.66E-02 | -0.47 | 0.351 | 0.632 | Fibroblasts | aPD-L1+aTGF-β vs aPD-L1 |
| Rps29 | 1.20E-08 | 2.13E-04 | -0.47 | 1 | 1 | DC | aPD-L1+aTGF-β vs aPD-L1 |
| Cdkn1c | 2.60E-06 | 4.64E-02 | -0.47 | 0.234 | 0.494 | Fibroblasts | aPD-L1+aTGF-β vs aPD-L1 |
| Lrrn4cl | 4.02E-11 | 7.17E-07 | -0.47 | 0.223 | 0.598 | Fibroblasts | aPD-L1+aTGF-β vs aPD-L1 |
| Cd248 | 3.88E-07 | 6.92E-03 | -0.47 | 0.569 | 0.793 | Fibroblasts | aPD-L1+aTGF-β vs aPD-L1 |
| Scara3 | 1.71E-10 | 3.05E-06 | -0.47 | 0.255 | 0.621 | Fibroblasts | aPD-L1+aTGF-β vs aPD-L1 |
| C4b | 1.14E-06 | 2.03E-02 | -0.47 | 0.803 | 0.897 | Fibroblasts | aPD-L1+aTGF-β vs aPD-L1 |
| Gm47283 | 8.09E-47 | 1.44E-42 | -0.47 | 0.612 | 0.861 | T cells | aPD-L1+aTGF-β vs aPD-L1 |
| Rpl37 | 1.38E-09 | 2.46E-05 | -0.48 | 1 | 1 | DC | aPD-L1+aTGF-β vs aPD-L1 |
| Gm42418 | 2.75E-12 | 4.90E-08 | -0.48 | 1 | 1 | ILC | aPD-L1+aTGF-β vs aPD-L1 |
| Klf4 | 5.32E-07 | 9.49E-03 | -0.48 | 0.516 | 0.805 | Fibroblasts | aPD-L1+aTGF-β vs aPD-L1 |
| mt-Nd2 | 5.27E-13 | 9.39E-09 | -0.48 | 0.929 | 0.978 | ILC | aPD-L1+aTGF-β vs aPD-L1 |
| Emilin2 | 6.62E-09 | 1.18E-04 | -0.50 | 0.383 | 0.69 | Fibroblasts | aPD-L1+aTGF-β vs aPD-L1 |
| Tnxb | 3.86E-08 | 6.89E-04 | -0.50 | 0.574 | 0.862 | Fibroblasts | aPD-L1+aTGF-β vs aPD-L1 |
| Creb5 | 2.76E-08 | 4.92E-04 | -0.50 | 0.266 | 0.563 | Fibroblasts | aPD-L1+aTGF-β vs aPD-L1 |
| Rps29 | 1.06E-58 | 1.88E-54 | -0.50 | 0.975 | 0.995 | Neutrophils | aPD-L1+aTGF-β vs aPD-L1 |
| Gm47283 | 5.15E-14 | 9.18E-10 | -0.51 | 0.378 | 0.663 | Monocytes | aPD-L1+aTGF-β vs aPD-L1 |
| Lars2 | 5.44E-11 | 9.70E-07 | -0.51 | 0.34 | 0.747 | Fibroblasts | aPD-L1+aTGF-β vs aPD-L1 |
| Fndc1 | 1.54E-08 | 2.74E-04 | -0.51 | 0.42 | 0.724 | Fibroblasts | aPD-L1+aTGF-β vs aPD-L1 |
| AY036118 | 6.97E-36 | 1.24E-31 | -0.52 | 0.588 | 0.76 | Tumor_cells | aPD-L1+aTGF-β vs aPD-L1 |
| Malat1 | 4.42E-19 | 7.88E-15 | -0.52 | 1 | 1 | NKT | aPD-L1+aTGF-β vs aPD-L1 |
| Lars2 | 1.38E-40 | 2.46E-36 | -0.53 | 0.409 | 0.699 | T cells | aPD-L1+aTGF-β vs aPD-L1 |
| Smad7 | 1.39E-07 | 2.48E-03 | -0.53 | 0.583 | 0.748 | NKT | aPD-L1+aTGF-β vs aPD-L1 |
| Gm42418 | 2.19E-145 | 3.91E-141 | -0.53 | 1 | 1 | B cells | aPD-L1+aTGF-β vs aPD-L1 |
| Lars2 | 3.74E-41 | 6.67E-37 | -0.54 | 0.266 | 0.571 | Neutrophils | aPD-L1+aTGF-β vs aPD-L1 |
| Gm42418 | 1.55E-15 | 2.76E-11 | -0.54 | 1 | 1 | Monocytes | aPD-L1+aTGF-β vs aPD-L1 |
| Timp2 | 2.91E-09 | 5.19E-05 | -0.54 | 0.883 | 0.954 | Fibroblasts | aPD-L1+aTGF-β vs aPD-L1 |
| Nov | 2.31E-08 | 4.11E-04 | -0.55 | 0.239 | 0.54 | Fibroblasts | aPD-L1+aTGF-β vs aPD-L1 |
| Gm42418 | 3.16E-19 | 5.63E-15 | -0.56 | 1 | 1 | Macrophages | aPD-L1+aTGF-β vs aPD-L1 |
| AC160336.1 | 3.66E-09 | 6.52E-05 | -0.56 | 0.181 | 0.403 | NKT | aPD-L1+aTGF-β vs aPD-L1 |
| Efemp1 | 1.55E-08 | 2.76E-04 | -0.57 | 0.537 | 0.782 | Fibroblasts | aPD-L1+aTGF-β vs aPD-L1 |
| Smad7 | 9.92E-12 | 1.77E-07 | -0.57 | 0.662 | 0.83 | ILC | aPD-L1+aTGF-β vs aPD-L1 |
| Pi16 | 7.87E-08 | 1.40E-03 | -0.58 | 0.665 | 0.885 | Fibroblasts | aPD-L1+aTGF-β vs aPD-L1 |
| Gm47283 | 1.41E-08 | 2.51E-04 | -0.58 | 0.575 | 0.741 | NKT | aPD-L1+aTGF-β vs aPD-L1 |
| Malat1 | 3.70E-17 | 6.60E-13 | -0.59 | 1 | 1 | ILC | aPD-L1+aTGF-β vs aPD-L1 |
| Smad7 | 1.84E-54 | 3.28E-50 | -0.60 | 0.674 | 0.882 | T cells | aPD-L1+aTGF-β vs aPD-L1 |
| Cd34 | 2.20E-11 | 3.92E-07 | -0.60 | 0.479 | 0.828 | Fibroblasts | aPD-L1+aTGF-β vs aPD-L1 |
| Gm42418 | 2.75E-91 | 4.91E-87 | -0.60 | 1 | 1 | Neutrophils | aPD-L1+aTGF-β vs aPD-L1 |
| Cd55 | 1.28E-06 | 2.29E-02 | -0.60 | 0.319 | 0.563 | Fibroblasts | aPD-L1+aTGF-β vs aPD-L1 |
| Dpt | 3.95E-10 | 7.05E-06 | -0.61 | 0.702 | 0.908 | Fibroblasts | aPD-L1+aTGF-β vs aPD-L1 |
| Gm47283 | 4.92E-12 | 8.77E-08 | -0.62 | 0.566 | 0.778 | ILC | aPD-L1+aTGF-β vs aPD-L1 |
| Adamts5 | 4.12E-09 | 7.35E-05 | -0.62 | 0.527 | 0.816 | Fibroblasts | aPD-L1+aTGF-β vs aPD-L1 |
| Sema3c | 8.50E-09 | 1.51E-04 | -0.62 | 0.314 | 0.632 | Fibroblasts | aPD-L1+aTGF-β vs aPD-L1 |
| Dmkn | 1.91E-06 | 3.40E-02 | -0.63 | 0.128 | 0.356 | Fibroblasts | aPD-L1+aTGF-β vs aPD-L1 |
| Lars2 | 1.14E-21 | 2.03E-17 | -0.63 | 0.476 | 0.728 | Macrophages | aPD-L1+aTGF-β vs aPD-L1 |
| Rps28 | 3.20E-10 | 5.71E-06 | -0.64 | 0.939 | 1 | DC | aPD-L1+aTGF-β vs aPD-L1 |

| | | | | | | | |
|---|---|---|---|---|---|---|---|
| Htra3 | 3.17E-12 | 5.65E-08 | -0.65 | 0.569 | 0.874 | Fibroblasts | aPD-L1+aTGF-β vs aPD-L1 |
| Col14a1 | 5.45E-12 | 9.71E-08 | -0.66 | 0.633 | 0.908 | Fibroblasts | aPD-L1+aTGF-β vs aPD-L1 |
| Clec3b | 1.71E-10 | 3.05E-06 | -0.67 | 0.676 | 0.897 | Fibroblasts | aPD-L1+aTGF-β vs aPD-L1 |
| Gm42418 | 3.94E-12 | 7.03E-08 | -0.67 | 1 | 1 | NKT | aPD-L1+aTGF-β vs aPD-L1 |
| Fstl1 | 2.73E-12 | 4.87E-08 | -0.68 | 0.941 | 0.954 | Fibroblasts | aPD-L1+aTGF-β vs aPD-L1 |
| Gm42418 | 6.48E-115 | 1.16E-110 | -0.69 | 1 | 1 | T cells | aPD-L1+aTGF-β vs aPD-L1 |
| Marcks | 8.67E-13 | 1.55E-08 | -0.69 | 0.644 | 0.897 | Fibroblasts | aPD-L1+aTGF-β vs aPD-L1 |
| Fn1 | 7.41E-12 | 1.32E-07 | -0.71 | 0.92 | 0.954 | Fibroblasts | aPD-L1+aTGF-β vs aPD-L1 |
| Lars2 | 1.30E-22 | 2.31E-18 | -0.71 | 0.535 | 0.842 | Monocytes | aPD-L1+aTGF-β vs aPD-L1 |
| Gas7 | 2.71E-10 | 4.83E-06 | -0.73 | 0.33 | 0.678 | Fibroblasts | aPD-L1+aTGF-β vs aPD-L1 |
| Timp3 | 6.50E-08 | 1.16E-03 | -0.74 | 0.287 | 0.575 | Fibroblasts | aPD-L1+aTGF-β vs aPD-L1 |
| Fbn1 | 2.61E-11 | 4.66E-07 | -0.76 | 0.782 | 0.92 | Fibroblasts | aPD-L1+aTGF-β vs aPD-L1 |
| Igfbp6 | 6.49E-11 | 1.16E-06 | -0.76 | 0.707 | 0.897 | Fibroblasts | aPD-L1+aTGF-β vs aPD-L1 |
| Gm42418 | 4.56E-35 | 8.14E-31 | -0.97 | 0.993 | 1 | Epithelial cells | aPD-L1+aTGF-β vs aPD-L1 |
| Gm42418 | 6.35E-24 | 1.13E-19 | -1.10 | 1 | 1 | Fibroblasts | aPD-L1+aTGF-β vs aPD-L1 |
| Gm42418 | 1.05E-176 | 1.87E-172 | -1.11 | 0.998 | 1 | Tumor_cells | aPD-L1+aTGF-β vs aPD-L1 |
| AY036118 | 4.87E-178 | 8.67E-174 | -1.16 | 0.866 | 0.988 | Neutrophils | aPD-L1+aTGF-β vs aPD-L1 |
| Igfbp5 | 4.74E-11 | 8.45E-07 | -1.17 | 0.319 | 0.667 | Fibroblasts | aPD-L1+aTGF-β vs aPD-L1 |
| AY036118 | 5.95E-300 | 1.06E-295 | -1.32 | 0.626 | 0.998 | B cells | aPD-L1+aTGF-β vs aPD-L1 |
| AY036118 | 9.76E-87 | 1.74E-82 | -1.34 | 0.652 | 0.989 | Macrophages | aPD-L1+aTGF-β vs aPD-L1 |
| AY036118 | 7.27E-45 | 1.30E-40 | -1.37 | 0.641 | 1 | ILC | aPD-L1+aTGF-β vs aPD-L1 |
| AY036118 | 2.01E-41 | 3.59E-37 | -1.44 | 0.739 | 1 | NKT | aPD-L1+aTGF-β vs aPD-L1 |
| AY036118 | 4.20E-13 | 7.48E-09 | -1.44 | 0.652 | 1 | DC | aPD-L1+aTGF-β vs aPD-L1 |
| AY036118 | 1.46E-213 | 2.61E-209 | -1.51 | 0.625 | 1 | T cells | aPD-L1+aTGF-β vs aPD-L1 |
| AY036118 | 6.71E-11 | 1.20E-06 | -1.55 | 0.698 | 1 | NK cells | aPD-L1+aTGF-β vs aPD-L1 |
| AY036118 | 4.52E-67 | 8.06E-63 | -1.56 | 0.835 | 1 | Monocytes | aPD-L1+aTGF-β vs aPD-L1 |
| Cxcl10 | 4.16E-17 | 7.42E-13 | 1.03 | 0.511 | 0.124 | Fibroblasts | aPD-L1+aTGF-β vs aTGF-β |
| Cxcl9 | 2.35E-08 | 4.19E-04 | 0.71 | 0.324 | 0.099 | Fibroblasts | aPD-L1+aTGF-β vs aTGF-β |
| Igkc | 3.24E-07 | 5.78E-03 | 0.64 | 0.162 | 0.081 | Macrophages | aPD-L1+aTGF-β vs aTGF-β |
| Klf2 | 9.01E-16 | 1.61E-11 | 0.62 | 0.883 | 0.743 | Epithelial cells | aPD-L1+aTGF-β vs aTGF-β |
| H2-Eb1 | 1.69E-18 | 3.01E-14 | 0.57 | 0.709 | 0.559 | Macrophages | aPD-L1+aTGF-β vs aTGF-β |
| Wfdc17 | 6.79E-08 | 1.21E-03 | 0.56 | 0.372 | 0.265 | Macrophages | aPD-L1+aTGF-β vs aTGF-β |
| H2-Aa | 4.52E-17 | 8.06E-13 | 0.55 | 0.712 | 0.588 | Macrophages | aPD-L1+aTGF-β vs aTGF-β |
| Irf1 | 1.18E-21 | 2.10E-17 | 0.54 | 0.806 | 0.562 | Monocytes | aPD-L1+aTGF-β vs aTGF-β |
| H2-Ab1 | 8.14E-17 | 1.45E-12 | 0.53 | 0.728 | 0.613 | Macrophages | aPD-L1+aTGF-β vs aTGF-β |
| Iglc1 | 9.25E-07 | 1.65E-02 | 0.49 | 0.209 | 0.284 | B cells | aPD-L1+aTGF-β vs aTGF-β |
| H2-D1 | 2.95E-12 | 5.26E-08 | 0.48 | 0.995 | 0.985 | Fibroblasts | aPD-L1+aTGF-β vs aTGF-β |
| Mt1 | 2.06E-09 | 3.68E-05 | 0.48 | 0.794 | 0.641 | Epithelial cells | aPD-L1+aTGF-β vs aTGF-β |
| Cxcl1 | 9.25E-82 | 1.65E-77 | 0.48 | 0.797 | 0.639 | Tumor_cells | aPD-L1+aTGF-β vs aTGF-β |
| Atf3 | 3.18E-13 | 5.67E-09 | 0.47 | 0.896 | 0.768 | Epithelial cells | aPD-L1+aTGF-β vs aTGF-β |
| Vps37b | 5.62E-23 | 1.00E-18 | 0.47 | 0.74 | 0.526 | T cells | aPD-L1+aTGF-β vs aTGF-β |
| Rpl37a | 1.72E-23 | 3.07E-19 | 0.47 | 1 | 1 | ILC | aPD-L1+aTGF-β vs aTGF-β |
| Irf1 | 7.17E-11 | 1.28E-06 | 0.47 | 0.437 | 0.221 | Epithelial cells | aPD-L1+aTGF-β vs aTGF-β |
| Cd74 | 1.16E-14 | 2.07E-10 | 0.47 | 0.868 | 0.836 | Macrophages | aPD-L1+aTGF-β vs aTGF-β |
| Vps37b | 6.62E-46 | 1.18E-41 | 0.47 | 0.593 | 0.35 | B cells | aPD-L1+aTGF-β vs aTGF-β |
| Irf1 | 1.23E-38 | 2.20E-34 | 0.46 | 0.548 | 0.257 | Macrophages | aPD-L1+aTGF-β vs aTGF-β |
| Cxcl10 | 1.76E-09 | 3.13E-05 | 0.46 | 0.593 | 0.481 | Macrophages | aPD-L1+aTGF-β vs aTGF-β |
| Dnajb1 | 5.59E-08 | 9.97E-04 | 0.45 | 0.687 | 0.522 | Epithelial cells | aPD-L1+aTGF-β vs aTGF-β |
| Rpl39 | 1.28E-17 | 2.28E-13 | 0.45 | 1 | 1 | ILC | aPD-L1+aTGF-β vs aTGF-β |
| Cxcl2 | 4.64E-42 | 8.26E-38 | 0.44 | 0.476 | 0.329 | Tumor_cells | aPD-L1+aTGF-β vs aTGF-β |
| Dennd4a | 5.38E-59 | 9.59E-55 | 0.44 | 0.779 | 0.528 | B cells | aPD-L1+aTGF-β vs aTGF-β |
| Gm10076 | 4.45E-40 | 7.93E-36 | 0.42 | 0.899 | 0.703 | T cells | aPD-L1+aTGF-β vs aTGF-β |
| Rps21 | 7.21E-21 | 1.29E-16 | 0.42 | 1 | 1 | ILC | aPD-L1+aTGF-β vs aTGF-β |
| Ccl5 | 1.15E-07 | 2.05E-03 | 0.42 | 0.178 | 0.092 | Macrophages | aPD-L1+aTGF-β vs aTGF-β |
| Irf1 | 1.29E-52 | 2.30E-48 | 0.42 | 0.477 | 0.212 | B cells | aPD-L1+aTGF-β vs aTGF-β |

| | | | | | | | |
|---|---|---|---|---|---|---|---|
| Socs1 | 3.19E-08 | 5.69E-04 | 0.41 | 0.356 | 0.124 | Fibroblasts | aPD-L1+aTGF-β vs aTGF-β |
| Dusp10 | 2.52E-16 | 4.50E-12 | 0.41 | 0.572 | 0.371 | T cells | aPD-L1+aTGF-β vs aTGF-β |
| AW112010 | 7.73E-09 | 1.38E-04 | 0.41 | 0.696 | 0.65 | Macrophages | aPD-L1+aTGF-β vs aTGF-β |
| Rps27 | 9.99E-23 | 1.78E-18 | 0.39 | 0.988 | 0.934 | Monocytes | aPD-L1+aTGF-β vs aTGF-β |
| Rel | 4.88E-51 | 8.70E-47 | 0.39 | 0.853 | 0.675 | B cells | aPD-L1+aTGF-β vs aTGF-β |
| Ubc | 9.24E-10 | 1.65E-05 | 0.39 | 0.903 | 0.866 | Epithelial cells | aPD-L1+aTGF-β vs aTGF-β |
| H2-K1 | 3.39E-09 | 6.05E-05 | 0.39 | 0.989 | 0.946 | Fibroblasts | aPD-L1+aTGF-β vs aTGF-β |
| Ly6a | 4.39E-09 | 7.82E-05 | 0.38 | 0.558 | 0.46 | Macrophages | aPD-L1+aTGF-β vs aTGF-β |
| H2-D1 | 4.06E-23 | 7.23E-19 | 0.38 | 1 | 0.993 | Epithelial cells | aPD-L1+aTGF-β vs aTGF-β |
| Vps37b | 2.21E-07 | 3.94E-03 | 0.38 | 0.764 | 0.557 | NKT | aPD-L1+aTGF-β vs aTGF-β |
| Cd274 | 1.32E-10 | 2.35E-06 | 0.38 | 0.666 | 0.478 | Monocytes | aPD-L1+aTGF-β vs aTGF-β |
| Oasl1 | 1.41E-13 | 2.52E-09 | 0.38 | 0.48 | 0.325 | Macrophages | aPD-L1+aTGF-β vs aTGF-β |
| Ifi27l2a | 3.04E-26 | 5.42E-22 | 0.37 | 0.533 | 0.353 | B cells | aPD-L1+aTGF-β vs aTGF-β |
| Kdm6b | 1.25E-46 | 2.23E-42 | 0.37 | 0.599 | 0.342 | B cells | aPD-L1+aTGF-β vs aTGF-β |
| Rps27 | 5.64E-19 | 1.00E-14 | 0.37 | 0.996 | 1 | ILC | aPD-L1+aTGF-β vs aTGF-β |
| Pim1 | 9.91E-34 | 1.77E-29 | 0.37 | 0.543 | 0.33 | B cells | aPD-L1+aTGF-β vs aTGF-β |
| Gm10076 | 1.64E-09 | 2.93E-05 | 0.36 | 0.854 | 0.667 | ILC | aPD-L1+aTGF-β vs aTGF-β |
| Gm10076 | 1.24E-56 | 2.21E-52 | 0.36 | 0.831 | 0.644 | B cells | aPD-L1+aTGF-β vs aTGF-β |
| Crip1 | 3.87E-17 | 6.89E-13 | 0.36 | 0.919 | 0.871 | Macrophages | aPD-L1+aTGF-β vs aTGF-β |
| Tmsb10 | 1.47E-21 | 2.62E-17 | 0.36 | 0.971 | 0.938 | Macrophages | aPD-L1+aTGF-β vs aTGF-β |
| Rpl39 | 1.94E-17 | 3.46E-13 | 0.36 | 1 | 1 | NKT | aPD-L1+aTGF-β vs aTGF-β |
| Irf1 | 7.46E-07 | 1.33E-02 | 0.36 | 0.633 | 0.431 | Fibroblasts | aPD-L1+aTGF-β vs aTGF-β |
| Rpl37 | 1.35E-16 | 2.40E-12 | 0.36 | 1 | 0.972 | ILC | aPD-L1+aTGF-β vs aTGF-β |
| Rpl37 | 1.69E-26 | 3.02E-22 | 0.35 | 0.99 | 0.939 | Monocytes | aPD-L1+aTGF-β vs aTGF-β |
| Nfkbia | 4.45E-84 | 7.94E-80 | 0.35 | 0.919 | 0.847 | Tumor_cells | aPD-L1+aTGF-β vs aTGF-β |
| Rpl38 | 1.60E-13 | 2.85E-09 | 0.35 | 0.996 | 0.993 | ILC | aPD-L1+aTGF-β vs aTGF-β |
| Rpl37a | 4.22E-18 | 7.52E-14 | 0.34 | 1 | 1 | NKT | aPD-L1+aTGF-β vs aTGF-β |
| Rpl39 | 1.46E-18 | 2.60E-14 | 0.34 | 0.978 | 0.946 | Monocytes | aPD-L1+aTGF-β vs aTGF-β |
| Rps21 | 2.70E-16 | 4.81E-12 | 0.34 | 1 | 0.995 | NKT | aPD-L1+aTGF-β vs aTGF-β |
| Rps27 | 3.12E-160 | 5.56E-156 | 0.34 | 0.999 | 1 | B cells | aPD-L1+aTGF-β vs aTGF-β |
| Gm10076 | 4.41E-47 | 7.86E-43 | 0.34 | 0.874 | 0.678 | Macrophages | aPD-L1+aTGF-β vs aTGF-β |
| Satb1 | 4.98E-41 | 8.88E-37 | 0.34 | 0.757 | 0.546 | B cells | aPD-L1+aTGF-β vs aTGF-β |
| Mt2 | 4.55E-69 | 8.11E-65 | 0.34 | 0.763 | 0.568 | Tumor_cells | aPD-L1+aTGF-β vs aTGF-β |
| Zfand5 | 1.96E-79 | 3.49E-75 | 0.34 | 0.712 | 0.523 | Tumor_cells | aPD-L1+aTGF-β vs aTGF-β |
| Ly6a | 1.53E-10 | 2.73E-06 | 0.34 | 0.687 | 0.428 | Epithelial cells | aPD-L1+aTGF-β vs aTGF-β |
| Irf1 | 4.38E-87 | 7.81E-83 | 0.34 | 0.639 | 0.398 | Tumor_cells | aPD-L1+aTGF-β vs aTGF-β |
| Gm10076 | 6.23E-21 | 1.11E-16 | 0.33 | 0.794 | 0.544 | Monocytes | aPD-L1+aTGF-β vs aTGF-β |
| H2-K1 | 2.44E-15 | 4.35E-11 | 0.32 | 0.998 | 0.978 | Epithelial cells | aPD-L1+aTGF-β vs aTGF-β |
| Rsad2 | 8.96E-07 | 1.60E-02 | 0.32 | 0.426 | 0.322 | Macrophages | aPD-L1+aTGF-β vs aTGF-β |
| Rpl37 | 1.81E-14 | 3.22E-10 | 0.32 | 1 | 0.975 | NKT | aPD-L1+aTGF-β vs aTGF-β |
| Iigp1 | 3.65E-07 | 6.51E-03 | 0.32 | 0.263 | 0.101 | Epithelial cells | aPD-L1+aTGF-β vs aTGF-β |
| Il1a | 3.03E-12 | 5.40E-08 | 0.32 | 0.181 | 0.095 | Neutrophils | aPD-L1+aTGF-β vs aTGF-β |
| B2m | 5.27E-13 | 9.40E-09 | 0.32 | 0.993 | 0.975 | Epithelial cells | aPD-L1+aTGF-β vs aTGF-β |
| Serpina3g | 9.02E-19 | 1.61E-14 | 0.32 | 0.308 | 0.136 | Macrophages | aPD-L1+aTGF-β vs aTGF-β |
| Arg2 | 8.30E-10 | 1.48E-05 | 0.31 | 0.603 | 0.44 | Monocytes | aPD-L1+aTGF-β vs aTGF-β |
| Dusp2 | 2.48E-06 | 4.42E-02 | 0.31 | 0.375 | 0.235 | Monocytes | aPD-L1+aTGF-β vs aTGF-β |
| Rpl39 | 2.48E-56 | 4.42E-52 | 0.31 | 1 | 1 | T cells | aPD-L1+aTGF-β vs aTGF-β |
| Gm10076 | 7.66E-09 | 1.37E-04 | 0.31 | 0.793 | 0.622 | NKT | aPD-L1+aTGF-β vs aTGF-β |
| Rps27 | 8.10E-56 | 1.44E-51 | 0.31 | 1 | 1 | T cells | aPD-L1+aTGF-β vs aTGF-β |
| Krt8 | 4.15E-07 | 7.40E-03 | 0.31 | 0.911 | 0.851 | Epithelial cells | aPD-L1+aTGF-β vs aTGF-β |
| Dnaja1 | 3.53E-13 | 6.30E-09 | 0.31 | 0.903 | 0.798 | T cells | aPD-L1+aTGF-β vs aTGF-β |
| Rpl36 | 1.18E-90 | 2.10E-86 | 0.31 | 0.997 | 0.998 | B cells | aPD-L1+aTGF-β vs aTGF-β |
| Klf6 | 4.17E-08 | 7.44E-04 | 0.31 | 0.933 | 0.851 | Epithelial cells | aPD-L1+aTGF-β vs aTGF-β |
| Fcgr4 | 2.93E-09 | 5.22E-05 | 0.31 | 0.593 | 0.504 | Macrophages | aPD-L1+aTGF-β vs aTGF-β |
| Rpl39 | 3.98E-98 | 7.10E-94 | 0.31 | 0.996 | 0.999 | B cells | aPD-L1+aTGF-β vs aTGF-β |

| | | | | | | | |
|---|---|---|---|---|---|---|---|
| Socs1 | 3.82E-12 | 6.81E-08 | 0.31 | 0.453 | 0.228 | Monocytes | aPD-L1+aTGF-β vs aTGF-β |
| Gm10260 | 6.06E-08 | 1.08E-03 | 0.31 | 0.94 | 0.861 | ILC | aPD-L1+aTGF-β vs aTGF-β |
| Plaur | 7.40E-08 | 1.32E-03 | 0.31 | 0.634 | 0.475 | Monocytes | aPD-L1+aTGF-β vs aTGF-β |
| Rpl37a | 3.16E-100 | 5.63E-96 | 0.31 | 0.998 | 0.998 | B cells | aPD-L1+aTGF-β vs aTGF-β |
| Icam1 | 1.06E-10 | 1.89E-06 | 0.30 | 0.443 | 0.247 | Monocytes | aPD-L1+aTGF-β vs aTGF-β |
| Rps21 | 1.32E-49 | 2.36E-45 | 0.30 | 1 | 1 | T cells | aPD-L1+aTGF-β vs aTGF-β |
| Rpl37 | 4.50E-45 | 8.02E-41 | 0.30 | 1 | 0.996 | T cells | aPD-L1+aTGF-β vs aTGF-β |
| Mgp | 1.86E-09 | 3.31E-05 | 0.30 | 0.339 | 0.266 | Tumor_cells | aPD-L1+aTGF-β vs aTGF-β |
| Rpl36 | 3.16E-15 | 5.63E-11 | 0.30 | 0.994 | 1 | NKT | aPD-L1+aTGF-β vs aTGF-β |
| Nr4a1 | 2.94E-35 | 5.24E-31 | 0.30 | 0.695 | 0.451 | B cells | aPD-L1+aTGF-β vs aTGF-β |
| Lmna | 2.69E-07 | 4.79E-03 | 0.30 | 0.886 | 0.793 | Epithelial cells | aPD-L1+aTGF-β vs aTGF-β |
| Cxcl10 | 1.13E-22 | 2.01E-18 | 0.30 | 0.176 | 0.088 | Tumor_cells | aPD-L1+aTGF-β vs aTGF-β |
| Rps21 | 3.58E-102 | 6.37E-98 | 0.30 | 0.998 | 1 | B cells | aPD-L1+aTGF-β vs aTGF-β |
| Rpl34 | 9.27E-10 | 1.65E-05 | 0.29 | 1 | 1 | ILC | aPD-L1+aTGF-β vs aTGF-β |
| Rps27 | 2.09E-10 | 3.72E-06 | 0.29 | 0.997 | 1 | NKT | aPD-L1+aTGF-β vs aTGF-β |
| Ifi27l2a | 3.57E-08 | 6.37E-04 | 0.29 | 0.65 | 0.524 | T cells | aPD-L1+aTGF-β vs aTGF-β |
| Socs3 | 3.18E-08 | 5.67E-04 | 0.29 | 0.765 | 0.619 | Monocytes | aPD-L1+aTGF-β vs aTGF-β |
| Rpl37a | 1.24E-43 | 2.21E-39 | 0.29 | 1 | 1 | T cells | aPD-L1+aTGF-β vs aTGF-β |
| Rpl37a | 3.61E-18 | 6.43E-14 | 0.29 | 0.99 | 0.984 | Monocytes | aPD-L1+aTGF-β vs aTGF-β |
| Rps27 | 6.90E-32 | 1.23E-27 | 0.29 | 0.987 | 0.972 | Macrophages | aPD-L1+aTGF-β vs aTGF-β |
| Crem | 1.84E-40 | 3.28E-36 | 0.29 | 0.323 | 0.115 | B cells | aPD-L1+aTGF-β vs aTGF-β |
| Hsp90aa1 | 5.57E-07 | 9.92E-03 | 0.29 | 0.973 | 0.964 | Epithelial cells | aPD-L1+aTGF-β vs aTGF-β |
| Rpl27 | 1.55E-06 | 2.77E-02 | 0.29 | 0.794 | 0.688 | ILC | aPD-L1+aTGF-β vs aTGF-β |
| Rps15a | 5.42E-09 | 9.65E-05 | 0.29 | 1 | 1 | ILC | aPD-L1+aTGF-β vs aTGF-β |
| Rpl39 | 3.41E-38 | 6.08E-34 | 0.29 | 0.991 | 0.991 | Macrophages | aPD-L1+aTGF-β vs aTGF-β |
| Rpl36 | 1.60E-15 | 2.85E-11 | 0.28 | 0.983 | 0.918 | Monocytes | aPD-L1+aTGF-β vs aTGF-β |
| Gadd45b | 2.56E-07 | 4.57E-03 | 0.28 | 0.396 | 0.287 | Macrophages | aPD-L1+aTGF-β vs aTGF-β |
| Rpl37 | 3.51E-81 | 6.25E-77 | 0.28 | 0.999 | 1 | B cells | aPD-L1+aTGF-β vs aTGF-β |
| Ly6a | 2.87E-23 | 5.12E-19 | 0.28 | 0.166 | 0.049 | B cells | aPD-L1+aTGF-β vs aTGF-β |
| Spp1 | 1.23E-23 | 2.19E-19 | 0.28 | 0.989 | 0.979 | Tumor_cells | aPD-L1+aTGF-β vs aTGF-β |
| Rpl36 | 2.42E-09 | 4.32E-05 | 0.28 | 1 | 1 | ILC | aPD-L1+aTGF-β vs aTGF-β |
| Uba52 | 1.66E-07 | 2.96E-03 | 0.28 | 0.591 | 0.326 | ILC | aPD-L1+aTGF-β vs aTGF-β |
| Psmb9 | 4.94E-08 | 8.80E-04 | 0.28 | 0.618 | 0.428 | Epithelial cells | aPD-L1+aTGF-β vs aTGF-β |
| Ccr7 | 4.64E-10 | 8.27E-06 | 0.28 | 0.69 | 0.544 | T cells | aPD-L1+aTGF-β vs aTGF-β |
| Irf1 | 2.64E-18 | 4.71E-14 | 0.28 | 0.686 | 0.563 | Neutrophils | aPD-L1+aTGF-β vs aTGF-β |
| Rpl36 | 3.52E-40 | 6.27E-36 | 0.28 | 1 | 0.998 | T cells | aPD-L1+aTGF-β vs aTGF-β |
| Plac8 | 1.21E-14 | 2.16E-10 | 0.28 | 0.338 | 0.214 | B cells | aPD-L1+aTGF-β vs aTGF-β |
| Gramd3 | 2.85E-14 | 5.09E-10 | 0.27 | 0.467 | 0.27 | T cells | aPD-L1+aTGF-β vs aTGF-β |
| Irf1 | 6.83E-12 | 1.22E-07 | 0.27 | 0.612 | 0.443 | T cells | aPD-L1+aTGF-β vs aTGF-β |
| Uba52 | 1.60E-07 | 2.84E-03 | 0.27 | 0.529 | 0.299 | NKT | aPD-L1+aTGF-β vs aTGF-β |
| Rps29 | 3.93E-11 | 7.01E-07 | 0.27 | 1 | 1 | ILC | aPD-L1+aTGF-β vs aTGF-β |
| Nfkbia | 1.50E-07 | 2.67E-03 | 0.27 | 0.944 | 0.868 | Monocytes | aPD-L1+aTGF-β vs aTGF-β |
| Rps21 | 5.22E-15 | 9.30E-11 | 0.27 | 0.961 | 0.953 | Monocytes | aPD-L1+aTGF-β vs aTGF-β |
| Sub1 | 8.72E-28 | 1.55E-23 | 0.27 | 0.824 | 0.672 | B cells | aPD-L1+aTGF-β vs aTGF-β |
| Gbp2b | 3.47E-13 | 6.18E-09 | 0.27 | 0.369 | 0.215 | Macrophages | aPD-L1+aTGF-β vs aTGF-β |
| H2-DMb1 | 1.18E-11 | 2.10E-07 | 0.27 | 0.515 | 0.374 | Macrophages | aPD-L1+aTGF-β vs aTGF-β |
| Gm10260 | 1.31E-08 | 2.34E-04 | 0.27 | 0.705 | 0.614 | Monocytes | aPD-L1+aTGF-β vs aTGF-β |
| Nfe2l2 | 7.67E-07 | 1.37E-02 | 0.27 | 0.554 | 0.402 | Monocytes | aPD-L1+aTGF-β vs aTGF-β |
| Cytip | 1.57E-28 | 2.81E-24 | 0.26 | 0.749 | 0.562 | B cells | aPD-L1+aTGF-β vs aTGF-β |
| Jun | 3.37E-08 | 6.01E-04 | 0.26 | 0.998 | 0.975 | Epithelial cells | aPD-L1+aTGF-β vs aTGF-β |
| Cd83 | 3.59E-07 | 6.39E-03 | 0.26 | 0.571 | 0.419 | Monocytes | aPD-L1+aTGF-β vs aTGF-β |
| Rel | 4.09E-08 | 7.30E-04 | 0.26 | 0.525 | 0.355 | Monocytes | aPD-L1+aTGF-β vs aTGF-β |
| Rpl38 | 1.00E-57 | 1.79E-53 | 0.26 | 0.997 | 0.986 | B cells | aPD-L1+aTGF-β vs aTGF-β |
| Gm10260 | 3.42E-23 | 6.09E-19 | 0.26 | 0.98 | 0.959 | T cells | aPD-L1+aTGF-β vs aTGF-β |
| Hal | 1.82E-10 | 3.24E-06 | 0.26 | 0.289 | 0.172 | Macrophages | aPD-L1+aTGF-β vs aTGF-β |

| | | | | | | | |
|---|---|---|---|---|---|---|---|
| Rpl34 | 4.42E-13 | 7.87E-09 | 0.26 | 0.985 | 0.967 | Monocytes | aPD-L1+aTGF-β vs aTGF-β |
| Tnfaip2 | 7.79E-08 | 1.39E-03 | 0.26 | 0.438 | 0.264 | Monocytes | aPD-L1+aTGF-β vs aTGF-β |
| Cd44 | 2.38E-08 | 4.25E-04 | 0.25 | 0.78 | 0.614 | Monocytes | aPD-L1+aTGF-β vs aTGF-β |
| Rpl35 | 1.04E-25 | 1.85E-21 | 0.25 | 0.997 | 0.981 | T cells | aPD-L1+aTGF-β vs aTGF-β |
| Rps28 | 1.33E-06 | 2.37E-02 | 0.25 | 0.951 | 0.866 | NKT | aPD-L1+aTGF-β vs aTGF-β |
| Tmsb10 | 3.00E-07 | 5.35E-03 | 0.25 | 1 | 1 | NKT | aPD-L1+aTGF-β vs aTGF-β |
| Fos | 1.04E-07 | 1.85E-03 | 0.25 | 0.995 | 0.975 | Epithelial cells | aPD-L1+aTGF-β vs aTGF-β |
| Rpl41 | 6.85E-08 | 1.22E-03 | 0.25 | 1 | 1 | ILC | aPD-L1+aTGF-β vs aTGF-β |
| Gadd45g | 7.49E-26 | 1.33E-21 | -0.25 | 0.32 | 0.441 | Tumor_cells | aPD-L1+aTGF-β vs aTGF-β |
| Trim30a | 2.11E-13 | 3.75E-09 | -0.25 | 0.475 | 0.586 | Neutrophils | aPD-L1+aTGF-β vs aTGF-β |
| Ly6e | 2.27E-10 | 4.04E-06 | -0.26 | 0.664 | 0.723 | Neutrophils | aPD-L1+aTGF-β vs aTGF-β |
| Ppp1r14b | 1.68E-07 | 3.00E-03 | -0.26 | 0.238 | 0.486 | ILC | aPD-L1+aTGF-β vs aTGF-β |
| Cryab | 1.84E-11 | 3.27E-07 | -0.26 | 0.443 | 0.505 | Tumor_cells | aPD-L1+aTGF-β vs aTGF-β |
| Slfn4 | 1.83E-11 | 3.26E-07 | -0.26 | 0.846 | 0.853 | Neutrophils | aPD-L1+aTGF-β vs aTGF-β |
| Il4i1 | 3.11E-21 | 5.55E-17 | -0.26 | 0.13 | 0.259 | B cells | aPD-L1+aTGF-β vs aTGF-β |
| Iglc2 | 6.11E-32 | 1.09E-27 | -0.26 | 0.768 | 0.876 | B cells | aPD-L1+aTGF-β vs aTGF-β |
| Tspo | 3.42E-19 | 6.11E-15 | -0.27 | 0.875 | 0.925 | Neutrophils | aPD-L1+aTGF-β vs aTGF-β |
| Tmx4 | 1.32E-10 | 2.35E-06 | -0.27 | 0.244 | 0.376 | Macrophages | aPD-L1+aTGF-β vs aTGF-β |
| Fos | 2.48E-08 | 4.42E-04 | -0.28 | 0.927 | 0.965 | Monocytes | aPD-L1+aTGF-β vs aTGF-β |
| Cdkn1a | 1.77E-16 | 3.15E-12 | -0.28 | 0.778 | 0.836 | Neutrophils | aPD-L1+aTGF-β vs aTGF-β |
| Slfn5 | 7.02E-11 | 1.25E-06 | -0.28 | 0.38 | 0.474 | Neutrophils | aPD-L1+aTGF-β vs aTGF-β |
| Pfn1 | 3.93E-42 | 7.01E-38 | -0.28 | 0.935 | 0.98 | B cells | aPD-L1+aTGF-β vs aTGF-β |
| Hspa1b | 2.78E-10 | 4.96E-06 | -0.29 | 0.624 | 0.757 | T cells | aPD-L1+aTGF-β vs aTGF-β |
| H3f3b | 1.17E-08 | 2.09E-04 | -0.29 | 1 | 1 | Monocytes | aPD-L1+aTGF-β vs aTGF-β |
| Dnajb1 | 1.67E-40 | 2.97E-36 | -0.29 | 0.69 | 0.803 | Tumor_cells | aPD-L1+aTGF-β vs aTGF-β |
| Slfn5 | 7.82E-08 | 1.39E-03 | -0.29 | 0.433 | 0.572 | Monocytes | aPD-L1+aTGF-β vs aTGF-β |
| Mx1 | 1.36E-07 | 2.42E-03 | -0.30 | 0.24 | 0.395 | Monocytes | aPD-L1+aTGF-β vs aTGF-β |
| Usp18 | 3.01E-19 | 5.36E-15 | -0.30 | 0.176 | 0.318 | Neutrophils | aPD-L1+aTGF-β vs aTGF-β |
| Actb | 2.71E-24 | 4.83E-20 | -0.30 | 1 | 1 | T cells | aPD-L1+aTGF-β vs aTGF-β |
| Hspa1b | 2.56E-54 | 4.57E-50 | -0.30 | 0.667 | 0.818 | Tumor_cells | aPD-L1+aTGF-β vs aTGF-β |
| Fos | 1.36E-18 | 2.42E-14 | -0.31 | 0.989 | 0.993 | Neutrophils | aPD-L1+aTGF-β vs aTGF-β |
| Jun | 3.62E-24 | 6.44E-20 | -0.31 | 0.922 | 0.954 | B cells | aPD-L1+aTGF-β vs aTGF-β |
| mt-Co2 | 1.84E-08 | 3.28E-04 | -0.31 | 0.995 | 1 | Fibroblasts | aPD-L1+aTGF-β vs aTGF-β |
| Itm2b | 3.01E-08 | 5.37E-04 | -0.32 | 0.96 | 0.986 | Epithelial cells | aPD-L1+aTGF-β vs aTGF-β |
| Ifi44 | 2.88E-07 | 5.13E-03 | -0.32 | 0.533 | 0.631 | Monocytes | aPD-L1+aTGF-β vs aTGF-β |
| Iglc3 | 6.56E-34 | 1.17E-29 | -0.33 | 0.535 | 0.696 | B cells | aPD-L1+aTGF-β vs aTGF-β |
| Cyr61 | 4.40E-33 | 7.84E-29 | -0.33 | 0.492 | 0.605 | Tumor_cells | aPD-L1+aTGF-β vs aTGF-β |
| Oasl1 | 8.34E-19 | 1.49E-14 | -0.33 | 0.395 | 0.537 | Neutrophils | aPD-L1+aTGF-β vs aTGF-β |
| Klf2 | 2.05E-17 | 3.65E-13 | -0.33 | 0.967 | 0.979 | Neutrophils | aPD-L1+aTGF-β vs aTGF-β |
| Ifit3 | 7.79E-08 | 1.39E-03 | -0.34 | 0.194 | 0.348 | Monocytes | aPD-L1+aTGF-β vs aTGF-β |
| mt-Co3 | 2.37E-09 | 4.23E-05 | -0.34 | 0.995 | 0.995 | Fibroblasts | aPD-L1+aTGF-β vs aTGF-β |
| Isg20 | 6.04E-08 | 1.08E-03 | -0.34 | 0.206 | 0.36 | Monocytes | aPD-L1+aTGF-β vs aTGF-β |
| Gm42418 | 2.13E-07 | 3.79E-03 | -0.34 | 1 | 1 | Macrophages | aPD-L1+aTGF-β vs aTGF-β |
| Cmpk2 | 1.56E-16 | 2.78E-12 | -0.35 | 0.179 | 0.303 | Neutrophils | aPD-L1+aTGF-β vs aTGF-β |
| Hspb1 | 1.00E-52 | 1.78E-48 | -0.35 | 0.878 | 0.924 | Tumor_cells | aPD-L1+aTGF-β vs aTGF-β |
| AY036118 | 9.84E-08 | 1.75E-03 | -0.35 | 0.835 | 0.861 | Monocytes | aPD-L1+aTGF-β vs aTGF-β |
| Actb | 6.49E-10 | 1.16E-05 | -0.35 | 1 | 1 | ILC | aPD-L1+aTGF-β vs aTGF-β |
| AY036118 | 8.02E-07 | 1.43E-02 | -0.35 | 0.648 | 0.779 | Epithelial cells | aPD-L1+aTGF-β vs aTGF-β |
| Lrg1 | 8.76E-07 | 1.56E-02 | -0.36 | 0.166 | 0.319 | Epithelial cells | aPD-L1+aTGF-β vs aTGF-β |
| Cd79b | 8.62E-46 | 1.54E-41 | -0.36 | 0.755 | 0.861 | B cells | aPD-L1+aTGF-β vs aTGF-β |
| Actb | 6.98E-84 | 1.24E-79 | -0.37 | 0.996 | 1 | B cells | aPD-L1+aTGF-β vs aTGF-β |
| Ifit3b | 1.23E-15 | 2.19E-11 | -0.37 | 0.186 | 0.308 | Neutrophils | aPD-L1+aTGF-β vs aTGF-β |
| Ltb | 1.95E-51 | 3.48E-47 | -0.38 | 0.603 | 0.792 | B cells | aPD-L1+aTGF-β vs aTGF-β |
| Isg20 | 4.35E-13 | 7.75E-09 | -0.39 | 0.546 | 0.618 | Neutrophils | aPD-L1+aTGF-β vs aTGF-β |
| Ifit2 | 1.69E-06 | 3.01E-02 | -0.40 | 0.235 | 0.369 | Monocytes | aPD-L1+aTGF-β vs aTGF-β |

| | | | | | | | |
|---|---|---|---|---|---|---|---|
| Ifit3 | 8.08E-18 | 1.44E-13 | -0.43 | 0.191 | 0.326 | Neutrophils | aPD-L1+aTGF-β vs aTGF-β |
| Bst2 | 5.91E-11 | 1.05E-06 | -0.44 | 0.634 | 0.767 | Monocytes | aPD-L1+aTGF-β vs aTGF-β |
| Gm42418 | 1.43E-09 | 2.55E-05 | -0.45 | 1 | 1 | Monocytes | aPD-L1+aTGF-β vs aTGF-β |
| Hspa1a | 1.18E-30 | 2.10E-26 | -0.48 | 0.383 | 0.576 | B cells | aPD-L1+aTGF-β vs aTGF-β |
| Irf7 | 1.20E-08 | 2.14E-04 | -0.50 | 0.743 | 0.816 | Monocytes | aPD-L1+aTGF-β vs aTGF-β |
| Isg15 | 1.59E-06 | 2.84E-02 | -0.51 | 0.76 | 0.814 | Monocytes | aPD-L1+aTGF-β vs aTGF-β |
| Slfn4 | 6.71E-10 | 1.20E-05 | -0.53 | 0.361 | 0.529 | Monocytes | aPD-L1+aTGF-β vs aTGF-β |
| Ifit2 | 3.45E-21 | 6.15E-17 | -0.54 | 0.201 | 0.349 | Neutrophils | aPD-L1+aTGF-β vs aTGF-β |
| Rsad2 | 4.70E-14 | 8.38E-10 | -0.54 | 0.665 | 0.716 | Neutrophils | aPD-L1+aTGF-β vs aTGF-β |
| Hspa1b | 6.26E-49 | 1.12E-44 | -0.59 | 0.408 | 0.653 | B cells | aPD-L1+aTGF-β vs aTGF-β |
| Ifit1 | 6.02E-23 | 1.07E-18 | -0.62 | 0.38 | 0.525 | Neutrophils | aPD-L1+aTGF-β vs aTGF-β |
| Apoe | 1.50E-07 | 2.68E-03 | -0.77 | 0.161 | 0.337 | Epithelial cells | aPD-L1+aTGF-β vs aTGF-β |
| Apoe | 1.69E-14 | 3.02E-10 | -0.78 | 0.542 | 0.725 | Monocytes | aPD-L1+aTGF-β vs aTGF-β |
| Cxcl10 | 7.55E-14 | 1.35E-09 | -0.81 | 0.172 | 0.282 | Neutrophils | aPD-L1+aTGF-β vs aTGF-β |
