## Additional file 2, Table S2 for "Single cell transcriptomics reveals the effect of PD-L1/TGF-β blockade on the tumor microenvironment"

Table S2. ***Enriched terms as determined by functional enrichment analysis using DE genes from the EMT6 single cell RNA-seq experiment.*** The functional enrichment analysis was performed using all upregulated or downregulated genes (indicated in the “genes_used” column) from the CD45+ or CD45- cells (indicated in the “cells” column), for a given treatment group comparison (indicated in the “comparison” column). The categories tested were biological process (BP), molecular function (MF), and cellular component (CC), as shown in the “ONTOLOGY” column. “ID” indicates gene ontology ID; “pvalue” indicates the significance of enrichment, while “p.adjust” shows adjusted p-values using the Benjamini–Hochberg method; “gene_count” indicates the number of genes in the tested gene lists that are members of the ontology term, and the genes are listed in the “geneID” column.

| ONTOLOGY | ID | Description | pvalue | p.adjust | geneID | gene_count | comparison | genes_used | cells |
|---|---|---|---|---|---|---|---|---|---|
| BP | GO:0009617 | response to bacterium | 7.20E-21 | 2.23E-17 | Rpl39/Cxcl9/Ly6a | 39 | aPD-L1 vs PBS | Up-regulated genes | CD45+ |
| BP | GO:0002237 | response to molecule of bacterial origin | 1.95E-18 | 3.01E-15 | Cxcl9/Cd274/Tnf/ | 28 | aPD-L1 vs PBS | Up-regulated genes | CD45+ |
| BP | GO:0032496 | response to lipopolysaccharide | 3.25E-18 | 3.35E-15 | Cxcl9/Cd274/Tnf/ | 27 | aPD-L1 vs PBS | Up-regulated genes | CD45+ |
| BP | GO:0071222 | cellular response to lipopolysaccharide | 3.32E-15 | 2.57E-12 | Cxcl9/Cd274/Tnf/ | 21 | aPD-L1 vs PBS | Up-regulated genes | CD45+ |
| BP | GO:0071219 | cellular response to molecule of bacterial origin | 7.99E-15 | 4.94E-12 | Cxcl9/Cd274/Tnf/ | 21 | aPD-L1 vs PBS | Up-regulated genes | CD45+ |
| BP | GO:0002181 | cytoplasmic translation | 7.14E-14 | 3.68E-11 | Rpl39/Rps21/Rpl | 15 | aPD-L1 vs PBS | Up-regulated genes | CD45+ |
| BP | GO:0071216 | cellular response to biotic stimulus | 8.78E-14 | 3.88E-11 | Cxcl9/Cd274/Tnf/ | 21 | aPD-L1 vs PBS | Up-regulated genes | CD45+ |
| BP | GO:0002819 | regulation of adaptive immune response | 2.55E-13 | 9.78E-11 | Cd74/Irf1/H2-Ab1 | 20 | aPD-L1 vs PBS | Up-regulated genes | CD45+ |
| BP | GO:0001819 | positive regulation of cytokine production | 2.84E-13 | 9.78E-11 | Cd74/Irf1/Hilpda/ | 28 | aPD-L1 vs PBS | Up-regulated genes | CD45+ |
| BP | GO:0002821 | positive regulation of adaptive immune response | 5.87E-12 | 1.82E-09 | Cd74/Irf1/H2-Ab1 | 16 | aPD-L1 vs PBS | Up-regulated genes | CD45+ |
| BP | GO:0034341 | response to interferon-gamma | 8.64E-12 | 2.43E-09 | H2-Eb1/H2-Aa/Irf | 16 | aPD-L1 vs PBS | Up-regulated genes | CD45+ |
| BP | GO:0002824 | positive regulation of adaptive immune response | 2.95E-11 | 7.16E-09 | Irf1/H2-Ab1/Cd27 | 15 | aPD-L1 vs PBS | Up-regulated genes | CD45+ |
| BP | GO:0098542 | defense response to other organism | 3.01E-11 | 7.16E-09 | Rpl39/Irf1/Cxcl9/T | 25 | aPD-L1 vs PBS | Up-regulated genes | CD45+ |
| BP | GO:0071396 | cellular response to lipid | 4.02E-11 | 8.87E-09 | Cxcl9/Dnaja1/Inh | 25 | aPD-L1 vs PBS | Up-regulated genes | CD45+ |
| BP | GO:0007159 | leukocyte cell-cell adhesion | 4.80E-11 | 9.85E-09 | H2-Aa/Cd74/Irf1/ | 22 | aPD-L1 vs PBS | Up-regulated genes | CD45+ |
| BP | GO:0002822 | regulation of adaptive immune response based | 5.10E-11 | 9.85E-09 | Irf1/H2-Ab1/Cd27 | 17 | aPD-L1 vs PBS | Up-regulated genes | CD45+ |
| BP | GO:0002697 | regulation of immune effector process | 7.22E-11 | 1.31E-08 | Cd74/Irf1/Tnf/Rsa | 24 | aPD-L1 vs PBS | Up-regulated genes | CD45+ |
| BP | GO:0048002 | antigen processing and presentation of peptide | 7.62E-11 | 1.31E-08 | H2-Eb1/H2-Aa/C | 11 | aPD-L1 vs PBS | Up-regulated genes | CD45+ |
| BP | GO:0002250 | adaptive immune response | 8.11E-11 | 1.32E-08 | Cd74/Irf1/H2-Ab1 | 24 | aPD-L1 vs PBS | Up-regulated genes | CD45+ |
| BP | GO:0019882 | antigen processing and presentation | 1.41E-10 | 2.18E-08 | H2-Eb1/H2-Aa/C | 13 | aPD-L1 vs PBS | Up-regulated genes | CD45+ |
| BP | GO:0022407 | regulation of cell-cell adhesion | 1.83E-10 | 2.70E-08 | H2-Aa/Cd74/Irf1/ | 23 | aPD-L1 vs PBS | Up-regulated genes | CD45+ |
| BP | GO:0002460 | adaptive immune response based on somatic re | 3.41E-10 | 4.80E-08 | Cd74/Irf1/H2-Ab1 | 20 | aPD-L1 vs PBS | Up-regulated genes | CD45+ |
| BP | GO:1903037 | regulation of leukocyte cell-cell adhesion | 3.66E-10 | 4.92E-08 | H2-Aa/Cd74/Irf1/ | 20 | aPD-L1 vs PBS | Up-regulated genes | CD45+ |
| BP | GO:0002478 | antigen processing and presentation of exogen | 5.86E-10 | 7.56E-08 | H2-Eb1/H2-Aa/C | 8 | aPD-L1 vs PBS | Up-regulated genes | CD45+ |
| BP | GO:0071346 | cellular response to interferon-gamma | 1.38E-09 | 1.70E-07 | Irf1/H2-Ab1/Irgm1 | 13 | aPD-L1 vs PBS | Up-regulated genes | CD45+ |
| BP | GO:0002711 | positive regulation of T cell mediated immunity | 2.28E-09 | 2.72E-07 | Rsad2/H2-T23/B | 10 | aPD-L1 vs PBS | Up-regulated genes | CD45+ |
| BP | GO:0014070 | response to organic cyclic compound | 2.66E-09 | 3.05E-07 | Dnaja1/Inhba/Tnf | 25 | aPD-L1 vs PBS | Up-regulated genes | CD45+ |
| BP | GO:0019884 | antigen processing and presentation of exogen | 3.23E-09 | 3.57E-07 | H2-Eb1/H2-Aa/C | 8 | aPD-L1 vs PBS | Up-regulated genes | CD45+ |
| BP | GO:0042742 | defense response to bacterium | 3.53E-09 | 3.77E-07 | Rpl39/Tnf/Arg2/R | 15 | aPD-L1 vs PBS | Up-regulated genes | CD45+ |
| BP | GO:0002456 | T cell mediated immunity | 3.89E-09 | 4.01E-07 | Rsad2/H2-T23/B | 13 | aPD-L1 vs PBS | Up-regulated genes | CD45+ |
| BP | GO:0002699 | positive regulation of immune effector process | 4.14E-09 | 4.13E-07 | Cd74/Irf1/Tnf/Rsa | 17 | aPD-L1 vs PBS | Up-regulated genes | CD45+ |
| BP | GO:1903039 | positive regulation of leukocyte cell-cell adhesio | 4.40E-09 | 4.26E-07 | H2-Aa/Cd74/Irf1/ | 16 | aPD-L1 vs PBS | Up-regulated genes | CD45+ |
| BP | GO:0002718 | regulation of cytokine production involved in imm | 4.72E-09 | 4.31E-07 | Cd74/Tnf/Rsad2/ | 11 | aPD-L1 vs PBS | Up-regulated genes | CD45+ |
| BP | GO:0002367 | cytokine production involved in immune respons | 4.73E-09 | 4.31E-07 | Cd74/Tnf/Rsad2/ | 12 | aPD-L1 vs PBS | Up-regulated genes | CD45+ |
| BP | GO:0022409 | positive regulation of cell-cell adhesion | 5.12E-09 | 4.53E-07 | H2-Aa/Cd74/Irf1/ | 17 | aPD-L1 vs PBS | Up-regulated genes | CD45+ |
| BP | GO:0002440 | production of molecular mediator of immune res | 5.99E-09 | 5.15E-07 | Cd74/H2-Ab1/Tnf | 16 | aPD-L1 vs PBS | Up-regulated genes | CD45+ |
| BP | GO:0002709 | regulation of T cell mediated immunity | 6.29E-09 | 5.26E-07 | Rsad2/H2-T23/B | 11 | aPD-L1 vs PBS | Up-regulated genes | CD45+ |
| BP | GO:0042110 | T cell activation | 1.50E-08 | 1.22E-06 | H2-Aa/Cd74/Irf1/ | 23 | aPD-L1 vs PBS | Up-regulated genes | CD45+ |
| BP | GO:0002720 | positive regulation of cytokine production involve | 1.60E-08 | 1.27E-06 | Cd74/Rsad2/H2- | 9 | aPD-L1 vs PBS | Up-regulated genes | CD45+ |
| BP | GO:1903706 | regulation of hemopoiesis | 1.94E-08 | 1.50E-06 | H2-Aa/Cd74/Irf1/ | 21 | aPD-L1 vs PBS | Up-regulated genes | CD45+ |
| BP | GO:1903428 | positive regulation of reactive oxygen species bi | 2.77E-08 | 2.09E-06 | Tnf/Cd36/Sod2/Ic | 9 | aPD-L1 vs PBS | Up-regulated genes | CD45+ |
| BP | GO:0002449 | lymphocyte mediated immunity | 3.19E-08 | 2.34E-06 | Cd74/H2-Ab1/Tnf | 17 | aPD-L1 vs PBS | Up-regulated genes | CD45+ |
| BP | GO:0019221 | cytokine-mediated signaling pathway | 3.26E-08 | 2.34E-06 | Cd74/Irf1/Cxcl9/Ir | 18 | aPD-L1 vs PBS | Up-regulated genes | CD45+ |
| BP | GO:1903708 | positive regulation of hemopoiesis | 4.35E-08 | 3.06E-06 | H2-Aa/Cd74/Irf1/ | 15 | aPD-L1 vs PBS | Up-regulated genes | CD45+ |
| BP | GO:0002443 | leukocyte mediated immunity | 4.70E-08 | 3.23E-06 | Cd74/H2-Ab1/Tnf | 19 | aPD-L1 vs PBS | Up-regulated genes | CD45+ |
| BP | GO:0045429 | positive regulation of nitric oxide biosynthetic pr | 6.26E-08 | 4.21E-06 | Tnf/Cd36/Sod2/Ic | 8 | aPD-L1 vs PBS | Up-regulated genes | CD45+ |
| BP | GO:0045785 | positive regulation of cell adhesion | 7.34E-08 | 4.83E-06 | H2-Aa/Cd74/Irf1/ | 20 | aPD-L1 vs PBS | Up-regulated genes | CD45+ |
| BP | GO:1904407 | positive regulation of nitric oxide metabolic proc | 7.65E-08 | 4.93E-06 | Tnf/Cd36/Sod2/Ic | 8 | aPD-L1 vs PBS | Up-regulated genes | CD45+ |
| BP | GO:0045088 | regulation of innate immune response | 8.11E-08 | 5.12E-06 | Cd74/Irf1/Irgm1/T | 17 | aPD-L1 vs PBS | Up-regulated genes | CD45+ |
| BP | GO:0001818 | negative regulation of cytokine production | 1.13E-07 | 7.02E-06 | Cd274/Tnf/Arg2/ | 16 | aPD-L1 vs PBS | Up-regulated genes | CD45+ |
| BP | GO:0044419 | interspecies interaction between organisms | 1.25E-07 | 7.45E-06 | Cd74/Irgm1/Tnf/R | 19 | aPD-L1 vs PBS | Up-regulated genes | CD45+ |
| BP | GO:1903409 | reactive oxygen species biosynthetic process | 1.25E-07 | 7.45E-06 | Tnf/Arg2/Cd36/S | 11 | aPD-L1 vs PBS | Up-regulated genes | CD45+ |
| BP | GO:0097530 | granulocyte migration | 1.45E-07 | 8.45E-06 | Cd74/Cxcl9/Ccl4/ | 12 | aPD-L1 vs PBS | Up-regulated genes | CD45+ |
| BP | GO:1990266 | neutrophil migration | 1.53E-07 | 8.77E-06 | Cd74/Cxcl9/Ccl4/ | 11 | aPD-L1 vs PBS | Up-regulated genes | CD45+ |
| BP | GO:0001660 | fever generation | 1.85E-07 | 1.03E-05 | Tnf/Ccl5/Il1a/Il1rn | 5 | aPD-L1 vs PBS | Up-regulated genes | CD45+ |
| BP | GO:0002700 | regulation of production of molecular mediator o | 1.87E-07 | 1.03E-05 | Cd74/Tnf/Rsad2/ | 12 | aPD-L1 vs PBS | Up-regulated genes | CD45+ |
| BP | GO:0071621 | granulocyte chemotaxis | 2.05E-07 | 1.11E-05 | Cd74/Cxcl9/Ccl4/ | 11 | aPD-L1 vs PBS | Up-regulated genes | CD45+ |
| BP | GO:0031663 | lipopolysaccharide-mediated signaling pathway | 2.28E-07 | 1.22E-05 | Tnf/Acod1/Tnfaip | 8 | aPD-L1 vs PBS | Up-regulated genes | CD45+ |
| BP | GO:0030593 | neutrophil chemotaxis | 2.60E-07 | 1.36E-05 | Cd74/Cxcl9/Ccl4/ | 10 | aPD-L1 vs PBS | Up-regulated genes | CD45+ |
| BP | GO:0006809 | nitric oxide biosynthetic process | 2.70E-07 | 1.39E-05 | Tnf/Cd36/Sod2/Ic | 9 | aPD-L1 vs PBS | Up-regulated genes | CD45+ |
| BP | GO:0050777 | negative regulation of immune response | 3.07E-07 | 1.56E-05 | Irf1/Tnf/Samsn1/A | 12 | aPD-L1 vs PBS | Up-regulated genes | CD45+ |
| BP | GO:1903426 | regulation of reactive oxygen species biosynthet | 3.23E-07 | 1.61E-05 | Tnf/Arg2/Cd36/S | 10 | aPD-L1 vs PBS | Up-regulated genes | CD45+ |
| BP | GO:0002708 | positive regulation of lymphocyte mediated imm | 3.59E-07 | 1.76E-05 | Tnf/Rsad2/H2-T2 | 11 | aPD-L1 vs PBS | Up-regulated genes | CD45+ |
| BP | GO:0046209 | nitric oxide metabolic process | 3.97E-07 | 1.92E-05 | Tnf/Cd36/Sod2/Ic | 9 | aPD-L1 vs PBS | Up-regulated genes | CD45+ |
| BP | GO:0048660 | regulation of smooth muscle cell proliferation | 4.21E-07 | 1.99E-05 | Tnf/Mdm2/Tnfaip | 12 | aPD-L1 vs PBS | Up-regulated genes | CD45+ |
| BP | GO:0031349 | positive regulation of defense response | 4.24E-07 | 1.99E-05 | Cd74/Irf1/Irgm1/T | 18 | aPD-L1 vs PBS | Up-regulated genes | CD45+ |
| BP | GO:0002705 | positive regulation of leukocyte mediated immun | 4.55E-07 | 2.10E-05 | Tnf/Rsad2/H2-T2 | 12 | aPD-L1 vs PBS | Up-regulated genes | CD45+ |
| BP | GO:2000379 | positive regulation of reactive oxygen species m | 4.89E-07 | 2.23E-05 | Tnf/Acod1/Cd36/ | 10 | aPD-L1 vs PBS | Up-regulated genes | CD45+ |
| BP | GO:0019886 | antigen processing and presentation of exogen | 5.05E-07 | 2.26E-05 | H2-Eb1/H2-Aa/C | 5 | aPD-L1 vs PBS | Up-regulated genes | CD45+ |
| BP | GO:0048659 | smooth muscle cell proliferation | 5.30E-07 | 2.34E-05 | Tnf/Mdm2/Tnfaip | 12 | aPD-L1 vs PBS | Up-regulated genes | CD45+ |
| BP | GO:2001057 | reactive nitrogen species metabolic process | 6.44E-07 | 2.81E-05 | Tnf/Cd36/Sod2/Ic | 9 | aPD-L1 vs PBS | Up-regulated genes | CD45+ |
| BP | GO:1901652 | response to peptide | 7.20E-07 | 3.06E-05 | Irf1/Nr4a1/Mdm2/ | 17 | aPD-L1 vs PBS | Up-regulated genes | CD45+ |
| BP | GO:0002521 | leukocyte differentiation | 7.21E-07 | 3.06E-05 | H2-Aa/Cd74/Irf1/ | 22 | aPD-L1 vs PBS | Up-regulated genes | CD45+ |
| BP | GO:0009615 | response to virus | 7.73E-07 | 3.23E-05 | Irf1/Cxcl9/Ifi27l2a | 15 | aPD-L1 vs PBS | Up-regulated genes | CD45+ |
| BP | GO:0043900 | regulation of multi-organism process | 7.84E-07 | 3.23E-05 | Cd74/Inhba/Tnf/A | 18 | aPD-L1 vs PBS | Up-regulated genes | CD45+ |
| BP | GO:0046634 | regulation of alpha-beta T cell activation | 8.00E-07 | 3.26E-05 | Irf1/H2-Ab1/Cd27 | 10 | aPD-L1 vs PBS | Up-regulated genes | CD45+ |
| BP | GO:0002369 | T cell cytokine production | 9.43E-07 | 3.79E-05 | Rsad2/B2m/Malt1 | 7 | aPD-L1 vs PBS | Up-regulated genes | CD45+ |
| BP | GO:0097529 | myeloid leukocyte migration | 9.90E-07 | 3.93E-05 | Cd74/Cxcl9/Ccl4/ | 13 | aPD-L1 vs PBS | Up-regulated genes | CD45+ |
| BP | GO:0009636 | response to toxic substance | 1.01E-06 | 3.94E-05 | Tnf/Ccl4/Slc7a11/ | 15 | aPD-L1 vs PBS | Up-regulated genes | CD45+ |
| BP | GO:0010035 | response to inorganic substance | 1.14E-06 | 4.42E-05 | Crip1/Tnf/Nr4a1/ | 18 | aPD-L1 vs PBS | Up-regulated genes | CD45+ |
| BP | GO:0030217 | T cell differentiation | 1.18E-06 | 4.49E-05 | H2-Aa/Cd74/Irf1/ | 15 | aPD-L1 vs PBS | Up-regulated genes | CD45+ |

| | | | | | | | | | |
|---|---|---|---|---|---|---|---|---|---|
| BP | GO:1902107 | positive regulation of leukocyte differentiation | 1.26E-06 | 4.70E-05 | H2-Aa/Cd74/Irf1/ | 12 | aPD-L1 vs PBS | Up-regulated genes | CD45+ |
| BP | GO:0042136 | neurotransmitter biosynthetic process | 1.26E-06 | 4.70E-05 | Tnf/Cd36/Sod2/Ic | 9 | aPD-L1 vs PBS | Up-regulated genes | CD45+ |
| BP | GO:0045089 | positive regulation of innate immune response | 1.31E-06 | 4.81E-05 | Cd74/Irf1/Irgm1/T | 14 | aPD-L1 vs PBS | Up-regulated genes | CD45+ |
| BP | GO:0045428 | regulation of nitric oxide biosynthetic process | 1.35E-06 | 4.90E-05 | Tnf/Cd36/Sod2/Ic | 8 | aPD-L1 vs PBS | Up-regulated genes | CD45+ |
| BP | GO:0097237 | cellular response to toxic substance | 1.47E-06 | 5.30E-05 | Tnf/Tnfaip3/Cd36/ | 11 | aPD-L1 vs PBS | Up-regulated genes | CD45+ |
| BP | GO:0042274 | ribosomal small subunit biogenesis | 1.53E-06 | 5.39E-05 | Rps27/Rps21/Rpl | 8 | aPD-L1 vs PBS | Up-regulated genes | CD45+ |
| BP | GO:2000514 | regulation of CD4-positive, alpha-beta T cell acti | 1.53E-06 | 5.39E-05 | Irf1/Cd274/Arg2/M | 8 | aPD-L1 vs PBS | Up-regulated genes | CD45+ |
| BP | GO:0002706 | regulation of lymphocyte mediated immunity | 1.65E-06 | 5.64E-05 | Tnf/Rsad2/H2-T2 | 12 | aPD-L1 vs PBS | Up-regulated genes | CD45+ |
| BP | GO:0022408 | negative regulation of cell-cell adhesion | 1.65E-06 | 5.64E-05 | H2-Aa/Cd74/Irf1/ | 12 | aPD-L1 vs PBS | Up-regulated genes | CD45+ |
| BP | GO:0031649 | heat generation | 1.66E-06 | 5.64E-05 | Tnf/Ccl5/Il1a/Il1rn | 5 | aPD-L1 vs PBS | Up-regulated genes | CD45+ |
| BP | GO:0070555 | response to interleukin-1 | 2.11E-06 | 7.11E-05 | Irf1/Ccl4/Acod1/C | 9 | aPD-L1 vs PBS | Up-regulated genes | CD45+ |
| BP | GO:0035458 | cellular response to interferon-beta | 2.53E-06 | 8.41E-05 | Irf1/Irgm1/Gbp2/G | 7 | aPD-L1 vs PBS | Up-regulated genes | CD45+ |
| BP | GO:0002724 | regulation of T cell cytokine production | 2.56E-06 | 8.41E-05 | Rsad2/B2m/Malt1 | 6 | aPD-L1 vs PBS | Up-regulated genes | CD45+ |
| BP | GO:0050727 | regulation of inflammatory response | 2.63E-06 | 8.52E-05 | Tnf/H2-T23/Dusp | 16 | aPD-L1 vs PBS | Up-regulated genes | CD45+ |
| BP | GO:0030098 | lymphocyte differentiation | 2.64E-06 | 8.52E-05 | H2-Aa/Cd74/Irf1/ | 17 | aPD-L1 vs PBS | Up-regulated genes | CD45+ |
| BP | GO:0050900 | leukocyte migration | 2.75E-06 | 8.77E-05 | Cd74/Cxcl9/Tnf/C | 16 | aPD-L1 vs PBS | Up-regulated genes | CD45+ |
| BP | GO:0051250 | negative regulation of lymphocyte activation | 2.85E-06 | 8.99E-05 | H2-Aa/Cd74/Irf1/ | 11 | aPD-L1 vs PBS | Up-regulated genes | CD45+ |
| BP | GO:1902105 | regulation of leukocyte differentiation | 2.98E-06 | 9.21E-05 | H2-Aa/Cd74/Irf1/ | 15 | aPD-L1 vs PBS | Up-regulated genes | CD45+ |
| BP | GO:0071236 | cellular response to antibiotic | 2.98E-06 | 9.21E-05 | Crip1/Mdm2/Tnfai | 10 | aPD-L1 vs PBS | Up-regulated genes | CD45+ |
| BP | GO:0050865 | regulation of cell activation | 3.03E-06 | 9.28E-05 | H2-Aa/Cd74/Irf1/ | 21 | aPD-L1 vs PBS | Up-regulated genes | CD45+ |
| BP | GO:0050830 | defense response to Gram-positive bacterium | 3.17E-06 | 9.47E-05 | Rpl39/Tnf/H2-T23 | 8 | aPD-L1 vs PBS | Up-regulated genes | CD45+ |
| BP | GO:0000028 | ribosomal small subunit assembly | 3.18E-06 | 9.47E-05 | Rps27/Rpl38/Rps | 5 | aPD-L1 vs PBS | Up-regulated genes | CD45+ |
| BP | GO:0002495 | antigen processing and presentation of peptide | 3.18E-06 | 9.47E-05 | H2-Eb1/H2-Aa/C | 5 | aPD-L1 vs PBS | Up-regulated genes | CD45+ |
| BP | GO:0002683 | negative regulation of immune system process | 3.24E-06 | 9.55E-05 | H2-Aa/Cd74/Irf1/ | 19 | aPD-L1 vs PBS | Up-regulated genes | CD45+ |
| BP | GO:0050870 | positive regulation of T cell activation | 3.53E-06 | 1.02E-04 | H2-Aa/Cd74/Irf1/ | 12 | aPD-L1 vs PBS | Up-regulated genes | CD45+ |
| BP | GO:0071347 | cellular response to interleukin-1 | 3.55E-06 | 1.02E-04 | Irf1/Ccl4/Acod1/C | 8 | aPD-L1 vs PBS | Up-regulated genes | CD45+ |
| BP | GO:0051249 | regulation of lymphocyte activation | 3.56E-06 | 1.02E-04 | H2-Aa/Cd74/Irf1/ | 18 | aPD-L1 vs PBS | Up-regulated genes | CD45+ |
| BP | GO:0002702 | positive regulation of production of molecular me | 3.75E-06 | 1.06E-04 | Cd74/Rsad2/H2- | 9 | aPD-L1 vs PBS | Up-regulated genes | CD45+ |
| BP | GO:0002694 | regulation of leukocyte activation | 4.01E-06 | 1.12E-04 | H2-Aa/Cd74/Irf1/ | 20 | aPD-L1 vs PBS | Up-regulated genes | CD45+ |
| BP | GO:0046631 | alpha-beta T cell activation | 4.02E-06 | 1.12E-04 | Irf1/H2-Ab1/Cd27 | 11 | aPD-L1 vs PBS | Up-regulated genes | CD45+ |
| BP | GO:0002428 | antigen processing and presentation of peptide | 4.27E-06 | 1.16E-04 | H2-T23/B2m/H2- | 5 | aPD-L1 vs PBS | Up-regulated genes | CD45+ |
| BP | GO:0002504 | antigen processing and presentation of peptide | 4.27E-06 | 1.16E-04 | H2-Eb1/H2-Aa/C | 5 | aPD-L1 vs PBS | Up-regulated genes | CD45+ |
| BP | GO:0002726 | positive regulation of T cell cytokine production | 4.27E-06 | 1.16E-04 | Rsad2/B2m/Malt1 | 5 | aPD-L1 vs PBS | Up-regulated genes | CD45+ |
| BP | GO:0035710 | CD4-positive, alpha-beta T cell activation | 4.50E-06 | 1.21E-04 | Irf1/Cd274/Arg2/ | 9 | aPD-L1 vs PBS | Up-regulated genes | CD45+ |
| BP | GO:0001916 | positive regulation of T cell mediated cytotoxicity | 4.61E-06 | 1.23E-04 | H2-T23/B2m/H2- | 6 | aPD-L1 vs PBS | Up-regulated genes | CD45+ |
| BP | GO:0002703 | regulation of leukocyte mediated immunity | 4.71E-06 | 1.24E-04 | Tnf/Rsad2/H2-T2 | 13 | aPD-L1 vs PBS | Up-regulated genes | CD45+ |
| BP | GO:1903038 | negative regulation of leukocyte cell-cell adhesi | 4.76E-06 | 1.24E-04 | H2-Aa/Cd74/Irf1/ | 10 | aPD-L1 vs PBS | Up-regulated genes | CD45+ |
| BP | GO:2000377 | regulation of reactive oxygen species metabolic | 4.76E-06 | 1.24E-04 | Tnf/Arg2/Acod1/C | 12 | aPD-L1 vs PBS | Up-regulated genes | CD45+ |
| BP | GO:0050863 | regulation of T cell activation | 4.91E-06 | 1.26E-04 | H2-Aa/Cd74/Irf1/ | 15 | aPD-L1 vs PBS | Up-regulated genes | CD45+ |
| BP | GO:0007249 | I-kappaB kinase/NF-kappaB signaling | 5.05E-06 | 1.29E-04 | Cd74/Irf1/Tnf/Tnfa | 12 | aPD-L1 vs PBS | Up-regulated genes | CD45+ |
| BP | GO:0071407 | cellular response to organic cyclic compound | 5.17E-06 | 1.31E-04 | Dnaja1/Inhba/Tnf/ | 16 | aPD-L1 vs PBS | Up-regulated genes | CD45+ |
| BP | GO:1900407 | regulation of cellular response to oxidative stres | 5.49E-06 | 1.38E-04 | Tnf/Slc7a11/Cd3 | 8 | aPD-L1 vs PBS | Up-regulated genes | CD45+ |
| BP | GO:0031348 | negative regulation of defense response | 5.67E-06 | 1.41E-04 | Arg2/H2-T23/Dus | 12 | aPD-L1 vs PBS | Up-regulated genes | CD45+ |
| BP | GO:0060326 | cell chemotaxis | 5.75E-06 | 1.42E-04 | Cd74/Cxcl9/Nr4a | 14 | aPD-L1 vs PBS | Up-regulated genes | CD45+ |
| BP | GO:0043901 | negative regulation of multi-organism process | 5.97E-06 | 1.47E-04 | Tnf/Arg2/Rsad2/C | 11 | aPD-L1 vs PBS | Up-regulated genes | CD45+ |
| BP | GO:0044403 | symbiont process | 6.59E-06 | 1.60E-04 | Cd74/Irgm1/Tnf/R | 16 | aPD-L1 vs PBS | Up-regulated genes | CD45+ |
| BP | GO:0071356 | cellular response to tumor necrosis factor | 6.89E-06 | 1.66E-04 | Irf1/Tnf/Ccl4/Acod | 10 | aPD-L1 vs PBS | Up-regulated genes | CD45+ |
| BP | GO:0048661 | positive regulation of smooth muscle cell prolifer | 7.55E-06 | 1.81E-04 | Tnf/Mdm2/Ccl5/N | 9 | aPD-L1 vs PBS | Up-regulated genes | CD45+ |
| BP | GO:1901031 | regulation of response to reactive oxygen specie | 7.84E-06 | 1.86E-04 | Tnf/Cd36/Hk3/Nr4 | 6 | aPD-L1 vs PBS | Up-regulated genes | CD45+ |
| BP | GO:0035690 | cellular response to drug | 7.96E-06 | 1.88E-04 | Tnf/Ccl4/Acod1/T | 14 | aPD-L1 vs PBS | Up-regulated genes | CD45+ |
| BP | GO:0035456 | response to interferon-beta | 8.65E-06 | 2.03E-04 | Irf1/Irgm1/Gbp2/G | 7 | aPD-L1 vs PBS | Up-regulated genes | CD45+ |
| BP | GO:0030595 | leukocyte chemotaxis | 8.87E-06 | 2.05E-04 | Cd74/Cxcl9/Ccl4/ | 12 | aPD-L1 vs PBS | Up-regulated genes | CD45+ |
| BP | GO:0033002 | muscle cell proliferation | 8.87E-06 | 2.05E-04 | Tnf/Mdm2/Tnfaip3 | 12 | aPD-L1 vs PBS | Up-regulated genes | CD45+ |
| BP | GO:0002475 | antigen processing and presentation via MHC cl | 9.36E-06 | 2.13E-04 | H2-T23/B2m/H2- | 5 | aPD-L1 vs PBS | Up-regulated genes | CD45+ |
| BP | GO:0046677 | response to antibiotic | 9.36E-06 | 2.13E-04 | Crip1/Mdm2/Tnfai | 12 | aPD-L1 vs PBS | Up-regulated genes | CD45+ |
| BP | GO:0001914 | regulation of T cell mediated cytotoxicity | 1.09E-05 | 2.45E-04 | H2-T23/B2m/H2- | 6 | aPD-L1 vs PBS | Up-regulated genes | CD45+ |
| BP | GO:1902882 | regulation of response to oxidative stress | 1.10E-05 | 2.47E-04 | Tnf/Slc7a11/Cd3 | 8 | aPD-L1 vs PBS | Up-regulated genes | CD45+ |
| BP | GO:0002695 | negative regulation of leukocyte activation | 1.24E-05 | 2.77E-04 | H2-Aa/Cd74/Irf1/ | 11 | aPD-L1 vs PBS | Up-regulated genes | CD45+ |
| BP | GO:0042255 | ribosome assembly | 1.39E-05 | 3.07E-04 | Rps27/Rpl38/Rps | 7 | aPD-L1 vs PBS | Up-regulated genes | CD45+ |
| BP | GO:0002483 | antigen processing and presentation of endoger | 1.48E-05 | 3.20E-04 | H2-T23/B2m/H2- | 5 | aPD-L1 vs PBS | Up-regulated genes | CD45+ |
| BP | GO:0019885 | antigen processing and presentation of endoger | 1.48E-05 | 3.20E-04 | H2-T23/B2m/H2- | 5 | aPD-L1 vs PBS | Up-regulated genes | CD45+ |
| BP | GO:0045619 | regulation of lymphocyte differentiation | 1.48E-05 | 3.20E-04 | H2-Aa/Cd74/Irf1/ | 11 | aPD-L1 vs PBS | Up-regulated genes | CD45+ |
| BP | GO:0045580 | regulation of T cell differentiation | 1.56E-05 | 3.35E-04 | H2-Aa/Cd74/Irf1/ | 10 | aPD-L1 vs PBS | Up-regulated genes | CD45+ |
| BP | GO:0050868 | negative regulation of T cell activation | 1.66E-05 | 3.50E-04 | H2-Aa/Cd74/Irf1/ | 9 | aPD-L1 vs PBS | Up-regulated genes | CD45+ |
| BP | GO:0051607 | defense response to virus | 1.66E-05 | 3.50E-04 | Irf1/Cxcl9/Rsad2/ | 12 | aPD-L1 vs PBS | Up-regulated genes | CD45+ |
| BP | GO:0001906 | cell killing | 1.66E-05 | 3.50E-04 | Rpl30/Hsp90ab1/ | 10 | aPD-L1 vs PBS | Up-regulated genes | CD45+ |
| BP | GO:0006953 | acute-phase response | 1.83E-05 | 3.82E-04 | Tnf/Ccl5/Il1a/Il1rn | 5 | aPD-L1 vs PBS | Up-regulated genes | CD45+ |
| BP | GO:0072593 | reactive oxygen species metabolic process | 1.85E-05 | 3.84E-04 | Tnf/Arg2/Acod1/C | 13 | aPD-L1 vs PBS | Up-regulated genes | CD45+ |
| BP | GO:0002526 | acute inflammatory response | 1.90E-05 | 3.92E-04 | Tnf/H2-T23/Ccl5/ | 8 | aPD-L1 vs PBS | Up-regulated genes | CD45+ |
| BP | GO:0034113 | heterotypic cell-cell adhesion | 1.97E-05 | 4.02E-04 | Tnf/Tnfaip3/Itgb7/ | 6 | aPD-L1 vs PBS | Up-regulated genes | CD45+ |
| BP | GO:1903202 | negative regulation of oxidative stress-induced c | 1.97E-05 | 4.02E-04 | Slc7a11/Hk3/Nr4 | 6 | aPD-L1 vs PBS | Up-regulated genes | CD45+ |
| BP | GO:0032870 | cellular response to hormone stimulus | 1.99E-05 | 4.03E-04 | Irf1/Dnaja1/Inhba/ | 17 | aPD-L1 vs PBS | Up-regulated genes | CD45+ |
| BP | GO:0044406 | adhesion of symbiont to host | 2.10E-05 | 4.23E-04 | Hsp90ab1/Gbp2/ | 4 | aPD-L1 vs PBS | Up-regulated genes | CD45+ |
| BP | GO:0034612 | response to tumor necrosis factor | 2.14E-05 | 4.27E-04 | Irf1/Tnf/Ccl4/Acod | 10 | aPD-L1 vs PBS | Up-regulated genes | CD45+ |
| BP | GO:0042133 | neurotransmitter metabolic process | 2.22E-05 | 4.38E-04 | Tnf/Cd36/Sod2/Ic | 9 | aPD-L1 vs PBS | Up-regulated genes | CD45+ |
| BP | GO:0019883 | antigen processing and presentation of endoger | 2.24E-05 | 4.38E-04 | H2-T23/B2m/H2- | 5 | aPD-L1 vs PBS | Up-regulated genes | CD45+ |
| BP | GO:0002286 | T cell activation involved in immune response | 2.26E-05 | 4.38E-04 | Irf1/Malt1/Icam1/F | 8 | aPD-L1 vs PBS | Up-regulated genes | CD45+ |
| BP | GO:0045582 | positive regulation of T cell differentiation | 2.26E-05 | 4.38E-04 | H2-Aa/Cd74/Irf1/ | 8 | aPD-L1 vs PBS | Up-regulated genes | CD45+ |
| BP | GO:1900408 | negative regulation of cellular response to oxida | 2.27E-05 | 4.38E-04 | Slc7a11/Hk3/Nr4 | 6 | aPD-L1 vs PBS | Up-regulated genes | CD45+ |
| BP | GO:0032623 | interleukin-2 production | 2.39E-05 | 4.60E-04 | Tnfaip3/Malt1/Cd | 7 | aPD-L1 vs PBS | Up-regulated genes | CD45+ |
| BP | GO:0019730 | antimicrobial humoral response | 2.60E-05 | 4.96E-04 | Rpl39/Cxcl9/Rpl3 | 6 | aPD-L1 vs PBS | Up-regulated genes | CD45+ |
| BP | GO:0046635 | positive regulation of alpha-beta T cell activation | 2.65E-05 | 5.03E-04 | Irf1/H2-Ab1/H2-T | 7 | aPD-L1 vs PBS | Up-regulated genes | CD45+ |
| BP | GO:0045639 | positive regulation of myeloid cell differentiation | 2.66E-05 | 5.03E-04 | Cd74/Inhba/Tnf/C | 8 | aPD-L1 vs PBS | Up-regulated genes | CD45+ |
| BP | GO:0050866 | negative regulation of cell activation | 2.85E-05 | 5.35E-04 | H2-Aa/Cd74/Irf1/ | 11 | aPD-L1 vs PBS | Up-regulated genes | CD45+ |
| BP | GO:2000341 | regulation of chemokine (C-X-C motif) ligand 2 p | 2.92E-05 | 5.43E-04 | Cd74/Tnf/Ccl5/Klf | 4 | aPD-L1 vs PBS | Up-regulated genes | CD45+ |
| BP | GO:0002696 | positive regulation of leukocyte activation | 2.94E-05 | 5.45E-04 | H2-Aa/Cd74/Irf1/ | 14 | aPD-L1 vs PBS | Up-regulated genes | CD45+ |
| BP | GO:0043372 | positive regulation of CD4-positive, alpha-beta T | 3.27E-05 | 5.99E-04 | Irf1/Malt1/Cd83/S | 5 | aPD-L1 vs PBS | Up-regulated genes | CD45+ |
| BP | GO:2001024 | negative regulation of response to drug | 3.27E-05 | 5.99E-04 | Tnf/Hk3/Nr4a3/Ps | 5 | aPD-L1 vs PBS | Up-regulated genes | CD45+ |
| BP | GO:1902883 | negative regulation of response to oxidative stre | 3.37E-05 | 6.13E-04 | Slc7a11/Hk3/Nr4 | 6 | aPD-L1 vs PBS | Up-regulated genes | CD45+ |

| | | | | | | | | | |
|---|---|---|---|---|---|---|---|---|---|
| BP | GO:0043370 | regulation of CD4-positive, alpha-beta T cell diffe | 3.82E-05 | 6.91E-04 | Irf1/Malt1/Cd83/S | 6 | aPD-L1 vs PBS | Up-regulated genes | CD45+ |
| BP | GO:0072567 | chemokine (C-X-C motif) ligand 2 production | 3.93E-05 | 7.08E-04 | Cd74/Tnf/Ccl5/Klf | 4 | aPD-L1 vs PBS | Up-regulated genes | CD45+ |
| BP | GO:0002262 | myeloid cell homeostasis | 4.09E-05 | 7.31E-04 | Inhba/Rps24/B2m | 10 | aPD-L1 vs PBS | Up-regulated genes | CD45+ |
| BP | GO:0045824 | negative regulation of innate immune response | 4.32E-05 | 7.67E-04 | H2-T23/Dusp10/A | 6 | aPD-L1 vs PBS | Up-regulated genes | CD45+ |
| BP | GO:0042035 | regulation of cytokine biosynthetic process | 4.62E-05 | 8.16E-04 | Irf1/Tnf/Zfp36/Rel/ | 8 | aPD-L1 vs PBS | Up-regulated genes | CD45+ |
| BP | GO:0050867 | positive regulation of cell activation | 5.00E-05 | 8.74E-04 | H2-Aa/Cd74/Irf1/I | 14 | aPD-L1 vs PBS | Up-regulated genes | CD45+ |
| BP | GO:0071417 | cellular response to organonitrogen compound | 5.00E-05 | 8.74E-04 | Irf1/Tnf/Nr4a1/Md | 16 | aPD-L1 vs PBS | Up-regulated genes | CD45+ |
| BP | GO:0045637 | regulation of myeloid cell differentiation | 5.21E-05 | 9.05E-04 | Cd74/Inhba/Tnf/B | 11 | aPD-L1 vs PBS | Up-regulated genes | CD45+ |
| BP | GO:0002828 | regulation of type 2 immune response | 5.47E-05 | 9.40E-04 | Cd74/Irf1/Arg2/Rs | 5 | aPD-L1 vs PBS | Up-regulated genes | CD45+ |
| BP | GO:0061756 | leukocyte adhesion to vascular endothelial cell | 5.47E-05 | 9.40E-04 | Tnf/Icam1/Itgb7/Kl | 5 | aPD-L1 vs PBS | Up-regulated genes | CD45+ |
| BP | GO:0042254 | ribosome biogenesis | 5.52E-05 | 9.43E-04 | Rps27/Rps21/Rpl | 13 | aPD-L1 vs PBS | Up-regulated genes | CD45+ |
| BP | GO:1902532 | negative regulation of intracellular signal transdu | 5.82E-05 | 9.89E-04 | Cd74/Irf1/Dnaja1/ | 18 | aPD-L1 vs PBS | Up-regulated genes | CD45+ |
| BP | GO:1901699 | cellular response to nitrogen compound | 5.93E-05 | 1.00E-03 | Irf1/Tnf/Nr4a1/Md | 17 | aPD-L1 vs PBS | Up-regulated genes | CD45+ |
| BP | GO:0045621 | positive regulation of lymphocyte differentiation | 6.19E-05 | 1.04E-03 | H2-Aa/Cd74/Irf1/D | 8 | aPD-L1 vs PBS | Up-regulated genes | CD45+ |
| BP | GO:0031341 | regulation of cell killing | 6.64E-05 | 1.09E-03 | Hsp90ab1/H2-T2 | 8 | aPD-L1 vs PBS | Up-regulated genes | CD45+ |
| BP | GO:1904018 | positive regulation of vasculature development | 6.65E-05 | 1.09E-03 | Ccl5/Hk2/Sod2/Kl | 10 | aPD-L1 vs PBS | Up-regulated genes | CD45+ |
| BP | GO:0034614 | cellular response to reactive oxygen species | 6.71E-05 | 1.09E-03 | Tnf/Tnfaip3/Cd36/ | 9 | aPD-L1 vs PBS | Up-regulated genes | CD45+ |
| BP | GO:0002486 | antigen processing and presentation of endoger | 6.71E-05 | 1.09E-03 | H2-T23/H2-D1/H2 | 4 | aPD-L1 vs PBS | Up-regulated genes | CD45+ |
| BP | GO:2001185 | regulation of CD8-positive, alpha-beta T cell acti | 6.71E-05 | 1.09E-03 | Irf1/Cd274/H2-T2 | 4 | aPD-L1 vs PBS | Up-regulated genes | CD45+ |
| BP | GO:0031343 | positive regulation of cell killing | 6.74E-05 | 1.09E-03 | H2-T23/B2m/H2-I | 7 | aPD-L1 vs PBS | Up-regulated genes | CD45+ |
| BP | GO:0043367 | CD4-positive, alpha-beta T cell differentiation | 6.74E-05 | 1.09E-03 | Irf1/Rsad2/Malt1/ | 7 | aPD-L1 vs PBS | Up-regulated genes | CD45+ |
| BP | GO:0002474 | antigen processing and presentation of peptide | 7.48E-05 | 1.20E-03 | H2-T23/B2m/H2-I | 5 | aPD-L1 vs PBS | Up-regulated genes | CD45+ |
| BP | GO:0006979 | response to oxidative stress | 7.98E-05 | 1.27E-03 | Tnf/Slc7a11/ND3/ | 15 | aPD-L1 vs PBS | Up-regulated genes | CD45+ |
| BP | GO:0032642 | regulation of chemokine production | 7.99E-05 | 1.27E-03 | Cd74/Tnf/Arg2/Cc | 7 | aPD-L1 vs PBS | Up-regulated genes | CD45+ |
| BP | GO:1901653 | cellular response to peptide | 8.12E-05 | 1.29E-03 | Irf1/Nr4a1/Mdm2/ | 12 | aPD-L1 vs PBS | Up-regulated genes | CD45+ |
| BP | GO:0060759 | regulation of response to cytokine stimulus | 8.18E-05 | 1.29E-03 | Cd74/Irgm1/Zbp1 | 8 | aPD-L1 vs PBS | Up-regulated genes | CD45+ |
| BP | GO:0002476 | antigen processing and presentation of endoger | 8.54E-05 | 1.32E-03 | H2-T23/H2-D1/H2 | 4 | aPD-L1 vs PBS | Up-regulated genes | CD45+ |
| BP | GO:0002484 | antigen processing and presentation of endoger | 8.54E-05 | 1.32E-03 | H2-T23/H2-D1/H2 | 4 | aPD-L1 vs PBS | Up-regulated genes | CD45+ |
| BP | GO:0002577 | regulation of antigen processing and presentatic | 8.54E-05 | 1.32E-03 | Cd74/H2-Ab1/Fgl | 4 | aPD-L1 vs PBS | Up-regulated genes | CD45+ |
| BP | GO:1904994 | regulation of leukocyte adhesion to vascular enc | 8.54E-05 | 1.32E-03 | Tnf/Icam1/Klf4/Itgl | 4 | aPD-L1 vs PBS | Up-regulated genes | CD45+ |
| BP | GO:2000516 | positive regulation of CD4-positive, alpha-beta T | 8.67E-05 | 1.33E-03 | Irf1/Malt1/Cd83/S | 5 | aPD-L1 vs PBS | Up-regulated genes | CD45+ |
| BP | GO:0007162 | negative regulation of cell adhesion | 8.81E-05 | 1.34E-03 | H2-Aa/Cd74/Irf1/I | 12 | aPD-L1 vs PBS | Up-regulated genes | CD45+ |
| BP | GO:0051251 | positive regulation of lymphocyte activation | 8.81E-05 | 1.34E-03 | H2-Aa/Cd74/Irf1/I | 12 | aPD-L1 vs PBS | Up-regulated genes | CD45+ |
| BP | GO:1901342 | regulation of vasculature development | 9.04E-05 | 1.37E-03 | Ccl5/Cd36/Stat1/I | 13 | aPD-L1 vs PBS | Up-regulated genes | CD45+ |
| BP | GO:0009144 | purine nucleoside triphosphate metabolic proces | 9.17E-05 | 1.38E-03 | ATP8/Atp5e/Hspa | 12 | aPD-L1 vs PBS | Up-regulated genes | CD45+ |
| BP | GO:0001913 | T cell mediated cytotoxicity | 9.43E-05 | 1.42E-03 | H2-T23/B2m/H2-I | 6 | aPD-L1 vs PBS | Up-regulated genes | CD45+ |
| BP | GO:0002831 | regulation of response to biotic stimulus | 9.52E-05 | 1.42E-03 | Cd274/Arg2/Acod | 9 | aPD-L1 vs PBS | Up-regulated genes | CD45+ |
| BP | GO:0048872 | homeostasis of number of cells | 9.73E-05 | 1.45E-03 | Cd74/Inhba/Rps2 | 13 | aPD-L1 vs PBS | Up-regulated genes | CD45+ |
| BP | GO:0042089 | cytokine biosynthetic process | 1.00E-04 | 1.48E-03 | Irf1/Tnf/Zfp36/Rel/ | 8 | aPD-L1 vs PBS | Up-regulated genes | CD45+ |
| BP | GO:0032663 | regulation of interleukin-2 production | 1.04E-04 | 1.54E-03 | Tnfaip3/Malt1/Cd | 6 | aPD-L1 vs PBS | Up-regulated genes | CD45+ |
| BP | GO:0046034 | ATP metabolic process | 1.08E-04 | 1.58E-03 | ATP8/Atp5e/Hspa | 11 | aPD-L1 vs PBS | Up-regulated genes | CD45+ |
| BP | GO:2001234 | negative regulation of apoptotic signaling pathw | 1.08E-04 | 1.58E-03 | Cd74/Lmna/Dnaja | 11 | aPD-L1 vs PBS | Up-regulated genes | CD45+ |
| BP | GO:0022613 | ribonucleoprotein complex biogenesis | 1.10E-04 | 1.60E-03 | Rps27/Rps21/Rpl | 16 | aPD-L1 vs PBS | Up-regulated genes | CD45+ |
| BP | GO:0032602 | chemokine production | 1.10E-04 | 1.60E-03 | Cd74/Tnf/Arg2/Cc | 7 | aPD-L1 vs PBS | Up-regulated genes | CD45+ |
| BP | GO:0045123 | cellular extravasation | 1.15E-04 | 1.66E-03 | Tnf/Ccl5/Icam1/Itg | 6 | aPD-L1 vs PBS | Up-regulated genes | CD45+ |
| BP | GO:0032635 | interleukin-6 production | 1.19E-04 | 1.70E-03 | Tnf/Tnfaip3/Cd36/ | 9 | aPD-L1 vs PBS | Up-regulated genes | CD45+ |
| BP | GO:0042107 | cytokine metabolic process | 1.29E-04 | 1.82E-03 | Irf1/Tnf/Zfp36/Rel/ | 8 | aPD-L1 vs PBS | Up-regulated genes | CD45+ |
| BP | GO:0042092 | type 2 immune response | 1.31E-04 | 1.82E-03 | Cd74/Irf1/Arg2/Rs | 5 | aPD-L1 vs PBS | Up-regulated genes | CD45+ |
| BP | GO:0015985 | energy coupled proton transport, down electroch | 1.32E-04 | 1.82E-03 | ATP8/Atp5e/ATP6 | 4 | aPD-L1 vs PBS | Up-regulated genes | CD45+ |
| BP | GO:0015986 | ATP synthesis coupled proton transport | 1.32E-04 | 1.82E-03 | ATP8/Atp5e/ATP6 | 4 | aPD-L1 vs PBS | Up-regulated genes | CD45+ |
| BP | GO:0034114 | regulation of heterotypic cell-cell adhesion | 1.32E-04 | 1.82E-03 | Tnf/Tnfaip3/Klf4/Il | 4 | aPD-L1 vs PBS | Up-regulated genes | CD45+ |
| BP | GO:0060333 | interferon-gamma-mediated signaling pathway | 1.32E-04 | 1.82E-03 | Irf1/Irgm1/Stat1/S | 4 | aPD-L1 vs PBS | Up-regulated genes | CD45+ |
| BP | GO:1901032 | negative regulation of response to reactive oxyg | 1.32E-04 | 1.82E-03 | Hk3/Nr4a3/Psap/ | 4 | aPD-L1 vs PBS | Up-regulated genes | CD45+ |
| BP | GO:1903206 | negative regulation of hydrogen peroxide-induce | 1.32E-04 | 1.82E-03 | Hk3/Nr4a3/Psap/ | 4 | aPD-L1 vs PBS | Up-regulated genes | CD45+ |
| BP | GO:2001039 | negative regulation of cellular response to drug | 1.32E-04 | 1.82E-03 | Hk3/Nr4a3/Psap/ | 4 | aPD-L1 vs PBS | Up-regulated genes | CD45+ |
| BP | GO:0000302 | response to reactive oxygen species | 1.33E-04 | 1.82E-03 | Tnf/Tnfaip3/Cd36/ | 10 | aPD-L1 vs PBS | Up-regulated genes | CD45+ |
| BP | GO:0050663 | cytokine secretion | 1.46E-04 | 1.98E-03 | Cd274/Tnf/Arg2/T | 11 | aPD-L1 vs PBS | Up-regulated genes | CD45+ |
| BP | GO:0006959 | humoral immune response | 1.47E-04 | 1.99E-03 | Rpl39/Cxcl9/H2-A | 8 | aPD-L1 vs PBS | Up-regulated genes | CD45+ |
| BP | GO:0002791 | regulation of peptide secretion | 1.55E-04 | 2.09E-03 | Cd74/Cd274/Tnf/ | 16 | aPD-L1 vs PBS | Up-regulated genes | CD45+ |
| BP | GO:0016032 | viral process | 1.58E-04 | 2.12E-03 | Cd74/Tnf/Rsad2/ | 12 | aPD-L1 vs PBS | Up-regulated genes | CD45+ |
| BP | GO:0010742 | macrophage derived foam cell differentiation | 1.62E-04 | 2.16E-03 | Cd36/Nfkbia/Stat | 4 | aPD-L1 vs PBS | Up-regulated genes | CD45+ |
| BP | GO:2000316 | regulation of T-helper 17 type immune response | 1.62E-04 | 2.16E-03 | Malt1/Nlrp3/Arid5 | 4 | aPD-L1 vs PBS | Up-regulated genes | CD45+ |
| BP | GO:0009141 | nucleoside triphosphate metabolic process | 1.70E-04 | 2.25E-03 | ATP8/Atp5e/Hspa | 12 | aPD-L1 vs PBS | Up-regulated genes | CD45+ |
| BP | GO:0050730 | regulation of peptidyl-tyrosine phosphorylation | 1.79E-04 | 2.36E-03 | Cd74/Irf1/Tnf/Sam | 11 | aPD-L1 vs PBS | Up-regulated genes | CD45+ |
| BP | GO:0046637 | regulation of alpha-beta T cell differentiation | 1.85E-04 | 2.43E-03 | Irf1/Malt1/Cd83/S | 6 | aPD-L1 vs PBS | Up-regulated genes | CD45+ |
| BP | GO:0046636 | negative regulation of alpha-beta T cell activatio | 1.92E-04 | 2.52E-03 | Irf1/Cd274/Arg2/S | 5 | aPD-L1 vs PBS | Up-regulated genes | CD45+ |
| BP | GO:0002827 | positive regulation of T-helper 1 type immune re | 1.96E-04 | 2.54E-03 | Irf1/H2-Ab1/Slc11 | 4 | aPD-L1 vs PBS | Up-regulated genes | CD45+ |
| BP | GO:0090077 | foam cell differentiation | 1.96E-04 | 2.54E-03 | Cd36/Nfkbia/Stat | 4 | aPD-L1 vs PBS | Up-regulated genes | CD45+ |
| BP | GO:0001912 | positive regulation of leukocyte mediated cytoto | 2.02E-04 | 2.61E-03 | H2-T23/B2m/H2-I | 6 | aPD-L1 vs PBS | Up-regulated genes | CD45+ |
| BP | GO:1903201 | regulation of oxidative stress-induced cell death | 2.02E-04 | 2.61E-03 | Slc7a11/Hk3/Nr4 | 6 | aPD-L1 vs PBS | Up-regulated genes | CD45+ |
| BP | GO:0050707 | regulation of cytokine secretion | 2.10E-04 | 2.69E-03 | Cd274/Tnf/Arg2/T | 10 | aPD-L1 vs PBS | Up-regulated genes | CD45+ |
| BP | GO:0032103 | positive regulation of response to external stimul | 2.18E-04 | 2.79E-03 | Cd74/Tnf/Ccl4/H2 | 12 | aPD-L1 vs PBS | Up-regulated genes | CD45+ |
| BP | GO:0050729 | positive regulation of inflammatory response | 2.22E-04 | 2.83E-03 | Tnf/H2-T23/Ccl5/ | 8 | aPD-L1 vs PBS | Up-regulated genes | CD45+ |
| BP | GO:0002366 | leukocyte activation involved in immune respons | 2.26E-04 | 2.87E-03 | Irf1/Tnfaip3/Malt1 | 11 | aPD-L1 vs PBS | Up-regulated genes | CD45+ |
| BP | GO:0032943 | mononuclear cell proliferation | 2.34E-04 | 2.93E-03 | H2-Aa/Cd74/Irf1/I | 12 | aPD-L1 vs PBS | Up-regulated genes | CD45+ |
| BP | GO:0002675 | positive regulation of acute inflammatory respon | 2.34E-04 | 2.93E-03 | Tnf/H2-T23/Ccl5/I | 4 | aPD-L1 vs PBS | Up-regulated genes | CD45+ |
| BP | GO:0002698 | negative regulation of immune effector process | 2.35E-04 | 2.93E-03 | Irf1/Tnf/H2-T23/D | 8 | aPD-L1 vs PBS | Up-regulated genes | CD45+ |
| BP | GO:0034101 | erythrocyte homeostasis | 2.35E-04 | 2.93E-03 | Inhba/Rps24/B2m | 8 | aPD-L1 vs PBS | Up-regulated genes | CD45+ |
| BP | GO:0043903 | regulation of symbiosis, encompassing mutualis | 2.39E-04 | 2.97E-03 | Cd74/Tnf/Rsad2/ | 10 | aPD-L1 vs PBS | Up-regulated genes | CD45+ |
| BP | GO:0042098 | T cell proliferation | 2.60E-04 | 3.22E-03 | H2-Aa/Irf1/H2-Ab | 10 | aPD-L1 vs PBS | Up-regulated genes | CD45+ |
| BP | GO:0001959 | regulation of cytokine-mediated signaling pathw | 2.62E-04 | 3.23E-03 | Cd74/Irgm1/Zbp1 | 7 | aPD-L1 vs PBS | Up-regulated genes | CD45+ |
| BP | GO:0050728 | negative regulation of inflammatory response | 2.63E-04 | 3.23E-03 | Dusp10/Acod1/Tr | 8 | aPD-L1 vs PBS | Up-regulated genes | CD45+ |
| BP | GO:0002263 | cell activation involved in immune response | 2.64E-04 | 3.23E-03 | Irf1/Tnfaip3/Malt1 | 11 | aPD-L1 vs PBS | Up-regulated genes | CD45+ |
| BP | GO:0002468 | dendritic cell antigen processing and presentatic | 2.72E-04 | 3.25E-03 | Cd74/Fgl2/Slc11a | 3 | aPD-L1 vs PBS | Up-regulated genes | CD45+ |
| BP | GO:0002579 | positive regulation of antigen processing and pr | 2.72E-04 | 3.25E-03 | Cd74/H2-Ab1/Slc | 3 | aPD-L1 vs PBS | Up-regulated genes | CD45+ |
| BP | GO:0031652 | positive regulation of heat generation | 2.72E-04 | 3.25E-03 | Tnf/Ccl5/Ptgs2 | 3 | aPD-L1 vs PBS | Up-regulated genes | CD45+ |
| BP | GO:0042090 | interleukin-12 biosynthetic process | 2.72E-04 | 3.25E-03 | Irf1/Rel/Nfkb1 | 3 | aPD-L1 vs PBS | Up-regulated genes | CD45+ |
| BP | GO:0045075 | regulation of interleukin-12 biosynthetic process | 2.72E-04 | 3.25E-03 | Irf1/Rel/Nfkb1 | 3 | aPD-L1 vs PBS | Up-regulated genes | CD45+ |
| BP | GO:0070586 | cell-cell adhesion involved in gastrulation | 2.72E-04 | 3.25E-03 | Tnfaip3/Klf4/Il1rn | 3 | aPD-L1 vs PBS | Up-regulated genes | CD45+ |

| | | | | | | | | | |
|---|---|---|---|---|---|---|---|---|---|
| BP | GO:0009126 | purine nucleoside monophosphate metabolic pr | 2.74E-04 | 3.25E-03 | ATP8/Atp5e/Hspa | 11 | aPD-L1 vs PBS | Up-regulated genes | CD45+ |
| BP | GO:0009167 | purine ribonucleoside monophosphate metaboli | 2.74E-04 | 3.25E-03 | ATP8/Atp5e/Hspa | 11 | aPD-L1 vs PBS | Up-regulated genes | CD45+ |
| BP | GO:0002292 | T cell differentiation involved in immune respons | 2.84E-04 | 3.35E-03 | Irf1/Malt1/Fgl2/Cle | 6 | aPD-L1 vs PBS | Up-regulated genes | CD45+ |
| BP | GO:0009205 | purine ribonucleoside triphosphate metabolic pr | 2.85E-04 | 3.35E-03 | ATP8/Atp5e/Hspa | 11 | aPD-L1 vs PBS | Up-regulated genes | CD45+ |
| BP | GO:0050792 | regulation of viral process | 2.96E-04 | 3.47E-03 | Cd74/Tnf/Rsad2/( | 9 | aPD-L1 vs PBS | Up-regulated genes | CD45+ |
| BP | GO:0045581 | negative regulation of T cell differentiation | 3.03E-04 | 3.54E-03 | Cd74/Irf1/Socs1/F | 5 | aPD-L1 vs PBS | Up-regulated genes | CD45+ |
| BP | GO:0030099 | myeloid cell differentiation | 3.08E-04 | 3.58E-03 | Cd74/Inhba/Tnf/B | 14 | aPD-L1 vs PBS | Up-regulated genes | CD45+ |
| BP | GO:0009161 | ribonucleoside monophosphate metabolic proce | 3.18E-04 | 3.69E-03 | ATP8/Atp5e/Hspa | 11 | aPD-L1 vs PBS | Up-regulated genes | CD45+ |
| BP | GO:0036037 | CD8-positive, alpha-beta T cell activation | 3.28E-04 | 3.79E-03 | Irf1/Cd274/H2-T2 | 4 | aPD-L1 vs PBS | Up-regulated genes | CD45+ |
| BP | GO:0009199 | ribonucleoside triphosphate metabolic process | 3.30E-04 | 3.80E-03 | ATP8/Atp5e/Hspa | 11 | aPD-L1 vs PBS | Up-regulated genes | CD45+ |
| BP | GO:0045444 | fat cell differentiation | 3.34E-04 | 3.83E-03 | Tnf/Nr4a1/Dusp1( | 10 | aPD-L1 vs PBS | Up-regulated genes | CD45+ |
| BP | GO:0046638 | positive regulation of alpha-beta T cell differentia | 3.38E-04 | 3.85E-03 | Irf1/Malt1/Cd83/S | 5 | aPD-L1 vs PBS | Up-regulated genes | CD45+ |
| BP | GO:0002224 | toll-like receptor signaling pathway | 3.40E-04 | 3.86E-03 | Irf1/Tnf/Rsad2/Ac | 7 | aPD-L1 vs PBS | Up-regulated genes | CD45+ |
| BP | GO:0002285 | lymphocyte activation involved in immune respo | 3.40E-04 | 3.86E-03 | Irf1/Tnfaip3/Malt1 | 9 | aPD-L1 vs PBS | Up-regulated genes | CD45+ |
| BP | GO:2000343 | positive regulation of chemokine (C-X-C motif) li | 3.71E-04 | 4.19E-03 | Cd74/Tnf/Ccl5 | 3 | aPD-L1 vs PBS | Up-regulated genes | CD45+ |
| BP | GO:0045646 | regulation of erythrocyte differentiation | 3.75E-04 | 4.22E-03 | Inhba/B2m/Stat1/ | 5 | aPD-L1 vs PBS | Up-regulated genes | CD45+ |
| BP | GO:0002221 | pattern recognition receptor signaling pathway | 3.83E-04 | 4.29E-03 | Irf1/Tnf/Rsad2/Ac | 8 | aPD-L1 vs PBS | Up-regulated genes | CD45+ |
| BP | GO:0001933 | negative regulation of protein phosphorylation | 3.87E-04 | 4.32E-03 | Irf1/Dnaja1/Sams | 14 | aPD-L1 vs PBS | Up-regulated genes | CD45+ |
| BP | GO:0008015 | blood circulation | 3.90E-04 | 4.34E-03 | Tnf/Ccl4/Mdm2/C | 13 | aPD-L1 vs PBS | Up-regulated genes | CD45+ |
| BP | GO:0070661 | leukocyte proliferation | 4.02E-04 | 4.46E-03 | H2-Aa/Cd74/Irf1/I | 12 | aPD-L1 vs PBS | Up-regulated genes | CD45+ |
| BP | GO:0009123 | nucleoside monophosphate metabolic process | 4.10E-04 | 4.53E-03 | ATP8/Atp5e/Hspa | 11 | aPD-L1 vs PBS | Up-regulated genes | CD45+ |
| BP | GO:0002758 | innate immune response-activating signal transc | 4.25E-04 | 4.66E-03 | Irf1/Tnf/Rsad2/Ac | 8 | aPD-L1 vs PBS | Up-regulated genes | CD45+ |
| BP | GO:0032675 | regulation of interleukin-6 production | 4.25E-04 | 4.66E-03 | Tnf/Tnfaip3/Cd36/ | 8 | aPD-L1 vs PBS | Up-regulated genes | CD45+ |
| BP | GO:0046632 | alpha-beta T cell differentiation | 4.62E-04 | 5.05E-03 | Irf1/Rsad2/Malt1/ | 7 | aPD-L1 vs PBS | Up-regulated genes | CD45+ |
| BP | GO:0003013 | circulatory system process | 4.79E-04 | 5.22E-03 | Tnf/Ccl4/Mdm2/C | 13 | aPD-L1 vs PBS | Up-regulated genes | CD45+ |
| BP | GO:0031650 | regulation of heat generation | 4.89E-04 | 5.30E-03 | Tnf/Ccl5/Ptgs2 | 3 | aPD-L1 vs PBS | Up-regulated genes | CD45+ |
| BP | GO:0006754 | ATP biosynthetic process | 4.90E-04 | 5.30E-03 | ATP8/Atp5e/Hk3/ | 7 | aPD-L1 vs PBS | Up-regulated genes | CD45+ |
| BP | GO:0022618 | ribonucleoprotein complex assembly | 4.95E-04 | 5.33E-03 | Rps27/Rpl38/Rps | 10 | aPD-L1 vs PBS | Up-regulated genes | CD45+ |
| BP | GO:0002673 | regulation of acute inflammatory response | 5.04E-04 | 5.42E-03 | Tnf/H2-T23/Ccl5/ | 5 | aPD-L1 vs PBS | Up-regulated genes | CD45+ |
| BP | GO:0043281 | regulation of cysteine-type endopeptidase activi | 5.07E-04 | 5.42E-03 | Tnf/Nr4a1/Plaur/N | 9 | aPD-L1 vs PBS | Up-regulated genes | CD45+ |
| BP | GO:2001233 | regulation of apoptotic signaling pathway | 5.08E-04 | 5.42E-03 | Cd74/Lmna/Dnaja | 14 | aPD-L1 vs PBS | Up-regulated genes | CD45+ |
| BP | GO:0002507 | tolerance induction | 5.14E-04 | 5.45E-03 | Cd274/H2-T23/Ac | 4 | aPD-L1 vs PBS | Up-regulated genes | CD45+ |
| BP | GO:1903205 | regulation of hydrogen peroxide-induced cell de | 5.14E-04 | 5.45E-03 | Hk3/Nr4a3/Psap/ | 4 | aPD-L1 vs PBS | Up-regulated genes | CD45+ |
| BP | GO:0043434 | response to peptide hormone | 5.22E-04 | 5.49E-03 | Irf1/Nr4a1/Serpin | 11 | aPD-L1 vs PBS | Up-regulated genes | CD45+ |
| BP | GO:0070371 | ERK1 and ERK2 cascade | 5.22E-04 | 5.49E-03 | Cd74/Ccl4/Atf3/D | 11 | aPD-L1 vs PBS | Up-regulated genes | CD45+ |
| BP | GO:0001659 | temperature homeostasis | 5.44E-04 | 5.71E-03 | Tnf/Ccl5/Cd36/Kd | 8 | aPD-L1 vs PBS | Up-regulated genes | CD45+ |
| BP | GO:0045765 | regulation of angiogenesis | 5.59E-04 | 5.84E-03 | Ccl5/Cd36/Stat1/ | 11 | aPD-L1 vs PBS | Up-regulated genes | CD45+ |
| BP | GO:2001023 | regulation of response to drug | 5.63E-04 | 5.86E-03 | Tnf/Arg2/Hk3/Nr4 | 6 | aPD-L1 vs PBS | Up-regulated genes | CD45+ |
| BP | GO:0002825 | regulation of T-helper 1 type immune response | 5.90E-04 | 6.13E-03 | Irf1/H2-Ab1/Slc11 | 4 | aPD-L1 vs PBS | Up-regulated genes | CD45+ |
| BP | GO:0032479 | regulation of type I interferon production | 6.03E-04 | 6.24E-03 | Irf1/Acod1/Gbp4/ | 6 | aPD-L1 vs PBS | Up-regulated genes | CD45+ |
| BP | GO:0006801 | superoxide metabolic process | 6.08E-04 | 6.25E-03 | Tnf/Cd36/Sod2/N | 5 | aPD-L1 vs PBS | Up-regulated genes | CD45+ |
| BP | GO:0045620 | negative regulation of lymphocyte differentiation | 6.08E-04 | 6.25E-03 | Cd74/Irf1/Socs1/F | 5 | aPD-L1 vs PBS | Up-regulated genes | CD45+ |
| BP | GO:0042542 | response to hydrogen peroxide | 6.17E-04 | 6.32E-03 | Tnfaip3/Hk3/Nr4a | 7 | aPD-L1 vs PBS | Up-regulated genes | CD45+ |
| BP | GO:1904950 | negative regulation of establishment of protein lo | 6.25E-04 | 6.38E-03 | Dnaja1/Tnf/Arg2/ | 9 | aPD-L1 vs PBS | Up-regulated genes | CD45+ |
| BP | GO:0042532 | negative regulation of tyrosine phosphorylation | 6.30E-04 | 6.41E-03 | Irf1/Socs3/Socs1 | 3 | aPD-L1 vs PBS | Up-regulated genes | CD45+ |
| BP | GO:1902600 | proton transmembrane transport | 6.46E-04 | 6.56E-03 | ATP8/Atp5e/ATP6 | 6 | aPD-L1 vs PBS | Up-regulated genes | CD45+ |
| BP | GO:0009612 | response to mechanical stimulus | 6.52E-04 | 6.59E-03 | Tnf/Cd36/Nfkbia/ | 7 | aPD-L1 vs PBS | Up-regulated genes | CD45+ |
| BP | GO:0072538 | T-helper 17 type immune response | 6.74E-04 | 6.77E-03 | Malt1/Nlrp3/Arid5 | 4 | aPD-L1 vs PBS | Up-regulated genes | CD45+ |
| BP | GO:2001038 | regulation of cellular response to drug | 6.74E-04 | 6.77E-03 | Hk3/Nr4a3/Psap/ | 4 | aPD-L1 vs PBS | Up-regulated genes | CD45+ |
| BP | GO:0070301 | cellular response to hydrogen peroxide | 6.92E-04 | 6.93E-03 | Tnfaip3/Hk3/Nr4a | 6 | aPD-L1 vs PBS | Up-regulated genes | CD45+ |
| BP | GO:0036473 | cell death in response to oxidative stress | 7.40E-04 | 7.38E-03 | Slc7a11/Hk3/Nr4 | 6 | aPD-L1 vs PBS | Up-regulated genes | CD45+ |
| BP | GO:0071826 | ribonucleoprotein complex subunit organization | 7.41E-04 | 7.38E-03 | Rps27/Rpl38/Rps | 10 | aPD-L1 vs PBS | Up-regulated genes | CD45+ |
| BP | GO:0032743 | positive regulation of interleukin-2 production | 7.66E-04 | 7.54E-03 | Malt1/Cd83/Pde4 | 4 | aPD-L1 vs PBS | Up-regulated genes | CD45+ |
| BP | GO:0036474 | cell death in response to hydrogen peroxide | 7.66E-04 | 7.54E-03 | Hk3/Nr4a3/Psap/ | 4 | aPD-L1 vs PBS | Up-regulated genes | CD45+ |
| BP | GO:2000515 | negative regulation of CD4-positive, alpha-beta | 7.66E-04 | 7.54E-03 | Irf1/Cd274/Arg2/S | 4 | aPD-L1 vs PBS | Up-regulated genes | CD45+ |
| BP | GO:0030218 | erythrocyte differentiation | 7.68E-04 | 7.54E-03 | Inhba/B2m/Rps1 | 7 | aPD-L1 vs PBS | Up-regulated genes | CD45+ |
| BP | GO:0032606 | type I interferon production | 7.90E-04 | 7.73E-03 | Irf1/Acod1/Gbp4/ | 6 | aPD-L1 vs PBS | Up-regulated genes | CD45+ |
| BP | GO:0001505 | regulation of neurotransmitter levels | 8.00E-04 | 7.81E-03 | Tnf/Slc7a11/Cd3 | 11 | aPD-L1 vs PBS | Up-regulated genes | CD45+ |
| BP | GO:0006935 | chemotaxis | 8.02E-04 | 7.81E-03 | Cd74/Cxcl9/Nr4a | 15 | aPD-L1 vs PBS | Up-regulated genes | CD45+ |
| BP | GO:0050708 | regulation of protein secretion | 8.10E-04 | 7.83E-03 | Cd274/Tnf/Arg2/F | 14 | aPD-L1 vs PBS | Up-regulated genes | CD45+ |
| BP | GO:0009206 | purine ribonucleoside triphosphate biosynthetic | 8.10E-04 | 7.83E-03 | ATP8/Atp5e/Hk3/ | 7 | aPD-L1 vs PBS | Up-regulated genes | CD45+ |
| BP | GO:0046651 | lymphocyte proliferation | 8.26E-04 | 7.96E-03 | H2-Aa/Cd74/Irf1/ | 11 | aPD-L1 vs PBS | Up-regulated genes | CD45+ |
| BP | GO:0001910 | regulation of leukocyte mediated cytotoxicity | 8.43E-04 | 8.10E-03 | H2-T23/B2m/H2- | 6 | aPD-L1 vs PBS | Up-regulated genes | CD45+ |
| BP | GO:0009145 | purine nucleoside triphosphate biosynthetic pro | 8.54E-04 | 8.18E-03 | ATP8/Atp5e/Hk3/ | 7 | aPD-L1 vs PBS | Up-regulated genes | CD45+ |
| BP | GO:0042330 | taxis | 8.60E-04 | 8.21E-03 | Cd74/Cxcl9/Nr4a | 15 | aPD-L1 vs PBS | Up-regulated genes | CD45+ |
| BP | GO:0045766 | positive regulation of angiogenesis | 8.66E-04 | 8.21E-03 | Ccl5/Hk2/Klf4/Itg | 8 | aPD-L1 vs PBS | Up-regulated genes | CD45+ |
| BP | GO:1903555 | regulation of tumor necrosis factor superfamily c | 8.66E-04 | 8.21E-03 | Cd274/Arg2/Ccl4 | 8 | aPD-L1 vs PBS | Up-regulated genes | CD45+ |
| BP | GO:0018108 | peptidyl-tyrosine phosphorylation | 9.36E-04 | 8.83E-03 | Cd74/Irf1/Tnf/Sam | 11 | aPD-L1 vs PBS | Up-regulated genes | CD45+ |
| BP | GO:0051384 | response to glucocorticoid | 9.36E-04 | 8.83E-03 | Tnf/ND3/Zfp36/Zf | 5 | aPD-L1 vs PBS | Up-regulated genes | CD45+ |
| BP | GO:0009201 | ribonucleoside triphosphate biosynthetic proces | 9.47E-04 | 8.90E-03 | ATP8/Atp5e/Hk3/ | 7 | aPD-L1 vs PBS | Up-regulated genes | CD45+ |
| BP | GO:0048525 | negative regulation of viral process | 9.57E-04 | 8.97E-03 | Tnf/Rsad2/Ccl4/C | 6 | aPD-L1 vs PBS | Up-regulated genes | CD45+ |
| BP | GO:1904996 | positive regulation of leukocyte adhesion to vas | 9.82E-04 | 9.18E-03 | Tnf/Icam1/Itgb2 | 3 | aPD-L1 vs PBS | Up-regulated genes | CD45+ |
| BP | GO:0043122 | regulation of I-kappaB kinase/NF-kappaB signal | 9.87E-04 | 9.20E-03 | Cd74/Tnf/Tnfaip3 | 8 | aPD-L1 vs PBS | Up-regulated genes | CD45+ |
| BP | GO:0042326 | negative regulation of phosphorylation | 1.01E-03 | 9.38E-03 | Irf1/Dnaja1/Sams | 14 | aPD-L1 vs PBS | Up-regulated genes | CD45+ |
| BP | GO:0031960 | response to corticosteroid | 1.02E-03 | 9.38E-03 | Tnf/ND3/Zfp36/Zf | 5 | aPD-L1 vs PBS | Up-regulated genes | CD45+ |
| BP | GO:0032722 | positive regulation of chemokine production | 1.02E-03 | 9.38E-03 | Cd74/Tnf/Ccl5/Eg | 5 | aPD-L1 vs PBS | Up-regulated genes | CD45+ |
| BP | GO:0032755 | positive regulation of interleukin-6 production | 1.02E-03 | 9.38E-03 | Tnf/Cd36/Il1f9/Ari | 6 | aPD-L1 vs PBS | Up-regulated genes | CD45+ |
| BP | GO:0018212 | peptidyl-tyrosine modification | 1.03E-03 | 9.42E-03 | Cd74/Irf1/Tnf/Sam | 11 | aPD-L1 vs PBS | Up-regulated genes | CD45+ |
| BP | GO:0071706 | tumor necrosis factor superfamily cytokine produ | 1.03E-03 | 9.43E-03 | Cd274/Arg2/Ccl4 | 8 | aPD-L1 vs PBS | Up-regulated genes | CD45+ |
| BP | GO:0048871 | multicellular organismal homeostasis | 1.06E-03 | 9.66E-03 | Tnf/Fcgr4/Ccl5/Cc | 13 | aPD-L1 vs PBS | Up-regulated genes | CD45+ |
| BP | GO:0044057 | regulation of system process | 1.09E-03 | 9.88E-03 | Lmna/Inhba/Tnf/A | 14 | aPD-L1 vs PBS | Up-regulated genes | CD45+ |
| BP | GO:0009611 | response to wounding | 1.11E-03 | 1.01E-02 | Ifrd1/Dusp10/Slc7 | 13 | aPD-L1 vs PBS | Up-regulated genes | CD45+ |
| BP | GO:2000116 | regulation of cysteine-type endopeptidase activi | 1.12E-03 | 1.01E-02 | Tnf/Nr4a1/Plaur/N | 9 | aPD-L1 vs PBS | Up-regulated genes | CD45+ |
| BP | GO:0002763 | positive regulation of myeloid leukocyte different | 1.19E-03 | 1.06E-02 | Cd74/Tnf/Ccl5/Cs | 5 | aPD-L1 vs PBS | Up-regulated genes | CD45+ |
| BP | GO:0002830 | positive regulation of type 2 immune response | 1.20E-03 | 1.06E-02 | Cd74/Rsad2/Nlrp | 3 | aPD-L1 vs PBS | Up-regulated genes | CD45+ |
| BP | GO:0019430 | removal of superoxide radicals | 1.20E-03 | 1.06E-02 | Tnf/Cd36/Sod2 | 3 | aPD-L1 vs PBS | Up-regulated genes | CD45+ |
| BP | GO:0071157 | negative regulation of cell cycle arrest | 1.20E-03 | 1.06E-02 | Hsp90ab1/Mdm2 | 3 | aPD-L1 vs PBS | Up-regulated genes | CD45+ |
| BP | GO:0071498 | cellular response to fluid shear stress | 1.20E-03 | 1.06E-02 | Klf4/Klf2/Ptgs2 | 3 | aPD-L1 vs PBS | Up-regulated genes | CD45+ |
| BP | GO:2000319 | regulation of T-helper 17 cell differentiation | 1.20E-03 | 1.06E-02 | Malt1/Nlrp3/Smac | 3 | aPD-L1 vs PBS | Up-regulated genes | CD45+ |

| | | | | | | | | | |
|---|---|---|---|---|---|---|---|---|---|
| BP | GO:0045622 | regulation of T-helper cell differentiation | 1.22E-03 | 1.08E-02 | Irf1/Malt1/Nlrp3/S | 4 | aPD-L1 vs PBS | Up-regulated genes | CD45+ |
| BP | GO:0061844 | antimicrobial humoral immune response mediat | 1.22E-03 | 1.08E-02 | Rpl39/Cxcl9/Rpl3 | 4 | aPD-L1 vs PBS | Up-regulated genes | CD45+ |
| BP | GO:0002218 | activation of innate immune response | 1.27E-03 | 1.12E-02 | Irf1/Tnf/Rsad2/Ac | 8 | aPD-L1 vs PBS | Up-regulated genes | CD45+ |
| BP | GO:0070372 | regulation of ERK1 and ERK2 cascade | 1.27E-03 | 1.12E-02 | Cd74/Ccl4/Atf3/D | 10 | aPD-L1 vs PBS | Up-regulated genes | CD45+ |
| BP | GO:0007369 | gastrulation | 1.28E-03 | 1.12E-02 | Inhba/Tnfaip3/Nr4 | 7 | aPD-L1 vs PBS | Up-regulated genes | CD45+ |
| BP | GO:0034121 | regulation of toll-like receptor signaling pathway | 1.28E-03 | 1.12E-02 | Irf1/Rsad2/Acod1 | 5 | aPD-L1 vs PBS | Up-regulated genes | CD45+ |
| BP | GO:2000106 | regulation of leukocyte apoptotic process | 1.29E-03 | 1.12E-02 | Cd74/Cd274/Arg2 | 6 | aPD-L1 vs PBS | Up-regulated genes | CD45+ |
| BP | GO:0006163 | purine nucleotide metabolic process | 1.30E-03 | 1.12E-02 | ATP8/Atp5e/Hspa | 13 | aPD-L1 vs PBS | Up-regulated genes | CD45+ |
| BP | GO:0046942 | carboxylic acid transport | 1.30E-03 | 1.12E-02 | Tnf/Arg2/Slc7a11 | 9 | aPD-L1 vs PBS | Up-regulated genes | CD45+ |
| BP | GO:0015849 | organic acid transport | 1.34E-03 | 1.16E-02 | Tnf/Arg2/Slc7a11 | 9 | aPD-L1 vs PBS | Up-regulated genes | CD45+ |
| BP | GO:0034599 | cellular response to oxidative stress | 1.36E-03 | 1.17E-02 | Tnf/Slc7a11/Tnfai | 10 | aPD-L1 vs PBS | Up-regulated genes | CD45+ |
| BP | GO:1902041 | regulation of extrinsic apoptotic signaling pathwa | 1.36E-03 | 1.17E-02 | Atf3/Tnfaip3/Icam | 4 | aPD-L1 vs PBS | Up-regulated genes | CD45+ |
| BP | GO:0042509 | regulation of tyrosine phosphorylation of STAT p | 1.38E-03 | 1.18E-02 | Irf1/Tnf/Ccl5/Socs | 5 | aPD-L1 vs PBS | Up-regulated genes | CD45+ |
| BP | GO:0000188 | inactivation of MAPK activity | 1.44E-03 | 1.22E-02 | Dusp10/Prkca/Du | 3 | aPD-L1 vs PBS | Up-regulated genes | CD45+ |
| BP | GO:0010743 | regulation of macrophage derived foam cell diffe | 1.44E-03 | 1.22E-02 | Cd36/Nfkbia/Csf1 | 3 | aPD-L1 vs PBS | Up-regulated genes | CD45+ |
| BP | GO:0061081 | positive regulation of myeloid leukocyte cytokine | 1.44E-03 | 1.22E-02 | Cd74/Cd36/Nr4a | 3 | aPD-L1 vs PBS | Up-regulated genes | CD45+ |
| BP | GO:0009127 | purine nucleoside monophosphate biosynthetic | 1.47E-03 | 1.24E-02 | ATP8/Atp5e/Hk3/ | 7 | aPD-L1 vs PBS | Up-regulated genes | CD45+ |
| BP | GO:0009168 | purine ribonucleoside monophosphate biosynth | 1.47E-03 | 1.24E-02 | ATP8/Atp5e/Hk3/ | 7 | aPD-L1 vs PBS | Up-regulated genes | CD45+ |
| BP | GO:0046640 | regulation of alpha-beta T cell proliferation | 1.51E-03 | 1.27E-02 | Irf1/Cd274/Arg2/H | 4 | aPD-L1 vs PBS | Up-regulated genes | CD45+ |
| BP | GO:2001243 | negative regulation of intrinsic apoptotic signalin | 1.54E-03 | 1.29E-02 | Cd74/Dnaja1/Pla | 6 | aPD-L1 vs PBS | Up-regulated genes | CD45+ |
| BP | GO:0002757 | immune response-activating signal transduction | 1.59E-03 | 1.33E-02 | Irf1/Tnf/Rsad2/Ac | 11 | aPD-L1 vs PBS | Up-regulated genes | CD45+ |
| BP | GO:0008625 | extrinsic apoptotic signaling pathway via death c | 1.60E-03 | 1.33E-02 | Tnf/Atf3/Tnfaip3/Ic | 5 | aPD-L1 vs PBS | Up-regulated genes | CD45+ |
| BP | GO:0042108 | positive regulation of cytokine biosynthetic proce | 1.60E-03 | 1.33E-02 | Irf1/Tnf/Rel/Egr1/Il | 5 | aPD-L1 vs PBS | Up-regulated genes | CD45+ |
| BP | GO:0009142 | nucleoside triphosphate biosynthetic process | 1.61E-03 | 1.34E-02 | ATP8/Atp5e/Hk3/ | 7 | aPD-L1 vs PBS | Up-regulated genes | CD45+ |
| BP | GO:0048545 | response to steroid hormone | 1.62E-03 | 1.34E-02 | Dnaja1/Tnf/Acod1 | 8 | aPD-L1 vs PBS | Up-regulated genes | CD45+ |
| BP | GO:0002832 | negative regulation of response to biotic stimulu | 1.67E-03 | 1.38E-02 | Arg2/Acod1/Tnfai | 4 | aPD-L1 vs PBS | Up-regulated genes | CD45+ |
| BP | GO:0031664 | regulation of lipopolysaccharide-mediated signa | 1.71E-03 | 1.40E-02 | Acod1/Tnfaip3/Pr | 3 | aPD-L1 vs PBS | Up-regulated genes | CD45+ |
| BP | GO:0032656 | regulation of interleukin-13 production | 1.71E-03 | 1.40E-02 | Arg2/H2-T23/Nlrp | 3 | aPD-L1 vs PBS | Up-regulated genes | CD45+ |
| BP | GO:0071450 | cellular response to oxygen radical | 1.71E-03 | 1.40E-02 | Tnf/Cd36/Sod2 | 3 | aPD-L1 vs PBS | Up-regulated genes | CD45+ |
| BP | GO:0071451 | cellular response to superoxide | 1.71E-03 | 1.40E-02 | Tnf/Cd36/Sod2 | 3 | aPD-L1 vs PBS | Up-regulated genes | CD45+ |
| BP | GO:0009156 | ribonucleoside monophosphate biosynthetic pro | 1.76E-03 | 1.44E-02 | ATP8/Atp5e/Hk3/ | 7 | aPD-L1 vs PBS | Up-regulated genes | CD45+ |
| BP | GO:0032944 | regulation of mononuclear cell proliferation | 1.83E-03 | 1.49E-02 | H2-Aa/Cd74/Irf1/ | 9 | aPD-L1 vs PBS | Up-regulated genes | CD45+ |
| BP | GO:0007260 | tyrosine phosphorylation of STAT protein | 1.84E-03 | 1.49E-02 | Irf1/Tnf/Ccl5/Socs | 5 | aPD-L1 vs PBS | Up-regulated genes | CD45+ |
| BP | GO:0042273 | ribosomal large subunit biogenesis | 1.84E-03 | 1.49E-02 | Rpl35a/Rpl38/Rp | 5 | aPD-L1 vs PBS | Up-regulated genes | CD45+ |
| BP | GO:0010563 | negative regulation of phosphorus metabolic pro | 1.85E-03 | 1.49E-02 | Irf1/Dnaja1/Tnf/S | 15 | aPD-L1 vs PBS | Up-regulated genes | CD45+ |
| BP | GO:0045936 | negative regulation of phosphate metabolic proc | 1.85E-03 | 1.49E-02 | Irf1/Dnaja1/Tnf/S | 15 | aPD-L1 vs PBS | Up-regulated genes | CD45+ |
| BP | GO:0052548 | regulation of endopeptidase activity | 1.91E-03 | 1.53E-02 | Tnf/Nr4a1/Plaur/ | 10 | aPD-L1 vs PBS | Up-regulated genes | CD45+ |
| BP | GO:0050715 | positive regulation of cytokine secretion | 1.93E-03 | 1.55E-02 | Cd274/Tnf/Cd36/ | 7 | aPD-L1 vs PBS | Up-regulated genes | CD45+ |
| BP | GO:0098869 | cellular oxidant detoxification | 2.01E-03 | 1.61E-02 | Tnf/Cd36/Sod2 | 3 | aPD-L1 vs PBS | Up-regulated genes | CD45+ |
| BP | GO:0051090 | regulation of DNA-binding transcription factor ac | 2.02E-03 | 1.61E-02 | Tnf/Ifrd1/Acod1/Tr | 12 | aPD-L1 vs PBS | Up-regulated genes | CD45+ |
| BP | GO:0046633 | alpha-beta T cell proliferation | 2.02E-03 | 1.61E-02 | Irf1/Cd274/Arg2/H | 4 | aPD-L1 vs PBS | Up-regulated genes | CD45+ |
| BP | GO:0035821 | modification of morphology or physiology of othe | 2.12E-03 | 1.68E-02 | Rpl30/Ccl4/Hspa | 6 | aPD-L1 vs PBS | Up-regulated genes | CD45+ |
| BP | GO:0009306 | protein secretion | 2.18E-03 | 1.72E-02 | Cd274/Tnf/Arg2/F | 15 | aPD-L1 vs PBS | Up-regulated genes | CD45+ |
| BP | GO:0051224 | negative regulation of protein transport | 2.19E-03 | 1.73E-02 | Tnf/Arg2/Rsad2/T | 8 | aPD-L1 vs PBS | Up-regulated genes | CD45+ |
| BP | GO:0009124 | nucleoside monophosphate biosynthetic proces | 2.20E-03 | 1.73E-02 | ATP8/Atp5e/Hk3/ | 7 | aPD-L1 vs PBS | Up-regulated genes | CD45+ |
| BP | GO:0002764 | immune response-regulating signaling pathway | 2.21E-03 | 1.73E-02 | Irf1/Tnf/Rsad2/Ac | 11 | aPD-L1 vs PBS | Up-regulated genes | CD45+ |
| BP | GO:0030490 | maturation of SSU-rRNA | 2.22E-03 | 1.74E-02 | Rps21/Rps28/Rp | 4 | aPD-L1 vs PBS | Up-regulated genes | CD45+ |
| BP | GO:2001236 | regulation of extrinsic apoptotic signaling pathwa | 2.29E-03 | 1.79E-02 | Lmna/Inhba/Tnf/A | 7 | aPD-L1 vs PBS | Up-regulated genes | CD45+ |
| BP | GO:0032616 | interleukin-13 production | 2.34E-03 | 1.81E-02 | Arg2/H2-T23/Nlrp | 3 | aPD-L1 vs PBS | Up-regulated genes | CD45+ |
| BP | GO:0032897 | negative regulation of viral transcription | 2.34E-03 | 1.81E-02 | Ccl4/Ccl5/Zfp36 | 3 | aPD-L1 vs PBS | Up-regulated genes | CD45+ |
| BP | GO:0045624 | positive regulation of T-helper cell differentiation | 2.34E-03 | 1.81E-02 | Irf1/Malt1/Nlrp3 | 3 | aPD-L1 vs PBS | Up-regulated genes | CD45+ |
| BP | GO:0070498 | interleukin-1-mediated signaling pathway | 2.34E-03 | 1.81E-02 | Egr1/Il1a/Il1rn | 3 | aPD-L1 vs PBS | Up-regulated genes | CD45+ |
| BP | GO:0032649 | regulation of interferon-gamma production | 2.35E-03 | 1.81E-02 | Cd274/Tnf/H2-Q7 | 6 | aPD-L1 vs PBS | Up-regulated genes | CD45+ |
| BP | GO:0043123 | positive regulation of I-kappaB kinase/NF-kappa | 2.35E-03 | 1.81E-02 | Cd74/Tnf/Cd36/Il | 6 | aPD-L1 vs PBS | Up-regulated genes | CD45+ |
| BP | GO:0070663 | regulation of leukocyte proliferation | 2.38E-03 | 1.83E-02 | H2-Aa/Cd74/Irf1/ | 9 | aPD-L1 vs PBS | Up-regulated genes | CD45+ |
| BP | GO:0043154 | negative regulation of cysteine-type endopeptid | 2.39E-03 | 1.83E-02 | Nr4a1/Plaur/Mdm | 5 | aPD-L1 vs PBS | Up-regulated genes | CD45+ |
| BP | GO:0050702 | interleukin-1 beta secretion | 2.43E-03 | 1.85E-02 | Arg2/Tnfaip3/Cd3 | 4 | aPD-L1 vs PBS | Up-regulated genes | CD45+ |
| BP | GO:0051591 | response to cAMP | 2.43E-03 | 1.85E-02 | Stat1/Jund/Junb/ | 4 | aPD-L1 vs PBS | Up-regulated genes | CD45+ |
| BP | GO:0072521 | purine-containing compound metabolic process | 2.45E-03 | 1.86E-02 | ATP8/Atp5e/Hspa | 13 | aPD-L1 vs PBS | Up-regulated genes | CD45+ |
| BP | GO:0019058 | viral life cycle | 2.53E-03 | 1.92E-02 | Cd74/Tnf/Rsad2/ | 8 | aPD-L1 vs PBS | Up-regulated genes | CD45+ |
| BP | GO:0001961 | positive regulation of cytokine-mediated signalin | 2.65E-03 | 2.00E-02 | Cd74/Irgm1/Zbp1 | 4 | aPD-L1 vs PBS | Up-regulated genes | CD45+ |
| BP | GO:0032102 | negative regulation of response to external stim | 2.70E-03 | 2.02E-02 | Arg2/Dusp10/Aco | 10 | aPD-L1 vs PBS | Up-regulated genes | CD45+ |
| BP | GO:0000303 | response to superoxide | 2.71E-03 | 2.02E-02 | Tnf/Cd36/Sod2 | 3 | aPD-L1 vs PBS | Up-regulated genes | CD45+ |
| BP | GO:0046823 | negative regulation of nucleocytoplasmic transp | 2.71E-03 | 2.02E-02 | Cd36/Gbp4/Txn1 | 3 | aPD-L1 vs PBS | Up-regulated genes | CD45+ |
| BP | GO:0031397 | negative regulation of protein ubiquitination | 2.71E-03 | 2.02E-02 | Dnaja1/Tnfaip3/G | 5 | aPD-L1 vs PBS | Up-regulated genes | CD45+ |
| BP | GO:0032088 | negative regulation of NF-kappaB transcription f | 2.71E-03 | 2.02E-02 | Acod1/Tnfaip3/Nf | 5 | aPD-L1 vs PBS | Up-regulated genes | CD45+ |
| BP | GO:0032729 | positive regulation of interferon-gamma producti | 2.71E-03 | 2.02E-02 | Tnf/H2-Q7/Pde4b | 5 | aPD-L1 vs PBS | Up-regulated genes | CD45+ |
| BP | GO:0006364 | rRNA processing | 2.81E-03 | 2.09E-02 | Rps21/Rpl35a/Rp | 8 | aPD-L1 vs PBS | Up-regulated genes | CD45+ |
| BP | GO:0002437 | inflammatory response to antigenic stimulus | 2.88E-03 | 2.13E-02 | Tnf/H2-T23/Icam1 | 4 | aPD-L1 vs PBS | Up-regulated genes | CD45+ |
| BP | GO:0015807 | L-amino acid transport | 2.88E-03 | 2.13E-02 | Tnf/Arg2/Slc7a11 | 4 | aPD-L1 vs PBS | Up-regulated genes | CD45+ |
| BP | GO:0006913 | nucleocytoplasmic transport | 2.93E-03 | 2.16E-02 | Lmna/Mdm2/Hspa | 10 | aPD-L1 vs PBS | Up-regulated genes | CD45+ |
| BP | GO:0051169 | nuclear transport | 2.93E-03 | 2.16E-02 | Lmna/Mdm2/Hspa | 10 | aPD-L1 vs PBS | Up-regulated genes | CD45+ |
| BP | GO:0045598 | regulation of fat cell differentiation | 3.00E-03 | 2.21E-02 | Tnf/Dusp10/Zfp36 | 6 | aPD-L1 vs PBS | Up-regulated genes | CD45+ |
| BP | GO:1903707 | negative regulation of hemopoiesis | 3.05E-03 | 2.23E-02 | Cd74/Irf1/Nfkbia/ | 7 | aPD-L1 vs PBS | Up-regulated genes | CD45+ |
| BP | GO:0000305 | response to oxygen radical | 3.10E-03 | 2.26E-02 | Tnf/Cd36/Sod2 | 3 | aPD-L1 vs PBS | Up-regulated genes | CD45+ |
| BP | GO:0034405 | response to fluid shear stress | 3.10E-03 | 2.26E-02 | Klf4/Klf2/Ptgs2 | 3 | aPD-L1 vs PBS | Up-regulated genes | CD45+ |
| BP | GO:1902042 | negative regulation of extrinsic apoptotic signali | 3.10E-03 | 2.26E-02 | Tnfaip3/Icam1/Hm | 3 | aPD-L1 vs PBS | Up-regulated genes | CD45+ |
| BP | GO:0090257 | regulation of muscle system process | 3.12E-03 | 2.26E-02 | Lmna/Arg2/Nr4a3 | 8 | aPD-L1 vs PBS | Up-regulated genes | CD45+ |
| BP | GO:0050829 | defense response to Gram-negative bacterium | 3.13E-03 | 2.26E-02 | Rpl30/Rps19/Slc | 4 | aPD-L1 vs PBS | Up-regulated genes | CD45+ |
| BP | GO:1904705 | regulation of vascular smooth muscle cell prolife | 3.13E-03 | 2.26E-02 | Tnf/Mdm2/Nr4a3/ | 4 | aPD-L1 vs PBS | Up-regulated genes | CD45+ |
| BP | GO:1990874 | vascular smooth muscle cell proliferation | 3.13E-03 | 2.26E-02 | Tnf/Mdm2/Nr4a3/ | 4 | aPD-L1 vs PBS | Up-regulated genes | CD45+ |
| BP | GO:0048514 | blood vessel morphogenesis | 3.15E-03 | 2.27E-02 | Tnfaip2/Nr4a1/Cc | 15 | aPD-L1 vs PBS | Up-regulated genes | CD45+ |
| BP | GO:0051348 | negative regulation of transferase activity | 3.24E-03 | 2.33E-02 | Dnaja1/Dusp10/T | 9 | aPD-L1 vs PBS | Up-regulated genes | CD45+ |
| BP | GO:0032611 | interleukin-1 beta production | 3.25E-03 | 2.33E-02 | Arg2/Tnfaip3/Cd3 | 5 | aPD-L1 vs PBS | Up-regulated genes | CD45+ |
| BP | GO:0097193 | intrinsic apoptotic signaling pathway | 3.36E-03 | 2.40E-02 | Cd74/Dnaja1/Crip | 10 | aPD-L1 vs PBS | Up-regulated genes | CD45+ |
| BP | GO:0042088 | T-helper 1 type immune response | 3.39E-03 | 2.42E-02 | Irf1/H2-Ab1/Slc11 | 4 | aPD-L1 vs PBS | Up-regulated genes | CD45+ |
| BP | GO:2000117 | negative regulation of cysteine-type endopeptid | 3.44E-03 | 2.45E-02 | Nr4a1/Plaur/Mdm | 5 | aPD-L1 vs PBS | Up-regulated genes | CD45+ |
| BP | GO:0035743 | CD4-positive, alpha-beta T cell cytokine product | 3.53E-03 | 2.50E-02 | Rsad2/Nlrp3/Arid | 3 | aPD-L1 vs PBS | Up-regulated genes | CD45+ |
| BP | GO:0046426 | negative regulation of JAK-STAT cascade | 3.53E-03 | 2.50E-02 | Irf1/Socs3/Socs1 | 3 | aPD-L1 vs PBS | Up-regulated genes | CD45+ |

| | | | | | | | | | |
|---|---|---|---|---|---|---|---|---|---|
| BP | GO:1904893 | negative regulation of STAT cascade | 3.53E-03 | 2.50E-02 | Irf1/Socs3/Socs1 | 3 | aPD-L1 vs PBS | Up-regulated genes | CD45+ |
| BP | GO:0032680 | regulation of tumor necrosis factor production | 3.56E-03 | 2.50E-02 | Arg2/Ccl4/H2-T23 | 7 | aPD-L1 vs PBS | Up-regulated genes | CD45+ |
| BP | GO:0045333 | cellular respiration | 3.56E-03 | 2.50E-02 | Nr4a3/ND1/Cox7 | 7 | aPD-L1 vs PBS | Up-regulated genes | CD45+ |
| BP | GO:0071887 | leukocyte apoptotic process | 3.61E-03 | 2.54E-02 | Cd74/Cd274/Arg2 | 6 | aPD-L1 vs PBS | Up-regulated genes | CD45+ |
| BP | GO:0007498 | mesoderm development | 3.65E-03 | 2.55E-02 | Rpl38/Inhba/Nr4a | 5 | aPD-L1 vs PBS | Up-regulated genes | CD45+ |
| BP | GO:0043618 | regulation of transcription from RNA polymerase | 3.67E-03 | 2.56E-02 | Atf3/Cd36/Egr1/Kl | 4 | aPD-L1 vs PBS | Up-regulated genes | CD45+ |
| BP | GO:0001909 | leukocyte mediated cytotoxicity | 3.78E-03 | 2.63E-02 | H2-T23/B2m/H2-I | 6 | aPD-L1 vs PBS | Up-regulated genes | CD45+ |
| BP | GO:0006164 | purine nucleotide biosynthetic process | 3.93E-03 | 2.73E-02 | ATP8/Atp5e/Hk3/I | 8 | aPD-L1 vs PBS | Up-regulated genes | CD45+ |
| BP | GO:0002377 | immunoglobulin production | 3.95E-03 | 2.74E-02 | H2-Ab1/Tnf/H2-T | 6 | aPD-L1 vs PBS | Up-regulated genes | CD45+ |
| BP | GO:0045661 | regulation of myoblast differentiation | 3.96E-03 | 2.74E-02 | Cxcl9/Tnf/Il1f9/Btg | 4 | aPD-L1 vs PBS | Up-regulated genes | CD45+ |
| BP | GO:0050701 | interleukin-1 secretion | 3.96E-03 | 2.74E-02 | Arg2/Tnfaip3/Cd3 | 4 | aPD-L1 vs PBS | Up-regulated genes | CD45+ |
| BP | GO:0032640 | tumor necrosis factor production | 3.98E-03 | 2.74E-02 | Arg2/Ccl4/H2-T2 | 7 | aPD-L1 vs PBS | Up-regulated genes | CD45+ |
| BP | GO:0045648 | positive regulation of erythrocyte differentiation | 4.00E-03 | 2.74E-02 | Inhba/Stat1/Hmgb | 3 | aPD-L1 vs PBS | Up-regulated genes | CD45+ |
| BP | GO:0045730 | respiratory burst | 4.00E-03 | 2.74E-02 | Dusp10/Rps19/Sl | 3 | aPD-L1 vs PBS | Up-regulated genes | CD45+ |
| BP | GO:1903792 | negative regulation of anion transport | 4.00E-03 | 2.74E-02 | Tnf/Il1rn/Prkca | 3 | aPD-L1 vs PBS | Up-regulated genes | CD45+ |
| BP | GO:0022904 | respiratory electron transport chain | 4.08E-03 | 2.79E-02 | Cox7c/ND4L/CYT | 5 | aPD-L1 vs PBS | Up-regulated genes | CD45+ |
| BP | GO:0009165 | nucleotide biosynthetic process | 4.09E-03 | 2.79E-02 | ATP8/Atp5e/Hk3/I | 9 | aPD-L1 vs PBS | Up-regulated genes | CD45+ |
| BP | GO:0010951 | negative regulation of endopeptidase activity | 4.13E-03 | 2.80E-02 | Nr4a1/Plaur/Mdm | 6 | aPD-L1 vs PBS | Up-regulated genes | CD45+ |
| BP | GO:0032609 | interferon-gamma production | 4.13E-03 | 2.80E-02 | Cd274/Tnf/H2-Q7 | 6 | aPD-L1 vs PBS | Up-regulated genes | CD45+ |
| BP | GO:0009117 | nucleotide metabolic process | 4.23E-03 | 2.87E-02 | ATP8/Atp5e/Hspa | 14 | aPD-L1 vs PBS | Up-regulated genes | CD45+ |
| BP | GO:0001525 | angiogenesis | 4.24E-03 | 2.87E-02 | Tnfaip2/Nr4a1/Cc | 13 | aPD-L1 vs PBS | Up-regulated genes | CD45+ |
| BP | GO:0051051 | negative regulation of transport | 4.41E-03 | 2.98E-02 | Cd74/Tnf/Arg2/Rs | 13 | aPD-L1 vs PBS | Up-regulated genes | CD45+ |
| BP | GO:0043902 | positive regulation of multi-organism process | 4.45E-03 | 2.99E-02 | Cd74/Inhba/Acod | 7 | aPD-L1 vs PBS | Up-regulated genes | CD45+ |
| BP | GO:0050678 | regulation of epithelial cell proliferation | 4.46E-03 | 3.00E-02 | Tnf/Nr4a1/Dusp1 | 10 | aPD-L1 vs PBS | Up-regulated genes | CD45+ |
| BP | GO:0032480 | negative regulation of type I interferon productio | 4.50E-03 | 3.00E-02 | Acod1/Gbp4/Rel | 3 | aPD-L1 vs PBS | Up-regulated genes | CD45+ |
| BP | GO:0032673 | regulation of interleukin-4 production | 4.50E-03 | 3.00E-02 | H2-T23/Cd83/Nlr | 3 | aPD-L1 vs PBS | Up-regulated genes | CD45+ |
| BP | GO:0046639 | negative regulation of alpha-beta T cell differenti | 4.50E-03 | 3.00E-02 | Irf1/Socs1/Smad7 | 3 | aPD-L1 vs PBS | Up-regulated genes | CD45+ |
| BP | GO:1903321 | negative regulation of protein modification by sm | 4.54E-03 | 3.02E-02 | Dnaja1/Tnfaip3/G | 5 | aPD-L1 vs PBS | Up-regulated genes | CD45+ |
| BP | GO:0050732 | negative regulation of peptidyl-tyrosine phospho | 4.59E-03 | 3.04E-02 | Irf1/Samsn1/Socs | 4 | aPD-L1 vs PBS | Up-regulated genes | CD45+ |
| BP | GO:0060760 | positive regulation of response to cytokine stimu | 4.59E-03 | 3.04E-02 | Cd74/Irgm1/Zbp1 | 4 | aPD-L1 vs PBS | Up-regulated genes | CD45+ |
| BP | GO:0072522 | purine-containing compound biosynthetic proces | 4.60E-03 | 3.04E-02 | ATP8/Atp5e/Hk3/I | 8 | aPD-L1 vs PBS | Up-regulated genes | CD45+ |
| BP | GO:0062013 | positive regulation of small molecule metabolic p | 4.70E-03 | 3.10E-02 | Tnf/Nr4a3/Nfkb1/ | 6 | aPD-L1 vs PBS | Up-regulated genes | CD45+ |
| BP | GO:0006753 | nucleoside phosphate metabolic process | 4.74E-03 | 3.12E-02 | ATP8/Atp5e/Hspa | 14 | aPD-L1 vs PBS | Up-regulated genes | CD45+ |
| BP | GO:0071375 | cellular response to peptide hormone stimulus | 4.75E-03 | 3.12E-02 | Irf1/Nr4a1/Nr4a3/ | 8 | aPD-L1 vs PBS | Up-regulated genes | CD45+ |
| BP | GO:0046328 | regulation of JNK cascade | 4.78E-03 | 3.12E-02 | Dnaja1/Tnf/Dusp | 7 | aPD-L1 vs PBS | Up-regulated genes | CD45+ |
| BP | GO:0022900 | electron transport chain | 4.79E-03 | 3.12E-02 | Cox7c/ND4L/CYT | 5 | aPD-L1 vs PBS | Up-regulated genes | CD45+ |
| BP | GO:0044070 | regulation of anion transport | 4.79E-03 | 3.12E-02 | Tnf/Arg2/Il1a/Il1rn | 5 | aPD-L1 vs PBS | Up-regulated genes | CD45+ |
| BP | GO:2001237 | negative regulation of extrinsic apoptotic signali | 4.79E-03 | 3.12E-02 | Lmna/Tnf/Tnfaip3 | 5 | aPD-L1 vs PBS | Up-regulated genes | CD45+ |
| BP | GO:1901293 | nucleoside phosphate biosynthetic process | 4.84E-03 | 3.14E-02 | ATP8/Atp5e/Hk3/I | 9 | aPD-L1 vs PBS | Up-regulated genes | CD45+ |
| BP | GO:0072539 | T-helper 17 cell differentiation | 5.03E-03 | 3.27E-02 | Malt1/Nlrp3/Smac | 3 | aPD-L1 vs PBS | Up-regulated genes | CD45+ |
| BP | GO:0052547 | regulation of peptidase activity | 5.05E-03 | 3.27E-02 | Tnf/Nr4a1/Plaur/ | 10 | aPD-L1 vs PBS | Up-regulated genes | CD45+ |
| BP | GO:0007259 | JAK-STAT cascade | 5.11E-03 | 3.29E-02 | Irf1/Tnf/Ccl5/Stat1 | 6 | aPD-L1 vs PBS | Up-regulated genes | CD45+ |
| BP | GO:0051092 | positive regulation of NF-kappaB transcription fa | 5.11E-03 | 3.29E-02 | Tnf/Cd36/Malt1/Ic | 6 | aPD-L1 vs PBS | Up-regulated genes | CD45+ |
| BP | GO:0042129 | regulation of T cell proliferation | 5.13E-03 | 3.30E-02 | H2-Aa/Irf1/H2-Ab | 7 | aPD-L1 vs PBS | Up-regulated genes | CD45+ |
| BP | GO:0032655 | regulation of interleukin-12 production | 5.29E-03 | 3.37E-02 | Irf1/Cd36/Rel/Nfk | 4 | aPD-L1 vs PBS | Up-regulated genes | CD45+ |
| BP | GO:0032890 | regulation of organic acid transport | 5.29E-03 | 3.37E-02 | Tnf/Arg2/Il1a/Il1rn | 4 | aPD-L1 vs PBS | Up-regulated genes | CD45+ |
| BP | GO:0043620 | regulation of DNA-templated transcription in res | 5.29E-03 | 3.37E-02 | Atf3/Cd36/Egr1/Kl | 4 | aPD-L1 vs PBS | Up-regulated genes | CD45+ |
| BP | GO:0050810 | regulation of steroid biosynthetic process | 5.29E-03 | 3.37E-02 | Tnf/Nfkb1/Egr1/Il1 | 4 | aPD-L1 vs PBS | Up-regulated genes | CD45+ |
| BP | GO:0072594 | establishment of protein localization to organelle | 5.53E-03 | 3.52E-02 | Lmna/Dnaja1/Hsp | 11 | aPD-L1 vs PBS | Up-regulated genes | CD45+ |
| BP | GO:0034504 | protein localization to nucleus | 5.54E-03 | 3.52E-02 | Lmna/Mdm2/Gbp | 9 | aPD-L1 vs PBS | Up-regulated genes | CD45+ |
| BP | GO:0010470 | regulation of gastrulation | 5.61E-03 | 3.53E-02 | Tnfaip3/Klf4/Il1rn | 3 | aPD-L1 vs PBS | Up-regulated genes | CD45+ |
| BP | GO:0042036 | negative regulation of cytokine biosynthetic proc | 5.61E-03 | 3.53E-02 | Zfp36/Nfkb1/Klf4 | 3 | aPD-L1 vs PBS | Up-regulated genes | CD45+ |
| BP | GO:0042119 | neutrophil activation | 5.61E-03 | 3.53E-02 | Fcgr4/Ccl5/Itgb2 | 3 | aPD-L1 vs PBS | Up-regulated genes | CD45+ |
| BP | GO:0051955 | regulation of amino acid transport | 5.61E-03 | 3.53E-02 | Tnf/Arg2/Il1rn | 3 | aPD-L1 vs PBS | Up-regulated genes | CD45+ |
| BP | GO:0061082 | myeloid leukocyte cytokine production | 5.61E-03 | 3.53E-02 | Cd74/Cd36/Nr4a | 3 | aPD-L1 vs PBS | Up-regulated genes | CD45+ |
| BP | GO:0014911 | positive regulation of smooth muscle cell migrati | 5.66E-03 | 3.54E-02 | Mdm2/Ccl5/Nr4a | 4 | aPD-L1 vs PBS | Up-regulated genes | CD45+ |
| BP | GO:0033627 | cell adhesion mediated by integrin | 5.66E-03 | 3.54E-02 | Ccl5/Icam1/Itgb7/ | 4 | aPD-L1 vs PBS | Up-regulated genes | CD45+ |
| BP | GO:0043388 | positive regulation of DNA binding | 5.66E-03 | 3.54E-02 | Plaur/Txn1/Klf4/H | 4 | aPD-L1 vs PBS | Up-regulated genes | CD45+ |
| BP | GO:0097191 | extrinsic apoptotic signaling pathway | 5.69E-03 | 3.55E-02 | Lmna/Inhba/Tnf/A | 8 | aPD-L1 vs PBS | Up-regulated genes | CD45+ |
| BP | GO:0097696 | STAT cascade | 5.77E-03 | 3.60E-02 | Irf1/Tnf/Ccl5/Stat1 | 6 | aPD-L1 vs PBS | Up-regulated genes | CD45+ |
| BP | GO:0032615 | interleukin-12 production | 6.05E-03 | 3.76E-02 | Irf1/Cd36/Rel/Nfk | 4 | aPD-L1 vs PBS | Up-regulated genes | CD45+ |
| BP | GO:0070374 | positive regulation of ERK1 and ERK2 cascade | 6.08E-03 | 3.76E-02 | Cd74/Ccl4/Ccl5/C | 7 | aPD-L1 vs PBS | Up-regulated genes | CD45+ |
| BP | GO:2001242 | regulation of intrinsic apoptotic signaling pathwa | 6.08E-03 | 3.76E-02 | Cd74/Dnaja1/Pla | 7 | aPD-L1 vs PBS | Up-regulated genes | CD45+ |
| BP | GO:0050670 | regulation of lymphocyte proliferation | 6.22E-03 | 3.80E-02 | H2-Aa/Cd74/Irf1/ | 8 | aPD-L1 vs PBS | Up-regulated genes | CD45+ |
| BP | GO:0000027 | ribosomal large subunit assembly | 6.22E-03 | 3.80E-02 | Rpl38/Rplp0/Rpl1 | 3 | aPD-L1 vs PBS | Up-regulated genes | CD45+ |
| BP | GO:0031640 | killing of cells of other organism | 6.22E-03 | 3.80E-02 | Rpl30/Rps19/Nos | 3 | aPD-L1 vs PBS | Up-regulated genes | CD45+ |
| BP | GO:0044364 | disruption of cells of other organism | 6.22E-03 | 3.80E-02 | Rpl30/Rps19/Nos | 3 | aPD-L1 vs PBS | Up-regulated genes | CD45+ |
| BP | GO:0045589 | regulation of regulatory T cell differentiation | 6.22E-03 | 3.80E-02 | Irf1/Dusp10/Socs | 3 | aPD-L1 vs PBS | Up-regulated genes | CD45+ |
| BP | GO:0046885 | regulation of hormone biosynthetic process | 6.22E-03 | 3.80E-02 | Tnf/Nfkb1/Egr1 | 3 | aPD-L1 vs PBS | Up-regulated genes | CD45+ |
| BP | GO:0007254 | JNK cascade | 6.50E-03 | 3.96E-02 | Dnaja1/Tnf/Dusp | 7 | aPD-L1 vs PBS | Up-regulated genes | CD45+ |
| BP | GO:0002253 | activation of immune response | 6.72E-03 | 4.09E-02 | Irf1/Tnf/Rsad2/Ac | 11 | aPD-L1 vs PBS | Up-regulated genes | CD45+ |
| BP | GO:0009150 | purine ribonucleotide metabolic process | 6.72E-03 | 4.09E-02 | ATP8/Atp5e/Hspa | 11 | aPD-L1 vs PBS | Up-regulated genes | CD45+ |
| BP | GO:0019233 | sensory perception of pain | 6.79E-03 | 4.12E-02 | Tnf/Fcgr4/Il1a/Il1 | 5 | aPD-L1 vs PBS | Up-regulated genes | CD45+ |
| BP | GO:0002931 | response to ischemia | 6.87E-03 | 4.13E-02 | Hk2/Egr1/Csf1 | 3 | aPD-L1 vs PBS | Up-regulated genes | CD45+ |
| BP | GO:0060142 | regulation of syncytium formation by plasma me | 6.87E-03 | 4.13E-02 | Cxcl9/Stat1/Il1f9 | 3 | aPD-L1 vs PBS | Up-regulated genes | CD45+ |
| BP | GO:0071385 | cellular response to glucocorticoid stimulus | 6.87E-03 | 4.13E-02 | ND3/Zfp36/Zfp36l | 3 | aPD-L1 vs PBS | Up-regulated genes | CD45+ |
| BP | GO:0090322 | regulation of superoxide metabolic process | 6.87E-03 | 4.13E-02 | Tnf/Cd36/Itgb2 | 3 | aPD-L1 vs PBS | Up-regulated genes | CD45+ |
| BP | GO:1990748 | cellular detoxification | 6.87E-03 | 4.13E-02 | Tnf/Cd36/Sod2 | 3 | aPD-L1 vs PBS | Up-regulated genes | CD45+ |
| BP | GO:0016072 | rRNA metabolic process | 6.97E-03 | 4.18E-02 | Rps21/Rpl35a/Rp | 8 | aPD-L1 vs PBS | Up-regulated genes | CD45+ |
| BP | GO:1903900 | regulation of viral life cycle | 7.02E-03 | 4.20E-02 | Cd74/Tnf/Rsad2/ | 6 | aPD-L1 vs PBS | Up-regulated genes | CD45+ |
| BP | GO:0032612 | interleukin-1 production | 7.12E-03 | 4.25E-02 | Arg2/Tnfaip3/Cd3 | 5 | aPD-L1 vs PBS | Up-regulated genes | CD45+ |
| BP | GO:0045672 | positive regulation of osteoclast differentiation | 7.56E-03 | 4.41E-02 | Tnf/Ccl5/Csf1 | 3 | aPD-L1 vs PBS | Up-regulated genes | CD45+ |
| BP | GO:0071384 | cellular response to corticosteroid stimulus | 7.56E-03 | 4.41E-02 | ND3/Zfp36/Zfp36l | 3 | aPD-L1 vs PBS | Up-regulated genes | CD45+ |
| BP | GO:0002792 | negative regulation of peptide secretion | 7.57E-03 | 4.41E-02 | Cd74/Tnf/Arg2/Rs | 6 | aPD-L1 vs PBS | Up-regulated genes | CD45+ |
| BP | GO:0051403 | stress-activated MAPK cascade | 7.58E-03 | 4.41E-02 | Dnaja1/Tnf/Dusp | 8 | aPD-L1 vs PBS | Up-regulated genes | CD45+ |
| BP | GO:0002676 | regulation of chronic inflammatory response | 7.62E-03 | 4.41E-02 | Tnf/Tnfaip3 | 2 | aPD-L1 vs PBS | Up-regulated genes | CD45+ |
| BP | GO:0002679 | respiratory burst involved in defense response | 7.62E-03 | 4.41E-02 | Dusp10/Rps19 | 2 | aPD-L1 vs PBS | Up-regulated genes | CD45+ |
| BP | GO:0010744 | positive regulation of macrophage derived foam | 7.62E-03 | 4.41E-02 | Cd36/Csf1 | 2 | aPD-L1 vs PBS | Up-regulated genes | CD45+ |
| BP | GO:0015802 | basic amino acid transport | 7.62E-03 | 4.41E-02 | Arg2/Slc11a1 | 2 | aPD-L1 vs PBS | Up-regulated genes | CD45+ |

| | | | | | | | | | |
|---|---|---|---|---|---|---|---|---|---|
| BP | GO:0032769 | negative regulation of monooxygenase activity | 7.62E-03 | 4.41E-02 | Arg2/Nfkb1 | 2 | aPD-L1 vs PBS | Up-regulated genes | CD45+ |
| BP | GO:0042359 | vitamin D metabolic process | 7.62E-03 | 4.41E-02 | Tnf/Nfkb1 | 2 | aPD-L1 vs PBS | Up-regulated genes | CD45+ |
| BP | GO:0045080 | positive regulation of chemokine biosynthetic pro | 7.62E-03 | 4.41E-02 | Tnf/Egr1 | 2 | aPD-L1 vs PBS | Up-regulated genes | CD45+ |
| BP | GO:0045414 | regulation of interleukin-8 biosynthetic process | 7.62E-03 | 4.41E-02 | Tnf/Klf4 | 2 | aPD-L1 vs PBS | Up-regulated genes | CD45+ |
| BP | GO:0047484 | regulation of response to osmotic stress | 7.62E-03 | 4.41E-02 | Prkca/Ptgs2 | 2 | aPD-L1 vs PBS | Up-regulated genes | CD45+ |
| BP | GO:0048333 | mesodermal cell differentiation | 7.62E-03 | 4.41E-02 | Inhba/Kdm6b | 2 | aPD-L1 vs PBS | Up-regulated genes | CD45+ |
| BP | GO:0051956 | negative regulation of amino acid transport | 7.62E-03 | 4.41E-02 | Tnf/Il1rn | 2 | aPD-L1 vs PBS | Up-regulated genes | CD45+ |
| BP | GO:0097709 | connective tissue replacement | 7.62E-03 | 4.41E-02 | Il1a/Klf6 | 2 | aPD-L1 vs PBS | Up-regulated genes | CD45+ |
| BP | GO:2000551 | regulation of T-helper 2 cell cytokine production | 7.62E-03 | 4.41E-02 | Rsad2/Nlrp3 | 2 | aPD-L1 vs PBS | Up-regulated genes | CD45+ |
| BP | GO:0043410 | positive regulation of MAPK cascade | 7.64E-03 | 4.41E-02 | Cd74/Tnf/Ccl4/Cc | 12 | aPD-L1 vs PBS | Up-regulated genes | CD45+ |
| BP | GO:0019915 | lipid storage | 7.78E-03 | 4.48E-02 | Hilpda/Tnf/Cd36/I | 4 | aPD-L1 vs PBS | Up-regulated genes | CD45+ |
| BP | GO:0045786 | negative regulation of cell cycle | 7.79E-03 | 4.48E-02 | Irf1/Inhba/Tnf/Nr4 | 12 | aPD-L1 vs PBS | Up-regulated genes | CD45+ |
| BP | GO:0006091 | generation of precursor metabolites and energy | 8.06E-03 | 4.63E-02 | Hk3/Nr4a3/ND1/H | 10 | aPD-L1 vs PBS | Up-regulated genes | CD45+ |
| BP | GO:0002687 | positive regulation of leukocyte migration | 8.15E-03 | 4.66E-02 | Cd74/Ccl4/Ccl5/Ic | 6 | aPD-L1 vs PBS | Up-regulated genes | CD45+ |
| BP | GO:1902106 | negative regulation of leukocyte differentiation | 8.17E-03 | 4.66E-02 | Cd74/Irf1/Socs1/F | 5 | aPD-L1 vs PBS | Up-regulated genes | CD45+ |
| BP | GO:0042093 | T-helper cell differentiation | 8.26E-03 | 4.66E-02 | Irf1/Malt1/Nlrp3/S | 4 | aPD-L1 vs PBS | Up-regulated genes | CD45+ |
| BP | GO:0042773 | ATP synthesis coupled electron transport | 8.26E-03 | 4.66E-02 | Cox7c/ND4L/CYT | 4 | aPD-L1 vs PBS | Up-regulated genes | CD45+ |
| BP | GO:0009259 | ribonucleotide metabolic process | 8.27E-03 | 4.66E-02 | ATP8/Atp5e/Hspa | 11 | aPD-L1 vs PBS | Up-regulated genes | CD45+ |
| BP | GO:0000462 | maturation of SSU-rRNA from tricistronic rRNA tr | 8.28E-03 | 4.66E-02 | Rps21/Rps19/Rp | 3 | aPD-L1 vs PBS | Up-regulated genes | CD45+ |
| BP | GO:0032633 | interleukin-4 production | 8.28E-03 | 4.66E-02 | H2-T23/Cd83/Nlr | 3 | aPD-L1 vs PBS | Up-regulated genes | CD45+ |
| BP | GO:0045066 | regulatory T cell differentiation | 8.28E-03 | 4.66E-02 | Irf1/Dusp10/Socs | 3 | aPD-L1 vs PBS | Up-regulated genes | CD45+ |
| BP | GO:0051085 | chaperone cofactor-dependent protein refolding | 8.28E-03 | 4.66E-02 | Cd74/H2-DMb1/H | 3 | aPD-L1 vs PBS | Up-regulated genes | CD45+ |
| BP | GO:0060337 | type I interferon signaling pathway | 8.28E-03 | 4.66E-02 | Zbp1/Samhd1/Sta | 3 | aPD-L1 vs PBS | Up-regulated genes | CD45+ |
| BP | GO:0071357 | cellular response to type I interferon | 8.28E-03 | 4.66E-02 | Zbp1/Samhd1/Sta | 3 | aPD-L1 vs PBS | Up-regulated genes | CD45+ |
| BP | GO:0006606 | protein import into nucleus | 8.45E-03 | 4.74E-02 | Lmna/Hspa8/Cd3 | 6 | aPD-L1 vs PBS | Up-regulated genes | CD45+ |
| BP | GO:0002294 | CD4-positive, alpha-beta T cell differentiation inv | 8.76E-03 | 4.89E-02 | Irf1/Malt1/Nlrp3/S | 4 | aPD-L1 vs PBS | Up-regulated genes | CD45+ |
| BP | GO:0050710 | negative regulation of cytokine secretion | 8.76E-03 | 4.89E-02 | Tnf/Arg2/Tnfaip3/I | 4 | aPD-L1 vs PBS | Up-regulated genes | CD45+ |
| BP | GO:0070228 | regulation of lymphocyte apoptotic process | 8.76E-03 | 4.89E-02 | Cd74/Cd274/Arg2 | 4 | aPD-L1 vs PBS | Up-regulated genes | CD45+ |
| BP | GO:0009152 | purine ribonucleotide biosynthetic process | 8.90E-03 | 4.96E-02 | ATP8/Atp5e/Hk3/I | 7 | aPD-L1 vs PBS | Up-regulated genes | CD45+ |
| CC | GO:0022626 | cytosolic ribosome | 3.27E-47 | 9.88E-45 | Rpl39/Rps27/Rps | 39 | aPD-L1 vs PBS | Up-regulated genes | CD45+ |
| CC | GO:0044391 | ribosomal subunit | 3.16E-36 | 4.78E-34 | Rpl39/Rps27/Rps | 39 | aPD-L1 vs PBS | Up-regulated genes | CD45+ |
| CC | GO:0044445 | cytosolic part | 6.36E-36 | 6.41E-34 | Rpl39/Rps27/Rps | 42 | aPD-L1 vs PBS | Up-regulated genes | CD45+ |
| CC | GO:0005840 | ribosome | 9.17E-35 | 6.92E-33 | Rpl39/Rps27/Rps | 40 | aPD-L1 vs PBS | Up-regulated genes | CD45+ |
| CC | GO:0022625 | cytosolic large ribosomal subunit | 4.56E-34 | 2.75E-32 | Rpl39/Rpl37/Rpl3 | 26 | aPD-L1 vs PBS | Up-regulated genes | CD45+ |
| CC | GO:0015934 | large ribosomal subunit | 9.85E-25 | 4.96E-23 | Rpl39/Rpl37/Rpl3 | 26 | aPD-L1 vs PBS | Up-regulated genes | CD45+ |
| CC | GO:0022627 | cytosolic small ribosomal subunit | 3.11E-18 | 1.34E-16 | Rps27/Rps21/Rp | 15 | aPD-L1 vs PBS | Up-regulated genes | CD45+ |
| CC | GO:0015935 | small ribosomal subunit | 2.58E-14 | 9.72E-13 | Rps27/Rps21/Rp | 15 | aPD-L1 vs PBS | Up-regulated genes | CD45+ |
| CC | GO:0042611 | MHC protein complex | 4.41E-14 | 1.48E-12 | H2-Eb1/H2-Aa/Cd | 10 | aPD-L1 vs PBS | Up-regulated genes | CD45+ |
| CC | GO:0042788 | polysomal ribosome | 1.51E-12 | 4.57E-11 | Rpl39/Rps21/Rps | 10 | aPD-L1 vs PBS | Up-regulated genes | CD45+ |
| CC | GO:0005844 | polysome | 2.11E-08 | 5.80E-07 | Rpl39/Rps21/Rps | 10 | aPD-L1 vs PBS | Up-regulated genes | CD45+ |
| CC | GO:0042612 | MHC class I protein complex | 1.75E-07 | 4.40E-06 | H2-T23/B2m/H2-I | 5 | aPD-L1 vs PBS | Up-regulated genes | CD45+ |
| CC | GO:0009897 | external side of plasma membrane | 4.65E-06 | 1.08E-04 | H2-Aa/Cd74/Cxcl | 16 | aPD-L1 vs PBS | Up-regulated genes | CD45+ |
| CC | GO:0042824 | MHC class I peptide loading complex | 2.78E-05 | 6.01E-04 | B2m/H2-D1/H2-K | 4 | aPD-L1 vs PBS | Up-regulated genes | CD45+ |
| CC | GO:0098800 | inner mitochondrial membrane protein complex | 4.51E-05 | 9.09E-04 | ATP8/Atp5e/ND3/ | 9 | aPD-L1 vs PBS | Up-regulated genes | CD45+ |
| CC | GO:0098797 | plasma membrane protein complex | 5.95E-05 | 1.12E-03 | H2-Eb1/H2-Aa/Cd | 15 | aPD-L1 vs PBS | Up-regulated genes | CD45+ |
| CC | GO:0005753 | mitochondrial proton-transporting ATP synthase | 8.16E-05 | 1.45E-03 | ATP8/Atp5e/ATP6 | 4 | aPD-L1 vs PBS | Up-regulated genes | CD45+ |
| CC | GO:0045259 | proton-transporting ATP synthase complex | 1.02E-04 | 1.72E-03 | ATP8/Atp5e/ATP6 | 4 | aPD-L1 vs PBS | Up-regulated genes | CD45+ |
| CC | GO:0044455 | mitochondrial membrane part | 4.52E-04 | 6.80E-03 | ATP8/Atp5e/ND3/ | 10 | aPD-L1 vs PBS | Up-regulated genes | CD45+ |
| CC | GO:0098552 | side of membrane | 4.53E-04 | 6.80E-03 | H2-Aa/Cd74/Cxcl | 16 | aPD-L1 vs PBS | Up-regulated genes | CD45+ |
| CC | GO:0045263 | proton-transporting ATP synthase complex, coup | 4.73E-04 | 6.80E-03 | ATP8/ATP6/Atp5k | 3 | aPD-L1 vs PBS | Up-regulated genes | CD45+ |
| CC | GO:0098803 | respiratory chain complex | 5.29E-04 | 7.26E-03 | ND3/ND1/ND4L/C | 6 | aPD-L1 vs PBS | Up-regulated genes | CD45+ |
| CC | GO:0019866 | organelle inner membrane | 6.48E-04 | 8.51E-03 | Rsad2/ATP8/Atp5 | 15 | aPD-L1 vs PBS | Up-regulated genes | CD45+ |
| CC | GO:0070469 | respiratory chain | 8.45E-04 | 1.06E-02 | ND3/ND1/ND4L/C | 6 | aPD-L1 vs PBS | Up-regulated genes | CD45+ |
| CC | GO:0005743 | mitochondrial inner membrane | 8.81E-04 | 1.06E-02 | Rsad2/ATP8/Atp5 | 14 | aPD-L1 vs PBS | Up-regulated genes | CD45+ |
| CC | GO:0016469 | proton-transporting two-sector ATPase complex | 2.13E-03 | 2.47E-02 | ATP8/Atp5e/ATP6 | 4 | aPD-L1 vs PBS | Up-regulated genes | CD45+ |
| CC | GO:0033177 | proton-transporting two-sector ATPase complex, | 2.26E-03 | 2.53E-02 | ATP8/ATP6/Atp5k | 3 | aPD-L1 vs PBS | Up-regulated genes | CD45+ |
| CC | GO:0098798 | mitochondrial protein complex | 2.41E-03 | 2.60E-02 | ATP8/Atp5e/ND3/ | 10 | aPD-L1 vs PBS | Up-regulated genes | CD45+ |
| CC | GO:0005797 | Golgi medial cisterna | 3.00E-03 | 3.12E-02 | H2-D1/H2-K1/H2- | 3 | aPD-L1 vs PBS | Up-regulated genes | CD45+ |
| CC | GO:0005667 | transcription factor complex | 3.26E-03 | 3.19E-02 | Nr4a1/Hsp90ab1/ | 11 | aPD-L1 vs PBS | Up-regulated genes | CD45+ |
| CC | GO:0005746 | mitochondrial respiratory chain | 3.27E-03 | 3.19E-02 | ND3/ND1/ND4L/C | 5 | aPD-L1 vs PBS | Up-regulated genes | CD45+ |
| CC | GO:0005747 | mitochondrial respiratory chain complex I | 3.80E-03 | 3.37E-02 | ND3/ND1/ND4L/N | 4 | aPD-L1 vs PBS | Up-regulated genes | CD45+ |
| CC | GO:0030964 | NADH dehydrogenase complex | 3.80E-03 | 3.37E-02 | ND3/ND1/ND4L/N | 4 | aPD-L1 vs PBS | Up-regulated genes | CD45+ |
| CC | GO:0045271 | respiratory chain complex I | 3.80E-03 | 3.37E-02 | ND3/ND1/ND4L/N | 4 | aPD-L1 vs PBS | Up-regulated genes | CD45+ |
| CC | GO:0043230 | extracellular organelle | 4.80E-03 | 4.09E-02 | Cd274/Gbp2/Gbp | 5 | aPD-L1 vs PBS | Up-regulated genes | CD45+ |
| CC | GO:0070971 | endoplasmic reticulum exit site | 4.87E-03 | 4.09E-02 | H2-D1/H2-K1/H2- | 3 | aPD-L1 vs PBS | Up-regulated genes | CD45+ |
| MF | GO:0003735 | structural constituent of ribosome | 4.32E-36 | 1.67E-33 | Rpl39/Rps27/Rps | 36 | aPD-L1 vs PBS | Up-regulated genes | CD45+ |
| MF | GO:0005198 | structural molecule activity | 3.19E-20 | 6.16E-18 | Rpl39/Rps27/Rps | 37 | aPD-L1 vs PBS | Up-regulated genes | CD45+ |
| MF | GO:0042277 | peptide binding | 1.14E-07 | 1.47E-05 | H2-Aa/Cd74/H2-A | 15 | aPD-L1 vs PBS | Up-regulated genes | CD45+ |
| MF | GO:0042605 | peptide antigen binding | 4.12E-07 | 3.32E-05 | H2-Aa/H2-Ab1/H2 | 6 | aPD-L1 vs PBS | Up-regulated genes | CD45+ |
| MF | GO:0033218 | amide binding | 4.30E-07 | 3.32E-05 | H2-Aa/Cd74/H2-A | 16 | aPD-L1 vs PBS | Up-regulated genes | CD45+ |
| MF | GO:0005125 | cytokine activity | 8.04E-06 | 5.01E-04 | Cxcl9/Inhba/Tnf/C | 10 | aPD-L1 vs PBS | Up-regulated genes | CD45+ |
| MF | GO:0030881 | beta-2-microglobulin binding | 9.08E-06 | 5.01E-04 | H2-T23/H2-D1/H2 | 4 | aPD-L1 vs PBS | Up-regulated genes | CD45+ |
| MF | GO:0003823 | antigen binding | 1.12E-05 | 5.39E-04 | H2-Aa/H2-Ab1/H2 | 6 | aPD-L1 vs PBS | Up-regulated genes | CD45+ |
| MF | GO:0042608 | T cell receptor binding | 2.67E-05 | 1.15E-03 | H2-T23/H2-D1/H2 | 4 | aPD-L1 vs PBS | Up-regulated genes | CD45+ |
| MF | GO:0019843 | rRNA binding | 3.40E-05 | 1.31E-03 | Rpl37/Rpl37a/Md | 7 | aPD-L1 vs PBS | Up-regulated genes | CD45+ |
| MF | GO:0015078 | proton transmembrane transporter activity | 3.94E-05 | 1.38E-03 | ATP8/Atp5e/Cox7 | 8 | aPD-L1 vs PBS | Up-regulated genes | CD45+ |
| MF | GO:0001228 | DNA-binding transcription activator activity, RNA | 8.03E-05 | 2.58E-03 | Irf1/Nr4a1/Sub1/N | 13 | aPD-L1 vs PBS | Up-regulated genes | CD45+ |
| MF | GO:0005126 | cytokine receptor binding | 1.47E-04 | 4.35E-03 | Cxcl9/Inhba/Tnf/C | 11 | aPD-L1 vs PBS | Up-regulated genes | CD45+ |
| MF | GO:0031072 | heat shock protein binding | 2.67E-04 | 7.35E-03 | Dnaja1/Hsp90ab1 | 8 | aPD-L1 vs PBS | Up-regulated genes | CD45+ |
| MF | GO:0030545 | receptor regulator activity | 3.07E-04 | 7.45E-03 | Cxcl9/Inhba/Tnf/C | 12 | aPD-L1 vs PBS | Up-regulated genes | CD45+ |
| MF | GO:0031386 | protein tag | 3.47E-04 | 7.45E-03 | Uba52/Ubc/Uba5 | 3 | aPD-L1 vs PBS | Up-regulated genes | CD45+ |
| MF | GO:0042610 | CD8 receptor binding | 3.47E-04 | 7.45E-03 | H2-D1/H2-K1/H2- | 3 | aPD-L1 vs PBS | Up-regulated genes | CD45+ |
| MF | GO:0046977 | TAP binding | 3.47E-04 | 7.45E-03 | H2-D1/H2-K1/H2- | 3 | aPD-L1 vs PBS | Up-regulated genes | CD45+ |
| MF | GO:0048018 | receptor ligand activity | 5.84E-04 | 1.18E-02 | Cxcl9/Inhba/Tnf/C | 11 | aPD-L1 vs PBS | Up-regulated genes | CD45+ |
| MF | GO:0044389 | ubiquitin-like protein ligase binding | 6.11E-04 | 1.18E-02 | H2-Ab1/Dnaja1/U | 12 | aPD-L1 vs PBS | Up-regulated genes | CD45+ |
| MF | GO:0005149 | interleukin-1 receptor binding | 7.44E-04 | 1.37E-02 | Il1f9/Il1a/Il1rn | 3 | aPD-L1 vs PBS | Up-regulated genes | CD45+ |
| MF | GO:0000981 | DNA-binding transcription factor activity, RNA po | 8.47E-04 | 1.49E-02 | Irf1/Nr4a1/Sub1/A | 16 | aPD-L1 vs PBS | Up-regulated genes | CD45+ |
| MF | GO:0030547 | receptor inhibitor activity | 9.21E-04 | 1.55E-02 | Ccl5/Ly6e/Il1rn | 3 | aPD-L1 vs PBS | Up-regulated genes | CD45+ |
| MF | GO:0031625 | ubiquitin protein ligase binding | 1.25E-03 | 2.02E-02 | H2-Ab1/Dnaja1/U | 11 | aPD-L1 vs PBS | Up-regulated genes | CD45+ |

| | | | | | | | | | |
|---|---|---|---|---|---|---|---|---|---|
| MF | GO:0035014 | phosphatidylinositol 3-kinase regulator activity | 1.60E-03 | 2.48E-02 | Socs3/Socs1/Klf4 | 3 | aPD-L1 vs PBS | Up-regulated genes | CD45+ |
| MF | GO:0035259 | glucocorticoid receptor binding | 1.89E-03 | 2.62E-02 | Nr4a1/Nr4a3/Ets2 | 3 | aPD-L1 vs PBS | Up-regulated genes | CD45+ |
| MF | GO:0050998 | nitric-oxide synthase binding | 1.89E-03 | 2.62E-02 | Cd74/Arg2/Nos2 | 3 | aPD-L1 vs PBS | Up-regulated genes | CD45+ |
| MF | GO:0019207 | kinase regulator activity | 1.90E-03 | 2.62E-02 | Hsp90ab1/Ccl5/M | 8 | aPD-L1 vs PBS | Up-regulated genes | CD45+ |
| MF | GO:0000978 | RNA polymerase II proximal promoter sequence | 2.05E-03 | 2.73E-02 | Irf1/Nr4a1/Atf3/Nr | 13 | aPD-L1 vs PBS | Up-regulated genes | CD45+ |
| MF | GO:0004129 | cytochrome-c oxidase activity | 2.54E-03 | 2.99E-02 | Cox7c/Cox6c/CO | 3 | aPD-L1 vs PBS | Up-regulated genes | CD45+ |
| MF | GO:0015002 | heme-copper terminal oxidase activity | 2.54E-03 | 2.99E-02 | Cox7c/Cox6c/CO | 3 | aPD-L1 vs PBS | Up-regulated genes | CD45+ |
| MF | GO:0016676 | oxidoreductase activity, acting on a heme group | 2.54E-03 | 2.99E-02 | Cox7c/Cox6c/CO | 3 | aPD-L1 vs PBS | Up-regulated genes | CD45+ |
| MF | GO:0000987 | proximal promoter sequence-specific DNA bindi | 2.56E-03 | 2.99E-02 | Irf1/Nr4a1/Atf3/Nr | 13 | aPD-L1 vs PBS | Up-regulated genes | CD45+ |
| MF | GO:0033293 | monocarboxylic acid binding | 2.89E-03 | 3.12E-02 | Hspa8/Cd36/S10 | 4 | aPD-L1 vs PBS | Up-regulated genes | CD45+ |
| MF | GO:0042379 | chemokine receptor binding | 2.89E-03 | 3.12E-02 | Cxcl9/Ccl4/Ccl5/S | 4 | aPD-L1 vs PBS | Up-regulated genes | CD45+ |
| MF | GO:0016675 | oxidoreductase activity, acting on a heme group | 2.91E-03 | 3.12E-02 | Cox7c/Cox6c/CO | 3 | aPD-L1 vs PBS | Up-regulated genes | CD45+ |
| BP | GO:0009617 | response to bacterium | 1.66E-09 | 2.23E-06 | Cxcl10/B2m/Gbp2 | 13 | aPD-L1 vs PBS | Up-regulated genes | CD45- |
| BP | GO:0042832 | defense response to protozoan | 3.25E-08 | 2.18E-05 | Gbp2b/Iigp1/Gbp2 | 5 | aPD-L1 vs PBS | Up-regulated genes | CD45- |
| BP | GO:0001562 | response to protozoan | 5.69E-08 | 2.55E-05 | Gbp2b/Iigp1/Gbp2 | 5 | aPD-L1 vs PBS | Up-regulated genes | CD45- |
| BP | GO:0044406 | adhesion of symbiont to host | 1.76E-07 | 5.92E-05 | Gbp2b/Gbp2/Gbp | 4 | aPD-L1 vs PBS | Up-regulated genes | CD45- |
| BP | GO:0035458 | cellular response to interferon-beta | 4.18E-07 | 1.12E-04 | Irf1/Gbp2b/Iigp1/G | 5 | aPD-L1 vs PBS | Up-regulated genes | CD45- |
| BP | GO:0048660 | regulation of smooth muscle cell proliferation | 7.21E-07 | 1.61E-04 | Igfbp5/Egr1/Trib1/ | 7 | aPD-L1 vs PBS | Up-regulated genes | CD45- |
| BP | GO:0048659 | smooth muscle cell proliferation | 9.03E-07 | 1.61E-04 | Igfbp5/Egr1/Trib1/ | 7 | aPD-L1 vs PBS | Up-regulated genes | CD45- |
| BP | GO:0071346 | cellular response to interferon-gamma | 1.04E-06 | 1.61E-04 | Irf1/Gbp2b/Gbp2/ | 6 | aPD-L1 vs PBS | Up-regulated genes | CD45- |
| BP | GO:0035456 | response to interferon-beta | 1.08E-06 | 1.61E-04 | Irf1/Gbp2b/Iigp1/G | 5 | aPD-L1 vs PBS | Up-regulated genes | CD45- |
| BP | GO:0098542 | defense response to other organism | 1.57E-06 | 2.11E-04 | Cxcl10/Irf1/Gbp2b | 9 | aPD-L1 vs PBS | Up-regulated genes | CD45- |
| BP | GO:0034341 | response to interferon-gamma | 2.82E-06 | 3.45E-04 | Irf1/Gbp2b/Gbp2/ | 6 | aPD-L1 vs PBS | Up-regulated genes | CD45- |
| BP | GO:1904707 | positive regulation of vascular smooth muscle ce | 5.70E-06 | 6.39E-04 | Igfbp5/Mmp2/Jun/ | 4 | aPD-L1 vs PBS | Up-regulated genes | CD45- |
| BP | GO:0033002 | muscle cell proliferation | 7.05E-06 | 7.30E-04 | Igfbp5/Egr1/Trib1/ | 7 | aPD-L1 vs PBS | Up-regulated genes | CD45- |
| BP | GO:0042742 | defense response to bacterium | 7.78E-06 | 7.47E-04 | Gbp2b/Iigp1/Gbp2 | 6 | aPD-L1 vs PBS | Up-regulated genes | CD45- |
| BP | GO:0016525 | negative regulation of angiogenesis | 1.67E-05 | 1.50E-03 | Klf2/Cxcl10/Dcn/A | 5 | aPD-L1 vs PBS | Up-regulated genes | CD45- |
| BP | GO:0043618 | regulation of transcription from RNA polymerase | 1.94E-05 | 1.62E-03 | Klf2/Atf3/Egr1/Jun | 4 | aPD-L1 vs PBS | Up-regulated genes | CD45- |
| BP | GO:2000181 | negative regulation of blood vessel morphogene | 2.05E-05 | 1.62E-03 | Klf2/Cxcl10/Dcn/A | 5 | aPD-L1 vs PBS | Up-regulated genes | CD45- |
| BP | GO:0071498 | cellular response to fluid shear stress | 2.19E-05 | 1.64E-03 | Klf2/Spp1/Klf4 | 3 | aPD-L1 vs PBS | Up-regulated genes | CD45- |
| BP | GO:0048661 | positive regulation of smooth muscle cell prolifer | 2.38E-05 | 1.66E-03 | Igfbp5/Egr1/Mmp2 | 5 | aPD-L1 vs PBS | Up-regulated genes | CD45- |
| BP | GO:0002428 | antigen processing and presentation of peptide | 2.66E-05 | 1.66E-03 | B2m/H2-D1/H2-K | 3 | aPD-L1 vs PBS | Up-regulated genes | CD45- |
| BP | GO:1904705 | regulation of vascular smooth muscle cell prolife | 2.72E-05 | 1.66E-03 | Igfbp5/Mmp2/Jun/ | 4 | aPD-L1 vs PBS | Up-regulated genes | CD45- |
| BP | GO:1990874 | vascular smooth muscle cell proliferation | 2.72E-05 | 1.66E-03 | Igfbp5/Mmp2/Jun/ | 4 | aPD-L1 vs PBS | Up-regulated genes | CD45- |
| BP | GO:0043620 | regulation of DNA-templated transcription in res | 2.94E-05 | 1.72E-03 | Klf2/Atf3/Egr1/Jun | 4 | aPD-L1 vs PBS | Up-regulated genes | CD45- |
| BP | GO:1901343 | negative regulation of vasculature development | 3.47E-05 | 1.94E-03 | Klf2/Cxcl10/Dcn/A | 5 | aPD-L1 vs PBS | Up-regulated genes | CD45- |
| BP | GO:0002475 | antigen processing and presentation via MHC cl | 4.42E-05 | 2.29E-03 | B2m/H2-D1/H2-K | 3 | aPD-L1 vs PBS | Up-regulated genes | CD45- |
| BP | GO:0071276 | cellular response to cadmium ion | 4.42E-05 | 2.29E-03 | Fos/Mt2/Jun | 3 | aPD-L1 vs PBS | Up-regulated genes | CD45- |
| BP | GO:0071248 | cellular response to metal ion | 5.32E-05 | 2.58E-03 | B2m/Fosb/Fos/Mt | 5 | aPD-L1 vs PBS | Up-regulated genes | CD45- |
| BP | GO:0050830 | defense response to Gram-positive bacterium | 5.60E-05 | 2.58E-03 | Gbp2b/Gbp2/Gbp | 4 | aPD-L1 vs PBS | Up-regulated genes | CD45- |
| BP | GO:0002483 | antigen processing and presentation of endoger | 5.94E-05 | 2.58E-03 | B2m/H2-D1/H2-K | 3 | aPD-L1 vs PBS | Up-regulated genes | CD45- |
| BP | GO:0019885 | antigen processing and presentation of endoger | 5.94E-05 | 2.58E-03 | B2m/H2-D1/H2-K | 3 | aPD-L1 vs PBS | Up-regulated genes | CD45- |
| BP | GO:0034405 | response to fluid shear stress | 5.94E-05 | 2.58E-03 | Klf2/Spp1/Klf4 | 3 | aPD-L1 vs PBS | Up-regulated genes | CD45- |
| BP | GO:0019883 | antigen processing and presentation of endoger | 7.77E-05 | 3.26E-03 | B2m/H2-D1/H2-K | 3 | aPD-L1 vs PBS | Up-regulated genes | CD45- |
| BP | GO:0002237 | response to molecule of bacterial origin | 1.09E-04 | 4.33E-03 | Cxcl10/B2m/Gbp2 | 6 | aPD-L1 vs PBS | Up-regulated genes | CD45- |
| BP | GO:0001916 | positive regulation of T cell mediated cytotoxicity | 1.11E-04 | 4.33E-03 | B2m/H2-D1/H2-K | 3 | aPD-L1 vs PBS | Up-regulated genes | CD45- |
| BP | GO:0071241 | cellular response to inorganic substance | 1.13E-04 | 4.33E-03 | B2m/Fosb/Fos/Mt | 5 | aPD-L1 vs PBS | Up-regulated genes | CD45- |
| BP | GO:0046686 | response to cadmium ion | 1.39E-04 | 5.18E-03 | Fos/Mt2/Jun | 3 | aPD-L1 vs PBS | Up-regulated genes | CD45- |
| BP | GO:0002474 | antigen processing and presentation of peptide | 1.54E-04 | 5.43E-03 | B2m/H2-D1/H2-K | 3 | aPD-L1 vs PBS | Up-regulated genes | CD45- |
| BP | GO:0048730 | epidermis morphogenesis | 1.54E-04 | 5.43E-03 | Igfbp5/Fst/Klf4 | 3 | aPD-L1 vs PBS | Up-regulated genes | CD45- |
| BP | GO:0001914 | regulation of T cell mediated cytotoxicity | 1.70E-04 | 5.85E-03 | B2m/H2-D1/H2-K | 3 | aPD-L1 vs PBS | Up-regulated genes | CD45- |
| BP | GO:0014910 | regulation of smooth muscle cell migration | 1.90E-04 | 6.39E-03 | Igfbp5/Egr1/Trib1/ | 4 | aPD-L1 vs PBS | Up-regulated genes | CD45- |
| BP | GO:0002824 | positive regulation of adaptive immune response | 2.28E-04 | 7.47E-03 | Irf1/B2m/H2-D1/H | 4 | aPD-L1 vs PBS | Up-regulated genes | CD45- |
| BP | GO:0001933 | negative regulation of protein phosphorylation | 2.37E-04 | 7.47E-03 | Irf1/Atf3/Trib1/Gbp | 7 | aPD-L1 vs PBS | Up-regulated genes | CD45- |
| BP | GO:0008585 | female gonad development | 2.39E-04 | 7.47E-03 | Mmp2/Fst/Ubb/Ac | 4 | aPD-L1 vs PBS | Up-regulated genes | CD45- |
| BP | GO:0014909 | smooth muscle cell migration | 2.61E-04 | 7.79E-03 | Igfbp5/Egr1/Trib1/ | 4 | aPD-L1 vs PBS | Up-regulated genes | CD45- |
| BP | GO:0046545 | development of primary female sexual character | 2.61E-04 | 7.79E-03 | Mmp2/Fst/Ubb/Ac | 4 | aPD-L1 vs PBS | Up-regulated genes | CD45- |
| BP | GO:0045087 | innate immune response | 2.85E-04 | 8.33E-03 | Irf1/Gbp2b/Iigp1/G | 8 | aPD-L1 vs PBS | Up-regulated genes | CD45- |
| BP | GO:0002821 | positive regulation of adaptive immune response | 3.08E-04 | 8.83E-03 | Irf1/B2m/H2-D1/H | 4 | aPD-L1 vs PBS | Up-regulated genes | CD45- |
| BP | GO:1901342 | regulation of vasculature development | 3.89E-04 | 1.09E-02 | Klf2/Cxcl10/Egr1/ | 6 | aPD-L1 vs PBS | Up-regulated genes | CD45- |
| BP | GO:0031103 | axon regeneration | 4.22E-04 | 1.14E-02 | Mmp2/Jun/Klf4 | 3 | aPD-L1 vs PBS | Up-regulated genes | CD45- |
| BP | GO:0046660 | female sex differentiation | 4.23E-04 | 1.14E-02 | Mmp2/Fst/Ubb/Ac | 4 | aPD-L1 vs PBS | Up-regulated genes | CD45- |
| BP | GO:0014812 | muscle cell migration | 4.39E-04 | 1.16E-02 | Igfbp5/Egr1/Trib1/ | 4 | aPD-L1 vs PBS | Up-regulated genes | CD45- |
| BP | GO:0042326 | negative regulation of phosphorylation | 4.49E-04 | 1.16E-02 | Irf1/Atf3/Trib1/Gbp | 7 | aPD-L1 vs PBS | Up-regulated genes | CD45- |
| BP | GO:0001912 | positive regulation of leukocyte mediated cytoto | 5.17E-04 | 1.29E-02 | B2m/H2-D1/H2-K | 3 | aPD-L1 vs PBS | Up-regulated genes | CD45- |
| BP | GO:0002711 | positive regulation of T cell mediated immunity | 5.17E-04 | 1.29E-02 | B2m/H2-D1/H2-K | 3 | aPD-L1 vs PBS | Up-regulated genes | CD45- |
| BP | GO:0043568 | positive regulation of insulin-like growth factor re | 5.32E-04 | 1.30E-02 | Igfbp5/Igfbp4 | 2 | aPD-L1 vs PBS | Up-regulated genes | CD45- |
| BP | GO:0048002 | antigen processing and presentation of peptide | 5.52E-04 | 1.30E-02 | B2m/H2-D1/H2-K | 3 | aPD-L1 vs PBS | Up-regulated genes | CD45- |
| BP | GO:0048662 | negative regulation of smooth muscle cell prolife | 5.52E-04 | 1.30E-02 | Igfbp5/Trib1/Klf4 | 3 | aPD-L1 vs PBS | Up-regulated genes | CD45- |
| BP | GO:0001913 | T cell mediated cytotoxicity | 5.88E-04 | 1.36E-02 | B2m/H2-D1/H2-K | 3 | aPD-L1 vs PBS | Up-regulated genes | CD45- |
| BP | GO:0002763 | positive regulation of myeloid leukocyte different | 6.26E-04 | 1.40E-02 | Fos/Trib1/Jun | 3 | aPD-L1 vs PBS | Up-regulated genes | CD45- |
| BP | GO:0061614 | pri-miRNA transcription by RNA polymerase II | 6.26E-04 | 1.40E-02 | Fos/Jun/Klf4 | 3 | aPD-L1 vs PBS | Up-regulated genes | CD45- |
| BP | GO:0031102 | neuron projection regeneration | 7.06E-04 | 1.49E-02 | Mmp2/Jun/Klf4 | 3 | aPD-L1 vs PBS | Up-regulated genes | CD45- |
| BP | GO:0001525 | angiogenesis | 7.10E-04 | 1.49E-02 | Klf2/Cxcl10/Dcn/M | 7 | aPD-L1 vs PBS | Up-regulated genes | CD45- |
| BP | GO:0010038 | response to metal ion | 7.10E-04 | 1.49E-02 | B2m/Fosb/Fos/Mt | 5 | aPD-L1 vs PBS | Up-regulated genes | CD45- |
| BP | GO:0032496 | response to lipopolysaccharide | 7.10E-04 | 1.49E-02 | Cxcl10/Gbp2/Trib | 5 | aPD-L1 vs PBS | Up-regulated genes | CD45- |
| BP | GO:0014745 | negative regulation of muscle adaptation | 7.77E-04 | 1.61E-02 | Igfbp5/Klf4 | 2 | aPD-L1 vs PBS | Up-regulated genes | CD45- |
| BP | GO:0031343 | positive regulation of cell killing | 7.92E-04 | 1.61E-02 | B2m/H2-D1/H2-K | 3 | aPD-L1 vs PBS | Up-regulated genes | CD45- |
| BP | GO:0071277 | cellular response to calcium ion | 8.84E-04 | 1.78E-02 | Fosb/Fos/Jun | 3 | aPD-L1 vs PBS | Up-regulated genes | CD45- |
| BP | GO:1902107 | positive regulation of leukocyte differentiation | 9.43E-04 | 1.87E-02 | Irf1/Fos/Trib1/Jun | 4 | aPD-L1 vs PBS | Up-regulated genes | CD45- |
| BP | GO:0002822 | regulation of adaptive immune response based | 1.03E-03 | 1.99E-02 | Irf1/B2m/H2-D1/H | 4 | aPD-L1 vs PBS | Up-regulated genes | CD45- |
| BP | GO:0048678 | response to axon injury | 1.04E-03 | 1.99E-02 | Mmp2/Jun/Klf4 | 3 | aPD-L1 vs PBS | Up-regulated genes | CD45- |
| BP | GO:0001910 | regulation of leukocyte mediated cytotoxicity | 1.14E-03 | 2.17E-02 | B2m/H2-D1/H2-K | 3 | aPD-L1 vs PBS | Up-regulated genes | CD45- |
| BP | GO:0014911 | positive regulation of smooth muscle cell migrati | 1.20E-03 | 2.20E-02 | Igfbp5/Egr1/Adam | 3 | aPD-L1 vs PBS | Up-regulated genes | CD45- |
| BP | GO:0035914 | skeletal muscle cell differentiation | 1.20E-03 | 2.20E-02 | Atf3/Egr1/Fos | 3 | aPD-L1 vs PBS | Up-regulated genes | CD45- |
| BP | GO:0002486 | antigen processing and presentation of endoger | 1.23E-03 | 2.20E-02 | H2-D1/H2-K1 | 2 | aPD-L1 vs PBS | Up-regulated genes | CD45- |
| BP | GO:0035994 | response to muscle stretch | 1.23E-03 | 2.20E-02 | Fos/Jun | 2 | aPD-L1 vs PBS | Up-regulated genes | CD45- |
| BP | GO:0002709 | regulation of T cell mediated immunity | 1.26E-03 | 2.23E-02 | B2m/H2-D1/H2-K | 3 | aPD-L1 vs PBS | Up-regulated genes | CD45- |
| BP | GO:0010563 | negative regulation of phosphorus metabolic pro | 1.33E-03 | 2.29E-02 | Irf1/Atf3/Trib1/Gbp | 7 | aPD-L1 vs PBS | Up-regulated genes | CD45- |

| | | | | | | | | | |
|---|---|---|---|---|---|---|---|---|---|
| BP | GO:0045936 | negative regulation of phosphate metabolic proc | 1.33E-03 | 2.29E-02 | Irf1/Atf3/Trib1/Gbp | 7 | aPD-L1 vs PBS | Up-regulated genes | CD45- |
| BP | GO:0070373 | negative regulation of ERK1 and ERK2 cascade | 1.38E-03 | 2.32E-02 | Atf3/Dusp6/Klf4 | 3 | aPD-L1 vs PBS | Up-regulated genes | CD45- |
| BP | GO:0002476 | antigen processing and presentation of endoger | 1.40E-03 | 2.32E-02 | H2-D1/H2-K1 | 2 | aPD-L1 vs PBS | Up-regulated genes | CD45- |
| BP | GO:0002484 | antigen processing and presentation of endoger | 1.40E-03 | 2.32E-02 | H2-D1/H2-K1 | 2 | aPD-L1 vs PBS | Up-regulated genes | CD45- |
| BP | GO:0010035 | response to inorganic substance | 1.42E-03 | 2.33E-02 | B2m/Fosb/Fos/Mr | 6 | aPD-L1 vs PBS | Up-regulated genes | CD45- |
| BP | GO:0002819 | regulation of adaptive immune response | 1.47E-03 | 2.39E-02 | Irf1/B2m/H2-D1/H | 4 | aPD-L1 vs PBS | Up-regulated genes | CD45- |
| BP | GO:0071363 | cellular response to growth factor stimulus | 1.53E-03 | 2.45E-02 | Htra3/Egr1/Dcn/F | 7 | aPD-L1 vs PBS | Up-regulated genes | CD45- |
| BP | GO:0045765 | regulation of angiogenesis | 1.67E-03 | 2.64E-02 | Klf2/Cxcl10/Dcn/A | 5 | aPD-L1 vs PBS | Up-regulated genes | CD45- |
| BP | GO:0071396 | cellular response to lipid | 1.98E-03 | 3.03E-02 | Cxcl10/Gbp2/Sgk | 6 | aPD-L1 vs PBS | Up-regulated genes | CD45- |
| BP | GO:0043567 | regulation of insulin-like growth factor receptor s | 1.98E-03 | 3.03E-02 | Igfbp5/Igfbp4 | 2 | aPD-L1 vs PBS | Up-regulated genes | CD45- |
| BP | GO:1904754 | positive regulation of vascular associated smoot | 1.98E-03 | 3.03E-02 | Igfbp5/Adamts1 | 2 | aPD-L1 vs PBS | Up-regulated genes | CD45- |
| BP | GO:0031341 | regulation of cell killing | 2.03E-03 | 3.07E-02 | B2m/H2-D1/H2-K | 3 | aPD-L1 vs PBS | Up-regulated genes | CD45- |
| BP | GO:0019882 | antigen processing and presentation | 2.12E-03 | 3.16E-02 | B2m/H2-D1/H2-K | 3 | aPD-L1 vs PBS | Up-regulated genes | CD45- |
| BP | GO:0008406 | gonad development | 2.34E-03 | 3.42E-02 | Mmp2/Fst/Ubb/Ad | 4 | aPD-L1 vs PBS | Up-regulated genes | CD45- |
| BP | GO:1903708 | positive regulation of hemopoiesis | 2.34E-03 | 3.42E-02 | Irf1/Fos/Trib1/Jun | 4 | aPD-L1 vs PBS | Up-regulated genes | CD45- |
| BP | GO:0042035 | regulation of cytokine biosynthetic process | 2.37E-03 | 3.43E-02 | Irf1/Egr1/Klf4 | 3 | aPD-L1 vs PBS | Up-regulated genes | CD45- |
| BP | GO:0007178 | transmembrane receptor protein serine/threonin | 2.42E-03 | 3.43E-02 | Htra3/Egr1/Fos/F | 5 | aPD-L1 vs PBS | Up-regulated genes | CD45- |
| BP | GO:0014912 | negative regulation of smooth muscle cell migrat | 2.42E-03 | 3.43E-02 | Igfbp5/Trib1 | 2 | aPD-L1 vs PBS | Up-regulated genes | CD45- |
| BP | GO:0045137 | development of primary sexual characteristics | 2.50E-03 | 3.51E-02 | Mmp2/Fst/Ubb/Ad | 4 | aPD-L1 vs PBS | Up-regulated genes | CD45- |
| BP | GO:0045639 | positive regulation of myeloid cell differentiation | 2.56E-03 | 3.55E-02 | Fos/Trib1/Jun | 3 | aPD-L1 vs PBS | Up-regulated genes | CD45- |
| BP | GO:0002478 | antigen processing and presentation of exogenc | 2.66E-03 | 3.65E-02 | B2m/H2-K1 | 2 | aPD-L1 vs PBS | Up-regulated genes | CD45- |
| BP | GO:0001909 | leukocyte mediated cytotoxicity | 2.85E-03 | 3.87E-02 | B2m/H2-D1/H2-K | 3 | aPD-L1 vs PBS | Up-regulated genes | CD45- |
| BP | GO:0060343 | trabecula formation | 2.91E-03 | 3.91E-02 | Mmp2/Adamts1 | 2 | aPD-L1 vs PBS | Up-regulated genes | CD45- |
| BP | GO:0002708 | positive regulation of lymphocyte mediated immu | 3.05E-03 | 3.94E-02 | B2m/H2-D1/H2-K | 3 | aPD-L1 vs PBS | Up-regulated genes | CD45- |
| BP | GO:0045637 | regulation of myeloid cell differentiation | 3.10E-03 | 3.94E-02 | B2m/Fos/Trib1/Ju | 4 | aPD-L1 vs PBS | Up-regulated genes | CD45- |
| BP | GO:0036003 | positive regulation of transcription from RNA pol | 3.16E-03 | 3.94E-02 | Klf2/Atf3 | 2 | aPD-L1 vs PBS | Up-regulated genes | CD45- |
| BP | GO:0044342 | type B pancreatic cell proliferation | 3.16E-03 | 3.94E-02 | Igfbp5/Igfbp4 | 2 | aPD-L1 vs PBS | Up-regulated genes | CD45- |
| BP | GO:1903671 | negative regulation of sprouting angiogenesis | 3.16E-03 | 3.94E-02 | Klf2/Klf4 | 2 | aPD-L1 vs PBS | Up-regulated genes | CD45- |
| BP | GO:1904738 | vascular associated smooth muscle cell migratio | 3.16E-03 | 3.94E-02 | Igfbp5/Adamts1 | 2 | aPD-L1 vs PBS | Up-regulated genes | CD45- |
| BP | GO:1904752 | regulation of vascular associated smooth muscle | 3.16E-03 | 3.94E-02 | Igfbp5/Adamts1 | 2 | aPD-L1 vs PBS | Up-regulated genes | CD45- |
| BP | GO:0002699 | positive regulation of immune effector process | 3.17E-03 | 3.94E-02 | Irf1/B2m/H2-D1/H | 4 | aPD-L1 vs PBS | Up-regulated genes | CD45- |
| BP | GO:0002521 | leukocyte differentiation | 3.19E-03 | 3.94E-02 | Irf1/B2m/Egr1/Fos | 6 | aPD-L1 vs PBS | Up-regulated genes | CD45- |
| BP | GO:0051607 | defense response to virus | 3.37E-03 | 4.09E-02 | Cxcl10/Irf1/Gbp2b | 4 | aPD-L1 vs PBS | Up-regulated genes | CD45- |
| BP | GO:0042089 | cytokine biosynthetic process | 3.37E-03 | 4.09E-02 | Irf1/Egr1/Klf4 | 3 | aPD-L1 vs PBS | Up-regulated genes | CD45- |
| BP | GO:0042107 | cytokine metabolic process | 3.60E-03 | 4.32E-02 | Irf1/Egr1/Klf4 | 3 | aPD-L1 vs PBS | Up-regulated genes | CD45- |
| BP | GO:0002697 | regulation of immune effector process | 3.70E-03 | 4.37E-02 | Irf1/B2m/H2-D1/H | 5 | aPD-L1 vs PBS | Up-regulated genes | CD45- |
| BP | GO:0031069 | hair follicle morphogenesis | 3.71E-03 | 4.37E-02 | Igfbp5/Fst | 2 | aPD-L1 vs PBS | Up-regulated genes | CD45- |
| BP | GO:0002456 | T cell mediated immunity | 3.84E-03 | 4.49E-02 | B2m/H2-D1/H2-K | 3 | aPD-L1 vs PBS | Up-regulated genes | CD45- |
| BP | GO:0019884 | antigen processing and presentation of exogenc | 4.00E-03 | 4.63E-02 | B2m/H2-K1 | 2 | aPD-L1 vs PBS | Up-regulated genes | CD45- |
| BP | GO:0051592 | response to calcium ion | 4.21E-03 | 4.83E-02 | Fosb/Fos/Jun | 3 | aPD-L1 vs PBS | Up-regulated genes | CD45- |
| CC | GO:0020005 | symbiont-containing vacuole membrane | 1.60E-10 | 1.25E-08 | Gbp2b/Iigp1/Gbp | 5 | aPD-L1 vs PBS | Up-regulated genes | CD45- |
| CC | GO:0020003 | symbiont-containing vacuole | 4.43E-10 | 1.73E-08 | Gbp2b/Iigp1/Gbp | 5 | aPD-L1 vs PBS | Up-regulated genes | CD45- |
| CC | GO:0065010 | extracellular membrane-bounded organelle | 6.87E-10 | 1.79E-08 | Gbp2b/Iigp1/Gbp | 5 | aPD-L1 vs PBS | Up-regulated genes | CD45- |
| CC | GO:0030430 | host cell cytoplasm | 1.49E-09 | 2.33E-08 | Gbp2b/Iigp1/Gbp | 5 | aPD-L1 vs PBS | Up-regulated genes | CD45- |
| CC | GO:0033655 | host cell cytoplasm part | 1.49E-09 | 2.33E-08 | Gbp2b/Iigp1/Gbp | 5 | aPD-L1 vs PBS | Up-regulated genes | CD45- |
| CC | GO:0033646 | host intracellular part | 3.94E-09 | 4.39E-08 | Gbp2b/Iigp1/Gbp | 5 | aPD-L1 vs PBS | Up-regulated genes | CD45- |
| CC | GO:0043656 | intracellular region of host | 3.94E-09 | 4.39E-08 | Gbp2b/Iigp1/Gbp | 5 | aPD-L1 vs PBS | Up-regulated genes | CD45- |
| CC | GO:0033643 | host cell part | 6.87E-09 | 6.69E-08 | Gbp2b/Iigp1/Gbp | 5 | aPD-L1 vs PBS | Up-regulated genes | CD45- |
| CC | GO:0018995 | host | 1.13E-08 | 8.81E-08 | Gbp2b/Iigp1/Gbp | 5 | aPD-L1 vs PBS | Up-regulated genes | CD45- |
| CC | GO:0043657 | host cell | 1.13E-08 | 8.81E-08 | Gbp2b/Iigp1/Gbp | 5 | aPD-L1 vs PBS | Up-regulated genes | CD45- |
| CC | GO:0044215 | other organism | 2.19E-08 | 1.31E-07 | Gbp2b/Iigp1/Gbp | 5 | aPD-L1 vs PBS | Up-regulated genes | CD45- |
| CC | GO:0044216 | other organism cell | 2.19E-08 | 1.31E-07 | Gbp2b/Iigp1/Gbp | 5 | aPD-L1 vs PBS | Up-regulated genes | CD45- |
| CC | GO:0044217 | other organism part | 2.19E-08 | 1.31E-07 | Gbp2b/Iigp1/Gbp | 5 | aPD-L1 vs PBS | Up-regulated genes | CD45- |
| CC | GO:0043230 | extracellular organelle | 4.06E-07 | 2.26E-06 | Gbp2b/Iigp1/Gbp | 6 | aPD-L1 vs PBS | Up-regulated genes | CD45- |
| CC | GO:0042612 | MHC class I protein complex | 5.84E-06 | 3.04E-05 | B2m/H2-D1/H2-K | 3 | aPD-L1 vs PBS | Up-regulated genes | CD45- |
| CC | GO:0042824 | MHC class I peptide loading complex | 1.28E-05 | 6.23E-05 | B2m/H2-D1/H2-K | 3 | aPD-L1 vs PBS | Up-regulated genes | CD45- |
| CC | GO:0042611 | MHC protein complex | 3.36E-05 | 1.54E-04 | B2m/H2-D1/H2-K | 3 | aPD-L1 vs PBS | Up-regulated genes | CD45- |
| CC | GO:0031012 | extracellular matrix | 1.13E-03 | 4.91E-03 | Timp2/Emilin2/Dc | 6 | aPD-L1 vs PBS | Up-regulated genes | CD45- |
| CC | GO:0005581 | collagen trimer | 1.30E-03 | 5.34E-03 | Emilin2/Dcn/Scar | 3 | aPD-L1 vs PBS | Up-regulated genes | CD45- |
| CC | GO:0009897 | external side of plasma membrane | 1.38E-03 | 5.39E-03 | Cxcl10/B2m/Ly6a | 5 | aPD-L1 vs PBS | Up-regulated genes | CD45- |
| CC | GO:0005797 | Golgi medial cisterna | 2.70E-03 | 1.00E-02 | H2-D1/H2-K1 | 2 | aPD-L1 vs PBS | Up-regulated genes | CD45- |
| CC | GO:0070971 | endoplasmic reticulum exit site | 3.44E-03 | 1.22E-02 | H2-D1/H2-K1 | 2 | aPD-L1 vs PBS | Up-regulated genes | CD45- |
| CC | GO:0005719 | nuclear euchromatin | 4.88E-03 | 1.65E-02 | Jun/Klf4 | 2 | aPD-L1 vs PBS | Up-regulated genes | CD45- |
| CC | GO:0000791 | euchromatin | 7.64E-03 | 2.48E-02 | Jun/Klf4 | 2 | aPD-L1 vs PBS | Up-regulated genes | CD45- |
| CC | GO:0030176 | integral component of endoplasmic reticulum me | 9.17E-03 | 2.86E-02 | B2m/H2-D1/H2-K | 3 | aPD-L1 vs PBS | Up-regulated genes | CD45- |
| CC | GO:0098552 | side of membrane | 1.01E-02 | 3.04E-02 | Cxcl10/B2m/Ly6a | 5 | aPD-L1 vs PBS | Up-regulated genes | CD45- |
| CC | GO:0031227 | intrinsic component of endoplasmic reticulum me | 1.06E-02 | 3.06E-02 | B2m/H2-D1/H2-K | 3 | aPD-L1 vs PBS | Up-regulated genes | CD45- |
| CC | GO:0062023 | collagen-containing extracellular matrix | 1.52E-02 | 4.23E-02 | Timp2/Emilin2/Dc | 4 | aPD-L1 vs PBS | Up-regulated genes | CD45- |
| CC | GO:0031985 | Golgi cisterna | 1.58E-02 | 4.26E-02 | H2-D1/H2-K1 | 2 | aPD-L1 vs PBS | Up-regulated genes | CD45- |
| MF | GO:0005520 | insulin-like growth factor binding | 9.01E-05 | 1.29E-02 | Igfbp5/Htra3/Igfbp | 3 | aPD-L1 vs PBS | Up-regulated genes | CD45- |
| MF | GO:0030881 | beta-2-microglobulin binding | 6.59E-04 | 1.55E-02 | H2-D1/H2-K1 | 2 | aPD-L1 vs PBS | Up-regulated genes | CD45- |
| MF | GO:0031386 | protein tag | 6.59E-04 | 1.55E-02 | Ubc/Ubb | 2 | aPD-L1 vs PBS | Up-regulated genes | CD45- |
| MF | GO:0031994 | insulin-like growth factor I binding | 6.59E-04 | 1.55E-02 | Igfbp5/Igfbp4 | 2 | aPD-L1 vs PBS | Up-regulated genes | CD45- |
| MF | GO:0042610 | CD8 receptor binding | 6.59E-04 | 1.55E-02 | H2-D1/H2-K1 | 2 | aPD-L1 vs PBS | Up-regulated genes | CD45- |
| MF | GO:0046977 | TAP binding | 6.59E-04 | 1.55E-02 | H2-D1/H2-K1 | 2 | aPD-L1 vs PBS | Up-regulated genes | CD45- |
| MF | GO:0001228 | DNA-binding transcription activator activity, RNA | 7.60E-04 | 1.55E-02 | Irf1/Fosb/Egr1/Fo | 6 | aPD-L1 vs PBS | Up-regulated genes | CD45- |
| MF | GO:0042608 | T cell receptor binding | 1.25E-03 | 2.23E-02 | H2-D1/H2-K1 | 2 | aPD-L1 vs PBS | Up-regulated genes | CD45- |
| MF | GO:0003924 | GTPase activity | 2.02E-03 | 3.20E-02 | Gbp2b/Iigp1/Gbp | 5 | aPD-L1 vs PBS | Up-regulated genes | CD45- |
| MF | GO:0042605 | peptide antigen binding | 2.46E-03 | 3.51E-02 | H2-D1/H2-K1 | 2 | aPD-L1 vs PBS | Up-regulated genes | CD45- |
| MF | GO:0070412 | R-SMAD binding | 2.70E-03 | 3.51E-02 | Fos/Jun | 2 | aPD-L1 vs PBS | Up-regulated genes | CD45- |
| MF | GO:0000978 | RNA polymerase II proximal promoter sequence | 3.73E-03 | 3.91E-02 | Irf1/Atf3/Fosb/Fos | 6 | aPD-L1 vs PBS | Up-regulated genes | CD45- |
| MF | GO:0000987 | proximal promoter sequence-specific DNA bindi | 4.18E-03 | 3.91E-02 | Irf1/Atf3/Fosb/Fos | 6 | aPD-L1 vs PBS | Up-regulated genes | CD45- |
| MF | GO:0005525 | GTP binding | 4.65E-03 | 3.91E-02 | Gbp2b/Iigp1/Gbp | 5 | aPD-L1 vs PBS | Up-regulated genes | CD45- |
| MF | GO:0001968 | fibronectin binding | 4.99E-03 | 3.91E-02 | Igfbp5/Mmp2 | 2 | aPD-L1 vs PBS | Up-regulated genes | CD45- |
| MF | GO:0032550 | purine ribonucleoside binding | 5.04E-03 | 3.91E-02 | Gbp2b/Iigp1/Gbp | 5 | aPD-L1 vs PBS | Up-regulated genes | CD45- |
| MF | GO:0001883 | purine nucleoside binding | 5.18E-03 | 3.91E-02 | Gbp2b/Iigp1/Gbp | 5 | aPD-L1 vs PBS | Up-regulated genes | CD45- |
| MF | GO:0032549 | ribonucleoside binding | 5.38E-03 | 3.91E-02 | Gbp2b/Iigp1/Gbp | 5 | aPD-L1 vs PBS | Up-regulated genes | CD45- |
| MF | GO:0019001 | guanyl nucleotide binding | 5.60E-03 | 3.91E-02 | Gbp2b/Iigp1/Gbp | 5 | aPD-L1 vs PBS | Up-regulated genes | CD45- |
| MF | GO:0032561 | guanyl ribonucleotide binding | 5.60E-03 | 3.91E-02 | Gbp2b/Iigp1/Gbp | 5 | aPD-L1 vs PBS | Up-regulated genes | CD45- |

| | | | | | | | | | |
|---|---|---|---|---|---|---|---|---|---|
| MF | GO:0001882 | nucleoside binding | 5.97E-03 | 3.91E-02 | Gbp2b/Iigp1/Gbp2 | 5 | aPD-L1 vs PBS | Up-regulated genes | CD45- |
| MF | GO:0001102 | RNA polymerase II activating transcription factor | 6.02E-03 | 3.91E-02 | Fos/Jun | 2 | aPD-L1 vs PBS | Up-regulated genes | CD45- |
| MF | GO:0000979 | RNA polymerase II core promoter sequence-spe | 6.76E-03 | 4.02E-02 | Egr1/Fos | 2 | aPD-L1 vs PBS | Up-regulated genes | CD45- |
| MF | GO:0003823 | antigen binding | 6.76E-03 | 4.02E-02 | H2-D1/H2-K1 | 2 | aPD-L1 vs PBS | Up-regulated genes | CD45- |
| BP | GO:0043902 | positive regulation of multi-organism process | 3.01E-07 | 3.61E-04 | Park7/Ppia/Apoe/ | 10 | aPD-L1 vs PBS | Down-regulated gen | CD45+ |
| BP | GO:0051651 | maintenance of location in cell | 3.05E-07 | 3.61E-04 | Park7/Fth1/Ftl1/Tr | 8 | aPD-L1 vs PBS | Down-regulated gen | CD45+ |
| BP | GO:2000379 | positive regulation of reactive oxygen species m | 6.72E-07 | 5.31E-04 | Park7/Klf2/Gnai2/ | 8 | aPD-L1 vs PBS | Down-regulated gen | CD45+ |
| BP | GO:0009126 | purine nucleoside monophosphate metabolic pr | 1.78E-06 | 5.79E-04 | Park7/Atp5f1/Pkm | 11 | aPD-L1 vs PBS | Down-regulated gen | CD45+ |
| BP | GO:0009167 | purine ribonucleoside monophosphate metaboli | 1.78E-06 | 5.79E-04 | Park7/Atp5f1/Pkm | 11 | aPD-L1 vs PBS | Down-regulated gen | CD45+ |
| BP | GO:0009205 | purine ribonucleoside triphosphate metabolic pr | 1.86E-06 | 5.79E-04 | Park7/Atp5f1/Pkm | 11 | aPD-L1 vs PBS | Down-regulated gen | CD45+ |
| BP | GO:0016052 | carbohydrate catabolic process | 1.96E-06 | 5.79E-04 | Hexb/Akr1a1/Pkm | 8 | aPD-L1 vs PBS | Down-regulated gen | CD45+ |
| BP | GO:0009161 | ribonucleoside monophosphate metabolic proce | 2.11E-06 | 5.79E-04 | Park7/Atp5f1/Pkm | 11 | aPD-L1 vs PBS | Down-regulated gen | CD45+ |
| BP | GO:0009199 | ribonucleoside triphosphate metabolic process | 2.20E-06 | 5.79E-04 | Park7/Atp5f1/Pkm | 11 | aPD-L1 vs PBS | Down-regulated gen | CD45+ |
| BP | GO:0009144 | purine nucleoside triphosphate metabolic proce | 2.59E-06 | 6.06E-04 | Park7/Atp5f1/Pkm | 11 | aPD-L1 vs PBS | Down-regulated gen | CD45+ |
| BP | GO:0009123 | nucleoside monophosphate metabolic process | 2.81E-06 | 6.06E-04 | Park7/Atp5f1/Pkm | 11 | aPD-L1 vs PBS | Down-regulated gen | CD45+ |
| BP | GO:0009141 | nucleoside triphosphate metabolic process | 4.86E-06 | 8.91E-04 | Park7/Atp5f1/Pkm | 11 | aPD-L1 vs PBS | Down-regulated gen | CD45+ |
| BP | GO:0046034 | ATP metabolic process | 4.89E-06 | 8.91E-04 | Park7/Atp5f1/Pkm | 10 | aPD-L1 vs PBS | Down-regulated gen | CD45+ |
| BP | GO:0009127 | purine nucleoside monophosphate biosynthetic | 5.94E-06 | 9.38E-04 | Atp5f1/Pkm/Atp5g | 8 | aPD-L1 vs PBS | Down-regulated gen | CD45+ |
| BP | GO:0009168 | purine ribonucleoside monophosphate biosynth | 5.94E-06 | 9.38E-04 | Atp5f1/Pkm/Atp5g | 8 | aPD-L1 vs PBS | Down-regulated gen | CD45+ |
| BP | GO:0009156 | ribonucleoside monophosphate biosynthetic pro | 7.51E-06 | 1.11E-03 | Atp5f1/Pkm/Atp5g | 8 | aPD-L1 vs PBS | Down-regulated gen | CD45+ |
| BP | GO:0009124 | nucleoside monophosphate biosynthetic proces | 9.97E-06 | 1.34E-03 | Atp5f1/Pkm/Atp5g | 8 | aPD-L1 vs PBS | Down-regulated gen | CD45+ |
| BP | GO:2000377 | regulation of reactive oxygen species metabolic | 1.02E-05 | 1.34E-03 | Park7/Klf2/Gnai2/ | 9 | aPD-L1 vs PBS | Down-regulated gen | CD45+ |
| BP | GO:0043900 | regulation of multi-organism process | 1.43E-05 | 1.79E-03 | Park7/Ppia/Ifitm3/ | 12 | aPD-L1 vs PBS | Down-regulated gen | CD45+ |
| BP | GO:0009150 | purine ribonucleotide metabolic process | 1.52E-05 | 1.80E-03 | Park7/Atp5f1/Pkm | 12 | aPD-L1 vs PBS | Down-regulated gen | CD45+ |
| BP | GO:0006754 | ATP biosynthetic process | 1.66E-05 | 1.87E-03 | Atp5f1/Pkm/Atp5g | 7 | aPD-L1 vs PBS | Down-regulated gen | CD45+ |
| BP | GO:0043903 | regulation of symbiosis, encompassing mutualis | 1.73E-05 | 1.87E-03 | Ppia/Ifitm3/Apoe/ | 9 | aPD-L1 vs PBS | Down-regulated gen | CD45+ |
| BP | GO:0009259 | ribonucleotide metabolic process | 2.03E-05 | 2.09E-03 | Park7/Atp5f1/Pkm | 12 | aPD-L1 vs PBS | Down-regulated gen | CD45+ |
| BP | GO:0006163 | purine nucleotide metabolic process | 2.75E-05 | 2.71E-03 | Park7/Atp5f1/Pkm | 12 | aPD-L1 vs PBS | Down-regulated gen | CD45+ |
| BP | GO:0009206 | purine ribonucleoside triphosphate biosynthetic | 2.86E-05 | 2.71E-03 | Atp5f1/Pkm/Atp5g | 7 | aPD-L1 vs PBS | Down-regulated gen | CD45+ |
| BP | GO:0009145 | purine nucleoside triphosphate biosynthetic pro | 3.03E-05 | 2.73E-03 | Atp5f1/Pkm/Atp5g | 7 | aPD-L1 vs PBS | Down-regulated gen | CD45+ |
| BP | GO:0048524 | positive regulation of viral process | 3.15E-05 | 2.73E-03 | Ppia/Apoe/Pfn1/C | 6 | aPD-L1 vs PBS | Down-regulated gen | CD45+ |
| BP | GO:0019693 | ribose phosphate metabolic process | 3.23E-05 | 2.73E-03 | Park7/Atp5f1/Pkm | 12 | aPD-L1 vs PBS | Down-regulated gen | CD45+ |
| BP | GO:0009201 | ribonucleoside triphosphate biosynthetic proces | 3.39E-05 | 2.75E-03 | Atp5f1/Pkm/Atp5g | 7 | aPD-L1 vs PBS | Down-regulated gen | CD45+ |
| BP | GO:0050792 | regulation of viral process | 3.48E-05 | 2.75E-03 | Ppia/Ifitm3/Apoe/ | 8 | aPD-L1 vs PBS | Down-regulated gen | CD45+ |
| BP | GO:0051235 | maintenance of location | 3.86E-05 | 2.95E-03 | Park7/Hexb/Fth1/ | 10 | aPD-L1 vs PBS | Down-regulated gen | CD45+ |
| BP | GO:0044403 | symbiont process | 4.12E-05 | 3.05E-03 | Ppia/Ifitm3/Apoe/ | 11 | aPD-L1 vs PBS | Down-regulated gen | CD45+ |
| BP | GO:0032370 | positive regulation of lipid transport | 4.44E-05 | 3.19E-03 | Apoe/Prelid1/Nfkb | 5 | aPD-L1 vs PBS | Down-regulated gen | CD45+ |
| BP | GO:0072521 | purine-containing compound metabolic process | 5.53E-05 | 3.86E-03 | Park7/Atp5f1/Pkm | 12 | aPD-L1 vs PBS | Down-regulated gen | CD45+ |
| BP | GO:0009142 | nucleoside triphosphate biosynthetic process | 6.09E-05 | 4.07E-03 | Atp5f1/Pkm/Atp5g | 7 | aPD-L1 vs PBS | Down-regulated gen | CD45+ |
| BP | GO:0044419 | interspecies interaction between organisms | 6.25E-05 | 4.07E-03 | Ppia/Ifitm3/Apoe/ | 11 | aPD-L1 vs PBS | Down-regulated gen | CD45+ |
| BP | GO:0009152 | purine ribonucleotide biosynthetic process | 6.36E-05 | 4.07E-03 | Atp5f1/Pkm/Atp5g | 8 | aPD-L1 vs PBS | Down-regulated gen | CD45+ |
| BP | GO:0071277 | cellular response to calcium ion | 7.07E-05 | 4.41E-03 | Lgmn/Alox5ap/Ju | 5 | aPD-L1 vs PBS | Down-regulated gen | CD45+ |
| BP | GO:0009260 | ribonucleotide biosynthetic process | 9.10E-05 | 5.53E-03 | Atp5f1/Pkm/Atp5g | 8 | aPD-L1 vs PBS | Down-regulated gen | CD45+ |
| BP | GO:0072593 | reactive oxygen species metabolic process | 9.89E-05 | 5.73E-03 | Park7/Klf2/Gnai2/ | 9 | aPD-L1 vs PBS | Down-regulated gen | CD45+ |
| BP | GO:0042255 | ribosome assembly | 9.92E-05 | 5.73E-03 | Rpl5/Rps2/Rpl3/F | 5 | aPD-L1 vs PBS | Down-regulated gen | CD45+ |
| BP | GO:0015850 | organic hydroxy compound transport | 1.08E-04 | 6.11E-03 | Park7/Fcgr3/Apoe | 7 | aPD-L1 vs PBS | Down-regulated gen | CD45+ |
| BP | GO:0006164 | purine nucleotide biosynthetic process | 1.14E-04 | 6.11E-03 | Atp5f1/Pkm/Atp5g | 8 | aPD-L1 vs PBS | Down-regulated gen | CD45+ |
| BP | GO:0045185 | maintenance of protein location | 1.16E-04 | 6.11E-03 | Park7/Tmsb4x/Pfn | 6 | aPD-L1 vs PBS | Down-regulated gen | CD45+ |
| BP | GO:0051851 | modification by host of symbiont morphology or | 1.16E-04 | 6.11E-03 | Apoe/Gapdh/Cfl1/ | 5 | aPD-L1 vs PBS | Down-regulated gen | CD45+ |
| BP | GO:0046390 | ribose phosphate biosynthetic process | 1.19E-04 | 6.11E-03 | Atp5f1/Pkm/Atp5g | 8 | aPD-L1 vs PBS | Down-regulated gen | CD45+ |
| BP | GO:0035821 | modification of morphology or physiology of othe | 1.22E-04 | 6.17E-03 | Apoe/Gapdh/Cfl1/ | 6 | aPD-L1 vs PBS | Down-regulated gen | CD45+ |
| BP | GO:0072522 | purine-containing compound biosynthetic proces | 1.37E-04 | 6.77E-03 | Atp5f1/Pkm/Atp5g | 8 | aPD-L1 vs PBS | Down-regulated gen | CD45+ |
| BP | GO:0042273 | ribosomal large subunit biogenesis | 1.58E-04 | 7.63E-03 | Rpl5/Rpl14/Rpl3/ | 5 | aPD-L1 vs PBS | Down-regulated gen | CD45+ |
| BP | GO:0051702 | interaction with symbiont | 1.70E-04 | 8.04E-03 | Apoe/Gapdh/Cfl1/ | 5 | aPD-L1 vs PBS | Down-regulated gen | CD45+ |
| BP | GO:1905954 | positive regulation of lipid localization | 1.82E-04 | 8.46E-03 | Apoe/Prelid1/Nfkb | 5 | aPD-L1 vs PBS | Down-regulated gen | CD45+ |
| BP | GO:0046031 | ADP metabolic process | 2.24E-04 | 1.02E-02 | Pkm/Pgam1/Ier3/ | 5 | aPD-L1 vs PBS | Down-regulated gen | CD45+ |
| BP | GO:0032870 | cellular response to hormone stimulus | 2.35E-04 | 1.05E-02 | Park7/Pkm/Junb/ | 11 | aPD-L1 vs PBS | Down-regulated gen | CD45+ |
| BP | GO:0051881 | regulation of mitochondrial membrane potential | 2.56E-04 | 1.12E-02 | Park7/Clic1/Pid1/ | 5 | aPD-L1 vs PBS | Down-regulated gen | CD45+ |
| BP | GO:0009117 | nucleotide metabolic process | 2.89E-04 | 1.18E-02 | Park7/Atp5f1/Pkm | 12 | aPD-L1 vs PBS | Down-regulated gen | CD45+ |
| BP | GO:0009135 | purine nucleoside diphosphate metabolic proce | 2.91E-04 | 1.18E-02 | Pkm/Pgam1/Ier3/ | 5 | aPD-L1 vs PBS | Down-regulated gen | CD45+ |
| BP | GO:0009179 | purine ribonucleoside diphosphate metabolic pr | 2.91E-04 | 1.18E-02 | Pkm/Pgam1/Ier3/ | 5 | aPD-L1 vs PBS | Down-regulated gen | CD45+ |
| BP | GO:1903578 | regulation of ATP metabolic process | 2.91E-04 | 1.18E-02 | Park7/Pgam1/Pid | 5 | aPD-L1 vs PBS | Down-regulated gen | CD45+ |
| BP | GO:0046394 | carboxylic acid biosynthetic process | 2.96E-04 | 1.18E-02 | Park7/Akr1a1/Pkr | 9 | aPD-L1 vs PBS | Down-regulated gen | CD45+ |
| BP | GO:0035994 | response to muscle stretch | 3.00E-04 | 1.18E-02 | Fos/Nfkbia/Jun | 3 | aPD-L1 vs PBS | Down-regulated gen | CD45+ |
| BP | GO:0016053 | organic acid biosynthetic process | 3.04E-04 | 1.18E-02 | Park7/Akr1a1/Pkr | 9 | aPD-L1 vs PBS | Down-regulated gen | CD45+ |
| BP | GO:0006753 | nucleoside phosphate metabolic process | 3.25E-04 | 1.21E-02 | Park7/Atp5f1/Pkm | 12 | aPD-L1 vs PBS | Down-regulated gen | CD45+ |
| BP | GO:0002181 | cytoplasmic translation | 3.29E-04 | 1.21E-02 | Rps2/Rpl18/Rpl8/ | 5 | aPD-L1 vs PBS | Down-regulated gen | CD45+ |
| BP | GO:0010563 | negative regulation of phosphorus metabolic pro | 3.31E-04 | 1.21E-02 | Park7/Dusp1/Apo | 12 | aPD-L1 vs PBS | Down-regulated gen | CD45+ |
| BP | GO:0045936 | negative regulation of phosphate metabolic proc | 3.31E-04 | 1.21E-02 | Park7/Dusp1/Apo | 12 | aPD-L1 vs PBS | Down-regulated gen | CD45+ |
| BP | GO:0009185 | ribonucleoside diphosphate metabolic process | 3.72E-04 | 1.33E-02 | Pkm/Pgam1/Ier3/ | 5 | aPD-L1 vs PBS | Down-regulated gen | CD45+ |
| BP | GO:0052547 | regulation of peptidase activity | 3.86E-04 | 1.36E-02 | Park7/Rps3/Lgmr | 9 | aPD-L1 vs PBS | Down-regulated gen | CD45+ |
| BP | GO:0010035 | response to inorganic substance | 4.19E-04 | 1.46E-02 | Park7/Rps3/Lgmr | 10 | aPD-L1 vs PBS | Down-regulated gen | CD45+ |
| BP | GO:0032368 | regulation of lipid transport | 4.43E-04 | 1.50E-02 | Apoe/Prelid1/Nfkb | 5 | aPD-L1 vs PBS | Down-regulated gen | CD45+ |
| BP | GO:0051817 | modification of morphology or physiology of othe | 4.43E-04 | 1.50E-02 | Apoe/Gapdh/Cfl1/ | 5 | aPD-L1 vs PBS | Down-regulated gen | CD45+ |
| BP | GO:0009165 | nucleotide biosynthetic process | 4.66E-04 | 1.54E-02 | Atp5f1/Pkm/Atp5g | 8 | aPD-L1 vs PBS | Down-regulated gen | CD45+ |
| BP | GO:1900542 | regulation of purine nucleotide metabolic proces | 4.68E-04 | 1.54E-02 | Park7/Pgam1/Pid | 5 | aPD-L1 vs PBS | Down-regulated gen | CD45+ |
| BP | GO:0006090 | pyruvate metabolic process | 5.23E-04 | 1.70E-02 | Pkm/Pgam1/Ier3/ | 5 | aPD-L1 vs PBS | Down-regulated gen | CD45+ |
| BP | GO:1901293 | nucleoside phosphate biosynthetic process | 5.53E-04 | 1.76E-02 | Atp5f1/Pkm/Atp5g | 8 | aPD-L1 vs PBS | Down-regulated gen | CD45+ |
| BP | GO:0034614 | cellular response to reactive oxygen species | 5.66E-04 | 1.76E-02 | Park7/Rps3/Rhob | 6 | aPD-L1 vs PBS | Down-regulated gen | CD45+ |
| BP | GO:0097237 | cellular response to toxic substance | 5.66E-04 | 1.76E-02 | Park7/Akr1a1/Rps | 6 | aPD-L1 vs PBS | Down-regulated gen | CD45+ |
| BP | GO:0006140 | regulation of nucleotide metabolic process | 5.83E-04 | 1.80E-02 | Park7/Pgam1/Pid | 5 | aPD-L1 vs PBS | Down-regulated gen | CD45+ |
| BP | GO:2001233 | regulation of apoptotic signaling pathway | 6.76E-04 | 2.05E-02 | Park7/Rps3/Mcl1/ | 10 | aPD-L1 vs PBS | Down-regulated gen | CD45+ |
| BP | GO:0034404 | nucleobase-containing small molecule biosynth | 6.90E-04 | 2.05E-02 | Pkm/Pgam1/Aprt/ | 6 | aPD-L1 vs PBS | Down-regulated gen | CD45+ |
| BP | GO:0030595 | leukocyte chemotaxis | 6.91E-04 | 2.05E-02 | Dusp1/Lgmn/Fcgr | 7 | aPD-L1 vs PBS | Down-regulated gen | CD45+ |
| BP | GO:0009132 | nucleoside diphosphate metabolic process | 7.18E-04 | 2.10E-02 | Pkm/Pgam1/Ier3/ | 5 | aPD-L1 vs PBS | Down-regulated gen | CD45+ |
| BP | GO:0052548 | regulation of endopeptidase activity | 7.26E-04 | 2.10E-02 | Park7/Rps3/Lgmr | 8 | aPD-L1 vs PBS | Down-regulated gen | CD45+ |
| BP | GO:0002675 | positive regulation of acute inflammatory respon | 7.54E-04 | 2.10E-02 | Park7/Alox5ap/Fc | 3 | aPD-L1 vs PBS | Down-regulated gen | CD45+ |
| BP | GO:1900543 | negative regulation of purine nucleotide metabo | 7.54E-04 | 2.10E-02 | Pid1/Ier3/Tspo | 3 | aPD-L1 vs PBS | Down-regulated gen | CD45+ |
| BP | GO:1903579 | negative regulation of ATP metabolic process | 7.54E-04 | 2.10E-02 | Pid1/Ier3/Tspo | 3 | aPD-L1 vs PBS | Down-regulated gen | CD45+ |

| | | | | | | | | | |
|---|---|---|---|---|---|---|---|---|---|
| BP | GO:0051592 | response to calcium ion | 7.93E-04 | 2.17E-02 | Lgmn/Alox5ap/Ju | 5 | aPD-L1 vs PBS | Down-regulated gen | CD45+ |
| BP | GO:0006081 | cellular aldehyde metabolic process | 7.96E-04 | 2.17E-02 | Park7/Akr1a1/Pga | 4 | aPD-L1 vs PBS | Down-regulated gen | CD45+ |
| BP | GO:0016032 | viral process | 8.27E-04 | 2.23E-02 | Ppia/Ifitm3/Apoe/ | 8 | aPD-L1 vs PBS | Down-regulated gen | CD45+ |
| BP | GO:0045980 | negative regulation of nucleotide metabolic proc | 8.56E-04 | 2.28E-02 | Pid1/Ier3/Tspo | 3 | aPD-L1 vs PBS | Down-regulated gen | CD45+ |
| BP | GO:0046596 | regulation of viral entry into host cell | 9.68E-04 | 2.55E-02 | Ifitm3/Ifitm2/Lgals | 3 | aPD-L1 vs PBS | Down-regulated gen | CD45+ |
| BP | GO:0010038 | response to metal ion | 9.95E-04 | 2.59E-02 | Park7/Lgmn/Alox | 7 | aPD-L1 vs PBS | Down-regulated gen | CD45+ |
| BP | GO:0042254 | ribosome biogenesis | 1.06E-03 | 2.71E-02 | Rps6/Rpl5/Rpl14/ | 8 | aPD-L1 vs PBS | Down-regulated gen | CD45+ |
| BP | GO:0070997 | neuron death | 1.06E-03 | 2.71E-02 | Park7/Mcl1/Lgmn | 9 | aPD-L1 vs PBS | Down-regulated gen | CD45+ |
| BP | GO:0042326 | negative regulation of phosphorylation | 1.16E-03 | 2.91E-02 | Park7/Dusp1/Apo | 10 | aPD-L1 vs PBS | Down-regulated gen | CD45+ |
| BP | GO:2001234 | negative regulation of apoptotic signaling pathw | 1.18E-03 | 2.95E-02 | Park7/Mcl1/Gnai2 | 7 | aPD-L1 vs PBS | Down-regulated gen | CD45+ |
| BP | GO:0043467 | regulation of generation of precursor metabolites | 1.21E-03 | 2.98E-02 | Park7/Ppp1r3b/P | 5 | aPD-L1 vs PBS | Down-regulated gen | CD45+ |
| BP | GO:0009166 | nucleotide catabolic process | 1.32E-03 | 3.18E-02 | Pkm/Pgam1/Ier3/ | 5 | aPD-L1 vs PBS | Down-regulated gen | CD45+ |
| BP | GO:0000027 | ribosomal large subunit assembly | 1.35E-03 | 3.18E-02 | Rpl5/Rpl3/Rpl12 | 3 | aPD-L1 vs PBS | Down-regulated gen | CD45+ |
| BP | GO:0031640 | killing of cells of other organism | 1.35E-03 | 3.18E-02 | Gapdh/Pglyrp1/P | 3 | aPD-L1 vs PBS | Down-regulated gen | CD45+ |
| BP | GO:0044364 | disruption of cells of other organism | 1.35E-03 | 3.18E-02 | Gapdh/Pglyrp1/P | 3 | aPD-L1 vs PBS | Down-regulated gen | CD45+ |
| BP | GO:0071353 | cellular response to interleukin-4 | 1.35E-03 | 3.18E-02 | Nfil3/Rps2/Rpl3 | 3 | aPD-L1 vs PBS | Down-regulated gen | CD45+ |
| BP | GO:0008630 | intrinsic apoptotic signaling pathway in response | 1.44E-03 | 3.34E-02 | Shisa5/Rps3/Mcl | 5 | aPD-L1 vs PBS | Down-regulated gen | CD45+ |
| BP | GO:1905952 | regulation of lipid localization | 1.50E-03 | 3.44E-02 | Apoe/Prelid1/Nfk | 5 | aPD-L1 vs PBS | Down-regulated gen | CD45+ |
| BP | GO:0006096 | glycolytic process | 1.53E-03 | 3.44E-02 | Pkm/Pgam1/Ier3/ | 4 | aPD-L1 vs PBS | Down-regulated gen | CD45+ |
| BP | GO:0006757 | ATP generation from ADP | 1.53E-03 | 3.44E-02 | Pkm/Pgam1/Ier3/ | 4 | aPD-L1 vs PBS | Down-regulated gen | CD45+ |
| BP | GO:0062012 | regulation of small molecule metabolic process | 1.59E-03 | 3.56E-02 | Park7/Apoe/Ppp1 | 8 | aPD-L1 vs PBS | Down-regulated gen | CD45+ |
| BP | GO:0031099 | regeneration | 1.63E-03 | 3.61E-02 | Pkm/Ndel1/Plau/J | 5 | aPD-L1 vs PBS | Down-regulated gen | CD45+ |
| BP | GO:0070670 | response to interleukin-4 | 1.66E-03 | 3.64E-02 | Nfil3/Rps2/Rpl3 | 3 | aPD-L1 vs PBS | Down-regulated gen | CD45+ |
| BP | GO:0042176 | regulation of protein catabolic process | 1.68E-03 | 3.64E-02 | Park7/Oaz1/Rpl5/ | 9 | aPD-L1 vs PBS | Down-regulated gen | CD45+ |
| BP | GO:0042866 | pyruvate biosynthetic process | 1.71E-03 | 3.69E-02 | Pkm/Pgam1/Ier3/ | 4 | aPD-L1 vs PBS | Down-regulated gen | CD45+ |
| BP | GO:0006979 | response to oxidative stress | 1.74E-03 | 3.72E-02 | Park7/Rps3/Mcl1/ | 9 | aPD-L1 vs PBS | Down-regulated gen | CD45+ |
| BP | GO:1901292 | nucleoside phosphate catabolic process | 1.91E-03 | 4.05E-02 | Pkm/Pgam1/Ier3/ | 5 | aPD-L1 vs PBS | Down-regulated gen | CD45+ |
| BP | GO:0045862 | positive regulation of proteolysis | 1.99E-03 | 4.16E-02 | Rps3/Lgmn/Apoe | 8 | aPD-L1 vs PBS | Down-regulated gen | CD45+ |
| BP | GO:0032507 | maintenance of protein location in cell | 2.14E-03 | 4.38E-02 | Park7/Tmsb4x/Pfr | 4 | aPD-L1 vs PBS | Down-regulated gen | CD45+ |
| BP | GO:0071248 | cellular response to metal ion | 2.15E-03 | 4.38E-02 | Lgmn/Alox5ap/Ju | 5 | aPD-L1 vs PBS | Down-regulated gen | CD45+ |
| BP | GO:0001933 | negative regulation of protein phosphorylation | 2.15E-03 | 4.38E-02 | Park7/Dusp1/Apo | 9 | aPD-L1 vs PBS | Down-regulated gen | CD45+ |
| BP | GO:0009991 | response to extracellular stimulus | 2.16E-03 | 4.38E-02 | Slc2a1/Apoe/Gatr | 8 | aPD-L1 vs PBS | Down-regulated gen | CD45+ |
| BP | GO:0051345 | positive regulation of hydrolase activity | 2.24E-03 | 4.49E-02 | Rps3/Lgmn/Rpl5/ | 10 | aPD-L1 vs PBS | Down-regulated gen | CD45+ |
| BP | GO:0010810 | regulation of cell-substrate adhesion | 2.33E-03 | 4.63E-02 | S100a10/Cfl1/Lga | 6 | aPD-L1 vs PBS | Down-regulated gen | CD45+ |
| BP | GO:0031397 | negative regulation of protein ubiquitination | 2.37E-03 | 4.68E-02 | Park7/Rps3/Rpl5/ | 4 | aPD-L1 vs PBS | Down-regulated gen | CD45+ |
| BP | GO:0051402 | neuron apoptotic process | 2.51E-03 | 4.92E-02 | Park7/Mcl1/Lgmn | 7 | aPD-L1 vs PBS | Down-regulated gen | CD45+ |
| BP | GO:0034599 | cellular response to oxidative stress | 2.57E-03 | 4.97E-02 | Park7/Rps3/Mcl1/ | 7 | aPD-L1 vs PBS | Down-regulated gen | CD45+ |
| BP | GO:0061844 | antimicrobial humoral immune response mediat | 2.60E-03 | 4.97E-02 | Gapdh/Pglyrp1/P | 3 | aPD-L1 vs PBS | Down-regulated gen | CD45+ |
| BP | GO:0000302 | response to reactive oxygen species | 2.61E-03 | 4.97E-02 | Park7/Rps3/Rhob | 6 | aPD-L1 vs PBS | Down-regulated gen | CD45+ |
| BP | GO:0043471 | regulation of cellular carbohydrate catabolic pro | 2.66E-03 | 4.97E-02 | Ppp1r3b/Pgam1 | 2 | aPD-L1 vs PBS | Down-regulated gen | CD45+ |
| BP | GO:0006091 | generation of precursor metabolites and energy | 2.66E-03 | 4.97E-02 | Park7/Pkm/Ppp1r | 8 | aPD-L1 vs PBS | Down-regulated gen | CD45+ |
| BP | GO:1901652 | response to peptide | 2.66E-03 | 4.97E-02 | Pkm/Lgmn/Ndel1/ | 8 | aPD-L1 vs PBS | Down-regulated gen | CD45+ |
| CC | GO:0022626 | cytosolic ribosome | 2.46E-15 | 6.51E-13 | Rps6/Rps3/Rpl5/ | 15 | aPD-L1 vs PBS | Down-regulated gen | CD45+ |
| CC | GO:0044391 | ribosomal subunit | 1.24E-11 | 1.64E-09 | Rps6/Rps3/Rpl5/ | 15 | aPD-L1 vs PBS | Down-regulated gen | CD45+ |
| CC | GO:0005840 | ribosome | 1.14E-10 | 1.01E-08 | Rps6/Rps3/Rpl5/ | 15 | aPD-L1 vs PBS | Down-regulated gen | CD45+ |
| CC | GO:0044445 | cytosolic part | 3.54E-10 | 2.35E-08 | Rps6/Rps3/Rpl5/ | 15 | aPD-L1 vs PBS | Down-regulated gen | CD45+ |
| CC | GO:0022625 | cytosolic large ribosomal subunit | 6.05E-10 | 3.21E-08 | Rpl5/Rpl14/Rpl1 | 9 | aPD-L1 vs PBS | Down-regulated gen | CD45+ |
| CC | GO:0015934 | large ribosomal subunit | 3.45E-07 | 1.52E-05 | Rpl5/Rpl14/Rpl1 | 9 | aPD-L1 vs PBS | Down-regulated gen | CD45+ |
| CC | GO:0022627 | cytosolic small ribosomal subunit | 8.33E-07 | 3.15E-05 | Rps6/Rps3/Rps3 | 6 | aPD-L1 vs PBS | Down-regulated gen | CD45+ |
| CC | GO:0005844 | polysome | 1.47E-05 | 4.88E-04 | Rps6/Rps3/Rpl18 | 6 | aPD-L1 vs PBS | Down-regulated gen | CD45+ |
| CC | GO:0015935 | small ribosomal subunit | 2.05E-05 | 6.04E-04 | Rps6/Rps3/Rps3 | 6 | aPD-L1 vs PBS | Down-regulated gen | CD45+ |
| CC | GO:0043209 | myelin sheath | 6.81E-04 | 1.81E-02 | Ppia/Atp5f1/Pkm/ | 7 | aPD-L1 vs PBS | Down-regulated gen | CD45+ |
| CC | GO:0099572 | postsynaptic specialization | 8.10E-04 | 1.95E-02 | Rps3/Rpl5/Rpl14/ | 9 | aPD-L1 vs PBS | Down-regulated gen | CD45+ |
| CC | GO:0042788 | polysomal ribosome | 1.32E-03 | 2.93E-02 | Rpl18/Rpl8/Rpl1 | 3 | aPD-L1 vs PBS | Down-regulated gen | CD45+ |
| CC | GO:0031012 | extracellular matrix | 1.55E-03 | 3.16E-02 | Pkm/S100a6/Apo | 8 | aPD-L1 vs PBS | Down-regulated gen | CD45+ |
| CC | GO:0005635 | nuclear envelope | 2.12E-03 | 3.65E-02 | Shisa5/Clic1/Alox | 9 | aPD-L1 vs PBS | Down-regulated gen | CD45+ |
| CC | GO:0062023 | collagen-containing extracellular matrix | 2.13E-03 | 3.65E-02 | Pkm/S100a6/S10 | 7 | aPD-L1 vs PBS | Down-regulated gen | CD45+ |
| CC | GO:0014069 | postsynaptic density | 2.25E-03 | 3.65E-02 | Rps3/Rpl5/Rpl14/ | 8 | aPD-L1 vs PBS | Down-regulated gen | CD45+ |
| CC | GO:0032279 | asymmetric synapse | 2.34E-03 | 3.65E-02 | Rps3/Rpl5/Rpl14/ | 8 | aPD-L1 vs PBS | Down-regulated gen | CD45+ |
| CC | GO:0000276 | mitochondrial proton-transporting ATP synthase | 2.62E-03 | 3.86E-02 | Atp5f1/Atp5g2 | 2 | aPD-L1 vs PBS | Down-regulated gen | CD45+ |
| CC | GO:0098984 | neuron to neuron synapse | 3.04E-03 | 4.10E-02 | Rps3/Rpl5/Rpl14/ | 8 | aPD-L1 vs PBS | Down-regulated gen | CD45+ |
| CC | GO:0044754 | autolysosome | 3.19E-03 | 4.10E-02 | Fth1/Ftl1 | 2 | aPD-L1 vs PBS | Down-regulated gen | CD45+ |
| CC | GO:0005938 | cell cortex | 3.25E-03 | 4.10E-02 | Cap1/Slc2a1/Pfn | 7 | aPD-L1 vs PBS | Down-regulated gen | CD45+ |
| CC | GO:0005767 | secondary lysosome | 3.81E-03 | 4.39E-02 | Fth1/Ftl1 | 2 | aPD-L1 vs PBS | Down-regulated gen | CD45+ |
| CC | GO:0045263 | proton-transporting ATP synthase complex, coup | 3.81E-03 | 4.39E-02 | Atp5f1/Atp5g2 | 2 | aPD-L1 vs PBS | Down-regulated gen | CD45+ |
| CC | GO:0098685 | Schaffer collateral - CA1 synapse | 4.06E-03 | 4.48E-02 | Pfn1/Srgn/Actb/A | 4 | aPD-L1 vs PBS | Down-regulated gen | CD45+ |
| MF | GO:0003735 | structural constituent of ribosome | 3.10E-13 | 9.88E-11 | Rps6/Rps3/Rpl5/ | 15 | aPD-L1 vs PBS | Down-regulated gen | CD45+ |
| MF | GO:0005198 | structural molecule activity | 4.65E-10 | 7.42E-08 | Rps6/Rps3/Arpc1 | 19 | aPD-L1 vs PBS | Down-regulated gen | CD45+ |
| MF | GO:0019843 | rRNA binding | 8.98E-07 | 9.54E-05 | Rps3/Rpl5/Rpl3/ | 7 | aPD-L1 vs PBS | Down-regulated gen | CD45+ |
| MF | GO:0005125 | cytokine activity | 4.55E-04 | 3.32E-02 | Ltb/Ccl6/Cxcl14/ | 6 | aPD-L1 vs PBS | Down-regulated gen | CD45+ |
| MF | GO:0070412 | R-SMAD binding | 5.20E-04 | 3.32E-02 | Fos/Jun/Pmepa1 | 3 | aPD-L1 vs PBS | Down-regulated gen | CD45+ |
| BP | GO:0009617 | response to bacterium | 1.21E-18 | 1.80E-15 | Cxcl10/Ifit1/Gbp2 | 24 | aTGF-β vs PBS | Up-regulated genes | CD45+ |
| BP | GO:0098542 | defense response to other organism | 3.96E-18 | 2.94E-15 | Cxcl10/Rsad2/Ifit | 22 | aTGF-β vs PBS | Up-regulated genes | CD45+ |
| BP | GO:0009615 | response to virus | 9.83E-17 | 4.02E-14 | Cxcl10/Rsad2/Ifit | 18 | aTGF-β vs PBS | Up-regulated genes | CD45+ |
| BP | GO:0051607 | defense response to virus | 1.08E-16 | 4.02E-14 | Cxcl10/Rsad2/Ifit | 17 | aTGF-β vs PBS | Up-regulated genes | CD45+ |
| BP | GO:0035456 | response to interferon-beta | 2.67E-15 | 7.94E-13 | Ifit1/Gbp2b/Gbp2/ | 11 | aTGF-β vs PBS | Up-regulated genes | CD45+ |
| BP | GO:0035458 | cellular response to interferon-beta | 2.15E-14 | 5.32E-12 | Ifit1/Gbp2b/Gbp2/ | 10 | aTGF-β vs PBS | Up-regulated genes | CD45+ |
| BP | GO:0034341 | response to interferon-gamma | 1.06E-12 | 2.25E-10 | Gbp2b/Parp14/G | 12 | aTGF-β vs PBS | Up-regulated genes | CD45+ |
| BP | GO:0071346 | cellular response to interferon-gamma | 4.10E-12 | 7.62E-10 | Gbp2b/Parp14/G | 11 | aTGF-β vs PBS | Up-regulated genes | CD45+ |
| BP | GO:0044406 | adhesion of symbiont to host | 2.37E-11 | 3.92E-09 | Gbp2b/Gbp2/Hsp | 6 | aTGF-β vs PBS | Up-regulated genes | CD45+ |
| BP | GO:0044403 | symbiont process | 4.82E-11 | 7.17E-09 | Rsad2/Gbp2b/Gb | 15 | aTGF-β vs PBS | Up-regulated genes | CD45+ |
| BP | GO:0044419 | interspecies interaction between organisms | 9.23E-11 | 1.25E-08 | Rsad2/Gbp2b/Gb | 15 | aTGF-β vs PBS | Up-regulated genes | CD45+ |
| BP | GO:0045088 | regulation of innate immune response | 5.15E-10 | 6.39E-08 | Rsad2/Parp14/G | 13 | aTGF-β vs PBS | Up-regulated genes | CD45+ |
| BP | GO:0019221 | cytokine-mediated signaling pathway | 1.25E-07 | 1.43E-05 | Cxcl10/Parp14/Ifi | 11 | aTGF-β vs PBS | Up-regulated genes | CD45+ |
| BP | GO:0016032 | viral process | 5.30E-07 | 5.63E-05 | Rsad2/Ifitm3/Isg1 | 10 | aTGF-β vs PBS | Up-regulated genes | CD45+ |
| BP | GO:0034340 | response to type I interferon | 1.00E-06 | 9.56E-05 | Ifitm3/Isg15/Irf7/S | 5 | aTGF-β vs PBS | Up-regulated genes | CD45+ |
| BP | GO:0002683 | negative regulation of immune system process | 1.03E-06 | 9.56E-05 | Parp14/Irf1/Samh | 12 | aTGF-β vs PBS | Up-regulated genes | CD45+ |
| BP | GO:0045089 | positive regulation of innate immune response | 1.37E-06 | 1.20E-04 | Rsad2/Gbp5/Irf1/ | 9 | aTGF-β vs PBS | Up-regulated genes | CD45+ |
| BP | GO:0019058 | viral life cycle | 2.76E-06 | 2.28E-04 | Rsad2/Ifitm3/Isg1 | 8 | aTGF-β vs PBS | Up-regulated genes | CD45+ |

| | | | | | | | | | |
|---|---|---|---|---|---|---|---|---|---|
| BP | GO:0002819 | regulation of adaptive immune response | 3.00E-06 | 2.35E-04 | Rsad2/Irf1/Irf7/Cd | 8 | aTGF-β vs PBS | Up-regulated genes | CD45+ |
| BP | GO:0048525 | negative regulation of viral process | 4.23E-06 | 3.07E-04 | Rsad2/Ifitm3/Isg1 | 6 | aTGF-β vs PBS | Up-regulated genes | CD45+ |
| BP | GO:0045824 | negative regulation of innate immune response | 4.33E-06 | 3.07E-04 | Parp14/Samhd1/ | 5 | aTGF-β vs PBS | Up-regulated genes | CD45+ |
| BP | GO:0035455 | response to interferon-alpha | 5.13E-06 | 3.47E-04 | Ifit1/Ifit2/Ifit3/Ifitm3 | 4 | aTGF-β vs PBS | Up-regulated genes | CD45+ |
| BP | GO:0050777 | negative regulation of immune response | 5.75E-06 | 3.64E-04 | Parp14/Irf1/Samh | 7 | aTGF-β vs PBS | Up-regulated genes | CD45+ |
| BP | GO:0045071 | negative regulation of viral genome replication | 5.88E-06 | 3.64E-04 | Rsad2/Ifitm3/Isg1 | 5 | aTGF-β vs PBS | Up-regulated genes | CD45+ |
| BP | GO:0002460 | adaptive immune response based on somatic re | 6.51E-06 | 3.87E-04 | Rsad2/Irf1/Irf7/Cd | 9 | aTGF-β vs PBS | Up-regulated genes | CD45+ |
| BP | GO:0043901 | negative regulation of multi-organism process | 1.10E-05 | 6.27E-04 | Rsad2/Ifitm3/Isg1 | 7 | aTGF-β vs PBS | Up-regulated genes | CD45+ |
| BP | GO:0002250 | adaptive immune response | 1.21E-05 | 6.67E-04 | Rsad2/Irf1/Irf7/Cd | 10 | aTGF-β vs PBS | Up-regulated genes | CD45+ |
| BP | GO:0034121 | regulation of toll-like receptor signaling pathway | 1.32E-05 | 7.00E-04 | Rsad2/Irf1/Irf7/Ac | 5 | aTGF-β vs PBS | Up-regulated genes | CD45+ |
| BP | GO:0042742 | defense response to bacterium | 1.54E-05 | 7.62E-04 | Gbp2b/Gbp2/Plac | 7 | aTGF-β vs PBS | Up-regulated genes | CD45+ |
| BP | GO:0050792 | regulation of viral process | 1.54E-05 | 7.62E-04 | Rsad2/Ifitm3/Isg1 | 7 | aTGF-β vs PBS | Up-regulated genes | CD45+ |
| BP | GO:0071222 | cellular response to lipopolysaccharide | 1.67E-05 | 7.97E-04 | Cxcl10/Gbp2/Cm | 7 | aTGF-β vs PBS | Up-regulated genes | CD45+ |
| BP | GO:0060337 | type I interferon signaling pathway | 1.77E-05 | 7.97E-04 | Ifitm3/Irf7/Samhd1 | 4 | aTGF-β vs PBS | Up-regulated genes | CD45+ |
| BP | GO:0071357 | cellular response to type I interferon | 1.77E-05 | 7.97E-04 | Ifitm3/Irf7/Samhd1 | 4 | aTGF-β vs PBS | Up-regulated genes | CD45+ |
| BP | GO:0071219 | cellular response to molecule of bacterial origin | 2.20E-05 | 9.63E-04 | Cxcl10/Gbp2/Cm | 7 | aTGF-β vs PBS | Up-regulated genes | CD45+ |
| BP | GO:0042832 | defense response to protozoan | 2.28E-05 | 9.67E-04 | Gbp2b/Gbp2/Gbp | 4 | aTGF-β vs PBS | Up-regulated genes | CD45+ |
| BP | GO:0002824 | positive regulation of adaptive immune response | 2.34E-05 | 9.67E-04 | Rsad2/Irf1/Cd274 | 6 | aTGF-β vs PBS | Up-regulated genes | CD45+ |
| BP | GO:0035457 | cellular response to interferon-alpha | 2.72E-05 | 1.09E-03 | Ifit1/Ifit2/Ifit3 | 3 | aTGF-β vs PBS | Up-regulated genes | CD45+ |
| BP | GO:0001562 | response to protozoan | 2.90E-05 | 1.13E-03 | Gbp2b/Gbp2/Gbp | 4 | aTGF-β vs PBS | Up-regulated genes | CD45+ |
| BP | GO:1903901 | negative regulation of viral life cycle | 3.01E-05 | 1.15E-03 | Rsad2/Ifitm3/Isg1 | 5 | aTGF-β vs PBS | Up-regulated genes | CD45+ |
| BP | GO:0002237 | response to molecule of bacterial origin | 3.17E-05 | 1.15E-03 | Cxcl10/Gbp2/Cm | 8 | aTGF-β vs PBS | Up-regulated genes | CD45+ |
| BP | GO:0002449 | lymphocyte mediated immunity | 3.17E-05 | 1.15E-03 | Rsad2/Irf7/H2-D1 | 8 | aTGF-β vs PBS | Up-regulated genes | CD45+ |
| BP | GO:0002821 | positive regulation of adaptive immune response | 3.29E-05 | 1.16E-03 | Rsad2/Irf1/Cd274 | 6 | aTGF-β vs PBS | Up-regulated genes | CD45+ |
| BP | GO:0031349 | positive regulation of defense response | 3.84E-05 | 1.33E-03 | Rsad2/Gbp5/Irf1/I | 9 | aTGF-β vs PBS | Up-regulated genes | CD45+ |
| BP | GO:1903900 | regulation of viral life cycle | 4.14E-05 | 1.40E-03 | Rsad2/Ifitm3/Isg1 | 6 | aTGF-β vs PBS | Up-regulated genes | CD45+ |
| BP | GO:0043903 | regulation of symbiosis, encompassing mutualis | 4.71E-05 | 1.52E-03 | Rsad2/Ifitm3/Isg1 | 7 | aTGF-β vs PBS | Up-regulated genes | CD45+ |
| BP | GO:0071216 | cellular response to biotic stimulus | 4.71E-05 | 1.52E-03 | Cxcl10/Gbp2/Cm | 7 | aTGF-β vs PBS | Up-regulated genes | CD45+ |
| BP | GO:0002221 | pattern recognition receptor signaling pathway | 5.16E-05 | 1.63E-03 | Rsad2/Irf1/Irf7/Ac | 6 | aTGF-β vs PBS | Up-regulated genes | CD45+ |
| BP | GO:0043900 | regulation of multi-organism process | 5.33E-05 | 1.65E-03 | Rsad2/Ifitm3/Isg1 | 9 | aTGF-β vs PBS | Up-regulated genes | CD45+ |
| BP | GO:0002758 | innate immune response-activating signal trans | 5.62E-05 | 1.69E-03 | Rsad2/Irf1/Irf7/Ac | 6 | aTGF-β vs PBS | Up-regulated genes | CD45+ |
| BP | GO:0045069 | regulation of viral genome replication | 5.69E-05 | 1.69E-03 | Rsad2/Ifitm3/Isg1 | 5 | aTGF-β vs PBS | Up-regulated genes | CD45+ |
| BP | GO:0001961 | positive regulation of cytokine-mediated signalin | 6.62E-05 | 1.93E-03 | Parp14/Irgm1/Irf7 | 4 | aTGF-β vs PBS | Up-regulated genes | CD45+ |
| BP | GO:0019882 | antigen processing and presentation | 8.02E-05 | 2.29E-03 | H2-D1/H2-K1/B2 | 5 | aTGF-β vs PBS | Up-regulated genes | CD45+ |
| BP | GO:0001959 | regulation of cytokine-mediated signaling pathw | 1.10E-04 | 3.03E-03 | Parp14/Irgm1/Irf7 | 5 | aTGF-β vs PBS | Up-regulated genes | CD45+ |
| BP | GO:2000379 | positive regulation of reactive oxygen species m | 1.10E-04 | 3.03E-03 | Acod1/Klf2/Sod2/ | 5 | aTGF-β vs PBS | Up-regulated genes | CD45+ |
| BP | GO:0002428 | antigen processing and presentation of peptide | 1.17E-04 | 3.04E-03 | H2-D1/H2-K1/B2 | 3 | aTGF-β vs PBS | Up-regulated genes | CD45+ |
| BP | GO:0060338 | regulation of type I interferon-mediated signaling | 1.17E-04 | 3.04E-03 | Irf7/Samhd1/Zbp1 | 3 | aTGF-β vs PBS | Up-regulated genes | CD45+ |
| BP | GO:1901623 | regulation of lymphocyte chemotaxis | 1.17E-04 | 3.04E-03 | Cxcl10/Ccl4/Ccl1 | 3 | aTGF-β vs PBS | Up-regulated genes | CD45+ |
| BP | GO:0060760 | positive regulation of response to cytokine stimu | 1.20E-04 | 3.08E-03 | Parp14/Irgm1/Irf7 | 4 | aTGF-β vs PBS | Up-regulated genes | CD45+ |
| BP | GO:0002224 | toll-like receptor signaling pathway | 1.34E-04 | 3.33E-03 | Rsad2/Irf1/Irf7/Ac | 5 | aTGF-β vs PBS | Up-regulated genes | CD45+ |
| BP | GO:0019079 | viral genome replication | 1.34E-04 | 3.33E-03 | Rsad2/Ifitm3/Isg1 | 5 | aTGF-β vs PBS | Up-regulated genes | CD45+ |
| BP | GO:0060333 | interferon-gamma-mediated signaling pathway | 1.37E-04 | 3.33E-03 | Parp14/Irf1/Irgm1 | 3 | aTGF-β vs PBS | Up-regulated genes | CD45+ |
| BP | GO:0001819 | positive regulation of cytokine production | 1.41E-04 | 3.34E-03 | Rsad2/Gbp5/Irf1/I | 9 | aTGF-β vs PBS | Up-regulated genes | CD45+ |
| BP | GO:0002218 | activation of innate immune response | 1.42E-04 | 3.34E-03 | Rsad2/Irf1/Irf7/Ac | 6 | aTGF-β vs PBS | Up-regulated genes | CD45+ |
| BP | GO:0032496 | response to lipopolysaccharide | 1.49E-04 | 3.47E-03 | Cxcl10/Gbp2/Cm | 7 | aTGF-β vs PBS | Up-regulated genes | CD45+ |
| BP | GO:0002711 | positive regulation of T cell mediated immunity | 1.63E-04 | 3.61E-03 | Rsad2/H2-D1/H2 | 4 | aTGF-β vs PBS | Up-regulated genes | CD45+ |
| BP | GO:0031341 | regulation of cell killing | 1.63E-04 | 3.61E-03 | Hsp90ab1/H2-D1 | 5 | aTGF-β vs PBS | Up-regulated genes | CD45+ |
| BP | GO:0002822 | regulation of adaptive immune response based | 1.63E-04 | 3.61E-03 | Rsad2/Irf1/Cd274 | 6 | aTGF-β vs PBS | Up-regulated genes | CD45+ |
| BP | GO:0002475 | antigen processing and presentation via MHC cl | 1.83E-04 | 3.97E-03 | H2-D1/H2-K1/B2 | 3 | aTGF-β vs PBS | Up-regulated genes | CD45+ |
| BP | GO:0002443 | leukocyte mediated immunity | 1.86E-04 | 3.97E-03 | Rsad2/Irf7/H2-D1 | 8 | aTGF-β vs PBS | Up-regulated genes | CD45+ |
| BP | GO:0060759 | regulation of response to cytokine stimulus | 1.87E-04 | 3.97E-03 | Parp14/Irgm1/Irf7 | 5 | aTGF-β vs PBS | Up-regulated genes | CD45+ |
| BP | GO:0002483 | antigen processing and presentation of endoger | 2.39E-04 | 4.94E-03 | H2-D1/H2-K1/B2 | 3 | aTGF-β vs PBS | Up-regulated genes | CD45+ |
| BP | GO:0019885 | antigen processing and presentation of endoger | 2.39E-04 | 4.94E-03 | H2-D1/H2-K1/B2 | 3 | aTGF-β vs PBS | Up-regulated genes | CD45+ |
| BP | GO:2000377 | regulation of reactive oxygen species metabolic | 2.73E-04 | 5.53E-03 | Acod1/Klf2/Trim3 | 6 | aTGF-β vs PBS | Up-regulated genes | CD45+ |
| BP | GO:0002456 | T cell mediated immunity | 2.75E-04 | 5.53E-03 | Rsad2/H2-D1/H2 | 5 | aTGF-β vs PBS | Up-regulated genes | CD45+ |
| BP | GO:0019883 | antigen processing and presentation of endoger | 3.05E-04 | 6.05E-03 | H2-D1/H2-K1/B2 | 3 | aTGF-β vs PBS | Up-regulated genes | CD45+ |
| BP | GO:0050830 | defense response to Gram-positive bacterium | 3.75E-04 | 7.34E-03 | Gbp2b/Gbp2/Gbp | 4 | aTGF-β vs PBS | Up-regulated genes | CD45+ |
| BP | GO:0071347 | cellular response to interleukin-1 | 3.97E-04 | 7.67E-03 | Irf1/Ccl4/Acod1/C | 4 | aTGF-β vs PBS | Up-regulated genes | CD45+ |
| BP | GO:0071674 | mononuclear cell migration | 4.20E-04 | 7.98E-03 | Cxcl10/Ccl4/Ccl1 | 4 | aTGF-β vs PBS | Up-regulated genes | CD45+ |
| BP | GO:0042026 | protein refolding | 4.24E-04 | 7.98E-03 | Hspa1a/Hspa1b/ | 3 | aTGF-β vs PBS | Up-regulated genes | CD45+ |
| BP | GO:0002687 | positive regulation of leukocyte migration | 5.07E-04 | 9.41E-03 | Cxcl10/Ccl4/Ccl1 | 5 | aTGF-β vs PBS | Up-regulated genes | CD45+ |
| BP | GO:0001916 | positive regulation of T cell mediated cytotoxicity | 6.24E-04 | 1.13E-02 | H2-D1/H2-K1/B2 | 3 | aTGF-β vs PBS | Up-regulated genes | CD45+ |
| BP | GO:0002474 | antigen processing and presentation of peptide | 6.24E-04 | 1.13E-02 | H2-D1/H2-K1/B2 | 3 | aTGF-β vs PBS | Up-regulated genes | CD45+ |
| BP | GO:0002709 | regulation of T cell mediated immunity | 6.38E-04 | 1.14E-02 | Rsad2/H2-D1/H2 | 4 | aTGF-β vs PBS | Up-regulated genes | CD45+ |
| BP | GO:0001906 | cell killing | 6.66E-04 | 1.18E-02 | Hsp90ab1/H2-D1 | 5 | aTGF-β vs PBS | Up-regulated genes | CD45+ |
| BP | GO:0002699 | positive regulation of immune effector process | 7.33E-04 | 1.28E-02 | Rsad2/Irf1/H2-D1 | 6 | aTGF-β vs PBS | Up-regulated genes | CD45+ |
| BP | GO:1902041 | regulation of extrinsic apoptotic signaling pathwa | 8.08E-04 | 1.38E-02 | Atf3/Icam1/Thbs1 | 3 | aTGF-β vs PBS | Up-regulated genes | CD45+ |
| BP | GO:0001910 | regulation of leukocyte mediated cytotoxicity | 8.08E-04 | 1.38E-02 | H2-D1/H2-K1/B2 | 4 | aTGF-β vs PBS | Up-regulated genes | CD45+ |
| BP | GO:0070555 | response to interleukin-1 | 9.25E-04 | 1.56E-02 | Irf1/Ccl4/Acod1/C | 4 | aTGF-β vs PBS | Up-regulated genes | CD45+ |
| BP | GO:0001914 | regulation of T cell mediated cytotoxicity | 9.47E-04 | 1.58E-02 | H2-D1/H2-K1/B2 | 3 | aTGF-β vs PBS | Up-regulated genes | CD45+ |
| BP | GO:0072676 | lymphocyte migration | 9.67E-04 | 1.60E-02 | Cxcl10/Ccl4/Ccl1 | 4 | aTGF-β vs PBS | Up-regulated genes | CD45+ |
| BP | GO:0045429 | positive regulation of nitric oxide biosynthetic pr | 1.18E-03 | 1.93E-02 | Klf2/Sod2/Icam1 | 3 | aTGF-β vs PBS | Up-regulated genes | CD45+ |
| BP | GO:0001818 | negative regulation of cytokine production | 1.22E-03 | 1.98E-02 | Hsp90ab1/Acod1 | 6 | aTGF-β vs PBS | Up-regulated genes | CD45+ |
| BP | GO:0072593 | reactive oxygen species metabolic process | 1.25E-03 | 2.00E-02 | Acod1/Klf2/Trim3 | 6 | aTGF-β vs PBS | Up-regulated genes | CD45+ |
| BP | GO:1904407 | positive regulation of nitric oxide metabolic proc | 1.27E-03 | 2.01E-02 | Klf2/Sod2/Icam1 | 3 | aTGF-β vs PBS | Up-regulated genes | CD45+ |
| BP | GO:0002706 | regulation of lymphocyte mediated immunity | 1.30E-03 | 2.04E-02 | Rsad2/H2-D1/H2 | 5 | aTGF-β vs PBS | Up-regulated genes | CD45+ |
| BP | GO:0072608 | interleukin-10 secretion | 1.37E-03 | 2.11E-02 | Isg15/Cd274 | 2 | aTGF-β vs PBS | Up-regulated genes | CD45+ |
| BP | GO:1902043 | positive regulation of extrinsic apoptotic signalin | 1.37E-03 | 2.11E-02 | Atf3/Thbs1 | 2 | aTGF-β vs PBS | Up-regulated genes | CD45+ |
| BP | GO:0048247 | lymphocyte chemotaxis | 1.46E-03 | 2.19E-02 | Cxcl10/Ccl4/Ccl1 | 3 | aTGF-β vs PBS | Up-regulated genes | CD45+ |
| BP | GO:2000351 | regulation of endothelial cell apoptotic process | 1.46E-03 | 2.19E-02 | Ccl12/Icam1/Thbs | 3 | aTGF-β vs PBS | Up-regulated genes | CD45+ |
| BP | GO:0002548 | monocyte chemotaxis | 1.55E-03 | 2.29E-02 | Cxcl10/Ccl4/Ccl1 | 3 | aTGF-β vs PBS | Up-regulated genes | CD45+ |
| BP | GO:0070098 | chemokine-mediated signaling pathway | 1.55E-03 | 2.29E-02 | Cxcl10/Ccl4/Ccl1 | 3 | aTGF-β vs PBS | Up-regulated genes | CD45+ |
| BP | GO:0006968 | cellular defense response | 1.64E-03 | 2.37E-02 | B2m/Clec2d | 2 | aTGF-β vs PBS | Up-regulated genes | CD45+ |
| BP | GO:0060340 | positive regulation of type I interferon-mediated | 1.64E-03 | 2.37E-02 | Irf7/Zbp1 | 2 | aTGF-β vs PBS | Up-regulated genes | CD45+ |
| BP | GO:0072577 | endothelial cell apoptotic process | 1.66E-03 | 2.37E-02 | Ccl12/Icam1/Thbs | 3 | aTGF-β vs PBS | Up-regulated genes | CD45+ |
| BP | GO:0002688 | regulation of leukocyte chemotaxis | 1.69E-03 | 2.40E-02 | Cxcl10/Ccl4/Ccl1 | 4 | aTGF-β vs PBS | Up-regulated genes | CD45+ |
| BP | GO:0002253 | activation of immune response | 1.73E-03 | 2.43E-02 | Rsad2/Irf1/Irf7/Ac | 7 | aTGF-β vs PBS | Up-regulated genes | CD45+ |
| BP | GO:0002833 | positive regulation of response to biotic stimulus | 1.87E-03 | 2.60E-02 | Acod1/Cd274/Dd | 3 | aTGF-β vs PBS | Up-regulated genes | CD45+ |

| | | | | | | | | | |
|---|---|---|---|---|---|---|---|---|---|
| BP | GO:0042532 | negative regulation of tyrosine phosphorylation c | 1.94E-03 | 2.64E-02 | Parp14/Irf1 | 2 | aTGF-β vs PBS | Up-regulated genes | CD45+ |
| BP | GO:1900225 | regulation of NLRP3 inflammasome complex as | 1.94E-03 | 2.64E-02 | Gbp5/Trim30a | 2 | aTGF-β vs PBS | Up-regulated genes | CD45+ |
| BP | GO:0006986 | response to unfolded protein | 1.95E-03 | 2.64E-02 | Hspa1a/Atf3/Hspa | 4 | aTGF-β vs PBS | Up-regulated genes | CD45+ |
| BP | GO:0032103 | positive regulation of response to external stimul | 2.09E-03 | 2.80E-02 | Cxcl10/Gbp5/Ccl4 | 6 | aTGF-β vs PBS | Up-regulated genes | CD45+ |
| BP | GO:0050732 | negative regulation of peptidyl-tyrosine phospho | 2.11E-03 | 2.80E-02 | Parp14/Irf1/Sams | 3 | aTGF-β vs PBS | Up-regulated genes | CD45+ |
| BP | GO:0002697 | regulation of immune effector process | 2.14E-03 | 2.81E-02 | Rsad2/Irf1/H2-D1 | 7 | aTGF-β vs PBS | Up-regulated genes | CD45+ |
| BP | GO:0071621 | granulocyte chemotaxis | 2.17E-03 | 2.83E-02 | Cxcl10/Ccl4/Ccl1 | 4 | aTGF-β vs PBS | Up-regulated genes | CD45+ |
| BP | GO:0031348 | negative regulation of defense response | 2.19E-03 | 2.83E-02 | Parp14/Samhd1/ | 5 | aTGF-β vs PBS | Up-regulated genes | CD45+ |
| BP | GO:0044546 | NLRP3 inflammasome complex assembly | 2.25E-03 | 2.84E-02 | Gbp5/Trim30a | 2 | aTGF-β vs PBS | Up-regulated genes | CD45+ |
| BP | GO:0060330 | regulation of response to interferon-gamma | 2.25E-03 | 2.84E-02 | Parp14/Irgm1 | 2 | aTGF-β vs PBS | Up-regulated genes | CD45+ |
| BP | GO:0060334 | regulation of interferon-gamma-mediated signali | 2.25E-03 | 2.84E-02 | Parp14/Irgm1 | 2 | aTGF-β vs PBS | Up-regulated genes | CD45+ |
| BP | GO:0002685 | regulation of leukocyte migration | 2.35E-03 | 2.90E-02 | Cxcl10/Ccl4/Ccl1 | 5 | aTGF-β vs PBS | Up-regulated genes | CD45+ |
| BP | GO:0048002 | antigen processing and presentation of peptide a | 2.36E-03 | 2.90E-02 | H2-D1/H2-K1/B2r | 3 | aTGF-β vs PBS | Up-regulated genes | CD45+ |
| BP | GO:1903428 | positive regulation of reactive oxygen species bi | 2.36E-03 | 2.90E-02 | Klf2/Sod2/Icam1 | 3 | aTGF-β vs PBS | Up-regulated genes | CD45+ |
| BP | GO:0001909 | leukocyte mediated cytotoxicity | 2.40E-03 | 2.92E-02 | H2-D1/H2-K1/B2r | 4 | aTGF-β vs PBS | Up-regulated genes | CD45+ |
| BP | GO:0002708 | positive regulation of lymphocyte mediated immu | 2.64E-03 | 3.20E-02 | Rsad2/H2-D1/H2 | 4 | aTGF-β vs PBS | Up-regulated genes | CD45+ |
| BP | GO:0001913 | T cell mediated cytotoxicity | 2.77E-03 | 3.24E-02 | H2-D1/H2-K1/B2r | 3 | aTGF-β vs PBS | Up-regulated genes | CD45+ |
| BP | GO:1990868 | response to chemokine | 2.77E-03 | 3.24E-02 | Cxcl10/Ccl4/Ccl1 | 3 | aTGF-β vs PBS | Up-regulated genes | CD45+ |
| BP | GO:1990869 | cellular response to chemokine | 2.77E-03 | 3.24E-02 | Cxcl10/Ccl4/Ccl1 | 3 | aTGF-β vs PBS | Up-regulated genes | CD45+ |
| BP | GO:2000401 | regulation of lymphocyte migration | 2.77E-03 | 3.24E-02 | Cxcl10/Ccl4/Ccl1 | 3 | aTGF-β vs PBS | Up-regulated genes | CD45+ |
| BP | GO:0071396 | cellular response to lipid | 2.95E-03 | 3.43E-02 | Cxcl10/Gbp2/Cmp | 7 | aTGF-β vs PBS | Up-regulated genes | CD45+ |
| BP | GO:0002757 | immune response-activating signal transduction | 3.13E-03 | 3.61E-02 | Rsad2/Irf1/Irf7/Ac | 6 | aTGF-β vs PBS | Up-regulated genes | CD45+ |
| BP | GO:0002486 | antigen processing and presentation of endoger | 3.33E-03 | 3.78E-02 | H2-D1/H2-K1 | 2 | aTGF-β vs PBS | Up-regulated genes | CD45+ |
| BP | GO:2001185 | regulation of CD8-positive, alpha-beta T cell acti | 3.33E-03 | 3.78E-02 | Irf1/Cd274 | 2 | aTGF-β vs PBS | Up-regulated genes | CD45+ |
| BP | GO:0007050 | cell cycle arrest | 3.38E-03 | 3.81E-02 | Irf1/Hsp90ab1/Slf | 4 | aTGF-β vs PBS | Up-regulated genes | CD45+ |
| BP | GO:0045428 | regulation of nitric oxide biosynthetic process | 3.55E-03 | 3.97E-02 | Klf2/Sod2/Icam1 | 3 | aTGF-β vs PBS | Up-regulated genes | CD45+ |
| BP | GO:0071356 | cellular response to tumor necrosis factor | 3.69E-03 | 4.02E-02 | Irf1/Ccl4/Acod1/C | 4 | aTGF-β vs PBS | Up-regulated genes | CD45+ |
| BP | GO:0002476 | antigen processing and presentation of endoger | 3.73E-03 | 4.02E-02 | H2-D1/H2-K1 | 2 | aTGF-β vs PBS | Up-regulated genes | CD45+ |
| BP | GO:0002484 | antigen processing and presentation of endoger | 3.73E-03 | 4.02E-02 | H2-D1/H2-K1 | 2 | aTGF-β vs PBS | Up-regulated genes | CD45+ |
| BP | GO:0009200 | deoxyribonucleoside triphosphate metabolic pro | 3.73E-03 | 4.02E-02 | Cmpk2/Samhd1 | 2 | aTGF-β vs PBS | Up-regulated genes | CD45+ |
| BP | GO:0048245 | eosinophil chemotaxis | 3.73E-03 | 4.02E-02 | Ccl4/Ccl12 | 2 | aTGF-β vs PBS | Up-regulated genes | CD45+ |
| BP | GO:0002764 | immune response-regulating signaling pathway | 3.84E-03 | 4.11E-02 | Rsad2/Irf1/Irf7/Ac | 6 | aTGF-β vs PBS | Up-regulated genes | CD45+ |
| BP | GO:0035966 | response to topologically incorrect protein | 3.91E-03 | 4.15E-02 | Hspa1a/Atf3/Hspa | 4 | aTGF-β vs PBS | Up-regulated genes | CD45+ |
| BP | GO:0097530 | granulocyte migration | 4.02E-03 | 4.21E-02 | Cxcl10/Ccl4/Ccl1 | 4 | aTGF-β vs PBS | Up-regulated genes | CD45+ |
| BP | GO:0001912 | positive regulation of leukocyte mediated cytoto | 4.07E-03 | 4.21E-02 | H2-D1/H2-K1/B2r | 3 | aTGF-β vs PBS | Up-regulated genes | CD45+ |
| BP | GO:0008625 | extrinsic apoptotic signaling pathway via death c | 4.07E-03 | 4.21E-02 | Atf3/Icam1/Thbs1 | 3 | aTGF-β vs PBS | Up-regulated genes | CD45+ |
| BP | GO:0002703 | regulation of leukocyte mediated immunity | 4.09E-03 | 4.21E-02 | Rsad2/H2-D1/H2 | 5 | aTGF-β vs PBS | Up-regulated genes | CD45+ |
| BP | GO:0002726 | positive regulation of T cell cytokine production | 4.16E-03 | 4.21E-02 | Rsad2/B2m | 2 | aTGF-β vs PBS | Up-regulated genes | CD45+ |
| BP | GO:0002755 | MyD88-dependent toll-like receptor signaling pa | 4.16E-03 | 4.21E-02 | Irf1/Irf7 | 2 | aTGF-β vs PBS | Up-regulated genes | CD45+ |
| BP | GO:2000353 | positive regulation of endothelial cell apoptotic p | 4.16E-03 | 4.21E-02 | Ccl12/Thbs1 | 2 | aTGF-β vs PBS | Up-regulated genes | CD45+ |
| BP | GO:0032897 | negative regulation of viral transcription | 4.61E-03 | 4.63E-02 | Ifitm3/Ccl4 | 2 | aTGF-β vs PBS | Up-regulated genes | CD45+ |
| BP | GO:0006809 | nitric oxide biosynthetic process | 4.84E-03 | 4.83E-02 | Klf2/Sod2/Icam1 | 3 | aTGF-β vs PBS | Up-regulated genes | CD45+ |
| BP | GO:0072677 | eosinophil migration | 5.07E-03 | 5.00E-02 | Ccl4/Ccl12 | 2 | aTGF-β vs PBS | Up-regulated genes | CD45+ |
| BP | GO:1901673 | regulation of mitotic spindle assembly | 5.07E-03 | 5.00E-02 | Hspa1a/Hspa1b | 2 | aTGF-β vs PBS | Up-regulated genes | CD45+ |
| CC | GO:0009897 | external side of plasma membrane | 3.62E-08 | 5.04E-06 | Cxcl10/Cd274/Cc | 12 | aTGF-β vs PBS | Up-regulated genes | CD45+ |
| CC | GO:0020005 | symbiont-containing vacuole membrane | 1.76E-07 | 1.22E-05 | Gbp2b/Gbp2/Gbp | 4 | aTGF-β vs PBS | Up-regulated genes | CD45+ |
| CC | GO:0098552 | side of membrane | 3.22E-07 | 1.32E-05 | Cxcl10/Cd274/Cc | 13 | aTGF-β vs PBS | Up-regulated genes | CD45+ |
| CC | GO:0020003 | symbiont-containing vacuole | 3.78E-07 | 1.32E-05 | Gbp2b/Gbp2/Gbp | 4 | aTGF-β vs PBS | Up-regulated genes | CD45+ |
| CC | GO:0065010 | extracellular membrane-bounded organelle | 7.17E-07 | 1.89E-05 | Gbp2b/Gbp2/Gbp | 4 | aTGF-β vs PBS | Up-regulated genes | CD45+ |
| CC | GO:0030430 | host cell cytoplasm | 9.53E-07 | 1.89E-05 | Gbp2b/Gbp2/Gbp | 4 | aTGF-β vs PBS | Up-regulated genes | CD45+ |
| CC | GO:0033655 | host cell cytoplasm part | 9.53E-07 | 1.89E-05 | Gbp2b/Gbp2/Gbp | 4 | aTGF-β vs PBS | Up-regulated genes | CD45+ |
| CC | GO:0033646 | host intracellular part | 2.01E-06 | 3.10E-05 | Gbp2b/Gbp2/Gbp | 4 | aTGF-β vs PBS | Up-regulated genes | CD45+ |
| CC | GO:0043656 | intracellular region of host | 2.01E-06 | 3.10E-05 | Gbp2b/Gbp2/Gbp | 4 | aTGF-β vs PBS | Up-regulated genes | CD45+ |
| CC | GO:0043230 | extracellular organelle | 3.05E-06 | 3.89E-05 | Gbp2b/Gbp2/Cd2 | 6 | aTGF-β vs PBS | Up-regulated genes | CD45+ |
| CC | GO:0033643 | host cell part | 3.08E-06 | 3.89E-05 | Gbp2b/Gbp2/Gbp | 4 | aTGF-β vs PBS | Up-regulated genes | CD45+ |
| CC | GO:0018995 | host | 4.52E-06 | 4.83E-05 | Gbp2b/Gbp2/Gbp | 4 | aTGF-β vs PBS | Up-regulated genes | CD45+ |
| CC | GO:0043657 | host cell | 4.52E-06 | 4.83E-05 | Gbp2b/Gbp2/Gbp | 4 | aTGF-β vs PBS | Up-regulated genes | CD45+ |
| CC | GO:0044215 | other organism | 6.40E-06 | 5.56E-05 | Gbp2b/Gbp2/Gbp | 4 | aTGF-β vs PBS | Up-regulated genes | CD45+ |
| CC | GO:0044216 | other organism cell | 6.40E-06 | 5.56E-05 | Gbp2b/Gbp2/Gbp | 4 | aTGF-β vs PBS | Up-regulated genes | CD45+ |
| CC | GO:0044217 | other organism part | 6.40E-06 | 5.56E-05 | Gbp2b/Gbp2/Gbp | 4 | aTGF-β vs PBS | Up-regulated genes | CD45+ |
| CC | GO:0042612 | MHC class I protein complex | 1.86E-05 | 1.52E-04 | H2-D1/H2-K1/B2r | 3 | aTGF-β vs PBS | Up-regulated genes | CD45+ |
| CC | GO:0042824 | MHC class I peptide loading complex | 4.05E-05 | 3.13E-04 | H2-D1/H2-K1/B2r | 3 | aTGF-β vs PBS | Up-regulated genes | CD45+ |
| CC | GO:0042611 | MHC protein complex | 1.44E-04 | 1.06E-03 | H2-D1/H2-K1/B2r | 3 | aTGF-β vs PBS | Up-regulated genes | CD45+ |
| CC | GO:0008180 | COP9 signalosome | 6.21E-04 | 4.31E-03 | Hspa1a/Hsp90ab | 3 | aTGF-β vs PBS | Up-regulated genes | CD45+ |
| CC | GO:0005789 | endoplasmic reticulum membrane | 1.40E-03 | 9.25E-03 | Rsad2/Ifi47/Irgm1 | 8 | aTGF-β vs PBS | Up-regulated genes | CD45+ |
| CC | GO:0098827 | endoplasmic reticulum subcompartment | 1.68E-03 | 1.06E-02 | Rsad2/Ifi47/Irgm1 | 8 | aTGF-β vs PBS | Up-regulated genes | CD45+ |
| CC | GO:0042175 | nuclear outer membrane-endoplasmic reticulum | 1.80E-03 | 1.09E-02 | Rsad2/Ifi47/Irgm1 | 8 | aTGF-β vs PBS | Up-regulated genes | CD45+ |
| CC | GO:0030176 | integral component of endoplasmic reticulum me | 4.12E-03 | 2.38E-02 | H2-D1/H2-K1/B2r | 4 | aTGF-β vs PBS | Up-regulated genes | CD45+ |
| CC | GO:0031227 | intrinsic component of endoplasmic reticulum me | 4.94E-03 | 2.75E-02 | H2-D1/H2-K1/B2r | 4 | aTGF-β vs PBS | Up-regulated genes | CD45+ |
| CC | GO:0005797 | Golgi medial cisterna | 5.23E-03 | 2.79E-02 | H2-D1/H2-K1 | 2 | aTGF-β vs PBS | Up-regulated genes | CD45+ |
| CC | GO:0070971 | endoplasmic reticulum exit site | 7.26E-03 | 3.74E-02 | H2-D1/H2-K1 | 2 | aTGF-β vs PBS | Up-regulated genes | CD45+ |
| MF | GO:0003924 | GTPase activity | 4.16E-05 | 5.37E-03 | Gbp2b/Gbp2/Gbp | 8 | aTGF-β vs PBS | Up-regulated genes | CD45+ |
| MF | GO:0042379 | chemokine receptor binding | 6.80E-05 | 5.37E-03 | Cxcl10/Ccl4/Ccl1 | 4 | aTGF-β vs PBS | Up-regulated genes | CD45+ |
| MF | GO:0005525 | GTP binding | 1.78E-04 | 5.37E-03 | Gbp2b/Gbp2/Gbp | 8 | aTGF-β vs PBS | Up-regulated genes | CD45+ |
| MF | GO:0032550 | purine ribonucleoside binding | 2.02E-04 | 5.37E-03 | Gbp2b/Gbp2/Gbp | 8 | aTGF-β vs PBS | Up-regulated genes | CD45+ |
| MF | GO:0001883 | purine nucleoside binding | 2.14E-04 | 5.37E-03 | Gbp2b/Gbp2/Gbp | 8 | aTGF-β vs PBS | Up-regulated genes | CD45+ |
| MF | GO:0032549 | ribonucleoside binding | 2.23E-04 | 5.37E-03 | Gbp2b/Gbp2/Gbp | 8 | aTGF-β vs PBS | Up-regulated genes | CD45+ |
| MF | GO:0019001 | guanyl nucleotide binding | 2.47E-04 | 5.37E-03 | Gbp2b/Gbp2/Gbp | 8 | aTGF-β vs PBS | Up-regulated genes | CD45+ |
| MF | GO:0032561 | guanyl ribonucleotide binding | 2.47E-04 | 5.37E-03 | Gbp2b/Gbp2/Gbp | 8 | aTGF-β vs PBS | Up-regulated genes | CD45+ |
| MF | GO:0001882 | nucleoside binding | 2.67E-04 | 5.37E-03 | Gbp2b/Gbp2/Gbp | 8 | aTGF-β vs PBS | Up-regulated genes | CD45+ |
| MF | GO:0048020 | CCR chemokine receptor binding | 3.78E-04 | 6.83E-03 | Ccl4/Ccl12/Ccrl2 | 3 | aTGF-β vs PBS | Up-regulated genes | CD45+ |
| MF | GO:0003725 | double-stranded RNA binding | 4.97E-04 | 8.17E-03 | Hspa1a/Hsp90ab | 4 | aTGF-β vs PBS | Up-regulated genes | CD45+ |
| MF | GO:0008009 | chemokine activity | 6.62E-04 | 9.99E-03 | Cxcl10/Ccl4/Ccl1 | 3 | aTGF-β vs PBS | Up-regulated genes | CD45+ |
| MF | GO:0030881 | beta-2-microglobulin binding | 1.27E-03 | 1.53E-02 | H2-D1/H2-K1 | 2 | aTGF-β vs PBS | Up-regulated genes | CD45+ |
| MF | GO:0042610 | CD8 receptor binding | 1.27E-03 | 1.53E-02 | H2-D1/H2-K1 | 2 | aTGF-β vs PBS | Up-regulated genes | CD45+ |
| MF | GO:0046977 | TAP binding | 1.27E-03 | 1.53E-02 | H2-D1/H2-K1 | 2 | aTGF-β vs PBS | Up-regulated genes | CD45+ |
| MF | GO:0016763 | transferase activity, transferring pentosyl groups | 1.48E-03 | 1.67E-02 | Parp14/Upp1/Nar | 3 | aTGF-β vs PBS | Up-regulated genes | CD45+ |
| MF | GO:0042608 | T cell receptor binding | 2.08E-03 | 2.22E-02 | H2-D1/H2-K1 | 2 | aTGF-β vs PBS | Up-regulated genes | CD45+ |
| MF | GO:0001664 | G protein-coupled receptor binding | 3.24E-03 | 3.26E-02 | Cxcl10/Hspa1a/C | 5 | aTGF-β vs PBS | Up-regulated genes | CD45+ |

| | | | | | | | | | |
|---|---|---|---|---|---|---|---|---|---|
| MF | GO:0005125 | cytokine activity | 3.69E-03 | 3.52E-02 | Cxcl10/Ccl4/Ccl1 | 4 | aTGF-β vs PBS | Up-regulated genes | CD45+ |
| MF | GO:0044183 | protein binding involved in protein folding | 4.70E-03 | 4.05E-02 | Hspa1a/Hspa1b | 2 | aTGF-β vs PBS | Up-regulated genes | CD45+ |
| MF | GO:0051787 | misfolded protein binding | 4.70E-03 | 4.05E-02 | Hspa1a/Hspa1b | 2 | aTGF-β vs PBS | Up-regulated genes | CD45+ |
| MF | GO:0042605 | peptide antigen binding | 5.63E-03 | 4.63E-02 | H2-D1/H2-K1 | 2 | aTGF-β vs PBS | Up-regulated genes | CD45+ |
| BP | GO:0061077 | chaperone-mediated protein folding | 5.76E-06 | 2.60E-03 | Dnajb1/Hspa1b/H | 3 | aTGF-β vs PBS | Up-regulated genes | CD45- |
| BP | GO:0006457 | protein folding | 1.27E-04 | 2.21E-02 | Dnajb1/Hspa1b/H | 3 | aTGF-β vs PBS | Up-regulated genes | CD45- |
| BP | GO:0051085 | chaperone cofactor-dependent protein refolding | 2.01E-04 | 2.21E-02 | Dnajb1/Hspa1b | 2 | aTGF-β vs PBS | Up-regulated genes | CD45- |
| BP | GO:0051084 | 'de novo' posttranslational protein folding | 2.78E-04 | 2.21E-02 | Dnajb1/Hspa1b | 2 | aTGF-β vs PBS | Up-regulated genes | CD45- |
| BP | GO:0006458 | 'de novo' protein folding | 2.94E-04 | 2.21E-02 | Dnajb1/Hspa1b | 2 | aTGF-β vs PBS | Up-regulated genes | CD45- |
| BP | GO:0038084 | vascular endothelial growth factor signaling path | 2.94E-04 | 2.21E-02 | Dcn/Hspb1 | 2 | aTGF-β vs PBS | Up-regulated genes | CD45- |
| BP | GO:0043618 | regulation of transcription from RNA polymerase | 5.10E-04 | 2.88E-02 | Dnajb1/Klf2 | 2 | aTGF-β vs PBS | Up-regulated genes | CD45- |
| BP | GO:0043620 | regulation of DNA-templated transcription in res | 6.26E-04 | 2.88E-02 | Dnajb1/Klf2 | 2 | aTGF-β vs PBS | Up-regulated genes | CD45- |
| BP | GO:0035924 | cellular response to vascular endothelial growth | 6.51E-04 | 2.88E-02 | Dcn/Hspb1 | 2 | aTGF-β vs PBS | Up-regulated genes | CD45- |
| BP | GO:0045765 | regulation of angiogenesis | 6.74E-04 | 2.88E-02 | Dcn/Klf2/Hspb1 | 3 | aTGF-β vs PBS | Up-regulated genes | CD45- |
| BP | GO:0071277 | cellular response to calcium ion | 7.02E-04 | 2.88E-02 | Fos/Fosb | 2 | aTGF-β vs PBS | Up-regulated genes | CD45- |
| BP | GO:1901342 | regulation of vasculature development | 9.41E-04 | 3.42E-02 | Dcn/Klf2/Hspb1 | 3 | aTGF-β vs PBS | Up-regulated genes | CD45- |
| BP | GO:0032272 | negative regulation of protein polymerization | 9.85E-04 | 3.42E-02 | Tmsb10/Sgk1 | 2 | aTGF-β vs PBS | Up-regulated genes | CD45- |
| MF | GO:0044183 | protein binding involved in protein folding | 8.65E-05 | 5.53E-03 | Hspa1b/Hspb1 | 2 | aTGF-β vs PBS | Up-regulated genes | CD45- |
| BP | GO:1903706 | regulation of hemopoiesis | 3.63E-07 | 5.69E-04 | Ccr7/Hsp90aa1/F | 9 | aTGF-β vs PBS | Down-regulated gen | CD45+ |
| BP | GO:0032496 | response to lipopolysaccharide | 2.04E-06 | 1.29E-03 | Ccr7/Zfp36/Jun/Ju | 7 | aTGF-β vs PBS | Down-regulated gen | CD45+ |
| BP | GO:0001818 | negative regulation of cytokine production | 2.57E-06 | 1.29E-03 | Zfp36/Klf2/Srgn/C | 7 | aTGF-β vs PBS | Down-regulated gen | CD45+ |
| BP | GO:0002237 | response to molecule of bacterial origin | 3.30E-06 | 1.29E-03 | Ccr7/Zfp36/Jun/Ju | 7 | aTGF-β vs PBS | Down-regulated gen | CD45+ |
| BP | GO:0009612 | response to mechanical stimulus | 9.90E-06 | 2.63E-03 | Cxcr4/Jun/Junb/J | 5 | aTGF-β vs PBS | Down-regulated gen | CD45+ |
| BP | GO:0043903 | regulation of symbiosis, encompassing mutualis | 1.01E-05 | 2.63E-03 | Zfp36/Cxcr4/Jun/ | 6 | aTGF-β vs PBS | Down-regulated gen | CD45+ |
| BP | GO:0002456 | T cell mediated immunity | 1.23E-05 | 2.76E-03 | Ctsc/Il7r/Hspa8/S | 5 | aTGF-β vs PBS | Down-regulated gen | CD45+ |
| BP | GO:0002710 | negative regulation of T cell mediated immunity | 1.71E-05 | 3.34E-03 | Il7r/Smad7/Arg1 | 3 | aTGF-β vs PBS | Down-regulated gen | CD45+ |
| BP | GO:0044403 | symbiont process | 1.92E-05 | 3.34E-03 | Zfp36/Cxcr4/Jun/ | 7 | aTGF-β vs PBS | Down-regulated gen | CD45+ |
| BP | GO:0009617 | response to bacterium | 2.22E-05 | 3.47E-03 | Ccr7/Zfp36/Jun/Ju | 8 | aTGF-β vs PBS | Down-regulated gen | CD45+ |
| BP | GO:0044419 | interspecies interaction between organisms | 2.58E-05 | 3.65E-03 | Zfp36/Cxcr4/Jun/ | 7 | aTGF-β vs PBS | Down-regulated gen | CD45+ |
| BP | GO:0051702 | interaction with symbiont | 2.89E-05 | 3.65E-03 | Jun/Hspa8/Fn1/A | 4 | aTGF-β vs PBS | Down-regulated gen | CD45+ |
| BP | GO:0045580 | regulation of T cell differentiation | 3.03E-05 | 3.65E-03 | Ccr7/Hsp90aa1/Il | 5 | aTGF-β vs PBS | Down-regulated gen | CD45+ |
| BP | GO:0034612 | response to tumor necrosis factor | 3.60E-05 | 4.03E-03 | Zfp36/Pid1/Cxcl1 | 5 | aTGF-β vs PBS | Down-regulated gen | CD45+ |
| BP | GO:0048514 | blood vessel morphogenesis | 4.36E-05 | 4.44E-03 | Cxcr4/Btg1/Jun/K | 8 | aTGF-β vs PBS | Down-regulated gen | CD45+ |
| BP | GO:0002709 | regulation of T cell mediated immunity | 5.30E-05 | 4.44E-03 | Il7r/Hspa8/Smad7 | 4 | aTGF-β vs PBS | Down-regulated gen | CD45+ |
| BP | GO:1902107 | positive regulation of leukocyte differentiation | 5.66E-05 | 4.44E-03 | Ccr7/Hsp90aa1/J | 5 | aTGF-β vs PBS | Down-regulated gen | CD45+ |
| BP | GO:0050792 | regulation of viral process | 6.20E-05 | 4.44E-03 | Zfp36/Cxcr4/Jun/ | 5 | aTGF-β vs PBS | Down-regulated gen | CD45+ |
| BP | GO:0002460 | adaptive immune response based on somatic re | 6.25E-05 | 4.44E-03 | Ccr7/Ctsc/Il7r/Hsp | 6 | aTGF-β vs PBS | Down-regulated gen | CD45+ |
| BP | GO:0032103 | positive regulation of response to external stimul | 6.25E-05 | 4.44E-03 | Ccr7/Ctsc/Cxcr4/F | 6 | aTGF-β vs PBS | Down-regulated gen | CD45+ |
| BP | GO:1903037 | regulation of leukocyte cell-cell adhesion | 6.39E-05 | 4.44E-03 | Ccr7/Hsp90aa1/Il | 6 | aTGF-β vs PBS | Down-regulated gen | CD45+ |
| BP | GO:0002822 | regulation of adaptive immune response based | 6.58E-05 | 4.44E-03 | Ccr7/Il7r/Hspa8/S | 5 | aTGF-β vs PBS | Down-regulated gen | CD45+ |
| BP | GO:0032870 | cellular response to hormone stimulus | 6.71E-05 | 4.44E-03 | Zfp36/Pid1/Jun/S | 7 | aTGF-β vs PBS | Down-regulated gen | CD45+ |
| BP | GO:1902105 | regulation of leukocyte differentiation | 6.81E-05 | 4.44E-03 | Ccr7/Hsp90aa1/J | 6 | aTGF-β vs PBS | Down-regulated gen | CD45+ |
| BP | GO:0045619 | regulation of lymphocyte differentiation | 7.62E-05 | 4.78E-03 | Ccr7/Hsp90aa1/Il | 5 | aTGF-β vs PBS | Down-regulated gen | CD45+ |
| BP | GO:0050863 | regulation of T cell activation | 8.53E-05 | 4.97E-03 | Ccr7/Hsp90aa1/Il | 6 | aTGF-β vs PBS | Down-regulated gen | CD45+ |
| BP | GO:0045582 | positive regulation of T cell differentiation | 8.56E-05 | 4.97E-03 | Ccr7/Hsp90aa1/Il | 4 | aTGF-β vs PBS | Down-regulated gen | CD45+ |
| BP | GO:0002819 | regulation of adaptive immune response | 1.03E-04 | 5.79E-03 | Ccr7/Il7r/Hspa8/S | 5 | aTGF-β vs PBS | Down-regulated gen | CD45+ |
| BP | GO:0007159 | leukocyte cell-cell adhesion | 1.10E-04 | 5.94E-03 | Ccr7/Hsp90aa1/Il | 6 | aTGF-β vs PBS | Down-regulated gen | CD45+ |
| BP | GO:0042110 | T cell activation | 1.22E-04 | 6.40E-03 | Ccr7/Hsp90aa1/C | 7 | aTGF-β vs PBS | Down-regulated gen | CD45+ |
| BP | GO:0045621 | positive regulation of lymphocyte differentiation | 1.47E-04 | 7.25E-03 | Ccr7/Hsp90aa1/Il | 4 | aTGF-β vs PBS | Down-regulated gen | CD45+ |
| BP | GO:1903708 | positive regulation of hemopoiesis | 1.48E-04 | 7.25E-03 | Ccr7/Hsp90aa1/J | 5 | aTGF-β vs PBS | Down-regulated gen | CD45+ |
| BP | GO:0014070 | response to organic cyclic compound | 1.92E-04 | 8.46E-03 | Pim1/Zfp36/Jun/J | 7 | aTGF-β vs PBS | Down-regulated gen | CD45+ |
| BP | GO:0002823 | negative regulation of adaptive immune respons | 1.94E-04 | 8.46E-03 | Il7r/Smad7/Arg1 | 3 | aTGF-β vs PBS | Down-regulated gen | CD45+ |
| BP | GO:0051591 | response to cAMP | 1.94E-04 | 8.46E-03 | Jun/Junb/Jund | 3 | aTGF-β vs PBS | Down-regulated gen | CD45+ |
| BP | GO:0002694 | regulation of leukocyte activation | 1.94E-04 | 8.46E-03 | Ccr7/Ctsc/Hsp90a | 7 | aTGF-β vs PBS | Down-regulated gen | CD45+ |
| BP | GO:0002707 | negative regulation of lymphocyte mediated imm | 2.08E-04 | 8.76E-03 | Il7r/Smad7/Arg1 | 3 | aTGF-β vs PBS | Down-regulated gen | CD45+ |
| BP | GO:0001909 | leukocyte mediated cytotoxicity | 2.12E-04 | 8.76E-03 | Ctsc/Il7r/Hspa8/A | 4 | aTGF-β vs PBS | Down-regulated gen | CD45+ |
| BP | GO:0048247 | lymphocyte chemotaxis | 2.23E-04 | 8.97E-03 | Ccr7/Cxcl16/Ccl6 | 3 | aTGF-β vs PBS | Down-regulated gen | CD45+ |
| BP | GO:0071248 | cellular response to metal ion | 2.52E-04 | 9.46E-03 | Jun/Junb/Jund/Fn | 4 | aTGF-β vs PBS | Down-regulated gen | CD45+ |
| BP | GO:0022409 | positive regulation of cell-cell adhesion | 2.63E-04 | 9.46E-03 | Ccr7/Hsp90aa1/Il | 5 | aTGF-β vs PBS | Down-regulated gen | CD45+ |
| BP | GO:0022407 | regulation of cell-cell adhesion | 2.67E-04 | 9.46E-03 | Ccr7/Hsp90aa1/Il | 6 | aTGF-β vs PBS | Down-regulated gen | CD45+ |
| BP | GO:0043900 | regulation of multi-organism process | 2.67E-04 | 9.46E-03 | Zfp36/Cxcr4/Jun/ | 6 | aTGF-β vs PBS | Down-regulated gen | CD45+ |
| BP | GO:0002820 | negative regulation of adaptive immune respons | 2.72E-04 | 9.46E-03 | Il7r/Smad7/Arg1 | 3 | aTGF-β vs PBS | Down-regulated gen | CD45+ |
| BP | GO:0043370 | regulation of CD4-positive, alpha-beta T cell diff | 2.72E-04 | 9.46E-03 | Ccr7/Cd83/Smad | 3 | aTGF-β vs PBS | Down-regulated gen | CD45+ |
| BP | GO:0002521 | leukocyte differentiation | 2.83E-04 | 9.64E-03 | Ccr7/Hsp90aa1/J | 7 | aTGF-β vs PBS | Down-regulated gen | CD45+ |
| BP | GO:0050865 | regulation of cell activation | 2.94E-04 | 9.79E-03 | Ccr7/Ctsc/Hsp90a | 7 | aTGF-β vs PBS | Down-regulated gen | CD45+ |
| BP | GO:0010035 | response to inorganic substance | 3.07E-04 | 1.00E-02 | Ccr7/Jun/Junb/Ju | 6 | aTGF-β vs PBS | Down-regulated gen | CD45+ |
| BP | GO:0060394 | negative regulation of pathway-restricted SMAD | 3.16E-04 | 1.01E-02 | Pmepa1/Smad7 | 2 | aTGF-β vs PBS | Down-regulated gen | CD45+ |
| BP | GO:0030099 | myeloid cell differentiation | 3.27E-04 | 1.01E-02 | Ccr7/Zfp36/Jun/K | 6 | aTGF-β vs PBS | Down-regulated gen | CD45+ |
| BP | GO:0071356 | cellular response to tumor necrosis factor | 3.36E-04 | 1.01E-02 | Zfp36/Pid1/Ccl6/N | 4 | aTGF-β vs PBS | Down-regulated gen | CD45+ |
| BP | GO:0002704 | negative regulation of leukocyte mediated immu | 3.46E-04 | 1.01E-02 | Il7r/Smad7/Arg1 | 3 | aTGF-β vs PBS | Down-regulated gen | CD45+ |
| BP | GO:0002250 | adaptive immune response | 3.47E-04 | 1.01E-02 | Ccr7/Ctsc/Il7r/Hsp | 6 | aTGF-β vs PBS | Down-regulated gen | CD45+ |
| BP | GO:0045785 | positive regulation of cell adhesion | 3.47E-04 | 1.01E-02 | Ccr7/Hsp90aa1/Il | 6 | aTGF-β vs PBS | Down-regulated gen | CD45+ |
| BP | GO:2000107 | negative regulation of leukocyte apoptotic proce | 3.67E-04 | 1.03E-02 | Ccr7/Tsc22d3/Il7r | 3 | aTGF-β vs PBS | Down-regulated gen | CD45+ |
| BP | GO:0001933 | negative regulation of protein phosphorylation | 3.69E-04 | 1.03E-02 | Pmepa1/Pid1/Du | 6 | aTGF-β vs PBS | Down-regulated gen | CD45+ |
| BP | GO:0001913 | T cell mediated cytotoxicity | 4.33E-04 | 1.17E-02 | Ctsc/Il7r/Hspa8 | 3 | aTGF-β vs PBS | Down-regulated gen | CD45+ |
| BP | GO:0071277 | cellular response to calcium ion | 4.33E-04 | 1.17E-02 | Jun/Junb/Jund | 3 | aTGF-β vs PBS | Down-regulated gen | CD45+ |
| BP | GO:0060213 | positive regulation of nuclear-transcribed mRNA | 4.62E-04 | 1.17E-02 | Btg2/Zfp36 | 2 | aTGF-β vs PBS | Down-regulated gen | CD45+ |
| BP | GO:0002449 | lymphocyte mediated immunity | 4.63E-04 | 1.17E-02 | Ctsc/Il7r/Hspa8/S | 5 | aTGF-β vs PBS | Down-regulated gen | CD45+ |
| BP | GO:0071241 | cellular response to inorganic substance | 4.64E-04 | 1.17E-02 | Jun/Junb/Jund/Fn | 4 | aTGF-β vs PBS | Down-regulated gen | CD45+ |
| BP | GO:0051249 | regulation of lymphocyte activation | 4.71E-04 | 1.17E-02 | Ccr7/Hsp90aa1/Il | 6 | aTGF-β vs PBS | Down-regulated gen | CD45+ |
| BP | GO:0060326 | cell chemotaxis | 4.71E-04 | 1.17E-02 | Ccr7/Cxcr4/Cxcl1 | 5 | aTGF-β vs PBS | Down-regulated gen | CD45+ |
| BP | GO:0030217 | T cell differentiation | 4.80E-04 | 1.18E-02 | Ccr7/Hsp90aa1/Il | 5 | aTGF-β vs PBS | Down-regulated gen | CD45+ |
| BP | GO:0001906 | cell killing | 4.90E-04 | 1.18E-02 | Ctsc/Il7r/Hspa8/A | 4 | aTGF-β vs PBS | Down-regulated gen | CD45+ |
| BP | GO:0071774 | response to fibroblast growth factor | 5.33E-04 | 1.25E-02 | Zfp36/Ier2/Nr4a1 | 3 | aTGF-β vs PBS | Down-regulated gen | CD45+ |
| BP | GO:0016032 | viral process | 5.36E-04 | 1.25E-02 | Zfp36/Cxcr4/Jun/ | 5 | aTGF-β vs PBS | Down-regulated gen | CD45+ |
| BP | GO:0051851 | modification by host of symbiont morphology or | 5.88E-04 | 1.33E-02 | Jun/Hspa8/Arg1 | 3 | aTGF-β vs PBS | Down-regulated gen | CD45+ |
| BP | GO:2000514 | regulation of CD4-positive, alpha-beta T cell acti | 5.88E-04 | 1.33E-02 | Ccr7/Cd83/Smad | 3 | aTGF-β vs PBS | Down-regulated gen | CD45+ |
| BP | GO:0046637 | regulation of alpha-beta T cell differentiation | 6.16E-04 | 1.37E-02 | Ccr7/Cd83/Smad | 3 | aTGF-β vs PBS | Down-regulated gen | CD45+ |
| BP | GO:0042326 | negative regulation of phosphorylation | 6.19E-04 | 1.37E-02 | Pmepa1/Pid1/Du | 6 | aTGF-β vs PBS | Down-regulated gen | CD45+ |

| | | | | | | | | | |
|---|---|---|---|---|---|---|---|---|---|
| BP | GO:0060211 | regulation of nuclear-transcribed mRNA poly(A) | 6.34E-04 | 1.38E-02 | Btg2/Zfp36 | 2 | aTGF-β vs PBS | Down-regulated gen | CD45+ |
| BP | GO:0046683 | response to organophosphorus | 6.76E-04 | 1.45E-02 | Jun/Junb/Jund | 3 | aTGF-β vs PBS | Down-regulated gen | CD45+ |
| BP | GO:0043902 | positive regulation of multi-organism process | 7.08E-04 | 1.50E-02 | Jun/Vps37b/Hspa | 4 | aTGF-β vs PBS | Down-regulated gen | CD45+ |
| BP | GO:0002683 | negative regulation of immune system process | 7.62E-04 | 1.59E-02 | Tsc22d3/Zfp36/Il7 | 6 | aTGF-β vs PBS | Down-regulated gen | CD45+ |
| BP | GO:0001525 | angiogenesis | 7.81E-04 | 1.61E-02 | Cxcr4/Btg1/Jun/Kl | 6 | aTGF-β vs PBS | Down-regulated gen | CD45+ |
| BP | GO:0048872 | homeostasis of number of cells | 8.24E-04 | 1.68E-02 | Ccr7/Tsc22d3/Zfp | 5 | aTGF-β vs PBS | Down-regulated gen | CD45+ |
| BP | GO:0002706 | regulation of lymphocyte mediated immunity | 8.58E-04 | 1.72E-02 | Il7r/Hspa8/Smad7 | 4 | aTGF-β vs PBS | Down-regulated gen | CD45+ |
| BP | GO:0002696 | positive regulation of leukocyte activation | 9.08E-04 | 1.80E-02 | Ccr7/Ctsc/Hsp90a | 5 | aTGF-β vs PBS | Down-regulated gen | CD45+ |
| BP | GO:0010818 | T cell chemotaxis | 9.43E-04 | 1.80E-02 | Ccr7/Cxcl16 | 2 | aTGF-β vs PBS | Down-regulated gen | CD45+ |
| BP | GO:0035994 | response to muscle stretch | 9.43E-04 | 1.80E-02 | Jun/Nfkbia | 2 | aTGF-β vs PBS | Down-regulated gen | CD45+ |
| BP | GO:1900153 | positive regulation of nuclear-transcribed mRNA | 9.43E-04 | 1.80E-02 | Btg2/Zfp36 | 2 | aTGF-β vs PBS | Down-regulated gen | CD45+ |
| BP | GO:0006935 | chemotaxis | 9.75E-04 | 1.84E-02 | Ccr7/Cxcr4/Cxcl1 | 6 | aTGF-β vs PBS | Down-regulated gen | CD45+ |
| BP | GO:0042330 | taxis | 1.01E-03 | 1.88E-02 | Ccr7/Cxcr4/Cxcl1 | 6 | aTGF-β vs PBS | Down-regulated gen | CD45+ |
| BP | GO:0043367 | CD4-positive, alpha-beta T cell differentiation | 1.07E-03 | 1.90E-02 | Ccr7/Cd83/Smad | 3 | aTGF-β vs PBS | Down-regulated gen | CD45+ |
| BP | GO:0007517 | muscle organ development | 1.11E-03 | 1.90E-02 | Btg2/Tsc22d3/Pin | 5 | aTGF-β vs PBS | Down-regulated gen | CD45+ |
| BP | GO:0001938 | positive regulation of endothelial cell proliferatio | 1.12E-03 | 1.90E-02 | Jun/Nr4a1/Arg1 | 3 | aTGF-β vs PBS | Down-regulated gen | CD45+ |
| BP | GO:0014074 | response to purine-containing compound | 1.12E-03 | 1.90E-02 | Jun/Junb/Jund | 3 | aTGF-β vs PBS | Down-regulated gen | CD45+ |
| BP | GO:1903313 | positive regulation of mRNA metabolic process | 1.12E-03 | 1.90E-02 | Btg2/Zfp36/Hspa8 | 3 | aTGF-β vs PBS | Down-regulated gen | CD45+ |
| BP | GO:0070848 | response to growth factor | 1.12E-03 | 1.90E-02 | Pmepa1/Zfp36/Ju | 6 | aTGF-β vs PBS | Down-regulated gen | CD45+ |
| BP | GO:0050870 | positive regulation of T cell activation | 1.13E-03 | 1.90E-02 | Ccr7/Hsp90aa1/Il | 4 | aTGF-β vs PBS | Down-regulated gen | CD45+ |
| BP | GO:0050727 | regulation of inflammatory response | 1.13E-03 | 1.90E-02 | Ccr7/Ctsc/Zfp36/S | 5 | aTGF-β vs PBS | Down-regulated gen | CD45+ |
| BP | GO:0050867 | positive regulation of cell activation | 1.13E-03 | 1.90E-02 | Ccr7/Ctsc/Hsp90a | 5 | aTGF-β vs PBS | Down-regulated gen | CD45+ |
| BP | GO:0048524 | positive regulation of viral process | 1.16E-03 | 1.93E-02 | Jun/Vps37b/Hspa | 3 | aTGF-β vs PBS | Down-regulated gen | CD45+ |
| BP | GO:1900151 | regulation of nuclear-transcribed mRNA cataboli | 1.18E-03 | 1.95E-02 | Btg2/Zfp36 | 2 | aTGF-β vs PBS | Down-regulated gen | CD45+ |
| BP | GO:0032897 | negative regulation of viral transcription | 1.31E-03 | 2.14E-02 | Zfp36/Jun | 2 | aTGF-β vs PBS | Down-regulated gen | CD45+ |
| BP | GO:0051817 | modification of morphology or physiology of othe | 1.34E-03 | 2.17E-02 | Jun/Hspa8/Arg1 | 3 | aTGF-β vs PBS | Down-regulated gen | CD45+ |
| BP | GO:0001910 | regulation of leukocyte mediated cytotoxicity | 1.39E-03 | 2.22E-02 | Il7r/Hspa8/Arg1 | 3 | aTGF-β vs PBS | Down-regulated gen | CD45+ |
| BP | GO:0002443 | leukocyte mediated immunity | 1.44E-03 | 2.27E-02 | Ctsc/Il7r/Hspa8/S | 5 | aTGF-β vs PBS | Down-regulated gen | CD45+ |
| BP | GO:0002719 | negative regulation of cytokine production involv | 1.45E-03 | 2.27E-02 | Smad7/Arg1 | 2 | aTGF-β vs PBS | Down-regulated gen | CD45+ |
| BP | GO:0010563 | negative regulation of phosphorus metabolic pro | 1.50E-03 | 2.29E-02 | Pmepa1/Pid1/Dus | 6 | aTGF-β vs PBS | Down-regulated gen | CD45+ |
| BP | GO:0045936 | negative regulation of phosphate metabolic proc | 1.50E-03 | 2.29E-02 | Pmepa1/Pid1/Dus | 6 | aTGF-β vs PBS | Down-regulated gen | CD45+ |
| BP | GO:0014706 | striated muscle tissue development | 1.51E-03 | 2.29E-02 | Btg2/Tsc22d3/Pin | 5 | aTGF-β vs PBS | Down-regulated gen | CD45+ |
| BP | GO:1903039 | positive regulation of leukocyte cell-cell adhesio | 1.53E-03 | 2.31E-02 | Ccr7/Hsp90aa1/Il | 4 | aTGF-β vs PBS | Down-regulated gen | CD45+ |
| BP | GO:0072676 | lymphocyte migration | 1.60E-03 | 2.38E-02 | Ccr7/Cxcl16/Ccl6 | 3 | aTGF-β vs PBS | Down-regulated gen | CD45+ |
| BP | GO:0031331 | positive regulation of cellular catabolic process | 1.62E-03 | 2.39E-02 | Btg2/Ctsc/Hsp90a | 5 | aTGF-β vs PBS | Down-regulated gen | CD45+ |
| BP | GO:0000289 | nuclear-transcribed mRNA poly(A) tail shortenin | 1.74E-03 | 2.52E-02 | Btg2/Zfp36 | 2 | aTGF-β vs PBS | Down-regulated gen | CD45+ |
| BP | GO:0070233 | negative regulation of T cell apoptotic process | 1.74E-03 | 2.52E-02 | Tsc22d3/Il7r | 2 | aTGF-β vs PBS | Down-regulated gen | CD45+ |
| BP | GO:2000106 | regulation of leukocyte apoptotic process | 1.76E-03 | 2.53E-02 | Ccr7/Tsc22d3/Il7r | 3 | aTGF-β vs PBS | Down-regulated gen | CD45+ |
| BP | GO:0030098 | lymphocyte differentiation | 1.87E-03 | 2.67E-02 | Ccr7/Hsp90aa1/Il | 5 | aTGF-β vs PBS | Down-regulated gen | CD45+ |
| BP | GO:0060537 | muscle tissue development | 1.90E-03 | 2.68E-02 | Btg2/Tsc22d3/Pin | 5 | aTGF-β vs PBS | Down-regulated gen | CD45+ |
| BP | GO:0051592 | response to calcium ion | 1.94E-03 | 2.71E-02 | Jun/Junb/Jund | 3 | aTGF-β vs PBS | Down-regulated gen | CD45+ |
| BP | GO:0010038 | response to metal ion | 1.97E-03 | 2.72E-02 | Jun/Junb/Jund/Fn | 4 | aTGF-β vs PBS | Down-regulated gen | CD45+ |
| BP | GO:0035710 | CD4-positive, alpha-beta T cell activation | 2.00E-03 | 2.72E-02 | Ccr7/Cd83/Smad | 3 | aTGF-β vs PBS | Down-regulated gen | CD45+ |
| BP | GO:2000379 | positive regulation of reactive oxygen species m | 2.00E-03 | 2.72E-02 | Hsp90aa1/Pid1/K | 3 | aTGF-β vs PBS | Down-regulated gen | CD45+ |
| BP | GO:0002703 | regulation of leukocyte mediated immunity | 2.27E-03 | 3.06E-02 | Il7r/Hspa8/Smad7 | 4 | aTGF-β vs PBS | Down-regulated gen | CD45+ |
| BP | GO:0035821 | modification of morphology or physiology of othe | 2.32E-03 | 3.08E-02 | Jun/Hspa8/Arg1 | 3 | aTGF-β vs PBS | Down-regulated gen | CD45+ |
| BP | GO:0046634 | regulation of alpha-beta T cell activation | 2.32E-03 | 3.08E-02 | Ccr7/Cd83/Smad | 3 | aTGF-β vs PBS | Down-regulated gen | CD45+ |
| BP | GO:0048535 | lymph node development | 2.39E-03 | 3.15E-02 | Ccr7/Il7r | 2 | aTGF-β vs PBS | Down-regulated gen | CD45+ |
| BP | GO:0002697 | regulation of immune effector process | 2.48E-03 | 3.24E-02 | Ccr7/Il7r/Hspa8/S | 5 | aTGF-β vs PBS | Down-regulated gen | CD45+ |
| BP | GO:0031341 | regulation of cell killing | 2.53E-03 | 3.27E-02 | Il7r/Hspa8/Arg1 | 3 | aTGF-β vs PBS | Down-regulated gen | CD45+ |
| BP | GO:0043372 | positive regulation of CD4-positive, alpha-beta T | 2.57E-03 | 3.31E-02 | Ccr7/Cd83 | 2 | aTGF-β vs PBS | Down-regulated gen | CD45+ |
| BP | GO:0046632 | alpha-beta T cell differentiation | 2.60E-03 | 3.31E-02 | Ccr7/Cd83/Smad | 3 | aTGF-β vs PBS | Down-regulated gen | CD45+ |
| BP | GO:0042026 | protein refolding | 2.76E-03 | 3.49E-02 | Hsp90aa1/Hspa8 | 2 | aTGF-β vs PBS | Down-regulated gen | CD45+ |
| BP | GO:0001936 | regulation of endothelial cell proliferation | 2.90E-03 | 3.63E-02 | Jun/Nr4a1/Arg1 | 3 | aTGF-β vs PBS | Down-regulated gen | CD45+ |
| BP | GO:0002724 | regulation of T cell cytokine production | 2.95E-03 | 3.67E-02 | Smad7/Arg1 | 2 | aTGF-β vs PBS | Down-regulated gen | CD45+ |
| BP | GO:0071887 | leukocyte apoptotic process | 3.14E-03 | 3.86E-02 | Ccr7/Tsc22d3/Il7r | 3 | aTGF-β vs PBS | Down-regulated gen | CD45+ |
| BP | GO:0002701 | negative regulation of production of molecular m | 3.15E-03 | 3.86E-02 | Smad7/Arg1 | 2 | aTGF-β vs PBS | Down-regulated gen | CD45+ |
| BP | GO:0071396 | cellular response to lipid | 3.18E-03 | 3.86E-02 | Pim1/Zfp36/Pid1/ | 5 | aTGF-β vs PBS | Down-regulated gen | CD45+ |
| BP | GO:0006623 | protein targeting to vacuole | 3.35E-03 | 4.01E-02 | Vps37b/Hspa8 | 2 | aTGF-β vs PBS | Down-regulated gen | CD45+ |
| BP | GO:0060674 | placenta blood vessel development | 3.35E-03 | 4.01E-02 | Socs3/Junb | 2 | aTGF-β vs PBS | Down-regulated gen | CD45+ |
| BP | GO:0009896 | positive regulation of catabolic process | 3.40E-03 | 4.04E-02 | Btg2/Ctsc/Hsp90a | 5 | aTGF-β vs PBS | Down-regulated gen | CD45+ |
| BP | GO:0051251 | positive regulation of lymphocyte activation | 3.64E-03 | 4.29E-02 | Ccr7/Hsp90aa1/Il | 4 | aTGF-β vs PBS | Down-regulated gen | CD45+ |
| BP | GO:0046627 | negative regulation of insulin receptor signaling | 3.78E-03 | 4.39E-02 | Pid1/Socs3 | 2 | aTGF-β vs PBS | Down-regulated gen | CD45+ |
| BP | GO:2000516 | positive regulation of CD4-positive, alpha-beta T | 3.78E-03 | 4.39E-02 | Ccr7/Cd83 | 2 | aTGF-β vs PBS | Down-regulated gen | CD45+ |
| BP | GO:0045927 | positive regulation of growth | 3.86E-03 | 4.45E-02 | Pim1/Cxcr4/Cxcl1 | 4 | aTGF-β vs PBS | Down-regulated gen | CD45+ |
| BP | GO:0001935 | endothelial cell proliferation | 3.92E-03 | 4.49E-02 | Jun/Nr4a1/Arg1 | 3 | aTGF-β vs PBS | Down-regulated gen | CD45+ |
| BP | GO:0045622 | regulation of T-helper cell differentiation | 4.00E-03 | 4.49E-02 | Ccr7/Smad7 | 2 | aTGF-β vs PBS | Down-regulated gen | CD45+ |
| BP | GO:0070229 | negative regulation of lymphocyte apoptotic proc | 4.00E-03 | 4.49E-02 | Tsc22d3/Il7r | 2 | aTGF-β vs PBS | Down-regulated gen | CD45+ |
| BP | GO:0050921 | positive regulation of chemotaxis | 4.02E-03 | 4.49E-02 | Ccr7/Cxcr4/Fn1 | 3 | aTGF-β vs PBS | Down-regulated gen | CD45+ |
| BP | GO:0050729 | positive regulation of inflammatory response | 4.11E-03 | 4.57E-02 | Ccr7/Ctsc/Nfkbia | 3 | aTGF-β vs PBS | Down-regulated gen | CD45+ |
| BP | GO:0002698 | negative regulation of immune effector process | 4.21E-03 | 4.64E-02 | Il7r/Smad7/Arg1 | 3 | aTGF-β vs PBS | Down-regulated gen | CD45+ |
| BP | GO:0030855 | epithelial cell differentiation | 4.28E-03 | 4.69E-02 | Zfp36/Cxcr4/Btg1/ | 5 | aTGF-β vs PBS | Down-regulated gen | CD45+ |
| BP | GO:0034341 | response to interferon-gamma | 4.41E-03 | 4.79E-02 | Cxcl16/Ccl6/Arg1 | 3 | aTGF-β vs PBS | Down-regulated gen | CD45+ |
| BP | GO:0031647 | regulation of protein stability | 4.47E-03 | 4.83E-02 | Hsp90aa1/Pim1/H | 4 | aTGF-β vs PBS | Down-regulated gen | CD45+ |
| BP | GO:1903364 | positive regulation of cellular protein catabolic p | 4.51E-03 | 4.84E-02 | Ctsc/Hsp90aa1/S | 3 | aTGF-β vs PBS | Down-regulated gen | CD45+ |
| BP | GO:0001774 | microglial cell activation | 4.71E-03 | 4.88E-02 | Ctsc/Jun | 2 | aTGF-β vs PBS | Down-regulated gen | CD45+ |
| BP | GO:0001914 | regulation of T cell mediated cytotoxicity | 4.71E-03 | 4.88E-02 | Il7r/Hspa8 | 2 | aTGF-β vs PBS | Down-regulated gen | CD45+ |
| BP | GO:0002269 | leukocyte activation involved in inflammatory res | 4.71E-03 | 4.88E-02 | Ctsc/Jun | 2 | aTGF-β vs PBS | Down-regulated gen | CD45+ |
| BP | GO:0072666 | establishment of protein localization to vacuole | 4.71E-03 | 4.88E-02 | Vps37b/Hspa8 | 2 | aTGF-β vs PBS | Down-regulated gen | CD45+ |
| BP | GO:1900077 | negative regulation of cellular response to insuli | 4.71E-03 | 4.88E-02 | Pid1/Socs3 | 2 | aTGF-β vs PBS | Down-regulated gen | CD45+ |
| MF | GO:0019957 | C-C chemokine binding | 2.38E-05 | 4.38E-03 | Ccr7/Zfp36/Cxcr4 | 3 | aTGF-β vs PBS | Down-regulated gen | CD45+ |
| MF | GO:0019956 | chemokine binding | 5.18E-05 | 4.76E-03 | Ccr7/Zfp36/Cxcr4 | 3 | aTGF-β vs PBS | Down-regulated gen | CD45+ |
| MF | GO:0031625 | ubiquitin protein ligase binding | 9.73E-05 | 5.97E-03 | Hsp90aa1/Cxcr4/ | 6 | aTGF-β vs PBS | Down-regulated gen | CD45+ |
| MF | GO:0044389 | ubiquitin-like protein ligase binding | 1.37E-04 | 6.30E-03 | Hsp90aa1/Cxcr4/ | 6 | aTGF-β vs PBS | Down-regulated gen | CD45+ |
| MF | GO:0000981 | DNA-binding transcription factor activity, RNA po | 3.12E-04 | 1.15E-02 | Btg2/Jun/Klf2/Ier2 | 7 | aTGF-β vs PBS | Down-regulated gen | CD45+ |
| MF | GO:0001228 | DNA-binding transcription activator activity, RNA | 8.60E-04 | 2.24E-02 | Jun/Ier2/Junb/Jun | 5 | aTGF-β vs PBS | Down-regulated gen | CD45+ |
| MF | GO:0046332 | SMAD binding | 1.01E-03 | 2.24E-02 | Pmepa1/Jun/Sma | 3 | aTGF-β vs PBS | Down-regulated gen | CD45+ |
| MF | GO:0016493 | C-C chemokine receptor activity | 1.33E-03 | 2.24E-02 | Ccr7/Cxcr4 | 2 | aTGF-β vs PBS | Down-regulated gen | CD45+ |
| MF | GO:0004896 | cytokine receptor activity | 1.41E-03 | 2.24E-02 | Ccr7/Cxcr4/Il7r | 3 | aTGF-β vs PBS | Down-regulated gen | CD45+ |

| | | | | | | | | | |
|---|---|---|---|---|---|---|---|---|---|
| MF | GO:0001637 | G protein-coupled chemoattractant receptor acti | 1.46E-03 | 2.24E-02 | Ccr7/Cxcr4 | 2 | aTGF-β vs PBS | Down-regulated gen | CD45+ |
| MF | GO:0004950 | chemokine receptor activity | 1.46E-03 | 2.24E-02 | Ccr7/Cxcr4 | 2 | aTGF-β vs PBS | Down-regulated gen | CD45+ |
| MF | GO:0070412 | R-SMAD binding | 1.46E-03 | 2.24E-02 | Pmepa1/Jun | 2 | aTGF-β vs PBS | Down-regulated gen | CD45+ |
| MF | GO:0051087 | chaperone binding | 1.73E-03 | 2.45E-02 | Ctsc/Hspa8/Fn1 | 3 | aTGF-β vs PBS | Down-regulated gen | CD45+ |
| MF | GO:0016504 | peptidase activator activity | 2.79E-03 | 3.67E-02 | Ctsc/Fn1 | 2 | aTGF-β vs PBS | Down-regulated gen | CD45+ |
| MF | GO:0019955 | cytokine binding | 3.27E-03 | 3.90E-02 | Ccr7/Zfp36/Cxcr4 | 3 | aTGF-β vs PBS | Down-regulated gen | CD45+ |
| MF | GO:0097718 | disordered domain specific binding | 3.39E-03 | 3.90E-02 | Hsp90aa1/Fn1 | 2 | aTGF-β vs PBS | Down-regulated gen | CD45+ |
| MF | GO:0008009 | chemokine activity | 4.05E-03 | 4.38E-02 | Cxcl16/Ccl6 | 2 | aTGF-β vs PBS | Down-regulated gen | CD45+ |
| MF | GO:0031072 | heat shock protein binding | 4.79E-03 | 4.90E-02 | Zfp36/Hspa8/Nfkb | 3 | aTGF-β vs PBS | Down-regulated gen | CD45+ |
| MF | GO:0061134 | peptidase regulator activity | 5.11E-03 | 4.95E-02 | Cstb/Ctsc/Fn1 | 3 | aTGF-β vs PBS | Down-regulated gen | CD45+ |
| MF | GO:0031681 | G-protein beta-subunit binding | 5.10E-03 | 2.10E-02 | Gng11 | 1 | aTGF-β vs PBS | Down-regulated gen | CD45- |
| MF | GO:0017147 | Wnt-protein binding | 9.59E-03 | 2.10E-02 | Cthrc1 | 1 | aTGF-β vs PBS | Down-regulated gen | CD45- |
| MF | GO:0005109 | frizzled binding | 1.05E-02 | 2.10E-02 | Cthrc1 | 1 | aTGF-β vs PBS | Down-regulated gen | CD45- |
| BP | GO:0034341 | response to interferon-gamma | 2.81E-22 | 6.93E-19 | Irf1/H2-Eb1/H2-Aa | 22 | aPD-L1+aTGF-β vs PBS | Up-regulated genes | CD45+ |
| BP | GO:0009617 | response to bacterium | 1.74E-21 | 2.15E-18 | Cxcl10/Ly6a/Rpl3 | 34 | aPD-L1+aTGF-β vs PBS | Up-regulated genes | CD45+ |
| BP | GO:0071346 | cellular response to interferon-gamma | 1.57E-19 | 1.29E-16 | Irf1/H2-Ab1/Gbp2 | 19 | aPD-L1+aTGF-β vs PBS | Up-regulated genes | CD45+ |
| BP | GO:0098542 | defense response to other organism | 8.56E-18 | 5.29E-15 | Irf1/Cxcl10/Samh | 28 | aPD-L1+aTGF-β vs PBS | Up-regulated genes | CD45+ |
| BP | GO:0009615 | response to virus | 8.63E-14 | 4.26E-11 | Irf1/Cxcl10/Samh | 20 | aPD-L1+aTGF-β vs PBS | Up-regulated genes | CD45+ |
| BP | GO:0019221 | cytokine-mediated signaling pathway | 1.32E-12 | 5.43E-10 | Irf1/Cd74/Cxcl10/ | 20 | aPD-L1+aTGF-β vs PBS | Up-regulated genes | CD45+ |
| BP | GO:0071222 | cellular response to lipopolysaccharide | 2.32E-12 | 8.17E-10 | Cxcl10/Cd274/Gb | 16 | aPD-L1+aTGF-β vs PBS | Up-regulated genes | CD45+ |
| BP | GO:0071219 | cellular response to molecule of bacterial origin | 4.50E-12 | 1.39E-09 | Cxcl10/Cd274/Gb | 16 | aPD-L1+aTGF-β vs PBS | Up-regulated genes | CD45+ |
| BP | GO:0044419 | interspecies interaction between organisms | 5.75E-12 | 1.47E-09 | Cd74/Rsad2/Gbp | 21 | aPD-L1+aTGF-β vs PBS | Up-regulated genes | CD45+ |
| BP | GO:0051607 | defense response to virus | 5.95E-12 | 1.47E-09 | Irf1/Cxcl10/Samh | 17 | aPD-L1+aTGF-β vs PBS | Up-regulated genes | CD45+ |
| BP | GO:0002237 | response to molecule of bacterial origin | 1.51E-11 | 3.39E-09 | Cxcl10/Cd274/Gb | 18 | aPD-L1+aTGF-β vs PBS | Up-regulated genes | CD45+ |
| BP | GO:1990266 | neutrophil migration | 1.71E-11 | 3.53E-09 | Cd74/Cxcl10/Il1b | 13 | aPD-L1+aTGF-β vs PBS | Up-regulated genes | CD45+ |
| BP | GO:0071216 | cellular response to biotic stimulus | 2.76E-11 | 5.24E-09 | Cxcl10/Cd274/Gb | 16 | aPD-L1+aTGF-β vs PBS | Up-regulated genes | CD45+ |
| BP | GO:0030593 | neutrophil chemotaxis | 3.26E-11 | 5.76E-09 | Cd74/Cxcl10/Il1b | 12 | aPD-L1+aTGF-β vs PBS | Up-regulated genes | CD45+ |
| BP | GO:0032496 | response to lipopolysaccharide | 4.55E-11 | 7.50E-09 | Cxcl10/Cd274/Gb | 17 | aPD-L1+aTGF-β vs PBS | Up-regulated genes | CD45+ |
| BP | GO:0035456 | response to interferon-beta | 8.91E-11 | 1.38E-08 | Irf1/Gbp2b/Gbp2/I | 10 | aPD-L1+aTGF-β vs PBS | Up-regulated genes | CD45+ |
| BP | GO:0097530 | granulocyte migration | 2.22E-10 | 3.22E-08 | Cd74/Cxcl10/Il1b | 13 | aPD-L1+aTGF-β vs PBS | Up-regulated genes | CD45+ |
| BP | GO:0007159 | leukocyte cell-cell adhesion | 2.50E-10 | 3.43E-08 | Irf1/H2-Aa/Cd74/ | 18 | aPD-L1+aTGF-β vs PBS | Up-regulated genes | CD45+ |
| BP | GO:0042742 | defense response to bacterium | 3.02E-10 | 3.93E-08 | Rpl39/Gbp2b/Gbp | 14 | aPD-L1+aTGF-β vs PBS | Up-regulated genes | CD45+ |
| BP | GO:0071621 | granulocyte chemotaxis | 3.88E-10 | 4.75E-08 | Cd74/Cxcl10/Il1b | 12 | aPD-L1+aTGF-β vs PBS | Up-regulated genes | CD45+ |
| BP | GO:0035458 | cellular response to interferon-beta | 4.04E-10 | 4.75E-08 | Irf1/Gbp2b/Gbp2/I | 9 | aPD-L1+aTGF-β vs PBS | Up-regulated genes | CD45+ |
| BP | GO:1903037 | regulation of leukocyte cell-cell adhesion | 4.34E-10 | 4.87E-08 | Irf1/H2-Aa/Cd74/ | 17 | aPD-L1+aTGF-β vs PBS | Up-regulated genes | CD45+ |
| BP | GO:0044403 | symbiont process | 1.24E-09 | 1.33E-07 | Cd74/Rsad2/Gbp | 18 | aPD-L1+aTGF-β vs PBS | Up-regulated genes | CD45+ |
| BP | GO:0060326 | cell chemotaxis | 1.34E-09 | 1.37E-07 | Cd74/Cxcl10/Il1b | 16 | aPD-L1+aTGF-β vs PBS | Up-regulated genes | CD45+ |
| BP | GO:0002821 | positive regulation of adaptive immune response | 1.49E-09 | 1.47E-07 | Irf1/Cd74/H2-Ab1 | 12 | aPD-L1+aTGF-β vs PBS | Up-regulated genes | CD45+ |
| BP | GO:0097529 | myeloid leukocyte migration | 1.69E-09 | 1.61E-07 | Cd74/Cxcl10/Il1b | 14 | aPD-L1+aTGF-β vs PBS | Up-regulated genes | CD45+ |
| BP | GO:0002478 | antigen processing and presentation of exogen | 1.85E-09 | 1.69E-07 | H2-Eb1/H2-Aa/C | 7 | aPD-L1+aTGF-β vs PBS | Up-regulated genes | CD45+ |
| BP | GO:0071356 | cellular response to tumor necrosis factor | 2.18E-09 | 1.92E-07 | Irf1/Ccl4/Tnf/Nfe2 | 12 | aPD-L1+aTGF-β vs PBS | Up-regulated genes | CD45+ |
| BP | GO:0030595 | leukocyte chemotaxis | 2.79E-09 | 2.37E-07 | Cd74/Cxcl10/Il1b | 14 | aPD-L1+aTGF-β vs PBS | Up-regulated genes | CD45+ |
| BP | GO:0022407 | regulation of cell-cell adhesion | 3.41E-09 | 2.81E-07 | Irf1/H2-Aa/Cd74/ | 18 | aPD-L1+aTGF-β vs PBS | Up-regulated genes | CD45+ |
| BP | GO:0001819 | positive regulation of cytokine production | 4.02E-09 | 3.20E-07 | Irf1/Cd74/Rsad2/ | 19 | aPD-L1+aTGF-β vs PBS | Up-regulated genes | CD45+ |
| BP | GO:0019730 | antimicrobial humoral response | 7.32E-09 | 5.44E-07 | Cxcl10/Rpl39/Cx | 8 | aPD-L1+aTGF-β vs PBS | Up-regulated genes | CD45+ |
| BP | GO:0002250 | adaptive immune response | 7.35E-09 | 5.44E-07 | Irf1/Cd74/H2-Ab1 | 18 | aPD-L1+aTGF-β vs PBS | Up-regulated genes | CD45+ |
| BP | GO:0070555 | response to interleukin-1 | 7.49E-09 | 5.44E-07 | Irf1/Il1b/Ccl4/Ccl5 | 10 | aPD-L1+aTGF-β vs PBS | Up-regulated genes | CD45+ |
| BP | GO:0019884 | antigen processing and presentation of exogen | 8.00E-09 | 5.65E-07 | H2-Eb1/H2-Aa/C | 7 | aPD-L1+aTGF-β vs PBS | Up-regulated genes | CD45+ |
| BP | GO:0002548 | monocyte chemotaxis | 8.84E-09 | 5.94E-07 | Cxcl10/Ccl4/Ccl5 | 8 | aPD-L1+aTGF-β vs PBS | Up-regulated genes | CD45+ |
| BP | GO:0019882 | antigen processing and presentation | 9.43E-09 | 5.94E-07 | H2-Eb1/H2-Aa/C | 10 | aPD-L1+aTGF-β vs PBS | Up-regulated genes | CD45+ |
| BP | GO:0034612 | response to tumor necrosis factor | 9.50E-09 | 5.94E-07 | Irf1/Ccl4/Tnf/Nfe2 | 12 | aPD-L1+aTGF-β vs PBS | Up-regulated genes | CD45+ |
| BP | GO:1903706 | regulation of hemopoiesis | 9.54E-09 | 5.94E-07 | Irf1/H2-Aa/Cd74/I | 18 | aPD-L1+aTGF-β vs PBS | Up-regulated genes | CD45+ |
| BP | GO:0002824 | positive regulation of adaptive immune response | 9.86E-09 | 5.94E-07 | Irf1/H2-Ab1/Rsad | 11 | aPD-L1+aTGF-β vs PBS | Up-regulated genes | CD45+ |
| BP | GO:0006959 | humoral immune response | 9.86E-09 | 5.94E-07 | H2-Ab1/Cxcl10/R | 11 | aPD-L1+aTGF-β vs PBS | Up-regulated genes | CD45+ |
| BP | GO:0002819 | regulation of adaptive immune response | 1.25E-08 | 7.24E-07 | Irf1/Cd74/H2-Ab1 | 13 | aPD-L1+aTGF-β vs PBS | Up-regulated genes | CD45+ |
| BP | GO:0002683 | negative regulation of immune system process | 1.26E-08 | 7.24E-07 | Irf1/H2-Aa/Cd74/ | 19 | aPD-L1+aTGF-β vs PBS | Up-regulated genes | CD45+ |
| BP | GO:0071347 | cellular response to interleukin-1 | 1.51E-08 | 8.50E-07 | Irf1/Ccl4/Ccl5/Acc | 9 | aPD-L1+aTGF-β vs PBS | Up-regulated genes | CD45+ |
| BP | GO:0043901 | negative regulation of multi-organism process | 1.91E-08 | 1.05E-06 | Rsad2/Oasl1/Ccl | 12 | aPD-L1+aTGF-β vs PBS | Up-regulated genes | CD45+ |
| BP | GO:0042110 | T cell activation | 2.08E-08 | 1.12E-06 | Irf1/H2-Aa/Cd74/ | 19 | aPD-L1+aTGF-β vs PBS | Up-regulated genes | CD45+ |
| BP | GO:0050900 | leukocyte migration | 2.23E-08 | 1.16E-06 | Cd74/Cxcl10/Il1b | 16 | aPD-L1+aTGF-β vs PBS | Up-regulated genes | CD45+ |
| BP | GO:0043900 | regulation of multi-organism process | 2.25E-08 | 1.16E-06 | Cd74/Rsad2/Oasl | 17 | aPD-L1+aTGF-β vs PBS | Up-regulated genes | CD45+ |
| BP | GO:1903039 | positive regulation of leukocyte cell-cell adhesio | 2.42E-08 | 1.20E-06 | Irf1/H2-Aa/Cd74/ | 13 | aPD-L1+aTGF-β vs PBS | Up-regulated genes | CD45+ |
| BP | GO:0045088 | regulation of innate immune response | 2.43E-08 | 1.20E-06 | Irf1/Cd74/Samhd1 | 15 | aPD-L1+aTGF-β vs PBS | Up-regulated genes | CD45+ |
| BP | GO:0048002 | antigen processing and presentation of peptide | 2.92E-08 | 1.41E-06 | H2-Eb1/H2-Aa/C | 8 | aPD-L1+aTGF-β vs PBS | Up-regulated genes | CD45+ |
| BP | GO:0061844 | antimicrobial humoral immune response mediat | 3.29E-08 | 1.56E-06 | Cxcl10/Rpl39/Cx | 7 | aPD-L1+aTGF-β vs PBS | Up-regulated genes | CD45+ |
| BP | GO:0002697 | regulation of immune effector process | 4.25E-08 | 1.98E-06 | Irf1/Cd74/Rsad2/I | 17 | aPD-L1+aTGF-β vs PBS | Up-regulated genes | CD45+ |
| BP | GO:0050863 | regulation of T cell activation | 5.38E-08 | 2.46E-06 | Irf1/H2-Aa/Cd74/ | 15 | aPD-L1+aTGF-β vs PBS | Up-regulated genes | CD45+ |
| BP | GO:0050777 | negative regulation of immune response | 6.86E-08 | 3.08E-06 | Irf1/Samhd1/Arg2 | 11 | aPD-L1+aTGF-β vs PBS | Up-regulated genes | CD45+ |
| BP | GO:0050868 | negative regulation of T cell activation | 8.42E-08 | 3.71E-06 | Irf1/H2-Aa/Cd74/ | 10 | aPD-L1+aTGF-β vs PBS | Up-regulated genes | CD45+ |
| BP | GO:0051250 | negative regulation of lymphocyte activation | 8.60E-08 | 3.72E-06 | Irf1/H2-Aa/Cd74/ | 11 | aPD-L1+aTGF-β vs PBS | Up-regulated genes | CD45+ |
| BP | GO:0019886 | antigen processing and presentation of exogen | 8.86E-08 | 3.77E-06 | H2-Eb1/H2-Aa/C | 5 | aPD-L1+aTGF-β vs PBS | Up-regulated genes | CD45+ |
| BP | GO:0071396 | cellular response to lipid | 9.79E-08 | 4.10E-06 | Cxcl10/Dnaja1/C | 17 | aPD-L1+aTGF-β vs PBS | Up-regulated genes | CD45+ |
| BP | GO:0051249 | regulation of lymphocyte activation | 1.06E-07 | 4.35E-06 | Irf1/H2-Aa/Cd74/ | 17 | aPD-L1+aTGF-β vs PBS | Up-regulated genes | CD45+ |
| BP | GO:0022409 | positive regulation of cell-cell adhesion | 1.34E-07 | 5.42E-06 | Irf1/H2-Aa/Cd74/ | 13 | aPD-L1+aTGF-β vs PBS | Up-regulated genes | CD45+ |
| BP | GO:0002460 | adaptive immune response based on somatic re | 1.80E-07 | 7.19E-06 | Irf1/Cd74/H2-Ab1 | 14 | aPD-L1+aTGF-β vs PBS | Up-regulated genes | CD45+ |
| BP | GO:1903038 | negative regulation of leukocyte cell-cell adhesi | 1.94E-07 | 7.60E-06 | Irf1/H2-Aa/Cd74/ | 10 | aPD-L1+aTGF-β vs PBS | Up-regulated genes | CD45+ |
| BP | GO:0070098 | chemokine-mediated signaling pathway | 2.05E-07 | 7.91E-06 | Cxcl10/Cxcl9/Ccl | 7 | aPD-L1+aTGF-β vs PBS | Up-regulated genes | CD45+ |
| BP | GO:0043903 | regulation of symbiosis, encompassing mutualis | 2.32E-07 | 8.83E-06 | Cd74/Rsad2/Oasl | 12 | aPD-L1+aTGF-β vs PBS | Up-regulated genes | CD45+ |
| BP | GO:1903708 | positive regulation of hemopoiesis | 2.61E-07 | 9.75E-06 | Irf1/H2-Aa/Cd74/I | 12 | aPD-L1+aTGF-β vs PBS | Up-regulated genes | CD45+ |
| BP | GO:0071674 | mononuclear cell migration | 2.83E-07 | 1.04E-05 | Cxcl10/Ccl4/Ccl5 | 8 | aPD-L1+aTGF-β vs PBS | Up-regulated genes | CD45+ |
| BP | GO:0050792 | regulation of viral process | 3.17E-07 | 1.15E-05 | Cd74/Rsad2/Oasl | 11 | aPD-L1+aTGF-β vs PBS | Up-regulated genes | CD45+ |
| BP | GO:0022408 | negative regulation of cell-cell adhesion | 3.38E-07 | 1.21E-05 | Irf1/H2-Aa/Cd74/ | 11 | aPD-L1+aTGF-β vs PBS | Up-regulated genes | CD45+ |
| BP | GO:0002822 | regulation of adaptive immune response based | 3.60E-07 | 1.27E-05 | Irf1/H2-Ab1/Rsad | 11 | aPD-L1+aTGF-β vs PBS | Up-regulated genes | CD45+ |
| BP | GO:0002695 | negative regulation of leukocyte activation | 4.08E-07 | 1.42E-05 | Irf1/H2-Aa/Cd74/ | 11 | aPD-L1+aTGF-β vs PBS | Up-regulated genes | CD45+ |
| BP | GO:0031349 | positive regulation of defense response | 4.69E-07 | 1.61E-05 | Irf1/Cd74/Rsad2/I | 15 | aPD-L1+aTGF-β vs PBS | Up-regulated genes | CD45+ |
| BP | GO:0002495 | antigen processing and presentation of peptide | 5.67E-07 | 1.92E-05 | H2-Eb1/H2-Aa/C | 5 | aPD-L1+aTGF-β vs PBS | Up-regulated genes | CD45+ |
| BP | GO:0050870 | positive regulation of T cell activation | 7.02E-07 | 2.34E-05 | Irf1/H2-Aa/Cd74/ | 11 | aPD-L1+aTGF-β vs PBS | Up-regulated genes | CD45+ |
| BP | GO:0002711 | positive regulation of T cell mediated immunity | 7.45E-07 | 2.45E-05 | Rsad2/Il1b/H2-K1 | 7 | aPD-L1+aTGF-β vs PBS | Up-regulated genes | CD45+ |
| BP | GO:0002504 | antigen processing and presentation of peptide | 7.64E-07 | 2.48E-05 | H2-Eb1/H2-Aa/C | 5 | aPD-L1+aTGF-β vs PBS | Up-regulated genes | CD45+ |

| | | | | | | | | | |
|---|---|---|---|---|---|---|---|---|---|
| BP | GO:0016032 | viral process | 7.91E-07 | 2.54E-05 | Cd74/Rsad2/Oasl | 13 | aPD-L1+aTGF-β vs PBS | Up-regulated genes | CD45+ |
| BP | GO:0002699 | positive regulation of immune effector process | 8.44E-07 | 2.62E-05 | Irf1/Cd74/Rsad2/I | 12 | aPD-L1+aTGF-β vs PBS | Up-regulated genes | CD45+ |
| BP | GO:1990868 | response to chemokine | 8.47E-07 | 2.62E-05 | Cxcl10/Cxcl9/Ccl4 | 7 | aPD-L1+aTGF-β vs PBS | Up-regulated genes | CD45+ |
| BP | GO:1990869 | cellular response to chemokine | 8.47E-07 | 2.62E-05 | Cxcl10/Cxcl9/Ccl4 | 7 | aPD-L1+aTGF-β vs PBS | Up-regulated genes | CD45+ |
| BP | GO:0050866 | negative regulation of cell activation | 9.88E-07 | 3.01E-05 | Irf1/H2-Aa/Cd74/H | 11 | aPD-L1+aTGF-β vs PBS | Up-regulated genes | CD45+ |
| BP | GO:0060333 | interferon-gamma-mediated signaling pathway | 1.01E-06 | 3.05E-05 | Irf1/Irgm1/Socs1/F | 5 | aPD-L1+aTGF-β vs PBS | Up-regulated genes | CD45+ |
| BP | GO:0032103 | positive regulation of response to external stimul | 1.18E-06 | 3.50E-05 | Cd74/Cxcl10/Il1b/ | 13 | aPD-L1+aTGF-β vs PBS | Up-regulated genes | CD45+ |
| BP | GO:0048525 | negative regulation of viral process | 1.31E-06 | 3.85E-05 | Rsad2/Oasl1/Ccl4 | 8 | aPD-L1+aTGF-β vs PBS | Up-regulated genes | CD45+ |
| BP | GO:1902105 | regulation of leukocyte differentiation | 1.40E-06 | 4.06E-05 | Irf1/H2-Aa/Cd74/Il | 13 | aPD-L1+aTGF-β vs PBS | Up-regulated genes | CD45+ |
| BP | GO:0002694 | regulation of leukocyte activation | 1.67E-06 | 4.78E-05 | Irf1/H2-Aa/Cd74/H | 17 | aPD-L1+aTGF-β vs PBS | Up-regulated genes | CD45+ |
| BP | GO:1902107 | positive regulation of leukocyte differentiation | 2.24E-06 | 6.36E-05 | Irf1/H2-Aa/Cd74/Il | 10 | aPD-L1+aTGF-β vs PBS | Up-regulated genes | CD45+ |
| BP | GO:0001660 | fever generation | 2.48E-06 | 6.96E-05 | Il1b/Tnf/Ccl5/Il1rn | 4 | aPD-L1+aTGF-β vs PBS | Up-regulated genes | CD45+ |
| BP | GO:0051251 | positive regulation of lymphocyte activation | 2.60E-06 | 7.23E-05 | Irf1/H2-Aa/Cd74/H | 12 | aPD-L1+aTGF-β vs PBS | Up-regulated genes | CD45+ |
| BP | GO:1903900 | regulation of viral life cycle | 2.98E-06 | 8.17E-05 | Cd74/Rsad2/Oasl | 9 | aPD-L1+aTGF-β vs PBS | Up-regulated genes | CD45+ |
| BP | GO:0050830 | defense response to Gram-positive bacterium | 3.30E-06 | 8.96E-05 | Rpl39/Gbp2b/Gbp | 7 | aPD-L1+aTGF-β vs PBS | Up-regulated genes | CD45+ |
| BP | GO:0030098 | lymphocyte differentiation | 3.35E-06 | 9.01E-05 | Irf1/H2-Aa/Cd74/H | 14 | aPD-L1+aTGF-β vs PBS | Up-regulated genes | CD45+ |
| BP | GO:0048247 | lymphocyte chemotaxis | 3.52E-06 | 9.26E-05 | Cxcl10/Ccl4/Ccl5/ | 6 | aPD-L1+aTGF-β vs PBS | Up-regulated genes | CD45+ |
| BP | GO:0002449 | lymphocyte mediated immunity | 3.53E-06 | 9.26E-05 | Cd74/H2-Ab1/Rsa | 12 | aPD-L1+aTGF-β vs PBS | Up-regulated genes | CD45+ |
| BP | GO:0006935 | chemotaxis | 3.56E-06 | 9.26E-05 | Cd74/Cxcl10/Il1b/ | 16 | aPD-L1+aTGF-β vs PBS | Up-regulated genes | CD45+ |
| BP | GO:0002521 | leukocyte differentiation | 3.81E-06 | 9.77E-05 | Irf1/H2-Aa/Cd74/H | 17 | aPD-L1+aTGF-β vs PBS | Up-regulated genes | CD45+ |
| BP | GO:0030217 | T cell differentiation | 3.84E-06 | 9.77E-05 | Irf1/H2-Aa/Cd74/F | 12 | aPD-L1+aTGF-β vs PBS | Up-regulated genes | CD45+ |
| BP | GO:0042330 | taxis | 3.89E-06 | 9.81E-05 | Cd74/Cxcl10/Il1b/ | 16 | aPD-L1+aTGF-β vs PBS | Up-regulated genes | CD45+ |
| BP | GO:0045619 | regulation of lymphocyte differentiation | 3.93E-06 | 9.81E-05 | Irf1/H2-Aa/Cd74/Il | 10 | aPD-L1+aTGF-β vs PBS | Up-regulated genes | CD45+ |
| BP | GO:0050865 | regulation of cell activation | 4.14E-06 | 1.02E-04 | Irf1/H2-Aa/Cd74/H | 17 | aPD-L1+aTGF-β vs PBS | Up-regulated genes | CD45+ |
| BP | GO:0050727 | regulation of inflammatory response | 4.94E-06 | 1.21E-04 | Il1b/Tnf/Nfkbia/Nl | 13 | aPD-L1+aTGF-β vs PBS | Up-regulated genes | CD45+ |
| BP | GO:0042532 | negative regulation of tyrosine phosphorylation c | 5.29E-06 | 1.27E-04 | Irf1/Socs1/Parp14 | 4 | aPD-L1+aTGF-β vs PBS | Up-regulated genes | CD45+ |
| BP | GO:0044406 | adhesion of symbiont to host | 5.29E-06 | 1.27E-04 | Gbp2b/Gbp2/Ican | 4 | aPD-L1+aTGF-β vs PBS | Up-regulated genes | CD45+ |
| BP | GO:0031663 | lipopolysaccharide-mediated signaling pathway | 5.93E-06 | 1.40E-04 | Il1b/Tnf/Nfkbia/Cc | 6 | aPD-L1+aTGF-β vs PBS | Up-regulated genes | CD45+ |
| BP | GO:0045580 | regulation of T cell differentiation | 5.97E-06 | 1.40E-04 | Irf1/H2-Aa/Cd74/Il | 9 | aPD-L1+aTGF-β vs PBS | Up-regulated genes | CD45+ |
| BP | GO:0019058 | viral life cycle | 6.31E-06 | 1.47E-04 | Cd74/Rsad2/Oasl | 10 | aPD-L1+aTGF-β vs PBS | Up-regulated genes | CD45+ |
| BP | GO:0045089 | positive regulation of innate immune response | 6.55E-06 | 1.51E-04 | Irf1/Cd74/Rsad2/I | 11 | aPD-L1+aTGF-β vs PBS | Up-regulated genes | CD45+ |
| BP | GO:0007249 | I-kappaB kinase/NF-kappaB signaling | 7.35E-06 | 1.68E-04 | Irf1/Cd74/Il1b/Tnf/ | 10 | aPD-L1+aTGF-β vs PBS | Up-regulated genes | CD45+ |
| BP | GO:0045785 | positive regulation of cell adhesion | 7.49E-06 | 1.70E-04 | Irf1/H2-Aa/Cd74/H | 14 | aPD-L1+aTGF-β vs PBS | Up-regulated genes | CD45+ |
| BP | GO:0002181 | cytoplasmic translation | 7.73E-06 | 1.74E-04 | Rpl39/Rpl35a/Rp | 7 | aPD-L1+aTGF-β vs PBS | Up-regulated genes | CD45+ |
| BP | GO:0031348 | negative regulation of defense response | 8.11E-06 | 1.81E-04 | Samhd1/Arg2/Nlr | 10 | aPD-L1+aTGF-β vs PBS | Up-regulated genes | CD45+ |
| BP | GO:0002709 | regulation of T cell mediated immunity | 8.44E-06 | 1.86E-04 | Rsad2/Il1b/H2-K1 | 7 | aPD-L1+aTGF-β vs PBS | Up-regulated genes | CD45+ |
| BP | GO:0045071 | negative regulation of viral genome replication | 8.50E-06 | 1.86E-04 | Rsad2/Oasl1/Tnf/ | 6 | aPD-L1+aTGF-β vs PBS | Up-regulated genes | CD45+ |
| BP | GO:0046631 | alpha-beta T cell activation | 9.50E-06 | 2.06E-04 | Irf1/H2-Ab1/Rsad | 9 | aPD-L1+aTGF-β vs PBS | Up-regulated genes | CD45+ |
| BP | GO:0002828 | regulation of type 2 immune response | 1.02E-05 | 2.19E-04 | Irf1/Cd74/Rsad2/ | 5 | aPD-L1+aTGF-β vs PBS | Up-regulated genes | CD45+ |
| BP | GO:0045069 | regulation of viral genome replication | 1.18E-05 | 2.52E-04 | Rsad2/Oasl1/Tnf/ | 7 | aPD-L1+aTGF-β vs PBS | Up-regulated genes | CD45+ |
| BP | GO:0002456 | T cell mediated immunity | 1.22E-05 | 2.56E-04 | Rsad2/Il1b/H2-K1 | 8 | aPD-L1+aTGF-β vs PBS | Up-regulated genes | CD45+ |
| BP | GO:0002708 | positive regulation of lymphocyte mediated immu | 1.22E-05 | 2.56E-04 | Rsad2/Il1b/H2-K1 | 8 | aPD-L1+aTGF-β vs PBS | Up-regulated genes | CD45+ |
| BP | GO:0031649 | heat generation | 1.32E-05 | 2.74E-04 | Il1b/Tnf/Ccl5/Il1rn | 4 | aPD-L1+aTGF-β vs PBS | Up-regulated genes | CD45+ |
| BP | GO:0002262 | myeloid cell homeostasis | 1.47E-05 | 3.03E-04 | Pde4b/B2m/Slc7a | 9 | aPD-L1+aTGF-β vs PBS | Up-regulated genes | CD45+ |
| BP | GO:0007162 | negative regulation of cell adhesion | 1.58E-05 | 3.22E-04 | Irf1/H2-Aa/Cd74/H | 11 | aPD-L1+aTGF-β vs PBS | Up-regulated genes | CD45+ |
| BP | GO:0002696 | positive regulation of leukocyte activation | 1.60E-05 | 3.23E-04 | Irf1/H2-Aa/Cd74/H | 12 | aPD-L1+aTGF-β vs PBS | Up-regulated genes | CD45+ |
| BP | GO:0042129 | regulation of T cell proliferation | 1.72E-05 | 3.45E-04 | Irf1/H2-Aa/H2-Ab | 9 | aPD-L1+aTGF-β vs PBS | Up-regulated genes | CD45+ |
| BP | GO:0072676 | lymphocyte migration | 1.76E-05 | 3.50E-04 | Cxcl10/Ccl4/Icam | 7 | aPD-L1+aTGF-β vs PBS | Up-regulated genes | CD45+ |
| BP | GO:0045582 | positive regulation of T cell differentiation | 1.90E-05 | 3.72E-04 | Irf1/H2-Aa/Cd74/Il | 7 | aPD-L1+aTGF-β vs PBS | Up-regulated genes | CD45+ |
| BP | GO:0050729 | positive regulation of inflammatory response | 1.90E-05 | 3.72E-04 | Il1b/Tnf/Nfkbia/Cc | 8 | aPD-L1+aTGF-β vs PBS | Up-regulated genes | CD45+ |
| BP | GO:0034340 | response to type I interferon | 2.18E-05 | 4.21E-04 | Samhd1/Zbp1/Sta | 5 | aPD-L1+aTGF-β vs PBS | Up-regulated genes | CD45+ |
| BP | GO:0048245 | eosinophil chemotaxis | 2.18E-05 | 4.21E-04 | Ccl4/Ccl5/Ccl2/Cc | 4 | aPD-L1+aTGF-β vs PBS | Up-regulated genes | CD45+ |
| BP | GO:0050670 | regulation of lymphocyte proliferation | 2.33E-05 | 4.46E-04 | Irf1/H2-Aa/Cd74/H | 10 | aPD-L1+aTGF-β vs PBS | Up-regulated genes | CD45+ |
| BP | GO:0042509 | regulation of tyrosine phosphorylation of STAT p | 2.43E-05 | 4.57E-04 | Irf1/Socs1/Tnf/Ccl | 6 | aPD-L1+aTGF-β vs PBS | Up-regulated genes | CD45+ |
| BP | GO:2000514 | regulation of CD4-positive, alpha-beta T cell acti | 2.43E-05 | 4.57E-04 | Irf1/Cd274/Arg2/S | 6 | aPD-L1+aTGF-β vs PBS | Up-regulated genes | CD45+ |
| BP | GO:0042092 | type 2 immune response | 2.50E-05 | 4.67E-04 | Irf1/Cd74/Rsad2/ | 5 | aPD-L1+aTGF-β vs PBS | Up-regulated genes | CD45+ |
| BP | GO:0032944 | regulation of mononuclear cell proliferation | 2.53E-05 | 4.70E-04 | Irf1/H2-Aa/Cd74/H | 10 | aPD-L1+aTGF-β vs PBS | Up-regulated genes | CD45+ |
| BP | GO:0050867 | positive regulation of cell activation | 2.58E-05 | 4.76E-04 | Irf1/H2-Aa/Cd74/H | 12 | aPD-L1+aTGF-β vs PBS | Up-regulated genes | CD45+ |
| BP | GO:0042130 | negative regulation of T cell proliferation | 2.66E-05 | 4.88E-04 | H2-Aa/H2-Ab1/Cc | 6 | aPD-L1+aTGF-β vs PBS | Up-regulated genes | CD45+ |
| BP | GO:0002726 | positive regulation of T cell cytokine production | 2.74E-05 | 4.98E-04 | Rsad2/Il1b/Nlrp3/ | 4 | aPD-L1+aTGF-β vs PBS | Up-regulated genes | CD45+ |
| BP | GO:0035690 | cellular response to drug | 2.80E-05 | 5.04E-04 | Il1b/Ccl4/Tnf/Nfe2 | 11 | aPD-L1+aTGF-β vs PBS | Up-regulated genes | CD45+ |
| BP | GO:0001959 | regulation of cytokine-mediated signaling pathw | 2.93E-05 | 5.17E-04 | Cd74/Samhd1/Irg | 7 | aPD-L1+aTGF-β vs PBS | Up-regulated genes | CD45+ |
| BP | GO:0035710 | CD4-positive, alpha-beta T cell activation | 2.93E-05 | 5.17E-04 | Irf1/Rsad2/Cd274 | 7 | aPD-L1+aTGF-β vs PBS | Up-regulated genes | CD45+ |
| BP | GO:2000379 | positive regulation of reactive oxygen species m | 2.93E-05 | 5.17E-04 | Il1b/Icam1/Tnf/Nfe | 7 | aPD-L1+aTGF-β vs PBS | Up-regulated genes | CD45+ |
| BP | GO:0014070 | response to organic cyclic compound | 3.05E-05 | 5.34E-04 | Dnaja1/Il1b/Socs | 15 | aPD-L1+aTGF-β vs PBS | Up-regulated genes | CD45+ |
| BP | GO:0007260 | tyrosine phosphorylation of STAT protein | 3.50E-05 | 6.08E-04 | Irf1/Socs1/Tnf/Ccl | 6 | aPD-L1+aTGF-β vs PBS | Up-regulated genes | CD45+ |
| BP | GO:0070663 | regulation of leukocyte proliferation | 3.52E-05 | 6.08E-04 | Irf1/H2-Aa/Cd74/H | 10 | aPD-L1+aTGF-β vs PBS | Up-regulated genes | CD45+ |
| BP | GO:0019079 | viral genome replication | 3.85E-05 | 6.61E-04 | Rsad2/Oasl1/Tnf/ | 7 | aPD-L1+aTGF-β vs PBS | Up-regulated genes | CD45+ |
| BP | GO:0046634 | regulation of alpha-beta T cell activation | 4.11E-05 | 7.01E-04 | Irf1/H2-Ab1/Cd27 | 7 | aPD-L1+aTGF-β vs PBS | Up-regulated genes | CD45+ |
| BP | GO:0045429 | positive regulation of nitric oxide biosynthetic pro | 4.16E-05 | 7.02E-04 | Il1b/Icam1/Tnf/So | 5 | aPD-L1+aTGF-β vs PBS | Up-regulated genes | CD45+ |
| BP | GO:0072677 | eosinophil migration | 4.18E-05 | 7.02E-04 | Ccl4/Ccl5/Ccl2/Cc | 4 | aPD-L1+aTGF-β vs PBS | Up-regulated genes | CD45+ |
| BP | GO:0002443 | leukocyte mediated immunity | 4.46E-05 | 7.44E-04 | Cd74/H2-Ab1/Rsa | 12 | aPD-L1+aTGF-β vs PBS | Up-regulated genes | CD45+ |
| BP | GO:0002831 | regulation of response to biotic stimulus | 4.68E-05 | 7.62E-04 | Cd274/Il1b/Arg2/( | 8 | aPD-L1+aTGF-β vs PBS | Up-regulated genes | CD45+ |
| BP | GO:0002688 | regulation of leukocyte chemotaxis | 4.69E-05 | 7.62E-04 | Cd74/Cxcl10/Il1b/ | 7 | aPD-L1+aTGF-β vs PBS | Up-regulated genes | CD45+ |
| BP | GO:0045621 | positive regulation of lymphocyte differentiation | 4.69E-05 | 7.62E-04 | Irf1/H2-Aa/Cd74/Il | 7 | aPD-L1+aTGF-β vs PBS | Up-regulated genes | CD45+ |
| BP | GO:1904407 | positive regulation of nitric oxide metabolic proce | 4.69E-05 | 7.62E-04 | Il1b/Icam1/Tnf/So | 5 | aPD-L1+aTGF-β vs PBS | Up-regulated genes | CD45+ |
| BP | GO:0090025 | regulation of monocyte chemotaxis | 5.07E-05 | 8.18E-04 | Cxcl10/Ccl5/Ccl2/ | 4 | aPD-L1+aTGF-β vs PBS | Up-regulated genes | CD45+ |
| BP | GO:1903901 | negative regulation of viral life cycle | 5.79E-05 | 9.28E-04 | Rsad2/Oasl1/Tnf/ | 6 | aPD-L1+aTGF-β vs PBS | Up-regulated genes | CD45+ |
| BP | GO:0045581 | negative regulation of T cell differentiation | 5.90E-05 | 9.40E-04 | Irf1/Cd74/Socs1/C | 5 | aPD-L1+aTGF-β vs PBS | Up-regulated genes | CD45+ |
| BP | GO:0060759 | regulation of response to cytokine stimulus | 6.03E-05 | 9.46E-04 | Cd74/Samhd1/Irg | 7 | aPD-L1+aTGF-β vs PBS | Up-regulated genes | CD45+ |
| BP | GO:0002705 | positive regulation of leukocyte mediated immun | 6.04E-05 | 9.46E-04 | Rsad2/Il1b/H2-K1 | 8 | aPD-L1+aTGF-β vs PBS | Up-regulated genes | CD45+ |
| BP | GO:0046426 | negative regulation of JAK-STAT cascade | 6.09E-05 | 9.46E-04 | Irf1/Socs1/Parp14 | 4 | aPD-L1+aTGF-β vs PBS | Up-regulated genes | CD45+ |
| BP | GO:1904893 | negative regulation of STAT cascade | 6.09E-05 | 9.46E-04 | Irf1/Socs1/Parp14 | 4 | aPD-L1+aTGF-β vs PBS | Up-regulated genes | CD45+ |
| BP | GO:0002718 | regulation of cytokine production involved in imm | 7.89E-05 | 1.20E-03 | Cd74/Rsad2/Il1b/ | 6 | aPD-L1+aTGF-β vs PBS | Up-regulated genes | CD45+ |
| BP | GO:0032945 | negative regulation of mononuclear cell prolifera | 7.89E-05 | 1.20E-03 | H2-Aa/H2-Ab1/Cc | 6 | aPD-L1+aTGF-β vs PBS | Up-regulated genes | CD45+ |
| BP | GO:0050672 | negative regulation of lymphocyte proliferation | 7.89E-05 | 1.20E-03 | H2-Aa/H2-Ab1/Cc | 6 | aPD-L1+aTGF-β vs PBS | Up-regulated genes | CD45+ |
| BP | GO:0001818 | negative regulation of cytokine production | 8.09E-05 | 1.23E-03 | Cd274/Arg2/Gbp4 | 10 | aPD-L1+aTGF-β vs PBS | Up-regulated genes | CD45+ |
| BP | GO:0002440 | production of molecular mediator of immune res | 8.49E-05 | 1.27E-03 | Cd74/H2-Ab1/Sai | 9 | aPD-L1+aTGF-β vs PBS | Up-regulated genes | CD45+ |
| BP | GO:0042098 | T cell proliferation | 8.49E-05 | 1.27E-03 | Irf1/H2-Aa/H2-Ab | 9 | aPD-L1+aTGF-β vs PBS | Up-regulated genes | CD45+ |

| | | | | | | | | | |
|---|---|---|---|---|---|---|---|---|---|
| BP | GO:0006953 | acute-phase response | 8.57E-05 | 1.28E-03 | Il1b/Tnf/Ccl5/Il1rn | 4 | aPD-L1+aTGF-β vs PBS | Up-regulated genes | CD45+ |
| BP | GO:0045824 | negative regulation of innate immune response | 9.02E-05 | 1.33E-03 | Samhd1/Acod1/P | 5 | aPD-L1+aTGF-β vs PBS | Up-regulated genes | CD45+ |
| BP | GO:0031652 | positive regulation of heat generation | 9.70E-05 | 1.39E-03 | Il1b/Tnf/Ccl5 | 3 | aPD-L1+aTGF-β vs PBS | Up-regulated genes | CD45+ |
| BP | GO:0042090 | interleukin-12 biosynthetic process | 9.70E-05 | 1.39E-03 | Irf1/Rel/Nfkb1 | 3 | aPD-L1+aTGF-β vs PBS | Up-regulated genes | CD45+ |
| BP | GO:0042359 | vitamin D metabolic process | 9.70E-05 | 1.39E-03 | Il1b/Tnf/Nfkb1 | 3 | aPD-L1+aTGF-β vs PBS | Up-regulated genes | CD45+ |
| BP | GO:0045075 | regulation of interleukin-12 biosynthetic process | 9.70E-05 | 1.39E-03 | Irf1/Rel/Nfkb1 | 3 | aPD-L1+aTGF-β vs PBS | Up-regulated genes | CD45+ |
| BP | GO:0051956 | negative regulation of amino acid transport | 9.70E-05 | 1.39E-03 | Il1b/Tnf/Il1rn | 3 | aPD-L1+aTGF-β vs PBS | Up-regulated genes | CD45+ |
| BP | GO:0070664 | negative regulation of leukocyte proliferation | 9.83E-05 | 1.40E-03 | H2-Aa/H2-Ab1/Cc | 6 | aPD-L1+aTGF-β vs PBS | Up-regulated genes | CD45+ |
| BP | GO:0002720 | positive regulation of cytokine production involve | 9.97E-05 | 1.41E-03 | Cd74/Rsad2/Il1b/ | 5 | aPD-L1+aTGF-β vs PBS | Up-regulated genes | CD45+ |
| BP | GO:0002690 | positive regulation of leukocyte chemotaxis | 1.06E-04 | 1.49E-03 | Cd74/Cxcl10/Il1b | 6 | aPD-L1+aTGF-β vs PBS | Up-regulated genes | CD45+ |
| BP | GO:0009636 | response to toxic substance | 1.07E-04 | 1.50E-03 | Ccl4/Tnf/Nfe2l2/C | 10 | aPD-L1+aTGF-β vs PBS | Up-regulated genes | CD45+ |
| BP | GO:0050732 | negative regulation of peptidyl-tyrosine phospho | 1.10E-04 | 1.53E-03 | Irf1/Socs1/Parp14 | 5 | aPD-L1+aTGF-β vs PBS | Up-regulated genes | CD45+ |
| BP | GO:0051955 | regulation of amino acid transport | 1.17E-04 | 1.63E-03 | Il1b/Arg2/Tnf/Il1rn | 4 | aPD-L1+aTGF-β vs PBS | Up-regulated genes | CD45+ |
| BP | GO:0045620 | negative regulation of lymphocyte differentiation | 1.21E-04 | 1.67E-03 | Irf1/Cd74/Socs1/C | 5 | aPD-L1+aTGF-β vs PBS | Up-regulated genes | CD45+ |
| BP | GO:0007259 | JAK-STAT cascade | 1.27E-04 | 1.74E-03 | Irf1/Socs1/Tnf/Ccl | 7 | aPD-L1+aTGF-β vs PBS | Up-regulated genes | CD45+ |
| BP | GO:0072683 | T cell extravasation | 1.32E-04 | 1.79E-03 | Icam1/Ccl5/Ccl2 | 3 | aPD-L1+aTGF-β vs PBS | Up-regulated genes | CD45+ |
| BP | GO:2000343 | positive regulation of chemokine (C-X-C motif) li | 1.32E-04 | 1.79E-03 | Cd74/Tnf/Ccl5 | 3 | aPD-L1+aTGF-β vs PBS | Up-regulated genes | CD45+ |
| BP | GO:1903428 | positive regulation of reactive oxygen species bi | 1.33E-04 | 1.79E-03 | Il1b/Icam1/Tnf/So | 5 | aPD-L1+aTGF-β vs PBS | Up-regulated genes | CD45+ |
| BP | GO:0043372 | positive regulation of CD4-positive, alpha-beta T | 1.36E-04 | 1.82E-03 | Irf1/Socs1/Nlrp3/C | 4 | aPD-L1+aTGF-β vs PBS | Up-regulated genes | CD45+ |
| BP | GO:1902532 | negative regulation of intracellular signal transdu | 1.36E-04 | 1.82E-03 | Irf1/Cd74/Dnaja1/ | 14 | aPD-L1+aTGF-β vs PBS | Up-regulated genes | CD45+ |
| BP | GO:0002706 | regulation of lymphocyte mediated immunity | 1.40E-04 | 1.86E-03 | Rsad2/Il1b/H2-K1 | 8 | aPD-L1+aTGF-β vs PBS | Up-regulated genes | CD45+ |
| BP | GO:0070374 | positive regulation of ERK1 and ERK2 cascade | 1.46E-04 | 1.93E-03 | Cd74/Il1b/Ccl4/Ica | 8 | aPD-L1+aTGF-β vs PBS | Up-regulated genes | CD45+ |
| BP | GO:1901652 | response to peptide | 1.47E-04 | 1.94E-03 | Irf1/Il1b/Icam1/So | 11 | aPD-L1+aTGF-β vs PBS | Up-regulated genes | CD45+ |
| BP | GO:0097696 | STAT cascade | 1.49E-04 | 1.94E-03 | Irf1/Socs1/Tnf/Ccl | 7 | aPD-L1+aTGF-β vs PBS | Up-regulated genes | CD45+ |
| BP | GO:0002526 | acute inflammatory response | 1.59E-04 | 2.06E-03 | Il1b/Icam1/Tnf/Nlr | 6 | aPD-L1+aTGF-β vs PBS | Up-regulated genes | CD45+ |
| BP | GO:0030656 | regulation of vitamin metabolic process | 1.75E-04 | 2.24E-03 | Il1b/Tnf/Nfkb1 | 3 | aPD-L1+aTGF-β vs PBS | Up-regulated genes | CD45+ |
| BP | GO:0031650 | regulation of heat generation | 1.75E-04 | 2.24E-03 | Il1b/Tnf/Ccl5 | 3 | aPD-L1+aTGF-β vs PBS | Up-regulated genes | CD45+ |
| BP | GO:0042362 | fat-soluble vitamin biosynthetic process | 1.75E-04 | 2.24E-03 | Il1b/Tnf/Nfkb1 | 3 | aPD-L1+aTGF-β vs PBS | Up-regulated genes | CD45+ |
| BP | GO:0002724 | regulation of T cell cytokine production | 1.79E-04 | 2.28E-03 | Rsad2/Il1b/Nlrp3/ | 4 | aPD-L1+aTGF-β vs PBS | Up-regulated genes | CD45+ |
| BP | GO:0046651 | lymphocyte proliferation | 1.92E-04 | 2.43E-03 | Irf1/H2-Aa/Cd74/H | 10 | aPD-L1+aTGF-β vs PBS | Up-regulated genes | CD45+ |
| BP | GO:0050663 | cytokine secretion | 1.95E-04 | 2.46E-03 | Cd274/Il1b/Arg2/S | 9 | aPD-L1+aTGF-β vs PBS | Up-regulated genes | CD45+ |
| BP | GO:0060337 | type I interferon signaling pathway | 2.04E-04 | 2.51E-03 | Samhd1/Zbp1/Sta | 4 | aPD-L1+aTGF-β vs PBS | Up-regulated genes | CD45+ |
| BP | GO:0071357 | cellular response to type I interferon | 2.04E-04 | 2.51E-03 | Samhd1/Zbp1/Sta | 4 | aPD-L1+aTGF-β vs PBS | Up-regulated genes | CD45+ |
| BP | GO:0045639 | positive regulation of myeloid cell differentiation | 2.04E-04 | 2.51E-03 | Cd74/Tnf/Ccl5/Sta | 6 | aPD-L1+aTGF-β vs PBS | Up-regulated genes | CD45+ |
| BP | GO:1903426 | regulation of reactive oxygen species biosynthet | 2.04E-04 | 2.51E-03 | Il1b/Arg2/Icam1/T | 6 | aPD-L1+aTGF-β vs PBS | Up-regulated genes | CD45+ |
| BP | GO:0032943 | mononuclear cell proliferation | 2.04E-04 | 2.51E-03 | Irf1/H2-Aa/Cd74/H | 10 | aPD-L1+aTGF-β vs PBS | Up-regulated genes | CD45+ |
| BP | GO:0002922 | positive regulation of humoral immune response | 2.26E-04 | 2.77E-03 | Il1b/Tnf/Acod1 | 3 | aPD-L1+aTGF-β vs PBS | Up-regulated genes | CD45+ |
| BP | GO:0050730 | regulation of peptidyl-tyrosine phosphorylation | 2.32E-04 | 2.82E-03 | Irf1/Cd74/Icam1/S | 9 | aPD-L1+aTGF-β vs PBS | Up-regulated genes | CD45+ |
| BP | GO:0002687 | positive regulation of leukocyte migration | 2.33E-04 | 2.82E-03 | Cd74/Cxcl10/Il1b | 7 | aPD-L1+aTGF-β vs PBS | Up-regulated genes | CD45+ |
| BP | GO:0097237 | cellular response to toxic substance | 2.44E-04 | 2.94E-03 | Tnf/Nfe2l2/Txn1/K | 7 | aPD-L1+aTGF-β vs PBS | Up-regulated genes | CD45+ |
| BP | GO:0071417 | cellular response to organonitrogen compound | 2.45E-04 | 2.94E-03 | Irf1/Il1b/Icam1/So | 12 | aPD-L1+aTGF-β vs PBS | Up-regulated genes | CD45+ |
| BP | GO:0001916 | positive regulation of T cell mediated cytotoxicity | 2.61E-04 | 3.12E-03 | H2-K1/H2-D1/Hsp | 4 | aPD-L1+aTGF-β vs PBS | Up-regulated genes | CD45+ |
| BP | GO:0045428 | regulation of nitric oxide biosynthetic process | 2.63E-04 | 3.12E-03 | Il1b/Icam1/Tnf/So | 5 | aPD-L1+aTGF-β vs PBS | Up-regulated genes | CD45+ |
| BP | GO:1904950 | negative regulation of establishment of protein lo | 2.70E-04 | 3.20E-03 | Dnaja1/Rsad2/Il1 | 8 | aPD-L1+aTGF-β vs PBS | Up-regulated genes | CD45+ |
| BP | GO:0002367 | cytokine production involved in immune respons | 2.75E-04 | 3.24E-03 | Cd74/Rsad2/Il1b/ | 6 | aPD-L1+aTGF-β vs PBS | Up-regulated genes | CD45+ |
| BP | GO:2000377 | regulation of reactive oxygen species metabolic | 2.81E-04 | 3.29E-03 | Il1b/Arg2/Icam1/T | 8 | aPD-L1+aTGF-β vs PBS | Up-regulated genes | CD45+ |
| BP | GO:0042274 | ribosomal small subunit biogenesis | 2.84E-04 | 3.29E-03 | Rps27/Rps21/Rpl | 5 | aPD-L1+aTGF-β vs PBS | Up-regulated genes | CD45+ |
| BP | GO:0060330 | regulation of response to interferon-gamma | 2.86E-04 | 3.29E-03 | Irgm1/Socs1/Parp | 3 | aPD-L1+aTGF-β vs PBS | Up-regulated genes | CD45+ |
| BP | GO:0060334 | regulation of interferon-gamma-mediated signali | 2.86E-04 | 3.29E-03 | Irgm1/Socs1/Parp | 3 | aPD-L1+aTGF-β vs PBS | Up-regulated genes | CD45+ |
| BP | GO:2000341 | regulation of chemokine (C-X-C motif) ligand 2 p | 2.86E-04 | 3.29E-03 | Cd74/Tnf/Ccl5 | 3 | aPD-L1+aTGF-β vs PBS | Up-regulated genes | CD45+ |
| BP | GO:0032768 | regulation of monooxygenase activity | 2.94E-04 | 3.35E-03 | Il1b/Arg2/Tnf/Nfkb | 4 | aPD-L1+aTGF-β vs PBS | Up-regulated genes | CD45+ |
| BP | GO:2000516 | positive regulation of CD4-positive, alpha-beta T | 2.94E-04 | 3.35E-03 | Irf1/Socs1/Nlrp3/C | 4 | aPD-L1+aTGF-β vs PBS | Up-regulated genes | CD45+ |
| BP | GO:0048872 | homeostasis of number of cells | 3.10E-04 | 3.51E-03 | Cd74/Pde4b/B2m | 10 | aPD-L1+aTGF-β vs PBS | Up-regulated genes | CD45+ |
| BP | GO:0002224 | toll-like receptor signaling pathway | 3.26E-04 | 3.68E-03 | Irf1/Rsad2/Tnf/Nfk | 6 | aPD-L1+aTGF-β vs PBS | Up-regulated genes | CD45+ |
| BP | GO:0070661 | leukocyte proliferation | 3.29E-04 | 3.69E-03 | Irf1/H2-Aa/Cd74/H | 10 | aPD-L1+aTGF-β vs PBS | Up-regulated genes | CD45+ |
| BP | GO:0001912 | positive regulation of leukocyte mediated cytotox | 3.30E-04 | 3.69E-03 | H2-K1/H2-D1/Hsp | 5 | aPD-L1+aTGF-β vs PBS | Up-regulated genes | CD45+ |
| BP | GO:0001906 | cell killing | 3.37E-04 | 3.75E-03 | H2-K1/H2-D1/Hsp | 7 | aPD-L1+aTGF-β vs PBS | Up-regulated genes | CD45+ |
| BP | GO:0035821 | modification of morphology or physiology of othe | 3.45E-04 | 3.82E-03 | Ccl4/Hspa8/Ccl5/ | 6 | aPD-L1+aTGF-β vs PBS | Up-regulated genes | CD45+ |
| BP | GO:0002685 | regulation of leukocyte migration | 3.52E-04 | 3.89E-03 | Cd74/Cxcl10/Il1b | 8 | aPD-L1+aTGF-β vs PBS | Up-regulated genes | CD45+ |
| BP | GO:0072567 | chemokine (C-X-C motif) ligand 2 production | 3.55E-04 | 3.90E-03 | Cd74/Tnf/Ccl5 | 3 | aPD-L1+aTGF-β vs PBS | Up-regulated genes | CD45+ |
| BP | GO:0046425 | regulation of JAK-STAT cascade | 3.64E-04 | 3.98E-03 | Irf1/Socs1/Tnf/Ccl | 6 | aPD-L1+aTGF-β vs PBS | Up-regulated genes | CD45+ |
| BP | GO:0046635 | positive regulation of alpha-beta T cell activation | 3.82E-04 | 4.16E-03 | Irf1/H2-Ab1/Socs | 5 | aPD-L1+aTGF-β vs PBS | Up-regulated genes | CD45+ |
| BP | GO:1903707 | negative regulation of hemopoiesis | 4.01E-04 | 4.34E-03 | Irf1/Cd74/Socs1/C | 7 | aPD-L1+aTGF-β vs PBS | Up-regulated genes | CD45+ |
| BP | GO:0043410 | positive regulation of MAPK cascade | 4.02E-04 | 4.34E-03 | Cd74/Il1b/Ccl4/Ica | 12 | aPD-L1+aTGF-β vs PBS | Up-regulated genes | CD45+ |
| BP | GO:1901653 | cellular response to peptide | 4.04E-04 | 4.34E-03 | Irf1/Il1b/Icam1/So | 9 | aPD-L1+aTGF-β vs PBS | Up-regulated genes | CD45+ |
| BP | GO:0046632 | alpha-beta T cell differentiation | 4.28E-04 | 4.56E-03 | Irf1/Rsad2/Socs1/ | 6 | aPD-L1+aTGF-β vs PBS | Up-regulated genes | CD45+ |
| BP | GO:1903409 | reactive oxygen species biosynthetic process | 4.28E-04 | 4.56E-03 | Il1b/Arg2/Icam1/T | 6 | aPD-L1+aTGF-β vs PBS | Up-regulated genes | CD45+ |
| BP | GO:0002830 | positive regulation of type 2 immune response | 4.34E-04 | 4.56E-03 | Cd74/Rsad2/Nlrp | 3 | aPD-L1+aTGF-β vs PBS | Up-regulated genes | CD45+ |
| BP | GO:0019430 | removal of superoxide radicals | 4.34E-04 | 4.56E-03 | Tnf/Nfe2l2/Sod2 | 3 | aPD-L1+aTGF-β vs PBS | Up-regulated genes | CD45+ |
| BP | GO:0051953 | negative regulation of amine transport | 4.34E-04 | 4.56E-03 | Il1b/Tnf/Il1rn | 3 | aPD-L1+aTGF-β vs PBS | Up-regulated genes | CD45+ |
| BP | GO:0006809 | nitric oxide biosynthetic process | 4.40E-04 | 4.61E-03 | Il1b/Icam1/Tnf/So | 5 | aPD-L1+aTGF-β vs PBS | Up-regulated genes | CD45+ |
| BP | GO:1904892 | regulation of STAT cascade | 4.51E-04 | 4.68E-03 | Irf1/Socs1/Tnf/Ccl | 6 | aPD-L1+aTGF-β vs PBS | Up-regulated genes | CD45+ |
| BP | GO:0045637 | regulation of myeloid cell differentiation | 4.54E-04 | 4.68E-03 | Cd74/Tnf/Nfkbia/C | 8 | aPD-L1+aTGF-β vs PBS | Up-regulated genes | CD45+ |
| BP | GO:0001914 | regulation of T cell mediated cytotoxicity | 4.54E-04 | 4.68E-03 | H2-K1/H2-D1/Hsp | 4 | aPD-L1+aTGF-β vs PBS | Up-regulated genes | CD45+ |
| BP | GO:0048246 | macrophage chemotaxis | 4.54E-04 | 4.68E-03 | Ccl5/Ccl3/Ccl2/Sl | 4 | aPD-L1+aTGF-β vs PBS | Up-regulated genes | CD45+ |
| BP | GO:0051341 | regulation of oxidoreductase activity | 5.05E-04 | 5.17E-03 | Il1b/Arg2/Tnf/Nfkb | 5 | aPD-L1+aTGF-β vs PBS | Up-regulated genes | CD45+ |
| BP | GO:0090026 | positive regulation of monocyte chemotaxis | 5.23E-04 | 5.32E-03 | Cxcl10/Ccl5/Ccl2 | 3 | aPD-L1+aTGF-β vs PBS | Up-regulated genes | CD45+ |
| BP | GO:2001185 | regulation of CD8-positive, alpha-beta T cell acti | 5.23E-04 | 5.32E-03 | Irf1/Cd274/Socs1 | 3 | aPD-L1+aTGF-β vs PBS | Up-regulated genes | CD45+ |
| BP | GO:0046209 | nitric oxide metabolic process | 5.39E-04 | 5.46E-03 | Il1b/Icam1/Tnf/So | 5 | aPD-L1+aTGF-β vs PBS | Up-regulated genes | CD45+ |
| BP | GO:0002369 | T cell cytokine production | 5.54E-04 | 5.54E-03 | Rsad2/Il1b/Nlrp3/ | 4 | aPD-L1+aTGF-β vs PBS | Up-regulated genes | CD45+ |
| BP | GO:0043331 | response to dsRNA | 5.54E-04 | 5.54E-03 | Nfkbia/Stat1/Nfkb | 4 | aPD-L1+aTGF-β vs PBS | Up-regulated genes | CD45+ |
| BP | GO:0046636 | negative regulation of alpha-beta T cell activatio | 5.54E-04 | 5.54E-03 | Irf1/Cd274/Arg2/S | 4 | aPD-L1+aTGF-β vs PBS | Up-regulated genes | CD45+ |
| BP | GO:0043122 | regulation of I-kappaB kinase/NF-kappaB signal | 5.81E-04 | 5.79E-03 | Cd74/Il1b/Tnf/Sta | 7 | aPD-L1+aTGF-β vs PBS | Up-regulated genes | CD45+ |
| BP | GO:0015800 | acidic amino acid transport | 6.10E-04 | 5.97E-03 | Il1b/Tnf/Slc7a11/Il | 4 | aPD-L1+aTGF-β vs PBS | Up-regulated genes | CD45+ |
| BP | GO:1900407 | regulation of cellular response to oxidative stres | 6.14E-04 | 5.97E-03 | Tnf/Nfe2l2/Txn1/S | 5 | aPD-L1+aTGF-β vs PBS | Up-regulated genes | CD45+ |
| BP | GO:0000028 | ribosomal small subunit assembly | 6.24E-04 | 5.97E-03 | Rps27/Rpl38/Rps | 3 | aPD-L1+aTGF-β vs PBS | Up-regulated genes | CD45+ |
| BP | GO:0002523 | leukocyte migration involved in inflammatory res | 6.24E-04 | 5.97E-03 | S100a8/Ccl2/Slar | 3 | aPD-L1+aTGF-β vs PBS | Up-regulated genes | CD45+ |
| BP | GO:0002577 | regulation of antigen processing and presentatio | 6.24E-04 | 5.97E-03 | Cd74/H2-Ab1/Fgl | 3 | aPD-L1+aTGF-β vs PBS | Up-regulated genes | CD45+ |
| BP | GO:0009110 | vitamin biosynthetic process | 6.24E-04 | 5.97E-03 | Il1b/Tnf/Nfkb1 | 3 | aPD-L1+aTGF-β vs PBS | Up-regulated genes | CD45+ |

| | | | | | | | | | |
|---|---|---|---|---|---|---|---|---|---|
| BP | GO:0032891 | negative regulation of organic acid transport | 6.24E-04 | 5.97E-03 | Il1b/Tnf/Il1rn | 3 | aPD-L1+aTGF-β vs PBS | Up-regulated genes | CD45+ |
| BP | GO:0051770 | positive regulation of nitric-oxide synthase biosy | 6.24E-04 | 5.97E-03 | Nampt/Stat1/Ccl2 | 3 | aPD-L1+aTGF-β vs PBS | Up-regulated genes | CD45+ |
| BP | GO:0071450 | cellular response to oxygen radical | 6.24E-04 | 5.97E-03 | Tnf/Nfe2l2/Sod2 | 3 | aPD-L1+aTGF-β vs PBS | Up-regulated genes | CD45+ |
| BP | GO:0071451 | cellular response to superoxide | 6.24E-04 | 5.97E-03 | Tnf/Nfe2l2/Sod2 | 3 | aPD-L1+aTGF-β vs PBS | Up-regulated genes | CD45+ |
| BP | GO:0050688 | regulation of defense response to virus | 6.54E-04 | 6.24E-03 | Il1b/Gbp4/Ccl5/Fç | 5 | aPD-L1+aTGF-β vs PBS | Up-regulated genes | CD45+ |
| BP | GO:0050702 | interleukin-1 beta secretion | 6.69E-04 | 6.33E-03 | Arg2/Nlrp3/Gbp5/ | 4 | aPD-L1+aTGF-β vs PBS | Up-regulated genes | CD45+ |
| BP | GO:1903202 | negative regulation of oxidative stress-induced c | 6.69E-04 | 6.33E-03 | Nfe2l2/Txn1/Slc7ε | 4 | aPD-L1+aTGF-β vs PBS | Up-regulated genes | CD45+ |
| BP | GO:0032611 | interleukin-1 beta production | 6.96E-04 | 6.54E-03 | Il1b/Arg2/Nlrp3/Gl | 5 | aPD-L1+aTGF-β vs PBS | Up-regulated genes | CD45+ |
| BP | GO:2001057 | reactive nitrogen species metabolic process | 6.96E-04 | 6.54E-03 | Il1b/Icam1/Tnf/So | 5 | aPD-L1+aTGF-β vs PBS | Up-regulated genes | CD45+ |
| BP | GO:1901699 | cellular response to nitrogen compound | 7.07E-04 | 6.62E-03 | Irf1/Il1b/Icam1/So | 12 | aPD-L1+aTGF-β vs PBS | Up-regulated genes | CD45+ |
| BP | GO:0001961 | positive regulation of cytokine-mediated signalin | 7.32E-04 | 6.72E-03 | Cd74/Irgm1/Zbp1 | 4 | aPD-L1+aTGF-β vs PBS | Up-regulated genes | CD45+ |
| BP | GO:1900408 | negative regulation of cellular response to oxida | 7.32E-04 | 6.72E-03 | Nfe2l2/Txn1/Slc7ε | 4 | aPD-L1+aTGF-β vs PBS | Up-regulated genes | CD45+ |
| BP | GO:0002428 | antigen processing and presentation of peptide a | 7.36E-04 | 6.72E-03 | H2-K1/H2-D1/B2r | 3 | aPD-L1+aTGF-β vs PBS | Up-regulated genes | CD45+ |
| BP | GO:0098869 | cellular oxidant detoxification | 7.36E-04 | 6.72E-03 | Tnf/Nfe2l2/Sod2 | 3 | aPD-L1+aTGF-β vs PBS | Up-regulated genes | CD45+ |
| BP | GO:1901623 | regulation of lymphocyte chemotaxis | 7.36E-04 | 6.72E-03 | Cxcl10/Ccl4/Ccl5 | 3 | aPD-L1+aTGF-β vs PBS | Up-regulated genes | CD45+ |
| BP | GO:0071236 | cellular response to antibiotic | 7.37E-04 | 6.72E-03 | Crip1/Nfe2l2/Txn' | 6 | aPD-L1+aTGF-β vs PBS | Up-regulated genes | CD45+ |
| BP | GO:0031343 | positive regulation of cell killing | 7.40E-04 | 6.72E-03 | H2-K1/H2-D1/Hsp | 5 | aPD-L1+aTGF-β vs PBS | Up-regulated genes | CD45+ |
| BP | GO:0043367 | CD4-positive, alpha-beta T cell differentiation | 7.40E-04 | 6.72E-03 | Irf1/Rsad2/Socs1/ | 5 | aPD-L1+aTGF-β vs PBS | Up-regulated genes | CD45+ |
| BP | GO:2001234 | negative regulation of apoptotic signaling pathw | 7.75E-04 | 7.01E-03 | Cd74/Dnaja1/Il1b | 8 | aPD-L1+aTGF-β vs PBS | Up-regulated genes | CD45+ |
| BP | GO:0043506 | regulation of JUN kinase activity | 7.87E-04 | 7.09E-03 | Dnaja1/Il1b/Tnf/S | 5 | aPD-L1+aTGF-β vs PBS | Up-regulated genes | CD45+ |
| BP | GO:0071675 | regulation of mononuclear cell migration | 7.99E-04 | 7.18E-03 | Cxcl10/Ccl5/Ccl2/ | 4 | aPD-L1+aTGF-β vs PBS | Up-regulated genes | CD45+ |
| BP | GO:0071407 | cellular response to organic cyclic compound | 8.08E-04 | 7.23E-03 | Dnaja1/Il1b/Socs' | 10 | aPD-L1+aTGF-β vs PBS | Up-regulated genes | CD45+ |
| BP | GO:0002703 | regulation of leukocyte mediated immunity | 8.25E-04 | 7.36E-03 | Rsad2/Il1b/H2-K1 | 8 | aPD-L1+aTGF-β vs PBS | Up-regulated genes | CD45+ |
| BP | GO:0032479 | regulation of type I interferon production | 8.35E-04 | 7.39E-03 | Irf1/Gbp4/Rel/Aco | 5 | aPD-L1+aTGF-β vs PBS | Up-regulated genes | CD45+ |
| BP | GO:0032642 | regulation of chemokine production | 8.35E-04 | 7.39E-03 | Cd74/Il1b/Arg2/Tr | 5 | aPD-L1+aTGF-β vs PBS | Up-regulated genes | CD45+ |
| BP | GO:0006775 | fat-soluble vitamin metabolic process | 8.59E-04 | 7.53E-03 | Il1b/Tnf/Nfkb1 | 3 | aPD-L1+aTGF-β vs PBS | Up-regulated genes | CD45+ |
| BP | GO:0032897 | negative regulation of viral transcription | 8.59E-04 | 7.53E-03 | Ccl4/Ccl5/Ifitm3 | 3 | aPD-L1+aTGF-β vs PBS | Up-regulated genes | CD45+ |
| BP | GO:0034114 | regulation of heterotypic cell-cell adhesion | 8.59E-04 | 7.53E-03 | Il1b/Tnf/Il1rn | 3 | aPD-L1+aTGF-β vs PBS | Up-regulated genes | CD45+ |
| BP | GO:0046638 | positive regulation of alpha-beta T cell differentia | 8.71E-04 | 7.60E-03 | Irf1/Socs1/Nlrp3/C | 4 | aPD-L1+aTGF-β vs PBS | Up-regulated genes | CD45+ |
| BP | GO:1902882 | regulation of response to oxidative stress | 9.38E-04 | 8.16E-03 | Tnf/Nfe2l2/Txn1/S | 5 | aPD-L1+aTGF-β vs PBS | Up-regulated genes | CD45+ |
| BP | GO:0071622 | regulation of granulocyte chemotaxis | 9.47E-04 | 8.18E-03 | Cd74/Il1b/Ccl5/Cc | 4 | aPD-L1+aTGF-β vs PBS | Up-regulated genes | CD45+ |
| BP | GO:1902883 | negative regulation of response to oxidative stre | 9.47E-04 | 8.18E-03 | Nfe2l2/Txn1/Slc7ε | 4 | aPD-L1+aTGF-β vs PBS | Up-regulated genes | CD45+ |
| BP | GO:0018108 | peptidyl-tyrosine phosphorylation | 9.60E-04 | 8.26E-03 | Irf1/Cd74/Icam1/S | 9 | aPD-L1+aTGF-β vs PBS | Up-regulated genes | CD45+ |
| BP | GO:0050921 | positive regulation of chemotaxis | 9.67E-04 | 8.29E-03 | Cd74/Cxcl10/Il1b/ | 6 | aPD-L1+aTGF-β vs PBS | Up-regulated genes | CD45+ |
| BP | GO:0042136 | neurotransmitter biosynthetic process | 9.93E-04 | 8.48E-03 | Il1b/Icam1/Tnf/So | 5 | aPD-L1+aTGF-β vs PBS | Up-regulated genes | CD45+ |
| BP | GO:0000303 | response to superoxide | 9.96E-04 | 8.48E-03 | Tnf/Nfe2l2/Sod2 | 3 | aPD-L1+aTGF-β vs PBS | Up-regulated genes | CD45+ |
| BP | GO:0043370 | regulation of CD4-positive, alpha-beta T cell diffe | 1.03E-03 | 8.72E-03 | Irf1/Socs1/Nlrp3/C | 4 | aPD-L1+aTGF-β vs PBS | Up-regulated genes | CD45+ |
| BP | GO:0018212 | peptidyl-tyrosine modification | 1.04E-03 | 8.79E-03 | Irf1/Cd74/Icam1/S | 9 | aPD-L1+aTGF-β vs PBS | Up-regulated genes | CD45+ |
| BP | GO:0032602 | chemokine production | 1.05E-03 | 8.82E-03 | Cd74/Il1b/Arg2/Tr | 5 | aPD-L1+aTGF-β vs PBS | Up-regulated genes | CD45+ |
| BP | GO:0032606 | type I interferon production | 1.05E-03 | 8.82E-03 | Irf1/Gbp4/Rel/Aco | 5 | aPD-L1+aTGF-β vs PBS | Up-regulated genes | CD45+ |
| BP | GO:0034101 | erythrocyte homeostasis | 1.05E-03 | 8.83E-03 | B2m/Stat1/Klf2/Isç | 6 | aPD-L1+aTGF-β vs PBS | Up-regulated genes | CD45+ |
| BP | GO:0001910 | regulation of leukocyte mediated cytotoxicity | 1.11E-03 | 9.25E-03 | H2-K1/H2-D1/Hsp | 5 | aPD-L1+aTGF-β vs PBS | Up-regulated genes | CD45+ |
| BP | GO:0050701 | interleukin-1 secretion | 1.11E-03 | 9.25E-03 | Arg2/Nlrp3/Gbp5/ | 4 | aPD-L1+aTGF-β vs PBS | Up-regulated genes | CD45+ |
| BP | GO:0000305 | response to oxygen radical | 1.14E-03 | 9.39E-03 | Tnf/Nfe2l2/Sod2 | 3 | aPD-L1+aTGF-β vs PBS | Up-regulated genes | CD45+ |
| BP | GO:0002475 | antigen processing and presentation via MHC cl | 1.14E-03 | 9.39E-03 | H2-K1/H2-D1/B2r | 3 | aPD-L1+aTGF-β vs PBS | Up-regulated genes | CD45+ |
| BP | GO:0002827 | positive regulation of T-helper 1 type immune re: | 1.14E-03 | 9.39E-03 | Irf1/H2-Ab1/Il1b | 3 | aPD-L1+aTGF-β vs PBS | Up-regulated genes | CD45+ |
| BP | GO:0071677 | positive regulation of mononuclear cell migratior | 1.14E-03 | 9.39E-03 | Cxcl10/Ccl5/Ccl2 | 3 | aPD-L1+aTGF-β vs PBS | Up-regulated genes | CD45+ |
| BP | GO:0050728 | negative regulation of inflammatory response | 1.15E-03 | 9.39E-03 | Nlrp3/Acod1/Socs | 6 | aPD-L1+aTGF-β vs PBS | Up-regulated genes | CD45+ |
| BP | GO:0002673 | regulation of acute inflammatory response | 1.20E-03 | 9.80E-03 | Il1b/Tnf/Nlrp3/Ccl | 4 | aPD-L1+aTGF-β vs PBS | Up-regulated genes | CD45+ |
| BP | GO:0051224 | negative regulation of protein transport | 1.21E-03 | 9.86E-03 | Rsad2/Il1b/Arg2/C | 7 | aPD-L1+aTGF-β vs PBS | Up-regulated genes | CD45+ |
| BP | GO:0006835 | dicarboxylic acid transport | 1.30E-03 | 1.04E-02 | Il1b/Tnf/Slc7a11/Il | 4 | aPD-L1+aTGF-β vs PBS | Up-regulated genes | CD45+ |
| BP | GO:0060760 | positive regulation of response to cytokine stimu | 1.30E-03 | 1.04E-02 | Cd74/Irgm1/Zbp1 | 4 | aPD-L1+aTGF-β vs PBS | Up-regulated genes | CD45+ |
| BP | GO:1905517 | macrophage migration | 1.30E-03 | 1.04E-02 | Ccl5/Ccl3/Ccl2/Sl | 4 | aPD-L1+aTGF-β vs PBS | Up-regulated genes | CD45+ |
| BP | GO:0002675 | positive regulation of acute inflammatory respon | 1.31E-03 | 1.04E-02 | Il1b/Tnf/Ccl5 | 3 | aPD-L1+aTGF-β vs PBS | Up-regulated genes | CD45+ |
| BP | GO:0035743 | CD4-positive, alpha-beta T cell cytokine product | 1.31E-03 | 1.04E-02 | Rsad2/Il1b/Nlrp3 | 3 | aPD-L1+aTGF-β vs PBS | Up-regulated genes | CD45+ |
| BP | GO:0051767 | nitric-oxide synthase biosynthetic process | 1.31E-03 | 1.04E-02 | Nampt/Stat1/Ccl2 | 3 | aPD-L1+aTGF-β vs PBS | Up-regulated genes | CD45+ |
| BP | GO:0051769 | regulation of nitric-oxide synthase biosynthetic p | 1.31E-03 | 1.04E-02 | Nampt/Stat1/Ccl2 | 3 | aPD-L1+aTGF-β vs PBS | Up-regulated genes | CD45+ |
| BP | GO:0002792 | negative regulation of peptide secretion | 1.35E-03 | 1.07E-02 | Cd74/Rsad2/Il1b/ | 6 | aPD-L1+aTGF-β vs PBS | Up-regulated genes | CD45+ |
| BP | GO:0050920 | regulation of chemotaxis | 1.44E-03 | 1.13E-02 | Cd74/Cxcl10/Il1b/ | 7 | aPD-L1+aTGF-β vs PBS | Up-regulated genes | CD45+ |
| BP | GO:0002483 | antigen processing and presentation of endoger | 1.48E-03 | 1.16E-02 | H2-K1/H2-D1/B2r | 3 | aPD-L1+aTGF-β vs PBS | Up-regulated genes | CD45+ |
| BP | GO:0019885 | antigen processing and presentation of endoger | 1.48E-03 | 1.16E-02 | H2-K1/H2-D1/B2r | 3 | aPD-L1+aTGF-β vs PBS | Up-regulated genes | CD45+ |
| BP | GO:1903792 | negative regulation of anion transport | 1.48E-03 | 1.16E-02 | Il1b/Tnf/Il1rn | 3 | aPD-L1+aTGF-β vs PBS | Up-regulated genes | CD45+ |
| BP | GO:0032890 | regulation of organic acid transport | 1.50E-03 | 1.17E-02 | Il1b/Arg2/Tnf/Il1rn | 4 | aPD-L1+aTGF-β vs PBS | Up-regulated genes | CD45+ |
| BP | GO:0002221 | pattern recognition receptor signaling pathway | 1.53E-03 | 1.18E-02 | Irf1/Rsad2/Tnf/Nfk | 6 | aPD-L1+aTGF-β vs PBS | Up-regulated genes | CD45+ |
| BP | GO:0002700 | regulation of production of molecular mediator o | 1.53E-03 | 1.18E-02 | Cd74/Rsad2/Il1b/ | 6 | aPD-L1+aTGF-β vs PBS | Up-regulated genes | CD45+ |
| BP | GO:0032612 | interleukin-1 production | 1.60E-03 | 1.23E-02 | Il1b/Arg2/Nlrp3/Gl | 5 | aPD-L1+aTGF-β vs PBS | Up-regulated genes | CD45+ |
| BP | GO:2000106 | regulation of leukocyte apoptotic process | 1.60E-03 | 1.23E-02 | Cd74/Cd274/Arg2 | 5 | aPD-L1+aTGF-β vs PBS | Up-regulated genes | CD45+ |
| BP | GO:0002758 | innate immune response-activating signal transc | 1.65E-03 | 1.27E-02 | Irf1/Rsad2/Tnf/Nfk | 6 | aPD-L1+aTGF-β vs PBS | Up-regulated genes | CD45+ |
| BP | GO:0032480 | negative regulation of type I interferon productior | 1.67E-03 | 1.28E-02 | Gbp4/Rel/Acod1 | 3 | aPD-L1+aTGF-β vs PBS | Up-regulated genes | CD45+ |
| BP | GO:0036037 | CD8-positive, alpha-beta T cell activation | 1.67E-03 | 1.28E-02 | Irf1/Cd274/Socs1 | 3 | aPD-L1+aTGF-β vs PBS | Up-regulated genes | CD45+ |
| BP | GO:0072678 | T cell migration | 1.73E-03 | 1.32E-02 | Cxcl10/Icam1/Ccl | 4 | aPD-L1+aTGF-β vs PBS | Up-regulated genes | CD45+ |
| BP | GO:0070372 | regulation of ERK1 and ERK2 cascade | 1.76E-03 | 1.33E-02 | Cd74/Il1b/Ccl4/Ica | 8 | aPD-L1+aTGF-β vs PBS | Up-regulated genes | CD45+ |
| BP | GO:0002702 | positive regulation of production of molecular me | 1.77E-03 | 1.33E-02 | Cd74/Rsad2/Il1b/ | 5 | aPD-L1+aTGF-β vs PBS | Up-regulated genes | CD45+ |
| BP | GO:0001913 | T cell mediated cytotoxicity | 1.85E-03 | 1.38E-02 | H2-K1/H2-D1/Hsp | 4 | aPD-L1+aTGF-β vs PBS | Up-regulated genes | CD45+ |
| BP | GO:1902106 | negative regulation of leukocyte differentiation | 1.85E-03 | 1.38E-02 | Irf1/Cd74/Socs1/C | 5 | aPD-L1+aTGF-β vs PBS | Up-regulated genes | CD45+ |
| BP | GO:2001243 | negative regulation of intrinsic apoptotic signalin | 1.85E-03 | 1.38E-02 | Cd74/Dnaja1/Nfe | 5 | aPD-L1+aTGF-β vs PBS | Up-regulated genes | CD45+ |
| BP | GO:0072593 | reactive oxygen species metabolic process | 1.86E-03 | 1.38E-02 | Il1b/Arg2/Icam1/T | 8 | aPD-L1+aTGF-β vs PBS | Up-regulated genes | CD45+ |
| BP | GO:0019883 | antigen processing and presentation of endoger | 1.88E-03 | 1.40E-02 | H2-K1/H2-D1/B2r | 3 | aPD-L1+aTGF-β vs PBS | Up-regulated genes | CD45+ |
| BP | GO:0050707 | regulation of cytokine secretion | 1.97E-03 | 1.46E-02 | Cd274/Il1b/Arg2/S | 7 | aPD-L1+aTGF-β vs PBS | Up-regulated genes | CD45+ |
| BP | GO:0002791 | regulation of peptide secretion | 1.99E-03 | 1.47E-02 | Cd74/Rsad2/Cd2 | 11 | aPD-L1+aTGF-β vs PBS | Up-regulated genes | CD45+ |
| BP | GO:0001659 | temperature homeostasis | 1.99E-03 | 1.47E-02 | Il1b/Tnf/Ccl5/Kdm | 6 | aPD-L1+aTGF-β vs PBS | Up-regulated genes | CD45+ |
| BP | GO:0030099 | myeloid cell differentiation | 1.99E-03 | 1.47E-02 | Cd74/Tnf/Nfkbia/C | 10 | aPD-L1+aTGF-β vs PBS | Up-regulated genes | CD45+ |
| BP | GO:0009306 | protein secretion | 2.09E-03 | 1.53E-02 | Rsad2/Cd274/Il1b | 12 | aPD-L1+aTGF-β vs PBS | Up-regulated genes | CD45+ |
| BP | GO:0032722 | positive regulation of chemokine production | 2.11E-03 | 1.54E-02 | Cd74/Il1b/Tnf/Ccl | 4 | aPD-L1+aTGF-β vs PBS | Up-regulated genes | CD45+ |
| BP | GO:0045123 | cellular extravasation | 2.11E-03 | 1.54E-02 | Icam1/Tnf/Ccl5/Cc | 4 | aPD-L1+aTGF-β vs PBS | Up-regulated genes | CD45+ |
| BP | GO:0046677 | response to antibiotic | 2.16E-03 | 1.57E-02 | Crip1/Nfe2l2/Txn' | 7 | aPD-L1+aTGF-β vs PBS | Up-regulated genes | CD45+ |
| BP | GO:0042035 | regulation of cytokine biosynthetic process | 2.24E-03 | 1.62E-02 | Irf1/Il1b/Tnf/Rel/N | 5 | aPD-L1+aTGF-β vs PBS | Up-regulated genes | CD45+ |
| BP | GO:0051048 | negative regulation of secretion | 2.29E-03 | 1.66E-02 | Cd74/Rsad2/Il1b/ | 7 | aPD-L1+aTGF-β vs PBS | Up-regulated genes | CD45+ |
| BP | GO:0010758 | regulation of macrophage chemotaxis | 2.34E-03 | 1.66E-02 | Ccl5/Ccl2/Slamf8 | 3 | aPD-L1+aTGF-β vs PBS | Up-regulated genes | CD45+ |

| | | | | | | | | | |
|---|---|---|---|---|---|---|---|---|---|
| BP | GO:0031640 | killing of cells of other organism | 2.34E-03 | 1.66E-02 | Rpl30/Ccl2/Rps1 | 3 | aPD-L1+aTGF-β vs PBS | Up-regulated genes | CD45+ |
| BP | GO:0044364 | disruption of cells of other organism | 2.34E-03 | 1.66E-02 | Rpl30/Ccl2/Rps1 | 3 | aPD-L1+aTGF-β vs PBS | Up-regulated genes | CD45+ |
| BP | GO:0045589 | regulation of regulatory T cell differentiation | 2.34E-03 | 1.66E-02 | Irf1/Socs1/Ctla4 | 3 | aPD-L1+aTGF-β vs PBS | Up-regulated genes | CD45+ |
| BP | GO:0046885 | regulation of hormone biosynthetic process | 2.34E-03 | 1.66E-02 | Il1b/Tnf/Nfkb1 | 3 | aPD-L1+aTGF-β vs PBS | Up-regulated genes | CD45+ |
| BP | GO:2001024 | negative regulation of response to drug | 2.34E-03 | 1.66E-02 | Tnf/Nfe2l2/Txn1 | 3 | aPD-L1+aTGF-β vs PBS | Up-regulated genes | CD45+ |
| BP | GO:0003158 | endothelium development | 2.34E-03 | 1.66E-02 | Cxcl10/Il1b/Icam1 | 5 | aPD-L1+aTGF-β vs PBS | Up-regulated genes | CD45+ |
| BP | GO:0001933 | negative regulation of protein phosphorylation | 2.35E-03 | 1.66E-02 | Irf1/Dnaja1/Il1b/G | 10 | aPD-L1+aTGF-β vs PBS | Up-regulated genes | CD45+ |
| BP | GO:0002763 | positive regulation of myeloid leukocyte different | 2.40E-03 | 1.69E-02 | Cd74/Tnf/Ccl5/Cc | 4 | aPD-L1+aTGF-β vs PBS | Up-regulated genes | CD45+ |
| BP | GO:0043434 | response to peptide hormone | 2.47E-03 | 1.73E-02 | Irf1/Il1b/Socs1/Se | 8 | aPD-L1+aTGF-β vs PBS | Up-regulated genes | CD45+ |
| BP | GO:0070371 | ERK1 and ERK2 cascade | 2.47E-03 | 1.73E-02 | Cd74/Il1b/Ccl4/Ica | 8 | aPD-L1+aTGF-β vs PBS | Up-regulated genes | CD45+ |
| BP | GO:0071375 | cellular response to peptide hormone stimulus | 2.50E-03 | 1.75E-02 | Irf1/Il1b/Socs1/Nr | 7 | aPD-L1+aTGF-β vs PBS | Up-regulated genes | CD45+ |
| BP | GO:0034121 | regulation of toll-like receptor signaling pathway | 2.55E-03 | 1.77E-02 | Irf1/Rsad2/Acod1 | 4 | aPD-L1+aTGF-β vs PBS | Up-regulated genes | CD45+ |
| BP | GO:0051952 | regulation of amine transport | 2.55E-03 | 1.77E-02 | Il1b/Arg2/Tnf/Il1rn | 4 | aPD-L1+aTGF-β vs PBS | Up-regulated genes | CD45+ |
| BP | GO:0070228 | regulation of lymphocyte apoptotic process | 2.55E-03 | 1.77E-02 | Cd74/Cd274/Arg2 | 4 | aPD-L1+aTGF-β vs PBS | Up-regulated genes | CD45+ |
| BP | GO:0002825 | regulation of T-helper 1 type immune response | 2.59E-03 | 1.78E-02 | Irf1/H2-Ab1/Il1b | 3 | aPD-L1+aTGF-β vs PBS | Up-regulated genes | CD45+ |
| BP | GO:0060142 | regulation of syncytium formation by plasma mer | 2.59E-03 | 1.78E-02 | Cxcl10/Cxcl9/Stat | 3 | aPD-L1+aTGF-β vs PBS | Up-regulated genes | CD45+ |
| BP | GO:1990748 | cellular detoxification | 2.59E-03 | 1.78E-02 | Tnf/Nfe2l2/Sod2 | 3 | aPD-L1+aTGF-β vs PBS | Up-regulated genes | CD45+ |
| BP | GO:0032649 | regulation of interferon-gamma production | 2.67E-03 | 1.82E-02 | Cd274/Il1b/Tnf/Pc | 5 | aPD-L1+aTGF-β vs PBS | Up-regulated genes | CD45+ |
| BP | GO:0043123 | positive regulation of I-kappaB kinase/NF-kappa | 2.67E-03 | 1.82E-02 | Cd74/Il1b/Tnf/Peli | 5 | aPD-L1+aTGF-β vs PBS | Up-regulated genes | CD45+ |
| BP | GO:0043405 | regulation of MAP kinase activity | 2.72E-03 | 1.85E-02 | Cd74/Dnaja1/Il1b | 8 | aPD-L1+aTGF-β vs PBS | Up-regulated genes | CD45+ |
| BP | GO:0006865 | amino acid transport | 2.79E-03 | 1.89E-02 | Il1b/Arg2/Tnf/Slc7 | 5 | aPD-L1+aTGF-β vs PBS | Up-regulated genes | CD45+ |
| BP | GO:0031341 | regulation of cell killing | 2.79E-03 | 1.89E-02 | H2-K1/H2-D1/Hsp | 5 | aPD-L1+aTGF-β vs PBS | Up-regulated genes | CD45+ |
| BP | GO:0002920 | regulation of humoral immune response | 2.86E-03 | 1.92E-02 | Il1b/Tnf/Acod1 | 3 | aPD-L1+aTGF-β vs PBS | Up-regulated genes | CD45+ |
| BP | GO:0045672 | positive regulation of osteoclast differentiation | 2.86E-03 | 1.92E-02 | Tnf/Ccl5/Ccl3 | 3 | aPD-L1+aTGF-β vs PBS | Up-regulated genes | CD45+ |
| BP | GO:0046427 | positive regulation of JAK-STAT cascade | 2.88E-03 | 1.92E-02 | Socs1/Tnf/Ccl5/Pa | 4 | aPD-L1+aTGF-β vs PBS | Up-regulated genes | CD45+ |
| BP | GO:0046637 | regulation of alpha-beta T cell differentiation | 2.88E-03 | 1.92E-02 | Irf1/Socs1/Nlrp3/C | 4 | aPD-L1+aTGF-β vs PBS | Up-regulated genes | CD45+ |
| BP | GO:0050766 | positive regulation of phagocytosis | 2.88E-03 | 1.92E-02 | Il1b/Tnf/Hspa8/Cc | 4 | aPD-L1+aTGF-β vs PBS | Up-regulated genes | CD45+ |
| BP | GO:0051262 | protein tetramerization | 3.03E-03 | 2.02E-02 | Cd74/Samhd1/Cc | 6 | aPD-L1+aTGF-β vs PBS | Up-regulated genes | CD45+ |
| BP | GO:0015837 | amine transport | 3.05E-03 | 2.02E-02 | Il1b/Arg2/Tnf/Il1rn | 4 | aPD-L1+aTGF-β vs PBS | Up-regulated genes | CD45+ |
| BP | GO:0042108 | positive regulation of cytokine biosynthetic proce | 3.05E-03 | 2.02E-02 | Irf1/Il1b/Tnf/Rel | 4 | aPD-L1+aTGF-β vs PBS | Up-regulated genes | CD45+ |
| BP | GO:1903201 | regulation of oxidative stress-induced cell death | 3.05E-03 | 2.02E-02 | Nfe2l2/Txn1/Slc7a | 4 | aPD-L1+aTGF-β vs PBS | Up-regulated genes | CD45+ |
| BP | GO:0032743 | positive regulation of interleukin-2 production | 3.14E-03 | 2.05E-02 | Il1b/Pde4b/Cd83 | 3 | aPD-L1+aTGF-β vs PBS | Up-regulated genes | CD45+ |
| BP | GO:0045066 | regulatory T cell differentiation | 3.14E-03 | 2.05E-02 | Irf1/Socs1/Ctla4 | 3 | aPD-L1+aTGF-β vs PBS | Up-regulated genes | CD45+ |
| BP | GO:0051085 | chaperone cofactor-dependent protein refolding | 3.14E-03 | 2.05E-02 | Cd74/Hspa8/H2-I | 3 | aPD-L1+aTGF-β vs PBS | Up-regulated genes | CD45+ |
| BP | GO:2000515 | negative regulation of CD4-positive, alpha-beta | 3.14E-03 | 2.05E-02 | Irf1/Cd274/Arg2 | 3 | aPD-L1+aTGF-β vs PBS | Up-regulated genes | CD45+ |
| BP | GO:0032102 | negative regulation of response to external stimu | 3.30E-03 | 2.15E-02 | Arg2/Nlrp3/Acod1 | 8 | aPD-L1+aTGF-β vs PBS | Up-regulated genes | CD45+ |
| BP | GO:0006919 | activation of cysteine-type endopeptidase activit | 3.42E-03 | 2.22E-02 | Tnf/Nlrp3/S100a8 | 4 | aPD-L1+aTGF-β vs PBS | Up-regulated genes | CD45+ |
| BP | GO:0034142 | toll-like receptor 4 signaling pathway | 3.44E-03 | 2.23E-02 | Nfkbia/Acod1/Peli | 3 | aPD-L1+aTGF-β vs PBS | Up-regulated genes | CD45+ |
| BP | GO:0046328 | regulation of JNK cascade | 3.46E-03 | 2.24E-02 | Dnaja1/Il1b/Tnf/S | 6 | aPD-L1+aTGF-β vs PBS | Up-regulated genes | CD45+ |
| BP | GO:0042089 | cytokine biosynthetic process | 3.58E-03 | 2.31E-02 | Irf1/Il1b/Tnf/Rel/N | 5 | aPD-L1+aTGF-β vs PBS | Up-regulated genes | CD45+ |
| BP | GO:0043507 | positive regulation of JUN kinase activity | 3.61E-03 | 2.32E-02 | Il1b/Tnf/Spag9/Il1 | 4 | aPD-L1+aTGF-β vs PBS | Up-regulated genes | CD45+ |
| BP | GO:1904894 | positive regulation of STAT cascade | 3.61E-03 | 2.32E-02 | Socs1/Tnf/Ccl5/Pa | 4 | aPD-L1+aTGF-β vs PBS | Up-regulated genes | CD45+ |
| BP | GO:0002474 | antigen processing and presentation of peptide | 3.76E-03 | 2.39E-02 | H2-K1/H2-D1/B2r | 3 | aPD-L1+aTGF-β vs PBS | Up-regulated genes | CD45+ |
| BP | GO:0042832 | defense response to protozoan | 3.76E-03 | 2.39E-02 | Gbp2b/Gbp2/Gbp | 3 | aPD-L1+aTGF-β vs PBS | Up-regulated genes | CD45+ |
| BP | GO:1902930 | regulation of alcohol biosynthetic process | 3.76E-03 | 2.39E-02 | Il1b/Tnf/Nfkb1 | 3 | aPD-L1+aTGF-β vs PBS | Up-regulated genes | CD45+ |
| BP | GO:0002218 | activation of innate immune response | 3.80E-03 | 2.42E-02 | Irf1/Rsad2/Tnf/Nfk | 6 | aPD-L1+aTGF-β vs PBS | Up-regulated genes | CD45+ |
| BP | GO:0002468 | dendritic cell antigen processing and presentatio | 3.87E-03 | 2.42E-02 | Cd74/Fgl2 | 2 | aPD-L1+aTGF-β vs PBS | Up-regulated genes | CD45+ |
| BP | GO:0002579 | positive regulation of antigen processing and pr | 3.87E-03 | 2.42E-02 | Cd74/H2-Ab1 | 2 | aPD-L1+aTGF-β vs PBS | Up-regulated genes | CD45+ |
| BP | GO:0002679 | respiratory burst involved in defense response | 3.87E-03 | 2.42E-02 | Slamf8/Rps19 | 2 | aPD-L1+aTGF-β vs PBS | Up-regulated genes | CD45+ |
| BP | GO:0032769 | negative regulation of monooxygenase activity | 3.87E-03 | 2.42E-02 | Arg2/Nfkb1 | 2 | aPD-L1+aTGF-β vs PBS | Up-regulated genes | CD45+ |
| BP | GO:0045080 | positive regulation of chemokine biosynthetic pr | 3.87E-03 | 2.42E-02 | Il1b/Tnf | 2 | aPD-L1+aTGF-β vs PBS | Up-regulated genes | CD45+ |
| BP | GO:2000551 | regulation of T-helper 2 cell cytokine production | 3.87E-03 | 2.42E-02 | Rsad2/Nlrp3 | 2 | aPD-L1+aTGF-β vs PBS | Up-regulated genes | CD45+ |
| BP | GO:0071887 | leukocyte apoptotic process | 3.88E-03 | 2.42E-02 | Cd74/Cd274/Arg2 | 5 | aPD-L1+aTGF-β vs PBS | Up-regulated genes | CD45+ |
| BP | GO:0097193 | intrinsic apoptotic signaling pathway | 3.97E-03 | 2.47E-02 | Cd74/Dnaja1/Crip | 8 | aPD-L1+aTGF-β vs PBS | Up-regulated genes | CD45+ |
| BP | GO:0001909 | leukocyte mediated cytotoxicity | 4.03E-03 | 2.50E-02 | H2-K1/H2-D1/Hsp | 5 | aPD-L1+aTGF-β vs PBS | Up-regulated genes | CD45+ |
| BP | GO:0046627 | negative regulation of insulin receptor signaling | 4.09E-03 | 2.53E-02 | Il1b/Socs1/Socs3 | 3 | aPD-L1+aTGF-β vs PBS | Up-regulated genes | CD45+ |
| BP | GO:2000403 | positive regulation of lymphocyte migration | 4.09E-03 | 2.53E-02 | Cxcl10/Ccl4/Ccl5 | 3 | aPD-L1+aTGF-β vs PBS | Up-regulated genes | CD45+ |
| BP | GO:0030218 | erythrocyte differentiation | 4.19E-03 | 2.57E-02 | B2m/Stat1/Klf2/Isg | 5 | aPD-L1+aTGF-β vs PBS | Up-regulated genes | CD45+ |
| BP | GO:0042107 | cytokine metabolic process | 4.19E-03 | 2.57E-02 | Irf1/Il1b/Tnf/Rel/N | 5 | aPD-L1+aTGF-β vs PBS | Up-regulated genes | CD45+ |
| BP | GO:2001242 | regulation of intrinsic apoptotic signaling pathwa | 4.31E-03 | 2.64E-02 | Cd74/Dnaja1/Nfe | 6 | aPD-L1+aTGF-β vs PBS | Up-regulated genes | CD45+ |
| BP | GO:0032609 | interferon-gamma production | 4.35E-03 | 2.66E-02 | Cd274/Il1b/Tnf/Pc | 5 | aPD-L1+aTGF-β vs PBS | Up-regulated genes | CD45+ |
| BP | GO:0001562 | response to protozoan | 4.45E-03 | 2.70E-02 | Gbp2b/Gbp2/Gbp | 3 | aPD-L1+aTGF-β vs PBS | Up-regulated genes | CD45+ |
| BP | GO:0050706 | regulation of interleukin-1 beta secretion | 4.45E-03 | 2.70E-02 | Arg2/Nlrp3/Ccl3 | 3 | aPD-L1+aTGF-β vs PBS | Up-regulated genes | CD45+ |
| BP | GO:0070229 | negative regulation of lymphocyte apoptotic proc | 4.45E-03 | 2.70E-02 | Cd74/Arg2/Ccl5 | 3 | aPD-L1+aTGF-β vs PBS | Up-regulated genes | CD45+ |
| BP | GO:0042133 | neurotransmitter metabolic process | 4.52E-03 | 2.74E-02 | Il1b/Icam1/Tnf/So | 5 | aPD-L1+aTGF-β vs PBS | Up-regulated genes | CD45+ |
| BP | GO:0007254 | JNK cascade | 4.57E-03 | 2.76E-02 | Dnaja1/Il1b/Tnf/S | 6 | aPD-L1+aTGF-β vs PBS | Up-regulated genes | CD45+ |
| BP | GO:0031397 | negative regulation of protein ubiquitination | 4.70E-03 | 2.77E-02 | Dnaja1/Gbp4/Rpl | 4 | aPD-L1+aTGF-β vs PBS | Up-regulated genes | CD45+ |
| BP | GO:0032088 | negative regulation of NF-kappaB transcription f | 4.70E-03 | 2.77E-02 | Nfkbia/Nlrp3/Acod | 4 | aPD-L1+aTGF-β vs PBS | Up-regulated genes | CD45+ |
| BP | GO:0006750 | glutathione biosynthetic process | 4.70E-03 | 2.77E-02 | Nfe2l2/Slc7a11 | 2 | aPD-L1+aTGF-β vs PBS | Up-regulated genes | CD45+ |
| BP | GO:0034116 | positive regulation of heterotypic cell-cell adhesi | 4.70E-03 | 2.77E-02 | Il1b/Tnf | 2 | aPD-L1+aTGF-β vs PBS | Up-regulated genes | CD45+ |
| BP | GO:0035739 | CD4-positive, alpha-beta T cell proliferation | 4.70E-03 | 2.77E-02 | Cd274/Arg2 | 2 | aPD-L1+aTGF-β vs PBS | Up-regulated genes | CD45+ |
| BP | GO:0060263 | regulation of respiratory burst | 4.70E-03 | 2.77E-02 | Slamf8/Rps19 | 2 | aPD-L1+aTGF-β vs PBS | Up-regulated genes | CD45+ |
| BP | GO:0072608 | interleukin-10 secretion | 4.70E-03 | 2.77E-02 | Cd274/Isg15 | 2 | aPD-L1+aTGF-β vs PBS | Up-regulated genes | CD45+ |
| BP | GO:1901550 | regulation of endothelial cell development | 4.70E-03 | 2.77E-02 | Il1b/Tnf | 2 | aPD-L1+aTGF-β vs PBS | Up-regulated genes | CD45+ |
| BP | GO:1903140 | regulation of establishment of endothelial barrie | 4.70E-03 | 2.77E-02 | Il1b/Tnf | 2 | aPD-L1+aTGF-β vs PBS | Up-regulated genes | CD45+ |
| BP | GO:2000561 | regulation of CD4-positive, alpha-beta T cell pro | 4.70E-03 | 2.77E-02 | Cd274/Arg2 | 2 | aPD-L1+aTGF-β vs PBS | Up-regulated genes | CD45+ |
| BP | GO:0042326 | negative regulation of phosphorylation | 4.71E-03 | 2.77E-02 | Irf1/Dnaja1/Il1b/G | 10 | aPD-L1+aTGF-β vs PBS | Up-regulated genes | CD45+ |
| BP | GO:1901342 | regulation of vasculature development | 4.74E-03 | 2.78E-02 | Cxcl10/Il1b/Nfe2l | 8 | aPD-L1+aTGF-β vs PBS | Up-regulated genes | CD45+ |
| BP | GO:0051084 | 'de novo' posttranslational protein folding | 4.82E-03 | 2.81E-02 | Cd74/Hspa8/H2-I | 3 | aPD-L1+aTGF-β vs PBS | Up-regulated genes | CD45+ |
| BP | GO:1901031 | regulation of response to reactive oxygen specie | 4.82E-03 | 2.81E-02 | Tnf/Nfe2l2/Txn1 | 3 | aPD-L1+aTGF-β vs PBS | Up-regulated genes | CD45+ |
| BP | GO:0030335 | positive regulation of cell migration | 4.99E-03 | 2.91E-02 | Cd74/Cxcl10/Cd2 | 11 | aPD-L1+aTGF-β vs PBS | Up-regulated genes | CD45+ |
| BP | GO:0006458 | 'de novo' protein folding | 5.21E-03 | 3.00E-02 | Cd74/Hspa8/H2-I | 3 | aPD-L1+aTGF-β vs PBS | Up-regulated genes | CD45+ |
| BP | GO:0032350 | regulation of hormone metabolic process | 5.21E-03 | 3.00E-02 | Il1b/Tnf/Nfkb1 | 3 | aPD-L1+aTGF-β vs PBS | Up-regulated genes | CD45+ |
| BP | GO:0046640 | regulation of alpha-beta T cell proliferation | 5.21E-03 | 3.00E-02 | Irf1/Cd274/Arg2 | 3 | aPD-L1+aTGF-β vs PBS | Up-regulated genes | CD45+ |
| BP | GO:0098754 | detoxification | 5.21E-03 | 3.00E-02 | Tnf/Nfe2l2/Sod2 | 3 | aPD-L1+aTGF-β vs PBS | Up-regulated genes | CD45+ |
| BP | GO:0002253 | activation of immune response | 5.38E-03 | 3.09E-02 | Irf1/Rsad2/Il1b/Ct | 9 | aPD-L1+aTGF-β vs PBS | Up-regulated genes | CD45+ |
| BP | GO:0010563 | negative regulation of phosphorus metabolic pro | 5.40E-03 | 3.09E-02 | Irf1/Dnaja1/Il1b/G | 11 | aPD-L1+aTGF-β vs PBS | Up-regulated genes | CD45+ |
| BP | GO:0045936 | negative regulation of phosphate metabolic proc | 5.40E-03 | 3.09E-02 | Irf1/Dnaja1/Il1b/G | 11 | aPD-L1+aTGF-β vs PBS | Up-regulated genes | CD45+ |
| BP | GO:0014048 | regulation of glutamate secretion | 5.60E-03 | 3.15E-02 | Il1b/Il1rn | 2 | aPD-L1+aTGF-β vs PBS | Up-regulated genes | CD45+ |

| | | | | | | | | | |
|---|---|---|---|---|---|---|---|---|---|
| BP | GO:0031269 | pseudopodium assembly | 5.60E-03 | 3.15E-02 | Ccl5/Cdc42ep2 | 2 | aPD-L1+aTGF-β vs PBS | Up-regulated genes | CD45+ |
| BP | GO:0042033 | chemokine biosynthetic process | 5.60E-03 | 3.15E-02 | Il1b/Tnf | 2 | aPD-L1+aTGF-β vs PBS | Up-regulated genes | CD45+ |
| BP | GO:0045073 | regulation of chemokine biosynthetic process | 5.60E-03 | 3.15E-02 | Il1b/Tnf | 2 | aPD-L1+aTGF-β vs PBS | Up-regulated genes | CD45+ |
| BP | GO:0098712 | L-glutamate import across plasma membrane | 5.60E-03 | 3.15E-02 | Tnf/Slc7a11 | 2 | aPD-L1+aTGF-β vs PBS | Up-regulated genes | CD45+ |
| BP | GO:0002832 | negative regulation of response to biotic stimulus | 5.61E-03 | 3.15E-02 | Arg2/Acod1/Fgl2 | 3 | aPD-L1+aTGF-β vs PBS | Up-regulated genes | CD45+ |
| BP | GO:1900077 | negative regulation of cellular response to insuli | 5.61E-03 | 3.15E-02 | Il1b/Socs1/Socs3 | 3 | aPD-L1+aTGF-β vs PBS | Up-regulated genes | CD45+ |
| BP | GO:1905521 | regulation of macrophage migration | 5.61E-03 | 3.15E-02 | Ccl5/Ccl2/Slamf8 | 3 | aPD-L1+aTGF-β vs PBS | Up-regulated genes | CD45+ |
| BP | GO:0046330 | positive regulation of JNK cascade | 5.62E-03 | 3.15E-02 | Il1b/Tnf/Spag9/Ga | 5 | aPD-L1+aTGF-β vs PBS | Up-regulated genes | CD45+ |
| BP | GO:0002757 | immune response-activating signal transduction | 5.62E-03 | 3.15E-02 | Irf1/Rsad2/Ctla4/I | 8 | aPD-L1+aTGF-β vs PBS | Up-regulated genes | CD45+ |
| BP | GO:0043281 | regulation of cysteine-type endopeptidase activit | 5.77E-03 | 3.22E-02 | Tnf/Nlrp3/Plaur/S | 6 | aPD-L1+aTGF-β vs PBS | Up-regulated genes | CD45+ |
| BP | GO:0050671 | positive regulation of lymphocyte proliferation | 5.82E-03 | 3.25E-02 | Cd74/Cd274/Il1b/ | 5 | aPD-L1+aTGF-β vs PBS | Up-regulated genes | CD45+ |
| BP | GO:2000147 | positive regulation of cell motility | 5.83E-03 | 3.25E-02 | Cd74/Cxcl10/Cd2 | 11 | aPD-L1+aTGF-β vs PBS | Up-regulated genes | CD45+ |
| BP | GO:2001023 | regulation of response to drug | 5.98E-03 | 3.31E-02 | Arg2/Tnf/Nfe2l2/T | 4 | aPD-L1+aTGF-β vs PBS | Up-regulated genes | CD45+ |
| BP | GO:0002698 | negative regulation of immune effector process | 6.03E-03 | 3.31E-02 | Irf1/Tnf/Fgl2/Slam | 5 | aPD-L1+aTGF-β vs PBS | Up-regulated genes | CD45+ |
| BP | GO:0001960 | negative regulation of cytokine-mediated signali | 6.04E-03 | 3.31E-02 | Samhd1/Parp14/I | 3 | aPD-L1+aTGF-β vs PBS | Up-regulated genes | CD45+ |
| BP | GO:0046782 | regulation of viral transcription | 6.04E-03 | 3.31E-02 | Ccl4/Ccl5/Ifitm3 | 3 | aPD-L1+aTGF-β vs PBS | Up-regulated genes | CD45+ |
| BP | GO:0061028 | establishment of endothelial barrier | 6.04E-03 | 3.31E-02 | Il1b/Icam1/Tnf | 3 | aPD-L1+aTGF-β vs PBS | Up-regulated genes | CD45+ |
| BP | GO:0070232 | regulation of T cell apoptotic process | 6.04E-03 | 3.31E-02 | Cd274/Arg2/Ccl5 | 3 | aPD-L1+aTGF-β vs PBS | Up-regulated genes | CD45+ |
| BP | GO:1902622 | regulation of neutrophil migration | 6.04E-03 | 3.31E-02 | Cd74/Il1b/Slamf8 | 3 | aPD-L1+aTGF-β vs PBS | Up-regulated genes | CD45+ |
| BP | GO:1903531 | negative regulation of secretion by cell | 6.10E-03 | 3.33E-02 | Rsad2/Il1b/Arg2/T | 6 | aPD-L1+aTGF-β vs PBS | Up-regulated genes | CD45+ |
| BP | GO:0010035 | response to inorganic substance | 6.13E-03 | 3.34E-02 | Selenom/Crip1/Tr | 9 | aPD-L1+aTGF-β vs PBS | Up-regulated genes | CD45+ |
| BP | GO:0032946 | positive regulation of mononuclear cell proliferat | 6.24E-03 | 3.39E-02 | Cd74/Cd274/Il1b/ | 5 | aPD-L1+aTGF-β vs PBS | Up-regulated genes | CD45+ |
| BP | GO:0050709 | negative regulation of protein secretion | 6.24E-03 | 3.39E-02 | Rsad2/Il1b/Arg2/T | 5 | aPD-L1+aTGF-β vs PBS | Up-regulated genes | CD45+ |
| BP | GO:0050864 | regulation of B cell activation | 6.45E-03 | 3.49E-02 | Cd74/Ctla4/Sams | 5 | aPD-L1+aTGF-β vs PBS | Up-regulated genes | CD45+ |
| BP | GO:0046633 | alpha-beta T cell proliferation | 6.48E-03 | 3.49E-02 | Irf1/Cd274/Arg2 | 3 | aPD-L1+aTGF-β vs PBS | Up-regulated genes | CD45+ |
| BP | GO:0050691 | regulation of defense response to virus by host | 6.48E-03 | 3.49E-02 | Il1b/Ccl5/Stat1 | 3 | aPD-L1+aTGF-β vs PBS | Up-regulated genes | CD45+ |
| BP | GO:0050704 | regulation of interleukin-1 secretion | 6.48E-03 | 3.49E-02 | Arg2/Nlrp3/Ccl3 | 3 | aPD-L1+aTGF-β vs PBS | Up-regulated genes | CD45+ |
| BP | GO:0070227 | lymphocyte apoptotic process | 6.55E-03 | 3.50E-02 | Cd74/Cd274/Arg2 | 4 | aPD-L1+aTGF-β vs PBS | Up-regulated genes | CD45+ |
| BP | GO:0010819 | regulation of T cell chemotaxis | 6.58E-03 | 3.50E-02 | Cxcl10/Ccl5 | 2 | aPD-L1+aTGF-β vs PBS | Up-regulated genes | CD45+ |
| BP | GO:0019184 | nonribosomal peptide biosynthetic process | 6.58E-03 | 3.50E-02 | Nfe2l2/Slc7a11 | 2 | aPD-L1+aTGF-β vs PBS | Up-regulated genes | CD45+ |
| BP | GO:0030730 | sequestering of triglyceride | 6.58E-03 | 3.50E-02 | Il1b/Tnf | 2 | aPD-L1+aTGF-β vs PBS | Up-regulated genes | CD45+ |
| BP | GO:0045628 | regulation of T-helper 2 cell differentiation | 6.58E-03 | 3.50E-02 | Irf1/Nlrp3 | 2 | aPD-L1+aTGF-β vs PBS | Up-regulated genes | CD45+ |
| BP | GO:0046642 | negative regulation of alpha-beta T cell prolifera | 6.58E-03 | 3.50E-02 | Cd274/Arg2 | 2 | aPD-L1+aTGF-β vs PBS | Up-regulated genes | CD45+ |
| BP | GO:0051051 | negative regulation of transport | 6.60E-03 | 3.50E-02 | Cd74/Rsad2/Il1b/ | 10 | aPD-L1+aTGF-β vs PBS | Up-regulated genes | CD45+ |
| BP | GO:0051272 | positive regulation of cellular component movem | 7.09E-03 | 3.75E-02 | Cd74/Cxcl10/Cd2 | 11 | aPD-L1+aTGF-β vs PBS | Up-regulated genes | CD45+ |
| BP | GO:0036473 | cell death in response to oxidative stress | 7.16E-03 | 3.77E-02 | Nfe2l2/Txn1/Slc7 | 4 | aPD-L1+aTGF-β vs PBS | Up-regulated genes | CD45+ |
| BP | GO:1903321 | negative regulation of protein modification by sm | 7.16E-03 | 3.77E-02 | Dnaja1/Gbp4/Rpl | 4 | aPD-L1+aTGF-β vs PBS | Up-regulated genes | CD45+ |
| BP | GO:0002764 | immune response-regulating signaling pathway | 7.17E-03 | 3.77E-02 | Irf1/Rsad2/Ctla4/I | 8 | aPD-L1+aTGF-β vs PBS | Up-regulated genes | CD45+ |
| BP | GO:0070665 | positive regulation of leukocyte proliferation | 7.37E-03 | 3.86E-02 | Cd74/Cd274/Il1b/ | 5 | aPD-L1+aTGF-β vs PBS | Up-regulated genes | CD45+ |
| BP | GO:0019083 | viral transcription | 7.43E-03 | 3.87E-02 | Ccl4/Ccl5/Ifitm3 | 3 | aPD-L1+aTGF-β vs PBS | Up-regulated genes | CD45+ |
| BP | GO:0034113 | heterotypic cell-cell adhesion | 7.43E-03 | 3.87E-02 | Il1b/Tnf/Il1rn | 3 | aPD-L1+aTGF-β vs PBS | Up-regulated genes | CD45+ |
| BP | GO:0044070 | regulation of anion transport | 7.48E-03 | 3.87E-02 | Il1b/Arg2/Tnf/Il1rn | 4 | aPD-L1+aTGF-β vs PBS | Up-regulated genes | CD45+ |
| BP | GO:0009266 | response to temperature stimulus | 7.61E-03 | 3.87E-02 | Dnaja1/Hspa8/Pd | 5 | aPD-L1+aTGF-β vs PBS | Up-regulated genes | CD45+ |
| BP | GO:0002227 | innate immune response in mucosa | 7.63E-03 | 3.87E-02 | Rpl39/Fau | 2 | aPD-L1+aTGF-β vs PBS | Up-regulated genes | CD45+ |
| BP | GO:0002829 | negative regulation of type 2 immune response | 7.63E-03 | 3.87E-02 | Irf1/Arg2 | 2 | aPD-L1+aTGF-β vs PBS | Up-regulated genes | CD45+ |
| BP | GO:0030213 | hyaluronan biosynthetic process | 7.63E-03 | 3.87E-02 | Il1b/Nfkb1 | 2 | aPD-L1+aTGF-β vs PBS | Up-regulated genes | CD45+ |
| BP | GO:0031268 | pseudopodium organization | 7.63E-03 | 3.87E-02 | Ccl5/Cdc42ep2 | 2 | aPD-L1+aTGF-β vs PBS | Up-regulated genes | CD45+ |
| BP | GO:0035745 | T-helper 2 cell cytokine production | 7.63E-03 | 3.87E-02 | Rsad2/Nlrp3 | 2 | aPD-L1+aTGF-β vs PBS | Up-regulated genes | CD45+ |
| BP | GO:0043374 | CD8-positive, alpha-beta T cell differentiation | 7.63E-03 | 3.87E-02 | Irf1/Socs1 | 2 | aPD-L1+aTGF-β vs PBS | Up-regulated genes | CD45+ |
| BP | GO:0044546 | NLRP3 inflammasome complex assembly | 7.63E-03 | 3.87E-02 | Nlrp3/Gbp5 | 2 | aPD-L1+aTGF-β vs PBS | Up-regulated genes | CD45+ |
| BP | GO:0046886 | positive regulation of hormone biosynthetic proc | 7.63E-03 | 3.87E-02 | Il1b/Tnf | 2 | aPD-L1+aTGF-β vs PBS | Up-regulated genes | CD45+ |
| BP | GO:0050755 | chemokine metabolic process | 7.63E-03 | 3.87E-02 | Il1b/Tnf | 2 | aPD-L1+aTGF-β vs PBS | Up-regulated genes | CD45+ |
| BP | GO:0051938 | L-glutamate import | 7.63E-03 | 3.87E-02 | Tnf/Slc7a11 | 2 | aPD-L1+aTGF-β vs PBS | Up-regulated genes | CD45+ |
| BP | GO:0070234 | positive regulation of T cell apoptotic process | 7.63E-03 | 3.87E-02 | Cd274/Ccl5 | 2 | aPD-L1+aTGF-β vs PBS | Up-regulated genes | CD45+ |
| BP | GO:0070486 | leukocyte aggregation | 7.63E-03 | 3.87E-02 | Il1b/S100a8 | 2 | aPD-L1+aTGF-β vs PBS | Up-regulated genes | CD45+ |
| BP | GO:0060761 | negative regulation of response to cytokine stimu | 7.93E-03 | 4.02E-02 | Samhd1/Parp14/I | 3 | aPD-L1+aTGF-β vs PBS | Up-regulated genes | CD45+ |
| BP | GO:0043406 | positive regulation of MAP kinase activity | 7.97E-03 | 4.03E-02 | Cd74/Il1b/Tnf/Spa | 6 | aPD-L1+aTGF-β vs PBS | Up-regulated genes | CD45+ |
| BP | GO:0034614 | cellular response to reactive oxygen species | 8.11E-03 | 4.08E-02 | Tnf/Nfe2l2/Txn1/K | 5 | aPD-L1+aTGF-β vs PBS | Up-regulated genes | CD45+ |
| BP | GO:0050715 | positive regulation of cytokine secretion | 8.11E-03 | 4.08E-02 | Cd274/Il1b/Tnf/Nl | 5 | aPD-L1+aTGF-β vs PBS | Up-regulated genes | CD45+ |
| BP | GO:0015807 | L-amino acid transport | 8.45E-03 | 4.23E-02 | Arg2/Tnf/Slc7a11 | 3 | aPD-L1+aTGF-β vs PBS | Up-regulated genes | CD45+ |
| BP | GO:0061900 | glial cell activation | 8.45E-03 | 4.23E-02 | Il1b/Tnf/Nampt | 3 | aPD-L1+aTGF-β vs PBS | Up-regulated genes | CD45+ |
| BP | GO:2000351 | regulation of endothelial cell apoptotic process | 8.45E-03 | 4.23E-02 | Icam1/Tnf/Nfe2l2 | 3 | aPD-L1+aTGF-β vs PBS | Up-regulated genes | CD45+ |
| BP | GO:0051345 | positive regulation of hydrolase activity | 8.63E-03 | 4.31E-02 | Ccl4/Icam1/Tnf/Nl | 10 | aPD-L1+aTGF-β vs PBS | Up-regulated genes | CD45+ |
| BP | GO:0035094 | response to nicotine | 8.75E-03 | 4.31E-02 | Tnf/Nfkb1 | 2 | aPD-L1+aTGF-β vs PBS | Up-regulated genes | CD45+ |
| BP | GO:0045410 | positive regulation of interleukin-6 biosynthetic p | 8.75E-03 | 4.31E-02 | Il1b/Tnf | 2 | aPD-L1+aTGF-β vs PBS | Up-regulated genes | CD45+ |
| BP | GO:0051044 | positive regulation of membrane protein ectodor | 8.75E-03 | 4.31E-02 | Il1b/Tnf | 2 | aPD-L1+aTGF-β vs PBS | Up-regulated genes | CD45+ |
| BP | GO:0140131 | positive regulation of lymphocyte chemotaxis | 8.75E-03 | 4.31E-02 | Ccl4/Ccl5 | 2 | aPD-L1+aTGF-β vs PBS | Up-regulated genes | CD45+ |
| BP | GO:1902176 | negative regulation of oxidative stress-induced i | 8.75E-03 | 4.31E-02 | Nfe2l2/Sod2 | 2 | aPD-L1+aTGF-β vs PBS | Up-regulated genes | CD45+ |
| BP | GO:1904996 | positive regulation of leukocyte adhesion to vasc | 8.75E-03 | 4.31E-02 | Icam1/Tnf | 2 | aPD-L1+aTGF-β vs PBS | Up-regulated genes | CD45+ |
| BP | GO:0009408 | response to heat | 8.84E-03 | 4.35E-02 | Dnaja1/Hspa8/Pd | 4 | aPD-L1+aTGF-β vs PBS | Up-regulated genes | CD45+ |
| BP | GO:2001233 | regulation of apoptotic signaling pathway | 8.99E-03 | 4.42E-02 | Cd74/Dnaja1/Il1b | 9 | aPD-L1+aTGF-β vs PBS | Up-regulated genes | CD45+ |
| BP | GO:0052548 | regulation of endopeptidase activity | 9.28E-03 | 4.55E-02 | Tnf/Serpina3g/Nlr | 7 | aPD-L1+aTGF-β vs PBS | Up-regulated genes | CD45+ |
| BP | GO:0042088 | T-helper 1 type immune response | 9.56E-03 | 4.64E-02 | Irf1/H2-Ab1/Il1b | 3 | aPD-L1+aTGF-β vs PBS | Up-regulated genes | CD45+ |
| BP | GO:0042446 | hormone biosynthetic process | 9.56E-03 | 4.64E-02 | Il1b/Tnf/Nfkb1 | 3 | aPD-L1+aTGF-β vs PBS | Up-regulated genes | CD45+ |
| BP | GO:0045646 | regulation of erythrocyte differentiation | 9.56E-03 | 4.64E-02 | B2m/Stat1/Isg15 | 3 | aPD-L1+aTGF-β vs PBS | Up-regulated genes | CD45+ |
| BP | GO:0072577 | endothelial cell apoptotic process | 9.56E-03 | 4.64E-02 | Icam1/Tnf/Nfe2l2 | 3 | aPD-L1+aTGF-β vs PBS | Up-regulated genes | CD45+ |
| BP | GO:0002286 | T cell activation involved in immune response | 9.58E-03 | 4.64E-02 | Irf1/Icam1/Nlrp3/F | 4 | aPD-L1+aTGF-β vs PBS | Up-regulated genes | CD45+ |
| BP | GO:0050764 | regulation of phagocytosis | 9.58E-03 | 4.64E-02 | Il1b/Tnf/Hspa8/Cc | 4 | aPD-L1+aTGF-β vs PBS | Up-regulated genes | CD45+ |
| BP | GO:0045765 | regulation of angiogenesis | 9.67E-03 | 4.68E-02 | Cxcl10/Il1b/Nfe2l | 7 | aPD-L1+aTGF-β vs PBS | Up-regulated genes | CD45+ |
| BP | GO:2000116 | regulation of cysteine-type endopeptidase activit | 9.74E-03 | 4.70E-02 | Tnf/Nlrp3/Plaur/S | 6 | aPD-L1+aTGF-β vs PBS | Up-regulated genes | CD45+ |
| BP | GO:0015813 | L-glutamate transmembrane transport | 9.94E-03 | 4.75E-02 | Tnf/Slc7a11 | 2 | aPD-L1+aTGF-β vs PBS | Up-regulated genes | CD45+ |
| BP | GO:0044130 | negative regulation of growth of symbiont in host | 9.94E-03 | 4.75E-02 | Tnf/Sqstm1 | 2 | aPD-L1+aTGF-β vs PBS | Up-regulated genes | CD45+ |
| BP | GO:0071498 | cellular response to fluid shear stress | 9.94E-03 | 4.75E-02 | Nfe2l2/Klf2 | 2 | aPD-L1+aTGF-β vs PBS | Up-regulated genes | CD45+ |
| BP | GO:0000187 | activation of MAPK activity | 9.96E-03 | 4.75E-02 | Cd74/Il1b/Tnf/Spa | 4 | aPD-L1+aTGF-β vs PBS | Up-regulated genes | CD45+ |
| BP | GO:0019233 | sensory perception of pain | 9.96E-03 | 4.75E-02 | Tnf/Fcgr4/Ccl2/Il1 | 4 | aPD-L1+aTGF-β vs PBS | Up-regulated genes | CD45+ |
| BP | GO:0045446 | endothelial cell differentiation | 9.96E-03 | 4.75E-02 | Il1b/Icam1/Tnf/Kd | 4 | aPD-L1+aTGF-β vs PBS | Up-regulated genes | CD45+ |
| BP | GO:0032872 | regulation of stress-activated MAPK cascade | 9.98E-03 | 4.75E-02 | Dnaja1/Il1b/Tnf/S | 6 | aPD-L1+aTGF-β vs PBS | Up-regulated genes | CD45+ |
| BP | GO:0042531 | positive regulation of tyrosine phosphorylation o | 1.01E-02 | 4.82E-02 | Tnf/Ccl5/Parp14 | 3 | aPD-L1+aTGF-β vs PBS | Up-regulated genes | CD45+ |
| BP | GO:0003018 | vascular process in circulatory system | 1.03E-02 | 4.90E-02 | Ccl4/Icam1/Tnf/Du | 5 | aPD-L1+aTGF-β vs PBS | Up-regulated genes | CD45+ |

| | | | | | | | | | |
|---|---|---|---|---|---|---|---|---|---|
| BP | GO:0070302 | regulation of stress-activated protein kinase sign | 1.05E-02 | 4.95E-02 | Dnaja1/Il1b/Tnf/S | 6 | aPD-L1+aTGF-β vs PBS | Up-regulated genes | CD45+ |
| CC | GO:0022626 | cytosolic ribosome | 2.77E-27 | 5.51E-25 | Rpl39/Rps27/Rpl | 24 | aPD-L1+aTGF-β vs PBS | Up-regulated genes | CD45+ |
| CC | GO:0044445 | cytosolic part | 3.33E-22 | 3.31E-20 | Rpl39/Rps27/Rpl | 27 | aPD-L1+aTGF-β vs PBS | Up-regulated genes | CD45+ |
| CC | GO:0044391 | ribosomal subunit | 5.03E-21 | 3.34E-19 | Rpl39/Rps27/Rpl | 24 | aPD-L1+aTGF-β vs PBS | Up-regulated genes | CD45+ |
| CC | GO:0005840 | ribosome | 1.26E-20 | 6.26E-19 | Rpl39/Rps27/Rpl | 25 | aPD-L1+aTGF-β vs PBS | Up-regulated genes | CD45+ |
| CC | GO:0022625 | cytosolic large ribosomal subunit | 7.79E-20 | 3.10E-18 | Rpl39/Rpl37/Rpl | 16 | aPD-L1+aTGF-β vs PBS | Up-regulated genes | CD45+ |
| CC | GO:0015934 | large ribosomal subunit | 1.42E-14 | 4.72E-13 | Rpl39/Rpl37/Rpl | 16 | aPD-L1+aTGF-β vs PBS | Up-regulated genes | CD45+ |
| CC | GO:0042611 | MHC protein complex | 8.80E-12 | 2.50E-10 | H2-Eb1/H2-Aa/C | 8 | aPD-L1+aTGF-β vs PBS | Up-regulated genes | CD45+ |
| CC | GO:0022627 | cytosolic small ribosomal subunit | 1.45E-10 | 3.60E-09 | Rps27/Rps21/Rp | 9 | aPD-L1+aTGF-β vs PBS | Up-regulated genes | CD45+ |
| CC | GO:0015935 | small ribosomal subunit | 2.18E-08 | 4.83E-07 | Rps27/Rps21/Rp | 9 | aPD-L1+aTGF-β vs PBS | Up-regulated genes | CD45+ |
| CC | GO:0009897 | external side of plasma membrane | 3.65E-08 | 7.27E-07 | H2-Aa/Cd74/H2- | 16 | aPD-L1+aTGF-β vs PBS | Up-regulated genes | CD45+ |
| CC | GO:0042788 | polysomal ribosome | 5.54E-06 | 1.00E-04 | Rpl39/Rps21/Rpl | 5 | aPD-L1+aTGF-β vs PBS | Up-regulated genes | CD45+ |
| CC | GO:0098552 | side of membrane | 6.18E-06 | 1.02E-04 | H2-Aa/Cd74/H2- | 16 | aPD-L1+aTGF-β vs PBS | Up-regulated genes | CD45+ |
| CC | GO:0043230 | extracellular organelle | 1.18E-04 | 1.67E-03 | Cd274/Gbp2b/Gb | 6 | aPD-L1+aTGF-β vs PBS | Up-regulated genes | CD45+ |
| CC | GO:0020005 | symbiont-containing vacuole membrane | 1.26E-04 | 1.67E-03 | Gbp2b/Gbp2/Gbp | 3 | aPD-L1+aTGF-β vs PBS | Up-regulated genes | CD45+ |
| CC | GO:0042612 | MHC class I protein complex | 1.26E-04 | 1.67E-03 | H2-K1/H2-D1/B2 | 3 | aPD-L1+aTGF-β vs PBS | Up-regulated genes | CD45+ |
| CC | GO:0020003 | symbiont-containing vacuole | 2.15E-04 | 2.68E-03 | Gbp2b/Gbp2/Gbp | 3 | aPD-L1+aTGF-β vs PBS | Up-regulated genes | CD45+ |
| CC | GO:0042824 | MHC class I peptide loading complex | 2.72E-04 | 3.18E-03 | H2-K1/H2-D1/B2 | 3 | aPD-L1+aTGF-β vs PBS | Up-regulated genes | CD45+ |
| CC | GO:0065010 | extracellular membrane-bounded organelle | 3.38E-04 | 3.73E-03 | Gbp2b/Gbp2/Gbp | 3 | aPD-L1+aTGF-β vs PBS | Up-regulated genes | CD45+ |
| CC | GO:0030430 | host cell cytoplasm | 4.13E-04 | 4.11E-03 | Gbp2b/Gbp2/Gbp | 3 | aPD-L1+aTGF-β vs PBS | Up-regulated genes | CD45+ |
| CC | GO:0033655 | host cell cytoplasm part | 4.13E-04 | 4.11E-03 | Gbp2b/Gbp2/Gbp | 3 | aPD-L1+aTGF-β vs PBS | Up-regulated genes | CD45+ |
| CC | GO:0005844 | polysome | 4.36E-04 | 4.13E-03 | Rpl39/Rps21/Rpl | 5 | aPD-L1+aTGF-β vs PBS | Up-regulated genes | CD45+ |
| CC | GO:0033646 | host intracellular part | 7.00E-04 | 6.05E-03 | Gbp2b/Gbp2/Gbp | 3 | aPD-L1+aTGF-β vs PBS | Up-regulated genes | CD45+ |
| CC | GO:0043656 | intracellular region of host | 7.00E-04 | 6.05E-03 | Gbp2b/Gbp2/Gbp | 3 | aPD-L1+aTGF-β vs PBS | Up-regulated genes | CD45+ |
| CC | GO:0033643 | host cell part | 9.47E-04 | 7.85E-03 | Gbp2b/Gbp2/Gbp | 3 | aPD-L1+aTGF-β vs PBS | Up-regulated genes | CD45+ |
| CC | GO:0018995 | host | 1.24E-03 | 9.52E-03 | Gbp2b/Gbp2/Gbp | 3 | aPD-L1+aTGF-β vs PBS | Up-regulated genes | CD45+ |
| CC | GO:0043657 | host cell | 1.24E-03 | 9.52E-03 | Gbp2b/Gbp2/Gbp | 3 | aPD-L1+aTGF-β vs PBS | Up-regulated genes | CD45+ |
| CC | GO:0044215 | other organism | 1.59E-03 | 1.09E-02 | Gbp2b/Gbp2/Gbp | 3 | aPD-L1+aTGF-β vs PBS | Up-regulated genes | CD45+ |
| CC | GO:0044216 | other organism cell | 1.59E-03 | 1.09E-02 | Gbp2b/Gbp2/Gbp | 3 | aPD-L1+aTGF-β vs PBS | Up-regulated genes | CD45+ |
| CC | GO:0044217 | other organism part | 1.59E-03 | 1.09E-02 | Gbp2b/Gbp2/Gbp | 3 | aPD-L1+aTGF-β vs PBS | Up-regulated genes | CD45+ |
| CC | GO:0044754 | autolysosome | 4.54E-03 | 3.01E-02 | Irgm1/Sqstm1 | 2 | aPD-L1+aTGF-β vs PBS | Up-regulated genes | CD45+ |
| CC | GO:0098797 | plasma membrane protein complex | 5.09E-03 | 3.27E-02 | H2-Eb1/H2-Aa/C | 9 | aPD-L1+aTGF-β vs PBS | Up-regulated genes | CD45+ |
| CC | GO:0005767 | secondary lysosome | 5.42E-03 | 3.37E-02 | Irgm1/Sqstm1 | 2 | aPD-L1+aTGF-β vs PBS | Up-regulated genes | CD45+ |
| MF | GO:0003735 | structural constituent of ribosome | 6.19E-21 | 1.83E-18 | Rpl39/Rps27/Rpl | 22 | aPD-L1+aTGF-β vs PBS | Up-regulated genes | CD45+ |
| MF | GO:0005198 | structural molecule activity | 5.32E-12 | 7.87E-10 | Rpl39/Rps27/Rpl | 23 | aPD-L1+aTGF-β vs PBS | Up-regulated genes | CD45+ |
| MF | GO:0042379 | chemokine receptor binding | 2.23E-10 | 2.20E-08 | Cxcl10/Cxcl9/Ccl | 9 | aPD-L1+aTGF-β vs PBS | Up-regulated genes | CD45+ |
| MF | GO:0048020 | CCR chemokine receptor binding | 5.94E-09 | 4.40E-07 | Ccl4/Ccl5/Stat1/C | 7 | aPD-L1+aTGF-β vs PBS | Up-regulated genes | CD45+ |
| MF | GO:0008009 | chemokine activity | 2.45E-08 | 1.25E-06 | Cxcl10/Cxcl9/Ccl | 7 | aPD-L1+aTGF-β vs PBS | Up-regulated genes | CD45+ |
| MF | GO:0005125 | cytokine activity | 2.54E-08 | 1.25E-06 | Cxcl10/Il1b/Cxcl9 | 11 | aPD-L1+aTGF-β vs PBS | Up-regulated genes | CD45+ |
| MF | GO:0005126 | cytokine receptor binding | 6.81E-07 | 2.88E-05 | Cxcl10/Il1b/Cxcl9 | 12 | aPD-L1+aTGF-β vs PBS | Up-regulated genes | CD45+ |
| MF | GO:0001664 | G protein-coupled receptor binding | 2.60E-06 | 9.63E-05 | Cxcl10/Dnaja1/C | 11 | aPD-L1+aTGF-β vs PBS | Up-regulated genes | CD45+ |
| MF | GO:0048018 | receptor ligand activity | 3.70E-06 | 1.22E-04 | Cxcl10/Il1b/Cxcl9 | 12 | aPD-L1+aTGF-β vs PBS | Up-regulated genes | CD45+ |
| MF | GO:0030545 | receptor regulator activity | 8.30E-06 | 2.46E-04 | Cxcl10/Il1b/Cxcl9 | 12 | aPD-L1+aTGF-β vs PBS | Up-regulated genes | CD45+ |
| MF | GO:0042605 | peptide antigen binding | 5.15E-05 | 1.39E-03 | H2-Aa/H2-Ab1/H2 | 4 | aPD-L1+aTGF-β vs PBS | Up-regulated genes | CD45+ |
| MF | GO:0031386 | protein tag | 1.17E-04 | 2.88E-03 | Uba52/Ubc/Isg15 | 3 | aPD-L1+aTGF-β vs PBS | Up-regulated genes | CD45+ |
| MF | GO:0044389 | ubiquitin-like protein ligase binding | 4.04E-04 | 9.04E-03 | H2-Ab1/Dnaja1/N | 10 | aPD-L1+aTGF-β vs PBS | Up-regulated genes | CD45+ |
| MF | GO:0003823 | antigen binding | 4.28E-04 | 9.04E-03 | H2-Aa/H2-Ab1/H2 | 4 | aPD-L1+aTGF-β vs PBS | Up-regulated genes | CD45+ |
| MF | GO:0042277 | peptide binding | 6.64E-04 | 1.31E-02 | H2-Aa/Cd74/H2- | 8 | aPD-L1+aTGF-β vs PBS | Up-regulated genes | CD45+ |
| MF | GO:0031625 | ubiquitin protein ligase binding | 1.07E-03 | 1.98E-02 | H2-Ab1/Dnaja1/N | 9 | aPD-L1+aTGF-β vs PBS | Up-regulated genes | CD45+ |
| MF | GO:0008201 | heparin binding | 2.32E-03 | 4.04E-02 | Cxcl10/Ccl5/Rpl2 | 5 | aPD-L1+aTGF-β vs PBS | Up-regulated genes | CD45+ |
| MF | GO:0003924 | GTPase activity | 2.86E-03 | 4.55E-02 | Gbp2b/Gbp2/Irgm | 8 | aPD-L1+aTGF-β vs PBS | Up-regulated genes | CD45+ |
| MF | GO:0033218 | amide binding | 2.92E-03 | 4.55E-02 | H2-Aa/Cd74/H2- | 8 | aPD-L1+aTGF-β vs PBS | Up-regulated genes | CD45+ |
| BP | GO:0009617 | response to bacterium | 9.31E-08 | 1.17E-04 | Cxcl10/Cxcl9/Ly6 | 10 | aPD-L1+aTGF-β vs PBS | Up-regulated genes | CD45- |
| BP | GO:1990868 | response to chemokine | 2.86E-07 | 1.19E-04 | Cxcl10/Cxcl9/Cxc | 5 | aPD-L1+aTGF-β vs PBS | Up-regulated genes | CD45- |
| BP | GO:1990869 | cellular response to chemokine | 2.86E-07 | 1.19E-04 | Cxcl10/Cxcl9/Cxc | 5 | aPD-L1+aTGF-β vs PBS | Up-regulated genes | CD45- |
| BP | GO:0030593 | neutrophil chemotaxis | 8.35E-07 | 2.62E-04 | Cxcl10/Cxcl9/Cxc | 5 | aPD-L1+aTGF-β vs PBS | Up-regulated genes | CD45- |
| BP | GO:0019882 | antigen processing and presentation | 1.19E-06 | 2.99E-04 | B2m/Psmb9/H2-D | 5 | aPD-L1+aTGF-β vs PBS | Up-regulated genes | CD45- |
| BP | GO:0061844 | antimicrobial humoral immune response mediat | 1.51E-06 | 3.07E-04 | Cxcl10/Cxcl9/Cxc | 4 | aPD-L1+aTGF-β vs PBS | Up-regulated genes | CD45- |
| BP | GO:1990266 | neutrophil migration | 2.26E-06 | 3.07E-04 | Cxcl10/Cxcl9/Cxc | 5 | aPD-L1+aTGF-β vs PBS | Up-regulated genes | CD45- |
| BP | GO:0030595 | leukocyte chemotaxis | 2.67E-06 | 3.07E-04 | Cxcl10/Cxcl9/Cxc | 6 | aPD-L1+aTGF-β vs PBS | Up-regulated genes | CD45- |
| BP | GO:0097529 | myeloid leukocyte migration | 2.67E-06 | 3.07E-04 | Cxcl10/Cxcl9/Cxc | 6 | aPD-L1+aTGF-β vs PBS | Up-regulated genes | CD45- |
| BP | GO:0002824 | positive regulation of adaptive immune respons | 2.70E-06 | 3.07E-04 | B2m/Irf1/H2-D1/H | 5 | aPD-L1+aTGF-β vs PBS | Up-regulated genes | CD45- |
| BP | GO:0071621 | granulocyte chemotaxis | 2.70E-06 | 3.07E-04 | Cxcl10/Cxcl9/Cxc | 5 | aPD-L1+aTGF-β vs PBS | Up-regulated genes | CD45- |
| BP | GO:0002821 | positive regulation of adaptive immune respons | 3.97E-06 | 3.97E-04 | B2m/Irf1/H2-D1/H | 5 | aPD-L1+aTGF-β vs PBS | Up-regulated genes | CD45- |
| BP | GO:0019221 | cytokine-mediated signaling pathway | 4.24E-06 | 3.97E-04 | Cxcl10/Cxcl9/Irf1/ | 7 | aPD-L1+aTGF-β vs PBS | Up-regulated genes | CD45- |
| BP | GO:0019730 | antimicrobial humoral response | 4.47E-06 | 3.97E-04 | Cxcl10/Cxcl9/Cxc | 4 | aPD-L1+aTGF-β vs PBS | Up-regulated genes | CD45- |
| BP | GO:0002699 | positive regulation of immune effector process | 5.22E-06 | 3.97E-04 | B2m/Irf1/Cxcl1/H2 | 6 | aPD-L1+aTGF-β vs PBS | Up-regulated genes | CD45- |
| BP | GO:0001912 | positive regulation of leukocyte mediated cytoto | 5.38E-06 | 3.97E-04 | B2m/Cxcl1/H2-D1 | 4 | aPD-L1+aTGF-β vs PBS | Up-regulated genes | CD45- |
| BP | GO:0070098 | chemokine-mediated signaling pathway | 5.38E-06 | 3.97E-04 | Cxcl10/Cxcl9/Cxc | 4 | aPD-L1+aTGF-β vs PBS | Up-regulated genes | CD45- |
| BP | GO:0097530 | granulocyte migration | 6.57E-06 | 4.57E-04 | Cxcl10/Cxcl9/Cxc | 5 | aPD-L1+aTGF-β vs PBS | Up-regulated genes | CD45- |
| BP | GO:0031343 | positive regulation of cell killing | 9.65E-06 | 6.36E-04 | B2m/Cxcl1/H2-D1 | 4 | aPD-L1+aTGF-β vs PBS | Up-regulated genes | CD45- |
| BP | GO:0002428 | antigen processing and presentation of peptide | 1.08E-05 | 6.78E-04 | B2m/H2-D1/H2-K | 3 | aPD-L1+aTGF-β vs PBS | Up-regulated genes | CD45- |
| BP | GO:0071248 | cellular response to metal ion | 1.23E-05 | 7.31E-04 | Mt1/B2m/Mt2/Fos | 5 | aPD-L1+aTGF-β vs PBS | Up-regulated genes | CD45- |
| BP | GO:0001910 | regulation of leukocyte mediated cytotoxicity | 1.60E-05 | 9.01E-04 | B2m/Cxcl1/H2-D1 | 4 | aPD-L1+aTGF-β vs PBS | Up-regulated genes | CD45- |
| BP | GO:0002475 | antigen processing and presentation via MHC cl | 1.80E-05 | 9.01E-04 | B2m/H2-D1/H2-K | 3 | aPD-L1+aTGF-β vs PBS | Up-regulated genes | CD45- |
| BP | GO:0071276 | cellular response to cadmium ion | 1.80E-05 | 9.01E-04 | Mt1/Mt2/Fos | 3 | aPD-L1+aTGF-β vs PBS | Up-regulated genes | CD45- |
| BP | GO:0060326 | cell chemotaxis | 1.89E-05 | 9.01E-04 | Cxcl10/Cxcl9/Cxc | 6 | aPD-L1+aTGF-β vs PBS | Up-regulated genes | CD45- |
| BP | GO:0002822 | regulation of adaptive immune response based | 1.89E-05 | 9.01E-04 | B2m/Irf1/H2-D1/H | 5 | aPD-L1+aTGF-β vs PBS | Up-regulated genes | CD45- |
| BP | GO:0002237 | response to molecule of bacterial origin | 1.94E-05 | 9.01E-04 | Cxcl10/Cxcl9/B2n | 6 | aPD-L1+aTGF-β vs PBS | Up-regulated genes | CD45- |
| BP | GO:0002483 | antigen processing and presentation of endoger | 2.43E-05 | 1.05E-03 | B2m/H2-D1/H2-K | 3 | aPD-L1+aTGF-β vs PBS | Up-regulated genes | CD45- |
| BP | GO:0019885 | antigen processing and presentation of endoger | 2.43E-05 | 1.05E-03 | B2m/H2-D1/H2-K | 3 | aPD-L1+aTGF-β vs PBS | Up-regulated genes | CD45- |
| BP | GO:0071241 | cellular response to inorganic substance | 2.63E-05 | 1.10E-03 | Mt1/B2m/Mt2/Fos | 5 | aPD-L1+aTGF-β vs PBS | Up-regulated genes | CD45- |
| BP | GO:0006954 | inflammatory response | 3.01E-05 | 1.17E-03 | Cxcl10/Cxcl9/Cxc | 8 | aPD-L1+aTGF-β vs PBS | Up-regulated genes | CD45- |
| BP | GO:0002819 | regulation of adaptive immune response | 3.02E-05 | 1.17E-03 | B2m/Irf1/H2-D1/H | 5 | aPD-L1+aTGF-β vs PBS | Up-regulated genes | CD45- |
| BP | GO:0071222 | cellular response to lipopolysaccharide | 3.13E-05 | 1.17E-03 | Cxcl10/Cxcl9/Cxc | 5 | aPD-L1+aTGF-β vs PBS | Up-regulated genes | CD45- |
| BP | GO:0019883 | antigen processing and presentation of endoger | 3.18E-05 | 1.17E-03 | B2m/H2-D1/H2-K | 3 | aPD-L1+aTGF-β vs PBS | Up-regulated genes | CD45- |
| BP | GO:0031341 | regulation of cell killing | 3.53E-05 | 1.27E-03 | B2m/Cxcl1/H2-D1 | 4 | aPD-L1+aTGF-β vs PBS | Up-regulated genes | CD45- |
| BP | GO:0071219 | cellular response to molecule of bacterial origin | 4.07E-05 | 1.42E-03 | Cxcl10/Cxcl9/Cxc | 5 | aPD-L1+aTGF-β vs PBS | Up-regulated genes | CD45- |
| BP | GO:0001916 | positive regulation of T cell mediated cytotoxicity | 4.57E-05 | 1.53E-03 | B2m/H2-D1/H2-K | 3 | aPD-L1+aTGF-β vs PBS | Up-regulated genes | CD45- |

| | | | | | | | | | |
|---|---|---|---|---|---|---|---|---|---|
| BP | GO:0071396 | cellular response to lipid | 4.64E-05 | 1.53E-03 | Cxcl10/Cxcl9/Cxc | 7 | aPD-L1+aTGF-β vs PBS | Up-regulated genes | CD45- |
| BP | GO:0050900 | leukocyte migration | 5.09E-05 | 1.64E-03 | Cxcl10/Cxcl9/Cxc | 6 | aPD-L1+aTGF-β vs PBS | Up-regulated genes | CD45- |
| BP | GO:0001909 | leukocyte mediated cytotoxicity | 5.62E-05 | 1.74E-03 | B2m/Cxcl1/H2-D1 | 4 | aPD-L1+aTGF-β vs PBS | Up-regulated genes | CD45- |
| BP | GO:0046686 | response to cadmium ion | 5.69E-05 | 1.74E-03 | Mt1/Mt2/Fos | 3 | aPD-L1+aTGF-β vs PBS | Up-regulated genes | CD45- |
| BP | GO:0002474 | antigen processing and presentation of peptide | 6.31E-05 | 1.88E-03 | B2m/H2-D1/H2-K | 3 | aPD-L1+aTGF-β vs PBS | Up-regulated genes | CD45- |
| BP | GO:0001914 | regulation of T cell mediated cytotoxicity | 6.97E-05 | 2.03E-03 | B2m/H2-D1/H2-K | 3 | aPD-L1+aTGF-β vs PBS | Up-regulated genes | CD45- |
| BP | GO:0071216 | cellular response to biotic stimulus | 7.38E-05 | 2.10E-03 | Cxcl10/Cxcl9/Cxc | 5 | aPD-L1+aTGF-β vs PBS | Up-regulated genes | CD45- |
| BP | GO:0002697 | regulation of immune effector process | 1.06E-04 | 2.96E-03 | B2m/Irf1/Cxcl1/H2 | 6 | aPD-L1+aTGF-β vs PBS | Up-regulated genes | CD45- |
| BP | GO:0006959 | humoral immune response | 1.33E-04 | 3.62E-03 | Cxcl10/Cxcl9/Cxc | 4 | aPD-L1+aTGF-β vs PBS | Up-regulated genes | CD45- |
| BP | GO:0001906 | cell killing | 1.43E-04 | 3.82E-03 | B2m/Cxcl1/H2-D1 | 4 | aPD-L1+aTGF-β vs PBS | Up-regulated genes | CD45- |
| BP | GO:1903706 | regulation of hemopoiesis | 1.52E-04 | 3.89E-03 | B2m/Irf1/Cxcl1/Nf | 6 | aPD-L1+aTGF-β vs PBS | Up-regulated genes | CD45- |
| BP | GO:0055065 | metal ion homeostasis | 1.52E-04 | 3.89E-03 | Cxcl10/Mt1/B2m/ | 7 | aPD-L1+aTGF-β vs PBS | Up-regulated genes | CD45- |
| BP | GO:0002705 | positive regulation of leukocyte mediated immun | 1.66E-04 | 4.16E-03 | B2m/Cxcl1/H2-D1 | 4 | aPD-L1+aTGF-β vs PBS | Up-regulated genes | CD45- |
| BP | GO:0010038 | response to metal ion | 1.74E-04 | 4.20E-03 | Mt1/B2m/Mt2/Fos | 5 | aPD-L1+aTGF-β vs PBS | Up-regulated genes | CD45- |
| BP | GO:0032496 | response to lipopolysaccharide | 1.74E-04 | 4.20E-03 | Cxcl10/Cxcl9/Cxc | 5 | aPD-L1+aTGF-β vs PBS | Up-regulated genes | CD45- |
| BP | GO:0002711 | positive regulation of T cell mediated immunity | 2.15E-04 | 5.05E-03 | B2m/H2-D1/H2-K | 3 | aPD-L1+aTGF-β vs PBS | Up-regulated genes | CD45- |
| BP | GO:0002460 | adaptive immune response based on somatic re | 2.18E-04 | 5.05E-03 | B2m/Irf1/H2-D1/H | 5 | aPD-L1+aTGF-β vs PBS | Up-regulated genes | CD45- |
| BP | GO:0048002 | antigen processing and presentation of peptide | 2.29E-04 | 5.22E-03 | B2m/H2-D1/H2-K | 3 | aPD-L1+aTGF-β vs PBS | Up-regulated genes | CD45- |
| BP | GO:0001913 | T cell mediated cytotoxicity | 2.44E-04 | 5.47E-03 | B2m/H2-D1/H2-K | 3 | aPD-L1+aTGF-β vs PBS | Up-regulated genes | CD45- |
| BP | GO:0032103 | positive regulation of response to external stimul | 3.29E-04 | 7.23E-03 | Cxcl10/Cxcl1/Nfkl | 5 | aPD-L1+aTGF-β vs PBS | Up-regulated genes | CD45- |
| BP | GO:0006875 | cellular metal ion homeostasis | 4.64E-04 | 1.00E-02 | Cxcl10/Mt1/Cxcl2 | 6 | aPD-L1+aTGF-β vs PBS | Up-regulated genes | CD45- |
| BP | GO:0035914 | skeletal muscle cell differentiation | 5.04E-04 | 1.04E-02 | Fos/Nupr1/Egr1 | 3 | aPD-L1+aTGF-β vs PBS | Up-regulated genes | CD45- |
| BP | GO:0045820 | negative regulation of glycolytic process | 5.07E-04 | 1.04E-02 | Nupr1/Ier3 | 2 | aPD-L1+aTGF-β vs PBS | Up-regulated genes | CD45- |
| BP | GO:0071280 | cellular response to copper ion | 5.07E-04 | 1.04E-02 | Mt1/Mt2 | 2 | aPD-L1+aTGF-β vs PBS | Up-regulated genes | CD45- |
| BP | GO:0002709 | regulation of T cell mediated immunity | 5.29E-04 | 1.07E-02 | B2m/H2-D1/H2-K | 3 | aPD-L1+aTGF-β vs PBS | Up-regulated genes | CD45- |
| BP | GO:0030812 | negative regulation of nucleotide catabolic proce | 5.91E-04 | 1.16E-02 | Nupr1/Ier3 | 2 | aPD-L1+aTGF-β vs PBS | Up-regulated genes | CD45- |
| BP | GO:0051198 | negative regulation of coenzyme metabolic proc | 5.91E-04 | 1.16E-02 | Nupr1/Ier3 | 2 | aPD-L1+aTGF-β vs PBS | Up-regulated genes | CD45- |
| BP | GO:0010977 | negative regulation of neuron projection develo | 6.33E-04 | 1.22E-02 | B2m/H2-D1/H2-K | 4 | aPD-L1+aTGF-β vs PBS | Up-regulated genes | CD45- |
| BP | GO:0002486 | antigen processing and presentation of endoger | 6.81E-04 | 1.27E-02 | H2-D1/H2-K1 | 2 | aPD-L1+aTGF-β vs PBS | Up-regulated genes | CD45- |
| BP | GO:0035994 | response to muscle stretch | 6.81E-04 | 1.27E-02 | Nfkbia/Fos | 2 | aPD-L1+aTGF-β vs PBS | Up-regulated genes | CD45- |
| BP | GO:0002250 | adaptive immune response | 6.87E-04 | 1.27E-02 | B2m/Irf1/H2-D1/H | 5 | aPD-L1+aTGF-β vs PBS | Up-regulated genes | CD45- |
| BP | GO:0050778 | positive regulation of immune response | 7.03E-04 | 1.27E-02 | B2m/Irf1/H2-D1/H | 6 | aPD-L1+aTGF-β vs PBS | Up-regulated genes | CD45- |
| BP | GO:0031349 | positive regulation of defense response | 7.10E-04 | 1.27E-02 | Irf1/Cxcl1/Nfkbia/ | 5 | aPD-L1+aTGF-β vs PBS | Up-regulated genes | CD45- |
| BP | GO:1903708 | positive regulation of hemopoiesis | 7.68E-04 | 1.30E-02 | Irf1/Cxcl1/Fos/Nfk | 4 | aPD-L1+aTGF-β vs PBS | Up-regulated genes | CD45- |
| BP | GO:0006935 | chemotaxis | 7.75E-04 | 1.30E-02 | Cxcl10/Cxcl9/Cxc | 6 | aPD-L1+aTGF-β vs PBS | Up-regulated genes | CD45- |
| BP | GO:0002476 | antigen processing and presentation of endoger | 7.77E-04 | 1.30E-02 | H2-D1/H2-K1 | 2 | aPD-L1+aTGF-β vs PBS | Up-regulated genes | CD45- |
| BP | GO:0002484 | antigen processing and presentation of endoger | 7.77E-04 | 1.30E-02 | H2-D1/H2-K1 | 2 | aPD-L1+aTGF-β vs PBS | Up-regulated genes | CD45- |
| BP | GO:0071498 | cellular response to fluid shear stress | 7.77E-04 | 1.30E-02 | Klf2/Spp1 | 2 | aPD-L1+aTGF-β vs PBS | Up-regulated genes | CD45- |
| BP | GO:0042330 | taxis | 8.04E-04 | 1.31E-02 | Cxcl10/Cxcl9/Cxc | 6 | aPD-L1+aTGF-β vs PBS | Up-regulated genes | CD45- |
| BP | GO:0030003 | cellular cation homeostasis | 8.14E-04 | 1.31E-02 | Cxcl10/Mt1/Cxcl2 | 6 | aPD-L1+aTGF-β vs PBS | Up-regulated genes | CD45- |
| BP | GO:0002703 | regulation of leukocyte mediated immunity | 8.23E-04 | 1.31E-02 | B2m/Cxcl1/H2-D1 | 4 | aPD-L1+aTGF-β vs PBS | Up-regulated genes | CD45- |
| BP | GO:0007263 | nitric oxide mediated signal transduction | 8.79E-04 | 1.31E-02 | Mt1/Mt2 | 2 | aPD-L1+aTGF-β vs PBS | Up-regulated genes | CD45- |
| BP | GO:0030809 | negative regulation of nucleotide biosynthetic pr | 8.79E-04 | 1.31E-02 | Nupr1/Ier3 | 2 | aPD-L1+aTGF-β vs PBS | Up-regulated genes | CD45- |
| BP | GO:0045624 | positive regulation of T-helper cell differentiation | 8.79E-04 | 1.31E-02 | Irf1/Nfkbiz | 2 | aPD-L1+aTGF-β vs PBS | Up-regulated genes | CD45- |
| BP | GO:1900372 | negative regulation of purine nucleotide biosynt | 8.79E-04 | 1.31E-02 | Nupr1/Ier3 | 2 | aPD-L1+aTGF-β vs PBS | Up-regulated genes | CD45- |
| BP | GO:1901739 | regulation of myoblast fusion | 8.79E-04 | 1.31E-02 | Cxcl10/Cxcl9 | 2 | aPD-L1+aTGF-β vs PBS | Up-regulated genes | CD45- |
| BP | GO:2001170 | negative regulation of ATP biosynthetic process | 8.79E-04 | 1.31E-02 | Nupr1/Ier3 | 2 | aPD-L1+aTGF-β vs PBS | Up-regulated genes | CD45- |
| BP | GO:0006873 | cellular ion homeostasis | 9.06E-04 | 1.34E-02 | Cxcl10/Mt1/Cxcl2 | 6 | aPD-L1+aTGF-β vs PBS | Up-regulated genes | CD45- |
| BP | GO:0006882 | cellular zinc ion homeostasis | 1.10E-03 | 1.61E-02 | Mt1/Mt2 | 2 | aPD-L1+aTGF-β vs PBS | Up-regulated genes | CD45- |
| BP | GO:0031345 | negative regulation of cell projection organizatio | 1.12E-03 | 1.61E-02 | B2m/H2-D1/H2-K | 4 | aPD-L1+aTGF-β vs PBS | Up-regulated genes | CD45- |
| BP | GO:0010043 | response to zinc ion | 1.22E-03 | 1.66E-02 | Mt1/Mt2 | 2 | aPD-L1+aTGF-β vs PBS | Up-regulated genes | CD45- |
| BP | GO:0046688 | response to copper ion | 1.22E-03 | 1.66E-02 | Mt1/Mt2 | 2 | aPD-L1+aTGF-β vs PBS | Up-regulated genes | CD45- |
| BP | GO:0051195 | negative regulation of cofactor metabolic proces | 1.22E-03 | 1.66E-02 | Nupr1/Ier3 | 2 | aPD-L1+aTGF-β vs PBS | Up-regulated genes | CD45- |
| BP | GO:0055069 | zinc ion homeostasis | 1.22E-03 | 1.66E-02 | Mt1/Mt2 | 2 | aPD-L1+aTGF-β vs PBS | Up-regulated genes | CD45- |
| BP | GO:0090025 | regulation of monocyte chemotaxis | 1.22E-03 | 1.66E-02 | Cxcl10/Dusp1 | 2 | aPD-L1+aTGF-β vs PBS | Up-regulated genes | CD45- |
| BP | GO:0002708 | positive regulation of lymphocyte mediated immu | 1.30E-03 | 1.75E-02 | B2m/H2-D1/H2-K | 3 | aPD-L1+aTGF-β vs PBS | Up-regulated genes | CD45- |
| BP | GO:0010499 | proteasomal ubiquitin-independent protein catal | 1.35E-03 | 1.77E-02 | Psmb9/Psmb8 | 2 | aPD-L1+aTGF-β vs PBS | Up-regulated genes | CD45- |
| BP | GO:0043372 | positive regulation of CD4-positive, alpha-beta T | 1.35E-03 | 1.77E-02 | Irf1/Nfkbiz | 2 | aPD-L1+aTGF-β vs PBS | Up-regulated genes | CD45- |
| BP | GO:0031347 | regulation of defense response | 1.36E-03 | 1.77E-02 | Irf1/Cxcl1/Nfkbia/ | 6 | aPD-L1+aTGF-β vs PBS | Up-regulated genes | CD45- |
| BP | GO:0098542 | defense response to other organism | 1.45E-03 | 1.86E-02 | Cxcl10/Cxcl9/Irf1/ | 5 | aPD-L1+aTGF-β vs PBS | Up-regulated genes | CD45- |
| BP | GO:0072503 | cellular divalent inorganic cation homeostasis | 1.47E-03 | 1.86E-02 | Cxcl10/Mt1/Cxcl2 | 5 | aPD-L1+aTGF-β vs PBS | Up-regulated genes | CD45- |
| BP | GO:0002478 | antigen processing and presentation of exogenc | 1.48E-03 | 1.86E-02 | B2m/H2-K1 | 2 | aPD-L1+aTGF-β vs PBS | Up-regulated genes | CD45- |
| BP | GO:0034405 | response to fluid shear stress | 1.48E-03 | 1.86E-02 | Klf2/Spp1 | 2 | aPD-L1+aTGF-β vs PBS | Up-regulated genes | CD45- |
| BP | GO:0060142 | regulation of syncytium formation by plasma mer | 1.62E-03 | 2.01E-02 | Cxcl10/Cxcl9 | 2 | aPD-L1+aTGF-β vs PBS | Up-regulated genes | CD45- |
| BP | GO:0002456 | T cell mediated immunity | 1.64E-03 | 2.02E-02 | B2m/H2-D1/H2-K | 3 | aPD-L1+aTGF-β vs PBS | Up-regulated genes | CD45- |
| BP | GO:0016525 | negative regulation of angiogenesis | 1.75E-03 | 2.12E-02 | Cxcl10/Klf2/Dcn | 3 | aPD-L1+aTGF-β vs PBS | Up-regulated genes | CD45- |
| BP | GO:2000516 | positive regulation of CD4-positive, alpha-beta T | 1.76E-03 | 2.12E-02 | Irf1/Nfkbiz | 2 | aPD-L1+aTGF-β vs PBS | Up-regulated genes | CD45- |
| BP | GO:0030217 | T cell differentiation | 1.89E-03 | 2.24E-02 | B2m/Irf1/Nfkbiz/E | 4 | aPD-L1+aTGF-β vs PBS | Up-regulated genes | CD45- |
| BP | GO:1900543 | negative regulation of purine nucleotide metabo | 1.91E-03 | 2.24E-02 | Nupr1/Ier3 | 2 | aPD-L1+aTGF-β vs PBS | Up-regulated genes | CD45- |
| BP | GO:1903579 | negative regulation of ATP metabolic process | 1.91E-03 | 2.24E-02 | Nupr1/Ier3 | 2 | aPD-L1+aTGF-β vs PBS | Up-regulated genes | CD45- |
| BP | GO:2000181 | negative regulation of blood vessel morphogene | 1.97E-03 | 2.28E-02 | Cxcl10/Klf2/Dcn | 3 | aPD-L1+aTGF-β vs PBS | Up-regulated genes | CD45- |
| BP | GO:0072507 | divalent inorganic cation homeostasis | 1.99E-03 | 2.28E-02 | Cxcl10/Mt1/Cxcl2 | 5 | aPD-L1+aTGF-β vs PBS | Up-regulated genes | CD45- |
| BP | GO:0045980 | negative regulation of nucleotide metabolic proc | 2.07E-03 | 2.36E-02 | Nupr1/Ier3 | 2 | aPD-L1+aTGF-β vs PBS | Up-regulated genes | CD45- |
| BP | GO:0045665 | negative regulation of neuron differentiation | 2.13E-03 | 2.39E-02 | B2m/H2-D1/H2-K | 4 | aPD-L1+aTGF-β vs PBS | Up-regulated genes | CD45- |
| BP | GO:0010035 | response to inorganic substance | 2.14E-03 | 2.39E-02 | Mt1/B2m/Mt2/Fos | 5 | aPD-L1+aTGF-β vs PBS | Up-regulated genes | CD45- |
| BP | GO:0055076 | transition metal ion homeostasis | 2.22E-03 | 2.45E-02 | Mt1/B2m/Mt2 | 3 | aPD-L1+aTGF-β vs PBS | Up-regulated genes | CD45- |
| BP | GO:0019884 | antigen processing and presentation of exogenc | 2.23E-03 | 2.45E-02 | B2m/H2-K1 | 2 | aPD-L1+aTGF-β vs PBS | Up-regulated genes | CD45- |
| BP | GO:0050729 | positive regulation of inflammatory response | 2.35E-03 | 2.56E-02 | Nfkbia/Nupr1/Nfkl | 3 | aPD-L1+aTGF-β vs PBS | Up-regulated genes | CD45- |
| BP | GO:0045622 | regulation of T-helper cell differentiation | 2.40E-03 | 2.59E-02 | Irf1/Nfkbiz | 2 | aPD-L1+aTGF-β vs PBS | Up-regulated genes | CD45- |
| BP | GO:0060537 | muscle tissue development | 2.50E-03 | 2.68E-02 | Ifrd1/Zfand5/Fos/ | 5 | aPD-L1+aTGF-β vs PBS | Up-regulated genes | CD45- |
| BP | GO:1901343 | negative regulation of vasculature development | 2.69E-03 | 2.86E-02 | Cxcl10/Klf2/Dcn | 3 | aPD-L1+aTGF-β vs PBS | Up-regulated genes | CD45- |
| BP | GO:0007616 | long-term memory | 2.94E-03 | 3.09E-02 | Sgk1/Egr1 | 2 | aPD-L1+aTGF-β vs PBS | Up-regulated genes | CD45- |
| BP | GO:0007520 | myoblast fusion | 3.13E-03 | 3.27E-02 | Cxcl10/Cxcl9 | 2 | aPD-L1+aTGF-β vs PBS | Up-regulated genes | CD45- |
| BP | GO:0002706 | regulation of lymphocyte mediated immunity | 3.72E-03 | 3.80E-02 | B2m/H2-D1/H2-K | 3 | aPD-L1+aTGF-β vs PBS | Up-regulated genes | CD45- |
| BP | GO:0006110 | regulation of glycolytic process | 3.73E-03 | 3.80E-02 | Nupr1/Ier3 | 2 | aPD-L1+aTGF-β vs PBS | Up-regulated genes | CD45- |
| BP | GO:0030811 | regulation of nucleotide catabolic process | 3.73E-03 | 3.80E-02 | Nupr1/Ier3 | 2 | aPD-L1+aTGF-β vs PBS | Up-regulated genes | CD45- |
| BP | GO:0002443 | leukocyte mediated immunity | 3.82E-03 | 3.86E-02 | B2m/Cxcl1/H2-D1 | 4 | aPD-L1+aTGF-β vs PBS | Up-regulated genes | CD45- |
| BP | GO:0010765 | positive regulation of sodium ion transport | 4.17E-03 | 4.08E-02 | Cxcl1/Sgk1 | 2 | aPD-L1+aTGF-β vs PBS | Up-regulated genes | CD45- |
| BP | GO:0043370 | regulation of CD4-positive, alpha-beta T cell diff | 4.17E-03 | 4.08E-02 | Irf1/Nfkbiz | 2 | aPD-L1+aTGF-β vs PBS | Up-regulated genes | CD45- |

| | | | | | | | | | |
|---|---|---|---|---|---|---|---|---|---|
| BP | GO:0046638 | positive regulation of alpha-beta T cell differentia | 4.17E-03 | 4.08E-02 | Irf1/Nfkbiz | 2 | aPD-L1+aTGF-β vs PBS | Up-regulated genes | CD45- |
| BP | GO:1902107 | positive regulation of leukocyte differentiation | 4.17E-03 | 4.08E-02 | Irf1/Fos/Nfkbiz | 3 | aPD-L1+aTGF-β vs PBS | Up-regulated genes | CD45- |
| BP | GO:0009636 | response to toxic substance | 4.27E-03 | 4.12E-02 | Mt1/Mt2/Nupr1/Eg | 4 | aPD-L1+aTGF-β vs PBS | Up-regulated genes | CD45- |
| BP | GO:0002521 | leukocyte differentiation | 4.28E-03 | 4.12E-02 | B2m/Irf1/Fos/Nfkb | 5 | aPD-L1+aTGF-β vs PBS | Up-regulated genes | CD45- |
| BP | GO:0002548 | monocyte chemotaxis | 4.62E-03 | 4.35E-02 | Cxcl10/Dusp1 | 2 | aPD-L1+aTGF-β vs PBS | Up-regulated genes | CD45- |
| BP | GO:0010677 | negative regulation of cellular carbohydrate met | 4.62E-03 | 4.35E-02 | Nupr1/Ier3 | 2 | aPD-L1+aTGF-β vs PBS | Up-regulated genes | CD45- |
| BP | GO:0071675 | regulation of mononuclear cell migration | 4.62E-03 | 4.35E-02 | Cxcl10/Dusp1 | 2 | aPD-L1+aTGF-β vs PBS | Up-regulated genes | CD45- |
| BP | GO:0050727 | regulation of inflammatory response | 4.83E-03 | 4.52E-02 | Nfkbia/Nupr1/Nfkl | 4 | aPD-L1+aTGF-β vs PBS | Up-regulated genes | CD45- |
| BP | GO:0050768 | negative regulation of neurogenesis | 5.09E-03 | 4.70E-02 | B2m/H2-D1/H2-K | 4 | aPD-L1+aTGF-β vs PBS | Up-regulated genes | CD45- |
| BP | GO:0098754 | detoxification | 5.10E-03 | 4.70E-02 | Mt1/Mt2 | 2 | aPD-L1+aTGF-β vs PBS | Up-regulated genes | CD45- |
| BP | GO:0051147 | regulation of muscle cell differentiation | 5.38E-03 | 4.92E-02 | Cxcl10/Cxcl9/Tms | 3 | aPD-L1+aTGF-β vs PBS | Up-regulated genes | CD45- |
| CC | GO:0042612 | MHC class I protein complex | 2.76E-06 | 1.85E-04 | B2m/H2-D1/H2-K | 3 | aPD-L1+aTGF-β vs PBS | Up-regulated genes | CD45- |
| CC | GO:0042824 | MHC class I peptide loading complex | 6.05E-06 | 2.03E-04 | B2m/H2-D1/H2-K | 3 | aPD-L1+aTGF-β vs PBS | Up-regulated genes | CD45- |
| CC | GO:0042611 | MHC protein complex | 1.60E-05 | 3.56E-04 | B2m/H2-D1/H2-K | 3 | aPD-L1+aTGF-β vs PBS | Up-regulated genes | CD45- |
| CC | GO:0009897 | external side of plasma membrane | 4.04E-05 | 6.77E-04 | Cxcl10/Cxcl9/Ly6 | 6 | aPD-L1+aTGF-β vs PBS | Up-regulated genes | CD45- |
| CC | GO:0098552 | side of membrane | 5.19E-04 | 6.96E-03 | Cxcl10/Cxcl9/Ly6 | 6 | aPD-L1+aTGF-β vs PBS | Up-regulated genes | CD45- |
| CC | GO:0005839 | proteasome core complex | 9.01E-04 | 1.01E-02 | Psmb9/Psmb8 | 2 | aPD-L1+aTGF-β vs PBS | Up-regulated genes | CD45- |
| CC | GO:0005797 | Golgi medial cisterna | 1.66E-03 | 1.59E-02 | H2-D1/H2-K1 | 2 | aPD-L1+aTGF-β vs PBS | Up-regulated genes | CD45- |
| CC | GO:0070971 | endoplasmic reticulum exit site | 2.12E-03 | 1.78E-02 | H2-D1/H2-K1 | 2 | aPD-L1+aTGF-β vs PBS | Up-regulated genes | CD45- |
| CC | GO:0030176 | integral component of endoplasmic reticulum me | 4.62E-03 | 3.44E-02 | B2m/H2-D1/H2-K | 3 | aPD-L1+aTGF-β vs PBS | Up-regulated genes | CD45- |
| CC | GO:0031227 | intrinsic component of endoplasmic reticulum me | 5.35E-03 | 3.59E-02 | B2m/H2-D1/H2-K | 3 | aPD-L1+aTGF-β vs PBS | Up-regulated genes | CD45- |
| MF | GO:0008009 | chemokine activity | 1.75E-06 | 1.68E-04 | Cxcl10/Cxcl9/Cxc | 4 | aPD-L1+aTGF-β vs PBS | Up-regulated genes | CD45- |
| MF | GO:0042379 | chemokine receptor binding | 5.18E-06 | 2.49E-04 | Cxcl10/Cxcl9/Cxc | 4 | aPD-L1+aTGF-β vs PBS | Up-regulated genes | CD45- |
| MF | GO:0005125 | cytokine activity | 1.47E-05 | 4.70E-04 | Cxcl10/Cxcl9/Cxc | 5 | aPD-L1+aTGF-β vs PBS | Up-regulated genes | CD45- |
| MF | GO:0030881 | beta-2-microglobulin binding | 3.86E-04 | 6.18E-03 | H2-D1/H2-K1 | 2 | aPD-L1+aTGF-β vs PBS | Up-regulated genes | CD45- |
| MF | GO:0042610 | CD8 receptor binding | 3.86E-04 | 6.18E-03 | H2-D1/H2-K1 | 2 | aPD-L1+aTGF-β vs PBS | Up-regulated genes | CD45- |
| MF | GO:0046977 | TAP binding | 3.86E-04 | 6.18E-03 | H2-D1/H2-K1 | 2 | aPD-L1+aTGF-β vs PBS | Up-regulated genes | CD45- |
| MF | GO:0042608 | T cell receptor binding | 7.32E-04 | 9.60E-03 | H2-D1/H2-K1 | 2 | aPD-L1+aTGF-β vs PBS | Up-regulated genes | CD45- |
| MF | GO:0048018 | receptor ligand activity | 8.00E-04 | 9.60E-03 | Cxcl10/Cxcl9/Cxc | 5 | aPD-L1+aTGF-β vs PBS | Up-regulated genes | CD45- |
| MF | GO:0004298 | threonine-type endopeptidase activity | 1.06E-03 | 1.02E-02 | Psmb9/Psmb8 | 2 | aPD-L1+aTGF-β vs PBS | Up-regulated genes | CD45- |
| MF | GO:0070003 | threonine-type peptidase activity | 1.06E-03 | 1.02E-02 | Psmb9/Psmb8 | 2 | aPD-L1+aTGF-β vs PBS | Up-regulated genes | CD45- |
| MF | GO:0030545 | receptor regulator activity | 1.22E-03 | 1.06E-02 | Cxcl10/Cxcl9/Cxc | 5 | aPD-L1+aTGF-β vs PBS | Up-regulated genes | CD45- |
| MF | GO:0042605 | peptide antigen binding | 1.45E-03 | 1.16E-02 | H2-D1/H2-K1 | 2 | aPD-L1+aTGF-β vs PBS | Up-regulated genes | CD45- |
| MF | GO:0005126 | cytokine receptor binding | 2.12E-03 | 1.56E-02 | Cxcl10/Cxcl9/Cxc | 4 | aPD-L1+aTGF-β vs PBS | Up-regulated genes | CD45- |
| MF | GO:0001664 | G protein-coupled receptor binding | 2.27E-03 | 1.56E-02 | Cxcl10/Cxcl9/Cxc | 4 | aPD-L1+aTGF-β vs PBS | Up-regulated genes | CD45- |
| MF | GO:0000979 | RNA polymerase II core promoter sequence-spe | 4.01E-03 | 2.41E-02 | Fos/Egr1 | 2 | aPD-L1+aTGF-β vs PBS | Up-regulated genes | CD45- |
| MF | GO:0003823 | antigen binding | 4.01E-03 | 2.41E-02 | H2-D1/H2-K1 | 2 | aPD-L1+aTGF-β vs PBS | Up-regulated genes | CD45- |
| MF | GO:0001046 | core promoter sequence-specific DNA binding | 6.84E-03 | 3.87E-02 | Fos/Egr1 | 2 | aPD-L1+aTGF-β vs PBS | Up-regulated genes | CD45- |
| MF | GO:0050840 | extracellular matrix binding | 9.34E-03 | 4.65E-02 | Spp1/Dcn | 2 | aPD-L1+aTGF-β vs PBS | Up-regulated genes | CD45- |
| MF | GO:0001047 | core promoter binding | 9.67E-03 | 4.65E-02 | Fos/Egr1 | 2 | aPD-L1+aTGF-β vs PBS | Up-regulated genes | CD45- |
| MF | GO:0001228 | DNA-binding transcription activator activity, RNA | 9.70E-03 | 4.65E-02 | Irf1/Fosb/Fos/Egr | 4 | aPD-L1+aTGF-β vs PBS | Up-regulated genes | CD45- |
| BP | GO:0044403 | symbiont process | 6.01E-09 | 7.26E-06 | Gsn/P4hb/Fn1/Cx | 11 | aPD-L1+aTGF-β vs PBS | Down-regulated gen | CD45+ |
| BP | GO:0044419 | interspecies interaction between organisms | 9.70E-09 | 7.26E-06 | Gsn/P4hb/Fn1/Cx | 11 | aPD-L1+aTGF-β vs PBS | Down-regulated gen | CD45+ |
| BP | GO:0043903 | regulation of symbiosis, encompassing mutualis | 1.06E-08 | 7.26E-06 | Gsn/P4hb/Cxcr4/ | 9 | aPD-L1+aTGF-β vs PBS | Down-regulated gen | CD45+ |
| BP | GO:0050792 | regulation of viral process | 4.91E-08 | 2.53E-05 | Gsn/P4hb/Cxcr4/ | 8 | aPD-L1+aTGF-β vs PBS | Down-regulated gen | CD45+ |
| BP | GO:0071248 | cellular response to metal ion | 1.04E-07 | 4.27E-05 | Fosb/Gsn/Fn1/Lg | 7 | aPD-L1+aTGF-β vs PBS | Down-regulated gen | CD45+ |
| BP | GO:0043900 | regulation of multi-organism process | 1.46E-07 | 5.00E-05 | Gsn/P4hb/Grn/Cx | 10 | aPD-L1+aTGF-β vs PBS | Down-regulated gen | CD45+ |
| BP | GO:0010038 | response to metal ion | 2.92E-07 | 8.01E-05 | Fosb/Gsn/Fn1/Ga | 8 | aPD-L1+aTGF-β vs PBS | Down-regulated gen | CD45+ |
| BP | GO:0071241 | cellular response to inorganic substance | 3.11E-07 | 8.01E-05 | Fosb/Gsn/Fn1/Lg | 7 | aPD-L1+aTGF-β vs PBS | Down-regulated gen | CD45+ |
| BP | GO:0071277 | cellular response to calcium ion | 1.08E-06 | 2.47E-04 | Fosb/Lgmn/Junb/ | 5 | aPD-L1+aTGF-β vs PBS | Down-regulated gen | CD45+ |
| BP | GO:0060326 | cell chemotaxis | 1.37E-06 | 2.81E-04 | Ccr2/Dusp1/Cxcr | 8 | aPD-L1+aTGF-β vs PBS | Down-regulated gen | CD45+ |
| BP | GO:0016032 | viral process | 1.69E-06 | 3.16E-04 | Gsn/P4hb/Cxcr4/ | 8 | aPD-L1+aTGF-β vs PBS | Down-regulated gen | CD45+ |
| BP | GO:0009611 | response to wounding | 2.40E-06 | 3.84E-04 | Fn1/Ccr2/Grn/Cd | 9 | aPD-L1+aTGF-β vs PBS | Down-regulated gen | CD45+ |
| BP | GO:0001933 | negative regulation of protein phosphorylation | 2.63E-06 | 3.84E-04 | Dusp1/Cdkn1a/D | 9 | aPD-L1+aTGF-β vs PBS | Down-regulated gen | CD45+ |
| BP | GO:0051702 | interaction with symbiont | 2.70E-06 | 3.84E-04 | Fn1/Jun/Arg1/Pf4 | 5 | aPD-L1+aTGF-β vs PBS | Down-regulated gen | CD45+ |
| BP | GO:0030595 | leukocyte chemotaxis | 2.80E-06 | 3.84E-04 | Ccr2/Dusp1/Lgmr | 7 | aPD-L1+aTGF-β vs PBS | Down-regulated gen | CD45+ |
| BP | GO:0048514 | blood vessel morphogenesis | 4.83E-06 | 6.21E-04 | Fn1/Ccr2/Grn/Cxc | 10 | aPD-L1+aTGF-β vs PBS | Down-regulated gen | CD45+ |
| BP | GO:0009991 | response to extracellular stimulus | 5.20E-06 | 6.30E-04 | Cdkn1a/Gatm/Ldl | 8 | aPD-L1+aTGF-β vs PBS | Down-regulated gen | CD45+ |
| BP | GO:0042326 | negative regulation of phosphorylation | 5.85E-06 | 6.69E-04 | Dusp1/Cdkn1a/D | 9 | aPD-L1+aTGF-β vs PBS | Down-regulated gen | CD45+ |
| BP | GO:0002683 | negative regulation of immune system process | 8.05E-06 | 8.72E-04 | Il7r/Ccr2/Dusp1/G | 9 | aPD-L1+aTGF-β vs PBS | Down-regulated gen | CD45+ |
| BP | GO:0051348 | negative regulation of transferase activity | 8.66E-06 | 8.92E-04 | Dusp1/Rpl5/Cdkn | 7 | aPD-L1+aTGF-β vs PBS | Down-regulated gen | CD45+ |
| BP | GO:0043902 | positive regulation of multi-organism process | 1.09E-05 | 1.07E-03 | P4hb/Grn/Jun/Lg | 6 | aPD-L1+aTGF-β vs PBS | Down-regulated gen | CD45+ |
| BP | GO:0006935 | chemotaxis | 1.18E-05 | 1.10E-03 | Fn1/Ccr2/Dusp1/ | 9 | aPD-L1+aTGF-β vs PBS | Down-regulated gen | CD45+ |
| BP | GO:0042330 | taxis | 1.24E-05 | 1.11E-03 | Fn1/Ccr2/Dusp1/ | 9 | aPD-L1+aTGF-β vs PBS | Down-regulated gen | CD45+ |
| BP | GO:0051592 | response to calcium ion | 1.39E-05 | 1.19E-03 | Fosb/Lgmn/Junb/ | 5 | aPD-L1+aTGF-β vs PBS | Down-regulated gen | CD45+ |
| BP | GO:0070848 | response to growth factor | 1.46E-05 | 1.21E-03 | Dab2/Lgmn/Pmep | 9 | aPD-L1+aTGF-β vs PBS | Down-regulated gen | CD45+ |
| BP | GO:0002548 | monocyte chemotaxis | 1.54E-05 | 1.22E-03 | Ccr2/Dusp1/Lgmr | 4 | aPD-L1+aTGF-β vs PBS | Down-regulated gen | CD45+ |
| BP | GO:0010035 | response to inorganic substance | 1.87E-05 | 1.41E-03 | Fosb/Gsn/Fn1/Ga | 8 | aPD-L1+aTGF-β vs PBS | Down-regulated gen | CD45+ |
| BP | GO:0030099 | myeloid cell differentiation | 2.03E-05 | 1.41E-03 | Dab2/Zfp36/Junb | 8 | aPD-L1+aTGF-β vs PBS | Down-regulated gen | CD45+ |
| BP | GO:0030335 | positive regulation of cell migration | 2.11E-05 | 1.41E-03 | Fn1/Ccr2/Grn/Cxc | 9 | aPD-L1+aTGF-β vs PBS | Down-regulated gen | CD45+ |
| BP | GO:0010563 | negative regulation of phosphorus metabolic pro | 2.30E-05 | 1.41E-03 | Dusp1/Cdkn1a/D | 9 | aPD-L1+aTGF-β vs PBS | Down-regulated gen | CD45+ |
| BP | GO:0045936 | negative regulation of phosphate metabolic proc | 2.30E-05 | 1.41E-03 | Dusp1/Cdkn1a/D | 9 | aPD-L1+aTGF-β vs PBS | Down-regulated gen | CD45+ |
| BP | GO:0032370 | positive regulation of lipid transport | 2.35E-05 | 1.41E-03 | Dab2/Nfkbia/Spp | 4 | aPD-L1+aTGF-β vs PBS | Down-regulated gen | CD45+ |
| BP | GO:0035994 | response to muscle stretch | 2.38E-05 | 1.41E-03 | Jun/Nfkbia/Fos | 3 | aPD-L1+aTGF-β vs PBS | Down-regulated gen | CD45+ |
| BP | GO:0032103 | positive regulation of response to external stimul | 2.41E-05 | 1.41E-03 | Fn1/Ccr2/Grn/Cxc | 7 | aPD-L1+aTGF-β vs PBS | Down-regulated gen | CD45+ |
| BP | GO:1903706 | regulation of hemopoiesis | 2.48E-05 | 1.41E-03 | Il7r/Ccr2/Zfp36/Ju | 8 | aPD-L1+aTGF-β vs PBS | Down-regulated gen | CD45+ |
| BP | GO:2000147 | positive regulation of cell motility | 2.50E-05 | 1.41E-03 | Fn1/Ccr2/Grn/Cxc | 9 | aPD-L1+aTGF-β vs PBS | Down-regulated gen | CD45+ |
| BP | GO:0002704 | negative regulation of leukocyte mediated immu | 2.54E-05 | 1.41E-03 | Il7r/Ccr2/Smad7/ | 4 | aPD-L1+aTGF-β vs PBS | Down-regulated gen | CD45+ |
| BP | GO:0097529 | myeloid leukocyte migration | 2.96E-05 | 1.50E-03 | Ccr2/Dusp1/Lgmr | 6 | aPD-L1+aTGF-β vs PBS | Down-regulated gen | CD45+ |
| BP | GO:0009612 | response to mechanical stimulus | 3.03E-05 | 1.50E-03 | Cxcr4/Junb/Jun/N | 5 | aPD-L1+aTGF-β vs PBS | Down-regulated gen | CD45+ |
| BP | GO:0002521 | leukocyte differentiation | 3.04E-05 | 1.50E-03 | Il7r/Ccr2/Junb/Jur | 9 | aPD-L1+aTGF-β vs PBS | Down-regulated gen | CD45+ |
| BP | GO:0051272 | positive regulation of cellular component movem | 3.09E-05 | 1.50E-03 | Fn1/Ccr2/Grn/Cxc | 9 | aPD-L1+aTGF-β vs PBS | Down-regulated gen | CD45+ |
| BP | GO:0050865 | regulation of cell activation | 3.19E-05 | 1.50E-03 | Gsn/Il7r/Ccr2/Grn | 9 | aPD-L1+aTGF-β vs PBS | Down-regulated gen | CD45+ |
| BP | GO:0007611 | learning or memory | 3.25E-05 | 1.50E-03 | Btg2/Gatm/Lgmn/ | 6 | aPD-L1+aTGF-β vs PBS | Down-regulated gen | CD45+ |
| BP | GO:0002710 | negative regulation of T cell mediated immunity | 3.37E-05 | 1.50E-03 | Il7r/Smad7/Arg1 | 3 | aPD-L1+aTGF-β vs PBS | Down-regulated gen | CD45+ |
| BP | GO:0019221 | cytokine-mediated signaling pathway | 3.38E-05 | 1.50E-03 | Ccr2/Cxcr4/Egr1/I | 7 | aPD-L1+aTGF-β vs PBS | Down-regulated gen | CD45+ |
| BP | GO:1990868 | response to chemokine | 3.43E-05 | 1.50E-03 | Dusp1/Cxcr4/Ccl | 4 | aPD-L1+aTGF-β vs PBS | Down-regulated gen | CD45+ |
| BP | GO:1990869 | cellular response to chemokine | 3.43E-05 | 1.50E-03 | Dusp1/Cxcr4/Ccl | 4 | aPD-L1+aTGF-β vs PBS | Down-regulated gen | CD45+ |
| BP | GO:0071276 | cellular response to cadmium ion | 3.96E-05 | 1.70E-03 | Gsn/Jun/Fos | 3 | aPD-L1+aTGF-β vs PBS | Down-regulated gen | CD45+ |

| | | | | | | | | | |
|---|---|---|---|---|---|---|---|---|---|
| BP | GO:0045862 | positive regulation of proteolysis | 4.64E-05 | 1.95E-03 | Gsn/Grn/Dab2/Lg | 7 | aPD-L1+aTGF-β vs PBS | Down-regulated gen | CD45+ |
| BP | GO:0051851 | modification by host of symbiont morphology or | 5.17E-05 | 2.10E-03 | Jun/Arg1/Pf4/Apo | 4 | aPD-L1+aTGF-β vs PBS | Down-regulated gen | CD45+ |
| BP | GO:0002698 | negative regulation of immune effector process | 5.23E-05 | 2.10E-03 | Il7r/Ccr2/Grn/Sma | 5 | aPD-L1+aTGF-β vs PBS | Down-regulated gen | CD45+ |
| BP | GO:0090025 | regulation of monocyte chemotaxis | 5.32E-05 | 2.10E-03 | Ccr2/Dusp1/Lgmr | 3 | aPD-L1+aTGF-β vs PBS | Down-regulated gen | CD45+ |
| BP | GO:1901214 | regulation of neuron death | 5.40E-05 | 2.10E-03 | Btg2/Grn/Lgmn/Ju | 7 | aPD-L1+aTGF-β vs PBS | Down-regulated gen | CD45+ |
| BP | GO:0050900 | leukocyte migration | 5.64E-05 | 2.12E-03 | Ccr2/Dusp1/Lgmr | 7 | aPD-L1+aTGF-β vs PBS | Down-regulated gen | CD45+ |
| BP | GO:0034341 | response to interferon-gamma | 5.65E-05 | 2.12E-03 | Gsn/Cxcl16/Bst2/ | 5 | aPD-L1+aTGF-β vs PBS | Down-regulated gen | CD45+ |
| BP | GO:0050890 | cognition | 6.30E-05 | 2.32E-03 | Btg2/Gatm/Lgmn/ | 6 | aPD-L1+aTGF-β vs PBS | Down-regulated gen | CD45+ |
| BP | GO:0043523 | regulation of neuron apoptotic process | 7.20E-05 | 2.60E-03 | Btg2/Grn/Lgmn/Ju | 6 | aPD-L1+aTGF-β vs PBS | Down-regulated gen | CD45+ |
| BP | GO:1905954 | positive regulation of lipid localization | 7.49E-05 | 2.66E-03 | Dab2/Nfkbia/Spp | 4 | aPD-L1+aTGF-β vs PBS | Down-regulated gen | CD45+ |
| BP | GO:0042110 | T cell activation | 7.86E-05 | 2.71E-03 | Gsn/Il7r/Ccr2/Cxc | 8 | aPD-L1+aTGF-β vs PBS | Down-regulated gen | CD45+ |
| BP | GO:0046596 | regulation of viral entry into host cell | 7.89E-05 | 2.71E-03 | Gsn/P4hb/Lgals1 | 3 | aPD-L1+aTGF-β vs PBS | Down-regulated gen | CD45+ |
| BP | GO:0071674 | mononuclear cell migration | 8.40E-05 | 2.84E-03 | Ccr2/Dusp1/Lgmr | 4 | aPD-L1+aTGF-β vs PBS | Down-regulated gen | CD45+ |
| BP | GO:0071363 | cellular response to growth factor stimulus | 8.81E-05 | 2.93E-03 | Dab2/Lgmn/Pmep | 8 | aPD-L1+aTGF-β vs PBS | Down-regulated gen | CD45+ |
| BP | GO:0050777 | negative regulation of immune response | 9.11E-05 | 2.98E-03 | Il7r/Ccr2/Grn/Sma | 5 | aPD-L1+aTGF-β vs PBS | Down-regulated gen | CD45+ |
| BP | GO:0043524 | negative regulation of neuron apoptotic process | 9.43E-05 | 3.04E-03 | Btg2/Grn/Lgmn/Ju | 5 | aPD-L1+aTGF-β vs PBS | Down-regulated gen | CD45+ |
| BP | GO:0070997 | neuron death | 9.93E-05 | 3.15E-03 | Btg2/Grn/Lgmn/Ju | 7 | aPD-L1+aTGF-β vs PBS | Down-regulated gen | CD45+ |
| BP | GO:0007179 | transforming growth factor beta receptor signalin | 1.01E-04 | 3.15E-03 | Dab2/Pmepa1/Ju | 5 | aPD-L1+aTGF-β vs PBS | Down-regulated gen | CD45+ |
| BP | GO:0032496 | response to lipopolysaccharide | 1.05E-04 | 3.23E-03 | Zfp36/Junb/Jun/C | 6 | aPD-L1+aTGF-β vs PBS | Down-regulated gen | CD45+ |
| BP | GO:0034612 | response to tumor necrosis factor | 1.08E-04 | 3.27E-03 | Pid1/Zfp36/Cxcl1 | 5 | aPD-L1+aTGF-β vs PBS | Down-regulated gen | CD45+ |
| BP | GO:0009617 | response to bacterium | 1.21E-04 | 3.57E-03 | Grn/Zfp36/Junb/J | 8 | aPD-L1+aTGF-β vs PBS | Down-regulated gen | CD45+ |
| BP | GO:0007178 | transmembrane receptor protein serine/threonin | 1.24E-04 | 3.57E-03 | Dab2/Pmepa1/Ju | 6 | aPD-L1+aTGF-β vs PBS | Down-regulated gen | CD45+ |
| BP | GO:0070372 | regulation of ERK1 and ERK2 cascade | 1.24E-04 | 3.57E-03 | Fn1/Dusp1/Dab2/ | 6 | aPD-L1+aTGF-β vs PBS | Down-regulated gen | CD45+ |
| BP | GO:0051402 | neuron apoptotic process | 1.27E-04 | 3.57E-03 | Btg2/Grn/Lgmn/Ju | 6 | aPD-L1+aTGF-β vs PBS | Down-regulated gen | CD45+ |
| BP | GO:0002709 | regulation of T cell mediated immunity | 1.29E-04 | 3.57E-03 | Il7r/Ccr2/Smad7/ | 4 | aPD-L1+aTGF-β vs PBS | Down-regulated gen | CD45+ |
| BP | GO:0048524 | positive regulation of viral process | 1.29E-04 | 3.57E-03 | P4hb/Jun/Lgals1/ | 4 | aPD-L1+aTGF-β vs PBS | Down-regulated gen | CD45+ |
| BP | GO:0014070 | response to organic cyclic compound | 1.30E-04 | 3.57E-03 | Grn/Dab2/Zfp36/J | 8 | aPD-L1+aTGF-β vs PBS | Down-regulated gen | CD45+ |
| BP | GO:0002694 | regulation of leukocyte activation | 1.32E-04 | 3.58E-03 | Gsn/Il7r/Ccr2/Grn | 8 | aPD-L1+aTGF-β vs PBS | Down-regulated gen | CD45+ |
| BP | GO:0002724 | regulation of T cell cytokine production | 1.38E-04 | 3.63E-03 | Ccr2/Smad7/Arg1 | 3 | aPD-L1+aTGF-β vs PBS | Down-regulated gen | CD45+ |
| BP | GO:0046686 | response to cadmium ion | 1.38E-04 | 3.63E-03 | Gsn/Jun/Fos | 3 | aPD-L1+aTGF-β vs PBS | Down-regulated gen | CD45+ |
| BP | GO:0002237 | response to molecule of bacterial origin | 1.56E-04 | 3.99E-03 | Zfp36/Junb/Jun/C | 6 | aPD-L1+aTGF-β vs PBS | Down-regulated gen | CD45+ |
| BP | GO:0032368 | regulation of lipid transport | 1.57E-04 | 3.99E-03 | Dab2/Nfkbia/Spp | 4 | aPD-L1+aTGF-β vs PBS | Down-regulated gen | CD45+ |
| BP | GO:0051817 | modification of morphology or physiology of othe | 1.57E-04 | 3.99E-03 | Jun/Arg1/Pf4/Apo | 4 | aPD-L1+aTGF-β vs PBS | Down-regulated gen | CD45+ |
| BP | GO:0070371 | ERK1 and ERK2 cascade | 1.67E-04 | 4.18E-03 | Fn1/Dusp1/Dab2/ | 6 | aPD-L1+aTGF-β vs PBS | Down-regulated gen | CD45+ |
| BP | GO:1902107 | positive regulation of leukocyte differentiation | 1.69E-04 | 4.18E-03 | Il7r/Ccr2/Jun/Fos/ | 5 | aPD-L1+aTGF-β vs PBS | Down-regulated gen | CD45+ |
| BP | GO:0045785 | positive regulation of cell adhesion | 1.75E-04 | 4.28E-03 | Il7r/Fn1/Ccr2/Dab | 7 | aPD-L1+aTGF-β vs PBS | Down-regulated gen | CD45+ |
| BP | GO:0048525 | negative regulation of viral process | 1.81E-04 | 4.38E-03 | Gsn/Zfp36/Jun/Bs | 4 | aPD-L1+aTGF-β vs PBS | Down-regulated gen | CD45+ |
| BP | GO:0071396 | cellular response to lipid | 2.40E-04 | 5.76E-03 | Dab2/Pid1/Zfp36/ | 7 | aPD-L1+aTGF-β vs PBS | Down-regulated gen | CD45+ |
| BP | GO:1902105 | regulation of leukocyte differentiation | 2.45E-04 | 5.80E-03 | Il7r/Ccr2/Jun/Sma | 6 | aPD-L1+aTGF-β vs PBS | Down-regulated gen | CD45+ |
| BP | GO:0051249 | regulation of lymphocyte activation | 2.48E-04 | 5.81E-03 | Gsn/Il7r/Ccr2/Cdk | 7 | aPD-L1+aTGF-β vs PBS | Down-regulated gen | CD45+ |
| BP | GO:0042060 | wound healing | 2.55E-04 | 5.91E-03 | Fn1/Ccr2/Cdkn1a | 6 | aPD-L1+aTGF-β vs PBS | Down-regulated gen | CD45+ |
| BP | GO:0031667 | response to nutrient levels | 2.71E-04 | 6.19E-03 | Cdkn1a/Gatm/Zfp | 6 | aPD-L1+aTGF-β vs PBS | Down-regulated gen | CD45+ |
| BP | GO:0071560 | cellular response to transforming growth factor b | 2.74E-04 | 6.19E-03 | Dab2/Pmepa1/Ju | 5 | aPD-L1+aTGF-β vs PBS | Down-regulated gen | CD45+ |
| BP | GO:0019058 | viral life cycle | 2.89E-04 | 6.46E-03 | Gsn/P4hb/Bst2/L | 5 | aPD-L1+aTGF-β vs PBS | Down-regulated gen | CD45+ |
| BP | GO:0050920 | regulation of chemotaxis | 2.96E-04 | 6.48E-03 | Fn1/Ccr2/Dusp1/ | 5 | aPD-L1+aTGF-β vs PBS | Down-regulated gen | CD45+ |
| BP | GO:1901342 | regulation of vasculature development | 2.99E-04 | 6.48E-03 | Ccr2/Grn/Cxcr4/E | 6 | aPD-L1+aTGF-β vs PBS | Down-regulated gen | CD45+ |
| BP | GO:0042035 | regulation of cytokine biosynthetic process | 3.02E-04 | 6.48E-03 | Ccr2/Ltb/Zfp36/E | 4 | aPD-L1+aTGF-β vs PBS | Down-regulated gen | CD45+ |
| BP | GO:0071559 | response to transforming growth factor beta | 3.04E-04 | 6.48E-03 | Dab2/Pmepa1/Ju | 5 | aPD-L1+aTGF-β vs PBS | Down-regulated gen | CD45+ |
| BP | GO:0050863 | regulation of T cell activation | 3.05E-04 | 6.48E-03 | Gsn/Il7r/Ccr2/Sma | 6 | aPD-L1+aTGF-β vs PBS | Down-regulated gen | CD45+ |
| BP | GO:0050866 | negative regulation of cell activation | 3.12E-04 | 6.56E-03 | Ccr2/Grn/Smad7/ | 5 | aPD-L1+aTGF-β vs PBS | Down-regulated gen | CD45+ |
| BP | GO:0050678 | regulation of epithelial cell proliferation | 3.17E-04 | 6.60E-03 | Grn/Dab2/Zfp36/J | 6 | aPD-L1+aTGF-β vs PBS | Down-regulated gen | CD45+ |
| BP | GO:0035821 | modification of morphology or physiology of othe | 3.27E-04 | 6.60E-03 | Jun/Arg1/Pf4/Apo | 4 | aPD-L1+aTGF-β vs PBS | Down-regulated gen | CD45+ |
| BP | GO:0002369 | T cell cytokine production | 3.27E-04 | 6.60E-03 | Ccr2/Smad7/Arg1 | 3 | aPD-L1+aTGF-β vs PBS | Down-regulated gen | CD45+ |
| BP | GO:0060393 | regulation of pathway-restricted SMAD protein p | 3.27E-04 | 6.60E-03 | Dab2/Pmepa1/Sn | 3 | aPD-L1+aTGF-β vs PBS | Down-regulated gen | CD45+ |
| BP | GO:0071901 | negative regulation of protein serine/threonine k | 3.53E-04 | 7.06E-03 | Dusp1/Cdkn1a/D | 4 | aPD-L1+aTGF-β vs PBS | Down-regulated gen | CD45+ |
| BP | GO:0002823 | negative regulation of adaptive immune respons | 3.78E-04 | 7.49E-03 | Il7r/Smad7/Arg1 | 3 | aPD-L1+aTGF-β vs PBS | Down-regulated gen | CD45+ |
| BP | GO:1901216 | positive regulation of neuron death | 3.95E-04 | 7.75E-03 | Grn/Jun/Egr1/Fos | 4 | aPD-L1+aTGF-β vs PBS | Down-regulated gen | CD45+ |
| BP | GO:0002707 | negative regulation of lymphocyte mediated imm | 4.06E-04 | 7.81E-03 | Il7r/Smad7/Arg1 | 3 | aPD-L1+aTGF-β vs PBS | Down-regulated gen | CD45+ |
| BP | GO:0060389 | pathway-restricted SMAD protein phosphorylatic | 4.06E-04 | 7.81E-03 | Dab2/Pmepa1/Sn | 3 | aPD-L1+aTGF-β vs PBS | Down-regulated gen | CD45+ |
| BP | GO:0045637 | regulation of myeloid cell differentiation | 4.22E-04 | 7.98E-03 | Zfp36/Jun/Nfkbia/ | 5 | aPD-L1+aTGF-β vs PBS | Down-regulated gen | CD45+ |
| BP | GO:1901215 | negative regulation of neuron death | 4.22E-04 | 7.98E-03 | Btg2/Grn/Lgmn/Ju | 5 | aPD-L1+aTGF-β vs PBS | Down-regulated gen | CD45+ |
| BP | GO:1903708 | positive regulation of hemopoiesis | 4.33E-04 | 7.99E-03 | Il7r/Ccr2/Jun/Fos/ | 5 | aPD-L1+aTGF-β vs PBS | Down-regulated gen | CD45+ |
| BP | GO:0048247 | lymphocyte chemotaxis | 4.34E-04 | 7.99E-03 | Ccr2/Cxcl16/Ccl6 | 3 | aPD-L1+aTGF-β vs PBS | Down-regulated gen | CD45+ |
| BP | GO:0071675 | regulation of mononuclear cell migration | 4.34E-04 | 7.99E-03 | Ccr2/Dusp1/Lgmr | 3 | aPD-L1+aTGF-β vs PBS | Down-regulated gen | CD45+ |
| BP | GO:1905952 | regulation of lipid localization | 4.41E-04 | 8.02E-03 | Dab2/Nfkbia/Spp | 4 | aPD-L1+aTGF-β vs PBS | Down-regulated gen | CD45+ |
| BP | GO:0001525 | angiogenesis | 4.44E-04 | 8.02E-03 | Fn1/Ccr2/Grn/Cxc | 7 | aPD-L1+aTGF-β vs PBS | Down-regulated gen | CD45+ |
| BP | GO:0042089 | cytokine biosynthetic process | 4.57E-04 | 8.17E-03 | Ccr2/Ltb/Zfp36/E | 4 | aPD-L1+aTGF-β vs PBS | Down-regulated gen | CD45+ |
| BP | GO:0046718 | viral entry into host cell | 4.64E-04 | 8.17E-03 | Gsn/P4hb/Lgals1 | 3 | aPD-L1+aTGF-β vs PBS | Down-regulated gen | CD45+ |
| BP | GO:0070098 | chemokine-mediated signaling pathway | 4.64E-04 | 8.17E-03 | Cxcr4/Ccl6/Pf4 | 3 | aPD-L1+aTGF-β vs PBS | Down-regulated gen | CD45+ |
| BP | GO:0060394 | negative regulation of pathway-restricted SMAD | 4.95E-04 | 8.63E-03 | Pmepa1/Smad7 | 2 | aPD-L1+aTGF-β vs PBS | Down-regulated gen | CD45+ |
| BP | GO:1903050 | regulation of proteolysis involved in cellular prot | 5.11E-04 | 8.83E-03 | Rpl5/Dab2/Hspa1 | 5 | aPD-L1+aTGF-β vs PBS | Down-regulated gen | CD45+ |
| BP | GO:0042107 | cytokine metabolic process | 5.25E-04 | 8.91E-03 | Ccr2/Ltb/Zfp36/E | 4 | aPD-L1+aTGF-β vs PBS | Down-regulated gen | CD45+ |
| BP | GO:0002820 | negative regulation of adaptive immune respons | 5.28E-04 | 8.91E-03 | Il7r/Smad7/Arg1 | 3 | aPD-L1+aTGF-β vs PBS | Down-regulated gen | CD45+ |
| BP | GO:0043618 | regulation of transcription from RNA polymerase | 5.28E-04 | 8.91E-03 | Jun/Egr1/Klf2 | 3 | aPD-L1+aTGF-β vs PBS | Down-regulated gen | CD45+ |
| BP | GO:0002456 | T cell mediated immunity | 5.62E-04 | 9.40E-03 | Il7r/Ccr2/Smad7/ | 4 | aPD-L1+aTGF-β vs PBS | Down-regulated gen | CD45+ |
| BP | GO:0030260 | entry into host cell | 5.97E-04 | 9.68E-03 | Gsn/P4hb/Lgals1 | 3 | aPD-L1+aTGF-β vs PBS | Down-regulated gen | CD45+ |
| BP | GO:0044409 | entry into host | 5.97E-04 | 9.68E-03 | Gsn/P4hb/Lgals1 | 3 | aPD-L1+aTGF-β vs PBS | Down-regulated gen | CD45+ |
| BP | GO:0051806 | entry into cell of other organism involved in symb | 5.97E-04 | 9.68E-03 | Gsn/P4hb/Lgals1 | 3 | aPD-L1+aTGF-β vs PBS | Down-regulated gen | CD45+ |
| BP | GO:0051828 | entry into other organism involved in symbiotic i | 5.97E-04 | 9.68E-03 | Gsn/P4hb/Lgals1 | 3 | aPD-L1+aTGF-β vs PBS | Down-regulated gen | CD45+ |
| BP | GO:0007610 | behavior | 6.38E-04 | 1.03E-02 | Btg2/Grn/Gatm/Lg | 7 | aPD-L1+aTGF-β vs PBS | Down-regulated gen | CD45+ |
| BP | GO:0034754 | cellular hormone metabolic process | 6.72E-04 | 1.06E-02 | Dab2/Egr1/Spp1 | 3 | aPD-L1+aTGF-β vs PBS | Down-regulated gen | CD45+ |
| BP | GO:0060395 | SMAD protein signal transduction | 6.72E-04 | 1.06E-02 | Dab2/Jun/Fos | 3 | aPD-L1+aTGF-β vs PBS | Down-regulated gen | CD45+ |
| BP | GO:0050921 | positive regulation of chemotaxis | 6.83E-04 | 1.06E-02 | Fn1/Ccr2/Cxcr4/L | 4 | aPD-L1+aTGF-β vs PBS | Down-regulated gen | CD45+ |
| BP | GO:0031331 | positive regulation of cellular catabolic process | 6.93E-04 | 1.06E-02 | Btg2/Dab2/Zfp36/ | 6 | aPD-L1+aTGF-β vs PBS | Down-regulated gen | CD45+ |
| BP | GO:0043620 | regulation of DNA-templated transcription in res | 7.11E-04 | 1.06E-02 | Jun/Egr1/Klf2 | 3 | aPD-L1+aTGF-β vs PBS | Down-regulated gen | CD45+ |
| BP | GO:0050810 | regulation of steroid biosynthetic process | 7.11E-04 | 1.06E-02 | Dab2/Egr1/Apoe | 3 | aPD-L1+aTGF-β vs PBS | Down-regulated gen | CD45+ |
| BP | GO:0006700 | C21-steroid hormone biosynthetic process | 7.22E-04 | 1.06E-02 | Dab2/Egr1 | 2 | aPD-L1+aTGF-β vs PBS | Down-regulated gen | CD45+ |
| BP | GO:0010566 | regulation of ketone biosynthetic process | 7.22E-04 | 1.06E-02 | Dab2/Egr1 | 2 | aPD-L1+aTGF-β vs PBS | Down-regulated gen | CD45+ |
| BP | GO:0010875 | positive regulation of cholesterol efflux | 7.22E-04 | 1.06E-02 | Nfkbia/Apoe | 2 | aPD-L1+aTGF-β vs PBS | Down-regulated gen | CD45+ |

| | | | | | | | | | |
|---|---|---|---|---|---|---|---|---|---|
| BP | GO:0051852 | disruption by host of symbiont cells | 7.22E-04 | 1.06E-02 | Arg1/Pf4 | 2 | aPD-L1+aTGF-β vs PBS | Down-regulated gen | CD45+ |
| BP | GO:0051873 | killing by host of symbiont cells | 7.22E-04 | 1.06E-02 | Arg1/Pf4 | 2 | aPD-L1+aTGF-β vs PBS | Down-regulated gen | CD45+ |
| BP | GO:0060213 | positive regulation of nuclear-transcribed mRNA | 7.22E-04 | 1.06E-02 | Btg2/Zfp36 | 2 | aPD-L1+aTGF-β vs PBS | Down-regulated gen | CD45+ |
| BP | GO:0050673 | epithelial cell proliferation | 7.76E-04 | 1.13E-02 | Grn/Dab2/Zfp36/J | 6 | aPD-L1+aTGF-β vs PBS | Down-regulated gen | CD45+ |
| BP | GO:0071356 | cellular response to tumor necrosis factor | 7.97E-04 | 1.15E-02 | Pid1/Zfp36/Nfkbia | 4 | aPD-L1+aTGF-β vs PBS | Down-regulated gen | CD45+ |
| BP | GO:1903364 | positive regulation of cellular protein catabolic p | 7.97E-04 | 1.15E-02 | Dab2/Hspa1b/Sm | 4 | aPD-L1+aTGF-β vs PBS | Down-regulated gen | CD45+ |
| BP | GO:1903900 | regulation of viral life cycle | 8.21E-04 | 1.17E-02 | Gsn/P4hb/Bst2/Lg | 4 | aPD-L1+aTGF-β vs PBS | Down-regulated gen | CD45+ |
| BP | GO:0048146 | positive regulation of fibroblast proliferation | 8.38E-04 | 1.18E-02 | Fn1/Cdkn1a/Jun | 3 | aPD-L1+aTGF-β vs PBS | Down-regulated gen | CD45+ |
| BP | GO:0001667 | ameboidal-type cell migration | 8.39E-04 | 1.18E-02 | Fn1/Grn/Cxcr4/Lg | 6 | aPD-L1+aTGF-β vs PBS | Down-regulated gen | CD45+ |
| BP | GO:0006706 | steroid catabolic process | 8.52E-04 | 1.19E-02 | Spp1/Apoe | 2 | aPD-L1+aTGF-β vs PBS | Down-regulated gen | CD45+ |
| BP | GO:0090594 | inflammatory response to wounding | 8.52E-04 | 1.19E-02 | Ccr2/Grn | 2 | aPD-L1+aTGF-β vs PBS | Down-regulated gen | CD45+ |
| BP | GO:0035914 | skeletal muscle cell differentiation | 8.84E-04 | 1.22E-02 | Btg2/Egr1/Fos | 3 | aPD-L1+aTGF-β vs PBS | Down-regulated gen | CD45+ |
| BP | GO:1903362 | regulation of cellular protein catabolic process | 8.94E-04 | 1.23E-02 | Rpl5/Dab2/Hspa1 | 5 | aPD-L1+aTGF-β vs PBS | Down-regulated gen | CD45+ |
| BP | GO:0002829 | negative regulation of type 2 immune response | 9.92E-04 | 1.33E-02 | Ccr2/Arg1 | 2 | aPD-L1+aTGF-β vs PBS | Down-regulated gen | CD45+ |
| BP | GO:0046886 | positive regulation of hormone biosynthetic proc | 9.92E-04 | 1.33E-02 | Dab2/Egr1 | 2 | aPD-L1+aTGF-β vs PBS | Down-regulated gen | CD45+ |
| BP | GO:0060211 | regulation of nuclear-transcribed mRNA poly(A) | 9.92E-04 | 1.33E-02 | Btg2/Zfp36 | 2 | aPD-L1+aTGF-β vs PBS | Down-regulated gen | CD45+ |
| BP | GO:1990000 | amyloid fibril formation | 9.92E-04 | 1.33E-02 | Gsn/Apoe | 2 | aPD-L1+aTGF-β vs PBS | Down-regulated gen | CD45+ |
| BP | GO:0001558 | regulation of cell growth | 1.01E-03 | 1.34E-02 | Fn1/Cdkn1a/Cxcr | 6 | aPD-L1+aTGF-β vs PBS | Down-regulated gen | CD45+ |
| BP | GO:0002763 | positive regulation of myeloid leukocyte different | 1.03E-03 | 1.36E-02 | Jun/Fos/Pf4 | 3 | aPD-L1+aTGF-β vs PBS | Down-regulated gen | CD45+ |
| BP | GO:0001818 | negative regulation of cytokine production | 1.11E-03 | 1.45E-02 | Fn1/Zfp36/Smad7 | 5 | aPD-L1+aTGF-β vs PBS | Down-regulated gen | CD45+ |
| BP | GO:1903034 | regulation of response to wounding | 1.12E-03 | 1.46E-02 | Grn/Cdkn1a/Cxcr | 4 | aPD-L1+aTGF-β vs PBS | Down-regulated gen | CD45+ |
| BP | GO:0043407 | negative regulation of MAP kinase activity | 1.13E-03 | 1.46E-02 | Dusp1/Dusp2/Apo | 3 | aPD-L1+aTGF-β vs PBS | Down-regulated gen | CD45+ |
| BP | GO:0051818 | disruption of cells of other organism involved in s | 1.14E-03 | 1.46E-02 | Arg1/Pf4 | 2 | aPD-L1+aTGF-β vs PBS | Down-regulated gen | CD45+ |
| BP | GO:0051883 | killing of cells in other organism involved in syml | 1.14E-03 | 1.46E-02 | Arg1/Pf4 | 2 | aPD-L1+aTGF-β vs PBS | Down-regulated gen | CD45+ |
| BP | GO:0001659 | temperature homeostasis | 1.15E-03 | 1.47E-02 | Ccr2/Gatm/Cxcr4/ | 4 | aPD-L1+aTGF-β vs PBS | Down-regulated gen | CD45+ |
| BP | GO:0015850 | organic hydroxy compound transport | 1.19E-03 | 1.50E-02 | Dab2/Nfkbia/Spp1 | 4 | aPD-L1+aTGF-β vs PBS | Down-regulated gen | CD45+ |
| BP | GO:0042108 | positive regulation of cytokine biosynthetic proce | 1.25E-03 | 1.56E-02 | Ccr2/Ltb/Egr1 | 3 | aPD-L1+aTGF-β vs PBS | Down-regulated gen | CD45+ |
| BP | GO:1903707 | negative regulation of hemopoiesis | 1.28E-03 | 1.59E-02 | Zfp36/Smad7/Nfk | 4 | aPD-L1+aTGF-β vs PBS | Down-regulated gen | CD45+ |
| BP | GO:2000058 | regulation of ubiquitin-dependent protein catabo | 1.28E-03 | 1.59E-02 | Rpl5/Dab2/Hspa1 | 4 | aPD-L1+aTGF-β vs PBS | Down-regulated gen | CD45+ |
| BP | GO:0071498 | cellular response to fluid shear stress | 1.30E-03 | 1.61E-02 | Klf2/Spp1 | 2 | aPD-L1+aTGF-β vs PBS | Down-regulated gen | CD45+ |
| BP | GO:0045765 | regulation of angiogenesis | 1.44E-03 | 1.74E-02 | Ccr2/Grn/Cxcr4/K | 5 | aPD-L1+aTGF-β vs PBS | Down-regulated gen | CD45+ |
| BP | GO:0000188 | inactivation of MAPK activity | 1.47E-03 | 1.74E-02 | Dusp1/Dusp2 | 2 | aPD-L1+aTGF-β vs PBS | Down-regulated gen | CD45+ |
| BP | GO:0010743 | regulation of macrophage derived foam cell diffe | 1.47E-03 | 1.74E-02 | Nfkbia/Pf4 | 2 | aPD-L1+aTGF-β vs PBS | Down-regulated gen | CD45+ |
| BP | GO:0010818 | T cell chemotaxis | 1.47E-03 | 1.74E-02 | Ccr2/Cxcl16 | 2 | aPD-L1+aTGF-β vs PBS | Down-regulated gen | CD45+ |
| BP | GO:0010874 | regulation of cholesterol efflux | 1.47E-03 | 1.74E-02 | Nfkbia/Apoe | 2 | aPD-L1+aTGF-β vs PBS | Down-regulated gen | CD45+ |
| BP | GO:0090026 | positive regulation of monocyte chemotaxis | 1.47E-03 | 1.74E-02 | Ccr2/Lgmn | 2 | aPD-L1+aTGF-β vs PBS | Down-regulated gen | CD45+ |
| BP | GO:1900153 | positive regulation of nuclear-transcribed mRNA | 1.47E-03 | 1.74E-02 | Btg2/Zfp36 | 2 | aPD-L1+aTGF-β vs PBS | Down-regulated gen | CD45+ |
| BP | GO:0043409 | negative regulation of MAPK cascade | 1.54E-03 | 1.81E-02 | Dusp1/Dab2/Dus | 4 | aPD-L1+aTGF-β vs PBS | Down-regulated gen | CD45+ |
| BP | GO:0043525 | positive regulation of neuron apoptotic process | 1.55E-03 | 1.81E-02 | Grn/Jun/Egr1 | 3 | aPD-L1+aTGF-β vs PBS | Down-regulated gen | CD45+ |
| BP | GO:0071347 | cellular response to interleukin-1 | 1.55E-03 | 1.81E-02 | Fn1/Egr1/Ccl6 | 3 | aPD-L1+aTGF-β vs PBS | Down-regulated gen | CD45+ |
| BP | GO:0043901 | negative regulation of multi-organism process | 1.61E-03 | 1.87E-02 | Gsn/Zfp36/Jun/Bs | 4 | aPD-L1+aTGF-β vs PBS | Down-regulated gen | CD45+ |
| BP | GO:0051444 | negative regulation of ubiquitin-protein transfera | 1.65E-03 | 1.90E-02 | Rpl5/Smad7 | 2 | aPD-L1+aTGF-β vs PBS | Down-regulated gen | CD45+ |
| BP | GO:0009896 | positive regulation of catabolic process | 1.68E-03 | 1.92E-02 | Btg2/Dab2/Zfp36/ | 6 | aPD-L1+aTGF-β vs PBS | Down-regulated gen | CD45+ |
| BP | GO:0032102 | negative regulation of response to external stimu | 1.71E-03 | 1.94E-02 | Dusp1/Grn/Cdkn1 | 5 | aPD-L1+aTGF-β vs PBS | Down-regulated gen | CD45+ |
| BP | GO:0019218 | regulation of steroid metabolic process | 1.76E-03 | 1.99E-02 | Dab2/Egr1/Apoe | 3 | aPD-L1+aTGF-β vs PBS | Down-regulated gen | CD45+ |
| BP | GO:0046651 | lymphocyte proliferation | 1.77E-03 | 1.99E-02 | Il7r/Ccr2/Cdkn1a/ | 5 | aPD-L1+aTGF-β vs PBS | Down-regulated gen | CD45+ |
| BP | GO:1903037 | regulation of leukocyte cell-cell adhesion | 1.80E-03 | 2.01E-02 | Il7r/Ccr2/Smad7/L | 5 | aPD-L1+aTGF-β vs PBS | Down-regulated gen | CD45+ |
| BP | GO:0032870 | cellular response to hormone stimulus | 1.82E-03 | 2.01E-02 | Dab2/Pid1/Zfp36/ | 6 | aPD-L1+aTGF-β vs PBS | Down-regulated gen | CD45+ |
| BP | GO:0032943 | mononuclear cell proliferation | 1.83E-03 | 2.01E-02 | Il7r/Ccr2/Cdkn1a/ | 5 | aPD-L1+aTGF-β vs PBS | Down-regulated gen | CD45+ |
| BP | GO:0008207 | C21-steroid hormone metabolic process | 1.84E-03 | 2.01E-02 | Dab2/Egr1 | 2 | aPD-L1+aTGF-β vs PBS | Down-regulated gen | CD45+ |
| BP | GO:0032352 | positive regulation of hormone metabolic proces | 1.84E-03 | 2.01E-02 | Dab2/Egr1 | 2 | aPD-L1+aTGF-β vs PBS | Down-regulated gen | CD45+ |
| BP | GO:1900151 | regulation of nuclear-transcribed mRNA cataboli | 1.84E-03 | 2.01E-02 | Btg2/Zfp36 | 2 | aPD-L1+aTGF-β vs PBS | Down-regulated gen | CD45+ |
| BP | GO:0030307 | positive regulation of cell growth | 1.95E-03 | 2.12E-02 | Fn1/Cxcr4/Cxcl16 | 4 | aPD-L1+aTGF-β vs PBS | Down-regulated gen | CD45+ |
| BP | GO:0002706 | regulation of lymphocyte mediated immunity | 2.00E-03 | 2.15E-02 | Il7r/Ccr2/Smad7/ | 4 | aPD-L1+aTGF-β vs PBS | Down-regulated gen | CD45+ |
| BP | GO:0002822 | regulation of adaptive immune response based | 2.04E-03 | 2.15E-02 | Il7r/Ccr2/Smad7/ | 4 | aPD-L1+aTGF-β vs PBS | Down-regulated gen | CD45+ |
| BP | GO:0071222 | cellular response to lipopolysaccharide | 2.04E-03 | 2.15E-02 | Zfp36/Cxcl16/Nfk | 4 | aPD-L1+aTGF-β vs PBS | Down-regulated gen | CD45+ |
| BP | GO:0032373 | positive regulation of sterol transport | 2.04E-03 | 2.15E-02 | Nfkbia/Apoe | 2 | aPD-L1+aTGF-β vs PBS | Down-regulated gen | CD45+ |
| BP | GO:0032376 | positive regulation of cholesterol transport | 2.04E-03 | 2.15E-02 | Nfkbia/Apoe | 2 | aPD-L1+aTGF-β vs PBS | Down-regulated gen | CD45+ |
| BP | GO:0032897 | negative regulation of viral transcription | 2.04E-03 | 2.15E-02 | Zfp36/Jun | 2 | aPD-L1+aTGF-β vs PBS | Down-regulated gen | CD45+ |
| BP | GO:0002718 | regulation of cytokine production involved in imn | 2.06E-03 | 2.15E-02 | Ccr2/Smad7/Arg1 | 3 | aPD-L1+aTGF-β vs PBS | Down-regulated gen | CD45+ |
| BP | GO:0002695 | negative regulation of leukocyte activation | 2.14E-03 | 2.22E-02 | Ccr2/Grn/Smad7/ | 4 | aPD-L1+aTGF-β vs PBS | Down-regulated gen | CD45+ |
| BP | GO:0032436 | positive regulation of proteasomal ubiquitin-dep | 2.22E-03 | 2.28E-02 | Dab2/Hspa1b/Sm | 3 | aPD-L1+aTGF-β vs PBS | Down-regulated gen | CD45+ |
| BP | GO:0120162 | positive regulation of cold-induced thermogenes | 2.22E-03 | 2.28E-02 | Ccr2/Gatm/Cxcr4 | 3 | aPD-L1+aTGF-β vs PBS | Down-regulated gen | CD45+ |
| BP | GO:0002719 | negative regulation of cytokine production involv | 2.25E-03 | 2.28E-02 | Smad7/Arg1 | 2 | aPD-L1+aTGF-β vs PBS | Down-regulated gen | CD45+ |
| BP | GO:0010742 | macrophage derived foam cell differentiation | 2.25E-03 | 2.28E-02 | Nfkbia/Pf4 | 2 | aPD-L1+aTGF-β vs PBS | Down-regulated gen | CD45+ |
| BP | GO:1903077 | negative regulation of protein localization to plas | 2.25E-03 | 2.28E-02 | Dab2/Pid1 | 2 | aPD-L1+aTGF-β vs PBS | Down-regulated gen | CD45+ |
| BP | GO:0016049 | cell growth | 2.26E-03 | 2.28E-02 | Fn1/Cdkn1a/Cxcr | 6 | aPD-L1+aTGF-β vs PBS | Down-regulated gen | CD45+ |
| BP | GO:0062012 | regulation of small molecule metabolic process | 2.37E-03 | 2.37E-02 | Dab2/Pid1/Ppp1r | 5 | aPD-L1+aTGF-β vs PBS | Down-regulated gen | CD45+ |
| BP | GO:0070661 | leukocyte proliferation | 2.37E-03 | 2.37E-02 | Il7r/Ccr2/Cdkn1a/ | 5 | aPD-L1+aTGF-β vs PBS | Down-regulated gen | CD45+ |
| BP | GO:0071219 | cellular response to molecule of bacterial origin | 2.39E-03 | 2.38E-02 | Zfp36/Cxcl16/Nfk | 4 | aPD-L1+aTGF-β vs PBS | Down-regulated gen | CD45+ |
| BP | GO:0034405 | response to fluid shear stress | 2.47E-03 | 2.41E-02 | Klf2/Spp1 | 2 | aPD-L1+aTGF-β vs PBS | Down-regulated gen | CD45+ |
| BP | GO:0045940 | positive regulation of steroid metabolic process | 2.47E-03 | 2.41E-02 | Dab2/Apoe | 2 | aPD-L1+aTGF-β vs PBS | Down-regulated gen | CD45+ |
| BP | GO:0071677 | positive regulation of mononuclear cell migratior | 2.47E-03 | 2.41E-02 | Ccr2/Lgmn | 2 | aPD-L1+aTGF-β vs PBS | Down-regulated gen | CD45+ |
| BP | GO:0090077 | foam cell differentiation | 2.47E-03 | 2.41E-02 | Nfkbia/Pf4 | 2 | aPD-L1+aTGF-β vs PBS | Down-regulated gen | CD45+ |
| BP | GO:0061136 | regulation of proteasomal protein catabolic proc | 2.55E-03 | 2.47E-02 | Dab2/Hspa1b/Sm | 4 | aPD-L1+aTGF-β vs PBS | Down-regulated gen | CD45+ |
| BP | GO:0010810 | regulation of cell-substrate adhesion | 2.60E-03 | 2.52E-02 | Fn1/Dab2/Lgals1 | 4 | aPD-L1+aTGF-β vs PBS | Down-regulated gen | CD45+ |
| BP | GO:0000289 | nuclear-transcribed mRNA poly(A) tail shortenin | 2.70E-03 | 2.59E-02 | Btg2/Zfp36 | 2 | aPD-L1+aTGF-β vs PBS | Down-regulated gen | CD45+ |
| BP | GO:1904376 | negative regulation of protein localization to cell | 2.70E-03 | 2.59E-02 | Dab2/Pid1 | 2 | aPD-L1+aTGF-β vs PBS | Down-regulated gen | CD45+ |
| BP | GO:0060840 | artery development | 2.75E-03 | 2.62E-02 | Cxcr4/Smad7/Apo | 3 | aPD-L1+aTGF-β vs PBS | Down-regulated gen | CD45+ |
| BP | GO:0007159 | leukocyte cell-cell adhesion | 2.76E-03 | 2.62E-02 | Il7r/Ccr2/Smad7/L | 5 | aPD-L1+aTGF-β vs PBS | Down-regulated gen | CD45+ |
| BP | GO:0002819 | regulation of adaptive immune response | 2.89E-03 | 2.73E-02 | Il7r/Ccr2/Smad7/ | 4 | aPD-L1+aTGF-β vs PBS | Down-regulated gen | CD45+ |
| BP | GO:0070555 | response to interleukin-1 | 2.94E-03 | 2.75E-02 | Fn1/Egr1/Ccl6 | 3 | aPD-L1+aTGF-β vs PBS | Down-regulated gen | CD45+ |
| BP | GO:0032801 | receptor catabolic process | 2.94E-03 | 2.75E-02 | Lgmn/Apoe | 2 | aPD-L1+aTGF-β vs PBS | Down-regulated gen | CD45+ |
| BP | GO:0006469 | negative regulation of protein kinase activity | 2.95E-03 | 2.75E-02 | Dusp1/Cdkn1a/D | 4 | aPD-L1+aTGF-β vs PBS | Down-regulated gen | CD45+ |
| BP | GO:0017015 | regulation of transforming growth factor beta rec | 3.04E-03 | 2.79E-02 | Dab2/Pmepa1/Sn | 3 | aPD-L1+aTGF-β vs PBS | Down-regulated gen | CD45+ |
| BP | GO:0045638 | negative regulation of myeloid cell differentiatior | 3.04E-03 | 2.79E-02 | Zfp36/Nfkbia/Pf4 | 3 | aPD-L1+aTGF-β vs PBS | Down-regulated gen | CD45+ |
| BP | GO:0072676 | lymphocyte migration | 3.04E-03 | 2.79E-02 | Ccr2/Cxcl16/Ccl6 | 3 | aPD-L1+aTGF-β vs PBS | Down-regulated gen | CD45+ |
| BP | GO:0031348 | negative regulation of defense response | 3.07E-03 | 2.80E-02 | Grn/Zfp36/Arg1/A | 4 | aPD-L1+aTGF-β vs PBS | Down-regulated gen | CD45+ |
| BP | GO:0048638 | regulation of developmental growth | 3.11E-03 | 2.80E-02 | Fn1/Cdkn1a/Cxcr | 5 | aPD-L1+aTGF-β vs PBS | Down-regulated gen | CD45+ |

| | | | | | | | | | |
|---|---|---|---|---|---|---|---|---|---|
| BP | GO:0050727 | regulation of inflammatory response | 3.11E-03 | 2.80E-02 | Ccr2/Grn/Zfp36/N | 5 | aPD-L1+aTGF-β vs PBS | Down-regulated gen | CD45+ |
| BP | GO:0030593 | neutrophil chemotaxis | 3.14E-03 | 2.80E-02 | Ccl6/Pf4/Spp1 | 3 | aPD-L1+aTGF-β vs PBS | Down-regulated gen | CD45+ |
| BP | GO:0090090 | negative regulation of canonical Wnt signaling p | 3.14E-03 | 2.80E-02 | Dab2/Egr1/Apoe | 3 | aPD-L1+aTGF-β vs PBS | Down-regulated gen | CD45+ |
| BP | GO:1903844 | regulation of cellular response to transforming g | 3.14E-03 | 2.80E-02 | Dab2/Pmepa1/Sn | 3 | aPD-L1+aTGF-β vs PBS | Down-regulated gen | CD45+ |
| BP | GO:2000060 | positive regulation of ubiquitin-dependent protei | 3.14E-03 | 2.80E-02 | Dab2/Hspa1b/Sm | 3 | aPD-L1+aTGF-β vs PBS | Down-regulated gen | CD45+ |
| BP | GO:0045639 | positive regulation of myeloid cell differentiation | 3.35E-03 | 2.96E-02 | Jun/Fos/Pf4 | 3 | aPD-L1+aTGF-β vs PBS | Down-regulated gen | CD45+ |
| BP | GO:0048145 | regulation of fibroblast proliferation | 3.35E-03 | 2.96E-02 | Fn1/Cdkn1a/Jun | 3 | aPD-L1+aTGF-β vs PBS | Down-regulated gen | CD45+ |
| BP | GO:1901652 | response to peptide | 3.38E-03 | 2.98E-02 | Gatm/Pid1/Lgmn/ | 5 | aPD-L1+aTGF-β vs PBS | Down-regulated gen | CD45+ |
| BP | GO:0033344 | cholesterol efflux | 3.45E-03 | 3.00E-02 | Nfkbia/Apoe | 2 | aPD-L1+aTGF-β vs PBS | Down-regulated gen | CD45+ |
| BP | GO:0071480 | cellular response to gamma radiation | 3.45E-03 | 3.00E-02 | Cdkn1a/Egr1 | 2 | aPD-L1+aTGF-β vs PBS | Down-regulated gen | CD45+ |
| BP | GO:0048144 | fibroblast proliferation | 3.46E-03 | 3.00E-02 | Fn1/Cdkn1a/Jun | 3 | aPD-L1+aTGF-β vs PBS | Down-regulated gen | CD45+ |
| BP | GO:0031668 | cellular response to extracellular stimulus | 3.52E-03 | 3.05E-02 | Cdkn1a/Ldha/Jun | 4 | aPD-L1+aTGF-β vs PBS | Down-regulated gen | CD45+ |
| BP | GO:0071216 | cellular response to biotic stimulus | 3.66E-03 | 3.15E-02 | Zfp36/Cxcl16/Nfkl | 4 | aPD-L1+aTGF-β vs PBS | Down-regulated gen | CD45+ |
| BP | GO:0016525 | negative regulation of angiogenesis | 3.68E-03 | 3.15E-02 | Ccr2/Klf2/Pf4 | 3 | aPD-L1+aTGF-β vs PBS | Down-regulated gen | CD45+ |
| BP | GO:0007616 | long-term memory | 3.72E-03 | 3.15E-02 | Egr1/Apoe | 2 | aPD-L1+aTGF-β vs PBS | Down-regulated gen | CD45+ |
| BP | GO:0048535 | lymph node development | 3.72E-03 | 3.15E-02 | Il7r/Ltb | 2 | aPD-L1+aTGF-β vs PBS | Down-regulated gen | CD45+ |
| BP | GO:1905476 | negative regulation of protein localization to mer | 3.72E-03 | 3.15E-02 | Dab2/Pid1 | 2 | aPD-L1+aTGF-β vs PBS | Down-regulated gen | CD45+ |
| BP | GO:0007613 | memory | 3.79E-03 | 3.18E-02 | Lgmn/Egr1/Apoe | 3 | aPD-L1+aTGF-β vs PBS | Down-regulated gen | CD45+ |
| BP | GO:2000379 | positive regulation of reactive oxygen species m | 3.79E-03 | 3.18E-02 | Cdkn1a/Pid1/Klf2 | 3 | aPD-L1+aTGF-β vs PBS | Down-regulated gen | CD45+ |
| BP | GO:0002440 | production of molecular mediator of immune res | 3.80E-03 | 3.18E-02 | Il7r/Ccr2/Smad7/ | 4 | aPD-L1+aTGF-β vs PBS | Down-regulated gen | CD45+ |
| BP | GO:0008202 | steroid metabolic process | 3.87E-03 | 3.21E-02 | Dab2/Egr1/Spp1/ | 4 | aPD-L1+aTGF-β vs PBS | Down-regulated gen | CD45+ |
| BP | GO:0010632 | regulation of epithelial cell migration | 3.87E-03 | 3.21E-02 | Grn/Lgmn/Jun/Ap | 4 | aPD-L1+aTGF-β vs PBS | Down-regulated gen | CD45+ |
| BP | GO:0071407 | cellular response to organic cyclic compound | 3.89E-03 | 3.22E-02 | Dab2/Zfp36/Egr1/ | 5 | aPD-L1+aTGF-β vs PBS | Down-regulated gen | CD45+ |
| BP | GO:0002367 | cytokine production involved in immune respons | 3.91E-03 | 3.22E-02 | Ccr2/Smad7/Arg1 | 3 | aPD-L1+aTGF-β vs PBS | Down-regulated gen | CD45+ |
| BP | GO:0031640 | killing of cells of other organism | 4.00E-03 | 3.23E-02 | Arg1/Pf4 | 2 | aPD-L1+aTGF-β vs PBS | Down-regulated gen | CD45+ |
| BP | GO:0032371 | regulation of sterol transport | 4.00E-03 | 3.23E-02 | Nfkbia/Apoe | 2 | aPD-L1+aTGF-β vs PBS | Down-regulated gen | CD45+ |
| BP | GO:0032374 | regulation of cholesterol transport | 4.00E-03 | 3.23E-02 | Nfkbia/Apoe | 2 | aPD-L1+aTGF-β vs PBS | Down-regulated gen | CD45+ |
| BP | GO:0044364 | disruption of cells of other organism | 4.00E-03 | 3.23E-02 | Arg1/Pf4 | 2 | aPD-L1+aTGF-β vs PBS | Down-regulated gen | CD45+ |
| BP | GO:0046885 | regulation of hormone biosynthetic process | 4.00E-03 | 3.23E-02 | Dab2/Egr1 | 2 | aPD-L1+aTGF-β vs PBS | Down-regulated gen | CD45+ |
| BP | GO:0045732 | positive regulation of protein catabolic process | 4.02E-03 | 3.23E-02 | Dab2/Hspa1b/Sm | 4 | aPD-L1+aTGF-β vs PBS | Down-regulated gen | CD45+ |
| BP | GO:0033673 | negative regulation of kinase activity | 4.09E-03 | 3.28E-02 | Dusp1/Cdkn1a/D | 4 | aPD-L1+aTGF-β vs PBS | Down-regulated gen | CD45+ |
| BP | GO:1901800 | positive regulation of proteasomal protein catab | 4.15E-03 | 3.30E-02 | Dab2/Hspa1b/Sm | 3 | aPD-L1+aTGF-β vs PBS | Down-regulated gen | CD45+ |
| BP | GO:2000181 | negative regulation of blood vessel morphogene | 4.15E-03 | 3.30E-02 | Ccr2/Klf2/Pf4 | 3 | aPD-L1+aTGF-β vs PBS | Down-regulated gen | CD45+ |
| BP | GO:0002573 | myeloid leukocyte differentiation | 4.24E-03 | 3.36E-02 | Junb/Jun/Fos/Pf4 | 4 | aPD-L1+aTGF-β vs PBS | Down-regulated gen | CD45+ |
| BP | GO:0002688 | regulation of leukocyte chemotaxis | 4.65E-03 | 3.67E-02 | Ccr2/Dusp1/Lgmr | 3 | aPD-L1+aTGF-β vs PBS | Down-regulated gen | CD45+ |
| BP | GO:0002701 | negative regulation of production of molecular m | 4.89E-03 | 3.82E-02 | Smad7/Arg1 | 2 | aPD-L1+aTGF-β vs PBS | Down-regulated gen | CD45+ |
| BP | GO:0002828 | regulation of type 2 immune response | 4.89E-03 | 3.82E-02 | Ccr2/Arg1 | 2 | aPD-L1+aTGF-β vs PBS | Down-regulated gen | CD45+ |
| BP | GO:0006694 | steroid biosynthetic process | 4.91E-03 | 3.82E-02 | Dab2/Egr1/Apoe | 3 | aPD-L1+aTGF-β vs PBS | Down-regulated gen | CD45+ |
| BP | GO:0007612 | learning | 4.91E-03 | 3.82E-02 | Btg2/Lgmn/Jun | 3 | aPD-L1+aTGF-β vs PBS | Down-regulated gen | CD45+ |
| BP | GO:0061041 | regulation of wound healing | 5.05E-03 | 3.90E-02 | Cdkn1a/Cxcr4/Ap | 3 | aPD-L1+aTGF-β vs PBS | Down-regulated gen | CD45+ |
| BP | GO:0030098 | lymphocyte differentiation | 5.06E-03 | 3.90E-02 | Il7r/Ccr2/Smad7/E | 5 | aPD-L1+aTGF-β vs PBS | Down-regulated gen | CD45+ |
| BP | GO:0002703 | regulation of leukocyte mediated immunity | 5.15E-03 | 3.94E-02 | Il7r/Ccr2/Smad7/ | 4 | aPD-L1+aTGF-β vs PBS | Down-regulated gen | CD45+ |
| BP | GO:0010975 | regulation of neuron projection development | 5.16E-03 | 3.94E-02 | Fn1/Grn/Cxcr4/Da | 6 | aPD-L1+aTGF-β vs PBS | Down-regulated gen | CD45+ |
| BP | GO:0071346 | cellular response to interferon-gamma | 5.19E-03 | 3.94E-02 | Gsn/Ccl6/Arg1 | 3 | aPD-L1+aTGF-β vs PBS | Down-regulated gen | CD45+ |
| BP | GO:1990266 | neutrophil migration | 5.19E-03 | 3.94E-02 | Ccl6/Pf4/Spp1 | 3 | aPD-L1+aTGF-β vs PBS | Down-regulated gen | CD45+ |
| BP | GO:1902895 | positive regulation of pri-miRNA transcription by | 5.20E-03 | 3.94E-02 | Jun/Fos | 2 | aPD-L1+aTGF-β vs PBS | Down-regulated gen | CD45+ |
| BP | GO:0010811 | positive regulation of cell-substrate adhesion | 5.33E-03 | 4.02E-02 | Fn1/Dab2/Spp1 | 3 | aPD-L1+aTGF-β vs PBS | Down-regulated gen | CD45+ |
| BP | GO:0001936 | regulation of endothelial cell proliferation | 5.47E-03 | 4.05E-02 | Jun/Arg1/Apoe | 3 | aPD-L1+aTGF-β vs PBS | Down-regulated gen | CD45+ |
| BP | GO:0030178 | negative regulation of Wnt signaling pathway | 5.47E-03 | 4.05E-02 | Dab2/Egr1/Apoe | 3 | aPD-L1+aTGF-β vs PBS | Down-regulated gen | CD45+ |
| BP | GO:1901343 | negative regulation of vasculature development | 5.47E-03 | 4.05E-02 | Ccr2/Klf2/Pf4 | 3 | aPD-L1+aTGF-β vs PBS | Down-regulated gen | CD45+ |
| BP | GO:0022407 | regulation of cell-cell adhesion | 5.52E-03 | 4.05E-02 | Il7r/Ccr2/Smad7/L | 5 | aPD-L1+aTGF-β vs PBS | Down-regulated gen | CD45+ |
| BP | GO:0036230 | granulocyte activation | 5.52E-03 | 4.05E-02 | Ccr2/Grn | 2 | aPD-L1+aTGF-β vs PBS | Down-regulated gen | CD45+ |
| BP | GO:0042181 | ketone biosynthetic process | 5.52E-03 | 4.05E-02 | Dab2/Egr1 | 2 | aPD-L1+aTGF-β vs PBS | Down-regulated gen | CD45+ |
| BP | GO:0042832 | defense response to protozoan | 5.52E-03 | 4.05E-02 | Arg1/Pf4 | 2 | aPD-L1+aTGF-β vs PBS | Down-regulated gen | CD45+ |
| BP | GO:1902930 | regulation of alcohol biosynthetic process | 5.52E-03 | 4.05E-02 | Dab2/Apoe | 2 | aPD-L1+aTGF-β vs PBS | Down-regulated gen | CD45+ |
| BP | GO:0051701 | interaction with host | 5.61E-03 | 4.07E-02 | Gsn/P4hb/Lgals1 | 3 | aPD-L1+aTGF-β vs PBS | Down-regulated gen | CD45+ |
| BP | GO:0071621 | granulocyte chemotaxis | 5.61E-03 | 4.07E-02 | Ccl6/Pf4/Spp1 | 3 | aPD-L1+aTGF-β vs PBS | Down-regulated gen | CD45+ |
| BP | GO:1903052 | positive regulation of proteolysis involved in cell | 5.61E-03 | 4.07E-02 | Dab2/Hspa1b/Sm | 3 | aPD-L1+aTGF-β vs PBS | Down-regulated gen | CD45+ |
| BP | GO:0031099 | regeneration | 5.76E-03 | 4.16E-02 | Grn/Cdkn1a/Jun | 3 | aPD-L1+aTGF-β vs PBS | Down-regulated gen | CD45+ |
| BP | GO:0022409 | positive regulation of cell-cell adhesion | 5.88E-03 | 4.22E-02 | Il7r/Ccr2/Smad7/L | 4 | aPD-L1+aTGF-β vs PBS | Down-regulated gen | CD45+ |
| BP | GO:0071496 | cellular response to external stimulus | 5.88E-03 | 4.22E-02 | Cdkn1a/Ldha/Jun | 4 | aPD-L1+aTGF-β vs PBS | Down-regulated gen | CD45+ |
| BP | GO:0042176 | regulation of protein catabolic process | 6.08E-03 | 4.35E-02 | Rpl5/Dab2/Hspa1 | 5 | aPD-L1+aTGF-β vs PBS | Down-regulated gen | CD45+ |
| BP | GO:0001562 | response to protozoan | 6.20E-03 | 4.38E-02 | Arg1/Pf4 | 2 | aPD-L1+aTGF-β vs PBS | Down-regulated gen | CD45+ |
| BP | GO:0045622 | regulation of T-helper cell differentiation | 6.20E-03 | 4.38E-02 | Ccr2/Smad7 | 2 | aPD-L1+aTGF-β vs PBS | Down-regulated gen | CD45+ |
| BP | GO:0061844 | antimicrobial humoral immune response mediat | 6.20E-03 | 4.38E-02 | Hist1h2bc/Pf4 | 2 | aPD-L1+aTGF-β vs PBS | Down-regulated gen | CD45+ |
| BP | GO:0002761 | regulation of myeloid leukocyte differentiation | 6.21E-03 | 4.38E-02 | Jun/Fos/Pf4 | 3 | aPD-L1+aTGF-β vs PBS | Down-regulated gen | CD45+ |
| BP | GO:0006959 | humoral immune response | 6.53E-03 | 4.56E-02 | Ccr2/Hist1h2bc/P | 3 | aPD-L1+aTGF-β vs PBS | Down-regulated gen | CD45+ |
| BP | GO:0106106 | cold-induced thermogenesis | 6.53E-03 | 4.56E-02 | Ccr2/Gatm/Cxcr4 | 3 | aPD-L1+aTGF-β vs PBS | Down-regulated gen | CD45+ |
| BP | GO:0120161 | regulation of cold-induced thermogenesis | 6.53E-03 | 4.56E-02 | Ccr2/Gatm/Cxcr4 | 3 | aPD-L1+aTGF-β vs PBS | Down-regulated gen | CD45+ |
| BP | GO:0002697 | regulation of immune effector process | 6.61E-03 | 4.57E-02 | Il7r/Ccr2/Grn/Sma | 5 | aPD-L1+aTGF-β vs PBS | Down-regulated gen | CD45+ |
| BP | GO:0048871 | multicellular organismal homeostasis | 6.61E-03 | 4.57E-02 | Ccr2/Gatm/Cxcr4/ | 5 | aPD-L1+aTGF-β vs PBS | Down-regulated gen | CD45+ |
| BP | GO:0061564 | axon development | 6.61E-03 | 4.57E-02 | Fn1/Grn/Cxcr4/Ju | 5 | aPD-L1+aTGF-β vs PBS | Down-regulated gen | CD45+ |
| BP | GO:0002366 | leukocyte activation involved in immune respons | 6.68E-03 | 4.60E-02 | Ccr2/Grn/Smad7/ | 4 | aPD-L1+aTGF-β vs PBS | Down-regulated gen | CD45+ |
| BP | GO:0032350 | regulation of hormone metabolic process | 6.91E-03 | 4.71E-02 | Dab2/Egr1 | 2 | aPD-L1+aTGF-β vs PBS | Down-regulated gen | CD45+ |
| BP | GO:0042092 | type 2 immune response | 6.91E-03 | 4.71E-02 | Ccr2/Arg1 | 2 | aPD-L1+aTGF-β vs PBS | Down-regulated gen | CD45+ |
| BP | GO:1900271 | regulation of long-term synaptic potentiation | 6.91E-03 | 4.71E-02 | Lgmn/Apoe | 2 | aPD-L1+aTGF-β vs PBS | Down-regulated gen | CD45+ |
| BP | GO:0002263 | cell activation involved in immune response | 7.11E-03 | 4.83E-02 | Ccr2/Grn/Smad7/ | 4 | aPD-L1+aTGF-β vs PBS | Down-regulated gen | CD45+ |
| BP | GO:0032434 | regulation of proteasomal ubiquitin-dependent p | 7.19E-03 | 4.85E-02 | Dab2/Hspa1b/Sm | 3 | aPD-L1+aTGF-β vs PBS | Down-regulated gen | CD45+ |
| BP | GO:0042445 | hormone metabolic process | 7.19E-03 | 4.85E-02 | Dab2/Egr1/Spp1 | 3 | aPD-L1+aTGF-β vs PBS | Down-regulated gen | CD45+ |
| BP | GO:0001774 | microglial cell activation | 7.28E-03 | 4.88E-02 | Grn/Jun | 2 | aPD-L1+aTGF-β vs PBS | Down-regulated gen | CD45+ |
| BP | GO:0002269 | leukocyte activation involved in inflammatory res | 7.28E-03 | 4.88E-02 | Grn/Jun | 2 | aPD-L1+aTGF-β vs PBS | Down-regulated gen | CD45+ |
| BP | GO:0001935 | endothelial cell proliferation | 7.36E-03 | 4.92E-02 | Jun/Arg1/Apoe | 3 | aPD-L1+aTGF-β vs PBS | Down-regulated gen | CD45+ |
| BP | GO:0006869 | lipid transport | 7.44E-03 | 4.96E-02 | Dab2/Nfkbia/Spp | 4 | aPD-L1+aTGF-β vs PBS | Down-regulated gen | CD45+ |
| MF | GO:0031625 | ubiquitin protein ligase binding | 3.98E-06 | 2.76E-04 | Rpl5/Cdkn1a/Cxc | 8 | aPD-L1+aTGF-β vs PBS | Down-regulated gen | CD45+ |
| MF | GO:0005125 | cytokine activity | 4.56E-06 | 2.76E-04 | Grn/Ltb/Cxcl16/C | 6 | aPD-L1+aTGF-β vs PBS | Down-regulated gen | CD45+ |
| MF | GO:0046332 | SMAD binding | 4.58E-06 | 2.76E-04 | Dab2/Pmepa1/Ju | 5 | aPD-L1+aTGF-β vs PBS | Down-regulated gen | CD45+ |
| MF | GO:0044389 | ubiquitin-like protein ligase binding | 6.30E-06 | 2.85E-04 | Rpl5/Cdkn1a/Cxc | 8 | aPD-L1+aTGF-β vs PBS | Down-regulated gen | CD45+ |
| MF | GO:0005178 | integrin binding | 2.59E-05 | 9.39E-04 | P4hb/Fn1/Dab2/T | 5 | aPD-L1+aTGF-β vs PBS | Down-regulated gen | CD45+ |
| MF | GO:0019957 | C-C chemokine binding | 4.69E-05 | 1.21E-03 | Ccr2/Cxcr4/Zfp36 | 3 | aPD-L1+aTGF-β vs PBS | Down-regulated gen | CD45+ |

| | | | | | | | | | |
|---|---|---|---|---|---|---|---|---|---|
| MF | GO:0070412 | R-SMAD binding | 4.69E-05 | 1.21E-03 | Pmepa1/Jun/Fos | 3 | aPD-L1+aTGF-β vs PBS | Down-regulated gen | CD45+ |
| MF | GO:0019956 | chemokine binding | 1.02E-04 | 2.30E-03 | Ccr2/Cxcr4/Zfp36 | 3 | aPD-L1+aTGF-β vs PBS | Down-regulated gen | CD45+ |
| MF | GO:0048018 | receptor ligand activity | 2.10E-04 | 3.69E-03 | Grn/Ltb/Cxcl16/C | 6 | aPD-L1+aTGF-β vs PBS | Down-regulated gen | CD45+ |
| MF | GO:0008009 | chemokine activity | 2.23E-04 | 3.69E-03 | Cxcl16/Ccl6/Pf4 | 3 | aPD-L1+aTGF-β vs PBS | Down-regulated gen | CD45+ |
| MF | GO:0000981 | DNA-binding transcription factor activity, RNA po | 2.25E-04 | 3.69E-03 | Btg2/Fosb/Ier2/Ju | 8 | aPD-L1+aTGF-β vs PBS | Down-regulated gen | CD45+ |
| MF | GO:0050839 | cell adhesion molecule binding | 3.05E-04 | 4.23E-03 | P4hb/Fn1/Dab2/T | 5 | aPD-L1+aTGF-β vs PBS | Down-regulated gen | CD45+ |
| MF | GO:0030545 | receptor regulator activity | 3.21E-04 | 4.23E-03 | Grn/Ltb/Cxcl16/C | 6 | aPD-L1+aTGF-β vs PBS | Down-regulated gen | CD45+ |
| MF | GO:0001228 | DNA-binding transcription activator activity, RNA | 3.27E-04 | 4.23E-03 | Fosb/Ier2/Junb/Ju | 6 | aPD-L1+aTGF-β vs PBS | Down-regulated gen | CD45+ |
| MF | GO:0050840 | extracellular matrix binding | 4.13E-04 | 4.98E-03 | Tgfbi/Lgals1/Spp | 3 | aPD-L1+aTGF-β vs PBS | Down-regulated gen | CD45+ |
| MF | GO:0042379 | chemokine receptor binding | 4.72E-04 | 5.34E-03 | Cxcl16/Ccl6/Pf4 | 3 | aPD-L1+aTGF-β vs PBS | Down-regulated gen | CD45+ |
| MF | GO:0017017 | MAP kinase tyrosine/serine/threonine phosphata | 7.31E-04 | 7.78E-03 | Dusp1/Dusp2 | 2 | aPD-L1+aTGF-β vs PBS | Down-regulated gen | CD45+ |
| MF | GO:0005126 | cytokine receptor binding | 8.12E-04 | 8.17E-03 | Ltb/Smad7/Cxcl1 | 5 | aPD-L1+aTGF-β vs PBS | Down-regulated gen | CD45+ |
| MF | GO:0033549 | MAP kinase phosphatase activity | 8.62E-04 | 8.21E-03 | Dusp1/Dusp2 | 2 | aPD-L1+aTGF-β vs PBS | Down-regulated gen | CD45+ |
| MF | GO:0016493 | C-C chemokine receptor activity | 2.07E-03 | 1.87E-02 | Ccr2/Cxcr4 | 2 | aPD-L1+aTGF-β vs PBS | Down-regulated gen | CD45+ |
| MF | GO:0001637 | G protein-coupled chemoattractant receptor acti | 2.28E-03 | 1.88E-02 | Ccr2/Cxcr4 | 2 | aPD-L1+aTGF-β vs PBS | Down-regulated gen | CD45+ |
| MF | GO:0004950 | chemokine receptor activity | 2.28E-03 | 1.88E-02 | Ccr2/Cxcr4 | 2 | aPD-L1+aTGF-β vs PBS | Down-regulated gen | CD45+ |
| MF | GO:0004896 | cytokine receptor activity | 2.70E-03 | 2.12E-02 | Il7r/Ccr2/Cxcr4 | 3 | aPD-L1+aTGF-β vs PBS | Down-regulated gen | CD45+ |
| MF | GO:0043394 | proteoglycan binding | 3.76E-03 | 2.84E-02 | Fn1/Apoe | 2 | aPD-L1+aTGF-β vs PBS | Down-regulated gen | CD45+ |
| MF | GO:0008201 | heparin binding | 4.86E-03 | 3.52E-02 | Fn1/Pf4/Apoe | 3 | aPD-L1+aTGF-β vs PBS | Down-regulated gen | CD45+ |
| MF | GO:0001102 | RNA polymerase II activating transcription factor | 5.26E-03 | 3.53E-02 | Jun/Fos | 2 | aPD-L1+aTGF-β vs PBS | Down-regulated gen | CD45+ |
| MF | GO:0008138 | protein tyrosine/serine/threonine phosphatase a | 5.26E-03 | 3.53E-02 | Dusp1/Dusp2 | 2 | aPD-L1+aTGF-β vs PBS | Down-regulated gen | CD45+ |
| MF | GO:0051019 | mitogen-activated protein kinase binding | 5.59E-03 | 3.61E-02 | Dusp1/Dusp2 | 2 | aPD-L1+aTGF-β vs PBS | Down-regulated gen | CD45+ |
| MF | GO:0000979 | RNA polymerase II core promoter sequence-spe | 5.92E-03 | 3.70E-02 | Egr1/Fos | 2 | aPD-L1+aTGF-β vs PBS | Down-regulated gen | CD45+ |
| MF | GO:0019955 | cytokine binding | 6.16E-03 | 3.72E-02 | Ccr2/Cxcr4/Zfp36 | 3 | aPD-L1+aTGF-β vs PBS | Down-regulated gen | CD45+ |
| BP | GO:0046034 | ATP metabolic process | 1.18E-05 | 3.27E-03 | Tpi1/Atp6v1b1/Pg | 6 | aPD-L1+aTGF-β vs PBS | Down-regulated gen | CD45- |
| BP | GO:0001503 | ossification | 1.36E-05 | 3.27E-03 | Atp6v1b1/Ccn1/G | 7 | aPD-L1+aTGF-β vs PBS | Down-regulated gen | CD45- |
| BP | GO:0009126 | purine nucleoside monophosphate metabolic pr | 2.10E-05 | 3.27E-03 | Tpi1/Atp6v1b1/Pg | 6 | aPD-L1+aTGF-β vs PBS | Down-regulated gen | CD45- |
| BP | GO:0009167 | purine ribonucleoside monophosphate metaboli | 2.10E-05 | 3.27E-03 | Tpi1/Atp6v1b1/Pg | 6 | aPD-L1+aTGF-β vs PBS | Down-regulated gen | CD45- |
| BP | GO:0009205 | purine ribonucleoside triphosphate metabolic pr | 2.21E-05 | 3.27E-03 | Tpi1/Atp6v1b1/Pg | 6 | aPD-L1+aTGF-β vs PBS | Down-regulated gen | CD45- |
| BP | GO:0009161 | ribonucleoside monophosphate metabolic proce | 2.32E-05 | 3.27E-03 | Tpi1/Atp6v1b1/Pg | 6 | aPD-L1+aTGF-β vs PBS | Down-regulated gen | CD45- |
| BP | GO:0030198 | extracellular matrix organization | 2.32E-05 | 3.27E-03 | P3h4/Ccn1/Mmp2 | 6 | aPD-L1+aTGF-β vs PBS | Down-regulated gen | CD45- |
| BP | GO:0009199 | ribonucleoside triphosphate metabolic process | 2.43E-05 | 3.27E-03 | Tpi1/Atp6v1b1/Pg | 6 | aPD-L1+aTGF-β vs PBS | Down-regulated gen | CD45- |
| BP | GO:0009144 | purine nucleoside triphosphate metabolic proce | 2.67E-05 | 3.27E-03 | Tpi1/Atp6v1b1/Pg | 6 | aPD-L1+aTGF-β vs PBS | Down-regulated gen | CD45- |
| BP | GO:0009123 | nucleoside monophosphate metabolic process | 2.74E-05 | 3.27E-03 | Tpi1/Atp6v1b1/Pg | 6 | aPD-L1+aTGF-β vs PBS | Down-regulated gen | CD45- |
| BP | GO:0009141 | nucleoside triphosphate metabolic process | 3.68E-05 | 4.00E-03 | Tpi1/Atp6v1b1/Pg | 6 | aPD-L1+aTGF-β vs PBS | Down-regulated gen | CD45- |
| BP | GO:0043062 | extracellular structure organization | 5.18E-05 | 5.17E-03 | P3h4/Ccn1/Mmp2 | 6 | aPD-L1+aTGF-β vs PBS | Down-regulated gen | CD45- |
| BP | GO:0001649 | osteoblast differentiation | 6.22E-05 | 5.72E-03 | Ccn1/Gja1/Cthrc1 | 5 | aPD-L1+aTGF-β vs PBS | Down-regulated gen | CD45- |
| BP | GO:0032963 | collagen metabolic process | 8.24E-05 | 7.04E-03 | P3h4/Mmp23/Col | 4 | aPD-L1+aTGF-β vs PBS | Down-regulated gen | CD45- |
| BP | GO:0006754 | ATP biosynthetic process | 1.38E-04 | 1.10E-02 | Tpi1/Pgk1/COX2/ | 4 | aPD-L1+aTGF-β vs PBS | Down-regulated gen | CD45- |
| BP | GO:0009206 | purine ribonucleoside triphosphate biosynthetic | 1.96E-04 | 1.25E-02 | Tpi1/Pgk1/COX2/ | 4 | aPD-L1+aTGF-β vs PBS | Down-regulated gen | CD45- |
| BP | GO:0009145 | purine nucleoside triphosphate biosynthetic pro | 2.02E-04 | 1.25E-02 | Tpi1/Pgk1/COX2/ | 4 | aPD-L1+aTGF-β vs PBS | Down-regulated gen | CD45- |
| BP | GO:0009150 | purine ribonucleotide metabolic process | 2.13E-04 | 1.25E-02 | Tpi1/Atp6v1b1/Pg | 6 | aPD-L1+aTGF-β vs PBS | Down-regulated gen | CD45- |
| BP | GO:0009201 | ribonucleoside triphosphate biosynthetic proces | 2.16E-04 | 1.25E-02 | Tpi1/Pgk1/COX2/ | 4 | aPD-L1+aTGF-β vs PBS | Down-regulated gen | CD45- |
| BP | GO:0045667 | regulation of osteoblast differentiation | 2.30E-04 | 1.25E-02 | Ccn1/Gja1/Cthrc1 | 4 | aPD-L1+aTGF-β vs PBS | Down-regulated gen | CD45- |
| BP | GO:0009259 | ribonucleotide metabolic process | 2.49E-04 | 1.25E-02 | Tpi1/Atp6v1b1/Pg | 6 | aPD-L1+aTGF-β vs PBS | Down-regulated gen | CD45- |
| BP | GO:0009127 | purine nucleoside monophosphate biosynthetic | 2.69E-04 | 1.25E-02 | Tpi1/Pgk1/COX2/ | 4 | aPD-L1+aTGF-β vs PBS | Down-regulated gen | CD45- |
| BP | GO:0009168 | purine ribonucleoside monophosphate biosynth | 2.69E-04 | 1.25E-02 | Tpi1/Pgk1/COX2/ | 4 | aPD-L1+aTGF-β vs PBS | Down-regulated gen | CD45- |
| BP | GO:0030199 | collagen fibril organization | 2.69E-04 | 1.25E-02 | P3h4/Col1a2/Col | 3 | aPD-L1+aTGF-β vs PBS | Down-regulated gen | CD45- |
| BP | GO:0043589 | skin morphogenesis | 2.78E-04 | 1.25E-02 | Col1a2/Col1a1 | 2 | aPD-L1+aTGF-β vs PBS | Down-regulated gen | CD45- |
| BP | GO:0006163 | purine nucleotide metabolic process | 2.84E-04 | 1.25E-02 | Tpi1/Atp6v1b1/Pg | 6 | aPD-L1+aTGF-β vs PBS | Down-regulated gen | CD45- |
| BP | GO:0009142 | nucleoside triphosphate biosynthetic process | 2.86E-04 | 1.25E-02 | Tpi1/Pgk1/COX2/ | 4 | aPD-L1+aTGF-β vs PBS | Down-regulated gen | CD45- |
| BP | GO:0009156 | ribonucleoside monophosphate biosynthetic pro | 3.04E-04 | 1.25E-02 | Tpi1/Pgk1/COX2/ | 4 | aPD-L1+aTGF-β vs PBS | Down-regulated gen | CD45- |
| BP | GO:0031214 | biomineral tissue development | 3.13E-04 | 1.25E-02 | Ccn1/Gja1/Col1a | 4 | aPD-L1+aTGF-β vs PBS | Down-regulated gen | CD45- |
| BP | GO:0019693 | ribose phosphate metabolic process | 3.15E-04 | 1.25E-02 | Tpi1/Atp6v1b1/Pg | 6 | aPD-L1+aTGF-β vs PBS | Down-regulated gen | CD45- |
| BP | GO:0009124 | nucleoside monophosphate biosynthetic proces | 3.51E-04 | 1.35E-02 | Tpi1/Pgk1/COX2/ | 4 | aPD-L1+aTGF-β vs PBS | Down-regulated gen | CD45- |
| BP | GO:0071417 | cellular response to organonitrogen compound | 3.99E-04 | 1.49E-02 | Actb/Grb10/Prnp/ | 6 | aPD-L1+aTGF-β vs PBS | Down-regulated gen | CD45- |
| BP | GO:0072521 | purine-containing compound metabolic process | 4.44E-04 | 1.61E-02 | Tpi1/Atp6v1b1/Pg | 6 | aPD-L1+aTGF-β vs PBS | Down-regulated gen | CD45- |
| BP | GO:0033690 | positive regulation of osteoblast proliferation | 4.79E-04 | 1.68E-02 | Ccn1/Cthrc1 | 2 | aPD-L1+aTGF-β vs PBS | Down-regulated gen | CD45- |
| BP | GO:0045669 | positive regulation of osteoblast differentiation | 6.12E-04 | 2.09E-02 | Ccn1/Gja1/Cthrc1 | 3 | aPD-L1+aTGF-β vs PBS | Down-regulated gen | CD45- |
| BP | GO:0048771 | tissue remodeling | 6.55E-04 | 2.17E-02 | P3h4/Gja1/Cthrc1 | 4 | aPD-L1+aTGF-β vs PBS | Down-regulated gen | CD45- |
| BP | GO:1901699 | cellular response to nitrogen compound | 6.93E-04 | 2.24E-02 | Actb/Grb10/Prnp/ | 6 | aPD-L1+aTGF-β vs PBS | Down-regulated gen | CD45- |
| BP | GO:0006006 | glucose metabolic process | 7.19E-04 | 2.26E-02 | Tpi1/Pgk1/Grb10/ | 4 | aPD-L1+aTGF-β vs PBS | Down-regulated gen | CD45- |
| BP | GO:0062012 | regulation of small molecule metabolic process | 8.23E-04 | 2.52E-02 | Pgk1/Grb10/Fdps | 5 | aPD-L1+aTGF-β vs PBS | Down-regulated gen | CD45- |
| BP | GO:0009117 | nucleotide metabolic process | 9.82E-04 | 2.89E-02 | Tpi1/Atp6v1b1/Pg | 6 | aPD-L1+aTGF-β vs PBS | Down-regulated gen | CD45- |
| BP | GO:0120162 | positive regulation of cold-induced thermogenes | 9.96E-04 | 2.89E-02 | Gadd45g/Gja1/Gr | 3 | aPD-L1+aTGF-β vs PBS | Down-regulated gen | CD45- |
| BP | GO:0009152 | purine ribonucleotide biosynthetic process | 1.02E-03 | 2.89E-02 | Tpi1/Pgk1/COX2/ | 4 | aPD-L1+aTGF-β vs PBS | Down-regulated gen | CD45- |
| BP | GO:0070208 | protein heterotrimerization | 1.04E-03 | 2.89E-02 | Col1a2/Col1a1 | 2 | aPD-L1+aTGF-β vs PBS | Down-regulated gen | CD45- |
| BP | GO:0006753 | nucleoside phosphate metabolic process | 1.06E-03 | 2.89E-02 | Tpi1/Atp6v1b1/Pg | 6 | aPD-L1+aTGF-β vs PBS | Down-regulated gen | CD45- |
| BP | GO:0006091 | generation of precursor metabolites and energy | 1.10E-03 | 2.93E-02 | Tpi1/Pgk1/Grb10/ | 5 | aPD-L1+aTGF-β vs PBS | Down-regulated gen | CD45- |
| BP | GO:0022898 | regulation of transmembrane transporter activity | 1.16E-03 | 2.99E-02 | Actb/Gja1/Prnp/F | 4 | aPD-L1+aTGF-β vs PBS | Down-regulated gen | CD45- |
| BP | GO:0046849 | bone remodeling | 1.19E-03 | 2.99E-02 | P3h4/Gja1/Cthrc1 | 3 | aPD-L1+aTGF-β vs PBS | Down-regulated gen | CD45- |
| BP | GO:0042692 | muscle cell differentiation | 1.22E-03 | 2.99E-02 | Fdps/Actg1/Tmsb | 5 | aPD-L1+aTGF-β vs PBS | Down-regulated gen | CD45- |
| BP | GO:0009260 | ribonucleotide biosynthetic process | 1.23E-03 | 2.99E-02 | Tpi1/Pgk1/COX2/ | 4 | aPD-L1+aTGF-β vs PBS | Down-regulated gen | CD45- |
| BP | GO:0019318 | hexose metabolic process | 1.30E-03 | 3.05E-02 | Tpi1/Pgk1/Grb10/ | 4 | aPD-L1+aTGF-β vs PBS | Down-regulated gen | CD45- |
| BP | GO:0006164 | purine nucleotide biosynthetic process | 1.33E-03 | 3.05E-02 | Tpi1/Pgk1/COX2/ | 4 | aPD-L1+aTGF-β vs PBS | Down-regulated gen | CD45- |
| BP | GO:0032409 | regulation of transporter activity | 1.33E-03 | 3.05E-02 | Actb/Gja1/Prnp/F | 4 | aPD-L1+aTGF-β vs PBS | Down-regulated gen | CD45- |
| BP | GO:0030278 | regulation of ossification | 1.38E-03 | 3.05E-02 | Ccn1/Gja1/Cthrc1 | 4 | aPD-L1+aTGF-β vs PBS | Down-regulated gen | CD45- |
| BP | GO:0046390 | ribose phosphate biosynthetic process | 1.38E-03 | 3.05E-02 | Tpi1/Pgk1/COX2/ | 4 | aPD-L1+aTGF-β vs PBS | Down-regulated gen | CD45- |
| BP | GO:0072522 | purine-containing compound biosynthetic proce | 1.46E-03 | 3.17E-02 | Tpi1/Pgk1/COX2/ | 4 | aPD-L1+aTGF-β vs PBS | Down-regulated gen | CD45- |
| BP | GO:0006119 | oxidative phosphorylation | 1.51E-03 | 3.22E-02 | Pgk1/COX2/COX | 3 | aPD-L1+aTGF-β vs PBS | Down-regulated gen | CD45- |
| BP | GO:0033688 | regulation of osteoblast proliferation | 1.66E-03 | 3.49E-02 | Ccn1/Cthrc1 | 2 | aPD-L1+aTGF-β vs PBS | Down-regulated gen | CD45- |
| BP | GO:0045778 | positive regulation of ossification | 1.76E-03 | 3.63E-02 | Ccn1/Gja1/Cthrc1 | 3 | aPD-L1+aTGF-β vs PBS | Down-regulated gen | CD45- |
| BP | GO:0005996 | monosaccharide metabolic process | 1.81E-03 | 3.68E-02 | Tpi1/Pgk1/Grb10/ | 4 | aPD-L1+aTGF-β vs PBS | Down-regulated gen | CD45- |
| BP | GO:0016055 | Wnt signaling pathway | 1.90E-03 | 3.77E-02 | Wls/Grb10/Celsr2 | 5 | aPD-L1+aTGF-β vs PBS | Down-regulated gen | CD45- |
| BP | GO:0198738 | cell-cell signaling by wnt | 1.92E-03 | 3.77E-02 | Wls/Grb10/Celsr2 | 5 | aPD-L1+aTGF-β vs PBS | Down-regulated gen | CD45- |
| BP | GO:0030282 | bone mineralization | 1.98E-03 | 3.82E-02 | Ccn1/Gja1/Col1a | 3 | aPD-L1+aTGF-β vs PBS | Down-regulated gen | CD45- |
| BP | GO:0033687 | osteoblast proliferation | 2.11E-03 | 4.00E-02 | Ccn1/Cthrc1 | 2 | aPD-L1+aTGF-β vs PBS | Down-regulated gen | CD45- |
| BP | GO:0048009 | insulin-like growth factor receptor signaling path | 2.60E-03 | 4.86E-02 | Grb10/Igfbp5 | 2 | aPD-L1+aTGF-β vs PBS | Down-regulated gen | CD45- |
| BP | GO:0009165 | nucleotide biosynthetic process | 2.70E-03 | 4.89E-02 | Tpi1/Pgk1/COX2/ | 4 | aPD-L1+aTGF-β vs PBS | Down-regulated gen | CD45- |

| | | | | | | | | | |
|---|---|---|---|---|---|---|---|---|---|
| BP | GO:0034762 | regulation of transmembrane transport | 2.70E-03 | 4.89E-02 | Actb/Gja1/Grb10/l | 5 | aPD-L1+aTGF-β vs PBS | Down-regulated gen | CD45- |
| CC | GO:0031012 | extracellular matrix | 2.40E-06 | 3.82E-04 | Ccn1/Mmp23/Ccc | 8 | aPD-L1+aTGF-β vs PBS | Down-regulated gen | CD45- |
| CC | GO:0043292 | contractile fiber | 5.77E-05 | 4.03E-03 | Myl9/Gja1/Actg1/ | 5 | aPD-L1+aTGF-β vs PBS | Down-regulated gen | CD45- |
| CC | GO:0062023 | collagen-containing extracellular matrix | 7.61E-05 | 4.03E-03 | Ccn1/Ccdc80/Ltb | 6 | aPD-L1+aTGF-β vs PBS | Down-regulated gen | CD45- |
| CC | GO:0044420 | extracellular matrix component | 1.92E-04 | 7.61E-03 | Col1a2/Fbn1/Col | 3 | aPD-L1+aTGF-β vs PBS | Down-regulated gen | CD45- |
| CC | GO:0098644 | complex of collagen trimers | 4.92E-04 | 1.06E-02 | Col1a2/Col1a1 | 2 | aPD-L1+aTGF-β vs PBS | Down-regulated gen | CD45- |
| CC | GO:0044449 | contractile fiber part | 5.26E-04 | 1.06E-02 | Myl9/Actg1/Acta2 | 4 | aPD-L1+aTGF-β vs PBS | Down-regulated gen | CD45- |
| CC | GO:0098871 | postsynaptic actin cytoskeleton | 5.73E-04 | 1.06E-02 | Actb/Actg1 | 2 | aPD-L1+aTGF-β vs PBS | Down-regulated gen | CD45- |
| CC | GO:0005581 | collagen trimer | 5.82E-04 | 1.06E-02 | Col1a2/Cthrc1/Cc | 3 | aPD-L1+aTGF-β vs PBS | Down-regulated gen | CD45- |
| CC | GO:0099571 | postsynaptic cytoskeleton | 6.60E-04 | 1.06E-02 | Actb/Actg1 | 2 | aPD-L1+aTGF-β vs PBS | Down-regulated gen | CD45- |
| CC | GO:0045277 | respiratory chain complex IV | 7.53E-04 | 1.06E-02 | COX2/COX3 | 2 | aPD-L1+aTGF-β vs PBS | Down-regulated gen | CD45- |
| CC | GO:0045121 | membrane raft | 8.38E-04 | 1.06E-02 | Eno1/Actb/Pgk1/C | 5 | aPD-L1+aTGF-β vs PBS | Down-regulated gen | CD45- |
| CC | GO:0098857 | membrane microdomain | 8.51E-04 | 1.06E-02 | Eno1/Actb/Pgk1/C | 5 | aPD-L1+aTGF-β vs PBS | Down-regulated gen | CD45- |
| CC | GO:0098589 | membrane region | 9.75E-04 | 1.06E-02 | Eno1/Actb/Pgk1/C | 5 | aPD-L1+aTGF-β vs PBS | Down-regulated gen | CD45- |
| CC | GO:0001725 | stress fiber | 9.98E-04 | 1.06E-02 | Myl9/Actb/Acta2 | 3 | aPD-L1+aTGF-β vs PBS | Down-regulated gen | CD45- |
| CC | GO:0097517 | contractile actin filament bundle | 9.98E-04 | 1.06E-02 | Myl9/Actb/Acta2 | 3 | aPD-L1+aTGF-β vs PBS | Down-regulated gen | CD45- |
| CC | GO:0044305 | calyx of Held | 1.07E-03 | 1.06E-02 | Actb/Actg1 | 2 | aPD-L1+aTGF-β vs PBS | Down-regulated gen | CD45- |
| CC | GO:0032432 | actin filament bundle | 1.33E-03 | 1.24E-02 | Myl9/Actb/Acta2 | 3 | aPD-L1+aTGF-β vs PBS | Down-regulated gen | CD45- |
| CC | GO:0042641 | actomyosin | 1.42E-03 | 1.26E-02 | Myl9/Actb/Acta2 | 3 | aPD-L1+aTGF-β vs PBS | Down-regulated gen | CD45- |
| CC | GO:0070069 | cytochrome complex | 2.01E-03 | 1.68E-02 | COX2/COX3 | 2 | aPD-L1+aTGF-β vs PBS | Down-regulated gen | CD45- |
| CC | GO:0016469 | proton-transporting two-sector ATPase complex | 5.18E-03 | 4.12E-02 | Atp6v1b1/ATP6 | 2 | aPD-L1+aTGF-β vs PBS | Down-regulated gen | CD45- |
| CC | GO:0016328 | lateral plasma membrane | 6.45E-03 | 4.88E-02 | Atp6v1b1/Gja1 | 2 | aPD-L1+aTGF-β vs PBS | Down-regulated gen | CD45- |
| CC | GO:0030016 | myofibril | 6.87E-03 | 4.96E-02 | Myl9/Actg1/Csrp2 | 3 | aPD-L1+aTGF-β vs PBS | Down-regulated gen | CD45- |
| MF | GO:0019838 | growth factor binding | 1.97E-05 | 2.40E-03 | Ccn1/Ltbp4/Col1 | 5 | aPD-L1+aTGF-β vs PBS | Down-regulated gen | CD45- |
| MF | GO:0015078 | proton transmembrane transporter activity | 8.75E-05 | 5.34E-03 | Atp6v1b1/COX2/C | 4 | aPD-L1+aTGF-β vs PBS | Down-regulated gen | CD45- |
| MF | GO:0005198 | structural molecule activity | 2.42E-04 | 8.64E-03 | Actb/Actg1/Ltbp4/ | 7 | aPD-L1+aTGF-β vs PBS | Down-regulated gen | CD45- |
| MF | GO:0005201 | extracellular matrix structural constituent | 2.83E-04 | 8.64E-03 | Ltbp4/Col1a2/Fbr | 4 | aPD-L1+aTGF-β vs PBS | Down-regulated gen | CD45- |
| MF | GO:0048407 | platelet-derived growth factor binding | 3.65E-04 | 8.90E-03 | Col1a2/Col1a1 | 2 | aPD-L1+aTGF-β vs PBS | Down-regulated gen | CD45- |
| MF | GO:0099186 | structural constituent of postsynapse | 7.89E-04 | 1.41E-02 | Actb/Actg1 | 2 | aPD-L1+aTGF-β vs PBS | Down-regulated gen | CD45- |
| MF | GO:0005539 | glycosaminoglycan binding | 8.09E-04 | 1.41E-02 | Ccn1/Ccdc80/Prn | 4 | aPD-L1+aTGF-β vs PBS | Down-regulated gen | CD45- |
| MF | GO:0004129 | cytochrome-c oxidase activity | 1.37E-03 | 1.53E-02 | COX2/COX3 | 2 | aPD-L1+aTGF-β vs PBS | Down-regulated gen | CD45- |
| MF | GO:0015002 | heme-copper terminal oxidase activity | 1.37E-03 | 1.53E-02 | COX2/COX3 | 2 | aPD-L1+aTGF-β vs PBS | Down-regulated gen | CD45- |
| MF | GO:0016676 | oxidoreductase activity, acting on a heme group | 1.37E-03 | 1.53E-02 | COX2/COX3 | 2 | aPD-L1+aTGF-β vs PBS | Down-regulated gen | CD45- |
| MF | GO:0016675 | oxidoreductase activity, acting on a heme group | 1.50E-03 | 1.53E-02 | COX2/COX3 | 2 | aPD-L1+aTGF-β vs PBS | Down-regulated gen | CD45- |
| MF | GO:0098918 | structural constituent of synapse | 1.50E-03 | 1.53E-02 | Actb/Actg1 | 2 | aPD-L1+aTGF-β vs PBS | Down-regulated gen | CD45- |
| MF | GO:0015077 | monovalent inorganic cation transmembrane tra | 1.87E-03 | 1.69E-02 | Atp6v1b1/COX2/C | 4 | aPD-L1+aTGF-β vs PBS | Down-regulated gen | CD45- |
| MF | GO:0005520 | insulin-like growth factor binding | 1.94E-03 | 1.69E-02 | Ccn1/Igfbp5 | 2 | aPD-L1+aTGF-β vs PBS | Down-regulated gen | CD45- |
| MF | GO:0001968 | fibronectin binding | 2.80E-03 | 2.27E-02 | Ccdc80/Igfbp5 | 2 | aPD-L1+aTGF-β vs PBS | Down-regulated gen | CD45- |
| MF | GO:0017147 | Wnt-protein binding | 3.18E-03 | 2.42E-02 | Wls/Cthrc1 | 2 | aPD-L1+aTGF-β vs PBS | Down-regulated gen | CD45- |
| MF | GO:0002020 | protease binding | 3.37E-03 | 2.42E-02 | Prnp/Col1a2/Col1 | 3 | aPD-L1+aTGF-β vs PBS | Down-regulated gen | CD45- |
| MF | GO:0030020 | extracellular matrix structural constituent conferri | 3.58E-03 | 2.43E-02 | Col1a2/Col1a1 | 2 | aPD-L1+aTGF-β vs PBS | Down-regulated gen | CD45- |
| MF | GO:0008201 | heparin binding | 4.17E-03 | 2.68E-02 | Ccn1/Ccdc80/Fbr | 3 | aPD-L1+aTGF-β vs PBS | Down-regulated gen | CD45- |
| MF | GO:0005507 | copper ion binding | 5.42E-03 | 3.31E-02 | Prnp/COX2 | 2 | aPD-L1+aTGF-β vs PBS | Down-regulated gen | CD45- |
| MF | GO:0005200 | structural constituent of cytoskeleton | 6.48E-03 | 3.76E-02 | Actb/Actg1 | 2 | aPD-L1+aTGF-β vs PBS | Down-regulated gen | CD45- |
| BP | GO:0043903 | regulation of symbiosis, encompassing mutualis | 2.25E-07 | 2.45E-04 | Pfn1/Gapdh/Ifitm2 | 8 | aPD-L1+aTGF-β vs aPD | Up-regulated genes | CD45+ |
| BP | GO:0050792 | regulation of viral process | 1.06E-06 | 5.75E-04 | Pfn1/Ifitm2/Ppia/C | 7 | aPD-L1+aTGF-β vs aPD | Up-regulated genes | CD45+ |
| BP | GO:0043900 | regulation of multi-organism process | 1.95E-06 | 7.07E-04 | Pfn1/Arhgdib/Gap | 9 | aPD-L1+aTGF-β vs aPD | Up-regulated genes | CD45+ |
| BP | GO:0044403 | symbiont process | 1.15E-05 | 3.13E-03 | Pfn1/Gapdh/Ifitm2 | 8 | aPD-L1+aTGF-β vs aPD | Up-regulated genes | CD45+ |
| BP | GO:0044419 | interspecies interaction between organisms | 1.61E-05 | 3.50E-03 | Pfn1/Gapdh/Ifitm2 | 8 | aPD-L1+aTGF-β vs aPD | Up-regulated genes | CD45+ |
| BP | GO:0016032 | viral process | 2.23E-05 | 4.05E-03 | Pfn1/Ifitm2/Ppia/C | 7 | aPD-L1+aTGF-β vs aPD | Up-regulated genes | CD45+ |
| BP | GO:0006958 | complement activation, classical pathway | 3.60E-05 | 5.60E-03 | C1qa/C1qc/C1qb | 3 | aPD-L1+aTGF-β vs aPD | Up-regulated genes | CD45+ |
| BP | GO:1903900 | regulation of viral life cycle | 6.80E-05 | 9.24E-03 | Ifitm2/Ppia/Lgals1 | 5 | aPD-L1+aTGF-β vs aPD | Up-regulated genes | CD45+ |
| BP | GO:0046596 | regulation of viral entry into host cell | 8.43E-05 | 1.02E-02 | Ifitm2/Lgals1/Snx | 3 | aPD-L1+aTGF-β vs aPD | Up-regulated genes | CD45+ |
| BP | GO:0002181 | cytoplasmic translation | 1.34E-04 | 1.39E-02 | Rpl18/Rpl18a/Rp | 4 | aPD-L1+aTGF-β vs aPD | Up-regulated genes | CD45+ |
| BP | GO:0048524 | positive regulation of viral process | 1.41E-04 | 1.39E-02 | Pfn1/Ppia/Cfl1/Lg | 4 | aPD-L1+aTGF-β vs aPD | Up-regulated genes | CD45+ |
| BP | GO:0043902 | positive regulation of multi-organism process | 1.66E-04 | 1.43E-02 | Pfn1/Gapdh/Ppia/ | 5 | aPD-L1+aTGF-β vs aPD | Up-regulated genes | CD45+ |
| BP | GO:0002455 | humoral immune response mediated by circulati | 1.79E-04 | 1.43E-02 | C1qa/C1qc/C1qb | 3 | aPD-L1+aTGF-β vs aPD | Up-regulated genes | CD45+ |
| BP | GO:0052548 | regulation of endopeptidase activity | 1.89E-04 | 1.43E-02 | Psme2/Gapdh/Ps | 6 | aPD-L1+aTGF-β vs aPD | Up-regulated genes | CD45+ |
| BP | GO:0051651 | maintenance of location in cell | 1.97E-04 | 1.43E-02 | Pfn1/Tmsb4x/Fth1 | 4 | aPD-L1+aTGF-β vs aPD | Up-regulated genes | CD45+ |
| BP | GO:0006956 | complement activation | 2.77E-04 | 1.88E-02 | C1qa/C1qc/C1qb | 3 | aPD-L1+aTGF-β vs aPD | Up-regulated genes | CD45+ |
| BP | GO:0010810 | regulation of cell-substrate adhesion | 2.96E-04 | 1.89E-02 | Spp1/Rac2/S100 | 5 | aPD-L1+aTGF-β vs aPD | Up-regulated genes | CD45+ |
| BP | GO:0019058 | viral life cycle | 3.21E-04 | 1.94E-02 | Ifitm2/Ppia/Lgals1 | 5 | aPD-L1+aTGF-β vs aPD | Up-regulated genes | CD45+ |
| BP | GO:0052547 | regulation of peptidase activity | 3.95E-04 | 2.26E-02 | Psme2/Gapdh/Ps | 6 | aPD-L1+aTGF-β vs aPD | Up-regulated genes | CD45+ |
| BP | GO:0072376 | protein activation cascade | 4.33E-04 | 2.36E-02 | C1qa/C1qc/C1qb | 3 | aPD-L1+aTGF-β vs aPD | Up-regulated genes | CD45+ |
| BP | GO:0046718 | viral entry into host cell | 4.96E-04 | 2.46E-02 | Ifitm2/Lgals1/Snx | 3 | aPD-L1+aTGF-β vs aPD | Up-regulated genes | CD45+ |
| BP | GO:0051701 | interaction with host | 4.97E-04 | 2.46E-02 | Gapdh/Ifitm2/Lgal | 4 | aPD-L1+aTGF-β vs aPD | Up-regulated genes | CD45+ |
| BP | GO:0006959 | humoral immune response | 6.11E-04 | 2.48E-02 | C1qa/C1qc/Gapd | 4 | aPD-L1+aTGF-β vs aPD | Up-regulated genes | CD45+ |
| BP | GO:0042989 | sequestering of actin monomers | 6.30E-04 | 2.48E-02 | Pfn1/Tmsb4x | 2 | aPD-L1+aTGF-β vs aPD | Up-regulated genes | CD45+ |
| BP | GO:0030260 | entry into host cell | 6.37E-04 | 2.48E-02 | Ifitm2/Lgals1/Snx | 3 | aPD-L1+aTGF-β vs aPD | Up-regulated genes | CD45+ |
| BP | GO:0044409 | entry into host | 6.37E-04 | 2.48E-02 | Ifitm2/Lgals1/Snx | 3 | aPD-L1+aTGF-β vs aPD | Up-regulated genes | CD45+ |
| BP | GO:0051806 | entry into cell of other organism involved in symb | 6.37E-04 | 2.48E-02 | Ifitm2/Lgals1/Snx | 3 | aPD-L1+aTGF-β vs aPD | Up-regulated genes | CD45+ |
| BP | GO:0051828 | entry into other organism involved in symbiotic ir | 6.37E-04 | 2.48E-02 | Ifitm2/Lgals1/Snx | 3 | aPD-L1+aTGF-β vs aPD | Up-regulated genes | CD45+ |
| BP | GO:0030837 | negative regulation of actin filament polymerizati | 6.76E-04 | 2.54E-02 | Pfn1/Tmsb4x/Cfl1 | 3 | aPD-L1+aTGF-β vs aPD | Up-regulated genes | CD45+ |
| BP | GO:0050808 | synapse organization | 7.08E-04 | 2.57E-02 | Pfn1/C1qa/C1qc/ | 6 | aPD-L1+aTGF-β vs aPD | Up-regulated genes | CD45+ |
| BP | GO:0098885 | modification of postsynaptic actin cytoskeleton | 7.55E-04 | 2.65E-02 | Pfn1/Cfl1 | 2 | aPD-L1+aTGF-β vs aPD | Up-regulated genes | CD45+ |
| BP | GO:0032272 | negative regulation of protein polymerization | 1.33E-03 | 4.51E-02 | Pfn1/Tmsb4x/Cfl1 | 3 | aPD-L1+aTGF-β vs aPD | Up-regulated genes | CD45+ |
| CC | GO:0022626 | cytosolic ribosome | 5.69E-11 | 8.71E-09 | Rps5/Rps4x/Rpl1 | 9 | aPD-L1+aTGF-β vs aPD | Up-regulated genes | CD45+ |
| CC | GO:0005840 | ribosome | 2.04E-09 | 1.56E-07 | Rps5/Rps4x/Rpl1 | 10 | aPD-L1+aTGF-β vs aPD | Up-regulated genes | CD45+ |
| CC | GO:0044391 | ribosomal subunit | 9.28E-09 | 4.73E-07 | Rps5/Rps4x/Rpl1 | 9 | aPD-L1+aTGF-β vs aPD | Up-regulated genes | CD45+ |
| CC | GO:0044445 | cytosolic part | 7.18E-08 | 2.74E-06 | Rps5/Rps4x/Rpl1 | 9 | aPD-L1+aTGF-β vs aPD | Up-regulated genes | CD45+ |
| CC | GO:0022625 | cytosolic large ribosomal subunit | 1.45E-06 | 4.43E-05 | Rpl18/Rpl18a/Rp | 5 | aPD-L1+aTGF-β vs aPD | Up-regulated genes | CD45+ |
| CC | GO:0042788 | polysomal ribosome | 2.25E-06 | 5.73E-05 | Rpl18/Rpl18a/Eif | 4 | aPD-L1+aTGF-β vs aPD | Up-regulated genes | CD45+ |
| CC | GO:0005844 | polysome | 3.21E-06 | 7.03E-05 | Rps4x/Rpl18/Rpl | 5 | aPD-L1+aTGF-β vs aPD | Up-regulated genes | CD45+ |
| CC | GO:0022627 | cytosolic small ribosomal subunit | 1.19E-05 | 2.27E-04 | Rps5/Rps4x/Rps | 4 | aPD-L1+aTGF-β vs aPD | Up-regulated genes | CD45+ |
| CC | GO:0015934 | large ribosomal subunit | 4.76E-05 | 8.08E-04 | Rpl18/Rpl18a/Rp | 5 | aPD-L1+aTGF-β vs aPD | Up-regulated genes | CD45+ |
| CC | GO:0015935 | small ribosomal subunit | 1.01E-04 | 1.55E-03 | Rps5/Rps4x/Rps | 4 | aPD-L1+aTGF-β vs aPD | Up-regulated genes | CD45+ |
| CC | GO:0043209 | myelin sheath | 4.23E-04 | 5.88E-03 | Gapdh/Actg1/Ppia | 5 | aPD-L1+aTGF-β vs aPD | Up-regulated genes | CD45+ |
| CC | GO:0005581 | collagen trimer | 6.06E-04 | 7.73E-03 | C1qa/C1qc/C1qb | 3 | aPD-L1+aTGF-β vs aPD | Up-regulated genes | CD45+ |
| CC | GO:0098685 | Schaffer collateral - CA1 synapse | 2.79E-03 | 3.28E-02 | Pfn1/Actg1/Srgn | 3 | aPD-L1+aTGF-β vs aPD | Up-regulated genes | CD45+ |

| | | | | | | | | | |
|---|---|---|---|---|---|---|---|---|---|
| MF | GO:0003735 | structural constituent of ribosome | 1.27E-09 | 1.92E-07 | Rps5/Rps4x/Rpl1 | 9 | aPD-L1+aTGF-β vs aPD | Up-regulated genes | CD45+ |
| MF | GO:0005198 | structural molecule activity | 1.31E-06 | 9.87E-05 | Rps5/Rps4x/Actg | 10 | aPD-L1+aTGF-β vs aPD | Up-regulated genes | CD45+ |
| MF | GO:0061135 | endopeptidase regulator activity | 3.75E-04 | 1.89E-02 | Psme2/Gapdh/Ps | 4 | aPD-L1+aTGF-β vs aPD | Up-regulated genes | CD45+ |
| MF | GO:0061134 | peptidase regulator activity | 9.44E-04 | 3.56E-02 | Psme2/Gapdh/Ps | 4 | aPD-L1+aTGF-β vs aPD | Up-regulated genes | CD45+ |
| MF | GO:0019843 | rRNA binding | 1.65E-03 | 4.98E-02 | Rps5/Rps4x/Rps | 3 | aPD-L1+aTGF-β vs aPD | Up-regulated genes | CD45+ |
| BP | GO:0061844 | antimicrobial humoral immune response mediat | 6.92E-07 | 3.84E-04 | Cxcl2/Cxcl1/Rpl3 | 4 | aPD-L1+aTGF-β vs aPD | Up-regulated genes | CD45- |
| BP | GO:0019730 | antimicrobial humoral response | 2.06E-06 | 5.70E-04 | Cxcl2/Cxcl1/Rpl3 | 4 | aPD-L1+aTGF-β vs aPD | Up-regulated genes | CD45- |
| BP | GO:0002181 | cytoplasmic translation | 2.04E-05 | 3.76E-03 | Rpl30/Rpl36/Rpl3 | 4 | aPD-L1+aTGF-β vs aPD | Up-regulated genes | CD45- |
| BP | GO:0006959 | humoral immune response | 6.24E-05 | 8.65E-03 | Cxcl2/Cxcl1/Rpl3 | 4 | aPD-L1+aTGF-β vs aPD | Up-regulated genes | CD45- |
| BP | GO:0009617 | response to bacterium | 2.67E-04 | 2.96E-02 | Cxcl2/Cxcl1/Rpl3 | 6 | aPD-L1+aTGF-β vs aPD | Up-regulated genes | CD45- |
| CC | GO:0022626 | cytosolic ribosome | 3.77E-16 | 2.49E-14 | Rpl22/Rps15a/Rp | 11 | aPD-L1+aTGF-β vs aPD | Up-regulated genes | CD45- |
| CC | GO:0022625 | cytosolic large ribosomal subunit | 6.25E-15 | 2.06E-13 | Rpl22/Rpl30/Rpl3 | 9 | aPD-L1+aTGF-β vs aPD | Up-regulated genes | CD45- |
| CC | GO:0005840 | ribosome | 3.01E-14 | 6.63E-13 | Rpl22/Rps15a/Rp | 12 | aPD-L1+aTGF-β vs aPD | Up-regulated genes | CD45- |
| CC | GO:0044391 | ribosomal subunit | 2.17E-13 | 3.58E-12 | Rpl22/Rps15a/Rp | 11 | aPD-L1+aTGF-β vs aPD | Up-regulated genes | CD45- |
| CC | GO:0044445 | cytosolic part | 2.60E-12 | 3.44E-11 | Rpl22/Rps15a/Rp | 11 | aPD-L1+aTGF-β vs aPD | Up-regulated genes | CD45- |
| CC | GO:0015934 | large ribosomal subunit | 4.79E-12 | 5.27E-11 | Rpl22/Rpl30/Rpl3 | 9 | aPD-L1+aTGF-β vs aPD | Up-regulated genes | CD45- |
| CC | GO:0070069 | cytochrome complex | 2.95E-05 | 2.78E-04 | Cox8a/Uqcrb/Ndu | 3 | aPD-L1+aTGF-β vs aPD | Up-regulated genes | CD45- |
| CC | GO:0045277 | respiratory chain complex IV | 6.29E-04 | 5.19E-03 | Cox8a/Ndufa4 | 2 | aPD-L1+aTGF-β vs aPD | Up-regulated genes | CD45- |
| CC | GO:0098803 | respiratory chain complex | 7.66E-04 | 5.62E-03 | Cox8a/Uqcrb/Ndu | 3 | aPD-L1+aTGF-β vs aPD | Up-regulated genes | CD45- |
| CC | GO:0070469 | respiratory chain | 9.86E-04 | 6.51E-03 | Cox8a/Uqcrb/Ndu | 3 | aPD-L1+aTGF-β vs aPD | Up-regulated genes | CD45- |
| CC | GO:0042788 | polysomal ribosome | 1.95E-03 | 1.17E-02 | Rpl30/Rpl39 | 2 | aPD-L1+aTGF-β vs aPD | Up-regulated genes | CD45- |
| CC | GO:0022627 | cytosolic small ribosomal subunit | 4.55E-03 | 2.50E-02 | Rps15a/Rps27l | 2 | aPD-L1+aTGF-β vs aPD | Up-regulated genes | CD45- |
| MF | GO:0003735 | structural constituent of ribosome | 9.42E-15 | 6.31E-13 | Rpl22/Rps15a/Rp | 11 | aPD-L1+aTGF-β vs aPD | Up-regulated genes | CD45- |
| MF | GO:0005198 | structural molecule activity | 2.59E-10 | 8.69E-09 | Rpl22/Rps15a/Rp | 12 | aPD-L1+aTGF-β vs aPD | Up-regulated genes | CD45- |
| MF | GO:0009055 | electron transfer activity | 3.58E-04 | 8.00E-03 | Cox8a/Uqcrb/Ndu | 3 | aPD-L1+aTGF-β vs aPD | Up-regulated genes | CD45- |
| MF | GO:0004129 | cytochrome-c oxidase activity | 1.01E-03 | 1.06E-02 | Cox8a/Ndufa4 | 2 | aPD-L1+aTGF-β vs aPD | Up-regulated genes | CD45- |
| MF | GO:0015002 | heme-copper terminal oxidase activity | 1.01E-03 | 1.06E-02 | Cox8a/Ndufa4 | 2 | aPD-L1+aTGF-β vs aPD | Up-regulated genes | CD45- |
| MF | GO:0016676 | oxidoreductase activity, acting on a heme group | 1.01E-03 | 1.06E-02 | Cox8a/Ndufa4 | 2 | aPD-L1+aTGF-β vs aPD | Up-regulated genes | CD45- |
| MF | GO:0016675 | oxidoreductase activity, acting on a heme group | 1.11E-03 | 1.06E-02 | Cox8a/Ndufa4 | 2 | aPD-L1+aTGF-β vs aPD | Up-regulated genes | CD45- |
| MF | GO:0008009 | chemokine activity | 2.49E-03 | 2.08E-02 | Cxcl2/Cxcl1 | 2 | aPD-L1+aTGF-β vs aPD | Up-regulated genes | CD45- |
| MF | GO:0042379 | chemokine receptor binding | 4.20E-03 | 3.12E-02 | Cxcl2/Cxcl1 | 2 | aPD-L1+aTGF-β vs aPD | Up-regulated genes | CD45- |
| MF | GO:0045182 | translation regulator activity | 5.64E-03 | 3.78E-02 | Rpl22/Rps27l | 2 | aPD-L1+aTGF-β vs aPD | Up-regulated genes | CD45- |
| MF | GO:0048306 | calcium-dependent protein binding | 8.03E-03 | 4.89E-02 | S100a6/S100a4 | 2 | aPD-L1+aTGF-β vs aPD | Up-regulated genes | CD45- |
| BP | GO:0002181 | cytoplasmic translation | 1.25E-18 | 1.09E-15 | Rplp0/Rpl35a/Rp | 13 | aPD-L1+aTGF-β vs aPD | Down-regulated gen | CD45+ |
| BP | GO:0042274 | ribosomal small subunit biogenesis | 3.40E-09 | 1.48E-06 | Rpsa/Rps19/Rps | 7 | aPD-L1+aTGF-β vs aPD | Down-regulated gen | CD45+ |
| BP | GO:0000028 | ribosomal small subunit assembly | 6.17E-09 | 1.79E-06 | Rpsa/Rps19/Rps | 5 | aPD-L1+aTGF-β vs aPD | Down-regulated gen | CD45+ |
| BP | GO:0022613 | ribonucleoprotein complex biogenesis | 3.30E-08 | 7.20E-06 | Rplp0/Rpl35a/Rp | 12 | aPD-L1+aTGF-β vs aPD | Down-regulated gen | CD45+ |
| BP | GO:0042254 | ribosome biogenesis | 5.44E-08 | 9.48E-06 | Rplp0/Rpl35a/Rp | 10 | aPD-L1+aTGF-β vs aPD | Down-regulated gen | CD45+ |
| BP | GO:0042255 | ribosome assembly | 1.02E-07 | 1.48E-05 | Rplp0/Rpsa/Rps1 | 6 | aPD-L1+aTGF-β vs aPD | Down-regulated gen | CD45+ |
| BP | GO:0022618 | ribonucleoprotein complex assembly | 1.21E-06 | 1.50E-04 | Rplp0/Rpsa/Rpl1 | 8 | aPD-L1+aTGF-β vs aPD | Down-regulated gen | CD45+ |
| BP | GO:0071826 | ribonucleoprotein complex subunit organization | 1.79E-06 | 1.95E-04 | Rplp0/Rpsa/Rpl1 | 8 | aPD-L1+aTGF-β vs aPD | Down-regulated gen | CD45+ |
| BP | GO:0061844 | antimicrobial humoral immune response mediat | 9.93E-06 | 9.62E-04 | Hist1h2bc/Rpl39/ | 4 | aPD-L1+aTGF-β vs aPD | Down-regulated gen | CD45+ |
| BP | GO:0046034 | ATP metabolic process | 1.38E-05 | 1.20E-03 | ATP8/COX3/COX | 7 | aPD-L1+aTGF-β vs aPD | Down-regulated gen | CD45+ |
| BP | GO:0019730 | antimicrobial humoral response | 2.51E-05 | 1.72E-03 | Hist1h2bc/Rpl39/ | 4 | aPD-L1+aTGF-β vs aPD | Down-regulated gen | CD45+ |
| BP | GO:0009126 | purine nucleoside monophosphate metabolic pr | 2.75E-05 | 1.72E-03 | ATP8/COX3/COX | 7 | aPD-L1+aTGF-β vs aPD | Down-regulated gen | CD45+ |
| BP | GO:0009167 | purine ribonucleoside monophosphate metaboli | 2.75E-05 | 1.72E-03 | ATP8/COX3/COX | 7 | aPD-L1+aTGF-β vs aPD | Down-regulated gen | CD45+ |
| BP | GO:0009205 | purine ribonucleoside triphosphate metabolic pr | 2.82E-05 | 1.72E-03 | ATP8/COX3/COX | 7 | aPD-L1+aTGF-β vs aPD | Down-regulated gen | CD45+ |
| BP | GO:0009161 | ribonucleoside monophosphate metabolic proce | 3.07E-05 | 1.72E-03 | ATP8/COX3/COX | 7 | aPD-L1+aTGF-β vs aPD | Down-regulated gen | CD45+ |
| BP | GO:0009199 | ribonucleoside triphosphate metabolic process | 3.15E-05 | 1.72E-03 | ATP8/COX3/COX | 7 | aPD-L1+aTGF-β vs aPD | Down-regulated gen | CD45+ |
| BP | GO:0009144 | purine nucleoside triphosphate metabolic proce | 3.52E-05 | 1.80E-03 | ATP8/COX3/COX | 7 | aPD-L1+aTGF-β vs aPD | Down-regulated gen | CD45+ |
| BP | GO:0009123 | nucleoside monophosphate metabolic process | 3.71E-05 | 1.80E-03 | ATP8/COX3/COX | 7 | aPD-L1+aTGF-β vs aPD | Down-regulated gen | CD45+ |
| BP | GO:0009141 | nucleoside triphosphate metabolic process | 5.33E-05 | 2.45E-03 | ATP8/COX3/COX | 7 | aPD-L1+aTGF-β vs aPD | Down-regulated gen | CD45+ |
| BP | GO:0015985 | energy coupled proton transport, down electroc | 6.12E-05 | 2.47E-03 | ATP8/Atp5k/ATP6 | 3 | aPD-L1+aTGF-β vs aPD | Down-regulated gen | CD45+ |
| BP | GO:0015986 | ATP synthesis coupled proton transport | 6.12E-05 | 2.47E-03 | ATP8/Atp5k/ATP6 | 3 | aPD-L1+aTGF-β vs aPD | Down-regulated gen | CD45+ |
| BP | GO:0006364 | rRNA processing | 6.24E-05 | 2.47E-03 | Rpl35a/Rpl35/Rp | 6 | aPD-L1+aTGF-β vs aPD | Down-regulated gen | CD45+ |
| BP | GO:0042773 | ATP synthesis coupled electron transport | 8.05E-05 | 3.05E-03 | COX3/COX2/CYT | 4 | aPD-L1+aTGF-β vs aPD | Down-regulated gen | CD45+ |
| BP | GO:0042273 | ribosomal large subunit biogenesis | 1.18E-04 | 4.28E-03 | Rplp0/Rpl35a/Rp | 4 | aPD-L1+aTGF-β vs aPD | Down-regulated gen | CD45+ |
| BP | GO:0016072 | rRNA metabolic process | 1.44E-04 | 5.02E-03 | Rpl35a/Rpl35/Rp | 6 | aPD-L1+aTGF-β vs aPD | Down-regulated gen | CD45+ |
| BP | GO:0022904 | respiratory electron transport chain | 2.39E-04 | 7.69E-03 | COX3/COX2/CYT | 4 | aPD-L1+aTGF-β vs aPD | Down-regulated gen | CD45+ |
| BP | GO:1902600 | proton transmembrane transport | 2.39E-04 | 7.69E-03 | ATP8/Atp5k/CYT | 4 | aPD-L1+aTGF-β vs aPD | Down-regulated gen | CD45+ |
| BP | GO:0045333 | cellular respiration | 2.47E-04 | 7.69E-03 | COX3/COX2/CYT | 5 | aPD-L1+aTGF-β vs aPD | Down-regulated gen | CD45+ |
| BP | GO:0022900 | electron transport chain | 2.76E-04 | 8.31E-03 | COX3/COX2/CYT | 4 | aPD-L1+aTGF-β vs aPD | Down-regulated gen | CD45+ |
| BP | GO:0009150 | purine ribonucleotide metabolic process | 3.39E-04 | 9.84E-03 | ATP8/COX3/COX | 7 | aPD-L1+aTGF-β vs aPD | Down-regulated gen | CD45+ |
| BP | GO:0042742 | defense response to bacterium | 3.67E-04 | 1.03E-02 | Hist1h2bc/Rpl39/ | 5 | aPD-L1+aTGF-β vs aPD | Down-regulated gen | CD45+ |
| BP | GO:0006119 | oxidative phosphorylation | 3.79E-04 | 1.03E-02 | COX3/COX2/CYT | 4 | aPD-L1+aTGF-β vs aPD | Down-regulated gen | CD45+ |
| BP | GO:0009259 | ribonucleotide metabolic process | 4.03E-04 | 1.07E-02 | ATP8/COX3/COX | 7 | aPD-L1+aTGF-β vs aPD | Down-regulated gen | CD45+ |
| BP | GO:0006163 | purine nucleotide metabolic process | 4.85E-04 | 1.24E-02 | ATP8/COX3/COX | 7 | aPD-L1+aTGF-β vs aPD | Down-regulated gen | CD45+ |
| BP | GO:0019693 | ribose phosphate metabolic process | 5.35E-04 | 1.31E-02 | ATP8/COX3/COX | 7 | aPD-L1+aTGF-β vs aPD | Down-regulated gen | CD45+ |
| BP | GO:0030490 | maturation of SSU-rRNA | 5.40E-04 | 1.31E-02 | Rps19/Rps21/Rp | 3 | aPD-L1+aTGF-β vs aPD | Down-regulated gen | CD45+ |
| BP | GO:0006754 | ATP biosynthetic process | 6.89E-04 | 1.62E-02 | ATP8/COX2/Atp5 | 4 | aPD-L1+aTGF-β vs aPD | Down-regulated gen | CD45+ |
| BP | GO:0050829 | defense response to Gram-negative bacterium | 7.11E-04 | 1.63E-02 | Rps19/Hmgb2/Rp | 3 | aPD-L1+aTGF-β vs aPD | Down-regulated gen | CD45+ |
| BP | GO:0072521 | purine-containing compound metabolic process | 7.44E-04 | 1.66E-02 | ATP8/COX3/COX | 7 | aPD-L1+aTGF-β vs aPD | Down-regulated gen | CD45+ |
| BP | GO:0009206 | purine ribonucleoside triphosphate biosynthetic | 9.43E-04 | 2.02E-02 | ATP8/COX2/Atp5 | 4 | aPD-L1+aTGF-β vs aPD | Down-regulated gen | CD45+ |
| BP | GO:0006959 | humoral immune response | 9.75E-04 | 2.02E-02 | Hist1h2bc/Rpl39/ | 4 | aPD-L1+aTGF-β vs aPD | Down-regulated gen | CD45+ |
| BP | GO:0009145 | purine nucleoside triphosphate biosynthetic pro | 9.75E-04 | 2.02E-02 | ATP8/COX2/Atp5 | 4 | aPD-L1+aTGF-β vs aPD | Down-regulated gen | CD45+ |
| BP | GO:0009201 | ribonucleoside triphosphate biosynthetic proces | 1.04E-03 | 2.11E-02 | ATP8/COX2/Atp5 | 4 | aPD-L1+aTGF-β vs aPD | Down-regulated gen | CD45+ |
| BP | GO:0034660 | ncRNA metabolic process | 1.25E-03 | 2.43E-02 | Rpl35a/Rpl35/Rp | 7 | aPD-L1+aTGF-β vs aPD | Down-regulated gen | CD45+ |
| BP | GO:0034101 | erythrocyte homeostasis | 1.26E-03 | 2.43E-02 | Zfp36/Rps19/Hm | 4 | aPD-L1+aTGF-β vs aPD | Down-regulated gen | CD45+ |
| BP | GO:0042775 | mitochondrial ATP synthesis coupled electron tr | 1.28E-03 | 2.43E-02 | COX3/CYTB/ND2 | 3 | aPD-L1+aTGF-β vs aPD | Down-regulated gen | CD45+ |
| BP | GO:0002227 | innate immune response in mucosa | 1.32E-03 | 2.45E-02 | Hist1h2bc/Rpl39 | 2 | aPD-L1+aTGF-β vs aPD | Down-regulated gen | CD45+ |
| BP | GO:0009127 | purine nucleoside monophosphate biosynthetic | 1.38E-03 | 2.45E-02 | ATP8/COX2/Atp5 | 4 | aPD-L1+aTGF-β vs aPD | Down-regulated gen | CD45+ |
| BP | GO:0009168 | purine ribonucleoside monophosphate biosynth | 1.38E-03 | 2.45E-02 | ATP8/COX2/Atp5 | 4 | aPD-L1+aTGF-β vs aPD | Down-regulated gen | CD45+ |
| BP | GO:0015980 | energy derivation by oxidation of organic compo | 1.44E-03 | 2.50E-02 | COX3/COX2/CYT | 5 | aPD-L1+aTGF-β vs aPD | Down-regulated gen | CD45+ |
| BP | GO:0009142 | nucleoside triphosphate biosynthetic process | 1.46E-03 | 2.50E-02 | ATP8/COX2/Atp5 | 4 | aPD-L1+aTGF-β vs aPD | Down-regulated gen | CD45+ |
| BP | GO:0034470 | ncRNA processing | 1.53E-03 | 2.55E-02 | Rpl35a/Rpl35/Rp | 6 | aPD-L1+aTGF-β vs aPD | Down-regulated gen | CD45+ |
| BP | GO:0009156 | ribonucleoside monophosphate biosynthetic pro | 1.55E-03 | 2.55E-02 | ATP8/COX2/Atp5 | 4 | aPD-L1+aTGF-β vs aPD | Down-regulated gen | CD45+ |
| BP | GO:0009124 | nucleoside monophosphate biosynthetic proces | 1.78E-03 | 2.85E-02 | ATP8/COX2/Atp5 | 4 | aPD-L1+aTGF-β vs aPD | Down-regulated gen | CD45+ |
| BP | GO:0009617 | response to bacterium | 1.80E-03 | 2.85E-02 | Zfp36/Rpl13a/His | 7 | aPD-L1+aTGF-β vs aPD | Down-regulated gen | CD45+ |
| BP | GO:0009117 | nucleotide metabolic process | 2.06E-03 | 3.21E-02 | ATP8/COX3/COX | 7 | aPD-L1+aTGF-β vs aPD | Down-regulated gen | CD45+ |

| | | | | | | | | | |
|---|---|---|---|---|---|---|---|---|---|
| BP | GO:0019731 | antibacterial humoral response | 2.20E-03 | 3.28E-02 | Hist1h2bc/Rpl39 | 2 | aPD-L1+aTGF-β vs aPD | Down-regulated gen | CD45+ |
| BP | GO:0051444 | negative regulation of ubiquitin-protein transfera | 2.20E-03 | 3.28E-02 | Rpl23/Smad7 | 2 | aPD-L1+aTGF-β vs aPD | Down-regulated gen | CD45+ |
| BP | GO:0006753 | nucleoside phosphate metabolic process | 2.22E-03 | 3.28E-02 | ATP8/COX3/COX | 7 | aPD-L1+aTGF-β vs aPD | Down-regulated gen | CD45+ |
| BP | GO:0050830 | defense response to Gram-positive bacterium | 2.26E-03 | 3.29E-02 | Hist1h2bc/Rpl39/ | 3 | aPD-L1+aTGF-β vs aPD | Down-regulated gen | CD45+ |
| BP | GO:0002262 | myeloid cell homeostasis | 2.89E-03 | 4.14E-02 | Zfp36/Rps19/Hmg | 4 | aPD-L1+aTGF-β vs aPD | Down-regulated gen | CD45+ |
| BP | GO:0002385 | mucosal immune response | 3.29E-03 | 4.63E-02 | Hist1h2bc/Rpl39 | 2 | aPD-L1+aTGF-β vs aPD | Down-regulated gen | CD45+ |
| BP | GO:0002251 | organ or tissue specific immune response | 3.60E-03 | 4.90E-02 | Hist1h2bc/Rpl39 | 2 | aPD-L1+aTGF-β vs aPD | Down-regulated gen | CD45+ |
| BP | GO:0010837 | regulation of keratinocyte proliferation | 3.60E-03 | 4.90E-02 | Zfp36/Ctsl | 2 | aPD-L1+aTGF-β vs aPD | Down-regulated gen | CD45+ |
| CC | GO:0022626 | cytosolic ribosome | 9.13E-47 | 1.00E-44 | Rplp2/Rplp0/Rpl3 | 28 | aPD-L1+aTGF-β vs aPD | Down-regulated gen | CD45+ |
| CC | GO:0044391 | ribosomal subunit | 4.31E-39 | 1.96E-37 | Rplp2/Rplp0/Rpl3 | 28 | aPD-L1+aTGF-β vs aPD | Down-regulated gen | CD45+ |
| CC | GO:0005840 | ribosome | 5.36E-39 | 1.96E-37 | Rplp2/Rpl41/Rplp | 29 | aPD-L1+aTGF-β vs aPD | Down-regulated gen | CD45+ |
| CC | GO:0044445 | cytosolic part | 4.72E-36 | 1.30E-34 | Rplp2/Rplp0/Rpl3 | 28 | aPD-L1+aTGF-β vs aPD | Down-regulated gen | CD45+ |
| CC | GO:0022625 | cytosolic large ribosomal subunit | 1.20E-28 | 2.63E-27 | Rplp2/Rplp0/Rpl3 | 17 | aPD-L1+aTGF-β vs aPD | Down-regulated gen | CD45+ |
| CC | GO:0015934 | large ribosomal subunit | 7.26E-23 | 1.33E-21 | Rplp2/Rplp0/Rpl3 | 17 | aPD-L1+aTGF-β vs aPD | Down-regulated gen | CD45+ |
| CC | GO:0022627 | cytosolic small ribosomal subunit | 4.27E-18 | 6.70E-17 | Rpsa/Rps3a1/Rp | 11 | aPD-L1+aTGF-β vs aPD | Down-regulated gen | CD45+ |
| CC | GO:0015935 | small ribosomal subunit | 2.76E-15 | 3.80E-14 | Rpsa/Rps3a1/Rp | 11 | aPD-L1+aTGF-β vs aPD | Down-regulated gen | CD45+ |
| CC | GO:0042788 | polysomal ribosome | 1.04E-11 | 1.27E-10 | Rpl39/Rps29/Rps | 7 | aPD-L1+aTGF-β vs aPD | Down-regulated gen | CD45+ |
| CC | GO:0005844 | polysome | 7.53E-09 | 8.29E-08 | Rpl39/Rps29/Rps | 7 | aPD-L1+aTGF-β vs aPD | Down-regulated gen | CD45+ |
| CC | GO:0098803 | respiratory chain complex | 4.76E-07 | 4.76E-06 | ND3/COX3/COX2 | 6 | aPD-L1+aTGF-β vs aPD | Down-regulated gen | CD45+ |
| CC | GO:0098800 | inner mitochondrial membrane protein complex | 5.23E-07 | 4.79E-06 | ATP8/ND3/Atp5k/ | 7 | aPD-L1+aTGF-β vs aPD | Down-regulated gen | CD45+ |
| CC | GO:0070469 | respiratory chain | 8.05E-07 | 6.81E-06 | ND3/COX3/COX2 | 6 | aPD-L1+aTGF-β vs aPD | Down-regulated gen | CD45+ |
| CC | GO:0045263 | proton-transporting ATP synthase complex, coup | 1.22E-05 | 9.55E-05 | ATP8/Atp5k/ATP6 | 3 | aPD-L1+aTGF-β vs aPD | Down-regulated gen | CD45+ |
| CC | GO:0044455 | mitochondrial membrane part | 1.45E-05 | 1.06E-04 | ATP8/ND3/Atp5k/ | 7 | aPD-L1+aTGF-β vs aPD | Down-regulated gen | CD45+ |
| CC | GO:0005743 | mitochondrial inner membrane | 2.02E-05 | 1.39E-04 | ATP8/ND3/COX3/ | 9 | aPD-L1+aTGF-β vs aPD | Down-regulated gen | CD45+ |
| CC | GO:0019866 | organelle inner membrane | 3.80E-05 | 2.46E-04 | ATP8/ND3/COX3/ | 9 | aPD-L1+aTGF-β vs aPD | Down-regulated gen | CD45+ |
| CC | GO:0005753 | mitochondrial proton-transporting ATP synthase | 4.43E-05 | 2.71E-04 | ATP8/Atp5k/ATP6 | 3 | aPD-L1+aTGF-β vs aPD | Down-regulated gen | CD45+ |
| CC | GO:0045259 | proton-transporting ATP synthase complex | 5.25E-05 | 3.04E-04 | ATP8/Atp5k/ATP6 | 3 | aPD-L1+aTGF-β vs aPD | Down-regulated gen | CD45+ |
| CC | GO:0098798 | mitochondrial protein complex | 5.98E-05 | 3.23E-04 | ATP8/ND3/Atp5k/ | 7 | aPD-L1+aTGF-β vs aPD | Down-regulated gen | CD45+ |
| CC | GO:0033177 | proton-transporting two-sector ATPase complex, | 6.16E-05 | 3.23E-04 | ATP8/Atp5k/ATP6 | 3 | aPD-L1+aTGF-β vs aPD | Down-regulated gen | CD45+ |
| CC | GO:0070069 | cytochrome complex | 1.38E-04 | 6.91E-04 | COX3/COX2/CYT | 3 | aPD-L1+aTGF-β vs aPD | Down-regulated gen | CD45+ |
| CC | GO:0014069 | postsynaptic density | 1.45E-04 | 6.91E-04 | Rplp0/Rpl23/Rps | 7 | aPD-L1+aTGF-β vs aPD | Down-regulated gen | CD45+ |
| CC | GO:0032279 | asymmetric synapse | 1.51E-04 | 6.91E-04 | Rplp0/Rpl23/Rps | 7 | aPD-L1+aTGF-β vs aPD | Down-regulated gen | CD45+ |
| CC | GO:0098984 | neuron to neuron synapse | 1.96E-04 | 8.63E-04 | Rplp0/Rpl23/Rps | 7 | aPD-L1+aTGF-β vs aPD | Down-regulated gen | CD45+ |
| CC | GO:0005746 | mitochondrial respiratory chain | 2.07E-04 | 8.65E-04 | ND3/CYTB/ND1/N | 4 | aPD-L1+aTGF-β vs aPD | Down-regulated gen | CD45+ |
| CC | GO:0099572 | postsynaptic specialization | 2.12E-04 | 8.65E-04 | Rplp0/Rpl23/Rps | 7 | aPD-L1+aTGF-β vs aPD | Down-regulated gen | CD45+ |
| CC | GO:0016469 | proton-transporting two-sector ATPase complex | 5.43E-04 | 2.13E-03 | ATP8/Atp5k/ATP6 | 3 | aPD-L1+aTGF-β vs aPD | Down-regulated gen | CD45+ |
| CC | GO:1990204 | oxidoreductase complex | 5.76E-04 | 2.18E-03 | ND3/CYTB/ND1/N | 4 | aPD-L1+aTGF-β vs aPD | Down-regulated gen | CD45+ |
| CC | GO:0000276 | mitochondrial proton-transporting ATP synthase | 6.63E-04 | 2.43E-03 | ATP8/Atp5k | 2 | aPD-L1+aTGF-β vs aPD | Down-regulated gen | CD45+ |
| CC | GO:0005747 | mitochondrial respiratory chain complex I | 8.65E-04 | 2.88E-03 | ND3/ND1/ND2 | 3 | aPD-L1+aTGF-β vs aPD | Down-regulated gen | CD45+ |
| CC | GO:0030964 | NADH dehydrogenase complex | 8.65E-04 | 2.88E-03 | ND3/ND1/ND2 | 3 | aPD-L1+aTGF-β vs aPD | Down-regulated gen | CD45+ |
| CC | GO:0045271 | respiratory chain complex I | 8.65E-04 | 2.88E-03 | ND3/ND1/ND2 | 3 | aPD-L1+aTGF-β vs aPD | Down-regulated gen | CD45+ |
| CC | GO:0045277 | respiratory chain complex IV | 1.74E-03 | 5.63E-03 | COX3/COX2 | 2 | aPD-L1+aTGF-β vs aPD | Down-regulated gen | CD45+ |
| MF | GO:0003735 | structural constituent of ribosome | 4.36E-42 | 5.27E-40 | Rplp2/Rpl41/Rplp | 28 | aPD-L1+aTGF-β vs aPD | Down-regulated gen | CD45+ |
| MF | GO:0005198 | structural molecule activity | 1.65E-28 | 1.00E-26 | Rplp2/Rpl41/Rplp | 28 | aPD-L1+aTGF-β vs aPD | Down-regulated gen | CD45+ |
| MF | GO:0015078 | proton transmembrane transporter activity | 3.60E-05 | 1.45E-03 | ATP8/COX3/COX | 5 | aPD-L1+aTGF-β vs aPD | Down-regulated gen | CD45+ |
| MF | GO:0019843 | rRNA binding | 1.52E-04 | 4.24E-03 | Rplp0/Rpl37a/Rp | 4 | aPD-L1+aTGF-β vs aPD | Down-regulated gen | CD45+ |
| MF | GO:0003954 | NADH dehydrogenase activity | 1.75E-04 | 4.24E-03 | ND3/ND1/ND2 | 3 | aPD-L1+aTGF-β vs aPD | Down-regulated gen | CD45+ |
| MF | GO:0031369 | translation initiation factor binding | 3.73E-04 | 7.53E-03 | Rps3a1/Rps19/R | 3 | aPD-L1+aTGF-β vs aPD | Down-regulated gen | CD45+ |
| MF | GO:0015077 | monovalent inorganic cation transmembrane tra | 1.31E-03 | 2.26E-02 | ATP8/COX3/COX | 5 | aPD-L1+aTGF-β vs aPD | Down-regulated gen | CD45+ |
| MF | GO:0003729 | mRNA binding | 1.67E-03 | 2.47E-02 | Zfp36/Rpl13a/Rp | 5 | aPD-L1+aTGF-β vs aPD | Down-regulated gen | CD45+ |
| MF | GO:0009055 | electron transfer activity | 1.84E-03 | 2.47E-02 | COX3/COX2/CYT | 3 | aPD-L1+aTGF-β vs aPD | Down-regulated gen | CD45+ |
| MF | GO:0043021 | ribonucleoprotein complex binding | 2.19E-03 | 2.62E-02 | Rpl35a/Rpsa/Rpl | 4 | aPD-L1+aTGF-β vs aPD | Down-regulated gen | CD45+ |
| MF | GO:0016651 | oxidoreductase activity, acting on NAD(P)H | 2.71E-03 | 2.62E-02 | ND3/ND1/ND2 | 3 | aPD-L1+aTGF-β vs aPD | Down-regulated gen | CD45+ |
| MF | GO:0004129 | cytochrome-c oxidase activity | 3.03E-03 | 2.62E-02 | COX3/COX2 | 2 | aPD-L1+aTGF-β vs aPD | Down-regulated gen | CD45+ |
| MF | GO:0015002 | heme-copper terminal oxidase activity | 3.03E-03 | 2.62E-02 | COX3/COX2 | 2 | aPD-L1+aTGF-β vs aPD | Down-regulated gen | CD45+ |
| MF | GO:0016676 | oxidoreductase activity, acting on a heme group | 3.03E-03 | 2.62E-02 | COX3/COX2 | 2 | aPD-L1+aTGF-β vs aPD | Down-regulated gen | CD45+ |
| MF | GO:0016675 | oxidoreductase activity, acting on a heme group | 3.33E-03 | 2.69E-02 | COX3/COX2 | 2 | aPD-L1+aTGF-β vs aPD | Down-regulated gen | CD45+ |
| MF | GO:0008137 | NADH dehydrogenase (ubiquinone) activity | 4.29E-03 | 3.05E-02 | ND3/ND2 | 2 | aPD-L1+aTGF-β vs aPD | Down-regulated gen | CD45+ |
| MF | GO:0050136 | NADH dehydrogenase (quinone) activity | 4.29E-03 | 3.05E-02 | ND3/ND2 | 2 | aPD-L1+aTGF-β vs aPD | Down-regulated gen | CD45+ |
| MF | GO:0016655 | oxidoreductase activity, acting on NAD(P)H, quir | 7.40E-03 | 4.98E-02 | ND3/ND2 | 2 | aPD-L1+aTGF-β vs aPD | Down-regulated gen | CD45+ |
| BP | GO:0043062 | extracellular structure organization | 1.23E-06 | 1.73E-03 | Pdpn/Antxr1/Ddr2 | 10 | aPD-L1+aTGF-β vs aPD | Down-regulated gen | CD45- |
| BP | GO:0001501 | skeletal system development | 2.16E-06 | 1.73E-03 | Zfp385a/Osr1/Osr | 12 | aPD-L1+aTGF-β vs aPD | Down-regulated gen | CD45- |
| BP | GO:0030198 | extracellular matrix organization | 3.21E-06 | 1.73E-03 | Pdpn/Antxr1/Ddr2 | 9 | aPD-L1+aTGF-β vs aPD | Down-regulated gen | CD45- |
| BP | GO:0002062 | chondrocyte differentiation | 1.40E-05 | 5.52E-03 | Osr1/Osr2/Tgfbr2/ | 6 | aPD-L1+aTGF-β vs aPD | Down-regulated gen | CD45- |
| BP | GO:0055021 | regulation of cardiac muscle tissue growth | 2.35E-05 | 5.52E-03 | Fdps/Wnt2/Tgfbr2 | 5 | aPD-L1+aTGF-β vs aPD | Down-regulated gen | CD45- |
| BP | GO:0061448 | connective tissue development | 2.36E-05 | 5.52E-03 | Osr1/Osr2/Thbs3/ | 8 | aPD-L1+aTGF-β vs aPD | Down-regulated gen | CD45- |
| BP | GO:0051216 | cartilage development | 2.75E-05 | 5.52E-03 | Osr1/Osr2/Thbs3/ | 7 | aPD-L1+aTGF-β vs aPD | Down-regulated gen | CD45- |
| BP | GO:0060420 | regulation of heart growth | 3.35E-05 | 5.52E-03 | Fdps/Wnt2/Tgfbr2 | 5 | aPD-L1+aTGF-β vs aPD | Down-regulated gen | CD45- |
| BP | GO:0003416 | endochondral bone growth | 3.54E-05 | 5.52E-03 | Thbs3/Ddr2/Tgfbr | 4 | aPD-L1+aTGF-β vs aPD | Down-regulated gen | CD45- |
| BP | GO:0035265 | organ growth | 3.70E-05 | 5.52E-03 | Fdps/Thbs3/Ddr2/ | 7 | aPD-L1+aTGF-β vs aPD | Down-regulated gen | CD45- |
| BP | GO:1905048 | regulation of metallopeptidase activity | 3.76E-05 | 5.52E-03 | Antxr1/Timp2/Tim | 3 | aPD-L1+aTGF-β vs aPD | Down-regulated gen | CD45- |
| BP | GO:0098868 | bone growth | 4.40E-05 | 5.93E-03 | Thbs3/Ddr2/Tgfbr | 4 | aPD-L1+aTGF-β vs aPD | Down-regulated gen | CD45- |
| BP | GO:0055024 | regulation of cardiac muscle tissue development | 8.44E-05 | 9.90E-03 | Fdps/Wnt2/Tgfbr2 | 5 | aPD-L1+aTGF-β vs aPD | Down-regulated gen | CD45- |
| BP | GO:0043502 | regulation of muscle adaptation | 8.91E-05 | 9.90E-03 | Fdps/Klf4/Pi16/Co | 5 | aPD-L1+aTGF-β vs aPD | Down-regulated gen | CD45- |
| BP | GO:1901861 | regulation of muscle tissue development | 9.19E-05 | 9.90E-03 | Fdps/Wnt2/Tgfbr2 | 6 | aPD-L1+aTGF-β vs aPD | Down-regulated gen | CD45- |
| BP | GO:0055017 | cardiac muscle tissue growth | 1.10E-04 | 1.11E-02 | Fdps/Wnt2/Tgfbr2 | 5 | aPD-L1+aTGF-β vs aPD | Down-regulated gen | CD45- |
| BP | GO:0030199 | collagen fibril organization | 1.31E-04 | 1.25E-02 | Ddr2/Tnxb/Dpt/Co | 4 | aPD-L1+aTGF-β vs aPD | Down-regulated gen | CD45- |
| BP | GO:0046620 | regulation of organ growth | 1.49E-04 | 1.27E-02 | Fdps/Wnt2/Tgfbr2 | 5 | aPD-L1+aTGF-β vs aPD | Down-regulated gen | CD45- |
| BP | GO:0060419 | heart growth | 1.49E-04 | 1.27E-02 | Fdps/Wnt2/Tgfbr2 | 5 | aPD-L1+aTGF-β vs aPD | Down-regulated gen | CD45- |
| BP | GO:0043500 | muscle adaptation | 1.81E-04 | 1.46E-02 | Fdps/Klf4/Pi16/Co | 5 | aPD-L1+aTGF-β vs aPD | Down-regulated gen | CD45- |
| BP | GO:1905207 | regulation of cardiocyte differentiation | 1.90E-04 | 1.46E-02 | Fdps/Pi16/Sema3 | 4 | aPD-L1+aTGF-β vs aPD | Down-regulated gen | CD45- |
| BP | GO:0045995 | regulation of embryonic development | 2.93E-04 | 2.15E-02 | Osr1/Lama4/Osr2 | 5 | aPD-L1+aTGF-β vs aPD | Down-regulated gen | CD45- |
| BP | GO:0061050 | regulation of cell growth involved in cardiac mus | 3.25E-04 | 2.29E-02 | Fdps/Pi16/Col14a | 3 | aPD-L1+aTGF-β vs aPD | Down-regulated gen | CD45- |
| BP | GO:0014743 | regulation of muscle hypertrophy | 3.42E-04 | 2.30E-02 | Fdps/Pi16/Col14a | 4 | aPD-L1+aTGF-β vs aPD | Down-regulated gen | CD45- |
| BP | GO:0048762 | mesenchymal cell differentiation | 4.63E-04 | 2.89E-02 | Pdpn/Osr1/Wnt2/ | 6 | aPD-L1+aTGF-β vs aPD | Down-regulated gen | CD45- |
| BP | GO:0060348 | bone development | 4.90E-04 | 2.89E-02 | Zfp385a/Osr2/Thb | 6 | aPD-L1+aTGF-β vs aPD | Down-regulated gen | CD45- |
| BP | GO:0003417 | growth plate cartilage development | 5.01E-04 | 2.89E-02 | Thbs3/Tgfbr2/Col | 3 | aPD-L1+aTGF-β vs aPD | Down-regulated gen | CD45- |
| BP | GO:0048009 | insulin-like growth factor receptor signaling path | 5.01E-04 | 2.89E-02 | Grb10/Igfbp6/Igfb | 3 | aPD-L1+aTGF-β vs aPD | Down-regulated gen | CD45- |
| BP | GO:0010470 | regulation of gastrulation | 5.52E-04 | 3.08E-02 | Osr1/Osr2/Klf4 | 3 | aPD-L1+aTGF-β vs aPD | Down-regulated gen | CD45- |

| | | | | | | | | | |
|---|---|---|---|---|---|---|---|---|---|
| BP | GO:0001667 | ameboidal-type cell migration | 6.29E-04 | 3.20E-02 | Myh9/Ddr2/Dcn/T | 8 | aPD-L1+aTGF-β vs aPD | Down-regulated gen | CD45- |
| BP | GO:0060537 | muscle tissue development | 6.40E-04 | 3.20E-02 | Fdps/Osr1/Wnt2/T | 8 | aPD-L1+aTGF-β vs aPD | Down-regulated gen | CD45- |
| BP | GO:0040019 | positive regulation of embryonic development | 6.65E-04 | 3.20E-02 | Osr1/Osr2/Wnt2 | 3 | aPD-L1+aTGF-β vs aPD | Down-regulated gen | CD45- |
| BP | GO:0007369 | gastrulation | 6.89E-04 | 3.20E-02 | Osr1/Osr2/Tgfbr2/ | 5 | aPD-L1+aTGF-β vs aPD | Down-regulated gen | CD45- |
| BP | GO:0033002 | muscle cell proliferation | 7.27E-04 | 3.20E-02 | Wnt2/Tgfbr2/Mmp | 6 | aPD-L1+aTGF-β vs aPD | Down-regulated gen | CD45- |
| BP | GO:0016202 | regulation of striated muscle tissue development | 7.62E-04 | 3.20E-02 | Fdps/Wnt2/Tgfbr2 | 5 | aPD-L1+aTGF-β vs aPD | Down-regulated gen | CD45- |
| BP | GO:0003298 | physiological muscle hypertrophy | 7.92E-04 | 3.20E-02 | Fdps/Pi16/Col14a | 3 | aPD-L1+aTGF-β vs aPD | Down-regulated gen | CD45- |
| BP | GO:0003301 | physiological cardiac muscle hypertrophy | 7.92E-04 | 3.20E-02 | Fdps/Pi16/Col14a | 3 | aPD-L1+aTGF-β vs aPD | Down-regulated gen | CD45- |
| BP | GO:0061049 | cell growth involved in cardiac muscle cell devel | 7.92E-04 | 3.20E-02 | Fdps/Pi16/Col14a | 3 | aPD-L1+aTGF-β vs aPD | Down-regulated gen | CD45- |
| BP | GO:0110110 | positive regulation of animal organ morphogene | 8.00E-04 | 3.20E-02 | Wnt2/Tgfbr2/Cd34 | 4 | aPD-L1+aTGF-β vs aPD | Down-regulated gen | CD45- |
| BP | GO:0048705 | skeletal system morphogenesis | 8.04E-04 | 3.20E-02 | Osr1/Osr2/Thbs3/ | 6 | aPD-L1+aTGF-β vs aPD | Down-regulated gen | CD45- |
| BP | GO:0048634 | regulation of muscle organ development | 8.13E-04 | 3.20E-02 | Fdps/Wnt2/Tgfbr2 | 5 | aPD-L1+aTGF-β vs aPD | Down-regulated gen | CD45- |
| BP | GO:0048588 | developmental cell growth | 9.75E-04 | 3.71E-02 | Fdps/Tgfbr2/Pi16/ | 6 | aPD-L1+aTGF-β vs aPD | Down-regulated gen | CD45- |
| BP | GO:0014897 | striated muscle hypertrophy | 1.00E-03 | 3.71E-02 | Fdps/Pi16/Col14a | 4 | aPD-L1+aTGF-β vs aPD | Down-regulated gen | CD45- |
| BP | GO:0006509 | membrane protein ectodomain proteolysis | 1.01E-03 | 3.71E-02 | Myh9/Timp2/Timp | 3 | aPD-L1+aTGF-β vs aPD | Down-regulated gen | CD45- |
| BP | GO:0014896 | muscle hypertrophy | 1.05E-03 | 3.77E-02 | Fdps/Pi16/Col14a | 4 | aPD-L1+aTGF-β vs aPD | Down-regulated gen | CD45- |
| BP | GO:1903053 | regulation of extracellular matrix organization | 1.09E-03 | 3.81E-02 | Pdpn/Antxr1/Ddr2 | 3 | aPD-L1+aTGF-β vs aPD | Down-regulated gen | CD45- |
| BP | GO:0048659 | smooth muscle cell proliferation | 1.18E-03 | 3.81E-02 | Tgfbr2/Mmp2/Klf4 | 5 | aPD-L1+aTGF-β vs aPD | Down-regulated gen | CD45- |
| BP | GO:0060346 | bone trabecula formation | 1.18E-03 | 3.81E-02 | Thbs3/Mmp2 | 2 | aPD-L1+aTGF-β vs aPD | Down-regulated gen | CD45- |
| BP | GO:0072498 | embryonic skeletal joint development | 1.18E-03 | 3.81E-02 | Osr1/Osr2 | 2 | aPD-L1+aTGF-β vs aPD | Down-regulated gen | CD45- |
| BP | GO:0090091 | positive regulation of extracellular matrix disass | 1.18E-03 | 3.81E-02 | Pdpn/Ddr2 | 2 | aPD-L1+aTGF-β vs aPD | Down-regulated gen | CD45- |
| BP | GO:0060351 | cartilage development involved in endochondral | 1.26E-03 | 4.00E-02 | Thbs3/Tgfbr2/Col | 3 | aPD-L1+aTGF-β vs aPD | Down-regulated gen | CD45- |
| BP | GO:0060485 | mesenchyme development | 1.34E-03 | 4.17E-02 | Pdpn/Osr1/Wnt2/ | 6 | aPD-L1+aTGF-β vs aPD | Down-regulated gen | CD45- |
| BP | GO:0001503 | ossification | 1.44E-03 | 4.37E-02 | Osr1/Osr2/Thbs3/ | 7 | aPD-L1+aTGF-β vs aPD | Down-regulated gen | CD45- |
| BP | GO:0060349 | bone morphogenesis | 1.46E-03 | 4.37E-02 | Osr2/Thbs3/Tgfbr | 4 | aPD-L1+aTGF-β vs aPD | Down-regulated gen | CD45- |
| BP | GO:0048638 | regulation of developmental growth | 1.52E-03 | 4.47E-02 | Fdps/Wnt2/Tgfbr2 | 7 | aPD-L1+aTGF-β vs aPD | Down-regulated gen | CD45- |
| BP | GO:2000725 | regulation of cardiac muscle cell differentiation | 1.55E-03 | 4.47E-02 | Fdps/Pi16/Col14a | 3 | aPD-L1+aTGF-β vs aPD | Down-regulated gen | CD45- |
| BP | GO:0014745 | negative regulation of muscle adaptation | 1.72E-03 | 4.75E-02 | Klf4/Igfbp5 | 2 | aPD-L1+aTGF-β vs aPD | Down-regulated gen | CD45- |
| BP | GO:0016049 | cell growth | 1.74E-03 | 4.75E-02 | Fdps/Tgfbr2/Ccn3 | 8 | aPD-L1+aTGF-β vs aPD | Down-regulated gen | CD45- |
| BP | GO:0033619 | membrane protein proteolysis | 1.76E-03 | 4.75E-02 | Myh9/Timp2/Timp | 3 | aPD-L1+aTGF-β vs aPD | Down-regulated gen | CD45- |
| BP | GO:0030282 | bone mineralization | 1.77E-03 | 4.75E-02 | Osr1/Osr2/Ddr2/C | 4 | aPD-L1+aTGF-β vs aPD | Down-regulated gen | CD45- |
| BP | GO:0001525 | angiogenesis | 1.79E-03 | 4.75E-02 | Myh9/Dcn/Tgfbr2/ | 8 | aPD-L1+aTGF-β vs aPD | Down-regulated gen | CD45- |
| BP | GO:0051146 | striated muscle cell differentiation | 1.88E-03 | 4.88E-02 | Myh9/Fdps/Ccn3/ | 6 | aPD-L1+aTGF-β vs aPD | Down-regulated gen | CD45- |
| BP | GO:0051153 | regulation of striated muscle cell differentiation | 1.90E-03 | 4.88E-02 | Fdps/Ccn3/Pi16/C | 4 | aPD-L1+aTGF-β vs aPD | Down-regulated gen | CD45- |
| CC | GO:0031012 | extracellular matrix | 1.27E-18 | 1.80E-16 | Pcsk6/Prelp/Lama | 22 | aPD-L1+aTGF-β vs aPD | Down-regulated gen | CD45- |
| CC | GO:0062023 | collagen-containing extracellular matrix | 1.15E-16 | 8.11E-15 | Pcsk6/Prelp/Lama | 19 | aPD-L1+aTGF-β vs aPD | Down-regulated gen | CD45- |
| CC | GO:0009897 | external side of plasma membrane | 3.90E-06 | 1.83E-04 | Pdpn/Antxr2/B2m | 9 | aPD-L1+aTGF-β vs aPD | Down-regulated gen | CD45- |
| CC | GO:0005604 | basement membrane | 6.44E-06 | 2.27E-04 | Lama4/Timp2/Efe | 6 | aPD-L1+aTGF-β vs aPD | Down-regulated gen | CD45- |
| CC | GO:0098552 | side of membrane | 1.55E-04 | 4.37E-03 | Pdpn/Antxr2/B2m | 9 | aPD-L1+aTGF-β vs aPD | Down-regulated gen | CD45- |
| CC | GO:0005581 | collagen trimer | 3.05E-04 | 7.16E-03 | Dcn/Scara3/Emili | 4 | aPD-L1+aTGF-β vs aPD | Down-regulated gen | CD45- |
| MF | GO:0005201 | extracellular matrix structural constituent | 1.39E-12 | 2.10E-10 | Prelp/Lama4/Dcn | 12 | aPD-L1+aTGF-β vs aPD | Down-regulated gen | CD45- |
| MF | GO:0005539 | glycosaminoglycan binding | 3.81E-11 | 2.88E-09 | Pcsk6/Prelp/Thbs | 12 | aPD-L1+aTGF-β vs aPD | Down-regulated gen | CD45- |
| MF | GO:0008201 | heparin binding | 8.53E-10 | 4.29E-08 | Pcsk6/Prelp/Thbs | 10 | aPD-L1+aTGF-β vs aPD | Down-regulated gen | CD45- |
| MF | GO:0005198 | structural molecule activity | 3.58E-09 | 1.35E-07 | Rpl17/Rpl5/Cldn3 | 16 | aPD-L1+aTGF-β vs aPD | Down-regulated gen | CD45- |
| MF | GO:1901681 | sulfur compound binding | 9.19E-09 | 2.78E-07 | Pcsk6/Prelp/Thbs | 11 | aPD-L1+aTGF-β vs aPD | Down-regulated gen | CD45- |
| MF | GO:0005520 | insulin-like growth factor binding | 8.19E-06 | 2.06E-04 | Ccn3/Htra3/Igfbp6 | 4 | aPD-L1+aTGF-β vs aPD | Down-regulated gen | CD45- |
| MF | GO:0019838 | growth factor binding | 5.60E-05 | 1.21E-03 | Pcsk6/Tgfbr2/Ccn | 6 | aPD-L1+aTGF-β vs aPD | Down-regulated gen | CD45- |
| MF | GO:0004252 | serine-type endopeptidase activity | 6.97E-05 | 1.32E-03 | Pcsk6/Pamr1/Mm | 5 | aPD-L1+aTGF-β vs aPD | Down-regulated gen | CD45- |
| MF | GO:0008236 | serine-type peptidase activity | 1.63E-04 | 2.73E-03 | Pcsk6/Pamr1/Mm | 5 | aPD-L1+aTGF-β vs aPD | Down-regulated gen | CD45- |
| MF | GO:0017171 | serine hydrolase activity | 1.96E-04 | 2.96E-03 | Pcsk6/Pamr1/Mm | 5 | aPD-L1+aTGF-β vs aPD | Down-regulated gen | CD45- |
| MF | GO:0005518 | collagen binding | 2.75E-04 | 3.51E-03 | Antxr1/Ddr2/Dcn/ | 4 | aPD-L1+aTGF-β vs aPD | Down-regulated gen | CD45- |
| MF | GO:0005178 | integrin binding | 2.79E-04 | 3.51E-03 | Myh9/Timp2/Ccn3 | 5 | aPD-L1+aTGF-β vs aPD | Down-regulated gen | CD45- |
| MF | GO:0001968 | fibronectin binding | 5.12E-04 | 5.95E-03 | Mmp2/Igfbp6/Igfb | 3 | aPD-L1+aTGF-β vs aPD | Down-regulated gen | CD45- |
| MF | GO:0005509 | calcium ion binding | 1.25E-03 | 1.35E-02 | Thbs3/Man1a/Pa | 8 | aPD-L1+aTGF-β vs aPD | Down-regulated gen | CD45- |
| MF | GO:0008191 | metalloendopeptidase inhibitor activity | 1.46E-03 | 1.38E-02 | Timp2/Timp3 | 2 | aPD-L1+aTGF-β vs aPD | Down-regulated gen | CD45- |
| MF | GO:0031994 | insulin-like growth factor I binding | 1.46E-03 | 1.38E-02 | Igfbp6/Igfbp5 | 2 | aPD-L1+aTGF-β vs aPD | Down-regulated gen | CD45- |
| MF | GO:0051015 | actin filament binding | 1.66E-03 | 1.47E-02 | Myh9/Ablim1/Anb | 5 | aPD-L1+aTGF-β vs aPD | Down-regulated gen | CD45- |
| MF | GO:0004857 | enzyme inhibitor activity | 2.04E-03 | 1.71E-02 | Rpl5/Cdkn1c/Tim | 6 | aPD-L1+aTGF-β vs aPD | Down-regulated gen | CD45- |
| MF | GO:0050840 | extracellular matrix binding | 2.86E-03 | 2.22E-02 | Dcn/Cd248/Adam | 3 | aPD-L1+aTGF-β vs aPD | Down-regulated gen | CD45- |
| MF | GO:0004175 | endopeptidase activity | 3.04E-03 | 2.22E-02 | Pcsk6/Pamr1/Mm | 6 | aPD-L1+aTGF-β vs aPD | Down-regulated gen | CD45- |
| MF | GO:0050839 | cell adhesion molecule binding | 3.08E-03 | 2.22E-02 | Myh9/Timp2/Ccn3 | 5 | aPD-L1+aTGF-β vs aPD | Down-regulated gen | CD45- |
| MF | GO:0019199 | transmembrane receptor protein kinase activity | 4.83E-03 | 3.32E-02 | Ddr2/Tgfbr2/Efem | 3 | aPD-L1+aTGF-β vs aPD | Down-regulated gen | CD45- |
| MF | GO:0019843 | rRNA binding | 5.48E-03 | 3.49E-02 | Rpl17/Rpl5/Cavin | 3 | aPD-L1+aTGF-β vs aPD | Down-regulated gen | CD45- |
| MF | GO:0061134 | peptidase regulator activity | 5.55E-03 | 3.49E-02 | Timp2/Pi16/Fn1/T | 4 | aPD-L1+aTGF-β vs aPD | Down-regulated gen | CD45- |
| BP | GO:1903039 | positive regulation of leukocyte cell-cell adhesio | 4.58E-11 | 7.54E-08 | H2-Aa/Irf1/H2-Ab | 12 | aPD-L1+aTGF-β vs aTG | Up-regulated genes | CD45+ |
| BP | GO:1903037 | regulation of leukocyte cell-cell adhesion | 1.67E-10 | 1.36E-07 | H2-Aa/Irf1/H2-Ab | 13 | aPD-L1+aTGF-β vs aTG | Up-regulated genes | CD45+ |
| BP | GO:0022409 | positive regulation of cell-cell adhesion | 2.47E-10 | 1.36E-07 | H2-Aa/Irf1/H2-Ab | 12 | aPD-L1+aTGF-β vs aTG | Up-regulated genes | CD45+ |
| BP | GO:0035710 | CD4-positive, alpha-beta T cell activation | 4.01E-10 | 1.65E-07 | Irf1/Cd274/Satb1/ | 9 | aPD-L1+aTGF-β vs aTG | Up-regulated genes | CD45+ |
| BP | GO:0007159 | leukocyte cell-cell adhesion | 5.62E-10 | 1.85E-07 | H2-Aa/Irf1/H2-Ab | 13 | aPD-L1+aTGF-β vs aTG | Up-regulated genes | CD45+ |
| BP | GO:0042110 | T cell activation | 6.98E-10 | 1.92E-07 | H2-Aa/Irf1/H2-Ab | 15 | aPD-L1+aTGF-β vs aTG | Up-regulated genes | CD45+ |
| BP | GO:0046631 | alpha-beta T cell activation | 1.07E-09 | 2.52E-07 | Irf1/H2-Ab1/Cd27 | 10 | aPD-L1+aTGF-β vs aTG | Up-regulated genes | CD45+ |
| BP | GO:0019886 | antigen processing and presentation of exogenc | 2.33E-09 | 4.80E-07 | H2-Eb1/H2-Aa/H2 | 5 | aPD-L1+aTGF-β vs aTG | Up-regulated genes | CD45+ |
| BP | GO:0022407 | regulation of cell-cell adhesion | 4.08E-09 | 6.78E-07 | H2-Aa/Irf1/H2-Ab | 13 | aPD-L1+aTGF-β vs aTG | Up-regulated genes | CD45+ |
| BP | GO:0050863 | regulation of T cell activation | 4.11E-09 | 6.78E-07 | H2-Aa/Irf1/H2-Ab | 12 | aPD-L1+aTGF-β vs aTG | Up-regulated genes | CD45+ |
| BP | GO:0050870 | positive regulation of T cell activation | 5.70E-09 | 8.54E-07 | H2-Aa/Irf1/H2-Ab | 10 | aPD-L1+aTGF-β vs aTG | Up-regulated genes | CD45+ |
| BP | GO:2000514 | regulation of CD4-positive, alpha-beta T cell acti | 1.21E-08 | 1.66E-06 | Irf1/Cd274/Arg2/S | 7 | aPD-L1+aTGF-β vs aTG | Up-regulated genes | CD45+ |
| BP | GO:0002495 | antigen processing and presentation of peptide | 1.52E-08 | 1.86E-06 | H2-Eb1/H2-Aa/H2 | 5 | aPD-L1+aTGF-β vs aTG | Up-regulated genes | CD45+ |
| BP | GO:0046634 | regulation of alpha-beta T cell activation | 1.58E-08 | 1.86E-06 | Irf1/H2-Ab1/Cd27 | 8 | aPD-L1+aTGF-β vs aTG | Up-regulated genes | CD45+ |
| BP | GO:0002504 | antigen processing and presentation of peptide | 2.06E-08 | 2.26E-06 | H2-Eb1/H2-Aa/H2 | 5 | aPD-L1+aTGF-β vs aTG | Up-regulated genes | CD45+ |
| BP | GO:0050868 | negative regulation of T cell activation | 3.66E-08 | 3.77E-06 | H2-Aa/Irf1/H2-Ab | 8 | aPD-L1+aTGF-β vs aTG | Up-regulated genes | CD45+ |
| BP | GO:0045785 | positive regulation of cell adhesion | 7.24E-08 | 6.42E-06 | H2-Aa/Irf1/H2-Ab | 12 | aPD-L1+aTGF-β vs aTG | Up-regulated genes | CD45+ |
| BP | GO:1903038 | negative regulation of leukocyte cell-cell adhesi | 7.29E-08 | 6.42E-06 | H2-Aa/Irf1/H2-Ab | 8 | aPD-L1+aTGF-β vs aTG | Up-regulated genes | CD45+ |
| BP | GO:0002478 | antigen processing and presentation of exogenc | 7.40E-08 | 6.42E-06 | H2-Eb1/H2-Aa/H2 | 5 | aPD-L1+aTGF-β vs aTG | Up-regulated genes | CD45+ |
| BP | GO:1903706 | regulation of hemopoiesis | 8.69E-08 | 7.16E-06 | H2-Aa/Irf1/Cd74/ | 12 | aPD-L1+aTGF-β vs aTG | Up-regulated genes | CD45+ |
| BP | GO:0051251 | positive regulation of lymphocyte activation | 1.22E-07 | 9.57E-06 | H2-Aa/Irf1/H2-Ab | 10 | aPD-L1+aTGF-β vs aTG | Up-regulated genes | CD45+ |
| BP | GO:0051249 | regulation of lymphocyte activation | 1.35E-07 | 1.01E-05 | H2-Aa/Irf1/H2-Ab | 12 | aPD-L1+aTGF-β vs aTG | Up-regulated genes | CD45+ |
| BP | GO:0019882 | antigen processing and presentation | 1.47E-07 | 1.01E-05 | H2-Eb1/H2-Aa/H2 | 7 | aPD-L1+aTGF-β vs aTG | Up-regulated genes | CD45+ |
| BP | GO:0045582 | positive regulation of T cell differentiation | 1.47E-07 | 1.01E-05 | H2-Aa/Irf1/Cd74/ | 7 | aPD-L1+aTGF-β vs aTG | Up-regulated genes | CD45+ |
| BP | GO:0030217 | T cell differentiation | 1.73E-07 | 1.14E-05 | H2-Aa/Irf1/Cd74/ | 10 | aPD-L1+aTGF-β vs aTG | Up-regulated genes | CD45+ |

| | | | | | | | | | |
|---|---|---|---|---|---|---|---|---|---|
| BP | GO:0019884 | antigen processing and presentation of exogenc | 2.03E-07 | 1.26E-05 | H2-Eb1/H2-Aa/H2 | 5 | aPD-L1+aTGF-β vs aTG | Up-regulated genes | CD45+ |
| BP | GO:0045580 | regulation of T cell differentiation | 2.06E-07 | 1.26E-05 | H2-Aa/Irf1/Cd74/[ | 8 | aPD-L1+aTGF-β vs aTG | Up-regulated genes | CD45+ |
| BP | GO:0042098 | T cell proliferation | 2.16E-07 | 1.27E-05 | H2-Aa/Irf1/H2-Ab | 9 | aPD-L1+aTGF-β vs aTG | Up-regulated genes | CD45+ |
| BP | GO:0051250 | negative regulation of lymphocyte activation | 2.43E-07 | 1.38E-05 | H2-Aa/Irf1/H2-Ab | 8 | aPD-L1+aTGF-β vs aTG | Up-regulated genes | CD45+ |
| BP | GO:0046651 | lymphocyte proliferation | 3.00E-07 | 1.65E-05 | H2-Aa/Irf1/H2-Ab | 10 | aPD-L1+aTGF-β vs aTG | Up-regulated genes | CD45+ |
| BP | GO:0032943 | mononuclear cell proliferation | 3.22E-07 | 1.66E-05 | H2-Aa/Irf1/H2-Ab | 10 | aPD-L1+aTGF-β vs aTG | Up-regulated genes | CD45+ |
| BP | GO:0030098 | lymphocyte differentiation | 3.22E-07 | 1.66E-05 | H2-Aa/Irf1/H2-Ab | 11 | aPD-L1+aTGF-β vs aTG | Up-regulated genes | CD45+ |
| BP | GO:0050670 | regulation of lymphocyte proliferation | 3.62E-07 | 1.81E-05 | H2-Aa/Irf1/H2-Ab | 9 | aPD-L1+aTGF-β vs aTG | Up-regulated genes | CD45+ |
| BP | GO:0045621 | positive regulation of lymphocyte differentiation | 3.84E-07 | 1.85E-05 | H2-Aa/Irf1/Cd74/[ | 7 | aPD-L1+aTGF-β vs aTG | Up-regulated genes | CD45+ |
| BP | GO:0032944 | regulation of mononuclear cell proliferation | 3.94E-07 | 1.85E-05 | H2-Aa/Irf1/H2-Ab | 9 | aPD-L1+aTGF-β vs aTG | Up-regulated genes | CD45+ |
| BP | GO:0070663 | regulation of leukocyte proliferation | 5.45E-07 | 2.37E-05 | H2-Aa/Irf1/H2-Ab | 9 | aPD-L1+aTGF-β vs aTG | Up-regulated genes | CD45+ |
| BP | GO:0070661 | leukocyte proliferation | 5.56E-07 | 2.37E-05 | H2-Aa/Irf1/H2-Ab | 10 | aPD-L1+aTGF-β vs aTG | Up-regulated genes | CD45+ |
| BP | GO:0042129 | regulation of T cell proliferation | 5.61E-07 | 2.37E-05 | H2-Aa/Irf1/H2-Ab | 8 | aPD-L1+aTGF-β vs aTG | Up-regulated genes | CD45+ |
| BP | GO:1902107 | positive regulation of leukocyte differentiation | 5.61E-07 | 2.37E-05 | H2-Aa/Irf1/Cd74/( | 8 | aPD-L1+aTGF-β vs aTG | Up-regulated genes | CD45+ |
| BP | GO:0002696 | positive regulation of leukocyte activation | 6.33E-07 | 2.59E-05 | H2-Aa/Irf1/H2-Ab | 10 | aPD-L1+aTGF-β vs aTG | Up-regulated genes | CD45+ |
| BP | GO:0002250 | adaptive immune response | 6.45E-07 | 2.59E-05 | Irf1/H2-Ab1/Cd74 | 11 | aPD-L1+aTGF-β vs aTG | Up-regulated genes | CD45+ |
| BP | GO:0022408 | negative regulation of cell-cell adhesion | 6.81E-07 | 2.67E-05 | H2-Aa/Irf1/H2-Ab | 8 | aPD-L1+aTGF-β vs aTG | Up-regulated genes | CD45+ |
| BP | GO:0002695 | negative regulation of leukocyte activation | 7.85E-07 | 3.01E-05 | H2-Aa/Irf1/H2-Ab | 8 | aPD-L1+aTGF-β vs aTG | Up-regulated genes | CD45+ |
| BP | GO:0045619 | regulation of lymphocyte differentiation | 9.02E-07 | 3.38E-05 | H2-Aa/Irf1/Cd74/[ | 8 | aPD-L1+aTGF-β vs aTG | Up-regulated genes | CD45+ |
| BP | GO:0009617 | response to bacterium | 9.64E-07 | 3.53E-05 | Cxcl10/Rpl39/Ccl | 12 | aPD-L1+aTGF-β vs aTG | Up-regulated genes | CD45+ |
| BP | GO:0050867 | positive regulation of cell activation | 9.85E-07 | 3.53E-05 | H2-Aa/Irf1/H2-Ab | 10 | aPD-L1+aTGF-β vs aTG | Up-regulated genes | CD45+ |
| BP | GO:0046636 | negative regulation of alpha-beta T cell activatio | 1.08E-06 | 3.79E-05 | Irf1/Cd274/Arg2/S | 5 | aPD-L1+aTGF-β vs aTG | Up-regulated genes | CD45+ |
| BP | GO:0002694 | regulation of leukocyte activation | 1.11E-06 | 3.80E-05 | H2-Aa/Irf1/H2-Ab | 12 | aPD-L1+aTGF-β vs aTG | Up-regulated genes | CD45+ |
| BP | GO:0043367 | CD4-positive, alpha-beta T cell differentiation | 1.25E-06 | 4.21E-05 | Irf1/Satb1/Rsad2/ | 6 | aPD-L1+aTGF-β vs aTG | Up-regulated genes | CD45+ |
| BP | GO:0002821 | positive regulation of adaptive immune response | 1.33E-06 | 4.37E-05 | Irf1/H2-Ab1/Cd74 | 7 | aPD-L1+aTGF-β vs aTG | Up-regulated genes | CD45+ |
| BP | GO:0002181 | cytoplasmic translation | 1.35E-06 | 4.38E-05 | Rpl39/Rps21/Rpl | 6 | aPD-L1+aTGF-β vs aTG | Up-regulated genes | CD45+ |
| BP | GO:0002819 | regulation of adaptive immune response | 1.47E-06 | 4.66E-05 | Irf1/H2-Ab1/Cd74 | 8 | aPD-L1+aTGF-β vs aTG | Up-regulated genes | CD45+ |
| BP | GO:0050866 | negative regulation of cell activation | 1.53E-06 | 4.77E-05 | H2-Aa/Irf1/H2-Ab | 8 | aPD-L1+aTGF-β vs aTG | Up-regulated genes | CD45+ |
| BP | GO:0034341 | response to interferon-gamma | 1.57E-06 | 4.78E-05 | H2-Eb1/H2-Aa/Irf | 7 | aPD-L1+aTGF-β vs aTG | Up-regulated genes | CD45+ |
| BP | GO:0002521 | leukocyte differentiation | 2.10E-06 | 6.30E-05 | H2-Aa/Irf1/H2-Ab | 12 | aPD-L1+aTGF-β vs aTG | Up-regulated genes | CD45+ |
| BP | GO:0050865 | regulation of cell activation | 2.24E-06 | 6.59E-05 | H2-Aa/Irf1/H2-Ab | 12 | aPD-L1+aTGF-β vs aTG | Up-regulated genes | CD45+ |
| BP | GO:1903708 | positive regulation of hemopoiesis | 2.61E-06 | 7.54E-05 | H2-Aa/Irf1/Cd74/( | 8 | aPD-L1+aTGF-β vs aTG | Up-regulated genes | CD45+ |
| BP | GO:0002683 | negative regulation of immune system process | 2.75E-06 | 7.82E-05 | H2-Aa/Irf1/H2-Ab | 11 | aPD-L1+aTGF-β vs aTG | Up-regulated genes | CD45+ |
| BP | GO:1902105 | regulation of leukocyte differentiation | 3.36E-06 | 9.39E-05 | H2-Aa/Irf1/Cd74/( | 9 | aPD-L1+aTGF-β vs aTG | Up-regulated genes | CD45+ |
| BP | GO:2000106 | regulation of leukocyte apoptotic process | 3.44E-06 | 9.44E-05 | Cd74/Ccl5/Cd274 | 6 | aPD-L1+aTGF-β vs aTG | Up-regulated genes | CD45+ |
| BP | GO:0048002 | antigen processing and presentation of peptide | 4.09E-06 | 1.09E-04 | H2-Eb1/H2-Aa/H2 | 5 | aPD-L1+aTGF-β vs aTG | Up-regulated genes | CD45+ |
| BP | GO:2000107 | negative regulation of leukocyte apoptotic proce | 4.09E-06 | 1.09E-04 | Cd74/Ccl5/Arg2/( | 5 | aPD-L1+aTGF-β vs aTG | Up-regulated genes | CD45+ |
| BP | GO:0036037 | CD8-positive, alpha-beta T cell activation | 5.02E-06 | 1.31E-04 | Irf1/Cd274/Satb1/ | 4 | aPD-L1+aTGF-β vs aTG | Up-regulated genes | CD45+ |
| BP | GO:0046632 | alpha-beta T cell differentiation | 7.60E-06 | 1.96E-04 | Irf1/Satb1/Rsad2/ | 6 | aPD-L1+aTGF-β vs aTG | Up-regulated genes | CD45+ |
| BP | GO:0043372 | positive regulation of CD4-positive, alpha-beta T | 8.04E-06 | 2.01E-04 | Irf1/Socs1/Ccr7/C | 4 | aPD-L1+aTGF-β vs aTG | Up-regulated genes | CD45+ |
| BP | GO:0045589 | regulation of regulatory T cell differentiation | 8.04E-06 | 2.01E-04 | Irf1/Dusp10/Socs | 4 | aPD-L1+aTGF-β vs aTG | Up-regulated genes | CD45+ |
| BP | GO:0070228 | regulation of lymphocyte apoptotic process | 8.36E-06 | 2.06E-04 | Cd74/Ccl5/Cd274 | 5 | aPD-L1+aTGF-β vs aTG | Up-regulated genes | CD45+ |
| BP | GO:0042130 | negative regulation of T cell proliferation | 9.83E-06 | 2.38E-04 | H2-Aa/H2-Ab1/Cc | 5 | aPD-L1+aTGF-β vs aTG | Up-regulated genes | CD45+ |
| BP | GO:0071887 | leukocyte apoptotic process | 1.11E-05 | 2.66E-04 | Cd74/Ccl5/Cd274 | 6 | aPD-L1+aTGF-β vs aTG | Up-regulated genes | CD45+ |
| BP | GO:0002579 | positive regulation of antigen processing and pre | 1.13E-05 | 2.66E-04 | H2-Ab1/Cd74/Ccr | 3 | aPD-L1+aTGF-β vs aTG | Up-regulated genes | CD45+ |
| BP | GO:0002828 | regulation of type 2 immune response | 1.22E-05 | 2.76E-04 | Irf1/Cd74/Rsad2/ | 4 | aPD-L1+aTGF-β vs aTG | Up-regulated genes | CD45+ |
| BP | GO:0045066 | regulatory T cell differentiation | 1.22E-05 | 2.76E-04 | Irf1/Dusp10/Socs | 4 | aPD-L1+aTGF-β vs aTG | Up-regulated genes | CD45+ |
| BP | GO:2000515 | negative regulation of CD4-positive, alpha-beta | 1.22E-05 | 2.76E-04 | Irf1/Cd274/Arg2/C | 4 | aPD-L1+aTGF-β vs aTG | Up-regulated genes | CD45+ |
| BP | GO:0046635 | positive regulation of alpha-beta T cell activation | 1.24E-05 | 2.76E-04 | Irf1/H2-Ab1/Socs | 5 | aPD-L1+aTGF-β vs aTG | Up-regulated genes | CD45+ |
| BP | GO:0007162 | negative regulation of cell adhesion | 1.29E-05 | 2.83E-04 | H2-Aa/Irf1/H2-Ab | 8 | aPD-L1+aTGF-β vs aTG | Up-regulated genes | CD45+ |
| BP | GO:0009615 | response to virus | 1.33E-05 | 2.88E-04 | Irf1/Cxcl10/Ccl5/C | 8 | aPD-L1+aTGF-β vs aTG | Up-regulated genes | CD45+ |
| BP | GO:0035739 | CD4-positive, alpha-beta T cell proliferation | 1.55E-05 | 3.27E-04 | Cd274/Arg2/Cd4 | 3 | aPD-L1+aTGF-β vs aTG | Up-regulated genes | CD45+ |
| BP | GO:2000561 | regulation of CD4-positive, alpha-beta T cell pro | 1.55E-05 | 3.27E-04 | Cd274/Arg2/Cd4 | 3 | aPD-L1+aTGF-β vs aTG | Up-regulated genes | CD45+ |
| BP | GO:2000516 | positive regulation of CD4-positive, alpha-beta T | 1.78E-05 | 3.72E-04 | Irf1/Socs1/Ccr7/C | 4 | aPD-L1+aTGF-β vs aTG | Up-regulated genes | CD45+ |
| BP | GO:0070229 | negative regulation of lymphocyte apoptotic proc | 2.01E-05 | 4.14E-04 | Cd74/Ccl5/Arg2/( | 4 | aPD-L1+aTGF-β vs aTG | Up-regulated genes | CD45+ |
| BP | GO:0032945 | negative regulation of mononuclear cell prolifera | 2.50E-05 | 4.93E-04 | H2-Aa/H2-Ab1/Cc | 5 | aPD-L1+aTGF-β vs aTG | Up-regulated genes | CD45+ |
| BP | GO:0050672 | negative regulation of lymphocyte proliferation | 2.50E-05 | 4.93E-04 | H2-Aa/H2-Ab1/Cc | 5 | aPD-L1+aTGF-β vs aTG | Up-regulated genes | CD45+ |
| BP | GO:0042092 | type 2 immune response | 2.51E-05 | 4.93E-04 | Irf1/Cd74/Rsad2/ | 4 | aPD-L1+aTGF-β vs aTG | Up-regulated genes | CD45+ |
| BP | GO:0046640 | regulation of alpha-beta T cell proliferation | 2.51E-05 | 4.93E-04 | Irf1/Cd274/Arg2/C | 4 | aPD-L1+aTGF-β vs aTG | Up-regulated genes | CD45+ |
| BP | GO:0010819 | regulation of T cell chemotaxis | 2.66E-05 | 5.05E-04 | Cxcl10/Ccl5/Ccr7 | 3 | aPD-L1+aTGF-β vs aTG | Up-regulated genes | CD45+ |
| BP | GO:0042532 | negative regulation of tyrosine phosphorylation | 2.66E-05 | 5.05E-04 | Irf1/Socs1/Socs3 | 3 | aPD-L1+aTGF-β vs aTG | Up-regulated genes | CD45+ |
| BP | GO:0046642 | negative regulation of alpha-beta T cell prolifera | 2.66E-05 | 5.05E-04 | Cd274/Arg2/Cd4 | 3 | aPD-L1+aTGF-β vs aTG | Up-regulated genes | CD45+ |
| BP | GO:0002687 | positive regulation of leukocyte migration | 2.91E-05 | 5.45E-04 | Cd74/Cxcl10/Ccl | 6 | aPD-L1+aTGF-β vs aTG | Up-regulated genes | CD45+ |
| BP | GO:0070227 | lymphocyte apoptotic process | 3.03E-05 | 5.55E-04 | Cd74/Ccl5/Cd274 | 5 | aPD-L1+aTGF-β vs aTG | Up-regulated genes | CD45+ |
| BP | GO:0070664 | negative regulation of leukocyte proliferation | 3.03E-05 | 5.55E-04 | H2-Aa/H2-Ab1/Cc | 5 | aPD-L1+aTGF-β vs aTG | Up-regulated genes | CD45+ |
| BP | GO:0043374 | CD8-positive, alpha-beta T cell differentiation | 3.38E-05 | 6.12E-04 | Irf1/Satb1/Socs1 | 3 | aPD-L1+aTGF-β vs aTG | Up-regulated genes | CD45+ |
| BP | GO:0046633 | alpha-beta T cell proliferation | 3.44E-05 | 6.17E-04 | Irf1/Cd274/Arg2/C | 4 | aPD-L1+aTGF-β vs aTG | Up-regulated genes | CD45+ |
| BP | GO:1903707 | negative regulation of hemopoiesis | 4.80E-05 | 8.51E-04 | Irf1/Cd74/Socs1/N | 6 | aPD-L1+aTGF-β vs aTG | Up-regulated genes | CD45+ |
| BP | GO:0098542 | defense response to other organism | 4.97E-05 | 8.72E-04 | Irf1/Cxcl10/Rpl39 | 9 | aPD-L1+aTGF-β vs aTG | Up-regulated genes | CD45+ |
| BP | GO:0045581 | negative regulation of T cell differentiation | 5.04E-05 | 8.74E-04 | Irf1/Cd74/Socs1/( | 4 | aPD-L1+aTGF-β vs aTG | Up-regulated genes | CD45+ |
| BP | GO:1902532 | negative regulation of intracellular signal transdu | 5.09E-05 | 8.74E-04 | Irf1/Cd74/Dusp10 | 10 | aPD-L1+aTGF-β vs aTG | Up-regulated genes | CD45+ |
| BP | GO:0046638 | positive regulation of alpha-beta T cell differentia | 5.52E-05 | 9.37E-04 | Irf1/Socs1/Ccr7/C | 4 | aPD-L1+aTGF-β vs aTG | Up-regulated genes | CD45+ |
| BP | GO:0010818 | T cell chemotaxis | 6.25E-05 | 1.04E-03 | Cxcl10/Ccl5/Ccr7 | 3 | aPD-L1+aTGF-β vs aTG | Up-regulated genes | CD45+ |
| BP | GO:2001185 | regulation of CD8-positive, alpha-beta T cell acti | 6.25E-05 | 1.04E-03 | Irf1/Cd274/Socs1 | 3 | aPD-L1+aTGF-β vs aTG | Up-regulated genes | CD45+ |
| BP | GO:0050730 | regulation of peptidyl-tyrosine phosphorylation | 6.40E-05 | 1.05E-03 | Irf1/Cd74/Ccl5/So | 7 | aPD-L1+aTGF-β vs aTG | Up-regulated genes | CD45+ |
| BP | GO:0043370 | regulation of CD4-positive, alpha-beta T cell diffe | 6.56E-05 | 1.07E-03 | Irf1/Socs1/Ccr7/C | 4 | aPD-L1+aTGF-β vs aTG | Up-regulated genes | CD45+ |
| BP | GO:2001243 | negative regulation of intrinsic apoptotic signalin | 6.71E-05 | 1.08E-03 | Cd74/Dnaja1/Pla | 5 | aPD-L1+aTGF-β vs aTG | Up-regulated genes | CD45+ |
| BP | GO:0001819 | positive regulation of cytokine production | 6.76E-05 | 1.08E-03 | Irf1/Cd74/Ccl5/Re | 9 | aPD-L1+aTGF-β vs aTG | Up-regulated genes | CD45+ |
| BP | GO:0000028 | ribosomal small subunit assembly | 7.48E-05 | 1.17E-03 | Rps27/Rpl38/Rps | 3 | aPD-L1+aTGF-β vs aTG | Up-regulated genes | CD45+ |
| BP | GO:0002577 | regulation of antigen processing and presentatic | 7.48E-05 | 1.17E-03 | H2-Ab1/Cd74/Ccr | 3 | aPD-L1+aTGF-β vs aTG | Up-regulated genes | CD45+ |
| BP | GO:0032496 | response to lipopolysaccharide | 8.26E-05 | 1.28E-03 | Cxcl10/Ccl5/Dusp | 7 | aPD-L1+aTGF-β vs aTG | Up-regulated genes | CD45+ |
| BP | GO:1901623 | regulation of lymphocyte chemotaxis | 8.85E-05 | 1.36E-03 | Cxcl10/Ccl5/Ccr7 | 3 | aPD-L1+aTGF-β vs aTG | Up-regulated genes | CD45+ |
| BP | GO:0045620 | negative regulation of lymphocyte differentiation | 9.06E-05 | 1.38E-03 | Irf1/Cd74/Socs1/( | 4 | aPD-L1+aTGF-β vs aTG | Up-regulated genes | CD45+ |
| BP | GO:0072678 | T cell migration | 1.13E-04 | 1.72E-03 | Cxcl10/Ccl5/Icam | 4 | aPD-L1+aTGF-β vs aTG | Up-regulated genes | CD45+ |
| BP | GO:0071346 | cellular response to interferon-gamma | 1.20E-04 | 1.79E-03 | Irf1/H2-Ab1/Ccl5/ | 5 | aPD-L1+aTGF-β vs aTG | Up-regulated genes | CD45+ |
| BP | GO:1990266 | neutrophil migration | 1.20E-04 | 1.79E-03 | Cd74/Cxcl10/Ccl | 5 | aPD-L1+aTGF-β vs aTG | Up-regulated genes | CD45+ |
| BP | GO:0002237 | response to molecule of bacterial origin | 1.30E-04 | 1.91E-03 | Cxcl10/Ccl5/Dusp | 7 | aPD-L1+aTGF-β vs aTG | Up-regulated genes | CD45+ |
| BP | GO:0002827 | positive regulation of T-helper 1 type immune re | 1.39E-04 | 2.03E-03 | Irf1/H2-Ab1/Ccr7 | 3 | aPD-L1+aTGF-β vs aTG | Up-regulated genes | CD45+ |
| BP | GO:0019058 | viral life cycle | 1.54E-04 | 2.22E-03 | Vps37b/Cd74/Ccl | 6 | aPD-L1+aTGF-β vs aTG | Up-regulated genes | CD45+ |

| | | | | | | | | | |
|---|---|---|---|---|---|---|---|---|---|
| BP | GO:0046426 | negative regulation of JAK-STAT cascade | 1.60E-04 | 2.27E-03 | Irf1/Socs1/Socs3 | 3 | aPD-L1+aTGF-β vs aTG | Up-regulated genes | CD45+ |
| BP | GO:1904893 | negative regulation of STAT cascade | 1.60E-04 | 2.27E-03 | Irf1/Socs1/Socs3 | 3 | aPD-L1+aTGF-β vs aTG | Up-regulated genes | CD45+ |
| BP | GO:0002824 | positive regulation of adaptive immune response | 1.78E-04 | 2.50E-03 | Irf1/H2-Ab1/Cd27 | 5 | aPD-L1+aTGF-β vs aTG | Up-regulated genes | CD45+ |
| BP | GO:0042274 | ribosomal small subunit biogenesis | 1.83E-04 | 2.53E-03 | Rps21/Rps27/Rpl | 4 | aPD-L1+aTGF-β vs aTG | Up-regulated genes | CD45+ |
| BP | GO:0042509 | regulation of tyrosine phosphorylation of STAT p | 1.83E-04 | 2.53E-03 | Irf1/Ccl5/Socs1/Sc | 4 | aPD-L1+aTGF-β vs aTG | Up-regulated genes | CD45+ |
| BP | GO:0046637 | regulation of alpha-beta T cell differentiation | 1.95E-04 | 2.66E-03 | Irf1/Socs1/Ccr7/C | 4 | aPD-L1+aTGF-β vs aTG | Up-regulated genes | CD45+ |
| BP | GO:0002685 | regulation of leukocyte migration | 1.96E-04 | 2.66E-03 | Cd74/Cxcl10/Ccl5 | 6 | aPD-L1+aTGF-β vs aTG | Up-regulated genes | CD45+ |
| BP | GO:0002460 | adaptive immune response based on somatic re | 1.97E-04 | 2.66E-03 | Irf1/H2-Ab1/Cd74 | 7 | aPD-L1+aTGF-β vs aTG | Up-regulated genes | CD45+ |
| BP | GO:0018108 | peptidyl-tyrosine phosphorylation | 2.16E-04 | 2.90E-03 | Irf1/Cd74/Ccl5/So | 7 | aPD-L1+aTGF-β vs aTG | Up-regulated genes | CD45+ |
| BP | GO:0018212 | peptidyl-tyrosine modification | 2.32E-04 | 3.08E-03 | Irf1/Cd74/Ccl5/So | 7 | aPD-L1+aTGF-β vs aTG | Up-regulated genes | CD45+ |
| BP | GO:0007260 | tyrosine phosphorylation of STAT protein | 2.34E-04 | 3.08E-03 | Irf1/Ccl5/Socs1/Sc | 4 | aPD-L1+aTGF-β vs aTG | Up-regulated genes | CD45+ |
| BP | GO:0019221 | cytokine-mediated signaling pathway | 2.72E-04 | 3.55E-03 | Irf1/Cd74/Cxcl10/ | 7 | aPD-L1+aTGF-β vs aTG | Up-regulated genes | CD45+ |
| BP | GO:1903900 | regulation of viral life cycle | 2.85E-04 | 3.69E-03 | Vps37b/Cd74/Ccl | 5 | aPD-L1+aTGF-β vs aTG | Up-regulated genes | CD45+ |
| BP | GO:0097530 | granulocyte migration | 3.06E-04 | 3.94E-03 | Cd74/Cxcl10/Ccl5 | 5 | aPD-L1+aTGF-β vs aTG | Up-regulated genes | CD45+ |
| BP | GO:0051607 | defense response to virus | 3.18E-04 | 4.06E-03 | Irf1/Cxcl10/Ccl5/C | 6 | aPD-L1+aTGF-β vs aTG | Up-regulated genes | CD45+ |
| BP | GO:0002825 | regulation of T-helper 1 type immune response | 3.23E-04 | 4.09E-03 | Irf1/H2-Ab1/Ccr7 | 3 | aPD-L1+aTGF-β vs aTG | Up-regulated genes | CD45+ |
| BP | GO:2001234 | negative regulation of apoptotic signaling pathw | 3.73E-04 | 4.69E-03 | Cd74/Dnaja1/Plau | 6 | aPD-L1+aTGF-β vs aTG | Up-regulated genes | CD45+ |
| BP | GO:1902624 | positive regulation of neutrophil migration | 3.94E-04 | 4.89E-03 | Cd74/Il1a/Ccr7 | 3 | aPD-L1+aTGF-β vs aTG | Up-regulated genes | CD45+ |
| BP | GO:2000406 | positive regulation of T cell migration | 3.94E-04 | 4.89E-03 | Cxcl10/Ccl5/Ccr7 | 3 | aPD-L1+aTGF-β vs aTG | Up-regulated genes | CD45+ |
| BP | GO:2000108 | positive regulation of leukocyte apoptotic proces | 4.34E-04 | 5.34E-03 | Ccl5/Cd274/Cd44 | 3 | aPD-L1+aTGF-β vs aTG | Up-regulated genes | CD45+ |
| BP | GO:0032642 | regulation of chemokine production | 4.48E-04 | 5.47E-03 | Cd74/Ccl5/Il1a/Ar | 4 | aPD-L1+aTGF-β vs aTG | Up-regulated genes | CD45+ |
| BP | GO:0032735 | positive regulation of interleukin-12 production | 4.76E-04 | 5.76E-03 | Irf1/Rel/Ccr7 | 3 | aPD-L1+aTGF-β vs aTG | Up-regulated genes | CD45+ |
| BP | GO:1901652 | response to peptide | 4.88E-04 | 5.88E-03 | Irf1/Serpina3g/So | 7 | aPD-L1+aTGF-β vs aTG | Up-regulated genes | CD45+ |
| BP | GO:0002690 | positive regulation of leukocyte chemotaxis | 4.94E-04 | 5.90E-03 | Cd74/Cxcl10/Ccl5 | 4 | aPD-L1+aTGF-β vs aTG | Up-regulated genes | CD45+ |
| BP | GO:2000403 | positive regulation of lymphocyte migration | 5.20E-04 | 6.16E-03 | Cxcl10/Ccl5/Ccr7 | 3 | aPD-L1+aTGF-β vs aTG | Up-regulated genes | CD45+ |
| BP | GO:0032602 | chemokine production | 5.44E-04 | 6.40E-03 | Cd74/Ccl5/Il1a/Ar | 4 | aPD-L1+aTGF-β vs aTG | Up-regulated genes | CD45+ |
| BP | GO:0002791 | regulation of peptide secretion | 6.27E-04 | 7.33E-03 | Cd74/Ccl5/Cd274 | 8 | aPD-L1+aTGF-β vs aTG | Up-regulated genes | CD45+ |
| BP | GO:0002526 | acute inflammatory response | 6.52E-04 | 7.57E-03 | Ccl5/Il1a/Icam1/C | 4 | aPD-L1+aTGF-β vs aTG | Up-regulated genes | CD45+ |
| BP | GO:0044403 | symbiont process | 6.64E-04 | 7.65E-03 | Vps37b/Cd74/Ccl | 7 | aPD-L1+aTGF-β vs aTG | Up-regulated genes | CD45+ |
| BP | GO:0072676 | lymphocyte migration | 6.82E-04 | 7.80E-03 | Cxcl10/Ccl5/Icam | 4 | aPD-L1+aTGF-β vs aTG | Up-regulated genes | CD45+ |
| BP | GO:0030593 | neutrophil chemotaxis | 7.12E-04 | 8.09E-03 | Cd74/Cxcl10/Ccl5 | 4 | aPD-L1+aTGF-β vs aTG | Up-regulated genes | CD45+ |
| BP | GO:0046328 | regulation of JNK cascade | 7.19E-04 | 8.11E-03 | Dusp10/Il1a/Dnaj | 5 | aPD-L1+aTGF-β vs aTG | Up-regulated genes | CD45+ |
| BP | GO:0070372 | regulation of ERK1 and ERK2 cascade | 7.33E-04 | 8.21E-03 | Cd74/Ccl5/Dusp1 | 6 | aPD-L1+aTGF-β vs aTG | Up-regulated genes | CD45+ |
| BP | GO:0070232 | regulation of T cell apoptotic process | 7.81E-04 | 8.64E-03 | Ccl5/Cd274/Arg2 | 3 | aPD-L1+aTGF-β vs aTG | Up-regulated genes | CD45+ |
| BP | GO:1902622 | regulation of neutrophil migration | 7.81E-04 | 8.64E-03 | Cd74/Il1a/Ccr7 | 3 | aPD-L1+aTGF-β vs aTG | Up-regulated genes | CD45+ |
| BP | GO:0050792 | regulation of viral process | 8.31E-04 | 9.13E-03 | Vps37b/Cd74/Ccl | 5 | aPD-L1+aTGF-β vs aTG | Up-regulated genes | CD45+ |
| BP | GO:0044419 | interspecies interaction between organisms | 8.71E-04 | 9.29E-03 | Vps37b/Cd74/Ccl | 7 | aPD-L1+aTGF-β vs aTG | Up-regulated genes | CD45+ |
| BP | GO:1902106 | negative regulation of leukocyte differentiation | 8.79E-04 | 9.29E-03 | Irf1/Cd74/Socs1/C | 4 | aPD-L1+aTGF-β vs aTG | Up-regulated genes | CD45+ |
| BP | GO:0002822 | regulation of adaptive immune response based | 8.79E-04 | 9.29E-03 | Irf1/H2-Ab1/Cd27 | 5 | aPD-L1+aTGF-β vs aTG | Up-regulated genes | CD45+ |
| BP | GO:0070374 | positive regulation of ERK1 and ERK2 cascade | 8.79E-04 | 9.29E-03 | Cd74/Ccl5/Il1a/Ic | 5 | aPD-L1+aTGF-β vs aTG | Up-regulated genes | CD45+ |
| BP | GO:0071222 | cellular response to lipopolysaccharide | 8.79E-04 | 9.29E-03 | Cxcl10/Ccl5/Cd27 | 5 | aPD-L1+aTGF-β vs aTG | Up-regulated genes | CD45+ |
| BP | GO:2001242 | regulation of intrinsic apoptotic signaling pathwa | 8.79E-04 | 9.29E-03 | Cd74/Dnaja1/Plau | 5 | aPD-L1+aTGF-β vs aTG | Up-regulated genes | CD45+ |
| BP | GO:0007254 | JNK cascade | 9.30E-04 | 9.66E-03 | Dusp10/Il1a/Dnaj | 5 | aPD-L1+aTGF-β vs aTG | Up-regulated genes | CD45+ |
| BP | GO:0002468 | dendritic cell antigen processing and presentatio | 9.38E-04 | 9.66E-03 | Cd74/Ccr7 | 2 | aPD-L1+aTGF-β vs aTG | Up-regulated genes | CD45+ |
| BP | GO:0042090 | interleukin-12 biosynthetic process | 9.38E-04 | 9.66E-03 | Irf1/Rel | 2 | aPD-L1+aTGF-β vs aTG | Up-regulated genes | CD45+ |
| BP | GO:0045075 | regulation of interleukin-12 biosynthetic process | 9.38E-04 | 9.66E-03 | Irf1/Rel | 2 | aPD-L1+aTGF-β vs aTG | Up-regulated genes | CD45+ |
| BP | GO:0070371 | ERK1 and ERK2 cascade | 9.72E-04 | 9.95E-03 | Cd74/Ccl5/Dusp1 | 6 | aPD-L1+aTGF-β vs aTG | Up-regulated genes | CD45+ |
| BP | GO:2000404 | regulation of T cell migration | 1.04E-03 | 1.06E-02 | Cxcl10/Ccl5/Ccr7 | 3 | aPD-L1+aTGF-β vs aTG | Up-regulated genes | CD45+ |
| BP | GO:0043405 | regulation of MAP kinase activity | 1.06E-03 | 1.07E-02 | Cd74/Dusp10/Du | 6 | aPD-L1+aTGF-β vs aTG | Up-regulated genes | CD45+ |
| BP | GO:0071219 | cellular response to molecule of bacterial origin | 1.07E-03 | 1.07E-02 | Cxcl10/Ccl5/Cd27 | 5 | aPD-L1+aTGF-β vs aTG | Up-regulated genes | CD45+ |
| BP | GO:0016032 | viral process | 1.08E-03 | 1.08E-02 | Vps37b/Cd74/Ccl | 6 | aPD-L1+aTGF-β vs aTG | Up-regulated genes | CD45+ |
| BP | GO:0048247 | lymphocyte chemotaxis | 1.11E-03 | 1.10E-02 | Cxcl10/Ccl5/Ccr7 | 3 | aPD-L1+aTGF-β vs aTG | Up-regulated genes | CD45+ |
| BP | GO:0071675 | regulation of mononuclear cell migration | 1.11E-03 | 1.10E-02 | Cxcl10/Ccl5/Ccr7 | 3 | aPD-L1+aTGF-β vs aTG | Up-regulated genes | CD45+ |
| BP | GO:0001660 | fever generation | 1.14E-03 | 1.10E-02 | Ccl5/Il1a | 2 | aPD-L1+aTGF-β vs aTG | Up-regulated genes | CD45+ |
| BP | GO:0045625 | regulation of T-helper 1 cell differentiation | 1.14E-03 | 1.10E-02 | Irf1/Ccr7 | 2 | aPD-L1+aTGF-β vs aTG | Up-regulated genes | CD45+ |
| BP | GO:0072683 | T cell extravasation | 1.14E-03 | 1.10E-02 | Ccl5/Icam1 | 2 | aPD-L1+aTGF-β vs aTG | Up-regulated genes | CD45+ |
| BP | GO:1902166 | negative regulation of intrinsic apoptotic signalin | 1.14E-03 | 1.10E-02 | Cd74/Cd44 | 2 | aPD-L1+aTGF-β vs aTG | Up-regulated genes | CD45+ |
| BP | GO:2000343 | positive regulation of chemokine (C-X-C motif) li | 1.14E-03 | 1.10E-02 | Cd74/Ccl5 | 2 | aPD-L1+aTGF-β vs aTG | Up-regulated genes | CD45+ |
| BP | GO:0046425 | regulation of JAK-STAT cascade | 1.16E-03 | 1.10E-02 | Irf1/Ccl5/Socs1/Sc | 4 | aPD-L1+aTGF-β vs aTG | Up-regulated genes | CD45+ |
| BP | GO:0002688 | regulation of leukocyte chemotaxis | 1.20E-03 | 1.14E-02 | Cd74/Cxcl10/Ccl5 | 4 | aPD-L1+aTGF-β vs aTG | Up-regulated genes | CD45+ |
| BP | GO:0045088 | regulation of innate immune response | 1.27E-03 | 1.18E-02 | Irf1/Cd74/Dusp10 | 6 | aPD-L1+aTGF-β vs aTG | Up-regulated genes | CD45+ |
| BP | GO:0042088 | T-helper 1 type immune response | 1.27E-03 | 1.18E-02 | Irf1/H2-Ab1/Ccr7 | 3 | aPD-L1+aTGF-β vs aTG | Up-regulated genes | CD45+ |
| BP | GO:0071622 | regulation of granulocyte chemotaxis | 1.27E-03 | 1.18E-02 | Cd74/Ccl5/Ccr7 | 3 | aPD-L1+aTGF-β vs aTG | Up-regulated genes | CD45+ |
| BP | GO:0042254 | ribosome biogenesis | 1.32E-03 | 1.22E-02 | Rps21/Rps27/Rpl | 6 | aPD-L1+aTGF-β vs aTG | Up-regulated genes | CD45+ |
| BP | GO:1904892 | regulation of STAT cascade | 1.34E-03 | 1.23E-02 | Irf1/Ccl5/Socs1/Sc | 4 | aPD-L1+aTGF-β vs aTG | Up-regulated genes | CD45+ |
| BP | GO:0002903 | negative regulation of B cell apoptotic process | 1.37E-03 | 1.23E-02 | Cd74/Cd44 | 2 | aPD-L1+aTGF-β vs aTG | Up-regulated genes | CD45+ |
| BP | GO:0010820 | positive regulation of T cell chemotaxis | 1.37E-03 | 1.23E-02 | Ccl5/Ccr7 | 2 | aPD-L1+aTGF-β vs aTG | Up-regulated genes | CD45+ |
| BP | GO:0030335 | positive regulation of cell migration | 1.37E-03 | 1.23E-02 | Cd74/Cxcl10/Ccl5 | 8 | aPD-L1+aTGF-β vs aTG | Up-regulated genes | CD45+ |
| BP | GO:0001933 | negative regulation of protein phosphorylation | 1.37E-03 | 1.23E-02 | Irf1/Dusp10/Dusp | 7 | aPD-L1+aTGF-β vs aTG | Up-regulated genes | CD45+ |
| BP | GO:0007249 | I-kappaB kinase/NF-kappaB signaling | 1.38E-03 | 1.24E-02 | Irf1/Cd74/Rel/Il1a | 5 | aPD-L1+aTGF-β vs aTG | Up-regulated genes | CD45+ |
| BP | GO:0097193 | intrinsic apoptotic signaling pathway | 1.45E-03 | 1.29E-02 | Cd74/Crip1/Dnaja | 6 | aPD-L1+aTGF-β vs aTG | Up-regulated genes | CD45+ |
| BP | GO:0097529 | myeloid leukocyte migration | 1.49E-03 | 1.32E-02 | Cd74/Cxcl10/Ccl5 | 5 | aPD-L1+aTGF-β vs aTG | Up-regulated genes | CD45+ |
| BP | GO:0071900 | regulation of protein serine/threonine kinase acti | 1.53E-03 | 1.35E-02 | Cd74/Dusp10/Pin | 7 | aPD-L1+aTGF-β vs aTG | Up-regulated genes | CD45+ |
| BP | GO:0071621 | granulocyte chemotaxis | 1.54E-03 | 1.35E-02 | Cd74/Cxcl10/Ccl5 | 4 | aPD-L1+aTGF-β vs aTG | Up-regulated genes | CD45+ |
| BP | GO:2000147 | positive regulation of cell motility | 1.57E-03 | 1.37E-02 | Cd74/Cxcl10/Ccl5 | 8 | aPD-L1+aTGF-β vs aTG | Up-regulated genes | CD45+ |
| BP | GO:0044406 | adhesion of symbiont to host | 1.61E-03 | 1.38E-02 | Icam1/Gbp2b | 2 | aPD-L1+aTGF-β vs aTG | Up-regulated genes | CD45+ |
| BP | GO:0045060 | negative thymic T cell selection | 1.61E-03 | 1.38E-02 | Cd74/Ccr7 | 2 | aPD-L1+aTGF-β vs aTG | Up-regulated genes | CD45+ |
| BP | GO:0090594 | inflammatory response to wounding | 1.61E-03 | 1.38E-02 | Il1a/Cd44 | 2 | aPD-L1+aTGF-β vs aTG | Up-regulated genes | CD45+ |
| BP | GO:0050732 | negative regulation of peptidyl-tyrosine phospho | 1.62E-03 | 1.38E-02 | Irf1/Socs1/Socs3 | 3 | aPD-L1+aTGF-β vs aTG | Up-regulated genes | CD45+ |
| BP | GO:0050707 | regulation of cytokine secretion | 1.68E-03 | 1.42E-02 | Cd274/Il1a/Arg2/S | 5 | aPD-L1+aTGF-β vs aTG | Up-regulated genes | CD45+ |
| BP | GO:0045071 | negative regulation of viral genome replication | 1.71E-03 | 1.45E-02 | Ccl5/Oasl1/Rsad2 | 3 | aPD-L1+aTGF-β vs aTG | Up-regulated genes | CD45+ |
| BP | GO:0071396 | cellular response to lipid | 1.73E-03 | 1.45E-02 | Cxcl10/Ccl5/Cd27 | 7 | aPD-L1+aTGF-β vs aTG | Up-regulated genes | CD45+ |
| BP | GO:0043903 | regulation of symbiosis, encompassing mutualis | 1.80E-03 | 1.50E-02 | Vps37b/Cd74/Ccl | 5 | aPD-L1+aTGF-β vs aTG | Up-regulated genes | CD45+ |
| BP | GO:0071216 | cellular response to biotic stimulus | 1.80E-03 | 1.50E-02 | Cxcl10/Ccl5/Cd27 | 5 | aPD-L1+aTGF-β vs aTG | Up-regulated genes | CD45+ |
| BP | GO:0032655 | regulation of interleukin-12 production | 1.81E-03 | 1.50E-02 | Irf1/Rel/Ccr7 | 3 | aPD-L1+aTGF-β vs aTG | Up-regulated genes | CD45+ |
| BP | GO:0010951 | negative regulation of endopeptidase activity | 1.82E-03 | 1.50E-02 | Serpina3g/Plaur/I | 4 | aPD-L1+aTGF-β vs aTG | Up-regulated genes | CD45+ |
| BP | GO:0051272 | positive regulation of cellular component mover | 1.86E-03 | 1.50E-02 | Cd74/Cxcl10/Ccl5 | 8 | aPD-L1+aTGF-β vs aTG | Up-regulated genes | CD45+ |
| BP | GO:0002829 | negative regulation of type 2 immune response | 1.87E-03 | 1.50E-02 | Irf1/Arg2 | 2 | aPD-L1+aTGF-β vs aTG | Up-regulated genes | CD45+ |
| BP | GO:0043383 | negative T cell selection | 1.87E-03 | 1.50E-02 | Cd74/Ccr7 | 2 | aPD-L1+aTGF-β vs aTG | Up-regulated genes | CD45+ |

| | | | | | | | | | |
|---|---|---|---|---|---|---|---|---|---|
| BP | GO:0045591 | positive regulation of regulatory T cell differentia | 1.87E-03 | 1.50E-02 | Dusp10/Socs1 | 2 | aPD-L1+aTGF-β vs aTG | Up-regulated genes | CD45+ |
| BP | GO:0070234 | positive regulation of T cell apoptotic process | 1.87E-03 | 1.50E-02 | Ccl5/Cd274 | 2 | aPD-L1+aTGF-β vs aTG | Up-regulated genes | CD45+ |
| BP | GO:2000341 | regulation of chemokine (C-X-C motif) ligand 2 p | 1.87E-03 | 1.50E-02 | Cd74/Ccl5 | 2 | aPD-L1+aTGF-β vs aTG | Up-regulated genes | CD45+ |
| BP | GO:0006959 | humoral immune response | 1.88E-03 | 1.50E-02 | H2-Ab1/Cxcl10/R | 4 | aPD-L1+aTGF-β vs aTG | Up-regulated genes | CD45+ |
| BP | GO:0032872 | regulation of stress-activated MAPK cascade | 1.93E-03 | 1.53E-02 | Dusp10/Il1a/Dnaj | 5 | aPD-L1+aTGF-β vs aTG | Up-regulated genes | CD45+ |
| BP | GO:0032615 | interleukin-12 production | 2.02E-03 | 1.58E-02 | Irf1/Rel/Ccr7 | 3 | aPD-L1+aTGF-β vs aTG | Up-regulated genes | CD45+ |
| BP | GO:0070231 | T cell apoptotic process | 2.02E-03 | 1.58E-02 | Ccl5/Cd274/Arg2 | 3 | aPD-L1+aTGF-β vs aTG | Up-regulated genes | CD45+ |
| BP | GO:0070302 | regulation of stress-activated protein kinase sign | 2.02E-03 | 1.58E-02 | Dusp10/Il1a/Dnaj | 5 | aPD-L1+aTGF-β vs aTG | Up-regulated genes | CD45+ |
| BP | GO:0050708 | regulation of protein secretion | 2.10E-03 | 1.63E-02 | Ccl5/Cd274/Rsad | 7 | aPD-L1+aTGF-β vs aTG | Up-regulated genes | CD45+ |
| BP | GO:2000401 | regulation of lymphocyte migration | 2.13E-03 | 1.63E-02 | Cxcl10/Ccl5/Ccr7 | 3 | aPD-L1+aTGF-β vs aTG | Up-regulated genes | CD45+ |
| BP | GO:0007259 | JAK-STAT cascade | 2.14E-03 | 1.63E-02 | Irf1/Ccl5/Socs1/S | 4 | aPD-L1+aTGF-β vs aTG | Up-regulated genes | CD45+ |
| BP | GO:0043518 | negative regulation of DNA damage response, s | 2.16E-03 | 1.63E-02 | Cd74/Cd44 | 2 | aPD-L1+aTGF-β vs aTG | Up-regulated genes | CD45+ |
| BP | GO:0072567 | chemokine (C-X-C motif) ligand 2 production | 2.16E-03 | 1.63E-02 | Cd74/Ccl5 | 2 | aPD-L1+aTGF-β vs aTG | Up-regulated genes | CD45+ |
| BP | GO:0140131 | positive regulation of lymphocyte chemotaxis | 2.16E-03 | 1.63E-02 | Ccl5/Ccr7 | 2 | aPD-L1+aTGF-β vs aTG | Up-regulated genes | CD45+ |
| BP | GO:1902165 | regulation of intrinsic apoptotic signaling pathwa | 2.16E-03 | 1.63E-02 | Cd74/Cd44 | 2 | aPD-L1+aTGF-β vs aTG | Up-regulated genes | CD45+ |
| BP | GO:0050921 | positive regulation of chemotaxis | 2.27E-03 | 1.71E-02 | Cd74/Cxcl10/Ccl | 4 | aPD-L1+aTGF-β vs aTG | Up-regulated genes | CD45+ |
| BP | GO:0050671 | positive regulation of lymphocyte proliferation | 2.34E-03 | 1.73E-02 | Cd74/Ccl5/Cd274 | 4 | aPD-L1+aTGF-β vs aTG | Up-regulated genes | CD45+ |
| BP | GO:0097696 | STAT cascade | 2.34E-03 | 1.73E-02 | Irf1/Ccl5/Socs1/S | 4 | aPD-L1+aTGF-β vs aTG | Up-regulated genes | CD45+ |
| BP | GO:0043410 | positive regulation of MAPK cascade | 2.35E-03 | 1.73E-02 | Cd74/Ccl5/Il1a/Ic | 7 | aPD-L1+aTGF-β vs aTG | Up-regulated genes | CD45+ |
| BP | GO:0032722 | positive regulation of chemokine production | 2.36E-03 | 1.73E-02 | Cd74/Ccl5/Il1a | 3 | aPD-L1+aTGF-β vs aTG | Up-regulated genes | CD45+ |
| BP | GO:0045123 | cellular extravasation | 2.36E-03 | 1.73E-02 | Ccl5/Il1a/Icam1 | 3 | aPD-L1+aTGF-β vs aTG | Up-regulated genes | CD45+ |
| BP | GO:0042326 | negative regulation of phosphorylation | 2.39E-03 | 1.75E-02 | Irf1/Dusp10/Dusp | 7 | aPD-L1+aTGF-β vs aTG | Up-regulated genes | CD45+ |
| BP | GO:0002830 | positive regulation of type 2 immune response | 2.46E-03 | 1.78E-02 | Cd74/Rsad2 | 2 | aPD-L1+aTGF-β vs aTG | Up-regulated genes | CD45+ |
| BP | GO:0031649 | heat generation | 2.46E-03 | 1.78E-02 | Ccl5/Il1a | 2 | aPD-L1+aTGF-β vs aTG | Up-regulated genes | CD45+ |
| BP | GO:0060009 | Sertoli cell development | 2.46E-03 | 1.78E-02 | Il1a/Icam1 | 2 | aPD-L1+aTGF-β vs aTG | Up-regulated genes | CD45+ |
| BP | GO:0050727 | regulation of inflammatory response | 2.49E-03 | 1.78E-02 | Ccl5/Dusp10/Soc | 6 | aPD-L1+aTGF-β vs aTG | Up-regulated genes | CD45+ |
| BP | GO:0032946 | positive regulation of mononuclear cell proliferat | 2.49E-03 | 1.78E-02 | Cd74/Ccl5/Cd274 | 4 | aPD-L1+aTGF-β vs aTG | Up-regulated genes | CD45+ |
| BP | GO:0050900 | leukocyte migration | 2.53E-03 | 1.81E-02 | Cd74/Cxcl10/Ccl | 6 | aPD-L1+aTGF-β vs aTG | Up-regulated genes | CD45+ |
| BP | GO:0042255 | ribosome assembly | 2.60E-03 | 1.85E-02 | Rps27/Rpl38/Rps | 3 | aPD-L1+aTGF-β vs aTG | Up-regulated genes | CD45+ |
| BP | GO:0071356 | cellular response to tumor necrosis factor | 2.64E-03 | 1.87E-02 | Irf1/Ccl5/Nfkbia/N | 4 | aPD-L1+aTGF-β vs aTG | Up-regulated genes | CD45+ |
| BP | GO:0000188 | inactivation of MAPK activity | 2.78E-03 | 1.95E-02 | Dusp10/Dusp2 | 2 | aPD-L1+aTGF-β vs aTG | Up-regulated genes | CD45+ |
| BP | GO:0090026 | positive regulation of monocyte chemotaxis | 2.78E-03 | 1.95E-02 | Cxcl10/Ccl5 | 2 | aPD-L1+aTGF-β vs aTG | Up-regulated genes | CD45+ |
| BP | GO:0051403 | stress-activated MAPK cascade | 2.85E-03 | 1.99E-02 | Dusp10/Il1a/Dnaj | 5 | aPD-L1+aTGF-β vs aTG | Up-regulated genes | CD45+ |
| BP | GO:0043407 | negative regulation of MAP kinase activity | 2.86E-03 | 1.99E-02 | Dusp10/Dusp2/D | 3 | aPD-L1+aTGF-β vs aTG | Up-regulated genes | CD45+ |
| BP | GO:0070665 | positive regulation of leukocyte proliferation | 2.88E-03 | 1.99E-02 | Cd74/Ccl5/Cd274 | 4 | aPD-L1+aTGF-β vs aTG | Up-regulated genes | CD45+ |
| BP | GO:0050663 | cytokine secretion | 3.03E-03 | 2.09E-02 | Cd274/Il1a/Arg2/ | 5 | aPD-L1+aTGF-β vs aTG | Up-regulated genes | CD45+ |
| BP | GO:0046688 | response to copper ion | 3.11E-03 | 2.13E-02 | Il1a/Nfe2l2 | 2 | aPD-L1+aTGF-β vs aTG | Up-regulated genes | CD45+ |
| BP | GO:0060008 | Sertoli cell differentiation | 3.11E-03 | 2.13E-02 | Il1a/Icam1 | 2 | aPD-L1+aTGF-β vs aTG | Up-regulated genes | CD45+ |
| BP | GO:0042108 | positive regulation of cytokine biosynthetic proce | 3.14E-03 | 2.13E-02 | Irf1/Rel/Il1a | 3 | aPD-L1+aTGF-β vs aTG | Up-regulated genes | CD45+ |
| BP | GO:0002438 | acute inflammatory response to antigenic stimul | 3.47E-03 | 2.33E-02 | Icam1/Ccr7 | 2 | aPD-L1+aTGF-β vs aTG | Up-regulated genes | CD45+ |
| BP | GO:0002643 | regulation of tolerance induction | 3.47E-03 | 2.33E-02 | Cd274/Ccr7 | 2 | aPD-L1+aTGF-β vs aTG | Up-regulated genes | CD45+ |
| BP | GO:0002902 | regulation of B cell apoptotic process | 3.47E-03 | 2.33E-02 | Cd74/Cd44 | 2 | aPD-L1+aTGF-β vs aTG | Up-regulated genes | CD45+ |
| BP | GO:0010759 | positive regulation of macrophage chemotaxis | 3.47E-03 | 2.33E-02 | Ccl5/Ccr7 | 2 | aPD-L1+aTGF-β vs aTG | Up-regulated genes | CD45+ |
| BP | GO:0001659 | temperature homeostasis | 3.78E-03 | 2.51E-02 | Ccl5/Kdm6b/Il1a/I | 4 | aPD-L1+aTGF-β vs aTG | Up-regulated genes | CD45+ |
| BP | GO:0045063 | T-helper 1 cell differentiation | 3.84E-03 | 2.51E-02 | Irf1/Ccr7 | 2 | aPD-L1+aTGF-β vs aTG | Up-regulated genes | CD45+ |
| BP | GO:0045624 | positive regulation of T-helper cell differentiation | 3.84E-03 | 2.51E-02 | Irf1/Ccr7 | 2 | aPD-L1+aTGF-β vs aTG | Up-regulated genes | CD45+ |
| BP | GO:0060333 | interferon-gamma-mediated signaling pathway | 3.84E-03 | 2.51E-02 | Irf1/Socs1 | 2 | aPD-L1+aTGF-β vs aTG | Up-regulated genes | CD45+ |
| BP | GO:0070230 | positive regulation of lymphocyte apoptotic proc | 3.84E-03 | 2.51E-02 | Ccl5/Cd274 | 2 | aPD-L1+aTGF-β vs aTG | Up-regulated genes | CD45+ |
| BP | GO:1902254 | negative regulation of intrinsic apoptotic signalin | 3.84E-03 | 2.51E-02 | Cd74/Cd44 | 2 | aPD-L1+aTGF-β vs aTG | Up-regulated genes | CD45+ |
| BP | GO:0010466 | negative regulation of peptidase activity | 3.88E-03 | 2.53E-02 | Serpina3g/Plaur/ | 4 | aPD-L1+aTGF-β vs aTG | Up-regulated genes | CD45+ |
| BP | GO:0071347 | cellular response to interleukin-1 | 3.89E-03 | 2.53E-02 | Irf1/Ccl5/Il1a | 3 | aPD-L1+aTGF-β vs aTG | Up-regulated genes | CD45+ |
| BP | GO:0031349 | positive regulation of defense response | 3.95E-03 | 2.55E-02 | Irf1/Cd74/Ccl5/Rs | 6 | aPD-L1+aTGF-β vs aTG | Up-regulated genes | CD45+ |
| BP | GO:0043154 | negative regulation of cysteine-type endopeptid | 4.06E-03 | 2.60E-02 | Plaur/Nr4a1/Cd4 | 3 | aPD-L1+aTGF-β vs aTG | Up-regulated genes | CD45+ |
| BP | GO:0071674 | mononuclear cell migration | 4.06E-03 | 2.60E-02 | Cxcl10/Ccl5/Ccr7 | 3 | aPD-L1+aTGF-β vs aTG | Up-regulated genes | CD45+ |
| BP | GO:0034612 | response to tumor necrosis factor | 4.19E-03 | 2.68E-02 | Irf1/Ccl5/Nfkbia/N | 4 | aPD-L1+aTGF-β vs aTG | Up-regulated genes | CD45+ |
| BP | GO:0002693 | positive regulation of cellular extravasation | 4.24E-03 | 2.70E-02 | Il1a/Icam1 | 2 | aPD-L1+aTGF-β vs aTG | Up-regulated genes | CD45+ |
| BP | GO:1903901 | negative regulation of viral life cycle | 4.39E-03 | 2.79E-02 | Ccl5/Oasl1/Rsad | 3 | aPD-L1+aTGF-β vs aTG | Up-regulated genes | CD45+ |
| BP | GO:1901653 | cellular response to peptide | 4.62E-03 | 2.91E-02 | Irf1/Socs1/Icam1/ | 5 | aPD-L1+aTGF-β vs aTG | Up-regulated genes | CD45+ |
| BP | GO:0071677 | positive regulation of mononuclear cell migratio | 4.65E-03 | 2.91E-02 | Cxcl10/Ccl5 | 2 | aPD-L1+aTGF-β vs aTG | Up-regulated genes | CD45+ |
| BP | GO:0090025 | regulation of monocyte chemotaxis | 4.65E-03 | 2.91E-02 | Cxcl10/Ccl5 | 2 | aPD-L1+aTGF-β vs aTG | Up-regulated genes | CD45+ |
| BP | GO:0043900 | regulation of multi-organism process | 4.85E-03 | 3.02E-02 | Vps37b/Cd74/Ccl | 6 | aPD-L1+aTGF-β vs aTG | Up-regulated genes | CD45+ |
| BP | GO:0031098 | stress-activated protein kinase signaling cascad | 4.96E-03 | 3.08E-02 | Dusp10/Il1a/Dnaj | 5 | aPD-L1+aTGF-β vs aTG | Up-regulated genes | CD45+ |
| BP | GO:0002675 | positive regulation of acute inflammatory respon | 5.07E-03 | 3.12E-02 | Ccl5/Ccr7 | 2 | aPD-L1+aTGF-β vs aTG | Up-regulated genes | CD45+ |
| BP | GO:0070233 | negative regulation of T cell apoptotic process | 5.07E-03 | 3.12E-02 | Ccl5/Arg2 | 2 | aPD-L1+aTGF-β vs aTG | Up-regulated genes | CD45+ |
| BP | GO:0090023 | positive regulation of neutrophil chemotaxis | 5.07E-03 | 3.12E-02 | Cd74/Ccr7 | 2 | aPD-L1+aTGF-β vs aTG | Up-regulated genes | CD45+ |
| BP | GO:2000117 | negative regulation of cysteine-type endopeptid | 5.12E-03 | 3.14E-02 | Plaur/Nr4a1/Cd4 | 3 | aPD-L1+aTGF-β vs aTG | Up-regulated genes | CD45+ |
| BP | GO:0043901 | negative regulation of multi-organism process | 5.21E-03 | 3.18E-02 | Ccl5/Oasl1/Rsad | 4 | aPD-L1+aTGF-β vs aTG | Up-regulated genes | CD45+ |
| BP | GO:0014070 | response to organic cyclic compound | 5.25E-03 | 3.19E-02 | Ccl5/Pim1/Dnaja | 7 | aPD-L1+aTGF-β vs aTG | Up-regulated genes | CD45+ |
| BP | GO:0043506 | regulation of JUN kinase activity | 5.32E-03 | 3.22E-02 | Dusp10/Dnaja1/C | 3 | aPD-L1+aTGF-β vs aTG | Up-regulated genes | CD45+ |
| BP | GO:0010035 | response to inorganic substance | 5.50E-03 | 3.26E-02 | Kdm6b/Crip1/Il1a | 6 | aPD-L1+aTGF-β vs aTG | Up-regulated genes | CD45+ |
| BP | GO:0060326 | cell chemotaxis | 5.50E-03 | 3.26E-02 | Cd74/Cxcl10/Ccl | 5 | aPD-L1+aTGF-β vs aTG | Up-regulated genes | CD45+ |
| BP | GO:0006925 | inflammatory cell apoptotic process | 5.52E-03 | 3.26E-02 | Ccl5/Cd44 | 2 | aPD-L1+aTGF-β vs aTG | Up-regulated genes | CD45+ |
| BP | GO:0045061 | thymic T cell selection | 5.52E-03 | 3.26E-02 | Cd74/Ccr7 | 2 | aPD-L1+aTGF-β vs aTG | Up-regulated genes | CD45+ |
| BP | GO:1902230 | negative regulation of intrinsic apoptotic signalin | 5.52E-03 | 3.26E-02 | Cd74/Cd44 | 2 | aPD-L1+aTGF-β vs aTG | Up-regulated genes | CD45+ |
| BP | GO:1905523 | positive regulation of macrophage migration | 5.52E-03 | 3.26E-02 | Ccl5/Ccr7 | 2 | aPD-L1+aTGF-β vs aTG | Up-regulated genes | CD45+ |
| BP | GO:2000352 | negative regulation of endothelial cell apoptotic | 5.52E-03 | 3.26E-02 | Icam1/Nfe2l2 | 2 | aPD-L1+aTGF-β vs aTG | Up-regulated genes | CD45+ |
| BP | GO:0043434 | response to peptide hormone | 5.69E-03 | 3.35E-02 | Irf1/Serpina3g/So | 5 | aPD-L1+aTGF-β vs aTG | Up-regulated genes | CD45+ |
| BP | GO:0002697 | regulation of immune effector process | 5.96E-03 | 3.47E-02 | Irf1/Cd74/Ccl5/Du | 6 | aPD-L1+aTGF-β vs aTG | Up-regulated genes | CD45+ |
| BP | GO:0006953 | acute-phase response | 5.98E-03 | 3.47E-02 | Ccl5/Il1a | 2 | aPD-L1+aTGF-β vs aTG | Up-regulated genes | CD45+ |
| BP | GO:0046639 | negative regulation of alpha-beta T cell differenti | 5.98E-03 | 3.47E-02 | Irf1/Socs1 | 2 | aPD-L1+aTGF-β vs aTG | Up-regulated genes | CD45+ |
| BP | GO:0071624 | positive regulation of granulocyte chemotaxis | 5.98E-03 | 3.47E-02 | Cd74/Ccr7 | 2 | aPD-L1+aTGF-β vs aTG | Up-regulated genes | CD45+ |
| BP | GO:0010563 | negative regulation of phosphorus metabolic pro | 6.10E-03 | 3.51E-02 | Irf1/Dusp10/Dusp | 7 | aPD-L1+aTGF-β vs aTG | Up-regulated genes | CD45+ |
| BP | GO:0045936 | negative regulation of phosphate metabolic proc | 6.10E-03 | 3.51E-02 | Irf1/Dusp10/Dusp | 7 | aPD-L1+aTGF-β vs aTG | Up-regulated genes | CD45+ |
| BP | GO:0050731 | positive regulation of peptidyl-tyrosine phosphor | 6.12E-03 | 3.51E-02 | Cd74/Ccl5/Icam1/ | 4 | aPD-L1+aTGF-β vs aTG | Up-regulated genes | CD45+ |
| BP | GO:0042742 | defense response to bacterium | 6.26E-03 | 3.58E-02 | Rpl39/Arg2/Plac8 | 4 | aPD-L1+aTGF-β vs aTG | Up-regulated genes | CD45+ |
| BP | GO:0045069 | regulation of viral genome replication | 6.35E-03 | 3.62E-02 | Ccl5/Oasl1/Rsad | 3 | aPD-L1+aTGF-β vs aTG | Up-regulated genes | CD45+ |
| BP | GO:0001783 | B cell apoptotic process | 6.46E-03 | 3.67E-02 | Cd74/Cd44 | 2 | aPD-L1+aTGF-β vs aTG | Up-regulated genes | CD45+ |
| BP | GO:0009306 | protein secretion | 6.67E-03 | 3.78E-02 | Ccl5/Cd274/Rsad | 7 | aPD-L1+aTGF-β vs aTG | Up-regulated genes | CD45+ |
| BP | GO:0031295 | T cell costimulation | 6.96E-03 | 3.91E-02 | Cd274/Ccr7 | 2 | aPD-L1+aTGF-β vs aTG | Up-regulated genes | CD45+ |

| | | | | | | | | | |
|---|---|---|---|---|---|---|---|---|---|
| BP | GO:0042119 | neutrophil activation | 6.96E-03 | 3.91E-02 | Ccl5/Fcgr4 | 2 | aPD-L1+aTGF-β vs aTG | Up-regulated genes | CD45+ |
| BP | GO:0048525 | negative regulation of viral process | 7.02E-03 | 3.94E-02 | Ccl5/Oasl1/Rsad2 | 3 | aPD-L1+aTGF-β vs aTG | Up-regulated genes | CD45+ |
| BP | GO:0032103 | positive regulation of response to external stimul | 7.16E-03 | 4.00E-02 | Cd74/Cxcl10/Ccl5 | 5 | aPD-L1+aTGF-β vs aTG | Up-regulated genes | CD45+ |
| BP | GO:0070555 | response to interleukin-1 | 7.25E-03 | 4.04E-02 | Irf1/Ccl5/Il1a | 3 | aPD-L1+aTGF-β vs aTG | Up-regulated genes | CD45+ |
| BP | GO:2001233 | regulation of apoptotic signaling pathway | 7.36E-03 | 4.08E-02 | Cd74/Dnaja1/Pla | 6 | aPD-L1+aTGF-β vs aTG | Up-regulated genes | CD45+ |
| BP | GO:0002507 | tolerance induction | 7.47E-03 | 4.10E-02 | Cd274/Ccr7 | 2 | aPD-L1+aTGF-β vs aTG | Up-regulated genes | CD45+ |
| BP | GO:0010758 | regulation of macrophage chemotaxis | 7.47E-03 | 4.10E-02 | Ccl5/Ccr7 | 2 | aPD-L1+aTGF-β vs aTG | Up-regulated genes | CD45+ |
| BP | GO:0046329 | negative regulation of JNK cascade | 7.47E-03 | 4.10E-02 | Dusp10/Dnaja1 | 2 | aPD-L1+aTGF-β vs aTG | Up-regulated genes | CD45+ |
| BP | GO:0002286 | T cell activation involved in immune response | 7.73E-03 | 4.23E-02 | Irf1/Icam1/Ccr7 | 3 | aPD-L1+aTGF-β vs aTG | Up-regulated genes | CD45+ |
| BP | GO:0002691 | regulation of cellular extravasation | 8.00E-03 | 4.31E-02 | Il1a/Icam1 | 2 | aPD-L1+aTGF-β vs aTG | Up-regulated genes | CD45+ |
| BP | GO:0031294 | lymphocyte costimulation | 8.00E-03 | 4.31E-02 | Cd274/Ccr7 | 2 | aPD-L1+aTGF-β vs aTG | Up-regulated genes | CD45+ |
| BP | GO:0033032 | regulation of myeloid cell apoptotic process | 8.00E-03 | 4.31E-02 | Ccl5/Cd44 | 2 | aPD-L1+aTGF-β vs aTG | Up-regulated genes | CD45+ |
| BP | GO:0090022 | regulation of neutrophil chemotaxis | 8.00E-03 | 4.31E-02 | Cd74/Ccr7 | 2 | aPD-L1+aTGF-β vs aTG | Up-regulated genes | CD45+ |
| BP | GO:1902253 | regulation of intrinsic apoptotic signaling pathwa | 8.00E-03 | 4.31E-02 | Cd74/Cd44 | 2 | aPD-L1+aTGF-β vs aTG | Up-regulated genes | CD45+ |
| BP | GO:0042102 | positive regulation of T cell proliferation | 8.23E-03 | 4.42E-02 | Ccl5/Cd274/Ccr7 | 3 | aPD-L1+aTGF-β vs aTG | Up-regulated genes | CD45+ |
| BP | GO:0051930 | regulation of sensory perception of pain | 8.54E-03 | 4.56E-02 | Il1a/Fcgr4 | 2 | aPD-L1+aTGF-β vs aTG | Up-regulated genes | CD45+ |
| BP | GO:1901797 | negative regulation of signal transduction by p5 | 8.54E-03 | 4.56E-02 | Cd74/Cd44 | 2 | aPD-L1+aTGF-β vs aTG | Up-regulated genes | CD45+ |
| BP | GO:0050920 | regulation of chemotaxis | 8.91E-03 | 4.74E-02 | Cd74/Cxcl10/Ccl5 | 4 | aPD-L1+aTGF-β vs aTG | Up-regulated genes | CD45+ |
| BP | GO:0032743 | positive regulation of interleukin-2 production | 9.11E-03 | 4.79E-02 | Il1a/Cd83 | 2 | aPD-L1+aTGF-β vs aTG | Up-regulated genes | CD45+ |
| BP | GO:0051085 | chaperone cofactor-dependent protein refolding | 9.11E-03 | 4.79E-02 | Cd74/H2-DMb1 | 2 | aPD-L1+aTGF-β vs aTG | Up-regulated genes | CD45+ |
| BP | GO:0051931 | regulation of sensory perception | 9.11E-03 | 4.79E-02 | Il1a/Fcgr4 | 2 | aPD-L1+aTGF-β vs aTG | Up-regulated genes | CD45+ |
| BP | GO:0006364 | rRNA processing | 9.27E-03 | 4.85E-02 | Rps21/Rpl27/Rpl | 4 | aPD-L1+aTGF-β vs aTG | Up-regulated genes | CD45+ |
| BP | GO:0006469 | negative regulation of protein kinase activity | 9.27E-03 | 4.85E-02 | Dusp10/Dusp2/D | 4 | aPD-L1+aTGF-β vs aTG | Up-regulated genes | CD45+ |
| CC | GO:0022626 | cytosolic ribosome | 3.51E-17 | 3.75E-15 | Rpl37a/Rpl39/Rp | 14 | aPD-L1+aTGF-β vs aTG | Up-regulated genes | CD45+ |
| CC | GO:0005840 | ribosome | 5.06E-14 | 2.71E-12 | Rpl37a/Rpl39/Rp | 15 | aPD-L1+aTGF-β vs aTG | Up-regulated genes | CD45+ |
| CC | GO:0044391 | ribosomal subunit | 1.18E-13 | 4.20E-12 | Rpl37a/Rpl39/Rp | 14 | aPD-L1+aTGF-β vs aTG | Up-regulated genes | CD45+ |
| CC | GO:0044445 | cytosolic part | 3.01E-12 | 8.06E-11 | Rpl37a/Rpl39/Rp | 14 | aPD-L1+aTGF-β vs aTG | Up-regulated genes | CD45+ |
| CC | GO:0022625 | cytosolic large ribosomal subunit | 5.83E-12 | 1.25E-10 | Rpl37a/Rpl39/Rp | 9 | aPD-L1+aTGF-β vs aTG | Up-regulated genes | CD45+ |
| CC | GO:0015934 | large ribosomal subunit | 3.92E-09 | 7.00E-08 | Rpl37a/Rpl39/Rp | 9 | aPD-L1+aTGF-β vs aTG | Up-regulated genes | CD45+ |
| CC | GO:0042611 | MHC protein complex | 3.86E-08 | 5.39E-07 | H2-Eb1/H2-Aa/H2 | 5 | aPD-L1+aTGF-β vs aTG | Up-regulated genes | CD45+ |
| CC | GO:0022627 | cytosolic small ribosomal subunit | 4.03E-08 | 5.39E-07 | Rps21/Rps27/Rp | 6 | aPD-L1+aTGF-β vs aTG | Up-regulated genes | CD45+ |
| CC | GO:0042788 | polysomal ribosome | 1.82E-07 | 2.16E-06 | Rpl39/Rps21/Rpl | 5 | aPD-L1+aTGF-β vs aTG | Up-regulated genes | CD45+ |
| CC | GO:0009897 | external side of plasma membrane | 2.11E-07 | 2.26E-06 | H2-Aa/H2-Ab1/Cc | 11 | aPD-L1+aTGF-β vs aTG | Up-regulated genes | CD45+ |
| CC | GO:0015935 | small ribosomal subunit | 1.07E-06 | 1.05E-05 | Rps21/Rps27/Rp | 6 | aPD-L1+aTGF-β vs aTG | Up-regulated genes | CD45+ |
| CC | GO:0098552 | side of membrane | 9.02E-06 | 8.04E-05 | H2-Aa/H2-Ab1/Cc | 11 | aPD-L1+aTGF-β vs aTG | Up-regulated genes | CD45+ |
| CC | GO:0005844 | polysome | 1.67E-05 | 1.37E-04 | Rpl39/Rps21/Rpl | 5 | aPD-L1+aTGF-β vs aTG | Up-regulated genes | CD45+ |
| CC | GO:0005942 | phosphatidylinositol 3-kinase complex | 6.15E-03 | 4.70E-02 | Socs1/Socs3 | 2 | aPD-L1+aTGF-β vs aTG | Up-regulated genes | CD45+ |
| CC | GO:0043230 | extracellular organelle | 6.84E-03 | 4.88E-02 | Cd274/Icam1/Gbp | 3 | aPD-L1+aTGF-β vs aTG | Up-regulated genes | CD45+ |
| MF | GO:0003735 | structural constituent of ribosome | 9.56E-14 | 1.56E-11 | Rpl37a/Rpl39/Rp | 13 | aPD-L1+aTGF-β vs aTG | Up-regulated genes | CD45+ |
| MF | GO:0005198 | structural molecule activity | 5.67E-08 | 4.62E-06 | Rpl37a/Rpl39/Rp | 13 | aPD-L1+aTGF-β vs aTG | Up-regulated genes | CD45+ |
| MF | GO:0017017 | MAP kinase tyrosine/serine/threonine phosphata | 1.34E-03 | 4.29E-02 | Dusp10/Dusp2 | 2 | aPD-L1+aTGF-β vs aTG | Up-regulated genes | CD45+ |
| MF | GO:0023023 | MHC protein complex binding | 1.34E-03 | 4.29E-02 | Cd74/H2-DMb1 | 2 | aPD-L1+aTGF-β vs aTG | Up-regulated genes | CD45+ |
| MF | GO:0033549 | MAP kinase phosphatase activity | 1.58E-03 | 4.29E-02 | Dusp10/Dusp2 | 2 | aPD-L1+aTGF-β vs aTG | Up-regulated genes | CD45+ |
| MF | GO:0046935 | 1-phosphatidylinositol-3-kinase regulator activity | 1.58E-03 | 4.29E-02 | Socs1/Socs3 | 2 | aPD-L1+aTGF-β vs aTG | Up-regulated genes | CD45+ |
| BP | GO:0009617 | response to bacterium | 1.95E-11 | 1.93E-08 | Cxcl10/Cxcl9/Cxc | 12 | aPD-L1+aTGF-β vs aTG | Up-regulated genes | CD45- |
| BP | GO:0019221 | cytokine-mediated signaling pathway | 6.61E-11 | 3.27E-08 | Cxcl10/Cxcl9/Cxc | 10 | aPD-L1+aTGF-β vs aTG | Up-regulated genes | CD45- |
| BP | GO:0071276 | cellular response to cadmium ion | 7.21E-08 | 2.38E-05 | Mt1/Mt2/Jun/Fos | 4 | aPD-L1+aTGF-β vs aTG | Up-regulated genes | CD45- |
| BP | GO:1903706 | regulation of hemopoiesis | 1.84E-07 | 4.28E-05 | Cxcl1/Irf1/Socs1/ | 8 | aPD-L1+aTGF-β vs aTG | Up-regulated genes | CD45- |
| BP | GO:0002237 | response to molecule of bacterial origin | 2.50E-07 | 4.28E-05 | Cxcl10/Cxcl9/Cxc | 7 | aPD-L1+aTGF-β vs aTG | Up-regulated genes | CD45- |
| BP | GO:0030593 | neutrophil chemotaxis | 2.60E-07 | 4.28E-05 | Cxcl10/Cxcl9/Cxc | 5 | aPD-L1+aTGF-β vs aTG | Up-regulated genes | CD45- |
| BP | GO:0046686 | response to cadmium ion | 3.49E-07 | 4.93E-05 | Mt1/Mt2/Jun/Fos | 4 | aPD-L1+aTGF-β vs aTG | Up-regulated genes | CD45- |
| BP | GO:0061844 | antimicrobial humoral immune response mediat | 5.98E-07 | 7.39E-05 | Cxcl10/Cxcl9/Cxc | 4 | aPD-L1+aTGF-β vs aTG | Up-regulated genes | CD45- |
| BP | GO:1990266 | neutrophil migration | 7.08E-07 | 7.61E-05 | Cxcl10/Cxcl9/Cxc | 5 | aPD-L1+aTGF-β vs aTG | Up-regulated genes | CD45- |
| BP | GO:1903708 | positive regulation of hemopoiesis | 7.99E-07 | 7.61E-05 | Cxcl1/Irf1/Socs1/ | 6 | aPD-L1+aTGF-β vs aTG | Up-regulated genes | CD45- |
| BP | GO:0071621 | granulocyte chemotaxis | 8.46E-07 | 7.61E-05 | Cxcl10/Cxcl9/Cxc | 5 | aPD-L1+aTGF-β vs aTG | Up-regulated genes | CD45- |
| BP | GO:0019730 | antimicrobial humoral response | 1.78E-06 | 1.41E-04 | Cxcl10/Cxcl9/Cxc | 4 | aPD-L1+aTGF-β vs aTG | Up-regulated genes | CD45- |
| BP | GO:0097530 | granulocyte migration | 2.08E-06 | 1.41E-04 | Cxcl10/Cxcl9/Cxc | 5 | aPD-L1+aTGF-β vs aTG | Up-regulated genes | CD45- |
| BP | GO:0001912 | positive regulation of leukocyte mediated cytoto | 2.14E-06 | 1.41E-04 | H2-D1/Cxcl1/H2-I | 4 | aPD-L1+aTGF-β vs aTG | Up-regulated genes | CD45- |
| BP | GO:0070098 | chemokine-mediated signaling pathway | 2.14E-06 | 1.41E-04 | Cxcl10/Cxcl9/Cxc | 4 | aPD-L1+aTGF-β vs aTG | Up-regulated genes | CD45- |
| BP | GO:0043618 | regulation of transcription from RNA polymerase | 2.34E-06 | 1.45E-04 | Klf2/Atf3/Dnajb1/ | 4 | aPD-L1+aTGF-β vs aTG | Up-regulated genes | CD45- |
| BP | GO:0032496 | response to lipopolysaccharide | 3.29E-06 | 1.83E-04 | Cxcl10/Cxcl9/Cxc | 6 | aPD-L1+aTGF-β vs aTG | Up-regulated genes | CD45- |
| BP | GO:0043620 | regulation of DNA-templated transcription in res | 3.56E-06 | 1.83E-04 | Klf2/Atf3/Dnajb1/ | 4 | aPD-L1+aTGF-β vs aTG | Up-regulated genes | CD45- |
| BP | GO:0035994 | response to muscle stretch | 3.65E-06 | 1.83E-04 | Nfkbia/Jun/Fos | 3 | aPD-L1+aTGF-β vs aTG | Up-regulated genes | CD45- |
| BP | GO:0031343 | positive regulation of cell killing | 3.85E-06 | 1.83E-04 | H2-D1/Cxcl1/H2-I | 4 | aPD-L1+aTGF-β vs aTG | Up-regulated genes | CD45- |
| BP | GO:0071248 | cellular response to metal ion | 3.89E-06 | 1.83E-04 | Mt1/Mt2/B2m/Jun | 5 | aPD-L1+aTGF-β vs aTG | Up-regulated genes | CD45- |
| BP | GO:1990868 | response to chemokine | 4.83E-06 | 2.08E-04 | Cxcl10/Cxcl9/Cxc | 4 | aPD-L1+aTGF-β vs aTG | Up-regulated genes | CD45- |
| BP | GO:1990869 | cellular response to chemokine | 4.83E-06 | 2.08E-04 | Cxcl10/Cxcl9/Cxc | 4 | aPD-L1+aTGF-β vs aTG | Up-regulated genes | CD45- |
| BP | GO:1902107 | positive regulation of leukocyte differentiation | 5.39E-06 | 2.15E-04 | Irf1/Socs1/Hsp90 | 5 | aPD-L1+aTGF-β vs aTG | Up-regulated genes | CD45- |
| BP | GO:0002428 | antigen processing and presentation of peptide | 5.44E-06 | 2.15E-04 | H2-D1/H2-K1/B2r | 3 | aPD-L1+aTGF-β vs aTG | Up-regulated genes | CD45- |
| BP | GO:0001910 | regulation of leukocyte mediated cytotoxicity | 6.41E-06 | 2.44E-04 | H2-D1/Cxcl1/H2-I | 4 | aPD-L1+aTGF-β vs aTG | Up-regulated genes | CD45- |
| BP | GO:0071241 | cellular response to inorganic substance | 8.44E-06 | 3.09E-04 | Mt1/Mt2/B2m/Jun | 5 | aPD-L1+aTGF-β vs aTG | Up-regulated genes | CD45- |
| BP | GO:0002475 | antigen processing and presentation via MHC cl | 9.09E-06 | 3.21E-04 | H2-D1/H2-K1/B2r | 3 | aPD-L1+aTGF-β vs aTG | Up-regulated genes | CD45- |
| BP | GO:0071222 | cellular response to lipopolysaccharide | 1.00E-05 | 3.43E-04 | Cxcl10/Cxcl9/Cxc | 5 | aPD-L1+aTGF-β vs aTG | Up-regulated genes | CD45- |
| BP | GO:0002483 | antigen processing and presentation of endoger | 1.22E-05 | 3.91E-04 | H2-D1/H2-K1/B2r | 3 | aPD-L1+aTGF-β vs aTG | Up-regulated genes | CD45- |
| BP | GO:0019885 | antigen processing and presentation of endoger | 1.22E-05 | 3.91E-04 | H2-D1/H2-K1/B2r | 3 | aPD-L1+aTGF-β vs aTG | Up-regulated genes | CD45- |
| BP | GO:0071219 | cellular response to molecule of bacterial origin | 1.31E-05 | 4.05E-04 | Cxcl10/Cxcl9/Cxc | 5 | aPD-L1+aTGF-β vs aTG | Up-regulated genes | CD45- |
| BP | GO:0031341 | regulation of cell killing | 1.42E-05 | 4.22E-04 | H2-D1/Cxcl1/H2-I | 4 | aPD-L1+aTGF-β vs aTG | Up-regulated genes | CD45- |
| BP | GO:0019882 | antigen processing and presentation | 1.50E-05 | 4.22E-04 | H2-D1/H2-K1/B2r | 4 | aPD-L1+aTGF-β vs aTG | Up-regulated genes | CD45- |
| BP | GO:0030595 | leukocyte chemotaxis | 1.54E-05 | 4.22E-04 | Cxcl10/Cxcl9/Cxc | 5 | aPD-L1+aTGF-β vs aTG | Up-regulated genes | CD45- |
| BP | GO:0097529 | myeloid leukocyte migration | 1.54E-05 | 4.22E-04 | Cxcl10/Cxcl9/Cxc | 5 | aPD-L1+aTGF-β vs aTG | Up-regulated genes | CD45- |
| BP | GO:0019883 | antigen processing and presentation of endoger | 1.60E-05 | 4.29E-04 | H2-D1/H2-K1/B2r | 3 | aPD-L1+aTGF-β vs aTG | Up-regulated genes | CD45- |
| BP | GO:0001909 | leukocyte mediated cytotoxicity | 2.27E-05 | 5.86E-04 | H2-D1/Cxcl1/H2-I | 4 | aPD-L1+aTGF-β vs aTG | Up-regulated genes | CD45- |
| BP | GO:0001916 | positive regulation of T cell mediated cytotoxicity | 2.31E-05 | 5.86E-04 | H2-D1/H2-K1/B2r | 3 | aPD-L1+aTGF-β vs aTG | Up-regulated genes | CD45- |
| BP | GO:0071216 | cellular response to biotic stimulus | 2.40E-05 | 5.94E-04 | Cxcl10/Cxcl9/Cxc | 5 | aPD-L1+aTGF-β vs aTG | Up-regulated genes | CD45- |
| BP | GO:0002699 | positive regulation of immune effector process | 2.69E-05 | 6.48E-04 | H2-D1/Cxcl1/Irf1/ | 5 | aPD-L1+aTGF-β vs aTG | Up-regulated genes | CD45- |
| BP | GO:0002824 | positive regulation of adaptive immune respons | 2.88E-05 | 6.77E-04 | H2-D1/Irf1/H2-K1 | 4 | aPD-L1+aTGF-β vs aTG | Up-regulated genes | CD45- |
| BP | GO:0002474 | antigen processing and presentation of peptide | 3.19E-05 | 7.34E-04 | H2-D1/H2-K1/B2r | 3 | aPD-L1+aTGF-β vs aTG | Up-regulated genes | CD45- |
| BP | GO:0001914 | regulation of T cell mediated cytotoxicity | 3.53E-05 | 7.93E-04 | H2-D1/H2-K1/B2r | 3 | aPD-L1+aTGF-β vs aTG | Up-regulated genes | CD45- |
| BP | GO:0002821 | positive regulation of adaptive immune respons | 3.92E-05 | 8.60E-04 | H2-D1/Irf1/H2-K1 | 4 | aPD-L1+aTGF-β vs aTG | Up-regulated genes | CD45- |

| | | | | | | | | | |
|---|---|---|---|---|---|---|---|---|---|
| BP | GO:0098542 | defense response to other organism | 4.66E-05 | 1.00E-03 | Cxcl10/Cxcl9/Cxc | 6 | aPD-L1+aTGF-β vs aTG | Up-regulated genes | CD45- |
| BP | GO:0006959 | humoral immune response | 5.41E-05 | 1.14E-03 | Cxcl10/Cxcl9/Cxc | 4 | aPD-L1+aTGF-β vs aTG | Up-regulated genes | CD45- |
| BP | GO:0010038 | response to metal ion | 5.75E-05 | 1.18E-03 | Mt1/Mt2/B2m/Jun/ | 5 | aPD-L1+aTGF-β vs aTG | Up-regulated genes | CD45- |
| BP | GO:0001906 | cell killing | 5.84E-05 | 1.18E-03 | H2-D1/Cxcl1/H2-I | 4 | aPD-L1+aTGF-β vs aTG | Up-regulated genes | CD45- |
| BP | GO:0002705 | positive regulation of leukocyte mediated immun | 6.78E-05 | 1.34E-03 | H2-D1/Cxcl1/H2-I | 4 | aPD-L1+aTGF-β vs aTG | Up-regulated genes | CD45- |
| BP | GO:0010035 | response to inorganic substance | 7.61E-05 | 1.48E-03 | Mt1/Mt2/B2m/Krt8 | 6 | aPD-L1+aTGF-β vs aTG | Up-regulated genes | CD45- |
| BP | GO:0060326 | cell chemotaxis | 7.88E-05 | 1.50E-03 | Cxcl10/Cxcl9/Cxc | 5 | aPD-L1+aTGF-β vs aTG | Up-regulated genes | CD45- |
| BP | GO:1902105 | regulation of leukocyte differentiation | 8.58E-05 | 1.60E-03 | Irf1/Socs1/Hsp90a | 5 | aPD-L1+aTGF-β vs aTG | Up-regulated genes | CD45- |
| BP | GO:0071396 | cellular response to lipid | 1.09E-04 | 1.96E-03 | Cxcl10/Cxcl9/Cxc | 6 | aPD-L1+aTGF-β vs aTG | Up-regulated genes | CD45- |
| BP | GO:0002711 | positive regulation of T cell mediated immunity | 1.09E-04 | 1.96E-03 | H2-D1/H2-K1/B2r | 3 | aPD-L1+aTGF-β vs aTG | Up-regulated genes | CD45- |
| BP | GO:0048002 | antigen processing and presentation of peptide a | 1.17E-04 | 2.06E-03 | H2-D1/H2-K1/B2r | 3 | aPD-L1+aTGF-β vs aTG | Up-regulated genes | CD45- |
| BP | GO:0001913 | T cell mediated cytotoxicity | 1.24E-04 | 2.16E-03 | H2-D1/H2-K1/B2r | 3 | aPD-L1+aTGF-β vs aTG | Up-regulated genes | CD45- |
| BP | GO:0002822 | regulation of adaptive immune response based o | 1.36E-04 | 2.32E-03 | H2-D1/Irf1/H2-K1/ | 4 | aPD-L1+aTGF-β vs aTG | Up-regulated genes | CD45- |
| BP | GO:0050900 | leukocyte migration | 1.80E-04 | 3.02E-03 | Cxcl10/Cxcl9/Cxc | 5 | aPD-L1+aTGF-β vs aTG | Up-regulated genes | CD45- |
| BP | GO:0002521 | leukocyte differentiation | 1.84E-04 | 3.04E-03 | Irf1/Socs1/B2m/H | 6 | aPD-L1+aTGF-β vs aTG | Up-regulated genes | CD45- |
| BP | GO:0002819 | regulation of adaptive immune response | 1.97E-04 | 3.20E-03 | H2-D1/Irf1/H2-K1/ | 4 | aPD-L1+aTGF-β vs aTG | Up-regulated genes | CD45- |
| BP | GO:0002709 | regulation of T cell mediated immunity | 2.71E-04 | 4.30E-03 | H2-D1/H2-K1/B2r | 3 | aPD-L1+aTGF-β vs aTG | Up-regulated genes | CD45- |
| BP | GO:0097201 | negative regulation of transcription from RNA po | 2.74E-04 | 4.30E-03 | Dnajb1/Jun | 2 | aPD-L1+aTGF-β vs aTG | Up-regulated genes | CD45- |
| BP | GO:0055065 | metal ion homeostasis | 3.06E-04 | 4.73E-03 | Cxcl10/Mt1/Cxcl1 | 6 | aPD-L1+aTGF-β vs aTG | Up-regulated genes | CD45- |
| BP | GO:0042532 | negative regulation of tyrosine phosphorylation o | 3.24E-04 | 4.78E-03 | Irf1/Socs1 | 2 | aPD-L1+aTGF-β vs aTG | Up-regulated genes | CD45- |
| BP | GO:0043374 | CD8-positive, alpha-beta T cell differentiation | 3.24E-04 | 4.78E-03 | Irf1/Socs1 | 2 | aPD-L1+aTGF-β vs aTG | Up-regulated genes | CD45- |
| BP | GO:0071280 | cellular response to copper ion | 3.24E-04 | 4.78E-03 | Mt1/Mt2 | 2 | aPD-L1+aTGF-β vs aTG | Up-regulated genes | CD45- |
| BP | GO:0002697 | regulation of immune effector process | 3.34E-04 | 4.86E-03 | H2-D1/Cxcl1/Irf1/I | 5 | aPD-L1+aTGF-β vs aTG | Up-regulated genes | CD45- |
| BP | GO:0002703 | regulation of leukocyte mediated immunity | 3.44E-04 | 4.93E-03 | H2-D1/Cxcl1/H2-I | 4 | aPD-L1+aTGF-β vs aTG | Up-regulated genes | CD45- |
| BP | GO:0045582 | positive regulation of T cell differentiation | 4.07E-04 | 5.75E-03 | Irf1/Socs1/Hsp90a | 3 | aPD-L1+aTGF-β vs aTG | Up-regulated genes | CD45- |
| BP | GO:0045637 | regulation of myeloid cell differentiation | 4.31E-04 | 5.89E-03 | Nfkbia/B2m/Jun/F | 4 | aPD-L1+aTGF-β vs aTG | Up-regulated genes | CD45- |
| BP | GO:0002486 | antigen processing and presentation of endoger | 4.35E-04 | 5.89E-03 | H2-D1/H2-K1 | 2 | aPD-L1+aTGF-β vs aTG | Up-regulated genes | CD45- |
| BP | GO:2001185 | regulation of CD8-positive, alpha-beta T cell acti | 4.35E-04 | 5.89E-03 | Irf1/Socs1 | 2 | aPD-L1+aTGF-β vs aTG | Up-regulated genes | CD45- |
| BP | GO:0006954 | inflammatory response | 4.44E-04 | 5.93E-03 | Cxcl10/Cxcl9/Cxc | 6 | aPD-L1+aTGF-β vs aTG | Up-regulated genes | CD45- |
| BP | GO:0002476 | antigen processing and presentation of endoger | 4.96E-04 | 6.37E-03 | H2-D1/H2-K1 | 2 | aPD-L1+aTGF-β vs aTG | Up-regulated genes | CD45- |
| BP | GO:0002484 | antigen processing and presentation of endoger | 4.96E-04 | 6.37E-03 | H2-D1/H2-K1 | 2 | aPD-L1+aTGF-β vs aTG | Up-regulated genes | CD45- |
| BP | GO:0071498 | cellular response to fluid shear stress | 4.96E-04 | 6.37E-03 | Klf2/Spp1 | 2 | aPD-L1+aTGF-β vs aTG | Up-regulated genes | CD45- |
| BP | GO:0072503 | cellular divalent inorganic cation homeostasis | 5.12E-04 | 6.49E-03 | Cxcl10/Mt1/Cxcl1 | 5 | aPD-L1+aTGF-β vs aTG | Up-regulated genes | CD45- |
| BP | GO:0030099 | myeloid cell differentiation | 5.56E-04 | 6.85E-03 | Klf2/Nfkbia/B2m/J | 5 | aPD-L1+aTGF-β vs aTG | Up-regulated genes | CD45- |
| BP | GO:0007263 | nitric oxide mediated signal transduction | 5.61E-04 | 6.85E-03 | Mt1/Mt2 | 2 | aPD-L1+aTGF-β vs aTG | Up-regulated genes | CD45- |
| BP | GO:1901739 | regulation of myoblast fusion | 5.61E-04 | 6.85E-03 | Cxcl10/Cxcl9 | 2 | aPD-L1+aTGF-β vs aTG | Up-regulated genes | CD45- |
| BP | GO:0045621 | positive regulation of lymphocyte differentiation | 6.25E-04 | 7.53E-03 | Irf1/Socs1/Hsp90a | 3 | aPD-L1+aTGF-β vs aTG | Up-regulated genes | CD45- |
| BP | GO:0002708 | positive regulation of lymphocyte mediated immu | 6.71E-04 | 7.99E-03 | H2-D1/H2-K1/B2r | 3 | aPD-L1+aTGF-β vs aTG | Up-regulated genes | CD45- |
| BP | GO:0072507 | divalent inorganic cation homeostasis | 6.98E-04 | 8.19E-03 | Cxcl10/Mt1/Cxcl1 | 5 | aPD-L1+aTGF-β vs aTG | Up-regulated genes | CD45- |
| BP | GO:0006882 | cellular zinc ion homeostasis | 7.04E-04 | 8.19E-03 | Mt1/Mt2 | 2 | aPD-L1+aTGF-β vs aTG | Up-regulated genes | CD45- |
| BP | GO:0010043 | response to zinc ion | 7.81E-04 | 8.59E-03 | Mt1/Mt2 | 2 | aPD-L1+aTGF-β vs aTG | Up-regulated genes | CD45- |
| BP | GO:0046639 | negative regulation of alpha-beta T cell differenti | 7.81E-04 | 8.59E-03 | Irf1/Socs1 | 2 | aPD-L1+aTGF-β vs aTG | Up-regulated genes | CD45- |
| BP | GO:0046688 | response to copper ion | 7.81E-04 | 8.59E-03 | Mt1/Mt2 | 2 | aPD-L1+aTGF-β vs aTG | Up-regulated genes | CD45- |
| BP | GO:0055069 | zinc ion homeostasis | 7.81E-04 | 8.59E-03 | Mt1/Mt2 | 2 | aPD-L1+aTGF-β vs aTG | Up-regulated genes | CD45- |
| BP | GO:0060333 | interferon-gamma-mediated signaling pathway | 7.81E-04 | 8.59E-03 | Irf1/Socs1 | 2 | aPD-L1+aTGF-β vs aTG | Up-regulated genes | CD45- |
| BP | GO:0030217 | T cell differentiation | 8.01E-04 | 8.71E-03 | Irf1/Socs1/B2m/H | 4 | aPD-L1+aTGF-β vs aTG | Up-regulated genes | CD45- |
| BP | GO:0002456 | T cell mediated immunity | 8.49E-04 | 8.88E-03 | H2-D1/H2-K1/B2r | 3 | aPD-L1+aTGF-β vs aTG | Up-regulated genes | CD45- |
| BP | GO:0036037 | CD8-positive, alpha-beta T cell activation | 8.62E-04 | 8.88E-03 | Irf1/Socs1 | 2 | aPD-L1+aTGF-β vs aTG | Up-regulated genes | CD45- |
| BP | GO:0043372 | positive regulation of CD4-positive, alpha-beta T | 8.62E-04 | 8.88E-03 | Irf1/Socs1 | 2 | aPD-L1+aTGF-β vs aTG | Up-regulated genes | CD45- |
| BP | GO:0046426 | negative regulation of JAK-STAT cascade | 8.62E-04 | 8.88E-03 | Irf1/Socs1 | 2 | aPD-L1+aTGF-β vs aTG | Up-regulated genes | CD45- |
| BP | GO:1904893 | negative regulation of STAT cascade | 8.62E-04 | 8.88E-03 | Irf1/Socs1 | 2 | aPD-L1+aTGF-β vs aTG | Up-regulated genes | CD45- |
| BP | GO:2000379 | positive regulation of reactive oxygen species m | 9.05E-04 | 9.23E-03 | Klf2/Cxcl1/Hsp90a | 3 | aPD-L1+aTGF-β vs aTG | Up-regulated genes | CD45- |
| BP | GO:0002478 | antigen processing and presentation of exogenc | 9.47E-04 | 9.37E-03 | H2-K1/B2m | 2 | aPD-L1+aTGF-β vs aTG | Up-regulated genes | CD45- |
| BP | GO:0034405 | response to fluid shear stress | 9.47E-04 | 9.37E-03 | Klf2/Spp1 | 2 | aPD-L1+aTGF-β vs aTG | Up-regulated genes | CD45- |
| BP | GO:0045589 | regulation of regulatory T cell differentiation | 9.47E-04 | 9.37E-03 | Irf1/Socs1 | 2 | aPD-L1+aTGF-β vs aTG | Up-regulated genes | CD45- |
| BP | GO:0002460 | adaptive immune response based on somatic re | 9.58E-04 | 9.38E-03 | H2-D1/Irf1/H2-K1/ | 4 | aPD-L1+aTGF-β vs aTG | Up-regulated genes | CD45- |
| BP | GO:0045066 | regulatory T cell differentiation | 1.04E-03 | 9.95E-03 | Irf1/Socs1 | 2 | aPD-L1+aTGF-β vs aTG | Up-regulated genes | CD45- |
| BP | GO:0060142 | regulation of syncytium formation by plasma mer | 1.04E-03 | 9.95E-03 | Cxcl10/Cxcl9 | 2 | aPD-L1+aTGF-β vs aTG | Up-regulated genes | CD45- |
| BP | GO:0036003 | positive regulation of transcription from RNA poly | 1.13E-03 | 1.06E-02 | Klf2/Atf3 | 2 | aPD-L1+aTGF-β vs aTG | Up-regulated genes | CD45- |
| BP | GO:2000516 | positive regulation of CD4-positive, alpha-beta T | 1.13E-03 | 1.06E-02 | Irf1/Socs1 | 2 | aPD-L1+aTGF-β vs aTG | Up-regulated genes | CD45- |
| BP | GO:0006875 | cellular metal ion homeostasis | 1.15E-03 | 1.07E-02 | Cxcl10/Mt1/Cxcl1 | 5 | aPD-L1+aTGF-β vs aTG | Up-regulated genes | CD45- |
| BP | GO:0055076 | transition metal ion homeostasis | 1.15E-03 | 1.07E-02 | Mt1/Mt2/B2m | 3 | aPD-L1+aTGF-β vs aTG | Up-regulated genes | CD45- |
| BP | GO:0009612 | response to mechanical stimulus | 1.22E-03 | 1.12E-02 | Nfkbia/Jun/Fos | 3 | aPD-L1+aTGF-β vs aTG | Up-regulated genes | CD45- |
| BP | GO:0019884 | antigen processing and presentation of exogenc | 1.43E-03 | 1.30E-02 | H2-K1/B2m | 2 | aPD-L1+aTGF-β vs aTG | Up-regulated genes | CD45- |
| BP | GO:0042026 | protein refolding | 1.54E-03 | 1.38E-02 | B2m/Hsp90aa1 | 2 | aPD-L1+aTGF-β vs aTG | Up-regulated genes | CD45- |
| BP | GO:0050778 | positive regulation of immune response | 1.63E-03 | 1.45E-02 | H2-D1/Irf1/H2-K1/ | 5 | aPD-L1+aTGF-β vs aTG | Up-regulated genes | CD45- |
| BP | GO:0045580 | regulation of T cell differentiation | 1.64E-03 | 1.45E-02 | Irf1/Socs1/Hsp90a | 3 | aPD-L1+aTGF-β vs aTG | Up-regulated genes | CD45- |
| BP | GO:0002443 | leukocyte mediated immunity | 1.65E-03 | 1.45E-02 | H2-D1/Cxcl1/H2-I | 4 | aPD-L1+aTGF-β vs aTG | Up-regulated genes | CD45- |
| BP | GO:0051146 | striated muscle cell differentiation | 1.68E-03 | 1.45E-02 | Cxcl10/Cxcl9/Krt8 | 4 | aPD-L1+aTGF-β vs aTG | Up-regulated genes | CD45- |
| BP | GO:0006935 | chemotaxis | 1.77E-03 | 1.52E-02 | Cxcl10/Cxcl9/Cxc | 5 | aPD-L1+aTGF-β vs aTG | Up-regulated genes | CD45- |
| BP | GO:0042330 | taxis | 1.83E-03 | 1.56E-02 | Cxcl10/Cxcl9/Cxc | 5 | aPD-L1+aTGF-β vs aTG | Up-regulated genes | CD45- |
| BP | GO:0030003 | cellular cation homeostasis | 1.85E-03 | 1.56E-02 | Cxcl10/Mt1/Cxcl1 | 5 | aPD-L1+aTGF-β vs aTG | Up-regulated genes | CD45- |
| BP | GO:0002706 | regulation of lymphocyte mediated immunity | 1.95E-03 | 1.63E-02 | H2-D1/H2-K1/B2r | 3 | aPD-L1+aTGF-β vs aTG | Up-regulated genes | CD45- |
| BP | GO:0007520 | myoblast fusion | 2.01E-03 | 1.64E-02 | Cxcl10/Cxcl9 | 2 | aPD-L1+aTGF-β vs aTG | Up-regulated genes | CD45- |
| BP | GO:0046636 | negative regulation of alpha-beta T cell activatio | 2.01E-03 | 1.64E-02 | Irf1/Socs1 | 2 | aPD-L1+aTGF-β vs aTG | Up-regulated genes | CD45- |
| BP | GO:1902895 | positive regulation of pri-miRNA transcription by | 2.01E-03 | 1.64E-02 | Jun/Fos | 2 | aPD-L1+aTGF-β vs aTG | Up-regulated genes | CD45- |
| BP | GO:0006873 | cellular ion homeostasis | 2.02E-03 | 1.64E-02 | Cxcl10/Mt1/Cxcl1 | 5 | aPD-L1+aTGF-β vs aTG | Up-regulated genes | CD45- |
| BP | GO:0071356 | cellular response to tumor necrosis factor | 2.24E-03 | 1.80E-02 | Irf1/Nfkbia/Krt8 | 3 | aPD-L1+aTGF-β vs aTG | Up-regulated genes | CD45- |
| BP | GO:0034105 | positive regulation of tissue remodeling | 2.27E-03 | 1.81E-02 | Klf6/Spp1 | 2 | aPD-L1+aTGF-β vs aTG | Up-regulated genes | CD45- |
| BP | GO:1903707 | negative regulation of hemopoiesis | 2.34E-03 | 1.85E-02 | Irf1/Socs1/Nfkbia | 3 | aPD-L1+aTGF-β vs aTG | Up-regulated genes | CD45- |
| BP | GO:0002250 | adaptive immune response | 2.41E-03 | 1.89E-02 | H2-D1/Irf1/H2-K1/ | 4 | aPD-L1+aTGF-β vs aTG | Up-regulated genes | CD45- |
| BP | GO:0030098 | lymphocyte differentiation | 2.50E-03 | 1.95E-02 | Irf1/Socs1/B2m/H | 4 | aPD-L1+aTGF-β vs aTG | Up-regulated genes | CD45- |
| BP | GO:0043370 | regulation of CD4-positive, alpha-beta T cell diffe | 2.68E-03 | 2.04E-02 | Irf1/Socs1 | 2 | aPD-L1+aTGF-β vs aTG | Up-regulated genes | CD45- |
| BP | GO:0045581 | negative regulation of T cell differentiation | 2.68E-03 | 2.04E-02 | Irf1/Socs1 | 2 | aPD-L1+aTGF-β vs aTG | Up-regulated genes | CD45- |
| BP | GO:0046638 | positive regulation of alpha-beta T cell differentia | 2.68E-03 | 2.04E-02 | Irf1/Socs1 | 2 | aPD-L1+aTGF-β vs aTG | Up-regulated genes | CD45- |
| BP | GO:0045619 | regulation of lymphocyte differentiation | 2.72E-03 | 2.06E-02 | Irf1/Socs1/Hsp90a | 3 | aPD-L1+aTGF-β vs aTG | Up-regulated genes | CD45- |
| BP | GO:0050870 | positive regulation of T cell activation | 2.78E-03 | 2.08E-02 | Irf1/Socs1/Hsp90a | 3 | aPD-L1+aTGF-β vs aTG | Up-regulated genes | CD45- |
| BP | GO:0006457 | protein folding | 3.02E-03 | 2.24E-02 | Dnajb1/B2m/Hsp9 | 3 | aPD-L1+aTGF-β vs aTG | Up-regulated genes | CD45- |
| BP | GO:0045429 | positive regulation of nitric oxide biosynthetic pro | 3.13E-03 | 2.30E-02 | Klf2/Hsp90aa1 | 2 | aPD-L1+aTGF-β vs aTG | Up-regulated genes | CD45- |

| | | | | | | | | | |
|---|---|---|---|---|---|---|---|---|---|
| BP | GO:0034612 | response to tumor necrosis factor | 3.14E-03 | 2.30E-02 | Irf1/Nfkbia/Krt8 | 3 | aPD-L1+aTGF-β vs aTG | Up-regulated genes | CD45- |
| BP | GO:0098754 | detoxification | 3.28E-03 | 2.35E-02 | Mt1/Mt2 | 2 | aPD-L1+aTGF-β vs aTG | Up-regulated genes | CD45- |
| BP | GO:1902893 | regulation of pri-miRNA transcription by RNA po | 3.28E-03 | 2.35E-02 | Jun/Fos | 2 | aPD-L1+aTGF-β vs aTG | Up-regulated genes | CD45- |
| BP | GO:1904407 | positive regulation of nitric oxide metabolic proc | 3.28E-03 | 2.35E-02 | Klf2/Hsp90aa1 | 2 | aPD-L1+aTGF-β vs aTG | Up-regulated genes | CD45- |
| BP | GO:0045620 | negative regulation of lymphocyte differentiation | 3.44E-03 | 2.45E-02 | Irf1/Socs1 | 2 | aPD-L1+aTGF-β vs aTG | Up-regulated genes | CD45- |
| BP | GO:0009991 | response to extracellular stimulus | 3.62E-03 | 2.56E-02 | Atf3/Spp1/Jun/Fos | 4 | aPD-L1+aTGF-β vs aTG | Up-regulated genes | CD45- |
| BP | GO:1903039 | positive regulation of leukocyte cell-cell adhesio | 3.74E-03 | 2.62E-02 | Irf1/Socs1/Hsp90a | 3 | aPD-L1+aTGF-β vs aTG | Up-regulated genes | CD45- |
| BP | GO:0035458 | cellular response to interferon-beta | 3.77E-03 | 2.63E-02 | Irf1/Iigp1 | 2 | aPD-L1+aTGF-β vs aTG | Up-regulated genes | CD45- |
| BP | GO:0010977 | negative regulation of neuron projection develop | 3.81E-03 | 2.63E-02 | H2-D1/H2-K1/B2r | 3 | aPD-L1+aTGF-β vs aTG | Up-regulated genes | CD45- |
| BP | GO:0000768 | syncytium formation by plasma membrane fusion | 3.94E-03 | 2.69E-02 | Cxcl10/Cxcl9 | 2 | aPD-L1+aTGF-β vs aTG | Up-regulated genes | CD45- |
| BP | GO:0140253 | cell-cell fusion | 3.94E-03 | 2.69E-02 | Cxcl10/Cxcl9 | 2 | aPD-L1+aTGF-β vs aTG | Up-regulated genes | CD45- |
| BP | GO:0045661 | regulation of myoblast differentiation | 4.12E-03 | 2.79E-02 | Cxcl10/Cxcl9 | 2 | aPD-L1+aTGF-β vs aTG | Up-regulated genes | CD45- |
| BP | GO:0042692 | muscle cell differentiation | 4.25E-03 | 2.83E-02 | Cxcl10/Cxcl9/Krt8 | 4 | aPD-L1+aTGF-β vs aTG | Up-regulated genes | CD45- |
| BP | GO:0006949 | syncytium formation | 4.29E-03 | 2.83E-02 | Cxcl10/Cxcl9 | 2 | aPD-L1+aTGF-β vs aTG | Up-regulated genes | CD45- |
| BP | GO:0010830 | regulation of myotube differentiation | 4.29E-03 | 2.83E-02 | Cxcl10/Cxcl9 | 2 | aPD-L1+aTGF-β vs aTG | Up-regulated genes | CD45- |
| BP | GO:2000514 | regulation of CD4-positive, alpha-beta T cell acti | 4.29E-03 | 2.83E-02 | Irf1/Socs1 | 2 | aPD-L1+aTGF-β vs aTG | Up-regulated genes | CD45- |
| BP | GO:0002763 | positive regulation of myeloid leukocyte different | 4.47E-03 | 2.89E-02 | Jun/Fos | 2 | aPD-L1+aTGF-β vs aTG | Up-regulated genes | CD45- |
| BP | GO:0050732 | negative regulation of peptidyl-tyrosine phospho | 4.47E-03 | 2.89E-02 | Irf1/Socs1 | 2 | aPD-L1+aTGF-β vs aTG | Up-regulated genes | CD45- |
| BP | GO:0061614 | pri-miRNA transcription by RNA polymerase II | 4.47E-03 | 2.89E-02 | Jun/Fos | 2 | aPD-L1+aTGF-β vs aTG | Up-regulated genes | CD45- |
| BP | GO:0046635 | positive regulation of alpha-beta T cell activation | 4.66E-03 | 2.99E-02 | Irf1/Socs1 | 2 | aPD-L1+aTGF-β vs aTG | Up-regulated genes | CD45- |
| BP | GO:0032370 | positive regulation of lipid transport | 4.85E-03 | 3.09E-02 | Nfkbia/Spp1 | 2 | aPD-L1+aTGF-β vs aTG | Up-regulated genes | CD45- |
| BP | GO:0046637 | regulation of alpha-beta T cell differentiation | 5.04E-03 | 3.17E-02 | Irf1/Socs1 | 2 | aPD-L1+aTGF-β vs aTG | Up-regulated genes | CD45- |
| BP | GO:1903428 | positive regulation of reactive oxygen species bi | 5.04E-03 | 3.17E-02 | Klf2/Hsp90aa1 | 2 | aPD-L1+aTGF-β vs aTG | Up-regulated genes | CD45- |
| BP | GO:0045785 | positive regulation of cell adhesion | 5.20E-03 | 3.26E-02 | Irf1/Socs1/Hsp90a | 4 | aPD-L1+aTGF-β vs aTG | Up-regulated genes | CD45- |
| BP | GO:0035456 | response to interferon-beta | 5.43E-03 | 3.36E-02 | Irf1/Iigp1 | 2 | aPD-L1+aTGF-β vs aTG | Up-regulated genes | CD45- |
| BP | GO:0042509 | regulation of tyrosine phosphorylation of STAT p | 5.43E-03 | 3.36E-02 | Irf1/Socs1 | 2 | aPD-L1+aTGF-β vs aTG | Up-regulated genes | CD45- |
| BP | GO:0070997 | neuron death | 5.48E-03 | 3.36E-02 | Mt1/Cxcl2/Jun/Fo | 4 | aPD-L1+aTGF-β vs aTG | Up-regulated genes | CD45- |
| BP | GO:0071277 | cellular response to calcium ion | 5.63E-03 | 3.42E-02 | Jun/Fos | 2 | aPD-L1+aTGF-β vs aTG | Up-regulated genes | CD45- |
| BP | GO:2000377 | regulation of reactive oxygen species metabolic | 5.64E-03 | 3.42E-02 | Klf2/Cxcl1/Hsp90 | 3 | aPD-L1+aTGF-β vs aTG | Up-regulated genes | CD45- |
| BP | GO:0042110 | T cell activation | 5.82E-03 | 3.47E-02 | Irf1/Socs1/B2m/H | 4 | aPD-L1+aTGF-β vs aTG | Up-regulated genes | CD45- |
| BP | GO:0031345 | negative regulation of cell projection organizatio | 5.82E-03 | 3.47E-02 | H2-D1/H2-K1/B2r | 3 | aPD-L1+aTGF-β vs aTG | Up-regulated genes | CD45- |
| BP | GO:0051607 | defense response to virus | 5.82E-03 | 3.47E-02 | Cxcl10/Cxcl9/Irf1 | 3 | aPD-L1+aTGF-β vs aTG | Up-regulated genes | CD45- |
| BP | GO:0007260 | tyrosine phosphorylation of STAT protein | 6.05E-03 | 3.58E-02 | Irf1/Socs1 | 2 | aPD-L1+aTGF-β vs aTG | Up-regulated genes | CD45- |
| BP | GO:0001836 | release of cytochrome c from mitochondria | 6.26E-03 | 3.66E-02 | Lmna/Jun | 2 | aPD-L1+aTGF-β vs aTG | Up-regulated genes | CD45- |
| BP | GO:0033209 | tumor necrosis factor-mediated signaling pathwa | 6.26E-03 | 3.66E-02 | Nfkbia/Krt8 | 2 | aPD-L1+aTGF-β vs aTG | Up-regulated genes | CD45- |
| BP | GO:0001933 | negative regulation of protein phosphorylation | 6.29E-03 | 3.66E-02 | Atf3/Irf1/Socs1/Ju | 4 | aPD-L1+aTGF-β vs aTG | Up-regulated genes | CD45- |
| BP | GO:0051851 | modification by host of symbiont morphology or | 6.47E-03 | 3.74E-02 | Cxcl1/Jun | 2 | aPD-L1+aTGF-β vs aTG | Up-regulated genes | CD45- |
| BP | GO:0045428 | regulation of nitric oxide biosynthetic process | 6.69E-03 | 3.82E-02 | Klf2/Hsp90aa1 | 2 | aPD-L1+aTGF-β vs aTG | Up-regulated genes | CD45- |
| BP | GO:0060395 | SMAD protein signal transduction | 6.69E-03 | 3.82E-02 | Jun/Fos | 2 | aPD-L1+aTGF-β vs aTG | Up-regulated genes | CD45- |
| BP | GO:0022409 | positive regulation of cell-cell adhesion | 6.77E-03 | 3.84E-02 | Irf1/Socs1/Hsp90a | 3 | aPD-L1+aTGF-β vs aTG | Up-regulated genes | CD45- |
| BP | GO:0060537 | muscle tissue development | 6.80E-03 | 3.84E-02 | Atf3/Zfand5/Lmna | 4 | aPD-L1+aTGF-β vs aTG | Up-regulated genes | CD45- |
| BP | GO:0017038 | protein import | 6.87E-03 | 3.84E-02 | Nfkbia/Lmna/Hsp | 3 | aPD-L1+aTGF-β vs aTG | Up-regulated genes | CD45- |
| BP | GO:0035914 | skeletal muscle cell differentiation | 6.91E-03 | 3.84E-02 | Atf3/Fos | 2 | aPD-L1+aTGF-β vs aTG | Up-regulated genes | CD45- |
| BP | GO:0043367 | CD4-positive, alpha-beta T cell differentiation | 6.91E-03 | 3.84E-02 | Irf1/Socs1 | 2 | aPD-L1+aTGF-β vs aTG | Up-regulated genes | CD45- |
| BP | GO:0002449 | lymphocyte mediated immunity | 7.08E-03 | 3.91E-02 | H2-D1/H2-K1/B2r | 3 | aPD-L1+aTGF-β vs aTG | Up-regulated genes | CD45- |
| BP | GO:0051251 | positive regulation of lymphocyte activation | 7.49E-03 | 4.12E-02 | Irf1/Socs1/Hsp90a | 3 | aPD-L1+aTGF-β vs aTG | Up-regulated genes | CD45- |
| BP | GO:0051702 | interaction with symbiont | 7.60E-03 | 4.15E-02 | Cxcl1/Jun | 2 | aPD-L1+aTGF-β vs aTG | Up-regulated genes | CD45- |
| BP | GO:0031668 | cellular response to extracellular stimulus | 7.71E-03 | 4.19E-02 | Atf3/Jun/Fos | 3 | aPD-L1+aTGF-β vs aTG | Up-regulated genes | CD45- |
| BP | GO:1905954 | positive regulation of lipid localization | 8.07E-03 | 4.36E-02 | Nfkbia/Spp1 | 2 | aPD-L1+aTGF-β vs aTG | Up-regulated genes | CD45- |
| BP | GO:0006809 | nitric oxide biosynthetic process | 8.31E-03 | 4.45E-02 | Klf2/Hsp90aa1 | 2 | aPD-L1+aTGF-β vs aTG | Up-regulated genes | CD45- |
| BP | GO:0007204 | positive regulation of cytosolic calcium ion conce | 8.37E-03 | 4.45E-02 | Cxcl10/Cxcl1/Cxc | 3 | aPD-L1+aTGF-β vs aTG | Up-regulated genes | CD45- |
| BP | GO:0097191 | extrinsic apoptotic signaling pathway | 8.37E-03 | 4.45E-02 | Atf3/Krt8/Lmna | 3 | aPD-L1+aTGF-β vs aTG | Up-regulated genes | CD45- |
| BP | GO:0001101 | response to acid chemical | 8.72E-03 | 4.61E-02 | Klf2/Socs1/Krt8 | 3 | aPD-L1+aTGF-β vs aTG | Up-regulated genes | CD45- |
| BP | GO:0034103 | regulation of tissue remodeling | 9.06E-03 | 4.70E-02 | Klf6/Spp1 | 2 | aPD-L1+aTGF-β vs aTG | Up-regulated genes | CD45- |
| BP | GO:0045445 | myoblast differentiation | 9.06E-03 | 4.70E-02 | Cxcl10/Cxcl9 | 2 | aPD-L1+aTGF-β vs aTG | Up-regulated genes | CD45- |
| BP | GO:0046209 | nitric oxide metabolic process | 9.06E-03 | 4.70E-02 | Klf2/Hsp90aa1 | 2 | aPD-L1+aTGF-β vs aTG | Up-regulated genes | CD45- |
| BP | GO:0042326 | negative regulation of phosphorylation | 9.07E-03 | 4.70E-02 | Atf3/Irf1/Socs1/Ju | 4 | aPD-L1+aTGF-β vs aTG | Up-regulated genes | CD45- |
| BP | GO:0045665 | negative regulation of neuron differentiation | 9.43E-03 | 4.86E-02 | H2-D1/H2-K1/B2r | 3 | aPD-L1+aTGF-β vs aTG | Up-regulated genes | CD45- |
| BP | GO:0009615 | response to virus | 9.55E-03 | 4.89E-02 | Cxcl10/Cxcl9/Irf1 | 3 | aPD-L1+aTGF-β vs aTG | Up-regulated genes | CD45- |
| BP | GO:0032870 | cellular response to hormone stimulus | 9.71E-03 | 4.95E-02 | Irf1/Socs1/Spp1/J | 4 | aPD-L1+aTGF-β vs aTG | Up-regulated genes | CD45- |
| CC | GO:0042612 | MHC class I protein complex | 1.27E-06 | 1.51E-04 | H2-D1/H2-K1/B2r | 3 | aPD-L1+aTGF-β vs aTG | Up-regulated genes | CD45- |
| CC | GO:0042824 | MHC class I peptide loading complex | 2.79E-06 | 1.66E-04 | H2-D1/H2-K1/B2r | 3 | aPD-L1+aTGF-β vs aTG | Up-regulated genes | CD45- |
| CC | GO:0042611 | MHC protein complex | 7.38E-06 | 2.62E-04 | H2-D1/H2-K1/B2r | 3 | aPD-L1+aTGF-β vs aTG | Up-regulated genes | CD45- |
| CC | GO:0009897 | external side of plasma membrane | 8.82E-06 | 2.62E-04 | Cxcl10/Cxcl9/H2- | 6 | aPD-L1+aTGF-β vs aTG | Up-regulated genes | CD45- |
| CC | GO:0098552 | side of membrane | 1.22E-04 | 2.91E-03 | Cxcl10/Cxcl9/H2- | 6 | aPD-L1+aTGF-β vs aTG | Up-regulated genes | CD45- |
| CC | GO:0005797 | Golgi medial cisterna | 1.00E-03 | 1.99E-02 | H2-D1/H2-K1 | 2 | aPD-L1+aTGF-β vs aTG | Up-regulated genes | CD45- |
| CC | GO:0070971 | endoplasmic reticulum exit site | 1.28E-03 | 2.18E-02 | H2-D1/H2-K1 | 2 | aPD-L1+aTGF-β vs aTG | Up-regulated genes | CD45- |
| CC | GO:0030176 | integral component of endoplasmic reticulum me | 2.24E-03 | 3.33E-02 | H2-D1/H2-K1/B2r | 3 | aPD-L1+aTGF-β vs aTG | Up-regulated genes | CD45- |
| CC | GO:0031227 | intrinsic component of endoplasmic reticulum me | 2.60E-03 | 3.44E-02 | H2-D1/H2-K1/B2r | 3 | aPD-L1+aTGF-β vs aTG | Up-regulated genes | CD45- |
| MF | GO:0008009 | chemokine activity | 6.17E-07 | 6.42E-05 | Cxcl10/Cxcl9/Cxc | 4 | aPD-L1+aTGF-β vs aTG | Up-regulated genes | CD45- |
| MF | GO:0042379 | chemokine receptor binding | 1.83E-06 | 9.54E-05 | Cxcl10/Cxcl9/Cxc | 4 | aPD-L1+aTGF-β vs aTG | Up-regulated genes | CD45- |
| MF | GO:0005125 | cytokine activity | 4.05E-06 | 1.40E-04 | Cxcl10/Cxcl9/Cxc | 5 | aPD-L1+aTGF-β vs aTG | Up-regulated genes | CD45- |
| MF | GO:0030881 | beta-2-microglobulin binding | 2.32E-04 | 3.55E-03 | H2-D1/H2-K1 | 2 | aPD-L1+aTGF-β vs aTG | Up-regulated genes | CD45- |
| MF | GO:0042610 | CD8 receptor binding | 2.32E-04 | 3.55E-03 | H2-D1/H2-K1 | 2 | aPD-L1+aTGF-β vs aTG | Up-regulated genes | CD45- |
| MF | GO:0046977 | TAP binding | 2.32E-04 | 3.55E-03 | H2-D1/H2-K1 | 2 | aPD-L1+aTGF-β vs aTG | Up-regulated genes | CD45- |
| MF | GO:0048018 | receptor ligand activity | 2.39E-04 | 3.55E-03 | Cxcl10/Cxcl9/Cxc | 5 | aPD-L1+aTGF-β vs aTG | Up-regulated genes | CD45- |
| MF | GO:0030545 | receptor regulator activity | 3.68E-04 | 4.79E-03 | Cxcl10/Cxcl9/Cxc | 5 | aPD-L1+aTGF-β vs aTG | Up-regulated genes | CD45- |
| MF | GO:0042608 | T cell receptor binding | 4.41E-04 | 5.10E-03 | H2-D1/H2-K1 | 2 | aPD-L1+aTGF-β vs aTG | Up-regulated genes | CD45- |
| MF | GO:0005126 | cytokine receptor binding | 8.10E-04 | 7.59E-03 | Cxcl10/Cxcl9/Cxc | 4 | aPD-L1+aTGF-β vs aTG | Up-regulated genes | CD45- |
| MF | GO:0001664 | G protein-coupled receptor binding | 8.71E-04 | 7.59E-03 | Cxcl10/Cxcl9/Cxc | 4 | aPD-L1+aTGF-β vs aTG | Up-regulated genes | CD45- |
| MF | GO:0042605 | peptide antigen binding | 8.76E-04 | 7.59E-03 | H2-D1/H2-K1 | 2 | aPD-L1+aTGF-β vs aTG | Up-regulated genes | CD45- |
| MF | GO:0070412 | R-SMAD binding | 9.62E-04 | 7.70E-03 | Jun/Fos | 2 | aPD-L1+aTGF-β vs aTG | Up-regulated genes | CD45- |
| MF | GO:0000978 | RNA polymerase II proximal promoter sequence | 1.78E-03 | 1.28E-02 | Atf3/Irf1/Klf6/Jun/F | 5 | aPD-L1+aTGF-β vs aTG | Up-regulated genes | CD45- |
| MF | GO:0046982 | protein heterodimerization activity | 1.87E-03 | 1.28E-02 | H2-D1/Atf3/H2-K1 | 5 | aPD-L1+aTGF-β vs aTG | Up-regulated genes | CD45- |
| MF | GO:0000987 | proximal promoter sequence-specific DNA bindi | 1.97E-03 | 1.28E-02 | Atf3/Irf1/Klf6/Jun/F | 5 | aPD-L1+aTGF-β vs aTG | Up-regulated genes | CD45- |
| MF | GO:0001102 | RNA polymerase II activating transcription factor | 2.17E-03 | 1.33E-02 | Jun/Fos | 2 | aPD-L1+aTGF-β vs aTG | Up-regulated genes | CD45- |
| MF | GO:0003823 | antigen binding | 2.44E-03 | 1.41E-02 | H2-D1/H2-K1 | 2 | aPD-L1+aTGF-β vs aTG | Up-regulated genes | CD45- |
| MF | GO:0031625 | ubiquitin protein ligase binding | 2.58E-03 | 1.41E-02 | Ubc/Nfkbia/Hsp9( | 4 | aPD-L1+aTGF-β vs aTG | Up-regulated genes | CD45- |
| MF | GO:0044389 | ubiquitin-like protein ligase binding | 3.15E-03 | 1.64E-02 | Ubc/Nfkbia/Hsp9( | 4 | aPD-L1+aTGF-β vs aTG | Up-regulated genes | CD45- |

| | | | | | | | | | |
|---|---|---|---|---|---|---|---|---|---|
| MF | GO:0001228 | DNA-binding transcription activator activity, RNA | 3.90E-03 | 1.93E-02 | Irf1/Klf6/Jun/Fos | 4 | aPD-L1+aTGF-β vs aTG | Up-regulated genes | CD45- |
| MF | GO:0097110 | scaffold protein binding | 6.57E-03 | 3.11E-02 | Krt8/Hsp90aa1 | 2 | aPD-L1+aTGF-β vs aTG | Up-regulated genes | CD45- |
| MF | GO:0033613 | activating transcription factor binding | 8.44E-03 | 3.82E-02 | Jun/Fos | 2 | aPD-L1+aTGF-β vs aTG | Up-regulated genes | CD45- |
| MF | GO:0042277 | peptide binding | 1.03E-02 | 4.45E-02 | H2-D1/H2-K1/Nfk | 3 | aPD-L1+aTGF-β vs aTG | Up-regulated genes | CD45- |
| MF | GO:0046332 | SMAD binding | 1.11E-02 | 4.60E-02 | Jun/Fos | 2 | aPD-L1+aTGF-β vs aTG | Up-regulated genes | CD45- |
| BP | GO:0051607 | defense response to virus | 1.40E-15 | 1.74E-12 | Trim30a/Mx1/Oas | 13 | aPD-L1+aTGF-β vs aTG | Down-regulated gen | CD45+ |
| BP | GO:0009615 | response to virus | 1.36E-14 | 8.51E-12 | Trim30a/Mx1/Oas | 13 | aPD-L1+aTGF-β vs aTG | Down-regulated gen | CD45+ |
| BP | GO:0098542 | defense response to other organism | 4.09E-12 | 1.70E-09 | Trim30a/Mx1/Oas | 13 | aPD-L1+aTGF-β vs aTG | Down-regulated gen | CD45+ |
| BP | GO:0050792 | regulation of viral process | 2.02E-10 | 6.32E-08 | Trim30a/Pfn1/Jun | 9 | aPD-L1+aTGF-β vs aTG | Down-regulated gen | CD45+ |
| BP | GO:0043903 | regulation of symbiosis, encompassing mutualis | 9.52E-10 | 2.38E-07 | Trim30a/Pfn1/Jun | 9 | aPD-L1+aTGF-β vs aTG | Down-regulated gen | CD45+ |
| BP | GO:0016032 | viral process | 1.21E-08 | 2.24E-06 | Trim30a/Pfn1/Jun | 9 | aPD-L1+aTGF-β vs aTG | Down-regulated gen | CD45+ |
| BP | GO:0009617 | response to bacterium | 1.26E-08 | 2.24E-06 | Trim30a/Slfn4/Us | 11 | aPD-L1+aTGF-β vs aTG | Down-regulated gen | CD45+ |
| BP | GO:0048525 | negative regulation of viral process | 7.40E-08 | 1.16E-05 | Jun/Oasl1/Isg20/E | 6 | aPD-L1+aTGF-β vs aTG | Down-regulated gen | CD45+ |
| BP | GO:0044403 | symbiont process | 9.28E-08 | 1.29E-05 | Trim30a/Pfn1/Jun | 9 | aPD-L1+aTGF-β vs aTG | Down-regulated gen | CD45+ |
| BP | GO:0044419 | interspecies interaction between organisms | 1.37E-07 | 1.72E-05 | Trim30a/Pfn1/Jun | 9 | aPD-L1+aTGF-β vs aTG | Down-regulated gen | CD45+ |
| BP | GO:0043900 | regulation of multi-organism process | 1.60E-07 | 1.82E-05 | Trim30a/Pfn1/Jun | 9 | aPD-L1+aTGF-β vs aTG | Down-regulated gen | CD45+ |
| BP | GO:0045071 | negative regulation of viral genome replication | 2.01E-07 | 2.09E-05 | Oasl1/Isg20/Bst2/ | 5 | aPD-L1+aTGF-β vs aTG | Down-regulated gen | CD45+ |
| BP | GO:0019058 | viral life cycle | 2.86E-07 | 2.75E-05 | Trim30a/Oasl1/Isg | 7 | aPD-L1+aTGF-β vs aTG | Down-regulated gen | CD45+ |
| BP | GO:0035455 | response to interferon-alpha | 3.43E-07 | 3.06E-05 | Ifit3/Ifit2/Bst2/Ifit1 | 4 | aPD-L1+aTGF-β vs aTG | Down-regulated gen | CD45+ |
| BP | GO:1903900 | regulation of viral life cycle | 7.91E-07 | 6.59E-05 | Trim30a/Oasl1/Isg | 6 | aPD-L1+aTGF-β vs aTG | Down-regulated gen | CD45+ |
| BP | GO:1903901 | negative regulation of viral life cycle | 1.07E-06 | 8.39E-05 | Oasl1/Isg20/Bst2/ | 5 | aPD-L1+aTGF-β vs aTG | Down-regulated gen | CD45+ |
| BP | GO:0045069 | regulation of viral genome replication | 2.07E-06 | 1.52E-04 | Oasl1/Isg20/Bst2/ | 5 | aPD-L1+aTGF-β vs aTG | Down-regulated gen | CD45+ |
| BP | GO:0043901 | negative regulation of multi-organism process | 2.30E-06 | 1.60E-04 | Jun/Oasl1/Isg20/E | 6 | aPD-L1+aTGF-β vs aTG | Down-regulated gen | CD45+ |
| BP | GO:0035457 | cellular response to interferon-alpha | 3.61E-06 | 2.37E-04 | Ifit3/Ifit2/Ifit1 | 3 | aPD-L1+aTGF-β vs aTG | Down-regulated gen | CD45+ |
| BP | GO:0019079 | viral genome replication | 5.06E-06 | 3.16E-04 | Oasl1/Isg20/Bst2/ | 5 | aPD-L1+aTGF-β vs aTG | Down-regulated gen | CD45+ |
| BP | GO:0048524 | positive regulation of viral process | 4.74E-05 | 2.82E-03 | Trim30a/Pfn1/Jun | 4 | aPD-L1+aTGF-β vs aTG | Down-regulated gen | CD45+ |
| BP | GO:0035456 | response to interferon-beta | 3.98E-04 | 2.26E-02 | Ifit3/Bst2/Ifit1 | 3 | aPD-L1+aTGF-β vs aTG | Down-regulated gen | CD45+ |
| BP | GO:0034121 | regulation of toll-like receptor signaling pathway | 5.15E-04 | 2.80E-02 | Trim30a/Irf7/Rsad | 3 | aPD-L1+aTGF-β vs aTG | Down-regulated gen | CD45+ |
| BP | GO:0043902 | positive regulation of multi-organism process | 6.34E-04 | 3.30E-02 | Trim30a/Pfn1/Jun | 4 | aPD-L1+aTGF-β vs aTG | Down-regulated gen | CD45+ |
| BP | GO:0035994 | response to muscle stretch | 8.92E-04 | 4.45E-02 | Fos/Jun | 2 | aPD-L1+aTGF-β vs aTG | Down-regulated gen | CD45+ |
| MF | GO:0031625 | ubiquitin protein ligase binding | 6.90E-04 | 3.68E-02 | Cdkn1a/Hspa1b/J | 5 | aPD-L1+aTGF-β vs aTG | Down-regulated gen | CD45+ |
| MF | GO:0044389 | ubiquitin-like protein ligase binding | 9.13E-04 | 3.68E-02 | Cdkn1a/Hspa1b/J | 5 | aPD-L1+aTGF-β vs aTG | Down-regulated gen | CD45+ |
| MF | GO:0001158 | enhancer sequence-specific DNA binding | 1.15E-03 | 3.68E-02 | H3f3b/Jun/Irf7 | 3 | aPD-L1+aTGF-β vs aTG | Down-regulated gen | CD45+ |
| MF | GO:0044183 | protein binding involved in protein folding | 1.31E-03 | 3.68E-02 | Hspa1b/Hspa1a | 2 | aPD-L1+aTGF-β vs aTG | Down-regulated gen | CD45+ |
| MF | GO:0051787 | misfolded protein binding | 1.31E-03 | 3.68E-02 | Hspa1b/Hspa1a | 2 | aPD-L1+aTGF-β vs aTG | Down-regulated gen | CD45+ |
| MF | GO:0070412 | R-SMAD binding | 1.31E-03 | 3.68E-02 | Fos/Jun | 2 | aPD-L1+aTGF-β vs aTG | Down-regulated gen | CD45+ |
| MF | GO:0035326 | enhancer binding | 1.56E-03 | 3.78E-02 | H3f3b/Jun/Irf7 | 3 | aPD-L1+aTGF-β vs aTG | Down-regulated gen | CD45+ |
| BP | GO:0006457 | protein folding | 3.56E-06 | 2.79E-03 | Cryab/Dnajb1/Hsp | 4 | aPD-L1+aTGF-β vs aTG | Down-regulated gen | CD45- |
| BP | GO:0061077 | chaperone-mediated protein folding | 7.89E-06 | 3.09E-03 | Dnajb1/Hspa1b/H | 3 | aPD-L1+aTGF-β vs aTG | Down-regulated gen | CD45- |
| BP | GO:1990000 | amyloid fibril formation | 5.54E-05 | 1.45E-02 | Cryab/Apoe | 2 | aPD-L1+aTGF-β vs aTG | Down-regulated gen | CD45- |
| BP | GO:0032462 | regulation of protein homooligomerization | 1.40E-04 | 2.48E-02 | Cryab/Apoe | 2 | aPD-L1+aTGF-β vs aTG | Down-regulated gen | CD45- |
| BP | GO:0060561 | apoptotic process involved in morphogenesis | 2.29E-04 | 2.48E-02 | Cryab/Ccn1 | 2 | aPD-L1+aTGF-β vs aTG | Down-regulated gen | CD45- |
| BP | GO:0051085 | chaperone cofactor-dependent protein refolding | 2.46E-04 | 2.48E-02 | Dnajb1/Hspa1b | 2 | aPD-L1+aTGF-β vs aTG | Down-regulated gen | CD45- |
| BP | GO:0043281 | regulation of cysteine-type endopeptidase activit | 3.18E-04 | 2.48E-02 | Cryab/Hspa1b/Cc | 3 | aPD-L1+aTGF-β vs aTG | Down-regulated gen | CD45- |
| BP | GO:0051084 | 'de novo' posttranslational protein folding | 3.39E-04 | 2.48E-02 | Dnajb1/Hspa1b | 2 | aPD-L1+aTGF-β vs aTG | Down-regulated gen | CD45- |
| BP | GO:1902742 | apoptotic process involved in development | 3.39E-04 | 2.48E-02 | Cryab/Ccn1 | 2 | aPD-L1+aTGF-β vs aTG | Down-regulated gen | CD45- |
| BP | GO:0006469 | negative regulation of protein kinase activity | 3.46E-04 | 2.48E-02 | Gadd45g/Hspb1/ | 3 | aPD-L1+aTGF-β vs aTG | Down-regulated gen | CD45- |
| BP | GO:0006458 | 'de novo' protein folding | 3.59E-04 | 2.48E-02 | Dnajb1/Hspa1b | 2 | aPD-L1+aTGF-β vs aTG | Down-regulated gen | CD45- |
| BP | GO:0032459 | regulation of protein oligomerization | 3.80E-04 | 2.48E-02 | Cryab/Apoe | 2 | aPD-L1+aTGF-β vs aTG | Down-regulated gen | CD45- |
| BP | GO:2000116 | regulation of cysteine-type endopeptidase activit | 4.32E-04 | 2.57E-02 | Cryab/Hspa1b/Cc | 3 | aPD-L1+aTGF-β vs aTG | Down-regulated gen | CD45- |
| BP | GO:0033673 | negative regulation of kinase activity | 4.59E-04 | 2.57E-02 | Gadd45g/Hspb1/ | 3 | aPD-L1+aTGF-β vs aTG | Down-regulated gen | CD45- |
| BP | GO:0051348 | negative regulation of transferase activity | 6.79E-04 | 3.55E-02 | Gadd45g/Hspb1/ | 3 | aPD-L1+aTGF-β vs aTG | Down-regulated gen | CD45- |
| BP | GO:0042982 | amyloid precursor protein metabolic process | 7.64E-04 | 3.74E-02 | Itm2b/Apoe | 2 | aPD-L1+aTGF-β vs aTG | Down-regulated gen | CD45- |
| BP | GO:0072593 | reactive oxygen species metabolic process | 8.11E-04 | 3.74E-02 | Cryab/COX2/Ccn | 3 | aPD-L1+aTGF-β vs aTG | Down-regulated gen | CD45- |
| BP | GO:0052548 | regulation of endopeptidase activity | 1.01E-03 | 4.21E-02 | Cryab/Hspa1b/Cc | 3 | aPD-L1+aTGF-β vs aTG | Down-regulated gen | CD45- |
| BP | GO:0042773 | ATP synthesis coupled electron transport | 1.02E-03 | 4.21E-02 | COX2/COX3 | 2 | aPD-L1+aTGF-β vs aTG | Down-regulated gen | CD45- |
| CC | GO:0031430 | M band | 3.25E-05 | 2.01E-03 | Cryab/Hspb1 | 2 | aPD-L1+aTGF-β vs aTG | Down-regulated gen | CD45- |
| CC | GO:0045277 | respiratory chain complex IV | 7.07E-05 | 2.19E-03 | COX2/COX3 | 2 | aPD-L1+aTGF-β vs aTG | Down-regulated gen | CD45- |
| CC | GO:0031672 | A band | 1.49E-04 | 2.96E-03 | Cryab/Hspb1 | 2 | aPD-L1+aTGF-β vs aTG | Down-regulated gen | CD45- |
| CC | GO:0070069 | cytochrome complex | 1.91E-04 | 2.96E-03 | COX2/COX3 | 2 | aPD-L1+aTGF-β vs aTG | Down-regulated gen | CD45- |
| CC | GO:0014069 | postsynaptic density | 1.64E-03 | 1.21E-02 | Cryab/Dnajb1/Hs | 3 | aPD-L1+aTGF-β vs aTG | Down-regulated gen | CD45- |
| CC | GO:0032279 | asymmetric synapse | 1.65E-03 | 1.21E-02 | Cryab/Dnajb1/Hs | 3 | aPD-L1+aTGF-β vs aTG | Down-regulated gen | CD45- |
| CC | GO:0098803 | respiratory chain complex | 1.68E-03 | 1.21E-02 | COX2/COX3 | 2 | aPD-L1+aTGF-β vs aTG | Down-regulated gen | CD45- |
| CC | GO:0098984 | neuron to neuron synapse | 1.89E-03 | 1.21E-02 | Cryab/Dnajb1/Hs | 3 | aPD-L1+aTGF-β vs aTG | Down-regulated gen | CD45- |
| CC | GO:0099572 | postsynaptic specialization | 1.89E-03 | 1.21E-02 | Cryab/Dnajb1/Hs | 3 | aPD-L1+aTGF-β vs aTG | Down-regulated gen | CD45- |
| CC | GO:0070469 | respiratory chain | 1.99E-03 | 1.21E-02 | COX2/COX3 | 2 | aPD-L1+aTGF-β vs aTG | Down-regulated gen | CD45- |
| CC | GO:0030018 | Z disc | 2.23E-03 | 1.21E-02 | Cryab/Hspb1 | 2 | aPD-L1+aTGF-β vs aTG | Down-regulated gen | CD45- |
| CC | GO:0043204 | perikaryon | 2.34E-03 | 1.21E-02 | Cryab/Hspb1 | 2 | aPD-L1+aTGF-β vs aTG | Down-regulated gen | CD45- |
| CC | GO:0031674 | I band | 2.54E-03 | 1.21E-02 | Cryab/Hspb1 | 2 | aPD-L1+aTGF-β vs aTG | Down-regulated gen | CD45- |
| CC | GO:0030017 | sarcomere | 4.91E-03 | 2.18E-02 | Cryab/Hspb1 | 2 | aPD-L1+aTGF-β vs aTG | Down-regulated gen | CD45- |
| CC | GO:0043197 | dendritic spine | 5.89E-03 | 2.19E-02 | Cryab/Dnajb1 | 2 | aPD-L1+aTGF-β vs aTG | Down-regulated gen | CD45- |
| CC | GO:0044449 | contractile fiber part | 5.97E-03 | 2.19E-02 | Cryab/Hspb1 | 2 | aPD-L1+aTGF-β vs aTG | Down-regulated gen | CD45- |
| CC | GO:0044309 | neuron spine | 6.13E-03 | 2.19E-02 | Cryab/Dnajb1 | 2 | aPD-L1+aTGF-β vs aTG | Down-regulated gen | CD45- |
| CC | GO:0030016 | myofibril | 6.37E-03 | 2.19E-02 | Cryab/Hspb1 | 2 | aPD-L1+aTGF-β vs aTG | Down-regulated gen | CD45- |
| CC | GO:0043292 | contractile fiber | 7.38E-03 | 2.41E-02 | Cryab/Hspb1 | 2 | aPD-L1+aTGF-β vs aTG | Down-regulated gen | CD45- |
| CC | GO:0061827 | sperm head | 8.85E-03 | 2.74E-02 | Dnajb1 | 1 | aPD-L1+aTGF-β vs aTG | Down-regulated gen | CD45- |
| CC | GO:0034364 | high-density lipoprotein particle | 1.05E-02 | 2.95E-02 | Apoe | 1 | aPD-L1+aTGF-β vs aTG | Down-regulated gen | CD45- |
| CC | GO:0043083 | synaptic cleft | 1.05E-02 | 2.95E-02 | Apoe | 1 | aPD-L1+aTGF-β vs aTG | Down-regulated gen | CD45- |
| CC | GO:0034358 | plasma lipoprotein particle | 1.92E-02 | 4.77E-02 | Apoe | 1 | aPD-L1+aTGF-β vs aTG | Down-regulated gen | CD45- |
| CC | GO:0099524 | postsynaptic cytosol | 1.92E-02 | 4.77E-02 | Dnajb1 | 1 | aPD-L1+aTGF-β vs aTG | Down-regulated gen | CD45- |
| CC | GO:1990777 | lipoprotein particle | 1.92E-02 | 4.77E-02 | Apoe | 1 | aPD-L1+aTGF-β vs aTG | Down-regulated gen | CD45- |
| CC | GO:0032994 | protein-lipid complex | 2.08E-02 | 4.96E-02 | Apoe | 1 | aPD-L1+aTGF-β vs aTG | Down-regulated gen | CD45- |
| MF | GO:0001540 | amyloid-beta binding | 6.70E-06 | 4.89E-04 | Cryab/Itm2b/Apoe | 3 | aPD-L1+aTGF-β vs aTG | Down-regulated gen | CD45- |
| MF | GO:0051082 | unfolded protein binding | 4.30E-05 | 1.49E-03 | Cryab/Dnajb1/Hs | 3 | aPD-L1+aTGF-β vs aTG | Down-regulated gen | CD45- |
| MF | GO:0044183 | protein binding involved in protein folding | 1.06E-04 | 1.49E-03 | Hspa1b/Hspb1 | 2 | aPD-L1+aTGF-β vs aTG | Down-regulated gen | CD45- |
| MF | GO:0004129 | cytochrome-c oxidase activity | 1.30E-04 | 1.49E-03 | COX2/COX3 | 2 | aPD-L1+aTGF-β vs aTG | Down-regulated gen | CD45- |
| MF | GO:0015002 | heme-copper terminal oxidase activity | 1.30E-04 | 1.49E-03 | COX2/COX3 | 2 | aPD-L1+aTGF-β vs aTG | Down-regulated gen | CD45- |
| MF | GO:0016676 | oxidoreductase activity, acting on a heme group | 1.30E-04 | 1.49E-03 | COX2/COX3 | 2 | aPD-L1+aTGF-β vs aTG | Down-regulated gen | CD45- |
| MF | GO:0016675 | oxidoreductase activity, acting on a heme group | 1.42E-04 | 1.49E-03 | COX2/COX3 | 2 | aPD-L1+aTGF-β vs aTG | Down-regulated gen | CD45- |

| | | | | | | | | | |
|---|---|---|---|---|---|---|---|---|---|
| MF | GO:0042277 | peptide binding | 6.33E-04 | 5.78E-03 | Cryab/Itm2b/Apoe | 3 | aPD-L1+aTGF-β vs aTG | Down-regulated gen | CD45- |
| MF | GO:0009055 | electron transfer activity | 1.15E-03 | 8.93E-03 | COX2/COX3 | 2 | aPD-L1+aTGF-β vs aTG | Down-regulated gen | CD45- |
| MF | GO:0033218 | amide binding | 1.22E-03 | 8.93E-03 | Cryab/Itm2b/Apoe | 3 | aPD-L1+aTGF-β vs aTG | Down-regulated gen | CD45- |
| MF | GO:0015078 | proton transmembrane transporter activity | 2.39E-03 | 1.59E-02 | COX2/COX3 | 2 | aPD-L1+aTGF-β vs aTG | Down-regulated gen | CD45- |
| MF | GO:0008201 | heparin binding | 4.49E-03 | 2.56E-02 | Ccn1/Apoe | 2 | aPD-L1+aTGF-β vs aTG | Down-regulated gen | CD45- |
| MF | GO:0031072 | heat shock protein binding | 4.56E-03 | 2.56E-02 | Dnajb1/Hspa1b | 2 | aPD-L1+aTGF-β vs aTG | Down-regulated gen | CD45- |
| MF | GO:0005539 | glycosaminoglycan binding | 7.46E-03 | 3.89E-02 | Ccn1/Apoe | 2 | aPD-L1+aTGF-β vs aTG | Down-regulated gen | CD45- |
| MF | GO:0005212 | structural constituent of eye lens | 1.07E-02 | 4.60E-02 | Cryab | 1 | aPD-L1+aTGF-β vs aTG | Down-regulated gen | CD45- |
| MF | GO:0017127 | cholesterol transporter activity | 1.07E-02 | 4.60E-02 | Apoe | 1 | aPD-L1+aTGF-β vs aTG | Down-regulated gen | CD45- |
| MF | GO:0043395 | heparan sulfate proteoglycan binding | 1.07E-02 | 4.60E-02 | Apoe | 1 | aPD-L1+aTGF-β vs aTG | Down-regulated gen | CD45- |
| MF | GO:0015077 | monovalent inorganic cation transmembrane tra | 1.15E-02 | 4.60E-02 | COX2/COX3 | 2 | aPD-L1+aTGF-β vs aTG | Down-regulated gen | CD45- |
| MF | GO:1901681 | sulfur compound binding | 1.20E-02 | 4.60E-02 | Ccn1/Apoe | 2 | aPD-L1+aTGF-β vs aTG | Down-regulated gen | CD45- |
